# Repeated truncation of a modular antimicrobial peptide gene for neural context

**DOI:** 10.1101/2021.02.24.432738

**Authors:** M.A. Hanson, B. Lemaitre

## Abstract

Antimicrobial peptides (AMPs) are host-encoded antibiotics that combat invading pathogens. These genes commonly encode multiple products as post-translationally cleaved polypeptides. Recent studies have highlighted roles for AMPs in neurological contexts suggesting functions for these defence molecules beyond infection. During our immune study characterizing the antimicrobial peptide gene *Baramicin,* we recovered multiple *Baramicin* paralogs in *Drosophila melanogaster* and other species, united by their N-terminal IM24 domain. Not all paralogs were immune-induced. Here, through careful dissection of the *Baramicin* family’s evolutionary history, we find that these non-immune paralogs result from repeated events of duplication and subsequent truncation of the coding sequence from an immune-inducible ancestor. These truncations leave only the IM24 domain as the prominent gene product. Surprisingly, using mutation and targeted gene silencing we demonstrate that two such genes are adapted for function in neural contexts in *D. melanogaster.* We also show enrichment in the head for independent *Baramicin* genes in other species. The *Baramicin* evolutionary history reveals that the IM24 *Baramicin* domain is not strictly useful in an immune context. We thus provide a case study for how an AMP-encoding gene might play dual roles in both immune and non-immune processes via its multiple peptide products. We reflect on these findings to highlight a blind spot in the way researchers approach AMP research in in vivo contexts.

**Significance statement:** Antimicrobial peptides are immune proteins recently implicated in neurological roles. To date little attention has been paid to the contributions of different gene products in this function. Here we show that an antimicrobial peptide gene encodes multiple products with either immune-specific or neurological roles.

## Introduction

Antimicrobial peptides (AMPs) are immune effectors best known for their role in defence against infection. These antimicrobials are commonly encoded as a polypeptide including both pro- and mature peptide domains (Zanetti 2005; Hanson and Lemaitre 2020). AMP genes frequently experience events of duplication and loss (Wang and Zhu 2011; Vilcinskas et al. 2013; Sackton et al. 2017; Hanson, Lemaitre, et al. 2019) and undergo rapid evolution at the sequence level (Tennessen 2005; Jiggins and Kim 2007; Hellgren et al. 2010; Halldórsdóttir and Árnason 2015; Hanson et al. 2016; Chapman et al. 2019). The selective pressures that drive these evolutionary outcomes are likely the consequence of host-pathogen interactions (Unckless et al. 2016). However AMPs and AMP-like genes in many species have also recently been implicated in non-immune roles in flies, nematodes, and humans. These new contexts suggest that the evolutionary forces acting on AMP genes may not be driven strictly by trade-offs in host defence, but rather by conflicts between roles in immunity and other non-immune functions.

For instance, Diptericins are membrane-disrupting antimicrobial peptides of flies (Diptera) that are required for defence against infection by *Providencia* bacteria (Unckless et al. 2016; Hanson, Dostálová, et al. 2019). It was therefore surprising that the *D. melanogaster* gene *Diptericin B (DptB)* affects memory processes (Barajas-azpeleta et al. 2018). In this study, *DptB* derived from the fly fat body (analogous to the mammalian liver) regulated the ability of the fly to form long-term memory associations (Barajas-azpeleta et al. 2018). Another AMP-like gene, nemuri, regulates fly sleep and promotes survival upon infection (Toda et al. 2019). Studies in nematodes have also shown that an immune-induced polypeptide (NLP-29) binds to a G-protein coupled receptor (NPR-12) triggering neurodegeneration through activation of the NPR-12-dependent autophagy pathway (Lezi et al. 2018), and injury triggers epidermal AMPs including NLP-29 to promote sleep (Sinner et al. 2021). *Drosophila* AMPs have also recently been shown to regulate behaviours after seeing parasitoid wasps (Ebrahim et al. 2021), during feeding with different bacteria (Kobler et al. 2020), or following infection (Hanson et al. 2021). In humans, the *Cathelicidin* gene encodes the AMP LL-37, which is implicated in glia-mediated neuroinflammation and Alzheimer’s disease (Lee et al. 2015; De Lorenzi et al. 2017). Indeed recent evidence suggests Alzheimer’s disease is an infectious syndrome (Dominy et al. 2019), though the importance of this process is debated (Abbott 2020). Notably, AMPs share a number of properties with classic neuropeptides (Brogden et al. 2005), further muddying the distinction between peptides of the immune and nervous systems.

We recently described a novel antifungal peptide gene of *Drosophila melanogaster* that we named *Baramicin A (BaraA)* (Hanson et al. 2021). A unique aspect of *BaraA* is its precursor protein structure, which encodes a polypeptide cleaved into multiple mature products by interspersed furin cleavage sites. The use of furin cleavage sites to produce more than one mature peptide from a single polypeptide precursor is widespread in animal AMP genes (Gerdol et al. 2020; Hanson and Lemaitre 2020), including multiple peptide repeats in bees and other flies (Casteels-Josson et al. 1993; Hanson et al. 2016). However, *BaraA* represents an exceptional case as many tandem repeat peptides are cleaved by furin from a single precursor protein, effectively resembling a “protein-based operon”. The immature precursor protein of *D. melanogaster BaraA* encodes three types of domains: an IM24 domain, three tandem repeats of IM10-like domains, and an IM22 domain. *BaraA* mutants are susceptible to infection by fungi, and *in vitro* experiments suggest the *BaraA* IM10-like peptides have antifungal activity (Hanson et al. 2021). The other *Baramicin* domains encoding IM22 and IM24 remain uncharacterized. Curiously, *BaraA* deficient flies also display an erect wing behavioural phenotype upon immune stimulation even in the absence of infection, suggesting that *BaraA* products could have non-microbial targets (Hanson et al. 2021).

In this study, we describe the evolution of the Drosophilid *Baramicin* gene family. Three unique *Baramicin* genes *(BaraA, B, and C*) are present in the genome of *D. melanogaster.* Surprisingly, only *BaraA* is immune-induced, while *BaraB* and *BaraC* are enriched in the nervous system. Both *BaraB* and *BaraC* have truncations compared to the ancestral *Baramicin* gene, and these two genes effectively encode just the *Baramicin* IM24 domain. We found similar truncations in other species, and confirmed loss of immune expression for IM24-specific *Baramicins* of other species.

We also confirmed enrichment in the head or nervous system for IM24-specific genes in *D. melanogaster* and other species. We resolved the genomic ancestry of the *Baramicins,* which confirmed that these repeated truncations creating IM24-specific genes came from independent events (convergent evolution). The complex ‘protein operon’ polypeptide nature of *Baramicin* draws attention to how different sub-peptides can be adapted to context-specific roles, like in immunity or neurology. Attention to the multiple peptide products of AMP genes could explain how these immune effectors affect both immune and neurological processes.

## Results

### Baramicin is an ancestral immune effector

The *Baramicin A* gene was only recently described as encoding antifungal effectors by our group (Hanson et al. 2021), and another recent study also confirmed *Baramicin’*s important contribution to Toll immune defence (Huang et al. 2020). These initial characterizations were done only in *D. melanogaster,* and focused on one *Baramicin* gene (*BaraA*). We will therefore first provide a basic description of the immune *Baramicins* of other species and also the larger *Baramicin* gene family of *D. melanogaster* to establish that this is a classically immune gene family. This is relevant to non-immune duplicate genes discussed later.

In *D. melanogaster, BaraA* is regulated by the Toll immune signalling pathway (Huang et al. 2020; Hanson et al. 2021). Using BLAST, we recovered *BaraA*-like genes encoding each Baramicin peptide (IM24, IM10-like, and IM22) across the genus *Drosophila* and in the outgroup *Scaptodrosophila lebanonensis.* In many species, this was the only *Baramicin* gene present, suggesting *Dmel\BaraA* resembles the ancestral *Baramicin* structure. We performed infection experiments to confirm that *BaraA*-like genes were immune-inducible in the diverse species *D. melanogaster, D. pseudoobscura, D. willistoni, D. virilis,* and *D. neotesteacea* (last common ancestor ~53mya (Suvorov et al. 2021)) with *Micrococcus luteus* and *Candida albicans,* two microbes that stimulate the Toll pathway (**Fig.** 1A). In all five species, *BaraA*-like genes were immune-induced (**Fig.** 1B-F). We therefore confirm the ancestral *Baramicin* was an immune-induced gene. Deviations from immune function derived.

**Figure 1:**
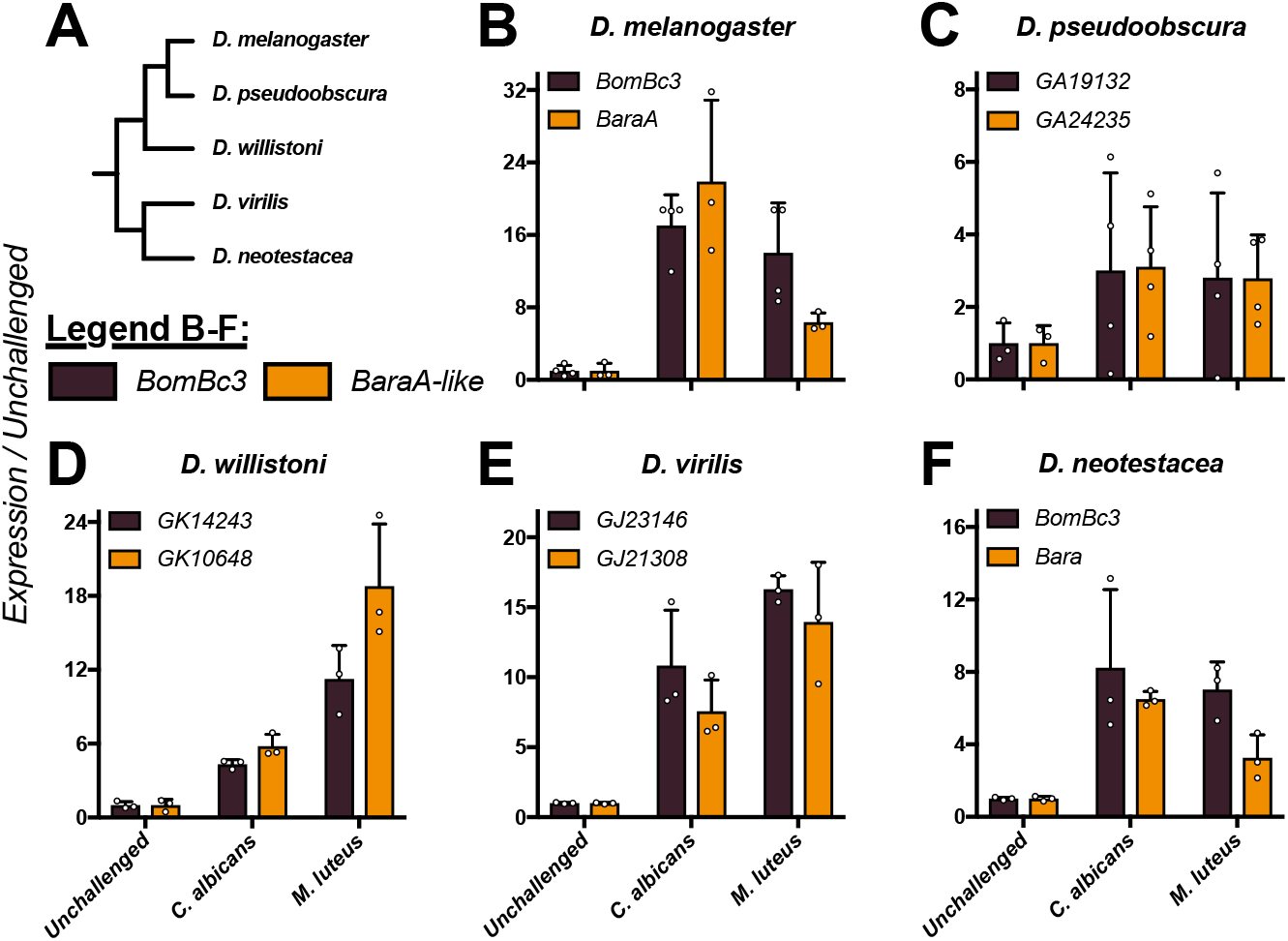
The ancestral *BaraA* gene was immune-induced. A) Cladogram of species used in B-F. B-F) Expression of the Toll-responsive gene *BomBc3* (brown) or *BaraA-like* genes (orange) in diverse *Drosophila* species upon infection. In all cases, both *BomBc3* and *BaraA-like* genes are induced upon infection by either *C. albicans* yeast or *M. luteus* bacteria.

### The D. melanogaster genome encodes up to four Baramicins: BaraAl, BaraA2, BaraB and BaraC

In *D. melanogaster,* we recovered four *Baramicin* genes. First, we realized that a duplication of *BaraA* is actively segregating in wild flies (**Fig. 2**A). The *D. melanogaster* R6 genome assembly encodes two 100% identical *BaraA* genes *(CG33470* and *CG18279, BaraA1* and *BaraA2* respectively). We screened 132 DGRP lines for the *BaraA* duplication event, finding only ~14% (18/132) of strains were PCR-positive for two *BaraA* copies (supplementary data file 1). Perhaps as a consequence of the identical sequences of these two genes, this genome region is poorly resolved in RNA sequencing studies and the Drosophila Genetic Reference Panel (DGRP, see **Fig.** S1) (Mackay et al. 2012; Leader et al. 2018). Because this region is poorly resolved, it is unclear if our PCR assay might be sensitive to cryptic sequence variation. However our PCR screen nevertheless confirms that this region is variable in the wild, and we additionally note that common fly strains seem to differ in their *BaraA* copy number, where extra gene copies correlates with increased expression after infection (see (Hanson et al. 2021) S10 Fig).

**Figure 2:**
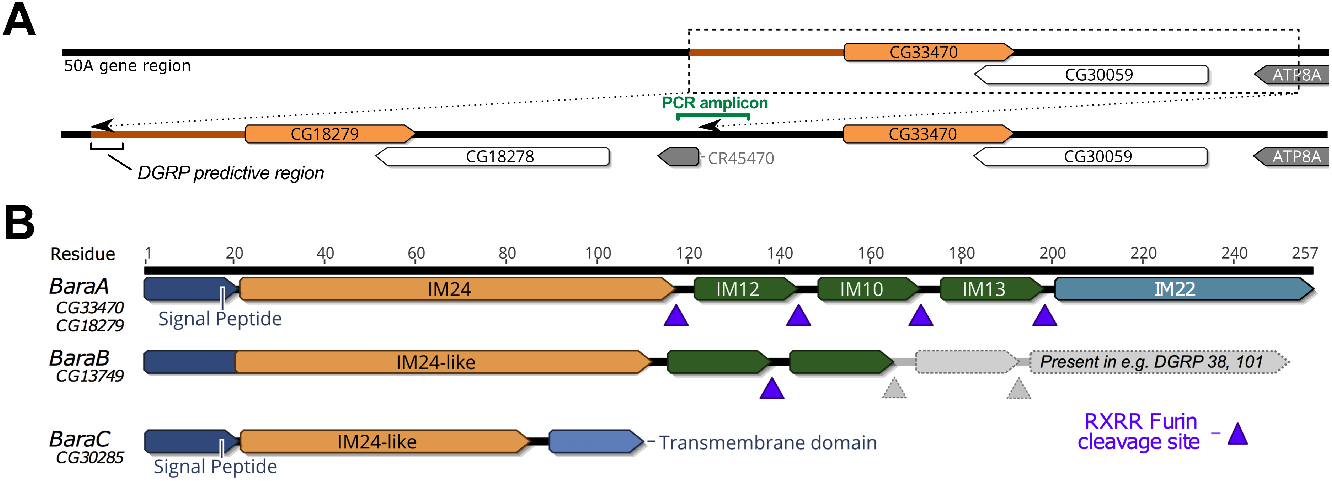
The *D. melanogaster Baramicin* genes. A) Schematic of the *BaraA* duplication. Using a PCR assay spanning the duplication-specific locus (PCR amplicon), we confirmed *BaraA* copy number is variable in various lab strains (Hanson et al. 2021) and wild-caught flies (Supplementary data file 1). B) *D. melanogaster* encodes two other *Baramicin* genes that we name *BaraB* and *BaraC.* These paralogs differ markedly in their precursor protein structure through truncation of the C-terminus relative to *BaraA.* The *BaraB* truncation is segregating in the DGRP (greyed out region, and see supplementary data file 2).

We also recovered two paralogous *Baramicin* genes in *D. melanogaster* through reciprocal BLAST searches: *CG13749* and *CG30285,* which we name *BaraB* and *BaraC* respectively (**Fig.** 2B). The three *Baramicin* gene loci are scattered on the right arm of chromosome II at cytological positions 44F9 (*BaraB*), 50A5 (*BaraA*), and 57F8 (*BaraC*). These paralogous *Baramicins* are united by the presence of the IM24 domain. In the case of *BaraB,* we additionally recovered a frameshift mutation (2R_4821599_INS) causing a premature stop segregating in the DGRP leading to the loss of IM13 and IM22 relative to the *BaraA* gene structure (**Fig. 2**B); this truncation is present in the Dmel_R6 genome assembly, but many DGRP strains encode a CDS with either a standard (e.g. DGRP38) or extended (e.g. DGRP101) IM22 domain (a DGRP *BaraB* alignment is provided in supplementary data file 2). Moreover, in contrast to BaraA, the initial IM10-like peptide of BaraB no longer follows a furin cleavage site, and encodes a serine (RSXR) in its IM10-like motif instead of the universal proline (RPXR) of *BaraA*-like IM10 peptides across the genus. Each of these mutations prevents the secretion of classical IM10-like and IM22 peptides by *BaraB.* Finally, *BaraC* encodes only IM24 tailed by a transmembrane domain at the C terminus (TMHMM v2.0 (Krogh et al. 2001)), and thus lacks both the IM10-like peptides and IM22 (**Fig. 2**B).

### BaraB and BaraC are not immune-inducible

*BaraA* is strongly induced following microbial challenge (Fig. 1), being predominantly regulated by the Toll pathway with a minor input from the Immune Deficiency (Imd) pathway (Huang et al. 2020; Hanson et al. 2021). We therefore assayed the expression of *BaraB* and *BaraC* in wild-type flies, and also flies with defective Toll *(spz^rm7^*) or Imd *(Rel^E20^)* signalling to see if their basal expression relied on these pathways. Surprisingly, neither gene was induced upon infection regardless of microbial challenge (**Fig. 3**A and **Fig.** S2A-B). Of note: *BaraC* levels were consistently reduced in *spz^rm7^* mutants regardless of treatment (cumulative data in **Fig.** S2C, p = .005), suggesting *BaraC* basal expression is affected by Toll signalling. We next generated a novel time course of development from egg to adult to monitor the expression of the three *Baramicin* genes. We found that expression of all genes increased over development and reached their highest level in young adults (**Fig.** 3B). Of note, *BaraB* expression approached the lower limit of our assay’s detection sensitivity at early life stages. However *BaraB* was robustly detected beginning at the pupal stage, indicating it is expressed during metamorphosis. *BaraC* expression also increased markedly between the L3 larval stage and pupal stage.

**Figure 3:**
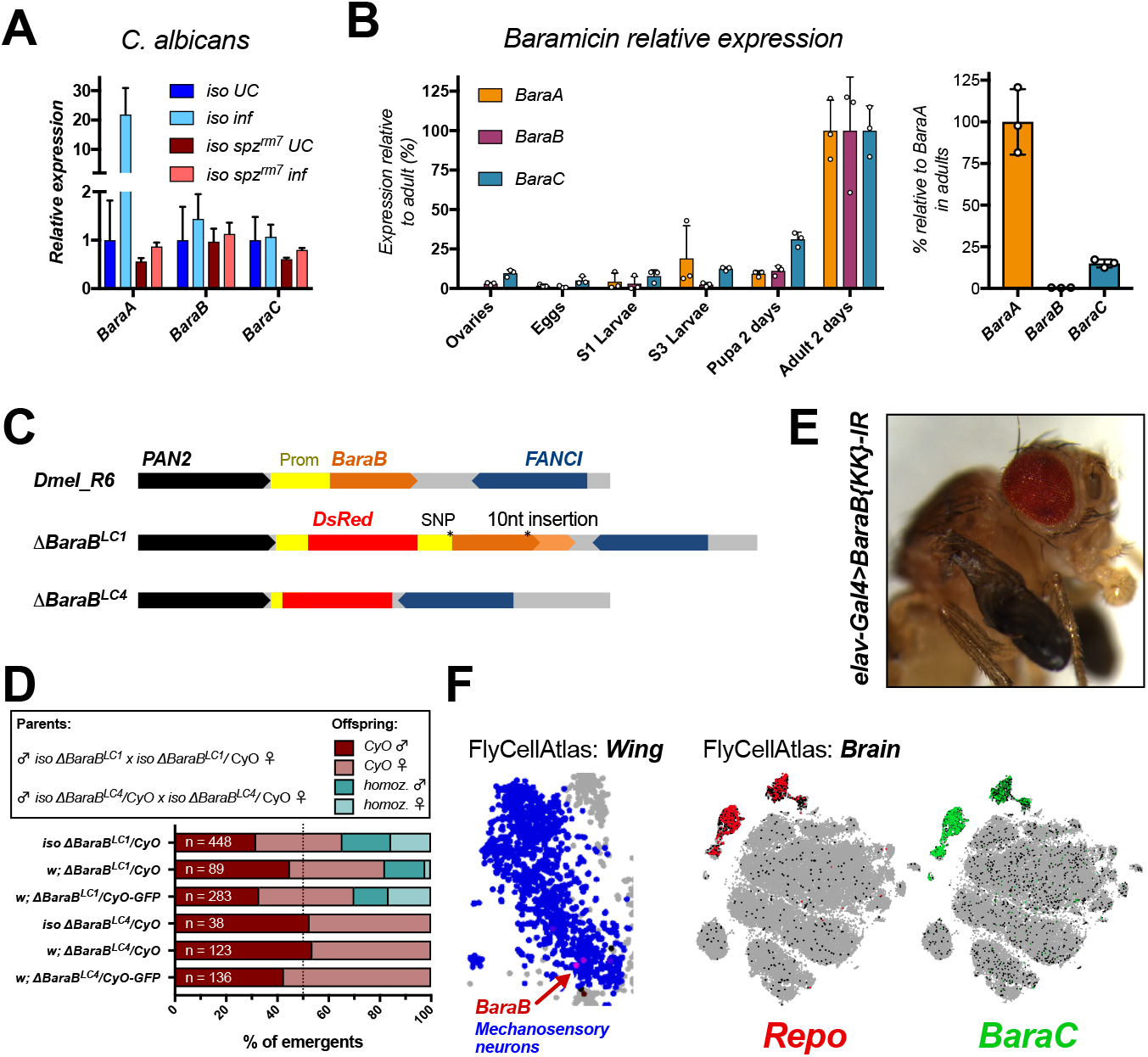
*D. melanogaster BaraB and BaraC* have neural functions. A) Only *BaraA* is immune-induced. *BaraB* and *BaraC* do not respond to infection, though basal *BaraC* expression relies on Toll signalling (Fig. S3C). B) Time series of whole animal *Baramicin* expression over the course of development. Expression values are normalized within each gene, with expression in the adult set to 100% (left panel). For context, normalizing each gene to *BaraA* in adults shows that *BaraC* and especially *BaraB* expression is much lower at a whole fly level (right panel). C) *ΔBaraB* mutations generated in this study. *ΔBaraB^LC1^* is an incidental hypomorph with reduced *BaraB* expression (Fig. S3A), while *ΔBaraB^LC4^* has the intended genetic replacement of *BaraB* with a DsRed cassette. Both mutations cause expression of DsRed in the eyes, ocelli, and abdomen. D) Partial lethality of *ΔBaraB^LC1^* hypomorphs or complete lethality of *ΔBaraB^LC4^* null flies in varied genetic backgrounds. E) The nubbin-like wing phenotype is copied by *BaraB* gene silencing using the neural driver *elav-Gal4.* F) Specific wing mechanosensory neurons seem to express *BaraB* in adults. Meanwhile the *BaraC* gene almost perfectly matches the expression pattern of glia specific *Æepo*-expressing cells in single cell RNA sequencing of the adult fly brain (Li et al. 2021).

Collectively we reveal that *BaraA* is part of a larger gene family. While the *BaraA* gene was first described as an immune effector, the two *Baramicin* paralogs *BaraB* and *BaraC* are not induced by infection in *D. melanogaster.* Both *BaraB* and *BaraC* first see increased expression during pupation, and are ultimately expressed at their highest levels in adults.

### Dmel\BaraB is required in the nervous system over the course of development

A simple interpretation of the truncated gene structure and low levels of *BaraB* expression is that this gene is undergoing pseudogenization. Indeed, AMP gene pseudogenization is common in insects including *Drosophila* (Quesada et al. 2005; Rolff and Schmid-Hempel 2016; Hanson, Lemaitre, et al. 2019). To explore *BaraB* function, we used two mutations for *BaraB (ΔBaraB^LC1^* and *ΔBaraB^LC4^,* generously gifted by S.A. Wasserman). These mutations were made using a CRISPR double gRNA approach to replace the *BaraB* locus with sequence from the pHD-DsRed vector. The *ΔBaraB^LC1^* and *ΔBaraB^LC4^* mutations differ in their ultimate effect, as *ΔBaraB^LC1^* is an incidental insertion of the DsRed cassette in the promoter of the gene. This disruption reduces gene expression, resulting in a hypomorph state (**Fig. S3** A). The *ΔBaraB^LC4^* mutation however deletes the locus as intended, leading to *BaraB* null flies (**Fig. 3**C).

We further introgressed both *ΔBaraB* mutations into the DrosDel isogenic background (referred to as *iso*) for seven generations according to Ferreira et al. (Ferreira et al. 2014). At the same time, we combined the original *ΔBaraB* chromosomes with a CyO-GFP balancer chromosome in a mixed genetic background to distinguish homozygous/heterozygous larvae. In all cases, *ΔBaraB^LC4^* homozygous individuals failed to develop to the adult stage, whereas homozygous *ΔBaraB^LC1^* adults emerged (**Fig. 3**D). We next assessed *BaraB* hypomorph viability by crossing *ΔBaraB^LC1^* homozygous males to *ΔBaraB^LC1^/CyO* heterozygous females. The resulting offspring ratio departed from mendelian inheritance, and was exacerbated by rearing at 29°C (**Fig. S3**B). Using our *CyO-GFP* reporter to track hetero-vs. homozygous larvae revealed that the major lethal phase occurs primarily in the late larval and pupal stages (**Fig. S3**C-F), consistent with a role for *BaraB* in the larva/ pupa stage as suggested by an increase in expression at this stage (**Fig.** 3B). Some *ΔBaraB^LC1^* homozygous flies also exhibited locomotor defects, and/or a nubbin-like wing phenotype (FlyBase: FBrf0220532 and e.g. in **Fig. 3**E) where the wings were stuck in a shrivelled state for the remainder of the fly’s lifespan. However, a proportion of *ΔBaraB^LC1^* homozygotes successfully emerged, and unlike their siblings, had no immediate morphological or locomotory defects. The lifespan of morphologically normal *iso ΔBaraB^LC1^* adults is nevertheless significantly shorter compared to wild-type flies and *iso ΔBaraB^LC1^/CyO* siblings (**Fig. S4**G). We confirmed these developmental defects using ubiquitous gene silencing with *Actin5C-Gal4 (Act-Gal4)* to drive two *BaraB* RNAi/interfering RNA (IR) constructs *(TRiP-IR* and *KK-IR*). Both constructs resulted in significant lethality and occurrence of nubbin-like wings (**Table** S1). Genomic deficiency crosses also confirmed significantly reduced numbers of eclosing *BaraB*-deficient flies at 25°C (n = 114, χ^2^ p < .001) and 29°C (n = 63, χ^2^ p < .001) (**Fig. S3**H). We therefore conclude that full gene deletion causes lethality at the larva-pupa transition stage, and *BaraB* hypomorph flies suffer significant costs to fitness during development, and have reduced lifespan even following successful eclosion.

These data indicate *BaraB* is unlikely to be pseudogenized. While whole-fly *BaraB* expression is low, *BaraB* appears to perform an integral developmental role. The fact that there is a bimodal outcome in hypomorph-like *ΔBaraB^LC1^* adults (either severe defects or generally healthy) suggests *BaraB* is involved in passing some checkpoint during larval/pupal development. Flies deficient for *BaraB* may be more likely to fail at this developmental checkpoint, resulting in either lethality or developmental defects.

### Baramicin B plays an important role in the nervous system

We next sought to determine in which tissue(s) *BaraB* is required. A previous screen using neural *elav-Gal4* driven RNA interference highlighted *BaraB* silencing for lethality effects (n = 15) (Neely et al. 2010). Given *BaraB* mutant locomotory defects, we started by silencing *BaraB* in the nervous system using the pan-neural *elav-Gal4* driver with both the *TRiP-IR* and *KK-IR BaraB-IR* lines *(IR = interfering RNA).* We also used a combination of *UAS-Dicer2 (Dcr2)* and/or 29°C for greater silencing efficiency. In all cases, *BaraB-IR* driven by *elav-Gal4* caused a significant (p < .02) departure from mendelian inheritance in lethality and nubbin-like wing presentation (Table S1). Moreover the frequency of both lethality and the nubbin-like wing phenotype was increased with increasing strength of gene silencing. This analysis suggests that *BaraB* plays an important role in the nervous system, explaining both the lethality and nubbin-like wing phenotypes. Interestingly, *BaraB* is expressed in specific subset of mechanosensory neuron cells in the wing in FlyCellAtlas (Li et al. 2021) (**Fig.** 3F), despite very low levels of *BaraB* expression in other FlyCellAtlas tissue datasets. We additionally investigated the effect of *BaraB* silencing in non-neural tissues including the fat body *(c564-Gal4),* hemocytes *(hml-Gal4),* the gut (esg-Gal4, MyO-Gal4), the wing disc *(nubbin-Gal4),* and in myocytes *(mef2-gal4),* which did not present with increased lethality or nubbin-like wings. We also screened neural drivers specific for glia *(Repo-Gal4),* motor neurons *(D42-, VGMN-,* and *OK6-Gal4),* and a *BaraA-Gal4* driver (Hanson et al. 2021) that could overlap *BaraB*-expressing cells. However all these *Gal4>BaraB-IR* flies were viable and never exhibited overt morphological defects.

### Baramicin C is expressed in glia

Tissue-specific transcriptomic data indicate that *BaraC* is expressed in various neural tissues including the eye, brain, and the thoracic abdominal ganglion (**Fig. S4**A), but also the hindgut and rectal pads pointing to a complex expression pattern (Hammonds et al. 2013; Leader et al. 2018). We next searched FlyCellAtlas (Li et al. 2021) to narrow down which neural subtypes *BaraB and BaraC* were expressed in. *BaraB* expressing cells were few and showed only low expression in this dataset. However *BaraC* was robustly expressed in all glial cell types, fully overlapping the glia marker *Repo* (**Fig. 3**F). To confirm the observation that *BaraC* was expressed in glia, we compared the effects of *BaraC* RNA silencing *(BaraC-IR)* using *Act-Gal4* (ubiquitous), *elav-Gal4* (neural) and *Repo-Gal4* (glia) drivers on *BaraC* expression. *Act-Gal4* reduced *BaraC* expression to just ~14% that of control flies (**Fig. S4**B). By comparison *elav-Gal4* reduced *BaraC* expression to ~63% that of controls, while *Repo-Gal4* led to *BaraC* levels only 57% that of controls (overall controls vs. neural/glia-IR, p = .002). We also screened for overt lethality, and locomotor or developmental defects upon *BaraC* silencing using ubiquitous *Act-Gal4* and neural *elav-Gal4>Dcr2* or *Repo-Gal4*. However *BaraC* silencing never produced overt phenotypes in morphology or locomotor activity (not shown).

Collectively, our results support the notion that *BaraC* is expressed in the nervous system, and are consistent with *BaraC* expression being most localized to glial cells.

### Repeated genomic turnover of the Baramicin gene family

Our results thus far show that *BaraA*-like genes are consistently immune-induced in all *Drosophila* species (**Fig.** 1), however the two paralogs *Dmel\BaraB* and *Dmel\BaraC* are not immune-induced, and are truncated in a fashion that deletes some or all of the antifungal IM10-like peptides (**Fig. 2**B). These two *Baramicins* are now enriched in the nervous system (**Fig. 3**E-F). In the case of *BaraB,* a role in the nervous system is evidenced by severe defects recapitulated using pan-neural RNA silencing. In the case of *BaraC,* nervous system expression is evidenced by a clear overlap with *Repo*-expressing cells.

While *BaraA*-like genes are conserved throughout the genus *Drosophila, BaraB* is conserved only in Melanogaster group flies, and *BaraC* is found only in Melanogaster and Obscura group flies, indicating that both paralogs stem from duplication events of a *BaraA*-like ancestor (**Fig. 4**). To determine the ancestry of each *D. melanogaster Baramicin* gene, we traced their evolutionary history by analyzing genomic synteny through hierarchical orthologous groups (Train et al. 2019). Ancestry tracing revealed that these three loci ultimately stem from a single-locus ancestor encoding only one *Baramicin* gene that resembled *Dmel\BaraA* (**Fig. 4**A). This is evidenced by the presence of only a single *BaraA*-like gene in the outgroup *S. lebanonensis,* and also in multiple lineages of the subgenus Drosophila (**Fig. 4**B). Indeed, the general *BaraA* gene structure encoding IM24, tandem repeats of IM10-like peptides, and IM22 is conserved in *S. lebanonensis* and all *Drosophila* species (**Fig. 4**C). On the other hand, the *Dmel\BaraC* gene comes from an ancient duplication restricted to the subgenus Sophophora, and *Dmel\BaraB* resulted from a more recent duplication found only in the Melanogaster group (**Fig. 4**B).

**Figure 4:**
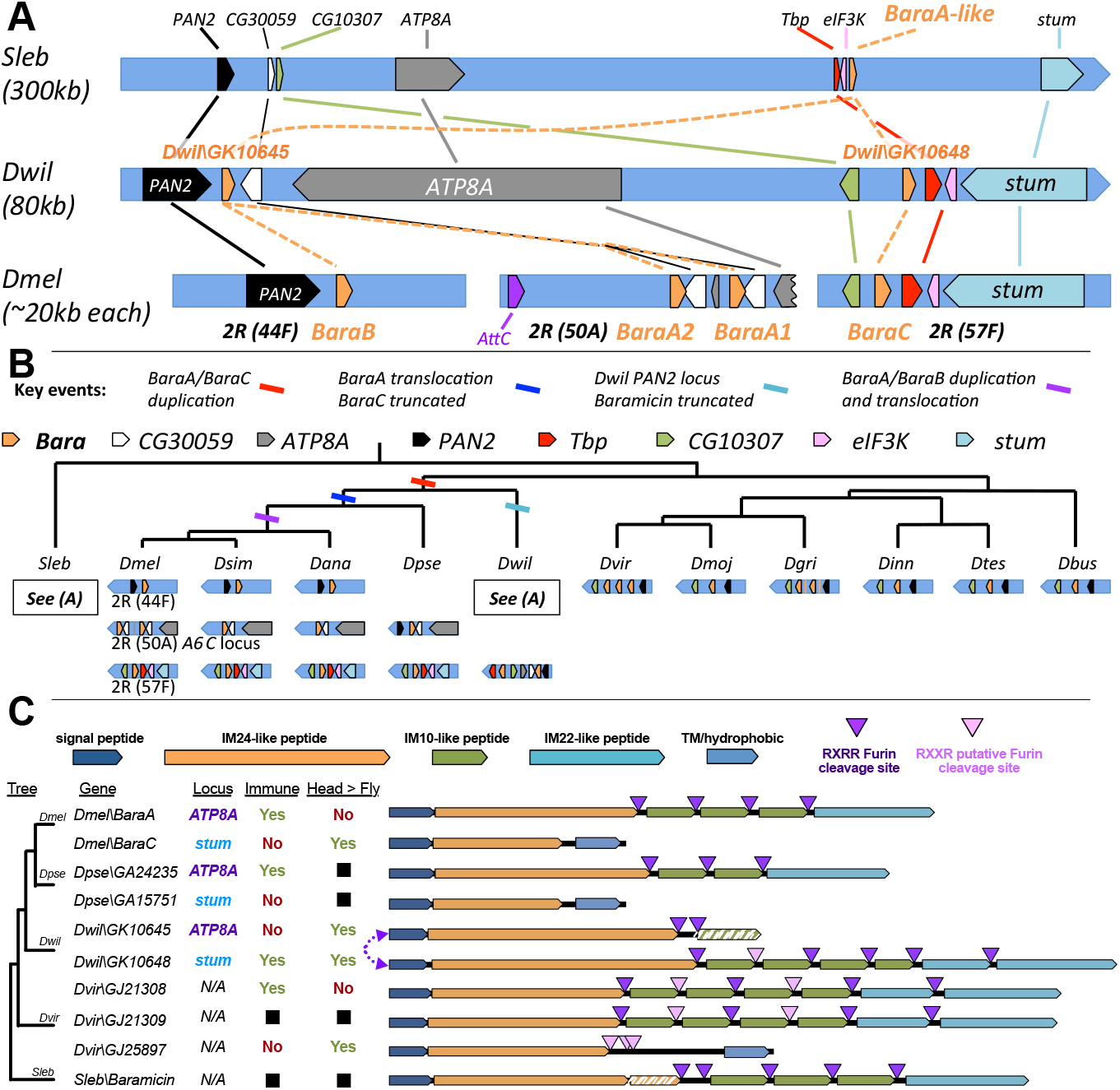
*Baramicin* evolutionary history. A) Detailed map of genomic neighbourhoods in the outgroup drosophilids *S. lebanonensis, D. willistoni,* and *D. melanogaster,* detailing inferred duplication, inversion, and translocation events. Gene names are given as found in *D. melanogaster.* B) Cladogram and genomic loci further detailing the series of events leading to the extant *Baramicin* loci of *Drosophila* species. Loci in *S. lebanonensis* and flies in the subgenus *Drosophila* encode only one *Baramicin* gene, indicating the ancestral drosophilid likely encoded only one *Baramicin.* C) IM24-specific *Baramicins* arose from convergent evolution both in gene structure and expression profile. Genomic loci are described here as *ATP8A* or *stum* reflecting prominent genes neighbouring the *Baramicins* (see Fig. 4A). Expression dynamics relating to immune-induction or enrichment in the head (checked boxes) are shown in Fig. 2 and Fig. S5. The *Baramicin* loci in *D. willistoni* are syntenic with *D. melanogaster,* but evolved in a vice versa fashion (purple arrow). The *D. virilis Baramicins GJ21309* and *GJ25897* are direct sister genes (100% identity at N-terminus).

We originally found outgroup *Baramicins* by reciprocal BLAST searches, and screened *BaraA-like* genes encoding the full suite of Baramicin peptides for immune induction (i.e. encoding IM24, IM10-likes, and IM22: expression in **Fig.** 1). However following genomic synteny analysis, we realized that the *D. willistoni BaraA-like* gene *Dwil\GK10648* is syntenic with the *Dmel\BaraC*locus (**Fig. 4**A), yet this gene is immune-induced (**Fig.** 1D) and retains a *BaraA*-like gene structure (**Fig. 4**C). On the other hand, *Dwil\GKIO645* is found at the locus syntenic with *BaraA,* but has undergone an independent truncation to encode just an IM24 peptide (similar to *Dmel\BaraC*). Thus these two *D. willistoni* genes have evolved similar to *D. melanogaster BaraA/BaraC,* but in a vice versa fashion. This suggests a pattern of convergent evolution with two key points: **i)** the duplication event producing *Dmel\BaraA* and *Dmel\BaraC* originally copied a full-length *BaraA*-like gene to the *BaraC* locus, and **ii)** the derivation of an IM24-specific gene structure has occurred more than once *(Dmel\BaraC*and *Dwil\GKIO645).* Indeed, another independent IM24-specific *Baramicin* gene is present in *D. virilis (Dvir\GJ25897),* which is a direct sister of the *BaraA*-like gene *Dvir\GJ2l3O9* (the signal peptides of these genes is identical at the nucleotide level, and see **Fig. 4**C). Thus *Baramicins* in both *D. willistoni* and *D. virilis* have convergently evolved towards an IM24-specific protein structure resembling *Dmel\BaraC.* We checked the expression of these truncated *Baramicins* in each species upon infection. As was the case for *Dmel\BaraC*, neither gene is immune-induced (**Fig. S5**A-C). Given the glial expression of *Dmel\BaraC,* we reasoned that the heads of adult flies (rich in nerve tissue) should be enriched in *BaraC* compared to whole animals. Indeed we saw a significant enrichment of *BaraC* in the heads of *D. melanogaster* males compared to whole flies, which was not the case for *BaraA* (**Fig. S5**D). When we checked the heads of *D. willistoni* and *D. virilis,* we indeed saw a consistent and significant enrichment in the head for the IM24-specific genes *Dwil\GK1O645* and *Dvir\GJ25897,* while *BaraA*-like genes were more variable in expression (**Fig. S5**E-F).

Genomic synteny shows the gene structure and immune expression of *BaraA* are the ancestral state, and *Dmel\BaraB* and *Dmel\BaraC* are paralogs derived from independent duplication events. Strikingly, we observe a parallel evolution of expression pattern and gene structure in *Baramicins* of *D. willistoni* and *D. virilis.* Moreover these independent IM24-specific *Baramicins* across species are not immune induced, and are enriched in the head. Expression data across genes and species are shown in **Fig.** S5 and summarized in **Fig. 4**C.

### Residue 29 in the IM24 domain evolves in lineage-specific fashions

Thus far we have shown that IM24-specific genes are expressed in the nervous system, yet IM24 is the only peptide domain conserved across all *Baramicin* genes. We were therefore interested to better understand the properties of IM24 to know if any evolutionary patterns might distinguish the IM24 domains of nervous system-expressed genes from IM24 domains of immune-induced genes. We were unable to model the protein satisfactorily with various protein prediction techniques, preventing a 3D comprehension of the IM24 peptide. Therefore we asked if we could highlight any residues in this traditionally immune peptide that might correlate with nervous system or immune-induced gene lineages to better understand what aspect of IM24 contributes to it being retained in neural contexts.

To do this, we screened for positive selection (elevated non-synonymous mutation rate) in the IM24 domain using the HyPhy package implemented in Datamonkey.org (Delport et al. 2010) using separate codon alignments of *Baramicin* IM24 domains beginning at their conserved Q^1^ starting residue. As is recommended with the HyPhy package (Delport et al. 2010), we employed multiple statistical approaches including Likelihood (FEL), Bayesian (FUBAR), and Count-based (SLAC) analyses to ensure patterns in selection analyses were robust to different methods of investigation. Specifically, we used locus-specific alignments (e.g. genes at the *stum* locus in **Fig. 4** B were all analyzed together) to ensure IM24 evolution reflected locus-specific evolution. FEL, FUBAR, and SLAC site-specific analyses each suggest strong purifying selection in many residues of the IM24 domain (p-adj < .05, data in **supplementary data file 3**), agreeing with the general protein structure of IM24 being broadly conserved (**Fig.** 5A). However one residue (site 29) was consistently highlighted as evolving under positive selection using each type of statistical approach for genes located at the Sophophora *ATP8A* locus *(BaraA* genes and *Dwil\GKlO645:* p-adj < .05; **Fig. 5**A). This site is universally Proline in *Baramicin* genes located at the stum locus (*BaraC*-like), in both *D. willistoni Baramicins,* and in the outgroup *S. lebanonensis*, suggesting Proline is the ancestral state. However this residue diverges in both the *BaraA* (commonly Threonine) and *BaraB* (commonly Valine) lineages. We also note that two sites on either side of site 29 (site 27 and site 31) similarly diverge by lineage in an otherwise highly conserved region of the IM24 domain. FUBAR analysis (but not FEL or SLAC) similarly found evidence of positive selection at site 31 in the *BaraA* locus genes (p-adj = .026). Thus this neighbouring site could also be evolving in a non-random fashion. Similar analyses of the *BaraB* and *stum* loci *Baramicins* did not find evidence of site-specific positive selection.

**Figure 5:**
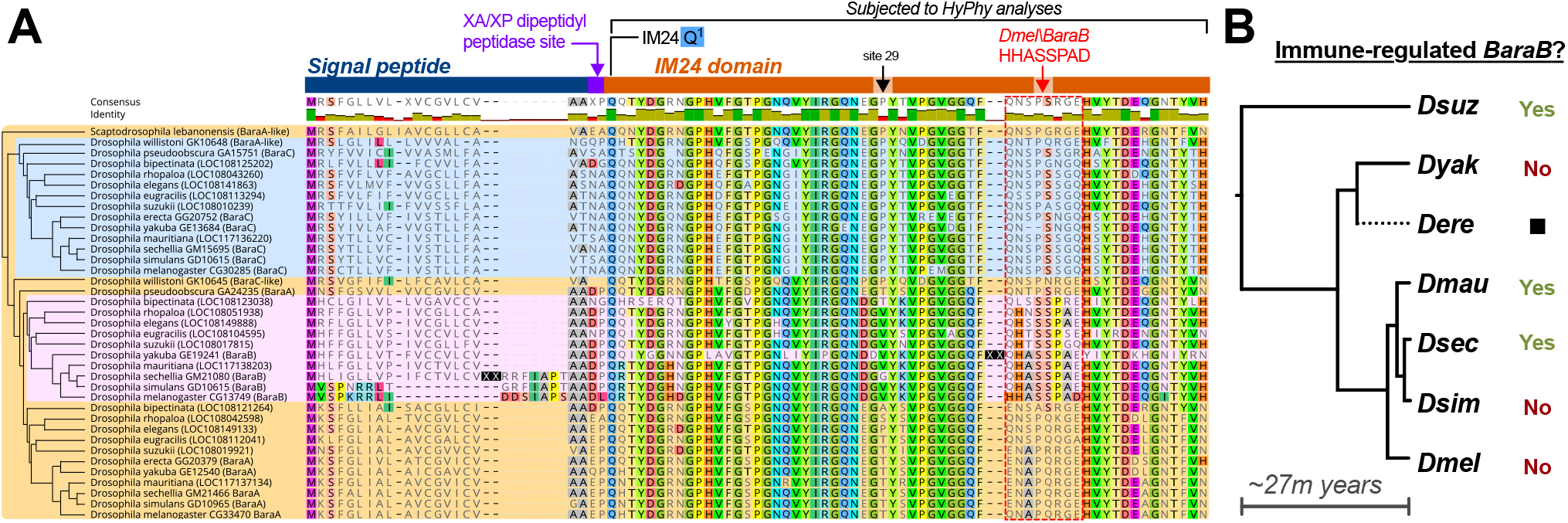
*BaraB* and IM24 rapid evolution. A) Evolution of the signal peptide and core IM24 domain. Residue highlighting indicates agreement with *Dmel\BaraB.* Arrows indicate site 29 and the *D. melanogaster* ^40^HHASSPAD^48^ domain. Insertion events in *D.yakuba* and *D. sechellia BaraB* are denoted as XX to save space. While longer than other Baramicin proteins, the *D. sechellia* signal peptide is predicted to remain functional (SignalP 5.0). The cladogram on the left shows genomic relatedness (by speciation and locus) independent of sequence similarity. Background colouring is included to show birth of novel *Baramicin* loci/lineages. B) *BaraB* immune or non-immune expression by phylogeny. The *D. erecta BaraB* gene is pseudogenized by multiple premature stop codons, and the *D. yakuba* gene is not immune-induced and encodes a 9-residue insertion in the IM24 peptide bordering the HHASSPAD domain (see XX site in A). However the *BaraB* genes of *D. suzukii,* and both *D. mauritiana* and *D. sechellia* remain inducible by infection, while the *BaraB* genes of *D. simulans* and *D. melanogaster* are not and are expressed at very low levels (Fig. S6). This pattern suggests that *BaraB* of *D. melanogaster* (and *D. simulans)* acquired its non-immune role only recently, and is correlated with the loss of the *BaraB* signal peptide.

We highlight site 29 as a key residue in IM24 that diverged in *Baramicin* in lineage-specific fashions. This ancestrally Proline residue has settled on a Threonine in most *BaraA*-like genes of Obscura and Melanogaster group flies, and a Valine in most *BaraB* genes, which are unique to the Melanogaster group. The ancestral Proline residue is found in both *D. willistoni Baramicins*, alongside significant enrichment of both genes in the head, despite only one gene being immune-induced (**Fig.** 4C, **Fig.** S5). Thus it is unclear how this site contributes to tissue-specific *Baramicin* functions, but Threonine and Valine residues evolved in the *BaraA* and *BaraB* lineages.

### Another IM24 domain in Baramicin lineages varies through relaxed selection

Visual inspection of aligned IM24 proteins makes it evident that the overall IM24 domain is broadly conserved, except in sites 40-48 (**Fig. 5**A). However a motif aligned to residues ^40^HHASSPAD^48^ of *Dmel\BaraB* departs in lineage-specific fashions; the three C-terminal residues of this motif are also diagnostic of each gene lineage *(BaraA, BaraB,* and *BaraC*have RGE, PXE, or (S/N)GQ respectively; **Fig. 5**A). However even with additional branch-site selection analyses (aBSREL and BUSTED (Murrell et al. 2015)), we found no evidence of positive selection at this motif, and in fact many residues also failed to show evidence of purifying selection. For instance, six of nine sites in the *BaraA* locus analysis failed to reach significance (p < .05) for purifying selection in SLAC analysis (supplementary data file 3).

Given an absence of positive selection, and many residues failing to reach significance for purifying selection at the residue 40-48 motif, we this motif is diversifying due to drift effects through relaxed selection. The high conservation of IM24 residues up- and downstream of this motif is nevertheless striking. One possible explanation may be that these residues act as a linker between the two functional parts of IM24 up- and downstream of residues 40-48. Perhaps supporting this interpretation, we found that *D. yakuba BaraB* independently lost immune induction alongside an insertion at site 40 (**Fig.** 5). Such speculation awaits validation by robust protein modelling efforts.

### Overt structural change best explains Baramicin loss of immune induction

We found that site 29 evolves rapidly in *Baramicin* lineages, but this site is common to both immune and non-immune Baramicin genes (e.g. in *D. willistoni*). Thus IM24 sequence variation does not explain why IM24-specific Baramicins lose immune inducibility. However within the *BaraA/BaraB* lineage, we observed that *BaraB* genes commonly encode Valine at IM24 site 29, compared to Threonine in *BaraA*. As the *BaraB* locus is derived from an ancestral immune-induced (**Fig.** 4A), it is unclear if other *BaraB* genes are immune-inducible, and thus what IM24 evolutionary patterns (like Valine at site 29) might predict *BaraB* functional divergence.

We therefore performed infection experiments in diverse species across the Melanogaster group to see if their *BaraB* genes had similarly lost immune induction (see **Fig.** S6 for qPCR data). Surprisingly, we found that the non-immune expression of *Dmel\BaraB* is extremely recent, as Melanogaster sister species like *D. sechellia* and *D. mauritiana* encode immune inducible *BaraB* loci (summary in **Fig.** 5B). However, we also found that *D. simulans BaraB* lacked immune induction, despite *D. simulans* being most closely related to *D. sechellia* (Chakraborty et al. 2021). Thus IM24 sequence evolution does not predict immune induction.

This drew our attention to the overall protein structure of the various extant *BaraB* genes. A striking feature of the Dmel\BaraB protein is the absence of a functional signal peptide (**Fig.** 2B). This signal peptide sequence is conserved in all *Baramicin* lineages, except in *Dmel/BaraB* and also *Dsim/BaraB.* Indeed despite *D. simulans* being more closely related to *D. sechellia* and *D. mauritiana,* both *Dmel\BaraB* and *Dsim\BaraB* encode a homologous N-terminus of parallel length (**Fig.** 5A). We also found that *Dyak\BaraB* is not immune-responsive, but note that *D.yakuba* has an insertion upstream of residue 40 that elongates the IM24 domain (**Fig.** 5B black X boxes), and its sister species *D. erecta* encodes multiple indels and premature stops suggesting *BaraB* is pseudogenized in this lineage. Loss of the BaraB signal peptide is therefore more specifically associated with loss of immune expression in the Melanogaster species complex (*D. simulans, D. sechellia, D. mauritiana,* and *D. melanogaster).* The last common ancestor of *D. simulans, D. sechellia,* and *D. mauritiana* is estimated to be just ~250,000 years ago, and these species diverged from *D. melanogaster* ~3 million years ago (Chakraborty et al. 2021). The fact that *D. simulans* uniquely encodes this *Dmel\BaraB-like* sequence suggests it was either introgressed from one species to the other prior to the complete development of hybrid inviability, or reflects incomplete lineage sorting of this locus in the Melanogaster species complex.

Overall, we find no evidence to suggest IM24 domain sequence has evolved drastically to allow for function in the nervous system. The Proline residue at site 29 of neural *Baramicin* genes appears to be the ancestral state. Rather than small sequence changes, overt structural changes like truncation to focus on the IM24 domain and loss of a signal peptide in *Dmel\BaraB* are associated with *Baramicins* expressed in the nervous system. After duplication, *Baramicin* daughter lineages have repeatedly derived neural-specific expression through subfunctionalization of the IM24 domain from the overall precursor protein. Importantly, this finding suggests that the ancestral *Baramicin* encoded peptides with distinct roles in either the immune response or the nervous system.

## Discussion

Recent studies have suggested AMP genes drive behaviour or disease through interactions with the nervous system. Many of these genes encode polypeptides with multiple mature products. To date, little attention has been paid to AMP genes in these neural contexts at the level of the sub-peptides they encode. Here we demonstrate that the *Baramicin* antimicrobial peptide gene of *Drosophila* ancestrally encodes distinct peptides involved with either the nervous system or the immune response. Given the ancestral *Baramicin* encoded all Baramicin peptide domains, this antimicrobial peptide gene has always produced sub-peptides that interact with either the nervous system (IM24) or invading pathogens (IM10-like, IM22). These peptides are maturated from a longer precursor protein, accomplished via furin cleavage. Importantly, this suggests that AMP genes can mediate these distinct neural and immune roles via specialized sub-peptides, and not necessarily due to dual action of a single peptide.

The goal of generating different gene products is commonly accomplished through alternate transcript isoforms. For instance, the insect *doublesex* gene determines sexual differentiation, with male- or female-specific isoforms mediated by alternative splicing. Select hymenopteran lineages have even lost coding sequence for the male-specific exon (Baral et al. 2019). These context-specific isoforms are similar in theme to loss of immune-relevant peptides in non-immune *Baramicin* genes, except that in *Baramicins* this is achieved through gene duplication. Moreover the ‘protein operon’ structure of immune-induced *Baramicins* can act as a mechanism to allow peptides with distinct roles to be produced simultaneously from a single mRNA transcript.

There is building evidence that immune-induced AMPs and AMP-like genes affect the nervous system. Loss of Metchnikowin protects flies from neurodegeneration after traumatic brain injury (Swanson et al. 2020), *Induced by infection (IBIN)* regulates behavioural changes in flies after seeing parasitoid wasps (Ebrahim et al. 2021), epidermal nematode AMPs trigger motor neuron autophagy (Lezi et al. 2018) and sleep (Sinner et al. 2021) after infection, and loss of *Diptericin B* produced by the fat body leads to memory deficits in *Drosophila* (Barajas-azpeleta et al. 2018). This last example is intriguing, as *Diptericin B* also encodes a polypeptide maturated by furin cleavage, and its effect on memory was derived from peptide secreted into the hemolymph by the fat body and not from neural expression. Similarly, we recently found that *BaraA* deletion causes infected flies to display an erect wing behavioural phenotype, which was independent of active infection, and could be rescued by priming the hemolymph with *BaraA* expressed by the fat body (Hanson et al. 2021). Thus some component of *BaraA* likely interacts with some host target(s) to prevent this behaviour during the immune response. The present study suggests this could be due to action of BaraA IM24, given evidence for IM10-like peptides as antifungal peptides (Hanson et al. 2021), and loss of these peptides in nervous system-specific *Baramicins.* However AMPs and neuropeptides have many similar features, including cationic charge and amphipathicity (Brogden et al. 2005). Thus while we present evidence that IM24 of different *Baramicin* genes underlies *Baramicin* interactions with the nervous system, we cannot exclude the possibility that IM24 is also antimicrobial.

One human AMP recently implicated in chronic neuroinflammatory disease is the Cathelicidin LL-37 (Lee et al. 2015; De Lorenzi et al. 2017; Moir et al. 2018). Like *Baramicin,* the *Cathelicidin* gene family is unified by its N-terminal “Cathelin” domain. However to date no one has described antimicrobial activity of the Cathelin domain in vitro (Zanetti 2005). Instead, *Cathelicidin* research has focused almost exclusively on the mature peptide LL-37 at the C-terminus of mammalian *Cathelicidin* genes. Reflecting on *Baramicin* evolution and the implication of *Cathelicidin* in neurodegenerative diseases, what does the Cathelin domain do? While this study was conducted in fruit flies, we hope we have emphasized the importance of considering each peptide of AMP genes for in vivo function. This is relevant to neural processes even if the gene is typically thought of for its role in innate immunity. Indeed, recent studies of *Drosophila* AMPs have emphasized that in vitro activity does not always predict the interactions that these genes can have in vivo (Clemmons et al. 2015; Hanson, Dostálová, et al. 2019). Care should be taken not to conflate in vitro activity with realized in vivo function. Most studies focus on AMPs specifically in an immune role, but this is akin to ‘looking for your keys under the streetlight.’ To understand AMP functions in vivo, genetic approaches will be necessary that allow a more global view of gene function.

In summary, we reflect on the structural characteristic of AMP genes through the lens of *Baramicins.* We found that one sub-peptide of the immune-induced *Baramicin* ancestor is readily adapted for functions relating to the nervous system. Meanwhile, other sub-peptides known to suppress fungi are repeatedly lost in daughter genes that lack immune inducibility, suggesting they are irrelevant to neural functions. As AMP genes commonly encode polypeptides maturated by furin cleavage (including *Baramicin),* it will be interesting to consider the functions of AMP genes in neural processes not simply at the level of the gene, but at the level of the mature peptides produced by that gene. This consideration may explain how some polypeptide immune effectors play dual roles in disparate contexts.

## Data availability statement

All data are included within the manuscript and supplementary files.

## Acknowledgements

We would like to thank Maria Litovchenko for advice, Ana Marija Jakšić for generously providing DGRP flies, Huang et al. (Huang et al. 2020) for collaborative cooperation, Brian McCabe for consultation, and Florent Masson, Hannah Westlake, and the anonymous reviewers and the editors at MBE for commentary on our initial manuscript. This research was supported by Sinergia grant CRSII5_186397 awarded to Bruno Lemaitre. The *BaraB^LC1^* and *BaraB^LC4^* mutations were graciously provided by Steven Wasserman and generated by Lianne Cohen, who we also thank for their critical involvement in characterizing *Baramicin A.*

## Materials and Methods

### DGRP population screening and bioinformatics analyses

Genomic sequence data were downloaded from GenBank default reference assemblies and Kim et al. (Kim et al. 2021), and DGRP sequence data from http://dgrp2.gnets.ncsu.edu/ (Mackay et al. 2012). Sequence comparisons and alignment figures were prepared using Geneious R10 (Kearse et al. 2012), Prism 7, and Inkscape. Alignments were performed using MUSCLE or MAFFT followed by manual curation, and phylogenetic analyses were performed to validate sequence patterns using the Neighbour Joining, PhyML, RaxML, and MrBayes plugins in Geneious. *BaraA* copy number screening was performed using primers specific to the duplication and *CG30059* control primers for DNA extraction (primers in supplementary data file 5). We found a significant correlation between *BaraA* PCR status and variant sites starting at 2R_9293471_SNP and extending to 2R_9293576_SNP (Pearson’s correlation matrix: 0.0001 < p-value < 0.005 at all nine sites), however the status of genetic variants at this site is poorly resolved and so we cannot be confident that our ~14% estimate for the *BaraA* duplication in the DGRP would hold true if long-read sequencing was employed. DGRP annotation of the *BaraA* locus in Fig. S1 was generated using the UCSC *D. melanogaster* DGRP2 genome browser. Selection analyses were performed using the HyPhy package implemented in datamonkey.org (Delport et al. 2010). Codon alignments of the IM24 domain used in **Fig. 5**A are included as a .fasta file in supplementary data file 3 alongside outputs from FEL, FUBAR, SLAC, and aBSREL selection analyses.

### Fly genetics

The *BaraB^LC1^* and *BaraB^LC4^* mutations were generated using CRISPR with two gRNAs and an HDR vector by cloning 5’ and 3’ region-homologous arms into the pHD-DsRed vector, and consequently *ΔBaraB* flies express DsRed in their eyes, ocelli, and abdomen. The following PAM sites were used for CRISPR bordering the *BaraB* region. Slashes indicate the cut site: 5’: GCGGGCAACAGATGTGTTCA/GGG 3’: GTCCATTGCTTATTCAAAAA/TGG. These mutants were generated in the laboratory of Steve Wasserman by Lianne Cohen, who graciously allowed their use in this study. All fly stocks including Gal4 and RNAi lines are listed in **supplementary data file 4**. Experiments were performed at 25°C unless otherwise indicated. When possible, genetic crosses of 6-8 males and 6-8 females were performed in both directions to test for an effect of the X or Y chromosomes on *BaraB*-mediated lethality; crosses in both directions yielded similar results in all cases and reported data are pooled results. Fly diet consisted of a nutrient-rich lab standard food: 3.72g agar, 35.28g cornmeal, 35.28g yeast, 36mL grape juice, 2.9mL propionic acid, 15.9mL moldex, and H2O to 600mL.

### Infection experiments

Bacteria and yeast were grown to mid-log phase shaking at 200rpm in their respective growth media (LB, BHI, or YPG) and temperature conditions, and then pelleted by centrifugation to concentrate microbes. Resulting cultures were diluted to OD = 200 at 600nm before infections to measure gene expression. The following microbes were grown at 37°C: *Escherichia coli strain 1106* (LB) and *Candida albicans* (YPG). *Micrococcus luteus* was grown at 29°C in LB. For **Fig.** 1 and S2, pooled fly samples were collected either 6 hours post-infection (*E. coli)* or 24 hours post-infection (*C. albicans, M. luteus)* prior to RNA extraction on pools of 5 adult males. These timepoints correspond to the maximal expression inputs of the Imd (6hpi) or Toll (24hpi) NF-κB signalling pathways, which are most specifically induced by Gram-negative bacteria (Imd) or Gram-positive bacteria or fungi (Toll) (Lemaitre et al. 1997). Flies were pricked in the thorax as described in (Hanson, Dostálová, et al. 2019).

RNA extractions were performed using TRIzol™, Ambion DNAse treatment, and PrimeScript RT according to manufacturer’s protocols. RT-qPCR was performed using PowerUP SYBR Green master mix with primers listed in supplementary data file 5. Gene expression differences were analyzed using the PFAFFL method (Pfaffl 2001). For gene expression experiments requiring dissection of heads, pools of 20 males were used for either whole flies or heads dissected in ice-cold PBS and transferred immediately to a tube kept on dry ice.

### Selection analysis using HyPhy package

Codon aligned nexus tree files were generated using either the Neighbourjoining (1000 bootstraps) or PhyML (100 bootstraps) methods including proteins beyond those shown in **Fig. 5**. These tree files were analyzed using the HyPhy package with only 174nt pertaining to just the IM24 domain codons included. The cladogram in **Fig. 5**A is manually drawn from known species divergences (Kim et al. 2021). Use of either tree building method was chosen for convenience to best reflect known lineage sorting, as use of just 174nt was too information poor to resolve exact phylogenetic relatedness reliably. Tree files were qualitatively screened to ensure topologies broadly matched known species sortings, and thus ensure only relevant comparisons were made given the genomic synteny analysis in **Fig. 4** is principally informative of true gene lineages. HyPhy analyses were run separately for each *Baramicin* lineage within their clade, defined by genomic synteny; i.e. based on locus (e.g. ATP8A locus), and not considering convergent gene structures. We used three site-specific analyses (FEL, FUBAR, and SLAC) that use three independent statistical approaches (Likelihood, Bayesian, and Count-based methods respectively). We also employed both BUSTED and aBSREL branch-site analyses, which are likelihood methods that differ in their approach of testing whole-phylogeny selection or branch-specific comparisons respectively; an anology might be performing analysis of variance (ANOVA) at the level of the entire ANOVA, or comparing multiple groups against each other and subsequently using multiple test correction. Each tree was rooted using the *Scaptodrosophila lebanonensis Baramicin* as an outgroup with ancestral characteristics; we did not include *Baramicins* of the subgenus Drosophila as including these resulted in long-branch attraction of the Willistoni group *Baramicins* to subgenus Drosophila lineages, which would confound relevant phylogenetic comparisons. When applicable, all internal branches were assessed for potential selection. For *Baramicins* of the *ATP8A* locus, one site (site 29) was highlighted as experiencing positive selection using FEL, FUBAR, and SLAC analyses (p-adj = .011, .013, and .039 respectively). Additionally, site 31 was also highlighted by FUBAR (p-adj = .026), but not FEL or SLAC analyses (p-adj > .05). BUSTED analysis also supported diversifying selection in the *BaraA* lineage (ATP8A locus, LRT p-adj = .008), indicating at least one site on at least one test branch has experienced diversifying selection within the ATP8A lineage. The aBSREL branchsite analysis specifically highlights the branch distinguishing the Willistoni group *Baramicins* from the other Sophophora species (p-adj = .0045), suggesting variation between these branches drives the signals of diversifying selection in the BUSTED analysis. This result is intuitive, as we find a parallel but opposite evolution of Baramicin protein structure in *Baramicins* of the *ATP8A* locus in *D. willistoni* compared with *Baramicins* of other Sophophora species. Furthermore, in whole-gene phylogenies, both *D. willistoni Baramicins* cluster together, supporting the notion that these two daughter genes have evolved independent from the selection that shaped the orthologues of *Dmel\BaraA* and *Dmel\BaraC,* also seen in qPCR data that showed both genes were significantly enriched in the head (**Fig.** S6). This phylogenetic clustering of the two *D. willistoni Baramicins* holds true when additional *Baramicins* from recently sequenced genomes of the Willistoni group are included (from (Kim et al. 2021) in supplementary data file 3), indicating this is characteristic of the Willistoni group lineage and not specific to *D. willistoni.*

