## supplementary data file 3 for "Repeated truncation of a modular antimicrobial peptide gene for neural context": Datamonkey Adaptive Evolution Server.html

Methods and Tools

aBSREL
SpiderMonkey/BGM
BUSTED
Contrast-FEL
FADE
FEL
FUBAR
GARD
HIV-TRACE
MULTI-HIT
MEME
RELAX
SLAC
All Methods

Job Queue
Usage statistics

API

API Info
Get API Key
Check Key Status

Citations
Help
COVID-19
Blog
 Classic

Methods and Tools

aBSREL
BUSTED
FADE Beta
FEL
FUBAR
GARD
HIV-TRACE
MEME
RELAX
SLAC
All Methods

Job Queue
Usage statistics
Citations
Help
 Classic

- summary
- tree
- table
- model fits

×Close**Error!**

##### adaptive Branch Site REL results summary

INPUT DATA |6095a660238adf71a515dce5|23 sequences |58 sites

Export

- Original file
- Analysis log
- Save JSON
- View JSON

###### Alignment viewer

×

Close

aBSREL **found evidence** of episodic diversifying selection on **1** out of **43** branches in your phylogeny.

A total of **43** branches were formally tested for diversifying selection. Significance was assessed using the Likelihood Ratio Test at a threshold of p ≤ 0.05, after correcting for multiple testing. Significance and number of rate categories inferred at each branch are provided in the detailed results table.

---

See here for more information about this method.  
Please cite PMID 25697341 if you use this result in a publication, presentation, or other scientific work.

###### Tree summary

| ω rate classes | # of branches | % of branches | % of tree length | # under selection |
| --- | --- | --- | --- | --- |
| 1 | 38 | 88% | 0.023% | 0 |
| 2 | 5 | 12% | 100% | 1 |

This table contains a summary of the inferred aBSREL model complexity. Each row provides information about the branches that were best described by the given number of ω rate categories.

###### Fitted tree

Options

- Models
- Full adaptive model
- Baseline MG94xREV

- Hide Legend
- GrayScale

Export 

- PNG
- SVG
- Newick File

00.010.10.512510ωLength = 0.04204643664228217Length = 0.05515285879763906Length = 0.02977772686265093Length = 0Length = 0.005959293660541262Length = 0.01812972194156237Length = 0.0147465975435195Length = 0.009658825134148272Length = 0.02070040883591088Length = 0.01743276220434956Length = 0.1025948919625218Length = 0.03150205394158095Length = 0.04359931517678639Length = 0.04756803423237001Length = 0.09063236712019877Length = 0.01761304822035577Length = 0.03330865114039226Length = 0.1109256507067189Length = 0Length = 0.1549972431366071Length = 0.07930281793902925Length = 0.06530527172814865Length = 0.04618248457816571Length = 0Length = 0.2106458167000396Length = 0.09433724526067282Length = 0.07833383753169616Length = 0.2076525437615734Length = 0.157148884843192Length = 0.033490228984756Length = 0.09466593222071644Length = 0.1885923485814873Length = 0Length = 0.005393601662584954Length = 303.1231304796415Length = 0.03529574502323375Length = 0.0095684171729065Length = 0.04621317206396958Length = 0.02222800688516957Length = 0.01159562376737461Length = 0.1897535427802949Length = 0.1094667291995619Length = 9026.448840319466DINS\_BARACDTRO\_BARACDWIL\_GK10645DPAU\_BARAC\_STR\_L06\_DPAU\_BARAC\_STR\_L12\_DPSE\_GA24235DSER\_XM\_020962164DBIP\_XM\_017235265DANA\_GF13690DRHO\_XM\_017120938DFIC\_XM\_017195800DEUG\_XM\_017221794DELE\_XM\_017276537DTAK\_XM\_017145699DSUZ\_XM\_017088027DBIA\_XM\_017101808DMEL\_BARAA\_CG33470\_AND\_CG18279\_DSIM\_GD10965\_W501\_DSEC\_GM21466\_XM\_002033668DMAU\_XM\_033298434\_LOC117137134DERE\_GG20379DYAK\_GE12540SLEB\_XM\_030519534

###### Detailed results

| Name | B | LRT | Test p-value | Uncorrected p-value | ω distribution over sites |
| --- | --- | --- | --- | --- | --- |
| Node12 | 0.0000 | 11.6549 | 0.0435 | 0.0010 | ω_1_ = 1.00 (94%) ω_2_ = 14400 (6.0%) |
| DANA\_GF13690 | 0.0000 | 5.4521 | 0.9892 | 0.0236 | ω_1_ = 0.771 (97%) ω_2_ = 225 (3.4%) |
| DBIA\_XM\_017101808 | 0.0000 | 0.0000 | 1.0000 | 1.0000 | ω_1_ = 0.117 (100%) |
| DBIP\_XM\_017235265 | 0.0000 | 0.0000 | 1.0000 | 1.0000 | ω_1_ = 0.0421 (100%) |
| DELE\_XM\_017276537 | 0.0000 | 0.0000 | 1.0000 | 1.0000 | ω_1_ = 0.0635 (100%) |
| DERE\_GG20379 | 0.0000 | 0.0000 | 1.0000 | 1.0000 | ω_1_ = 0.0424 (100%) |
| DEUG\_XM\_017221794 | 0.0000 | 0.0000 | 1.0000 | 1.0000 | ω_1_ = 0.0433 (100%) |
| DFIC\_XM\_017195800 | 0.0000 | 0.0000 | 1.0000 | 1.0000 | ω_1_ = 0.142 (100%) |
| DINS\_BARAC | 0.0000 | 0.4061 | 1.0000 | 0.3497 | ω_1_ = 10000000000 (100%) |
| DMAU\_XM\_033298434\_LOC117137134 | 0.0000 | 0.0000 | 1.0000 | 1.0000 | ω_1_ = 0.486 (100%) |
| DMEL\_BARAA\_CG33470\_AND\_CG18279\_ | 0.0000 | 0.0000 | 1.0000 | 1.0000 | ω_1_ = 0.505 (100%) |
| DPAU\_BARAC\_STR\_L06\_ | 0.0000 | 0.0000 | 1.0000 | 1.0000 | ω_1_ = 0.00 (100%) |
| DPAU\_BARAC\_STR\_L12\_ | 0.0000 | 0.0000 | 1.0000 | 1.0000 | ω_1_ = 1.00 (100%) |
| DPSE\_GA24235 | 0.0000 | 0.0000 | 1.0000 | 1.0000 | ω_1_ = 0.0642 (100%) |
| DRHO\_XM\_017120938 | 0.0000 | 0.0000 | 1.0000 | 1.0000 | ω_1_ = 0.190 (100%) |
| DSEC\_GM21466\_XM\_002033668 | 0.0000 | 0.0000 | 1.0000 | 1.0000 | ω_1_ = 1.00 (100%) |
| DSER\_XM\_020962164 | 0.0000 | 0.0000 | 1.0000 | 1.0000 | ω_1_ = 0.427 (100%) |
| DSIM\_GD10965\_W501\_ | 0.0000 | 0.0000 | 1.0000 | 1.0000 | ω_1_ = 0.159 (100%) |
| DSUZ\_XM\_017088027 | 0.0000 | 0.0000 | 1.0000 | 1.0000 | ω_1_ = 0.141 (100%) |
| DTAK\_XM\_017145699 | 0.0000 | 0.0000 | 1.0000 | 1.0000 | ω_1_ = 0.198 (100%) |
| DTRO\_BARAC | 0.0000 | 0.0000 | 1.0000 | 1.0000 | ω_1_ = 0.201 (100%) |
| DWIL\_GK10645 | 0.0000 | 0.0000 | 1.0000 | 1.0000 | ω_1_ = 0.0757 (100%) |
| DYAK\_GE12540 | 0.0000 | 0.0000 | 1.0000 | 1.0000 | ω_1_ = 0.130 (100%) |
| Node14 | 0.0000 | 0.0000 | 1.0000 | 1.0000 | ω_1_ = 0.00 (100%) |
| Node16 | 0.0000 | 0.0000 | 1.0000 | 1.0000 | ω_1_ = 0.0258 (100%) |
| Node17 | 0.0000 | 0.0000 | 1.0000 | 1.0000 | ω_1_ = 0.0376 (100%) |
| Node20 | 0.0000 | 3.4672 | 1.0000 | 0.0655 | ω_1_ = 0.00 (96%) ω_2_ = 18.4 (3.8%) |
| Node21 | 0.0000 | 0.0000 | 1.0000 | 1.0000 | ω_1_ = 1.00 (100%) |
| Node24 | 0.0000 | 0.0000 | 1.0000 | 1.0000 | ω_1_ = 0.00 (100%) |
| Node26 | 0.0000 | 0.0000 | 1.0000 | 1.0000 | ω_1_ = 1.00 (100%) |
| Node28 | 0.0000 | 0.0000 | 1.0000 | 1.0000 | ω_1_ = 0.386 (100%) |
| Node29 | 0.0000 | 0.1495 | 1.0000 | 0.4182 | ω_1_ = 10000000000 (100%) |
| Node3 | 0.0000 | 1.8525 | 1.0000 | 0.1535 | ω_1_ = 0.0125 (86%) ω_2_ = 100000 (14%) |
| Node31 | 0.0000 | 0.0000 | 1.0000 | 1.0000 | ω_1_ = 0.00 (100%) |
| Node34 | 0.0000 | 0.0000 | 1.0000 | 1.0000 | ω_1_ = 0.00 (100%) |
| Node35 | 0.0000 | 1.0099 | 1.0000 | 0.2440 | ω_1_ = 10000000000 (100%) |
| Node37 | 0.0000 | 0.0000 | 1.0000 | 1.0000 | ω_1_ = 0.00 (100%) |
| Node39 | 0.0000 | 0.0000 | 1.0000 | 1.0000 | ω_1_ = 0.00 (100%) |
| Node42 | 0.0000 | 0.0000 | 1.0000 | 1.0000 | ω_1_ = 0.128 (100%) |
| Node5 | 0.0000 | 0.0000 | 1.0000 | 1.0000 | ω_1_ = 0.00 (100%) |
| Node7 | 0.0000 | 0.8331 | 1.0000 | 0.2701 | ω_1_ = 10000000000 (100%) |
| Node9 | 0.0000 | 3.7017 | 1.0000 | 0.0580 | ω_1_ = 0.0535 (96%) ω_2_ = 100000 (4.2%) |
| SLEB\_XM\_030519534 | 0.0000 | 0.0000 | 1.0000 | 1.0000 | ω_1_ = 0.0212 (100%) |

###### aBSREL Site Proportion Chart

×

###### ω distribution

### **DANA\_GF13690**

SVG PNG

Neutrality (ω=1)ω0.000010.00010.0010.010.1110100100010000Proportion of sites0%10%20%30%40%50%60%70%80%90%100%

Close

###### Model fits

| Model | AIC_C_ | log L | Parameters |
| --- | --- | --- | --- |
| Nucleotide GTR | 3702.90 | -1799.78 | 51 |
| Baseline MG94xREV | 3393.49 | -1588.55 | 100 |
| Full adaptive model | 3323.58 | -1541.81 | 110 |

This table reports a statistical summary of the models fit to the data. Here, **Baseline MG94xREV** refers to the MG94xREV baseline model that infers a single ω rate category per branch. **Full adaptive model** refers to the adaptive aBSREL model that infers an optimized number of ω rate categories per branch.

×

###### Error

This is my error message

Close

Datamonkey is funded jointly by MIDAS and NIH award R01 GM093939
