## supplementary data file 3 for "Repeated truncation of a modular antimicrobial peptide gene for neural context": datamonkey.js

```
!function(e){function t(t){for(var n,i,l=t[0],f=t[1],c=t[2],m=0,d=[];m<l.length;m++)i=l[m],Object.prototype.hasOwnProperty.call(r,i)&&r[i]&&d.push(r[i][0]),r[i]=0;for(n in f)Object.prototype.hasOwnProperty.call(f,n)&&(e[n]=f[n]);for(s&&s(t);d.length;)d.shift()();return a.push.apply(a,c||[]),o()}function o(){for(var e,t=0;t<a.length;t++){for(var o=a[t],n=!0,l=1;l<o.length;l++){var f=o[l];0!==r[f]&&(n=!1)}n&&(a.splice(t--,1),e=i(i.s=o[0]))}return e}var n={},r={0:0},a=[];function i(t){if(n[t])return n[t].exports;var o=n[t]={i:t,l:!1,exports:{}};return e[t].call(o.exports,o,o.exports,i),o.l=!0,o.exports}i.m=e,i.c=n,i.d=function(e,t,o){i.o(e,t)||Object.defineProperty(e,t,{enumerable:!0,get:o})},i.r=function(e){"undefined"!=typeof Symbol&&Symbol.toStringTag&&Object.defineProperty(e,Symbol.toStringTag,{value:"Module"}),Object.defineProperty(e,"__esModule",{value:!0})},i.t=function(e,t){if(1&t&&(e=i(e)),8&t)return e;if(4&t&&"object"==typeof e&&e&&e.__esModule)return e;var o=Object.create(null);if(i.r(o),Object.defineProperty(o,"default",{enumerable:!0,value:e}),2&t&&"string"!=typeof e)for(var n in e)i.d(o,n,function(t){return e[t]}.bind(null,n));return o},i.n=function(e){var t=e&&e.__esModule?function(){return e.default}:function(){return e};return i.d(t,"a",t),t},i.o=function(e,t){return Object.prototype.hasOwnProperty.call(e,t)},i.p="";var l=window.webpackJsonp=window.webpackJsonp||[],f=l.push.bind(l);l.push=t,l=l.slice();for(var c=0;c<l.length;c++)t(l[c]);var s=f;a.push([469,1]),o()}({230:function(e,t,o){var n=o(177),r=o(505);(t=n(!1)).push([e.i,"@import url(https://fonts.googleapis.com/css?family=Montserrat:400,400i,500,500i,600,600i,700,700i,800,800i,900,900i|Noto+Sans:400,400i,700,700i);"]),t.i(r),t.push([e.i,"@import url(https://fonts.googleapis.com/css?family=Exo+2:100,100i,200,200i,300,300i,400,400i,500,500i,600,600i,700,700i,800,800i,900,900i|Exo:100,100i,200,200i,300,300i,400,400i,500,500i,600,600i,700,700i,800,800i,900,900i|Montserrat:100,100i,200,200i,300,300i,400,400i,500,500i,600,600i,700,700i,800,800i,900,900i|Noto+Sans:400,400i,700,700i|IBM+Plex+Mono:400,500,600,700);"]),t.push([e.i,'/*!\n * Bootstrap v4.6.0 (https://getbootstrap.com/)\n * Copyright 2011-2021 The Bootstrap Authors\n * Copyright 2011-2021 Twitter, Inc.\n * Licensed under MIT (https://github.com/twbs/bootstrap/blob/main/LICENSE)\n */:root{--blue: #007bff;--indigo: #6610f2;--purple: #6f42c1;--pink: #e83e8c;--red: #dc3545;--orange: #fd7e14;--yellow: #ffc107;--green: #28a745;--teal: #20c997;--cyan: #17a2b8;--white: #fff;--gray: #6c757d;--gray-dark: #343a40;--primary: #00a99d;--secondary: #6c757d;--success: #28a745;--info: #17a2b8;--warning: #ffc107;--danger: #dc3545;--light: #f8f9fa;--dark: #343a40;--breakpoint-xs: 0;--breakpoint-sm: 576px;--breakpoint-md: 768px;--breakpoint-lg: 992px;--breakpoint-xl: 1200px;--font-family-sans-serif: -apple-system, BlinkMacSystemFont, "Segoe UI", Roboto, "Helvetica Neue", Arial, "Noto Sans", "Liberation Sans", sans-serif, "Apple Color Emoji", "Segoe UI Emoji", "Segoe UI Symbol", "Noto Color Emoji";--font-family-monospace: SFMono-Regular, Menlo, Monaco, Consolas, "Liberation Mono", "Courier New", monospace}*,*::before,*::after{box-sizing:border-box}html{font-family:sans-serif;line-height:1.15;-webkit-text-size-adjust:100%;-webkit-tap-highlight-color:rgba(0,0,0,0)}article,aside,figcaption,figure,footer,header,hgroup,main,nav,section{display:block}body{margin:0;font-family:-apple-system,BlinkMacSystemFont,"Segoe UI",Roboto,"Helvetica Neue",Arial,"Noto Sans","Liberation Sans",sans-serif,"Apple Color Emoji","Segoe UI Emoji","Segoe UI Symbol","Noto Color Emoji";font-size:1rem;font-weight:400;line-height:1.5;color:#212529;text-align:left;background-color:#fff}[tabindex="-1"]:focus:not(:focus-visible){outline:0 !important}hr{box-sizing:content-box;height:0;overflow:visible}h1,h2,h3,h4,h5,h6{margin-top:0;margin-bottom:.5rem}p{margin-top:0;margin-bottom:1rem}abbr[title],abbr[data-original-title]{text-decoration:underline;text-decoration:underline dotted;cursor:help;border-bottom:0;text-decoration-skip-ink:none}address{margin-bottom:1rem;font-style:normal;line-height:inherit}ol,ul,dl{margin-top:0;margin-bottom:1rem}ol ol,ul ul,ol ul,ul ol{margin-bottom:0}dt{font-weight:700}dd{margin-bottom:.5rem;margin-left:0}blockquote{margin:0 0 1rem}b,strong{font-weight:bolder}small{font-size:80%}sub,sup{position:relative;font-size:75%;line-height:0;vertical-align:baseline}sub{bottom:-0.25em}sup{top:-0.5em}a{color:#00a99d;text-decoration:none;background-color:transparent}a:hover{color:#005d56;text-decoration:underline}a:not([href]):not([class]){color:inherit;text-decoration:none}a:not([href]):not([class]):hover{color:inherit;text-decoration:none}pre,code,kbd,samp{font-family:SFMono-Regular,Menlo,Monaco,Consolas,"Liberation Mono","Courier New",monospace;font-size:1em}pre{margin-top:0;margin-bottom:1rem;overflow:auto;-ms-overflow-style:scrollbar}figure{margin:0 0 1rem}img{vertical-align:middle;border-style:none}svg{overflow:hidden;vertical-align:middle}table{border-collapse:collapse}caption{padding-top:.75rem;padding-bottom:.75rem;color:#6c757d;text-align:left;caption-side:bottom}th{text-align:inherit;text-align:-webkit-match-parent}label{display:inline-block;margin-bottom:.5rem}button{border-radius:0}button:focus:not(:focus-visible){outline:0}input,button,select,optgroup,textarea{margin:0;font-family:inherit;font-size:inherit;line-height:inherit}button,input{overflow:visible}button,select{text-transform:none}[role=button]{cursor:pointer}select{word-wrap:normal}button,[type=button],[type=reset],[type=submit]{-webkit-appearance:button}button:not(:disabled),[type=button]:not(:disabled),[type=reset]:not(:disabled),[type=submit]:not(:disabled){cursor:pointer}button::-moz-focus-inner,[type=button]::-moz-focus-inner,[type=reset]::-moz-focus-inner,[type=submit]::-moz-focus-inner{padding:0;border-style:none}input[type=radio],input[type=checkbox]{box-sizing:border-box;padding:0}textarea{overflow:auto;resize:vertical}fieldset{min-width:0;padding:0;margin:0;border:0}legend{display:block;width:100%;max-width:100%;padding:0;margin-bottom:.5rem;font-size:1.5rem;line-height:inherit;color:inherit;white-space:normal}progress{vertical-align:baseline}[type=number]::-webkit-inner-spin-button,[type=number]::-webkit-outer-spin-button{height:auto}[type=search]{outline-offset:-2px;-webkit-appearance:none}[type=search]::-webkit-search-decoration{-webkit-appearance:none}::-webkit-file-upload-button{font:inherit;-webkit-appearance:button}output{display:inline-block}summary{display:list-item;cursor:pointer}template{display:none}[hidden]{display:none !important}h1,h2,h3,h4,h5,h6,.h1,.h2,.h3,.h4,.h5,.h6{margin-bottom:.5rem;font-weight:500;line-height:1.2}h1,.h1{font-size:2.5rem}h2,.h2{font-size:2rem}h3,.h3{font-size:1.75rem}h4,.h4{font-size:1.5rem}h5,.h5{font-size:1.25rem}h6,.h6{font-size:1rem}.lead{font-size:1.25rem;font-weight:300}.display-1{font-size:6rem;font-weight:300;line-height:1.2}.display-2{font-size:5.5rem;font-weight:300;line-height:1.2}.display-3{font-size:4.5rem;font-weight:300;line-height:1.2}.display-4{font-size:3.5rem;font-weight:300;line-height:1.2}hr{margin-top:1rem;margin-bottom:1rem;border:0;border-top:1px solid rgba(0,0,0,.1)}small,.small{font-size:80%;font-weight:400}mark,.mark{padding:.2em;background-color:#fcf8e3}.list-unstyled{padding-left:0;list-style:none}.list-inline{padding-left:0;list-style:none}.list-inline-item{display:inline-block}.list-inline-item:not(:last-child){margin-right:.5rem}.initialism{font-size:90%;text-transform:uppercase}.blockquote{margin-bottom:1rem;font-size:1.25rem}.blockquote-footer{display:block;font-size:80%;color:#6c757d}.blockquote-footer::before{content:"— "}.img-fluid{max-width:100%;height:auto}.img-thumbnail{padding:.25rem;background-color:#fff;border:1px solid #dee2e6;border-radius:.25rem;max-width:100%;height:auto}.figure{display:inline-block}.figure-img{margin-bottom:.5rem;line-height:1}.figure-caption{font-size:90%;color:#6c757d}code{font-size:87.5%;color:#e83e8c;word-wrap:break-word}a>code{color:inherit}kbd{padding:.2rem .4rem;font-size:87.5%;color:#fff;background-color:#212529;border-radius:.2rem}kbd kbd{padding:0;font-size:100%;font-weight:700}pre{display:block;font-size:87.5%;color:#212529}pre code{font-size:inherit;color:inherit;word-break:normal}.pre-scrollable{max-height:340px;overflow-y:scroll}.container,.container-fluid,.container-xl,.container-lg,.container-md,.container-sm{width:100%;padding-right:15px;padding-left:15px;margin-right:auto;margin-left:auto}@media(min-width: 576px){.container-sm,.container{max-width:540px}}@media(min-width: 768px){.container-md,.container-sm,.container{max-width:720px}}@media(min-width: 992px){.container-lg,.container-md,.container-sm,.container{max-width:960px}}@media(min-width: 1200px){.container-xl,.container-lg,.container-md,.container-sm,.container{max-width:1140px}}.row{display:flex;flex-wrap:wrap;margin-right:-15px;margin-left:-15px}.no-gutters{margin-right:0;margin-left:0}.no-gutters>.col,.no-gutters>[class*=col-]{padding-right:0;padding-left:0}.col-xl,.col-xl-auto,.col-xl-12,.col-xl-11,.col-xl-10,.col-xl-9,.col-xl-8,.col-xl-7,.col-xl-6,.col-xl-5,.col-xl-4,.col-xl-3,.col-xl-2,.col-xl-1,.col-lg,.col-lg-auto,.col-lg-12,.col-lg-11,.col-lg-10,.col-lg-9,.col-lg-8,.col-lg-7,.col-lg-6,.col-lg-5,.col-lg-4,.col-lg-3,.col-lg-2,.col-lg-1,.col-md,.col-md-auto,.col-md-12,.col-md-11,.col-md-10,.col-md-9,.col-md-8,.col-md-7,.col-md-6,.col-md-5,.col-md-4,.col-md-3,.col-md-2,.col-md-1,.col-sm,.col-sm-auto,.col-sm-12,.col-sm-11,.col-sm-10,.col-sm-9,.col-sm-8,.col-sm-7,.col-sm-6,.col-sm-5,.col-sm-4,.col-sm-3,.col-sm-2,.col-sm-1,.col,.col-auto,.col-12,.col-11,.col-10,.col-9,.col-8,.col-7,.col-6,.col-5,.col-4,.col-3,.col-2,.col-1{position:relative;width:100%;padding-right:15px;padding-left:15px}.col{flex-basis:0;flex-grow:1;max-width:100%}.row-cols-1>*{flex:0 0 100%;max-width:100%}.row-cols-2>*{flex:0 0 50%;max-width:50%}.row-cols-3>*{flex:0 0 33.3333333333%;max-width:33.3333333333%}.row-cols-4>*{flex:0 0 25%;max-width:25%}.row-cols-5>*{flex:0 0 20%;max-width:20%}.row-cols-6>*{flex:0 0 16.6666666667%;max-width:16.6666666667%}.col-auto{flex:0 0 auto;width:auto;max-width:100%}.col-1{flex:0 0 8.3333333333%;max-width:8.3333333333%}.col-2{flex:0 0 16.6666666667%;max-width:16.6666666667%}.col-3{flex:0 0 25%;max-width:25%}.col-4{flex:0 0 33.3333333333%;max-width:33.3333333333%}.col-5{flex:0 0 41.6666666667%;max-width:41.6666666667%}.col-6{flex:0 0 50%;max-width:50%}.col-7{flex:0 0 58.3333333333%;max-width:58.3333333333%}.col-8{flex:0 0 66.6666666667%;max-width:66.6666666667%}.col-9{flex:0 0 75%;max-width:75%}.col-10{flex:0 0 83.3333333333%;max-width:83.3333333333%}.col-11{flex:0 0 91.6666666667%;max-width:91.6666666667%}.col-12{flex:0 0 100%;max-width:100%}.order-first{order:-1}.order-last{order:13}.order-0{order:0}.order-1{order:1}.order-2{order:2}.order-3{order:3}.order-4{order:4}.order-5{order:5}.order-6{order:6}.order-7{order:7}.order-8{order:8}.order-9{order:9}.order-10{order:10}.order-11{order:11}.order-12{order:12}.offset-1{margin-left:8.3333333333%}.offset-2{margin-left:16.6666666667%}.offset-3{margin-left:25%}.offset-4{margin-left:33.3333333333%}.offset-5{margin-left:41.6666666667%}.offset-6{margin-left:50%}.offset-7{margin-left:58.3333333333%}.offset-8{margin-left:66.6666666667%}.offset-9{margin-left:75%}.offset-10{margin-left:83.3333333333%}.offset-11{margin-left:91.6666666667%}@media(min-width: 576px){.col-sm{flex-basis:0;flex-grow:1;max-width:100%}.row-cols-sm-1>*{flex:0 0 100%;max-width:100%}.row-cols-sm-2>*{flex:0 0 50%;max-width:50%}.row-cols-sm-3>*{flex:0 0 33.3333333333%;max-width:33.3333333333%}.row-cols-sm-4>*{flex:0 0 25%;max-width:25%}.row-cols-sm-5>*{flex:0 0 20%;max-width:20%}.row-cols-sm-6>*{flex:0 0 16.6666666667%;max-width:16.6666666667%}.col-sm-auto{flex:0 0 auto;width:auto;max-width:100%}.col-sm-1{flex:0 0 8.3333333333%;max-width:8.3333333333%}.col-sm-2{flex:0 0 16.6666666667%;max-width:16.6666666667%}.col-sm-3{flex:0 0 25%;max-width:25%}.col-sm-4{flex:0 0 33.3333333333%;max-width:33.3333333333%}.col-sm-5{flex:0 0 41.6666666667%;max-width:41.6666666667%}.col-sm-6{flex:0 0 50%;max-width:50%}.col-sm-7{flex:0 0 58.3333333333%;max-width:58.3333333333%}.col-sm-8{flex:0 0 66.6666666667%;max-width:66.6666666667%}.col-sm-9{flex:0 0 75%;max-width:75%}.col-sm-10{flex:0 0 83.3333333333%;max-width:83.3333333333%}.col-sm-11{flex:0 0 91.6666666667%;max-width:91.6666666667%}.col-sm-12{flex:0 0 100%;max-width:100%}.order-sm-first{order:-1}.order-sm-last{order:13}.order-sm-0{order:0}.order-sm-1{order:1}.order-sm-2{order:2}.order-sm-3{order:3}.order-sm-4{order:4}.order-sm-5{order:5}.order-sm-6{order:6}.order-sm-7{order:7}.order-sm-8{order:8}.order-sm-9{order:9}.order-sm-10{order:10}.order-sm-11{order:11}.order-sm-12{order:12}.offset-sm-0{margin-left:0}.offset-sm-1{margin-left:8.3333333333%}.offset-sm-2{margin-left:16.6666666667%}.offset-sm-3{margin-left:25%}.offset-sm-4{margin-left:33.3333333333%}.offset-sm-5{margin-left:41.6666666667%}.offset-sm-6{margin-left:50%}.offset-sm-7{margin-left:58.3333333333%}.offset-sm-8{margin-left:66.6666666667%}.offset-sm-9{margin-left:75%}.offset-sm-10{margin-left:83.3333333333%}.offset-sm-11{margin-left:91.6666666667%}}@media(min-width: 768px){.col-md{flex-basis:0;flex-grow:1;max-width:100%}.row-cols-md-1>*{flex:0 0 100%;max-width:100%}.row-cols-md-2>*{flex:0 0 50%;max-width:50%}.row-cols-md-3>*{flex:0 0 33.3333333333%;max-width:33.3333333333%}.row-cols-md-4>*{flex:0 0 25%;max-width:25%}.row-cols-md-5>*{flex:0 0 20%;max-width:20%}.row-cols-md-6>*{flex:0 0 16.6666666667%;max-width:16.6666666667%}.col-md-auto{flex:0 0 auto;width:auto;max-width:100%}.col-md-1{flex:0 0 8.3333333333%;max-width:8.3333333333%}.col-md-2{flex:0 0 16.6666666667%;max-width:16.6666666667%}.col-md-3{flex:0 0 25%;max-width:25%}.col-md-4{flex:0 0 33.3333333333%;max-width:33.3333333333%}.col-md-5{flex:0 0 41.6666666667%;max-width:41.6666666667%}.col-md-6{flex:0 0 50%;max-width:50%}.col-md-7{flex:0 0 58.3333333333%;max-width:58.3333333333%}.col-md-8{flex:0 0 66.6666666667%;max-width:66.6666666667%}.col-md-9{flex:0 0 75%;max-width:75%}.col-md-10{flex:0 0 83.3333333333%;max-width:83.3333333333%}.col-md-11{flex:0 0 91.6666666667%;max-width:91.6666666667%}.col-md-12{flex:0 0 100%;max-width:100%}.order-md-first{order:-1}.order-md-last{order:13}.order-md-0{order:0}.order-md-1{order:1}.order-md-2{order:2}.order-md-3{order:3}.order-md-4{order:4}.order-md-5{order:5}.order-md-6{order:6}.order-md-7{order:7}.order-md-8{order:8}.order-md-9{order:9}.order-md-10{order:10}.order-md-11{order:11}.order-md-12{order:12}.offset-md-0{margin-left:0}.offset-md-1{margin-left:8.3333333333%}.offset-md-2{margin-left:16.6666666667%}.offset-md-3{margin-left:25%}.offset-md-4{margin-left:33.3333333333%}.offset-md-5{margin-left:41.6666666667%}.offset-md-6{margin-left:50%}.offset-md-7{margin-left:58.3333333333%}.offset-md-8{margin-left:66.6666666667%}.offset-md-9{margin-left:75%}.offset-md-10{margin-left:83.3333333333%}.offset-md-11{margin-left:91.6666666667%}}@media(min-width: 992px){.col-lg{flex-basis:0;flex-grow:1;max-width:100%}.row-cols-lg-1>*{flex:0 0 100%;max-width:100%}.row-cols-lg-2>*{flex:0 0 50%;max-width:50%}.row-cols-lg-3>*{flex:0 0 33.3333333333%;max-width:33.3333333333%}.row-cols-lg-4>*{flex:0 0 25%;max-width:25%}.row-cols-lg-5>*{flex:0 0 20%;max-width:20%}.row-cols-lg-6>*{flex:0 0 16.6666666667%;max-width:16.6666666667%}.col-lg-auto{flex:0 0 auto;width:auto;max-width:100%}.col-lg-1{flex:0 0 8.3333333333%;max-width:8.3333333333%}.col-lg-2{flex:0 0 16.6666666667%;max-width:16.6666666667%}.col-lg-3{flex:0 0 25%;max-width:25%}.col-lg-4{flex:0 0 33.3333333333%;max-width:33.3333333333%}.col-lg-5{flex:0 0 41.6666666667%;max-width:41.6666666667%}.col-lg-6{flex:0 0 50%;max-width:50%}.col-lg-7{flex:0 0 58.3333333333%;max-width:58.3333333333%}.col-lg-8{flex:0 0 66.6666666667%;max-width:66.6666666667%}.col-lg-9{flex:0 0 75%;max-width:75%}.col-lg-10{flex:0 0 83.3333333333%;max-width:83.3333333333%}.col-lg-11{flex:0 0 91.6666666667%;max-width:91.6666666667%}.col-lg-12{flex:0 0 100%;max-width:100%}.order-lg-first{order:-1}.order-lg-last{order:13}.order-lg-0{order:0}.order-lg-1{order:1}.order-lg-2{order:2}.order-lg-3{order:3}.order-lg-4{order:4}.order-lg-5{order:5}.order-lg-6{order:6}.order-lg-7{order:7}.order-lg-8{order:8}.order-lg-9{order:9}.order-lg-10{order:10}.order-lg-11{order:11}.order-lg-12{order:12}.offset-lg-0{margin-left:0}.offset-lg-1{margin-left:8.3333333333%}.offset-lg-2{margin-left:16.6666666667%}.offset-lg-3{margin-left:25%}.offset-lg-4{margin-left:33.3333333333%}.offset-lg-5{margin-left:41.6666666667%}.offset-lg-6{margin-left:50%}.offset-lg-7{margin-left:58.3333333333%}.offset-lg-8{margin-left:66.6666666667%}.offset-lg-9{margin-left:75%}.offset-lg-10{margin-left:83.3333333333%}.offset-lg-11{margin-left:91.6666666667%}}@media(min-width: 1200px){.col-xl{flex-basis:0;flex-grow:1;max-width:100%}.row-cols-xl-1>*{flex:0 0 100%;max-width:100%}.row-cols-xl-2>*{flex:0 0 50%;max-width:50%}.row-cols-xl-3>*{flex:0 0 33.3333333333%;max-width:33.3333333333%}.row-cols-xl-4>*{flex:0 0 25%;max-width:25%}.row-cols-xl-5>*{flex:0 0 20%;max-width:20%}.row-cols-xl-6>*{flex:0 0 16.6666666667%;max-width:16.6666666667%}.col-xl-auto{flex:0 0 auto;width:auto;max-width:100%}.col-xl-1{flex:0 0 8.3333333333%;max-width:8.3333333333%}.col-xl-2{flex:0 0 16.6666666667%;max-width:16.6666666667%}.col-xl-3{flex:0 0 25%;max-width:25%}.col-xl-4{flex:0 0 33.3333333333%;max-width:33.3333333333%}.col-xl-5{flex:0 0 41.6666666667%;max-width:41.6666666667%}.col-xl-6{flex:0 0 50%;max-width:50%}.col-xl-7{flex:0 0 58.3333333333%;max-width:58.3333333333%}.col-xl-8{flex:0 0 66.6666666667%;max-width:66.6666666667%}.col-xl-9{flex:0 0 75%;max-width:75%}.col-xl-10{flex:0 0 83.3333333333%;max-width:83.3333333333%}.col-xl-11{flex:0 0 91.6666666667%;max-width:91.6666666667%}.col-xl-12{flex:0 0 100%;max-width:100%}.order-xl-first{order:-1}.order-xl-last{order:13}.order-xl-0{order:0}.order-xl-1{order:1}.order-xl-2{order:2}.order-xl-3{order:3}.order-xl-4{order:4}.order-xl-5{order:5}.order-xl-6{order:6}.order-xl-7{order:7}.order-xl-8{order:8}.order-xl-9{order:9}.order-xl-10{order:10}.order-xl-11{order:11}.order-xl-12{order:12}.offset-xl-0{margin-left:0}.offset-xl-1{margin-left:8.3333333333%}.offset-xl-2{margin-left:16.6666666667%}.offset-xl-3{margin-left:25%}.offset-xl-4{margin-left:33.3333333333%}.offset-xl-5{margin-left:41.6666666667%}.offset-xl-6{margin-left:50%}.offset-xl-7{margin-left:58.3333333333%}.offset-xl-8{margin-left:66.6666666667%}.offset-xl-9{margin-left:75%}.offset-xl-10{margin-left:83.3333333333%}.offset-xl-11{margin-left:91.6666666667%}}.table{width:100%;margin-bottom:1rem;color:#212529}.table th,.table td{padding:.75rem;vertical-align:top;border-top:1px solid #dee2e6}.table thead th{vertical-align:bottom;border-bottom:2px solid #dee2e6}.table tbody+tbody{border-top:2px solid #dee2e6}.table-sm th,.table-sm td{padding:.3rem}.table-bordered{border:1px solid #dee2e6}.table-bordered th,.table-bordered td{border:1px solid #dee2e6}.table-bordered thead th,.table-bordered thead td{border-bottom-width:2px}.table-borderless th,.table-borderless td,.table-borderless thead th,.table-borderless tbody+tbody{border:0}.table-striped tbody tr:nth-of-type(odd){background-color:rgba(0,0,0,.05)}.table-hover tbody tr:hover{color:#212529;background-color:rgba(0,0,0,.075)}.table-primary,.table-primary>th,.table-primary>td{background-color:#b8e7e4}.table-primary th,.table-primary td,.table-primary thead th,.table-primary tbody+tbody{border-color:#7ad2cc}.table-hover .table-primary:hover{background-color:#a5e1dd}.table-hover .table-primary:hover>td,.table-hover .table-primary:hover>th{background-color:#a5e1dd}.table-secondary,.table-secondary>th,.table-secondary>td{background-color:#d6d8db}.table-secondary th,.table-secondary td,.table-secondary thead th,.table-secondary tbody+tbody{border-color:#b3b7bb}.table-hover .table-secondary:hover{background-color:#c8cbcf}.table-hover .table-secondary:hover>td,.table-hover .table-secondary:hover>th{background-color:#c8cbcf}.table-success,.table-success>th,.table-success>td{background-color:#c3e6cb}.table-success th,.table-success td,.table-success thead th,.table-success tbody+tbody{border-color:#8fd19e}.table-hover .table-success:hover{background-color:#b1dfbb}.table-hover .table-success:hover>td,.table-hover .table-success:hover>th{background-color:#b1dfbb}.table-info,.table-info>th,.table-info>td{background-color:#bee5eb}.table-info th,.table-info td,.table-info thead th,.table-info tbody+tbody{border-color:#86cfda}.table-hover .table-info:hover{background-color:#abdde5}.table-hover .table-info:hover>td,.table-hover .table-info:hover>th{background-color:#abdde5}.table-warning,.table-warning>th,.table-warning>td{background-color:#ffeeba}.table-warning th,.table-warning td,.table-warning thead th,.table-warning tbody+tbody{border-color:#ffdf7e}.table-hover .table-warning:hover{background-color:#ffe8a1}.table-hover .table-warning:hover>td,.table-hover .table-warning:hover>th{background-color:#ffe8a1}.table-danger,.table-danger>th,.table-danger>td{background-color:#f5c6cb}.table-danger th,.table-danger td,.table-danger thead th,.table-danger tbody+tbody{border-color:#ed969e}.table-hover .table-danger:hover{background-color:#f1b0b7}.table-hover .table-danger:hover>td,.table-hover .table-danger:hover>th{background-color:#f1b0b7}.table-light,.table-light>th,.table-light>td{background-color:#fdfdfe}.table-light th,.table-light td,.table-light thead th,.table-light tbody+tbody{border-color:#fbfcfc}.table-hover .table-light:hover{background-color:#ececf6}.table-hover .table-light:hover>td,.table-hover .table-light:hover>th{background-color:#ececf6}.table-dark,.table-dark>th,.table-dark>td{background-color:#c6c8ca}.table-dark th,.table-dark td,.table-dark thead th,.table-dark tbody+tbody{border-color:#95999c}.table-hover .table-dark:hover{background-color:#b9bbbe}.table-hover .table-dark:hover>td,.table-hover .table-dark:hover>th{background-color:#b9bbbe}.table-active,.table-active>th,.table-active>td{background-color:rgba(0,0,0,.075)}.table-hover .table-active:hover{background-color:rgba(0,0,0,.075)}.table-hover .table-active:hover>td,.table-hover .table-active:hover>th{background-color:rgba(0,0,0,.075)}.table .thead-dark th{color:#fff;background-color:#343a40;border-color:#454d55}.table .thead-light th{color:#495057;background-color:#e9ecef;border-color:#dee2e6}.table-dark{color:#fff;background-color:#343a40}.table-dark th,.table-dark td,.table-dark thead th{border-color:#454d55}.table-dark.table-bordered{border:0}.table-dark.table-striped tbody tr:nth-of-type(odd){background-color:rgba(255,255,255,.05)}.table-dark.table-hover tbody tr:hover{color:#fff;background-color:rgba(255,255,255,.075)}@media(max-width: 575.98px){.table-responsive-sm{display:block;width:100%;overflow-x:auto;-webkit-overflow-scrolling:touch}.table-responsive-sm>.table-bordered{border:0}}@media(max-width: 767.98px){.table-responsive-md{display:block;width:100%;overflow-x:auto;-webkit-overflow-scrolling:touch}.table-responsive-md>.table-bordered{border:0}}@media(max-width: 991.98px){.table-responsive-lg{display:block;width:100%;overflow-x:auto;-webkit-overflow-scrolling:touch}.table-responsive-lg>.table-bordered{border:0}}@media(max-width: 1199.98px){.table-responsive-xl{display:block;width:100%;overflow-x:auto;-webkit-overflow-scrolling:touch}.table-responsive-xl>.table-bordered{border:0}}.table-responsive{display:block;width:100%;overflow-x:auto;-webkit-overflow-scrolling:touch}.table-responsive>.table-bordered{border:0}.form-control{display:block;width:100%;height:calc(1.5em + 0.75rem + 2px);padding:.375rem .75rem;font-size:1rem;font-weight:400;line-height:1.5;color:#495057;background-color:#fff;background-clip:padding-box;border:1px solid #ced4da;border-radius:.25rem;transition:border-color .15s ease-in-out,box-shadow .15s ease-in-out}@media(prefers-reduced-motion: reduce){.form-control{transition:none}}.form-control::-ms-expand{background-color:transparent;border:0}.form-control:-moz-focusring{color:transparent;text-shadow:0 0 0 #495057}.form-control:focus{color:#495057;background-color:#fff;border-color:#2afff0;outline:0;box-shadow:0 0 0 .2rem rgba(0,169,157,.25)}.form-control::placeholder{color:#6c757d;opacity:1}.form-control:disabled,.form-control[readonly]{background-color:#e9ecef;opacity:1}input[type=date].form-control,input[type=time].form-control,input[type=datetime-local].form-control,input[type=month].form-control{appearance:none}select.form-control:focus::-ms-value{color:#495057;background-color:#fff}.form-control-file,.form-control-range{display:block;width:100%}.col-form-label{padding-top:calc(0.375rem + 1px);padding-bottom:calc(0.375rem + 1px);margin-bottom:0;font-size:inherit;line-height:1.5}.col-form-label-lg{padding-top:calc(0.5rem + 1px);padding-bottom:calc(0.5rem + 1px);font-size:1.25rem;line-height:1.5}.col-form-label-sm{padding-top:calc(0.25rem + 1px);padding-bottom:calc(0.25rem + 1px);font-size:0.875rem;line-height:1.5}.form-control-plaintext{display:block;width:100%;padding:.375rem 0;margin-bottom:0;font-size:1rem;line-height:1.5;color:#212529;background-color:transparent;border:solid transparent;border-width:1px 0}.form-control-plaintext.form-control-sm,.form-control-plaintext.form-control-lg{padding-right:0;padding-left:0}.form-control-sm{height:calc(1.5em + 0.5rem + 2px);padding:.25rem .5rem;font-size:0.875rem;line-height:1.5;border-radius:.2rem}.form-control-lg{height:calc(1.5em + 1rem + 2px);padding:.5rem 1rem;font-size:1.25rem;line-height:1.5;border-radius:.3rem}select.form-control[size],select.form-control[multiple]{height:auto}textarea.form-control{height:auto}.form-group{margin-bottom:1rem}.form-text{display:block;margin-top:.25rem}.form-row{display:flex;flex-wrap:wrap;margin-right:-5px;margin-left:-5px}.form-row>.col,.form-row>[class*=col-]{padding-right:5px;padding-left:5px}.form-check{position:relative;display:block;padding-left:1.25rem}.form-check-input{position:absolute;margin-top:.3rem;margin-left:-1.25rem}.form-check-input[disabled]~.form-check-label,.form-check-input:disabled~.form-check-label{color:#6c757d}.form-check-label{margin-bottom:0}.form-check-inline{display:inline-flex;align-items:center;padding-left:0;margin-right:.75rem}.form-check-inline .form-check-input{position:static;margin-top:0;margin-right:.3125rem;margin-left:0}.valid-feedback{display:none;width:100%;margin-top:.25rem;font-size:80%;color:#28a745}.valid-tooltip{position:absolute;top:100%;left:0;z-index:5;display:none;max-width:100%;padding:.25rem .5rem;margin-top:.1rem;font-size:0.875rem;line-height:1.5;color:#fff;background-color:rgba(40,167,69,.9);border-radius:.25rem}.form-row>.col>.valid-tooltip,.form-row>[class*=col-]>.valid-tooltip{left:5px}.was-validated :valid~.valid-feedback,.was-validated :valid~.valid-tooltip,.is-valid~.valid-feedback,.is-valid~.valid-tooltip{display:block}.was-validated .form-control:valid,.form-control.is-valid{border-color:#28a745;padding-right:calc(1.5em + 0.75rem);background-image:url("data:image/svg+xml,%3csvg xmlns=\'http://www.w3.org/2000/svg\' width=\'8\' height=\'8\' viewBox=\'0 0 8 8\'%3e%3cpath fill=\'%2328a745\' d=\'M2.3 6.73L.6 4.53c-.4-1.04.46-1.4 1.1-.8l1.1 1.4 3.4-3.8c.6-.63 1.6-.27 1.2.7l-4 4.6c-.43.5-.8.4-1.1.1z\'/%3e%3c/svg%3e");background-repeat:no-repeat;background-position:right calc(0.375em + 0.1875rem) center;background-size:calc(0.75em + 0.375rem) calc(0.75em + 0.375rem)}.was-validated .form-control:valid:focus,.form-control.is-valid:focus{border-color:#28a745;box-shadow:0 0 0 .2rem rgba(40,167,69,.25)}.was-validated textarea.form-control:valid,textarea.form-control.is-valid{padding-right:calc(1.5em + 0.75rem);background-position:top calc(0.375em + 0.1875rem) right calc(0.375em + 0.1875rem)}.was-validated .custom-select:valid,.custom-select.is-valid{border-color:#28a745;padding-right:calc(0.75em + 2.3125rem);background:url("data:image/svg+xml,%3csvg xmlns=\'http://www.w3.org/2000/svg\' width=\'4\' height=\'5\' viewBox=\'0 0 4 5\'%3e%3cpath fill=\'%23343a40\' d=\'M2 0L0 2h4zm0 5L0 3h4z\'/%3e%3c/svg%3e") right .75rem center/8px 10px no-repeat,#fff url("data:image/svg+xml,%3csvg xmlns=\'http://www.w3.org/2000/svg\' width=\'8\' height=\'8\' viewBox=\'0 0 8 8\'%3e%3cpath fill=\'%2328a745\' d=\'M2.3 6.73L.6 4.53c-.4-1.04.46-1.4 1.1-.8l1.1 1.4 3.4-3.8c.6-.63 1.6-.27 1.2.7l-4 4.6c-.43.5-.8.4-1.1.1z\'/%3e%3c/svg%3e") center right 1.75rem/calc(0.75em + 0.375rem) calc(0.75em + 0.375rem) no-repeat}.was-validated .custom-select:valid:focus,.custom-select.is-valid:focus{border-color:#28a745;box-shadow:0 0 0 .2rem rgba(40,167,69,.25)}.was-validated .form-check-input:valid~.form-check-label,.form-check-input.is-valid~.form-check-label{color:#28a745}.was-validated .form-check-input:valid~.valid-feedback,.was-validated .form-check-input:valid~.valid-tooltip,.form-check-input.is-valid~.valid-feedback,.form-check-input.is-valid~.valid-tooltip{display:block}.was-validated .custom-control-input:valid~.custom-control-label,.custom-control-input.is-valid~.custom-control-label{color:#28a745}.was-validated .custom-control-input:valid~.custom-control-label::before,.custom-control-input.is-valid~.custom-control-label::before{border-color:#28a745}.was-validated .custom-control-input:valid:checked~.custom-control-label::before,.custom-control-input.is-valid:checked~.custom-control-label::before{border-color:#34ce57;background-color:#34ce57}.was-validated .custom-control-input:valid:focus~.custom-control-label::before,.custom-control-input.is-valid:focus~.custom-control-label::before{box-shadow:0 0 0 .2rem rgba(40,167,69,.25)}.was-validated .custom-control-input:valid:focus:not(:checked)~.custom-control-label::before,.custom-control-input.is-valid:focus:not(:checked)~.custom-control-label::before{border-color:#28a745}.was-validated .custom-file-input:valid~.custom-file-label,.custom-file-input.is-valid~.custom-file-label{border-color:#28a745}.was-validated .custom-file-input:valid:focus~.custom-file-label,.custom-file-input.is-valid:focus~.custom-file-label{border-color:#28a745;box-shadow:0 0 0 .2rem rgba(40,167,69,.25)}.invalid-feedback{display:none;width:100%;margin-top:.25rem;font-size:80%;color:#dc3545}.invalid-tooltip{position:absolute;top:100%;left:0;z-index:5;display:none;max-width:100%;padding:.25rem .5rem;margin-top:.1rem;font-size:0.875rem;line-height:1.5;color:#fff;background-color:rgba(220,53,69,.9);border-radius:.25rem}.form-row>.col>.invalid-tooltip,.form-row>[class*=col-]>.invalid-tooltip{left:5px}.was-validated :invalid~.invalid-feedback,.was-validated :invalid~.invalid-tooltip,.is-invalid~.invalid-feedback,.is-invalid~.invalid-tooltip{display:block}.was-validated .form-control:invalid,.form-control.is-invalid{border-color:#dc3545;padding-right:calc(1.5em + 0.75rem);background-image:url("data:image/svg+xml,%3csvg xmlns=\'http://www.w3.org/2000/svg\' width=\'12\' height=\'12\' fill=\'none\' stroke=\'%23dc3545\' viewBox=\'0 0 12 12\'%3e%3ccircle cx=\'6\' cy=\'6\' r=\'4.5\'/%3e%3cpath stroke-linejoin=\'round\' d=\'M5.8 3.6h.4L6 6.5z\'/%3e%3ccircle cx=\'6\' cy=\'8.2\' r=\'.6\' fill=\'%23dc3545\' stroke=\'none\'/%3e%3c/svg%3e");background-repeat:no-repeat;background-position:right calc(0.375em + 0.1875rem) center;background-size:calc(0.75em + 0.375rem) calc(0.75em + 0.375rem)}.was-validated .form-control:invalid:focus,.form-control.is-invalid:focus{border-color:#dc3545;box-shadow:0 0 0 .2rem rgba(220,53,69,.25)}.was-validated textarea.form-control:invalid,textarea.form-control.is-invalid{padding-right:calc(1.5em + 0.75rem);background-position:top calc(0.375em + 0.1875rem) right calc(0.375em + 0.1875rem)}.was-validated .custom-select:invalid,.custom-select.is-invalid{border-color:#dc3545;padding-right:calc(0.75em + 2.3125rem);background:url("data:image/svg+xml,%3csvg xmlns=\'http://www.w3.org/2000/svg\' width=\'4\' height=\'5\' viewBox=\'0 0 4 5\'%3e%3cpath fill=\'%23343a40\' d=\'M2 0L0 2h4zm0 5L0 3h4z\'/%3e%3c/svg%3e") right .75rem center/8px 10px no-repeat,#fff url("data:image/svg+xml,%3csvg xmlns=\'http://www.w3.org/2000/svg\' width=\'12\' height=\'12\' fill=\'none\' stroke=\'%23dc3545\' viewBox=\'0 0 12 12\'%3e%3ccircle cx=\'6\' cy=\'6\' r=\'4.5\'/%3e%3cpath stroke-linejoin=\'round\' d=\'M5.8 3.6h.4L6 6.5z\'/%3e%3ccircle cx=\'6\' cy=\'8.2\' r=\'.6\' fill=\'%23dc3545\' stroke=\'none\'/%3e%3c/svg%3e") center right 1.75rem/calc(0.75em + 0.375rem) calc(0.75em + 0.375rem) no-repeat}.was-validated .custom-select:invalid:focus,.custom-select.is-invalid:focus{border-color:#dc3545;box-shadow:0 0 0 .2rem rgba(220,53,69,.25)}.was-validated .form-check-input:invalid~.form-check-label,.form-check-input.is-invalid~.form-check-label{color:#dc3545}.was-validated .form-check-input:invalid~.invalid-feedback,.was-validated .form-check-input:invalid~.invalid-tooltip,.form-check-input.is-invalid~.invalid-feedback,.form-check-input.is-invalid~.invalid-tooltip{display:block}.was-validated .custom-control-input:invalid~.custom-control-label,.custom-control-input.is-invalid~.custom-control-label{color:#dc3545}.was-validated .custom-control-input:invalid~.custom-control-label::before,.custom-control-input.is-invalid~.custom-control-label::before{border-color:#dc3545}.was-validated .custom-control-input:invalid:checked~.custom-control-label::before,.custom-control-input.is-invalid:checked~.custom-control-label::before{border-color:#e4606d;background-color:#e4606d}.was-validated .custom-control-input:invalid:focus~.custom-control-label::before,.custom-control-input.is-invalid:focus~.custom-control-label::before{box-shadow:0 0 0 .2rem rgba(220,53,69,.25)}.was-validated .custom-control-input:invalid:focus:not(:checked)~.custom-control-label::before,.custom-control-input.is-invalid:focus:not(:checked)~.custom-control-label::before{border-color:#dc3545}.was-validated .custom-file-input:invalid~.custom-file-label,.custom-file-input.is-invalid~.custom-file-label{border-color:#dc3545}.was-validated .custom-file-input:invalid:focus~.custom-file-label,.custom-file-input.is-invalid:focus~.custom-file-label{border-color:#dc3545;box-shadow:0 0 0 .2rem rgba(220,53,69,.25)}.form-inline{display:flex;flex-flow:row wrap;align-items:center}.form-inline .form-check{width:100%}@media(min-width: 576px){.form-inline label{display:flex;align-items:center;justify-content:center;margin-bottom:0}.form-inline .form-group{display:flex;flex:0 0 auto;flex-flow:row wrap;align-items:center;margin-bottom:0}.form-inline .form-control{display:inline-block;width:auto;vertical-align:middle}.form-inline .form-control-plaintext{display:inline-block}.form-inline .input-group,.form-inline .custom-select{width:auto}.form-inline .form-check{display:flex;align-items:center;justify-content:center;width:auto;padding-left:0}.form-inline .form-check-input{position:relative;flex-shrink:0;margin-top:0;margin-right:.25rem;margin-left:0}.form-inline .custom-control{align-items:center;justify-content:center}.form-inline .custom-control-label{margin-bottom:0}}.btn{display:inline-block;font-weight:400;color:#212529;text-align:center;vertical-align:middle;user-select:none;background-color:transparent;border:1px solid transparent;padding:.375rem .75rem;font-size:1rem;line-height:1.5;border-radius:.25rem;transition:color .15s ease-in-out,background-color .15s ease-in-out,border-color .15s ease-in-out,box-shadow .15s ease-in-out}@media(prefers-reduced-motion: reduce){.btn{transition:none}}.btn:hover{color:#212529;text-decoration:none}.btn:focus,.btn.focus{outline:0;box-shadow:0 0 0 .2rem rgba(0,169,157,.25)}.btn.disabled,.btn:disabled{opacity:.65}.btn:not(:disabled):not(.disabled){cursor:pointer}a.btn.disabled,fieldset:disabled a.btn{pointer-events:none}.btn-primary{color:#fff;background-color:#00a99d;border-color:#00a99d}.btn-primary:hover{color:#fff;background-color:#008379;border-color:#00766e}.btn-primary:focus,.btn-primary.focus{color:#fff;background-color:#008379;border-color:#00766e;box-shadow:0 0 0 .2rem rgba(38,182,172,.5)}.btn-primary.disabled,.btn-primary:disabled{color:#fff;background-color:#00a99d;border-color:#00a99d}.btn-primary:not(:disabled):not(.disabled):active,.btn-primary:not(:disabled):not(.disabled).active,.show>.btn-primary.dropdown-toggle{color:#fff;background-color:#00766e;border-color:#006962}.btn-primary:not(:disabled):not(.disabled):active:focus,.btn-primary:not(:disabled):not(.disabled).active:focus,.show>.btn-primary.dropdown-toggle:focus{box-shadow:0 0 0 .2rem rgba(38,182,172,.5)}.btn-secondary{color:#fff;background-color:#6c757d;border-color:#6c757d}.btn-secondary:hover{color:#fff;background-color:#5a6268;border-color:#545b62}.btn-secondary:focus,.btn-secondary.focus{color:#fff;background-color:#5a6268;border-color:#545b62;box-shadow:0 0 0 .2rem rgba(130,138,145,.5)}.btn-secondary.disabled,.btn-secondary:disabled{color:#fff;background-color:#6c757d;border-color:#6c757d}.btn-secondary:not(:disabled):not(.disabled):active,.btn-secondary:not(:disabled):not(.disabled).active,.show>.btn-secondary.dropdown-toggle{color:#fff;background-color:#545b62;border-color:#4e555b}.btn-secondary:not(:disabled):not(.disabled):active:focus,.btn-secondary:not(:disabled):not(.disabled).active:focus,.show>.btn-secondary.dropdown-toggle:focus{box-shadow:0 0 0 .2rem rgba(130,138,145,.5)}.btn-success{color:#fff;background-color:#28a745;border-color:#28a745}.btn-success:hover{color:#fff;background-color:#218838;border-color:#1e7e34}.btn-success:focus,.btn-success.focus{color:#fff;background-color:#218838;border-color:#1e7e34;box-shadow:0 0 0 .2rem rgba(72,180,97,.5)}.btn-success.disabled,.btn-success:disabled{color:#fff;background-color:#28a745;border-color:#28a745}.btn-success:not(:disabled):not(.disabled):active,.btn-success:not(:disabled):not(.disabled).active,.show>.btn-success.dropdown-toggle{color:#fff;background-color:#1e7e34;border-color:#1c7430}.btn-success:not(:disabled):not(.disabled):active:focus,.btn-success:not(:disabled):not(.disabled).active:focus,.show>.btn-success.dropdown-toggle:focus{box-shadow:0 0 0 .2rem rgba(72,180,97,.5)}.btn-info{color:#fff;background-color:#17a2b8;border-color:#17a2b8}.btn-info:hover{color:#fff;background-color:#138496;border-color:#117a8b}.btn-info:focus,.btn-info.focus{color:#fff;background-color:#138496;border-color:#117a8b;box-shadow:0 0 0 .2rem rgba(58,176,195,.5)}.btn-info.disabled,.btn-info:disabled{color:#fff;background-color:#17a2b8;border-color:#17a2b8}.btn-info:not(:disabled):not(.disabled):active,.btn-info:not(:disabled):not(.disabled).active,.show>.btn-info.dropdown-toggle{color:#fff;background-color:#117a8b;border-color:#10707f}.btn-info:not(:disabled):not(.disabled):active:focus,.btn-info:not(:disabled):not(.disabled).active:focus,.show>.btn-info.dropdown-toggle:focus{box-shadow:0 0 0 .2rem rgba(58,176,195,.5)}.btn-warning{color:#212529;background-color:#ffc107;border-color:#ffc107}.btn-warning:hover{color:#212529;background-color:#e0a800;border-color:#d39e00}.btn-warning:focus,.btn-warning.focus{color:#212529;background-color:#e0a800;border-color:#d39e00;box-shadow:0 0 0 .2rem rgba(222,170,12,.5)}.btn-warning.disabled,.btn-warning:disabled{color:#212529;background-color:#ffc107;border-color:#ffc107}.btn-warning:not(:disabled):not(.disabled):active,.btn-warning:not(:disabled):not(.disabled).active,.show>.btn-warning.dropdown-toggle{color:#212529;background-color:#d39e00;border-color:#c69500}.btn-warning:not(:disabled):not(.disabled):active:focus,.btn-warning:not(:disabled):not(.disabled).active:focus,.show>.btn-warning.dropdown-toggle:focus{box-shadow:0 0 0 .2rem rgba(222,170,12,.5)}.btn-danger{color:#fff;background-color:#dc3545;border-color:#dc3545}.btn-danger:hover{color:#fff;background-color:#c82333;border-color:#bd2130}.btn-danger:focus,.btn-danger.focus{color:#fff;background-color:#c82333;border-color:#bd2130;box-shadow:0 0 0 .2rem rgba(225,83,97,.5)}.btn-danger.disabled,.btn-danger:disabled{color:#fff;background-color:#dc3545;border-color:#dc3545}.btn-danger:not(:disabled):not(.disabled):active,.btn-danger:not(:disabled):not(.disabled).active,.show>.btn-danger.dropdown-toggle{color:#fff;background-color:#bd2130;border-color:#b21f2d}.btn-danger:not(:disabled):not(.disabled):active:focus,.btn-danger:not(:disabled):not(.disabled).active:focus,.show>.btn-danger.dropdown-toggle:focus{box-shadow:0 0 0 .2rem rgba(225,83,97,.5)}.btn-light{color:#212529;background-color:#f8f9fa;border-color:#f8f9fa}.btn-light:hover{color:#212529;background-color:#e2e6ea;border-color:#dae0e5}.btn-light:focus,.btn-light.focus{color:#212529;background-color:#e2e6ea;border-color:#dae0e5;box-shadow:0 0 0 .2rem rgba(216,217,219,.5)}.btn-light.disabled,.btn-light:disabled{color:#212529;background-color:#f8f9fa;border-color:#f8f9fa}.btn-light:not(:disabled):not(.disabled):active,.btn-light:not(:disabled):not(.disabled).active,.show>.btn-light.dropdown-toggle{color:#212529;background-color:#dae0e5;border-color:#d3d9df}.btn-light:not(:disabled):not(.disabled):active:focus,.btn-light:not(:disabled):not(.disabled).active:focus,.show>.btn-light.dropdown-toggle:focus{box-shadow:0 0 0 .2rem rgba(216,217,219,.5)}.btn-dark{color:#fff;background-color:#343a40;border-color:#343a40}.btn-dark:hover{color:#fff;background-color:#23272b;border-color:#1d2124}.btn-dark:focus,.btn-dark.focus{color:#fff;background-color:#23272b;border-color:#1d2124;box-shadow:0 0 0 .2rem rgba(82,88,93,.5)}.btn-dark.disabled,.btn-dark:disabled{color:#fff;background-color:#343a40;border-color:#343a40}.btn-dark:not(:disabled):not(.disabled):active,.btn-dark:not(:disabled):not(.disabled).active,.show>.btn-dark.dropdown-toggle{color:#fff;background-color:#1d2124;border-color:#171a1d}.btn-dark:not(:disabled):not(.disabled):active:focus,.btn-dark:not(:disabled):not(.disabled).active:focus,.show>.btn-dark.dropdown-toggle:focus{box-shadow:0 0 0 .2rem rgba(82,88,93,.5)}.btn-outline-primary{color:#00a99d;border-color:#00a99d}.btn-outline-primary:hover{color:#fff;background-color:#00a99d;border-color:#00a99d}.btn-outline-primary:focus,.btn-outline-primary.focus{box-shadow:0 0 0 .2rem rgba(0,169,157,.5)}.btn-outline-primary.disabled,.btn-outline-primary:disabled{color:#00a99d;background-color:transparent}.btn-outline-primary:not(:disabled):not(.disabled):active,.btn-outline-primary:not(:disabled):not(.disabled).active,.show>.btn-outline-primary.dropdown-toggle{color:#fff;background-color:#00a99d;border-color:#00a99d}.btn-outline-primary:not(:disabled):not(.disabled):active:focus,.btn-outline-primary:not(:disabled):not(.disabled).active:focus,.show>.btn-outline-primary.dropdown-toggle:focus{box-shadow:0 0 0 .2rem rgba(0,169,157,.5)}.btn-outline-secondary{color:#6c757d;border-color:#6c757d}.btn-outline-secondary:hover{color:#fff;background-color:#6c757d;border-color:#6c757d}.btn-outline-secondary:focus,.btn-outline-secondary.focus{box-shadow:0 0 0 .2rem rgba(108,117,125,.5)}.btn-outline-secondary.disabled,.btn-outline-secondary:disabled{color:#6c757d;background-color:transparent}.btn-outline-secondary:not(:disabled):not(.disabled):active,.btn-outline-secondary:not(:disabled):not(.disabled).active,.show>.btn-outline-secondary.dropdown-toggle{color:#fff;background-color:#6c757d;border-color:#6c757d}.btn-outline-secondary:not(:disabled):not(.disabled):active:focus,.btn-outline-secondary:not(:disabled):not(.disabled).active:focus,.show>.btn-outline-secondary.dropdown-toggle:focus{box-shadow:0 0 0 .2rem rgba(108,117,125,.5)}.btn-outline-success{color:#28a745;border-color:#28a745}.btn-outline-success:hover{color:#fff;background-color:#28a745;border-color:#28a745}.btn-outline-success:focus,.btn-outline-success.focus{box-shadow:0 0 0 .2rem rgba(40,167,69,.5)}.btn-outline-success.disabled,.btn-outline-success:disabled{color:#28a745;background-color:transparent}.btn-outline-success:not(:disabled):not(.disabled):active,.btn-outline-success:not(:disabled):not(.disabled).active,.show>.btn-outline-success.dropdown-toggle{color:#fff;background-color:#28a745;border-color:#28a745}.btn-outline-success:not(:disabled):not(.disabled):active:focus,.btn-outline-success:not(:disabled):not(.disabled).active:focus,.show>.btn-outline-success.dropdown-toggle:focus{box-shadow:0 0 0 .2rem rgba(40,167,69,.5)}.btn-outline-info{color:#17a2b8;border-color:#17a2b8}.btn-outline-info:hover{color:#fff;background-color:#17a2b8;border-color:#17a2b8}.btn-outline-info:focus,.btn-outline-info.focus{box-shadow:0 0 0 .2rem rgba(23,162,184,.5)}.btn-outline-info.disabled,.btn-outline-info:disabled{color:#17a2b8;background-color:transparent}.btn-outline-info:not(:disabled):not(.disabled):active,.btn-outline-info:not(:disabled):not(.disabled).active,.show>.btn-outline-info.dropdown-toggle{color:#fff;background-color:#17a2b8;border-color:#17a2b8}.btn-outline-info:not(:disabled):not(.disabled):active:focus,.btn-outline-info:not(:disabled):not(.disabled).active:focus,.show>.btn-outline-info.dropdown-toggle:focus{box-shadow:0 0 0 .2rem rgba(23,162,184,.5)}.btn-outline-warning{color:#ffc107;border-color:#ffc107}.btn-outline-warning:hover{color:#212529;background-color:#ffc107;border-color:#ffc107}.btn-outline-warning:focus,.btn-outline-warning.focus{box-shadow:0 0 0 .2rem rgba(255,193,7,.5)}.btn-outline-warning.disabled,.btn-outline-warning:disabled{color:#ffc107;background-color:transparent}.btn-outline-warning:not(:disabled):not(.disabled):active,.btn-outline-warning:not(:disabled):not(.disabled).active,.show>.btn-outline-warning.dropdown-toggle{color:#212529;background-color:#ffc107;border-color:#ffc107}.btn-outline-warning:not(:disabled):not(.disabled):active:focus,.btn-outline-warning:not(:disabled):not(.disabled).active:focus,.show>.btn-outline-warning.dropdown-toggle:focus{box-shadow:0 0 0 .2rem rgba(255,193,7,.5)}.btn-outline-danger{color:#dc3545;border-color:#dc3545}.btn-outline-danger:hover{color:#fff;background-color:#dc3545;border-color:#dc3545}.btn-outline-danger:focus,.btn-outline-danger.focus{box-shadow:0 0 0 .2rem rgba(220,53,69,.5)}.btn-outline-danger.disabled,.btn-outline-danger:disabled{color:#dc3545;background-color:transparent}.btn-outline-danger:not(:disabled):not(.disabled):active,.btn-outline-danger:not(:disabled):not(.disabled).active,.show>.btn-outline-danger.dropdown-toggle{color:#fff;background-color:#dc3545;border-color:#dc3545}.btn-outline-danger:not(:disabled):not(.disabled):active:focus,.btn-outline-danger:not(:disabled):not(.disabled).active:focus,.show>.btn-outline-danger.dropdown-toggle:focus{box-shadow:0 0 0 .2rem rgba(220,53,69,.5)}.btn-outline-light{color:#f8f9fa;border-color:#f8f9fa}.btn-outline-light:hover{color:#212529;background-color:#f8f9fa;border-color:#f8f9fa}.btn-outline-light:focus,.btn-outline-light.focus{box-shadow:0 0 0 .2rem rgba(248,249,250,.5)}.btn-outline-light.disabled,.btn-outline-light:disabled{color:#f8f9fa;background-color:transparent}.btn-outline-light:not(:disabled):not(.disabled):active,.btn-outline-light:not(:disabled):not(.disabled).active,.show>.btn-outline-light.dropdown-toggle{color:#212529;background-color:#f8f9fa;border-color:#f8f9fa}.btn-outline-light:not(:disabled):not(.disabled):active:focus,.btn-outline-light:not(:disabled):not(.disabled).active:focus,.show>.btn-outline-light.dropdown-toggle:focus{box-shadow:0 0 0 .2rem rgba(248,249,250,.5)}.btn-outline-dark{color:#343a40;border-color:#343a40}.btn-outline-dark:hover{color:#fff;background-color:#343a40;border-color:#343a40}.btn-outline-dark:focus,.btn-outline-dark.focus{box-shadow:0 0 0 .2rem rgba(52,58,64,.5)}.btn-outline-dark.disabled,.btn-outline-dark:disabled{color:#343a40;background-color:transparent}.btn-outline-dark:not(:disabled):not(.disabled):active,.btn-outline-dark:not(:disabled):not(.disabled).active,.show>.btn-outline-dark.dropdown-toggle{color:#fff;background-color:#343a40;border-color:#343a40}.btn-outline-dark:not(:disabled):not(.disabled):active:focus,.btn-outline-dark:not(:disabled):not(.disabled).active:focus,.show>.btn-outline-dark.dropdown-toggle:focus{box-shadow:0 0 0 .2rem rgba(52,58,64,.5)}.btn-link{font-weight:400;color:#00a99d;text-decoration:none}.btn-link:hover{color:#005d56;text-decoration:underline}.btn-link:focus,.btn-link.focus{text-decoration:underline}.btn-link:disabled,.btn-link.disabled{color:#6c757d;pointer-events:none}.btn-lg,.btn-group-lg>.btn{padding:.5rem 1rem;font-size:1.25rem;line-height:1.5;border-radius:.3rem}.btn-sm,.btn-group-sm>.btn{padding:.25rem .5rem;font-size:0.875rem;line-height:1.5;border-radius:.2rem}.btn-block{display:block;width:100%}.btn-block+.btn-block{margin-top:.5rem}input[type=submit].btn-block,input[type=reset].btn-block,input[type=button].btn-block{width:100%}.fade{transition:opacity .15s linear}@media(prefers-reduced-motion: reduce){.fade{transition:none}}.fade:not(.show){opacity:0}.collapse:not(.show){display:none}.collapsing{position:relative;height:0;overflow:hidden;transition:height .35s ease}@media(prefers-reduced-motion: reduce){.collapsing{transition:none}}.dropup,.dropright,.dropdown,.dropleft{position:relative}.dropdown-toggle{white-space:nowrap}.dropdown-toggle::after{display:inline-block;margin-left:.255em;vertical-align:.255em;content:"";border-top:.3em solid;border-right:.3em solid transparent;border-bottom:0;border-left:.3em solid transparent}.dropdown-toggle:empty::after{margin-left:0}.dropdown-menu{position:absolute;top:100%;left:0;z-index:1000;display:none;float:left;min-width:10rem;padding:.5rem 0;margin:.125rem 0 0;font-size:1rem;color:#212529;text-align:left;list-style:none;background-color:#fff;background-clip:padding-box;border:1px solid rgba(0,0,0,.15);border-radius:.25rem}.dropdown-menu-left{right:auto;left:0}.dropdown-menu-right{right:0;left:auto}@media(min-width: 576px){.dropdown-menu-sm-left{right:auto;left:0}.dropdown-menu-sm-right{right:0;left:auto}}@media(min-width: 768px){.dropdown-menu-md-left{right:auto;left:0}.dropdown-menu-md-right{right:0;left:auto}}@media(min-width: 992px){.dropdown-menu-lg-left{right:auto;left:0}.dropdown-menu-lg-right{right:0;left:auto}}@media(min-width: 1200px){.dropdown-menu-xl-left{right:auto;left:0}.dropdown-menu-xl-right{right:0;left:auto}}.dropup .dropdown-menu{top:auto;bottom:100%;margin-top:0;margin-bottom:.125rem}.dropup .dropdown-toggle::after{display:inline-block;margin-left:.255em;vertical-align:.255em;content:"";border-top:0;border-right:.3em solid transparent;border-bottom:.3em solid;border-left:.3em solid transparent}.dropup .dropdown-toggle:empty::after{margin-left:0}.dropright .dropdown-menu{top:0;right:auto;left:100%;margin-top:0;margin-left:.125rem}.dropright .dropdown-toggle::after{display:inline-block;margin-left:.255em;vertical-align:.255em;content:"";border-top:.3em solid transparent;border-right:0;border-bottom:.3em solid transparent;border-left:.3em solid}.dropright .dropdown-toggle:empty::after{margin-left:0}.dropright .dropdown-toggle::after{vertical-align:0}.dropleft .dropdown-menu{top:0;right:100%;left:auto;margin-top:0;margin-right:.125rem}.dropleft .dropdown-toggle::after{display:inline-block;margin-left:.255em;vertical-align:.255em;content:""}.dropleft .dropdown-toggle::after{display:none}.dropleft .dropdown-toggle::before{display:inline-block;margin-right:.255em;vertical-align:.255em;content:"";border-top:.3em solid transparent;border-right:.3em solid;border-bottom:.3em solid transparent}.dropleft .dropdown-toggle:empty::after{margin-left:0}.dropleft .dropdown-toggle::before{vertical-align:0}.dropdown-menu[x-placement^=top],.dropdown-menu[x-placement^=right],.dropdown-menu[x-placement^=bottom],.dropdown-menu[x-placement^=left]{right:auto;bottom:auto}.dropdown-divider{height:0;margin:.5rem 0;overflow:hidden;border-top:1px solid #e9ecef}.dropdown-item{display:block;width:100%;padding:.25rem 1.5rem;clear:both;font-weight:400;color:#212529;text-align:inherit;white-space:nowrap;background-color:transparent;border:0}.dropdown-item:hover,.dropdown-item:focus{color:#16181b;text-decoration:none;background-color:#e9ecef}.dropdown-item.active,.dropdown-item:active{color:#fff;text-decoration:none;background-color:#00a99d}.dropdown-item.disabled,.dropdown-item:disabled{color:#adb5bd;pointer-events:none;background-color:transparent}.dropdown-menu.show{display:block}.dropdown-header{display:block;padding:.5rem 1.5rem;margin-bottom:0;font-size:0.875rem;color:#6c757d;white-space:nowrap}.dropdown-item-text{display:block;padding:.25rem 1.5rem;color:#212529}.btn-group,.btn-group-vertical{position:relative;display:inline-flex;vertical-align:middle}.btn-group>.btn,.btn-group-vertical>.btn{position:relative;flex:1 1 auto}.btn-group>.btn:hover,.btn-group-vertical>.btn:hover{z-index:1}.btn-group>.btn:focus,.btn-group>.btn:active,.btn-group>.btn.active,.btn-group-vertical>.btn:focus,.btn-group-vertical>.btn:active,.btn-group-vertical>.btn.active{z-index:1}.btn-toolbar{display:flex;flex-wrap:wrap;justify-content:flex-start}.btn-toolbar .input-group{width:auto}.btn-group>.btn:not(:first-child),.btn-group>.btn-group:not(:first-child){margin-left:-1px}.btn-group>.btn:not(:last-child):not(.dropdown-toggle),.btn-group>.btn-group:not(:last-child)>.btn{border-top-right-radius:0;border-bottom-right-radius:0}.btn-group>.btn:not(:first-child),.btn-group>.btn-group:not(:first-child)>.btn{border-top-left-radius:0;border-bottom-left-radius:0}.dropdown-toggle-split{padding-right:.5625rem;padding-left:.5625rem}.dropdown-toggle-split::after,.dropup .dropdown-toggle-split::after,.dropright .dropdown-toggle-split::after{margin-left:0}.dropleft .dropdown-toggle-split::before{margin-right:0}.btn-sm+.dropdown-toggle-split,.btn-group-sm>.btn+.dropdown-toggle-split{padding-right:.375rem;padding-left:.375rem}.btn-lg+.dropdown-toggle-split,.btn-group-lg>.btn+.dropdown-toggle-split{padding-right:.75rem;padding-left:.75rem}.btn-group-vertical{flex-direction:column;align-items:flex-start;justify-content:center}.btn-group-vertical>.btn,.btn-group-vertical>.btn-group{width:100%}.btn-group-vertical>.btn:not(:first-child),.btn-group-vertical>.btn-group:not(:first-child){margin-top:-1px}.btn-group-vertical>.btn:not(:last-child):not(.dropdown-toggle),.btn-group-vertical>.btn-group:not(:last-child)>.btn{border-bottom-right-radius:0;border-bottom-left-radius:0}.btn-group-vertical>.btn:not(:first-child),.btn-group-vertical>.btn-group:not(:first-child)>.btn{border-top-left-radius:0;border-top-right-radius:0}.btn-group-toggle>.btn,.btn-group-toggle>.btn-group>.btn{margin-bottom:0}.btn-group-toggle>.btn input[type=radio],.btn-group-toggle>.btn input[type=checkbox],.btn-group-toggle>.btn-group>.btn input[type=radio],.btn-group-toggle>.btn-group>.btn input[type=checkbox]{position:absolute;clip:rect(0, 0, 0, 0);pointer-events:none}.input-group{position:relative;display:flex;flex-wrap:wrap;align-items:stretch;width:100%}.input-group>.form-control,.input-group>.form-control-plaintext,.input-group>.custom-select,.input-group>.custom-file{position:relative;flex:1 1 auto;width:1%;min-width:0;margin-bottom:0}.input-group>.form-control+.form-control,.input-group>.form-control+.custom-select,.input-group>.form-control+.custom-file,.input-group>.form-control-plaintext+.form-control,.input-group>.form-control-plaintext+.custom-select,.input-group>.form-control-plaintext+.custom-file,.input-group>.custom-select+.form-control,.input-group>.custom-select+.custom-select,.input-group>.custom-select+.custom-file,.input-group>.custom-file+.form-control,.input-group>.custom-file+.custom-select,.input-group>.custom-file+.custom-file{margin-left:-1px}.input-group>.form-control:focus,.input-group>.custom-select:focus,.input-group>.custom-file .custom-file-input:focus~.custom-file-label{z-index:3}.input-group>.custom-file .custom-file-input:focus{z-index:4}.input-group>.form-control:not(:first-child),.input-group>.custom-select:not(:first-child){border-top-left-radius:0;border-bottom-left-radius:0}.input-group>.custom-file{display:flex;align-items:center}.input-group>.custom-file:not(:last-child) .custom-file-label,.input-group>.custom-file:not(:first-child) .custom-file-label{border-top-left-radius:0;border-bottom-left-radius:0}.input-group:not(.has-validation)>.form-control:not(:last-child),.input-group:not(.has-validation)>.custom-select:not(:last-child),.input-group:not(.has-validation)>.custom-file:not(:last-child) .custom-file-label::after{border-top-right-radius:0;border-bottom-right-radius:0}.input-group.has-validation>.form-control:nth-last-child(n+3),.input-group.has-validation>.custom-select:nth-last-child(n+3),.input-group.has-validation>.custom-file:nth-last-child(n+3) .custom-file-label::after{border-top-right-radius:0;border-bottom-right-radius:0}.input-group-prepend,.input-group-append{display:flex}.input-group-prepend .btn,.input-group-append .btn{position:relative;z-index:2}.input-group-prepend .btn:focus,.input-group-append .btn:focus{z-index:3}.input-group-prepend .btn+.btn,.input-group-prepend .btn+.input-group-text,.input-group-prepend .input-group-text+.input-group-text,.input-group-prepend .input-group-text+.btn,.input-group-append .btn+.btn,.input-group-append .btn+.input-group-text,.input-group-append .input-group-text+.input-group-text,.input-group-append .input-group-text+.btn{margin-left:-1px}.input-group-prepend{margin-right:-1px}.input-group-append{margin-left:-1px}.input-group-text{display:flex;align-items:center;padding:.375rem .75rem;margin-bottom:0;font-size:1rem;font-weight:400;line-height:1.5;color:#495057;text-align:center;white-space:nowrap;background-color:#e9ecef;border:1px solid #ced4da;border-radius:.25rem}.input-group-text input[type=radio],.input-group-text input[type=checkbox]{margin-top:0}.input-group-lg>.form-control:not(textarea),.input-group-lg>.custom-select{height:calc(1.5em + 1rem + 2px)}.input-group-lg>.form-control,.input-group-lg>.custom-select,.input-group-lg>.input-group-prepend>.input-group-text,.input-group-lg>.input-group-append>.input-group-text,.input-group-lg>.input-group-prepend>.btn,.input-group-lg>.input-group-append>.btn{padding:.5rem 1rem;font-size:1.25rem;line-height:1.5;border-radius:.3rem}.input-group-sm>.form-control:not(textarea),.input-group-sm>.custom-select{height:calc(1.5em + 0.5rem + 2px)}.input-group-sm>.form-control,.input-group-sm>.custom-select,.input-group-sm>.input-group-prepend>.input-group-text,.input-group-sm>.input-group-append>.input-group-text,.input-group-sm>.input-group-prepend>.btn,.input-group-sm>.input-group-append>.btn{padding:.25rem .5rem;font-size:0.875rem;line-height:1.5;border-radius:.2rem}.input-group-lg>.custom-select,.input-group-sm>.custom-select{padding-right:1.75rem}.input-group>.input-group-prepend>.btn,.input-group>.input-group-prepend>.input-group-text,.input-group:not(.has-validation)>.input-group-append:not(:last-child)>.btn,.input-group:not(.has-validation)>.input-group-append:not(:last-child)>.input-group-text,.input-group.has-validation>.input-group-append:nth-last-child(n+3)>.btn,.input-group.has-validation>.input-group-append:nth-last-child(n+3)>.input-group-text,.input-group>.input-group-append:last-child>.btn:not(:last-child):not(.dropdown-toggle),.input-group>.input-group-append:last-child>.input-group-text:not(:last-child){border-top-right-radius:0;border-bottom-right-radius:0}.input-group>.input-group-append>.btn,.input-group>.input-group-append>.input-group-text,.input-group>.input-group-prepend:not(:first-child)>.btn,.input-group>.input-group-prepend:not(:first-child)>.input-group-text,.input-group>.input-group-prepend:first-child>.btn:not(:first-child),.input-group>.input-group-prepend:first-child>.input-group-text:not(:first-child){border-top-left-radius:0;border-bottom-left-radius:0}.custom-control{position:relative;z-index:1;display:block;min-height:1.5rem;padding-left:1.5rem;color-adjust:exact}.custom-control-inline{display:inline-flex;margin-right:1rem}.custom-control-input{position:absolute;left:0;z-index:-1;width:1rem;height:1.25rem;opacity:0}.custom-control-input:checked~.custom-control-label::before{color:#fff;border-color:#00a99d;background-color:#00a99d}.custom-control-input:focus~.custom-control-label::before{box-shadow:0 0 0 .2rem rgba(0,169,157,.25)}.custom-control-input:focus:not(:checked)~.custom-control-label::before{border-color:#2afff0}.custom-control-input:not(:disabled):active~.custom-control-label::before{color:#fff;background-color:#5dfff3;border-color:#5dfff3}.custom-control-input[disabled]~.custom-control-label,.custom-control-input:disabled~.custom-control-label{color:#6c757d}.custom-control-input[disabled]~.custom-control-label::before,.custom-control-input:disabled~.custom-control-label::before{background-color:#e9ecef}.custom-control-label{position:relative;margin-bottom:0;vertical-align:top}.custom-control-label::before{position:absolute;top:.25rem;left:-1.5rem;display:block;width:1rem;height:1rem;pointer-events:none;content:"";background-color:#fff;border:#adb5bd solid 1px}.custom-control-label::after{position:absolute;top:.25rem;left:-1.5rem;display:block;width:1rem;height:1rem;content:"";background:50%/50% 50% no-repeat}.custom-checkbox .custom-control-label::before{border-radius:.25rem}.custom-checkbox .custom-control-input:checked~.custom-control-label::after{background-image:url("data:image/svg+xml,%3csvg xmlns=\'http://www.w3.org/2000/svg\' width=\'8\' height=\'8\' viewBox=\'0 0 8 8\'%3e%3cpath fill=\'%23fff\' d=\'M6.564.75l-3.59 3.612-1.538-1.55L0 4.26l2.974 2.99L8 2.193z\'/%3e%3c/svg%3e")}.custom-checkbox .custom-control-input:indeterminate~.custom-control-label::before{border-color:#00a99d;background-color:#00a99d}.custom-checkbox .custom-control-input:indeterminate~.custom-control-label::after{background-image:url("data:image/svg+xml,%3csvg xmlns=\'http://www.w3.org/2000/svg\' width=\'4\' height=\'4\' viewBox=\'0 0 4 4\'%3e%3cpath stroke=\'%23fff\' d=\'M0 2h4\'/%3e%3c/svg%3e")}.custom-checkbox .custom-control-input:disabled:checked~.custom-control-label::before{background-color:rgba(0,169,157,.5)}.custom-checkbox .custom-control-input:disabled:indeterminate~.custom-control-label::before{background-color:rgba(0,169,157,.5)}.custom-radio .custom-control-label::before{border-radius:50%}.custom-radio .custom-control-input:checked~.custom-control-label::after{background-image:url("data:image/svg+xml,%3csvg xmlns=\'http://www.w3.org/2000/svg\' width=\'12\' height=\'12\' viewBox=\'-4 -4 8 8\'%3e%3ccircle r=\'3\' fill=\'%23fff\'/%3e%3c/svg%3e")}.custom-radio .custom-control-input:disabled:checked~.custom-control-label::before{background-color:rgba(0,169,157,.5)}.custom-switch{padding-left:2.25rem}.custom-switch .custom-control-label::before{left:-2.25rem;width:1.75rem;pointer-events:all;border-radius:.5rem}.custom-switch .custom-control-label::after{top:calc(0.25rem + 2px);left:calc(-2.25rem + 2px);width:calc(1rem - 4px);height:calc(1rem - 4px);background-color:#adb5bd;border-radius:.5rem;transition:transform .15s ease-in-out,background-color .15s ease-in-out,border-color .15s ease-in-out,box-shadow .15s ease-in-out}@media(prefers-reduced-motion: reduce){.custom-switch .custom-control-label::after{transition:none}}.custom-switch .custom-control-input:checked~.custom-control-label::after{background-color:#fff;transform:translateX(0.75rem)}.custom-switch .custom-control-input:disabled:checked~.custom-control-label::before{background-color:rgba(0,169,157,.5)}.custom-select{display:inline-block;width:100%;height:calc(1.5em + 0.75rem + 2px);padding:.375rem 1.75rem .375rem .75rem;font-size:1rem;font-weight:400;line-height:1.5;color:#495057;vertical-align:middle;background:#fff url("data:image/svg+xml,%3csvg xmlns=\'http://www.w3.org/2000/svg\' width=\'4\' height=\'5\' viewBox=\'0 0 4 5\'%3e%3cpath fill=\'%23343a40\' d=\'M2 0L0 2h4zm0 5L0 3h4z\'/%3e%3c/svg%3e") right .75rem center/8px 10px no-repeat;border:1px solid #ced4da;border-radius:.25rem;appearance:none}.custom-select:focus{border-color:#2afff0;outline:0;box-shadow:0 0 0 .2rem rgba(0,169,157,.25)}.custom-select:focus::-ms-value{color:#495057;background-color:#fff}.custom-select[multiple],.custom-select[size]:not([size="1"]){height:auto;padding-right:.75rem;background-image:none}.custom-select:disabled{color:#6c757d;background-color:#e9ecef}.custom-select::-ms-expand{display:none}.custom-select:-moz-focusring{color:transparent;text-shadow:0 0 0 #495057}.custom-select-sm{height:calc(1.5em + 0.5rem + 2px);padding-top:.25rem;padding-bottom:.25rem;padding-left:.5rem;font-size:0.875rem}.custom-select-lg{height:calc(1.5em + 1rem + 2px);padding-top:.5rem;padding-bottom:.5rem;padding-left:1rem;font-size:1.25rem}.custom-file{position:relative;display:inline-block;width:100%;height:calc(1.5em + 0.75rem + 2px);margin-bottom:0}.custom-file-input{position:relative;z-index:2;width:100%;height:calc(1.5em + 0.75rem + 2px);margin:0;overflow:hidden;opacity:0}.custom-file-input:focus~.custom-file-label{border-color:#2afff0;box-shadow:0 0 0 .2rem rgba(0,169,157,.25)}.custom-file-input[disabled]~.custom-file-label,.custom-file-input:disabled~.custom-file-label{background-color:#e9ecef}.custom-file-input:lang(en)~.custom-file-label::after{content:"Browse"}.custom-file-input~.custom-file-label[data-browse]::after{content:attr(data-browse)}.custom-file-label{position:absolute;top:0;right:0;left:0;z-index:1;height:calc(1.5em + 0.75rem + 2px);padding:.375rem .75rem;overflow:hidden;font-weight:400;line-height:1.5;color:#495057;background-color:#fff;border:1px solid #ced4da;border-radius:.25rem}.custom-file-label::after{position:absolute;top:0;right:0;bottom:0;z-index:3;display:block;height:calc(1.5em + 0.75rem);padding:.375rem .75rem;line-height:1.5;color:#495057;content:"Browse";background-color:#e9ecef;border-left:inherit;border-radius:0 .25rem .25rem 0}.custom-range{width:100%;height:1.4rem;padding:0;background-color:transparent;appearance:none}.custom-range:focus{outline:0}.custom-range:focus::-webkit-slider-thumb{box-shadow:0 0 0 1px #fff,0 0 0 .2rem rgba(0,169,157,.25)}.custom-range:focus::-moz-range-thumb{box-shadow:0 0 0 1px #fff,0 0 0 .2rem rgba(0,169,157,.25)}.custom-range:focus::-ms-thumb{box-shadow:0 0 0 1px #fff,0 0 0 .2rem rgba(0,169,157,.25)}.custom-range::-moz-focus-outer{border:0}.custom-range::-webkit-slider-thumb{width:1rem;height:1rem;margin-top:-0.25rem;background-color:#00a99d;border:0;border-radius:1rem;transition:background-color .15s ease-in-out,border-color .15s ease-in-out,box-shadow .15s ease-in-out;appearance:none}@media(prefers-reduced-motion: reduce){.custom-range::-webkit-slider-thumb{transition:none}}.custom-range::-webkit-slider-thumb:active{background-color:#5dfff3}.custom-range::-webkit-slider-runnable-track{width:100%;height:.5rem;color:transparent;cursor:pointer;background-color:#dee2e6;border-color:transparent;border-radius:1rem}.custom-range::-moz-range-thumb{width:1rem;height:1rem;background-color:#00a99d;border:0;border-radius:1rem;transition:background-color .15s ease-in-out,border-color .15s ease-in-out,box-shadow .15s ease-in-out;appearance:none}@media(prefers-reduced-motion: reduce){.custom-range::-moz-range-thumb{transition:none}}.custom-range::-moz-range-thumb:active{background-color:#5dfff3}.custom-range::-moz-range-track{width:100%;height:.5rem;color:transparent;cursor:pointer;background-color:#dee2e6;border-color:transparent;border-radius:1rem}.custom-range::-ms-thumb{width:1rem;height:1rem;margin-top:0;margin-right:.2rem;margin-left:.2rem;background-color:#00a99d;border:0;border-radius:1rem;transition:background-color .15s ease-in-out,border-color .15s ease-in-out,box-shadow .15s ease-in-out;appearance:none}@media(prefers-reduced-motion: reduce){.custom-range::-ms-thumb{transition:none}}.custom-range::-ms-thumb:active{background-color:#5dfff3}.custom-range::-ms-track{width:100%;height:.5rem;color:transparent;cursor:pointer;background-color:transparent;border-color:transparent;border-width:.5rem}.custom-range::-ms-fill-lower{background-color:#dee2e6;border-radius:1rem}.custom-range::-ms-fill-upper{margin-right:15px;background-color:#dee2e6;border-radius:1rem}.custom-range:disabled::-webkit-slider-thumb{background-color:#adb5bd}.custom-range:disabled::-webkit-slider-runnable-track{cursor:default}.custom-range:disabled::-moz-range-thumb{background-color:#adb5bd}.custom-range:disabled::-moz-range-track{cursor:default}.custom-range:disabled::-ms-thumb{background-color:#adb5bd}.custom-control-label::before,.custom-file-label,.custom-select{transition:background-color .15s ease-in-out,border-color .15s ease-in-out,box-shadow .15s ease-in-out}@media(prefers-reduced-motion: reduce){.custom-control-label::before,.custom-file-label,.custom-select{transition:none}}.nav{display:flex;flex-wrap:wrap;padding-left:0;margin-bottom:0;list-style:none}.nav-link{display:block;padding:.5rem 1rem}.nav-link:hover,.nav-link:focus{text-decoration:none}.nav-link.disabled{color:#6c757d;pointer-events:none;cursor:default}.nav-tabs{border-bottom:1px solid #dee2e6}.nav-tabs .nav-link{margin-bottom:-1px;border:1px solid transparent;border-top-left-radius:.25rem;border-top-right-radius:.25rem}.nav-tabs .nav-link:hover,.nav-tabs .nav-link:focus{border-color:#e9ecef #e9ecef #dee2e6}.nav-tabs .nav-link.disabled{color:#6c757d;background-color:transparent;border-color:transparent}.nav-tabs .nav-link.active,.nav-tabs .nav-item.show .nav-link{color:#495057;background-color:#fff;border-color:#dee2e6 #dee2e6 #fff}.nav-tabs .dropdown-menu{margin-top:-1px;border-top-left-radius:0;border-top-right-radius:0}.nav-pills .nav-link{border-radius:.25rem}.nav-pills .nav-link.active,.nav-pills .show>.nav-link{color:#fff;background-color:#00a99d}.nav-fill>.nav-link,.nav-fill .nav-item{flex:1 1 auto;text-align:center}.nav-justified>.nav-link,.nav-justified .nav-item{flex-basis:0;flex-grow:1;text-align:center}.tab-content>.tab-pane{display:none}.tab-content>.active{display:block}.navbar{position:relative;display:flex;flex-wrap:wrap;align-items:center;justify-content:space-between;padding:.5rem 1rem}.navbar .container,.navbar .container-fluid,.navbar .container-sm,.navbar .container-md,.navbar .container-lg,.navbar .container-xl{display:flex;flex-wrap:wrap;align-items:center;justify-content:space-between}.navbar-brand{display:inline-block;padding-top:.3125rem;padding-bottom:.3125rem;margin-right:1rem;font-size:1.25rem;line-height:inherit;white-space:nowrap}.navbar-brand:hover,.navbar-brand:focus{text-decoration:none}.navbar-nav{display:flex;flex-direction:column;padding-left:0;margin-bottom:0;list-style:none}.navbar-nav .nav-link{padding-right:0;padding-left:0}.navbar-nav .dropdown-menu{position:static;float:none}.navbar-text{display:inline-block;padding-top:.5rem;padding-bottom:.5rem}.navbar-collapse{flex-basis:100%;flex-grow:1;align-items:center}.navbar-toggler{padding:.25rem .75rem;font-size:1.25rem;line-height:1;background-color:transparent;border:1px solid transparent;border-radius:.25rem}.navbar-toggler:hover,.navbar-toggler:focus{text-decoration:none}.navbar-toggler-icon{display:inline-block;width:1.5em;height:1.5em;vertical-align:middle;content:"";background:50%/100% 100% no-repeat}.navbar-nav-scroll{max-height:75vh;overflow-y:auto}@media(max-width: 575.98px){.navbar-expand-sm>.container,.navbar-expand-sm>.container-fluid,.navbar-expand-sm>.container-sm,.navbar-expand-sm>.container-md,.navbar-expand-sm>.container-lg,.navbar-expand-sm>.container-xl{padding-right:0;padding-left:0}}@media(min-width: 576px){.navbar-expand-sm{flex-flow:row nowrap;justify-content:flex-start}.navbar-expand-sm .navbar-nav{flex-direction:row}.navbar-expand-sm .navbar-nav .dropdown-menu{position:absolute}.navbar-expand-sm .navbar-nav .nav-link{padding-right:.5rem;padding-left:.5rem}.navbar-expand-sm>.container,.navbar-expand-sm>.container-fluid,.navbar-expand-sm>.container-sm,.navbar-expand-sm>.container-md,.navbar-expand-sm>.container-lg,.navbar-expand-sm>.container-xl{flex-wrap:nowrap}.navbar-expand-sm .navbar-nav-scroll{overflow:visible}.navbar-expand-sm .navbar-collapse{display:flex !important;flex-basis:auto}.navbar-expand-sm .navbar-toggler{display:none}}@media(max-width: 767.98px){.navbar-expand-md>.container,.navbar-expand-md>.container-fluid,.navbar-expand-md>.container-sm,.navbar-expand-md>.container-md,.navbar-expand-md>.container-lg,.navbar-expand-md>.container-xl{padding-right:0;padding-left:0}}@media(min-width: 768px){.navbar-expand-md{flex-flow:row nowrap;justify-content:flex-start}.navbar-expand-md .navbar-nav{flex-direction:row}.navbar-expand-md .navbar-nav .dropdown-menu{position:absolute}.navbar-expand-md .navbar-nav .nav-link{padding-right:.5rem;padding-left:.5rem}.navbar-expand-md>.container,.navbar-expand-md>.container-fluid,.navbar-expand-md>.container-sm,.navbar-expand-md>.container-md,.navbar-expand-md>.container-lg,.navbar-expand-md>.container-xl{flex-wrap:nowrap}.navbar-expand-md .navbar-nav-scroll{overflow:visible}.navbar-expand-md .navbar-collapse{display:flex !important;flex-basis:auto}.navbar-expand-md .navbar-toggler{display:none}}@media(max-width: 991.98px){.navbar-expand-lg>.container,.navbar-expand-lg>.container-fluid,.navbar-expand-lg>.container-sm,.navbar-expand-lg>.container-md,.navbar-expand-lg>.container-lg,.navbar-expand-lg>.container-xl{padding-right:0;padding-left:0}}@media(min-width: 992px){.navbar-expand-lg{flex-flow:row nowrap;justify-content:flex-start}.navbar-expand-lg .navbar-nav{flex-direction:row}.navbar-expand-lg .navbar-nav .dropdown-menu{position:absolute}.navbar-expand-lg .navbar-nav .nav-link{padding-right:.5rem;padding-left:.5rem}.navbar-expand-lg>.container,.navbar-expand-lg>.container-fluid,.navbar-expand-lg>.container-sm,.navbar-expand-lg>.container-md,.navbar-expand-lg>.container-lg,.navbar-expand-lg>.container-xl{flex-wrap:nowrap}.navbar-expand-lg .navbar-nav-scroll{overflow:visible}.navbar-expand-lg .navbar-collapse{display:flex !important;flex-basis:auto}.navbar-expand-lg .navbar-toggler{display:none}}@media(max-width: 1199.98px){.navbar-expand-xl>.container,.navbar-expand-xl>.container-fluid,.navbar-expand-xl>.container-sm,.navbar-expand-xl>.container-md,.navbar-expand-xl>.container-lg,.navbar-expand-xl>.container-xl{padding-right:0;padding-left:0}}@media(min-width: 1200px){.navbar-expand-xl{flex-flow:row nowrap;justify-content:flex-start}.navbar-expand-xl .navbar-nav{flex-direction:row}.navbar-expand-xl .navbar-nav .dropdown-menu{position:absolute}.navbar-expand-xl .navbar-nav .nav-link{padding-right:.5rem;padding-left:.5rem}.navbar-expand-xl>.container,.navbar-expand-xl>.container-fluid,.navbar-expand-xl>.container-sm,.navbar-expand-xl>.container-md,.navbar-expand-xl>.container-lg,.navbar-expand-xl>.container-xl{flex-wrap:nowrap}.navbar-expand-xl .navbar-nav-scroll{overflow:visible}.navbar-expand-xl .navbar-collapse{display:flex !important;flex-basis:auto}.navbar-expand-xl .navbar-toggler{display:none}}.navbar-expand{flex-flow:row nowrap;justify-content:flex-start}.navbar-expand>.container,.navbar-expand>.container-fluid,.navbar-expand>.container-sm,.navbar-expand>.container-md,.navbar-expand>.container-lg,.navbar-expand>.container-xl{padding-right:0;padding-left:0}.navbar-expand .navbar-nav{flex-direction:row}.navbar-expand .navbar-nav .dropdown-menu{position:absolute}.navbar-expand .navbar-nav .nav-link{padding-right:.5rem;padding-left:.5rem}.navbar-expand>.container,.navbar-expand>.container-fluid,.navbar-expand>.container-sm,.navbar-expand>.container-md,.navbar-expand>.container-lg,.navbar-expand>.container-xl{flex-wrap:nowrap}.navbar-expand .navbar-nav-scroll{overflow:visible}.navbar-expand .navbar-collapse{display:flex !important;flex-basis:auto}.navbar-expand .navbar-toggler{display:none}.navbar-light .navbar-brand{color:rgba(0,0,0,.9)}.navbar-light .navbar-brand:hover,.navbar-light .navbar-brand:focus{color:rgba(0,0,0,.9)}.navbar-light .navbar-nav .nav-link{color:rgba(0,0,0,.5)}.navbar-light .navbar-nav .nav-link:hover,.navbar-light .navbar-nav .nav-link:focus{color:rgba(0,0,0,.7)}.navbar-light .navbar-nav .nav-link.disabled{color:rgba(0,0,0,.3)}.navbar-light .navbar-nav .show>.nav-link,.navbar-light .navbar-nav .active>.nav-link,.navbar-light .navbar-nav .nav-link.show,.navbar-light .navbar-nav .nav-link.active{color:rgba(0,0,0,.9)}.navbar-light .navbar-toggler{color:rgba(0,0,0,.5);border-color:rgba(0,0,0,.1)}.navbar-light .navbar-toggler-icon{background-image:url("data:image/svg+xml,%3csvg xmlns=\'http://www.w3.org/2000/svg\' width=\'30\' height=\'30\' viewBox=\'0 0 30 30\'%3e%3cpath stroke=\'rgba%280, 0, 0, 0.5%29\' stroke-linecap=\'round\' stroke-miterlimit=\'10\' stroke-width=\'2\' d=\'M4 7h22M4 15h22M4 23h22\'/%3e%3c/svg%3e")}.navbar-light .navbar-text{color:rgba(0,0,0,.5)}.navbar-light .navbar-text a{color:rgba(0,0,0,.9)}.navbar-light .navbar-text a:hover,.navbar-light .navbar-text a:focus{color:rgba(0,0,0,.9)}.navbar-dark .navbar-brand{color:#fff}.navbar-dark .navbar-brand:hover,.navbar-dark .navbar-brand:focus{color:#fff}.navbar-dark .navbar-nav .nav-link{color:rgba(255,255,255,.5)}.navbar-dark .navbar-nav .nav-link:hover,.navbar-dark .navbar-nav .nav-link:focus{color:rgba(255,255,255,.75)}.navbar-dark .navbar-nav .nav-link.disabled{color:rgba(255,255,255,.25)}.navbar-dark .navbar-nav .show>.nav-link,.navbar-dark .navbar-nav .active>.nav-link,.navbar-dark .navbar-nav .nav-link.show,.navbar-dark .navbar-nav .nav-link.active{color:#fff}.navbar-dark .navbar-toggler{color:rgba(255,255,255,.5);border-color:rgba(255,255,255,.1)}.navbar-dark .navbar-toggler-icon{background-image:url("data:image/svg+xml,%3csvg xmlns=\'http://www.w3.org/2000/svg\' width=\'30\' height=\'30\' viewBox=\'0 0 30 30\'%3e%3cpath stroke=\'rgba%28255, 255, 255, 0.5%29\' stroke-linecap=\'round\' stroke-miterlimit=\'10\' stroke-width=\'2\' d=\'M4 7h22M4 15h22M4 23h22\'/%3e%3c/svg%3e")}.navbar-dark .navbar-text{color:rgba(255,255,255,.5)}.navbar-dark .navbar-text a{color:#fff}.navbar-dark .navbar-text a:hover,.navbar-dark .navbar-text a:focus{color:#fff}.card{position:relative;display:flex;flex-direction:column;min-width:0;word-wrap:break-word;background-color:#fff;background-clip:border-box;border:1px solid rgba(0,0,0,.125);border-radius:.25rem}.card>hr{margin-right:0;margin-left:0}.card>.list-group{border-top:inherit;border-bottom:inherit}.card>.list-group:first-child{border-top-width:0;border-top-left-radius:calc(0.25rem - 1px);border-top-right-radius:calc(0.25rem - 1px)}.card>.list-group:last-child{border-bottom-width:0;border-bottom-right-radius:calc(0.25rem - 1px);border-bottom-left-radius:calc(0.25rem - 1px)}.card>.card-header+.list-group,.card>.list-group+.card-footer{border-top:0}.card-body{flex:1 1 auto;min-height:1px;padding:1.25rem}.card-title{margin-bottom:.75rem}.card-subtitle{margin-top:-0.375rem;margin-bottom:0}.card-text:last-child{margin-bottom:0}.card-link:hover{text-decoration:none}.card-link+.card-link{margin-left:1.25rem}.card-header{padding:.75rem 1.25rem;margin-bottom:0;background-color:rgba(0,0,0,.03);border-bottom:1px solid rgba(0,0,0,.125)}.card-header:first-child{border-radius:calc(0.25rem - 1px) calc(0.25rem - 1px) 0 0}.card-footer{padding:.75rem 1.25rem;background-color:rgba(0,0,0,.03);border-top:1px solid rgba(0,0,0,.125)}.card-footer:last-child{border-radius:0 0 calc(0.25rem - 1px) calc(0.25rem - 1px)}.card-header-tabs{margin-right:-0.625rem;margin-bottom:-0.75rem;margin-left:-0.625rem;border-bottom:0}.card-header-pills{margin-right:-0.625rem;margin-left:-0.625rem}.card-img-overlay{position:absolute;top:0;right:0;bottom:0;left:0;padding:1.25rem;border-radius:calc(0.25rem - 1px)}.card-img,.card-img-top,.card-img-bottom{flex-shrink:0;width:100%}.card-img,.card-img-top{border-top-left-radius:calc(0.25rem - 1px);border-top-right-radius:calc(0.25rem - 1px)}.card-img,.card-img-bottom{border-bottom-right-radius:calc(0.25rem - 1px);border-bottom-left-radius:calc(0.25rem - 1px)}.card-deck .card{margin-bottom:15px}@media(min-width: 576px){.card-deck{display:flex;flex-flow:row wrap;margin-right:-15px;margin-left:-15px}.card-deck .card{flex:1 0 0%;margin-right:15px;margin-bottom:0;margin-left:15px}}.card-group>.card{margin-bottom:15px}@media(min-width: 576px){.card-group{display:flex;flex-flow:row wrap}.card-group>.card{flex:1 0 0%;margin-bottom:0}.card-group>.card+.card{margin-left:0;border-left:0}.card-group>.card:not(:last-child){border-top-right-radius:0;border-bottom-right-radius:0}.card-group>.card:not(:last-child) .card-img-top,.card-group>.card:not(:last-child) .card-header{border-top-right-radius:0}.card-group>.card:not(:last-child) .card-img-bottom,.card-group>.card:not(:last-child) .card-footer{border-bottom-right-radius:0}.card-group>.card:not(:first-child){border-top-left-radius:0;border-bottom-left-radius:0}.card-group>.card:not(:first-child) .card-img-top,.card-group>.card:not(:first-child) .card-header{border-top-left-radius:0}.card-group>.card:not(:first-child) .card-img-bottom,.card-group>.card:not(:first-child) .card-footer{border-bottom-left-radius:0}}.card-columns .card{margin-bottom:.75rem}@media(min-width: 576px){.card-columns{column-count:3;column-gap:1.25rem;orphans:1;widows:1}.card-columns .card{display:inline-block;width:100%}}.accordion{overflow-anchor:none}.accordion>.card{overflow:hidden}.accordion>.card:not(:last-of-type){border-bottom:0;border-bottom-right-radius:0;border-bottom-left-radius:0}.accordion>.card:not(:first-of-type){border-top-left-radius:0;border-top-right-radius:0}.accordion>.card>.card-header{border-radius:0;margin-bottom:-1px}.breadcrumb{display:flex;flex-wrap:wrap;padding:.75rem 1rem;margin-bottom:1rem;list-style:none;background-color:#e9ecef;border-radius:.25rem}.breadcrumb-item+.breadcrumb-item{padding-left:.5rem}.breadcrumb-item+.breadcrumb-item::before{float:left;padding-right:.5rem;color:#6c757d;content:"/"}.breadcrumb-item+.breadcrumb-item:hover::before{text-decoration:underline}.breadcrumb-item+.breadcrumb-item:hover::before{text-decoration:none}.breadcrumb-item.active{color:#6c757d}.pagination{display:flex;padding-left:0;list-style:none;border-radius:.25rem}.page-link{position:relative;display:block;padding:.5rem .75rem;margin-left:-1px;line-height:1.25;color:#00a99d;background-color:#fff;border:1px solid #dee2e6}.page-link:hover{z-index:2;color:#005d56;text-decoration:none;background-color:#e9ecef;border-color:#dee2e6}.page-link:focus{z-index:3;outline:0;box-shadow:0 0 0 .2rem rgba(0,169,157,.25)}.page-item:first-child .page-link{margin-left:0;border-top-left-radius:.25rem;border-bottom-left-radius:.25rem}.page-item:last-child .page-link{border-top-right-radius:.25rem;border-bottom-right-radius:.25rem}.page-item.active .page-link{z-index:3;color:#fff;background-color:#00a99d;border-color:#00a99d}.page-item.disabled .page-link{color:#6c757d;pointer-events:none;cursor:auto;background-color:#fff;border-color:#dee2e6}.pagination-lg .page-link{padding:.75rem 1.5rem;font-size:1.25rem;line-height:1.5}.pagination-lg .page-item:first-child .page-link{border-top-left-radius:.3rem;border-bottom-left-radius:.3rem}.pagination-lg .page-item:last-child .page-link{border-top-right-radius:.3rem;border-bottom-right-radius:.3rem}.pagination-sm .page-link{padding:.25rem .5rem;font-size:0.875rem;line-height:1.5}.pagination-sm .page-item:first-child .page-link{border-top-left-radius:.2rem;border-bottom-left-radius:.2rem}.pagination-sm .page-item:last-child .page-link{border-top-right-radius:.2rem;border-bottom-right-radius:.2rem}.badge{display:inline-block;padding:.25em .4em;font-size:75%;font-weight:700;line-height:1;text-align:center;white-space:nowrap;vertical-align:baseline;border-radius:.25rem;transition:color .15s ease-in-out,background-color .15s ease-in-out,border-color .15s ease-in-out,box-shadow .15s ease-in-out}@media(prefers-reduced-motion: reduce){.badge{transition:none}}a.badge:hover,a.badge:focus{text-decoration:none}.badge:empty{display:none}.btn .badge{position:relative;top:-1px}.badge-pill{padding-right:.6em;padding-left:.6em;border-radius:10rem}.badge-primary{color:#fff;background-color:#00a99d}a.badge-primary:hover,a.badge-primary:focus{color:#fff;background-color:#00766e}a.badge-primary:focus,a.badge-primary.focus{outline:0;box-shadow:0 0 0 .2rem rgba(0,169,157,.5)}.badge-secondary{color:#fff;background-color:#6c757d}a.badge-secondary:hover,a.badge-secondary:focus{color:#fff;background-color:#545b62}a.badge-secondary:focus,a.badge-secondary.focus{outline:0;box-shadow:0 0 0 .2rem rgba(108,117,125,.5)}.badge-success{color:#fff;background-color:#28a745}a.badge-success:hover,a.badge-success:focus{color:#fff;background-color:#1e7e34}a.badge-success:focus,a.badge-success.focus{outline:0;box-shadow:0 0 0 .2rem rgba(40,167,69,.5)}.badge-info{color:#fff;background-color:#17a2b8}a.badge-info:hover,a.badge-info:focus{color:#fff;background-color:#117a8b}a.badge-info:focus,a.badge-info.focus{outline:0;box-shadow:0 0 0 .2rem rgba(23,162,184,.5)}.badge-warning{color:#212529;background-color:#ffc107}a.badge-warning:hover,a.badge-warning:focus{color:#212529;background-color:#d39e00}a.badge-warning:focus,a.badge-warning.focus{outline:0;box-shadow:0 0 0 .2rem rgba(255,193,7,.5)}.badge-danger{color:#fff;background-color:#dc3545}a.badge-danger:hover,a.badge-danger:focus{color:#fff;background-color:#bd2130}a.badge-danger:focus,a.badge-danger.focus{outline:0;box-shadow:0 0 0 .2rem rgba(220,53,69,.5)}.badge-light{color:#212529;background-color:#f8f9fa}a.badge-light:hover,a.badge-light:focus{color:#212529;background-color:#dae0e5}a.badge-light:focus,a.badge-light.focus{outline:0;box-shadow:0 0 0 .2rem rgba(248,249,250,.5)}.badge-dark{color:#fff;background-color:#343a40}a.badge-dark:hover,a.badge-dark:focus{color:#fff;background-color:#1d2124}a.badge-dark:focus,a.badge-dark.focus{outline:0;box-shadow:0 0 0 .2rem rgba(52,58,64,.5)}.jumbotron{padding:2rem 1rem;margin-bottom:2rem;background-color:#e9ecef;border-radius:.3rem}@media(min-width: 576px){.jumbotron{padding:4rem 2rem}}.jumbotron-fluid{padding-right:0;padding-left:0;border-radius:0}.alert{position:relative;padding:.75rem 1.25rem;margin-bottom:1rem;border:1px solid transparent;border-radius:.25rem}.alert-heading{color:inherit}.alert-link{font-weight:700}.alert-dismissible{padding-right:4rem}.alert-dismissible .close{position:absolute;top:0;right:0;z-index:2;padding:.75rem 1.25rem;color:inherit}.alert-primary{color:#005852;background-color:#cceeeb;border-color:#b8e7e4}.alert-primary hr{border-top-color:#a5e1dd}.alert-primary .alert-link{color:#002522}.alert-secondary{color:#383d41;background-color:#e2e3e5;border-color:#d6d8db}.alert-secondary hr{border-top-color:#c8cbcf}.alert-secondary .alert-link{color:#202326}.alert-success{color:#155724;background-color:#d4edda;border-color:#c3e6cb}.alert-success hr{border-top-color:#b1dfbb}.alert-success .alert-link{color:#0b2e13}.alert-info{color:#0c5460;background-color:#d1ecf1;border-color:#bee5eb}.alert-info hr{border-top-color:#abdde5}.alert-info .alert-link{color:#062c33}.alert-warning{color:#856404;background-color:#fff3cd;border-color:#ffeeba}.alert-warning hr{border-top-color:#ffe8a1}.alert-warning .alert-link{color:#533f03}.alert-danger{color:#721c24;background-color:#f8d7da;border-color:#f5c6cb}.alert-danger hr{border-top-color:#f1b0b7}.alert-danger .alert-link{color:#491217}.alert-light{color:#818182;background-color:#fefefe;border-color:#fdfdfe}.alert-light hr{border-top-color:#ececf6}.alert-light .alert-link{color:#686868}.alert-dark{color:#1b1e21;background-color:#d6d8d9;border-color:#c6c8ca}.alert-dark hr{border-top-color:#b9bbbe}.alert-dark .alert-link{color:#040505}@keyframes progress-bar-stripes{from{background-position:1rem 0}to{background-position:0 0}}.progress{display:flex;height:1rem;overflow:hidden;line-height:0;font-size:0.75rem;background-color:#e9ecef;border-radius:.25rem}.progress-bar{display:flex;flex-direction:column;justify-content:center;overflow:hidden;color:#fff;text-align:center;white-space:nowrap;background-color:#00a99d;transition:width .6s ease}@media(prefers-reduced-motion: reduce){.progress-bar{transition:none}}.progress-bar-striped{background-image:linear-gradient(45deg, rgba(255, 255, 255, 0.15) 25%, transparent 25%, transparent 50%, rgba(255, 255, 255, 0.15) 50%, rgba(255, 255, 255, 0.15) 75%, transparent 75%, transparent);background-size:1rem 1rem}.progress-bar-animated{animation:1s linear infinite progress-bar-stripes}@media(prefers-reduced-motion: reduce){.progress-bar-animated{animation:none}}.media{display:flex;align-items:flex-start}.media-body{flex:1}.list-group{display:flex;flex-direction:column;padding-left:0;margin-bottom:0;border-radius:.25rem}.list-group-item-action{width:100%;color:#495057;text-align:inherit}.list-group-item-action:hover,.list-group-item-action:focus{z-index:1;color:#495057;text-decoration:none;background-color:#f8f9fa}.list-group-item-action:active{color:#212529;background-color:#e9ecef}.list-group-item{position:relative;display:block;padding:.75rem 1.25rem;background-color:#fff;border:1px solid rgba(0,0,0,.125)}.list-group-item:first-child{border-top-left-radius:inherit;border-top-right-radius:inherit}.list-group-item:last-child{border-bottom-right-radius:inherit;border-bottom-left-radius:inherit}.list-group-item.disabled,.list-group-item:disabled{color:#6c757d;pointer-events:none;background-color:#fff}.list-group-item.active{z-index:2;color:#fff;background-color:#00a99d;border-color:#00a99d}.list-group-item+.list-group-item{border-top-width:0}.list-group-item+.list-group-item.active{margin-top:-1px;border-top-width:1px}.list-group-horizontal{flex-direction:row}.list-group-horizontal>.list-group-item:first-child{border-bottom-left-radius:.25rem;border-top-right-radius:0}.list-group-horizontal>.list-group-item:last-child{border-top-right-radius:.25rem;border-bottom-left-radius:0}.list-group-horizontal>.list-group-item.active{margin-top:0}.list-group-horizontal>.list-group-item+.list-group-item{border-top-width:1px;border-left-width:0}.list-group-horizontal>.list-group-item+.list-group-item.active{margin-left:-1px;border-left-width:1px}@media(min-width: 576px){.list-group-horizontal-sm{flex-direction:row}.list-group-horizontal-sm>.list-group-item:first-child{border-bottom-left-radius:.25rem;border-top-right-radius:0}.list-group-horizontal-sm>.list-group-item:last-child{border-top-right-radius:.25rem;border-bottom-left-radius:0}.list-group-horizontal-sm>.list-group-item.active{margin-top:0}.list-group-horizontal-sm>.list-group-item+.list-group-item{border-top-width:1px;border-left-width:0}.list-group-horizontal-sm>.list-group-item+.list-group-item.active{margin-left:-1px;border-left-width:1px}}@media(min-width: 768px){.list-group-horizontal-md{flex-direction:row}.list-group-horizontal-md>.list-group-item:first-child{border-bottom-left-radius:.25rem;border-top-right-radius:0}.list-group-horizontal-md>.list-group-item:last-child{border-top-right-radius:.25rem;border-bottom-left-radius:0}.list-group-horizontal-md>.list-group-item.active{margin-top:0}.list-group-horizontal-md>.list-group-item+.list-group-item{border-top-width:1px;border-left-width:0}.list-group-horizontal-md>.list-group-item+.list-group-item.active{margin-left:-1px;border-left-width:1px}}@media(min-width: 992px){.list-group-horizontal-lg{flex-direction:row}.list-group-horizontal-lg>.list-group-item:first-child{border-bottom-left-radius:.25rem;border-top-right-radius:0}.list-group-horizontal-lg>.list-group-item:last-child{border-top-right-radius:.25rem;border-bottom-left-radius:0}.list-group-horizontal-lg>.list-group-item.active{margin-top:0}.list-group-horizontal-lg>.list-group-item+.list-group-item{border-top-width:1px;border-left-width:0}.list-group-horizontal-lg>.list-group-item+.list-group-item.active{margin-left:-1px;border-left-width:1px}}@media(min-width: 1200px){.list-group-horizontal-xl{flex-direction:row}.list-group-horizontal-xl>.list-group-item:first-child{border-bottom-left-radius:.25rem;border-top-right-radius:0}.list-group-horizontal-xl>.list-group-item:last-child{border-top-right-radius:.25rem;border-bottom-left-radius:0}.list-group-horizontal-xl>.list-group-item.active{margin-top:0}.list-group-horizontal-xl>.list-group-item+.list-group-item{border-top-width:1px;border-left-width:0}.list-group-horizontal-xl>.list-group-item+.list-group-item.active{margin-left:-1px;border-left-width:1px}}.list-group-flush{border-radius:0}.list-group-flush>.list-group-item{border-width:0 0 1px}.list-group-flush>.list-group-item:last-child{border-bottom-width:0}.list-group-item-primary{color:#005852;background-color:#b8e7e4}.list-group-item-primary.list-group-item-action:hover,.list-group-item-primary.list-group-item-action:focus{color:#005852;background-color:#a5e1dd}.list-group-item-primary.list-group-item-action.active{color:#fff;background-color:#005852;border-color:#005852}.list-group-item-secondary{color:#383d41;background-color:#d6d8db}.list-group-item-secondary.list-group-item-action:hover,.list-group-item-secondary.list-group-item-action:focus{color:#383d41;background-color:#c8cbcf}.list-group-item-secondary.list-group-item-action.active{color:#fff;background-color:#383d41;border-color:#383d41}.list-group-item-success{color:#155724;background-color:#c3e6cb}.list-group-item-success.list-group-item-action:hover,.list-group-item-success.list-group-item-action:focus{color:#155724;background-color:#b1dfbb}.list-group-item-success.list-group-item-action.active{color:#fff;background-color:#155724;border-color:#155724}.list-group-item-info{color:#0c5460;background-color:#bee5eb}.list-group-item-info.list-group-item-action:hover,.list-group-item-info.list-group-item-action:focus{color:#0c5460;background-color:#abdde5}.list-group-item-info.list-group-item-action.active{color:#fff;background-color:#0c5460;border-color:#0c5460}.list-group-item-warning{color:#856404;background-color:#ffeeba}.list-group-item-warning.list-group-item-action:hover,.list-group-item-warning.list-group-item-action:focus{color:#856404;background-color:#ffe8a1}.list-group-item-warning.list-group-item-action.active{color:#fff;background-color:#856404;border-color:#856404}.list-group-item-danger{color:#721c24;background-color:#f5c6cb}.list-group-item-danger.list-group-item-action:hover,.list-group-item-danger.list-group-item-action:focus{color:#721c24;background-color:#f1b0b7}.list-group-item-danger.list-group-item-action.active{color:#fff;background-color:#721c24;border-color:#721c24}.list-group-item-light{color:#818182;background-color:#fdfdfe}.list-group-item-light.list-group-item-action:hover,.list-group-item-light.list-group-item-action:focus{color:#818182;background-color:#ececf6}.list-group-item-light.list-group-item-action.active{color:#fff;background-color:#818182;border-color:#818182}.list-group-item-dark{color:#1b1e21;background-color:#c6c8ca}.list-group-item-dark.list-group-item-action:hover,.list-group-item-dark.list-group-item-action:focus{color:#1b1e21;background-color:#b9bbbe}.list-group-item-dark.list-group-item-action.active{color:#fff;background-color:#1b1e21;border-color:#1b1e21}.close{float:right;font-size:1.5rem;font-weight:700;line-height:1;color:#000;text-shadow:0 1px 0 #fff;opacity:.5}.close:hover{color:#000;text-decoration:none}.close:not(:disabled):not(.disabled):hover,.close:not(:disabled):not(.disabled):focus{opacity:.75}button.close{padding:0;background-color:transparent;border:0}a.close.disabled{pointer-events:none}.toast{flex-basis:350px;max-width:350px;font-size:0.875rem;background-color:rgba(255,255,255,.85);background-clip:padding-box;border:1px solid rgba(0,0,0,.1);box-shadow:0 .25rem .75rem rgba(0,0,0,.1);opacity:0;border-radius:.25rem}.toast:not(:last-child){margin-bottom:.75rem}.toast.showing{opacity:1}.toast.show{display:block;opacity:1}.toast.hide{display:none}.toast-header{display:flex;align-items:center;padding:.25rem .75rem;color:#6c757d;background-color:rgba(255,255,255,.85);background-clip:padding-box;border-bottom:1px solid rgba(0,0,0,.05);border-top-left-radius:calc(0.25rem - 1px);border-top-right-radius:calc(0.25rem - 1px)}.toast-body{padding:.75rem}.modal-open{overflow:hidden}.modal-open .modal{overflow-x:hidden;overflow-y:auto}.modal{position:fixed;top:0;left:0;z-index:1050;display:none;width:100%;height:100%;overflow:hidden;outline:0}.modal-dialog{position:relative;width:auto;margin:.5rem;pointer-events:none}.modal.fade .modal-dialog{transition:transform .3s ease-out;transform:translate(0, -50px)}@media(prefers-reduced-motion: reduce){.modal.fade .modal-dialog{transition:none}}.modal.show .modal-dialog{transform:none}.modal.modal-static .modal-dialog{transform:scale(1.02)}.modal-dialog-scrollable{display:flex;max-height:calc(100% - 1rem)}.modal-dialog-scrollable .modal-content{max-height:calc(100vh - 1rem);overflow:hidden}.modal-dialog-scrollable .modal-header,.modal-dialog-scrollable .modal-footer{flex-shrink:0}.modal-dialog-scrollable .modal-body{overflow-y:auto}.modal-dialog-centered{display:flex;align-items:center;min-height:calc(100% - 1rem)}.modal-dialog-centered::before{display:block;height:calc(100vh - 1rem);height:min-content;content:""}.modal-dialog-centered.modal-dialog-scrollable{flex-direction:column;justify-content:center;height:100%}.modal-dialog-centered.modal-dialog-scrollable .modal-content{max-height:none}.modal-dialog-centered.modal-dialog-scrollable::before{content:none}.modal-content{position:relative;display:flex;flex-direction:column;width:100%;pointer-events:auto;background-color:#fff;background-clip:padding-box;border:1px solid rgba(0,0,0,.2);border-radius:.3rem;outline:0}.modal-backdrop{position:fixed;top:0;left:0;z-index:1040;width:100vw;height:100vh;background-color:#000}.modal-backdrop.fade{opacity:0}.modal-backdrop.show{opacity:.5}.modal-header{display:flex;align-items:flex-start;justify-content:space-between;padding:1rem 1rem;border-bottom:1px solid #dee2e6;border-top-left-radius:calc(0.3rem - 1px);border-top-right-radius:calc(0.3rem - 1px)}.modal-header .close{padding:1rem 1rem;margin:-1rem -1rem -1rem auto}.modal-title{margin-bottom:0;line-height:1.5}.modal-body{position:relative;flex:1 1 auto;padding:1rem}.modal-footer{display:flex;flex-wrap:wrap;align-items:center;justify-content:flex-end;padding:.75rem;border-top:1px solid #dee2e6;border-bottom-right-radius:calc(0.3rem - 1px);border-bottom-left-radius:calc(0.3rem - 1px)}.modal-footer>*{margin:.25rem}.modal-scrollbar-measure{position:absolute;top:-9999px;width:50px;height:50px;overflow:scroll}@media(min-width: 576px){.modal-dialog{max-width:500px;margin:1.75rem auto}.modal-dialog-scrollable{max-height:calc(100% - 3.5rem)}.modal-dialog-scrollable .modal-content{max-height:calc(100vh - 3.5rem)}.modal-dialog-centered{min-height:calc(100% - 3.5rem)}.modal-dialog-centered::before{height:calc(100vh - 3.5rem);height:min-content}.modal-sm{max-width:300px}}@media(min-width: 992px){.modal-lg,.modal-xl{max-width:800px}}@media(min-width: 1200px){.modal-xl{max-width:1140px}}.tooltip{position:absolute;z-index:1070;display:block;margin:0;font-family:-apple-system,BlinkMacSystemFont,"Segoe UI",Roboto,"Helvetica Neue",Arial,"Noto Sans","Liberation Sans",sans-serif,"Apple Color Emoji","Segoe UI Emoji","Segoe UI Symbol","Noto Color Emoji";font-style:normal;font-weight:400;line-height:1.5;text-align:left;text-align:start;text-decoration:none;text-shadow:none;text-transform:none;letter-spacing:normal;word-break:normal;word-spacing:normal;white-space:normal;line-break:auto;font-size:0.875rem;word-wrap:break-word;opacity:0}.tooltip.show{opacity:.9}.tooltip .arrow{position:absolute;display:block;width:.8rem;height:.4rem}.tooltip .arrow::before{position:absolute;content:"";border-color:transparent;border-style:solid}.bs-tooltip-top,.bs-tooltip-auto[x-placement^=top]{padding:.4rem 0}.bs-tooltip-top .arrow,.bs-tooltip-auto[x-placement^=top] .arrow{bottom:0}.bs-tooltip-top .arrow::before,.bs-tooltip-auto[x-placement^=top] .arrow::before{top:0;border-width:.4rem .4rem 0;border-top-color:#000}.bs-tooltip-right,.bs-tooltip-auto[x-placement^=right]{padding:0 .4rem}.bs-tooltip-right .arrow,.bs-tooltip-auto[x-placement^=right] .arrow{left:0;width:.4rem;height:.8rem}.bs-tooltip-right .arrow::before,.bs-tooltip-auto[x-placement^=right] .arrow::before{right:0;border-width:.4rem .4rem .4rem 0;border-right-color:#000}.bs-tooltip-bottom,.bs-tooltip-auto[x-placement^=bottom]{padding:.4rem 0}.bs-tooltip-bottom .arrow,.bs-tooltip-auto[x-placement^=bottom] .arrow{top:0}.bs-tooltip-bottom .arrow::before,.bs-tooltip-auto[x-placement^=bottom] .arrow::before{bottom:0;border-width:0 .4rem .4rem;border-bottom-color:#000}.bs-tooltip-left,.bs-tooltip-auto[x-placement^=left]{padding:0 .4rem}.bs-tooltip-left .arrow,.bs-tooltip-auto[x-placement^=left] .arrow{right:0;width:.4rem;height:.8rem}.bs-tooltip-left .arrow::before,.bs-tooltip-auto[x-placement^=left] .arrow::before{left:0;border-width:.4rem 0 .4rem .4rem;border-left-color:#000}.tooltip-inner{max-width:200px;padding:.25rem .5rem;color:#fff;text-align:center;background-color:#000;border-radius:.25rem}.popover{position:absolute;top:0;left:0;z-index:1060;display:block;max-width:276px;font-family:-apple-system,BlinkMacSystemFont,"Segoe UI",Roboto,"Helvetica Neue",Arial,"Noto Sans","Liberation Sans",sans-serif,"Apple Color Emoji","Segoe UI Emoji","Segoe UI Symbol","Noto Color Emoji";font-style:normal;font-weight:400;line-height:1.5;text-align:left;text-align:start;text-decoration:none;text-shadow:none;text-transform:none;letter-spacing:normal;word-break:normal;word-spacing:normal;white-space:normal;line-break:auto;font-size:0.875rem;word-wrap:break-word;background-color:#fff;background-clip:padding-box;border:1px solid rgba(0,0,0,.2);border-radius:.3rem}.popover .arrow{position:absolute;display:block;width:1rem;height:.5rem;margin:0 .3rem}.popover .arrow::before,.popover .arrow::after{position:absolute;display:block;content:"";border-color:transparent;border-style:solid}.bs-popover-top,.bs-popover-auto[x-placement^=top]{margin-bottom:.5rem}.bs-popover-top>.arrow,.bs-popover-auto[x-placement^=top]>.arrow{bottom:calc(-0.5rem - 1px)}.bs-popover-top>.arrow::before,.bs-popover-auto[x-placement^=top]>.arrow::before{bottom:0;border-width:.5rem .5rem 0;border-top-color:rgba(0,0,0,.25)}.bs-popover-top>.arrow::after,.bs-popover-auto[x-placement^=top]>.arrow::after{bottom:1px;border-width:.5rem .5rem 0;border-top-color:#fff}.bs-popover-right,.bs-popover-auto[x-placement^=right]{margin-left:.5rem}.bs-popover-right>.arrow,.bs-popover-auto[x-placement^=right]>.arrow{left:calc(-0.5rem - 1px);width:.5rem;height:1rem;margin:.3rem 0}.bs-popover-right>.arrow::before,.bs-popover-auto[x-placement^=right]>.arrow::before{left:0;border-width:.5rem .5rem .5rem 0;border-right-color:rgba(0,0,0,.25)}.bs-popover-right>.arrow::after,.bs-popover-auto[x-placement^=right]>.arrow::after{left:1px;border-width:.5rem .5rem .5rem 0;border-right-color:#fff}.bs-popover-bottom,.bs-popover-auto[x-placement^=bottom]{margin-top:.5rem}.bs-popover-bottom>.arrow,.bs-popover-auto[x-placement^=bottom]>.arrow{top:calc(-0.5rem - 1px)}.bs-popover-bottom>.arrow::before,.bs-popover-auto[x-placement^=bottom]>.arrow::before{top:0;border-width:0 .5rem .5rem .5rem;border-bottom-color:rgba(0,0,0,.25)}.bs-popover-bottom>.arrow::after,.bs-popover-auto[x-placement^=bottom]>.arrow::after{top:1px;border-width:0 .5rem .5rem .5rem;border-bottom-color:#fff}.bs-popover-bottom .popover-header::before,.bs-popover-auto[x-placement^=bottom] .popover-header::before{position:absolute;top:0;left:50%;display:block;width:1rem;margin-left:-0.5rem;content:"";border-bottom:1px solid #f7f7f7}.bs-popover-left,.bs-popover-auto[x-placement^=left]{margin-right:.5rem}.bs-popover-left>.arrow,.bs-popover-auto[x-placement^=left]>.arrow{right:calc(-0.5rem - 1px);width:.5rem;height:1rem;margin:.3rem 0}.bs-popover-left>.arrow::before,.bs-popover-auto[x-placement^=left]>.arrow::before{right:0;border-width:.5rem 0 .5rem .5rem;border-left-color:rgba(0,0,0,.25)}.bs-popover-left>.arrow::after,.bs-popover-auto[x-placement^=left]>.arrow::after{right:1px;border-width:.5rem 0 .5rem .5rem;border-left-color:#fff}.popover-header{padding:.5rem .75rem;margin-bottom:0;font-size:1rem;background-color:#f7f7f7;border-bottom:1px solid #ebebeb;border-top-left-radius:calc(0.3rem - 1px);border-top-right-radius:calc(0.3rem - 1px)}.popover-header:empty{display:none}.popover-body{padding:.5rem .75rem;color:#212529}.carousel{position:relative}.carousel.pointer-event{touch-action:pan-y}.carousel-inner{position:relative;width:100%;overflow:hidden}.carousel-inner::after{display:block;clear:both;content:""}.carousel-item{position:relative;display:none;float:left;width:100%;margin-right:-100%;backface-visibility:hidden;transition:transform .6s ease-in-out}@media(prefers-reduced-motion: reduce){.carousel-item{transition:none}}.carousel-item.active,.carousel-item-next,.carousel-item-prev{display:block}.carousel-item-next:not(.carousel-item-left),.active.carousel-item-right{transform:translateX(100%)}.carousel-item-prev:not(.carousel-item-right),.active.carousel-item-left{transform:translateX(-100%)}.carousel-fade .carousel-item{opacity:0;transition-property:opacity;transform:none}.carousel-fade .carousel-item.active,.carousel-fade .carousel-item-next.carousel-item-left,.carousel-fade .carousel-item-prev.carousel-item-right{z-index:1;opacity:1}.carousel-fade .active.carousel-item-left,.carousel-fade .active.carousel-item-right{z-index:0;opacity:0;transition:opacity 0s .6s}@media(prefers-reduced-motion: reduce){.carousel-fade .active.carousel-item-left,.carousel-fade .active.carousel-item-right{transition:none}}.carousel-control-prev,.carousel-control-next{position:absolute;top:0;bottom:0;z-index:1;display:flex;align-items:center;justify-content:center;width:15%;color:#fff;text-align:center;opacity:.5;transition:opacity .15s ease}@media(prefers-reduced-motion: reduce){.carousel-control-prev,.carousel-control-next{transition:none}}.carousel-control-prev:hover,.carousel-control-prev:focus,.carousel-control-next:hover,.carousel-control-next:focus{color:#fff;text-decoration:none;outline:0;opacity:.9}.carousel-control-prev{left:0}.carousel-control-next{right:0}.carousel-control-prev-icon,.carousel-control-next-icon{display:inline-block;width:20px;height:20px;background:50%/100% 100% no-repeat}.carousel-control-prev-icon{background-image:url("data:image/svg+xml,%3csvg xmlns=\'http://www.w3.org/2000/svg\' fill=\'%23fff\' width=\'8\' height=\'8\' viewBox=\'0 0 8 8\'%3e%3cpath d=\'M5.25 0l-4 4 4 4 1.5-1.5L4.25 4l2.5-2.5L5.25 0z\'/%3e%3c/svg%3e")}.carousel-control-next-icon{background-image:url("data:image/svg+xml,%3csvg xmlns=\'http://www.w3.org/2000/svg\' fill=\'%23fff\' width=\'8\' height=\'8\' viewBox=\'0 0 8 8\'%3e%3cpath d=\'M2.75 0l-1.5 1.5L3.75 4l-2.5 2.5L2.75 8l4-4-4-4z\'/%3e%3c/svg%3e")}.carousel-indicators{position:absolute;right:0;bottom:0;left:0;z-index:15;display:flex;justify-content:center;padding-left:0;margin-right:15%;margin-left:15%;list-style:none}.carousel-indicators li{box-sizing:content-box;flex:0 1 auto;width:30px;height:3px;margin-right:3px;margin-left:3px;text-indent:-999px;cursor:pointer;background-color:#fff;background-clip:padding-box;border-top:10px solid transparent;border-bottom:10px solid transparent;opacity:.5;transition:opacity .6s ease}@media(prefers-reduced-motion: reduce){.carousel-indicators li{transition:none}}.carousel-indicators .active{opacity:1}.carousel-caption{position:absolute;right:15%;bottom:20px;left:15%;z-index:10;padding-top:20px;padding-bottom:20px;color:#fff;text-align:center}@keyframes spinner-border{to{transform:rotate(360deg)}}.spinner-border{display:inline-block;width:2rem;height:2rem;vertical-align:text-bottom;border:.25em solid currentColor;border-right-color:transparent;border-radius:50%;animation:.75s linear infinite spinner-border}.spinner-border-sm{width:1rem;height:1rem;border-width:.2em}@keyframes spinner-grow{0%{transform:scale(0)}50%{opacity:1;transform:none}}.spinner-grow{display:inline-block;width:2rem;height:2rem;vertical-align:text-bottom;background-color:currentColor;border-radius:50%;opacity:0;animation:.75s linear infinite spinner-grow}.spinner-grow-sm{width:1rem;height:1rem}@media(prefers-reduced-motion: reduce){.spinner-border,.spinner-grow{animation-duration:1.5s}}.align-baseline{vertical-align:baseline !important}.align-top{vertical-align:top !important}.align-middle{vertical-align:middle !important}.align-bottom{vertical-align:bottom !important}.align-text-bottom{vertical-align:text-bottom !important}.align-text-top{vertical-align:text-top !important}.bg-primary{background-color:#00a99d !important}a.bg-primary:hover,a.bg-primary:focus,button.bg-primary:hover,button.bg-primary:focus{background-color:#00766e !important}.bg-secondary{background-color:#6c757d !important}a.bg-secondary:hover,a.bg-secondary:focus,button.bg-secondary:hover,button.bg-secondary:focus{background-color:#545b62 !important}.bg-success{background-color:#28a745 !important}a.bg-success:hover,a.bg-success:focus,button.bg-success:hover,button.bg-success:focus{background-color:#1e7e34 !important}.bg-info{background-color:#17a2b8 !important}a.bg-info:hover,a.bg-info:focus,button.bg-info:hover,button.bg-info:focus{background-color:#117a8b !important}.bg-warning{background-color:#ffc107 !important}a.bg-warning:hover,a.bg-warning:focus,button.bg-warning:hover,button.bg-warning:focus{background-color:#d39e00 !important}.bg-danger{background-color:#dc3545 !important}a.bg-danger:hover,a.bg-danger:focus,button.bg-danger:hover,button.bg-danger:focus{background-color:#bd2130 !important}.bg-light{background-color:#f8f9fa !important}a.bg-light:hover,a.bg-light:focus,button.bg-light:hover,button.bg-light:focus{background-color:#dae0e5 !important}.bg-dark{background-color:#343a40 !important}a.bg-dark:hover,a.bg-dark:focus,button.bg-dark:hover,button.bg-dark:focus{background-color:#1d2124 !important}.bg-white{background-color:#fff !important}.bg-transparent{background-color:transparent !important}.border{border:1px solid #dee2e6 !important}.border-top{border-top:1px solid #dee2e6 !important}.border-right{border-right:1px solid #dee2e6 !important}.border-bottom{border-bottom:1px solid #dee2e6 !important}.border-left{border-left:1px solid #dee2e6 !important}.border-0{border:0 !important}.border-top-0{border-top:0 !important}.border-right-0{border-right:0 !important}.border-bottom-0{border-bottom:0 !important}.border-left-0{border-left:0 !important}.border-primary{border-color:#00a99d !important}.border-secondary{border-color:#6c757d !important}.border-success{border-color:#28a745 !important}.border-info{border-color:#17a2b8 !important}.border-warning{border-color:#ffc107 !important}.border-danger{border-color:#dc3545 !important}.border-light{border-color:#f8f9fa !important}.border-dark{border-color:#343a40 !important}.border-white{border-color:#fff !important}.rounded-sm{border-radius:.2rem !important}.rounded{border-radius:.25rem !important}.rounded-top{border-top-left-radius:.25rem !important;border-top-right-radius:.25rem !important}.rounded-right{border-top-right-radius:.25rem !important;border-bottom-right-radius:.25rem !important}.rounded-bottom{border-bottom-right-radius:.25rem !important;border-bottom-left-radius:.25rem !important}.rounded-left{border-top-left-radius:.25rem !important;border-bottom-left-radius:.25rem !important}.rounded-lg{border-radius:.3rem !important}.rounded-circle{border-radius:50% !important}.rounded-pill{border-radius:50rem !important}.rounded-0{border-radius:0 !important}.clearfix::after{display:block;clear:both;content:""}.d-none{display:none !important}.d-inline{display:inline !important}.d-inline-block{display:inline-block !important}.d-block{display:block !important}.d-table{display:table !important}.d-table-row{display:table-row !important}.d-table-cell{display:table-cell !important}.d-flex{display:flex !important}.d-inline-flex{display:inline-flex !important}@media(min-width: 576px){.d-sm-none{display:none !important}.d-sm-inline{display:inline !important}.d-sm-inline-block{display:inline-block !important}.d-sm-block{display:block !important}.d-sm-table{display:table !important}.d-sm-table-row{display:table-row !important}.d-sm-table-cell{display:table-cell !important}.d-sm-flex{display:flex !important}.d-sm-inline-flex{display:inline-flex !important}}@media(min-width: 768px){.d-md-none{display:none !important}.d-md-inline{display:inline !important}.d-md-inline-block{display:inline-block !important}.d-md-block{display:block !important}.d-md-table{display:table !important}.d-md-table-row{display:table-row !important}.d-md-table-cell{display:table-cell !important}.d-md-flex{display:flex !important}.d-md-inline-flex{display:inline-flex !important}}@media(min-width: 992px){.d-lg-none{display:none !important}.d-lg-inline{display:inline !important}.d-lg-inline-block{display:inline-block !important}.d-lg-block{display:block !important}.d-lg-table{display:table !important}.d-lg-table-row{display:table-row !important}.d-lg-table-cell{display:table-cell !important}.d-lg-flex{display:flex !important}.d-lg-inline-flex{display:inline-flex !important}}@media(min-width: 1200px){.d-xl-none{display:none !important}.d-xl-inline{display:inline !important}.d-xl-inline-block{display:inline-block !important}.d-xl-block{display:block !important}.d-xl-table{display:table !important}.d-xl-table-row{display:table-row !important}.d-xl-table-cell{display:table-cell !important}.d-xl-flex{display:flex !important}.d-xl-inline-flex{display:inline-flex !important}}@media print{.d-print-none{display:none !important}.d-print-inline{display:inline !important}.d-print-inline-block{display:inline-block !important}.d-print-block{display:block !important}.d-print-table{display:table !important}.d-print-table-row{display:table-row !important}.d-print-table-cell{display:table-cell !important}.d-print-flex{display:flex !important}.d-print-inline-flex{display:inline-flex !important}}.embed-responsive{position:relative;display:block;width:100%;padding:0;overflow:hidden}.embed-responsive::before{display:block;content:""}.embed-responsive .embed-responsive-item,.embed-responsive iframe,.embed-responsive embed,.embed-responsive object,.embed-responsive video{position:absolute;top:0;bottom:0;left:0;width:100%;height:100%;border:0}.embed-responsive-21by9::before{padding-top:42.8571428571%}.embed-responsive-16by9::before{padding-top:56.25%}.embed-responsive-4by3::before{padding-top:75%}.embed-responsive-1by1::before{padding-top:100%}.flex-row{flex-direction:row !important}.flex-column{flex-direction:column !important}.flex-row-reverse{flex-direction:row-reverse !important}.flex-column-reverse{flex-direction:column-reverse !important}.flex-wrap{flex-wrap:wrap !important}.flex-nowrap{flex-wrap:nowrap !important}.flex-wrap-reverse{flex-wrap:wrap-reverse !important}.flex-fill{flex:1 1 auto !important}.flex-grow-0{flex-grow:0 !important}.flex-grow-1{flex-grow:1 !important}.flex-shrink-0{flex-shrink:0 !important}.flex-shrink-1{flex-shrink:1 !important}.justify-content-start{justify-content:flex-start !important}.justify-content-end{justify-content:flex-end !important}.justify-content-center{justify-content:center !important}.justify-content-between{justify-content:space-between !important}.justify-content-around{justify-content:space-around !important}.align-items-start{align-items:flex-start !important}.align-items-end{align-items:flex-end !important}.align-items-center{align-items:center !important}.align-items-baseline{align-items:baseline !important}.align-items-stretch{align-items:stretch !important}.align-content-start{align-content:flex-start !important}.align-content-end{align-content:flex-end !important}.align-content-center{align-content:center !important}.align-content-between{align-content:space-between !important}.align-content-around{align-content:space-around !important}.align-content-stretch{align-content:stretch !important}.align-self-auto{align-self:auto !important}.align-self-start{align-self:flex-start !important}.align-self-end{align-self:flex-end !important}.align-self-center{align-self:center !important}.align-self-baseline{align-self:baseline !important}.align-self-stretch{align-self:stretch !important}@media(min-width: 576px){.flex-sm-row{flex-direction:row !important}.flex-sm-column{flex-direction:column !important}.flex-sm-row-reverse{flex-direction:row-reverse !important}.flex-sm-column-reverse{flex-direction:column-reverse !important}.flex-sm-wrap{flex-wrap:wrap !important}.flex-sm-nowrap{flex-wrap:nowrap !important}.flex-sm-wrap-reverse{flex-wrap:wrap-reverse !important}.flex-sm-fill{flex:1 1 auto !important}.flex-sm-grow-0{flex-grow:0 !important}.flex-sm-grow-1{flex-grow:1 !important}.flex-sm-shrink-0{flex-shrink:0 !important}.flex-sm-shrink-1{flex-shrink:1 !important}.justify-content-sm-start{justify-content:flex-start !important}.justify-content-sm-end{justify-content:flex-end !important}.justify-content-sm-center{justify-content:center !important}.justify-content-sm-between{justify-content:space-between !important}.justify-content-sm-around{justify-content:space-around !important}.align-items-sm-start{align-items:flex-start !important}.align-items-sm-end{align-items:flex-end !important}.align-items-sm-center{align-items:center !important}.align-items-sm-baseline{align-items:baseline !important}.align-items-sm-stretch{align-items:stretch !important}.align-content-sm-start{align-content:flex-start !important}.align-content-sm-end{align-content:flex-end !important}.align-content-sm-center{align-content:center !important}.align-content-sm-between{align-content:space-between !important}.align-content-sm-around{align-content:space-around !important}.align-content-sm-stretch{align-content:stretch !important}.align-self-sm-auto{align-self:auto !important}.align-self-sm-start{align-self:flex-start !important}.align-self-sm-end{align-self:flex-end !important}.align-self-sm-center{align-self:center !important}.align-self-sm-baseline{align-self:baseline !important}.align-self-sm-stretch{align-self:stretch !important}}@media(min-width: 768px){.flex-md-row{flex-direction:row !important}.flex-md-column{flex-direction:column !important}.flex-md-row-reverse{flex-direction:row-reverse !important}.flex-md-column-reverse{flex-direction:column-reverse !important}.flex-md-wrap{flex-wrap:wrap !important}.flex-md-nowrap{flex-wrap:nowrap !important}.flex-md-wrap-reverse{flex-wrap:wrap-reverse !important}.flex-md-fill{flex:1 1 auto !important}.flex-md-grow-0{flex-grow:0 !important}.flex-md-grow-1{flex-grow:1 !important}.flex-md-shrink-0{flex-shrink:0 !important}.flex-md-shrink-1{flex-shrink:1 !important}.justify-content-md-start{justify-content:flex-start !important}.justify-content-md-end{justify-content:flex-end !important}.justify-content-md-center{justify-content:center !important}.justify-content-md-between{justify-content:space-between !important}.justify-content-md-around{justify-content:space-around !important}.align-items-md-start{align-items:flex-start !important}.align-items-md-end{align-items:flex-end !important}.align-items-md-center{align-items:center !important}.align-items-md-baseline{align-items:baseline !important}.align-items-md-stretch{align-items:stretch !important}.align-content-md-start{align-content:flex-start !important}.align-content-md-end{align-content:flex-end !important}.align-content-md-center{align-content:center !important}.align-content-md-between{align-content:space-between !important}.align-content-md-around{align-content:space-around !important}.align-content-md-stretch{align-content:stretch !important}.align-self-md-auto{align-self:auto !important}.align-self-md-start{align-self:flex-start !important}.align-self-md-end{align-self:flex-end !important}.align-self-md-center{align-self:center !important}.align-self-md-baseline{align-self:baseline !important}.align-self-md-stretch{align-self:stretch !important}}@media(min-width: 992px){.flex-lg-row{flex-direction:row !important}.flex-lg-column{flex-direction:column !important}.flex-lg-row-reverse{flex-direction:row-reverse !important}.flex-lg-column-reverse{flex-direction:column-reverse !important}.flex-lg-wrap{flex-wrap:wrap !important}.flex-lg-nowrap{flex-wrap:nowrap !important}.flex-lg-wrap-reverse{flex-wrap:wrap-reverse !important}.flex-lg-fill{flex:1 1 auto !important}.flex-lg-grow-0{flex-grow:0 !important}.flex-lg-grow-1{flex-grow:1 !important}.flex-lg-shrink-0{flex-shrink:0 !important}.flex-lg-shrink-1{flex-shrink:1 !important}.justify-content-lg-start{justify-content:flex-start !important}.justify-content-lg-end{justify-content:flex-end !important}.justify-content-lg-center{justify-content:center !important}.justify-content-lg-between{justify-content:space-between !important}.justify-content-lg-around{justify-content:space-around !important}.align-items-lg-start{align-items:flex-start !important}.align-items-lg-end{align-items:flex-end !important}.align-items-lg-center{align-items:center !important}.align-items-lg-baseline{align-items:baseline !important}.align-items-lg-stretch{align-items:stretch !important}.align-content-lg-start{align-content:flex-start !important}.align-content-lg-end{align-content:flex-end !important}.align-content-lg-center{align-content:center !important}.align-content-lg-between{align-content:space-between !important}.align-content-lg-around{align-content:space-around !important}.align-content-lg-stretch{align-content:stretch !important}.align-self-lg-auto{align-self:auto !important}.align-self-lg-start{align-self:flex-start !important}.align-self-lg-end{align-self:flex-end !important}.align-self-lg-center{align-self:center !important}.align-self-lg-baseline{align-self:baseline !important}.align-self-lg-stretch{align-self:stretch !important}}@media(min-width: 1200px){.flex-xl-row{flex-direction:row !important}.flex-xl-column{flex-direction:column !important}.flex-xl-row-reverse{flex-direction:row-reverse !important}.flex-xl-column-reverse{flex-direction:column-reverse !important}.flex-xl-wrap{flex-wrap:wrap !important}.flex-xl-nowrap{flex-wrap:nowrap !important}.flex-xl-wrap-reverse{flex-wrap:wrap-reverse !important}.flex-xl-fill{flex:1 1 auto !important}.flex-xl-grow-0{flex-grow:0 !important}.flex-xl-grow-1{flex-grow:1 !important}.flex-xl-shrink-0{flex-shrink:0 !important}.flex-xl-shrink-1{flex-shrink:1 !important}.justify-content-xl-start{justify-content:flex-start !important}.justify-content-xl-end{justify-content:flex-end !important}.justify-content-xl-center{justify-content:center !important}.justify-content-xl-between{justify-content:space-between !important}.justify-content-xl-around{justify-content:space-around !important}.align-items-xl-start{align-items:flex-start !important}.align-items-xl-end{align-items:flex-end !important}.align-items-xl-center{align-items:center !important}.align-items-xl-baseline{align-items:baseline !important}.align-items-xl-stretch{align-items:stretch !important}.align-content-xl-start{align-content:flex-start !important}.align-content-xl-end{align-content:flex-end !important}.align-content-xl-center{align-content:center !important}.align-content-xl-between{align-content:space-between !important}.align-content-xl-around{align-content:space-around !important}.align-content-xl-stretch{align-content:stretch !important}.align-self-xl-auto{align-self:auto !important}.align-self-xl-start{align-self:flex-start !important}.align-self-xl-end{align-self:flex-end !important}.align-self-xl-center{align-self:center !important}.align-self-xl-baseline{align-self:baseline !important}.align-self-xl-stretch{align-self:stretch !important}}.float-left{float:left !important}.float-right{float:right !important}.float-none{float:none !important}@media(min-width: 576px){.float-sm-left{float:left !important}.float-sm-right{float:right !important}.float-sm-none{float:none !important}}@media(min-width: 768px){.float-md-left{float:left !important}.float-md-right{float:right !important}.float-md-none{float:none !important}}@media(min-width: 992px){.float-lg-left{float:left !important}.float-lg-right{float:right !important}.float-lg-none{float:none !important}}@media(min-width: 1200px){.float-xl-left{float:left !important}.float-xl-right{float:right !important}.float-xl-none{float:none !important}}.user-select-all{user-select:all !important}.user-select-auto{user-select:auto !important}.user-select-none{user-select:none !important}.overflow-auto{overflow:auto !important}.overflow-hidden{overflow:hidden !important}.position-static{position:static !important}.position-relative{position:relative !important}.position-absolute{position:absolute !important}.position-fixed{position:fixed !important}.position-sticky{position:sticky !important}.fixed-top{position:fixed;top:0;right:0;left:0;z-index:1030}.fixed-bottom{position:fixed;right:0;bottom:0;left:0;z-index:1030}@supports(position: sticky){.sticky-top{position:sticky;top:0;z-index:1020}}.sr-only{position:absolute;width:1px;height:1px;padding:0;margin:-1px;overflow:hidden;clip:rect(0, 0, 0, 0);white-space:nowrap;border:0}.sr-only-focusable:active,.sr-only-focusable:focus{position:static;width:auto;height:auto;overflow:visible;clip:auto;white-space:normal}.shadow-sm{box-shadow:0 .125rem .25rem rgba(0,0,0,.075) !important}.shadow{box-shadow:0 .5rem 1rem rgba(0,0,0,.15) !important}.shadow-lg{box-shadow:0 1rem 3rem rgba(0,0,0,.175) !important}.shadow-none{box-shadow:none !important}.w-25{width:25% !important}.w-50{width:50% !important}.w-75{width:75% !important}.w-100{width:100% !important}.w-auto{width:auto !important}.h-25{height:25% !important}.h-50{height:50% !important}.h-75{height:75% !important}.h-100{height:100% !important}.h-auto{height:auto !important}.mw-100{max-width:100% !important}.mh-100{max-height:100% !important}.min-vw-100{min-width:100vw !important}.min-vh-100{min-height:100vh !important}.vw-100{width:100vw !important}.vh-100{height:100vh !important}.m-0{margin:0 !important}.mt-0,.my-0{margin-top:0 !important}.mr-0,.mx-0{margin-right:0 !important}.mb-0,.my-0{margin-bottom:0 !important}.ml-0,.mx-0{margin-left:0 !important}.m-1{margin:.25rem !important}.mt-1,.my-1{margin-top:.25rem !important}.mr-1,.mx-1{margin-right:.25rem !important}.mb-1,.my-1{margin-bottom:.25rem !important}.ml-1,.mx-1{margin-left:.25rem !important}.m-2{margin:.5rem !important}.mt-2,.my-2{margin-top:.5rem !important}.mr-2,.mx-2{margin-right:.5rem !important}.mb-2,.my-2{margin-bottom:.5rem !important}.ml-2,.mx-2{margin-left:.5rem !important}.m-3{margin:1rem !important}.mt-3,.my-3{margin-top:1rem !important}.mr-3,.mx-3{margin-right:1rem !important}.mb-3,.my-3{margin-bottom:1rem !important}.ml-3,.mx-3{margin-left:1rem !important}.m-4{margin:1.5rem !important}.mt-4,.my-4{margin-top:1.5rem !important}.mr-4,.mx-4{margin-right:1.5rem !important}.mb-4,.my-4{margin-bottom:1.5rem !important}.ml-4,.mx-4{margin-left:1.5rem !important}.m-5{margin:3rem !important}.mt-5,.my-5{margin-top:3rem !important}.mr-5,.mx-5{margin-right:3rem !important}.mb-5,.my-5{margin-bottom:3rem !important}.ml-5,.mx-5{margin-left:3rem !important}.p-0{padding:0 !important}.pt-0,.py-0{padding-top:0 !important}.pr-0,.px-0{padding-right:0 !important}.pb-0,.py-0{padding-bottom:0 !important}.pl-0,.px-0{padding-left:0 !important}.p-1{padding:.25rem !important}.pt-1,.py-1{padding-top:.25rem !important}.pr-1,.px-1{padding-right:.25rem !important}.pb-1,.py-1{padding-bottom:.25rem !important}.pl-1,.px-1{padding-left:.25rem !important}.p-2{padding:.5rem !important}.pt-2,.py-2{padding-top:.5rem !important}.pr-2,.px-2{padding-right:.5rem !important}.pb-2,.py-2{padding-bottom:.5rem !important}.pl-2,.px-2{padding-left:.5rem !important}.p-3{padding:1rem !important}.pt-3,.py-3{padding-top:1rem !important}.pr-3,.px-3{padding-right:1rem !important}.pb-3,.py-3{padding-bottom:1rem !important}.pl-3,.px-3{padding-left:1rem !important}.p-4{padding:1.5rem !important}.pt-4,.py-4{padding-top:1.5rem !important}.pr-4,.px-4{padding-right:1.5rem !important}.pb-4,.py-4{padding-bottom:1.5rem !important}.pl-4,.px-4{padding-left:1.5rem !important}.p-5{padding:3rem !important}.pt-5,.py-5{padding-top:3rem !important}.pr-5,.px-5{padding-right:3rem !important}.pb-5,.py-5{padding-bottom:3rem !important}.pl-5,.px-5{padding-left:3rem !important}.m-n1{margin:-0.25rem !important}.mt-n1,.my-n1{margin-top:-0.25rem !important}.mr-n1,.mx-n1{margin-right:-0.25rem !important}.mb-n1,.my-n1{margin-bottom:-0.25rem !important}.ml-n1,.mx-n1{margin-left:-0.25rem !important}.m-n2{margin:-0.5rem !important}.mt-n2,.my-n2{margin-top:-0.5rem !important}.mr-n2,.mx-n2{margin-right:-0.5rem !important}.mb-n2,.my-n2{margin-bottom:-0.5rem !important}.ml-n2,.mx-n2{margin-left:-0.5rem !important}.m-n3{margin:-1rem !important}.mt-n3,.my-n3{margin-top:-1rem !important}.mr-n3,.mx-n3{margin-right:-1rem !important}.mb-n3,.my-n3{margin-bottom:-1rem !important}.ml-n3,.mx-n3{margin-left:-1rem !important}.m-n4{margin:-1.5rem !important}.mt-n4,.my-n4{margin-top:-1.5rem !important}.mr-n4,.mx-n4{margin-right:-1.5rem !important}.mb-n4,.my-n4{margin-bottom:-1.5rem !important}.ml-n4,.mx-n4{margin-left:-1.5rem !important}.m-n5{margin:-3rem !important}.mt-n5,.my-n5{margin-top:-3rem !important}.mr-n5,.mx-n5{margin-right:-3rem !important}.mb-n5,.my-n5{margin-bottom:-3rem !important}.ml-n5,.mx-n5{margin-left:-3rem !important}.m-auto{margin:auto !important}.mt-auto,.my-auto{margin-top:auto !important}.mr-auto,.mx-auto{margin-right:auto !important}.mb-auto,.my-auto{margin-bottom:auto !important}.ml-auto,.mx-auto{margin-left:auto !important}@media(min-width: 576px){.m-sm-0{margin:0 !important}.mt-sm-0,.my-sm-0{margin-top:0 !important}.mr-sm-0,.mx-sm-0{margin-right:0 !important}.mb-sm-0,.my-sm-0{margin-bottom:0 !important}.ml-sm-0,.mx-sm-0{margin-left:0 !important}.m-sm-1{margin:.25rem !important}.mt-sm-1,.my-sm-1{margin-top:.25rem !important}.mr-sm-1,.mx-sm-1{margin-right:.25rem !important}.mb-sm-1,.my-sm-1{margin-bottom:.25rem !important}.ml-sm-1,.mx-sm-1{margin-left:.25rem !important}.m-sm-2{margin:.5rem !important}.mt-sm-2,.my-sm-2{margin-top:.5rem !important}.mr-sm-2,.mx-sm-2{margin-right:.5rem !important}.mb-sm-2,.my-sm-2{margin-bottom:.5rem !important}.ml-sm-2,.mx-sm-2{margin-left:.5rem !important}.m-sm-3{margin:1rem !important}.mt-sm-3,.my-sm-3{margin-top:1rem !important}.mr-sm-3,.mx-sm-3{margin-right:1rem !important}.mb-sm-3,.my-sm-3{margin-bottom:1rem !important}.ml-sm-3,.mx-sm-3{margin-left:1rem !important}.m-sm-4{margin:1.5rem !important}.mt-sm-4,.my-sm-4{margin-top:1.5rem !important}.mr-sm-4,.mx-sm-4{margin-right:1.5rem !important}.mb-sm-4,.my-sm-4{margin-bottom:1.5rem !important}.ml-sm-4,.mx-sm-4{margin-left:1.5rem !important}.m-sm-5{margin:3rem !important}.mt-sm-5,.my-sm-5{margin-top:3rem !important}.mr-sm-5,.mx-sm-5{margin-right:3rem !important}.mb-sm-5,.my-sm-5{margin-bottom:3rem !important}.ml-sm-5,.mx-sm-5{margin-left:3rem !important}.p-sm-0{padding:0 !important}.pt-sm-0,.py-sm-0{padding-top:0 !important}.pr-sm-0,.px-sm-0{padding-right:0 !important}.pb-sm-0,.py-sm-0{padding-bottom:0 !important}.pl-sm-0,.px-sm-0{padding-left:0 !important}.p-sm-1{padding:.25rem !important}.pt-sm-1,.py-sm-1{padding-top:.25rem !important}.pr-sm-1,.px-sm-1{padding-right:.25rem !important}.pb-sm-1,.py-sm-1{padding-bottom:.25rem !important}.pl-sm-1,.px-sm-1{padding-left:.25rem !important}.p-sm-2{padding:.5rem !important}.pt-sm-2,.py-sm-2{padding-top:.5rem !important}.pr-sm-2,.px-sm-2{padding-right:.5rem !important}.pb-sm-2,.py-sm-2{padding-bottom:.5rem !important}.pl-sm-2,.px-sm-2{padding-left:.5rem !important}.p-sm-3{padding:1rem !important}.pt-sm-3,.py-sm-3{padding-top:1rem !important}.pr-sm-3,.px-sm-3{padding-right:1rem !important}.pb-sm-3,.py-sm-3{padding-bottom:1rem !important}.pl-sm-3,.px-sm-3{padding-left:1rem !important}.p-sm-4{padding:1.5rem !important}.pt-sm-4,.py-sm-4{padding-top:1.5rem !important}.pr-sm-4,.px-sm-4{padding-right:1.5rem !important}.pb-sm-4,.py-sm-4{padding-bottom:1.5rem !important}.pl-sm-4,.px-sm-4{padding-left:1.5rem !important}.p-sm-5{padding:3rem !important}.pt-sm-5,.py-sm-5{padding-top:3rem !important}.pr-sm-5,.px-sm-5{padding-right:3rem !important}.pb-sm-5,.py-sm-5{padding-bottom:3rem !important}.pl-sm-5,.px-sm-5{padding-left:3rem !important}.m-sm-n1{margin:-0.25rem !important}.mt-sm-n1,.my-sm-n1{margin-top:-0.25rem !important}.mr-sm-n1,.mx-sm-n1{margin-right:-0.25rem !important}.mb-sm-n1,.my-sm-n1{margin-bottom:-0.25rem !important}.ml-sm-n1,.mx-sm-n1{margin-left:-0.25rem !important}.m-sm-n2{margin:-0.5rem !important}.mt-sm-n2,.my-sm-n2{margin-top:-0.5rem !important}.mr-sm-n2,.mx-sm-n2{margin-right:-0.5rem !important}.mb-sm-n2,.my-sm-n2{margin-bottom:-0.5rem !important}.ml-sm-n2,.mx-sm-n2{margin-left:-0.5rem !important}.m-sm-n3{margin:-1rem !important}.mt-sm-n3,.my-sm-n3{margin-top:-1rem !important}.mr-sm-n3,.mx-sm-n3{margin-right:-1rem !important}.mb-sm-n3,.my-sm-n3{margin-bottom:-1rem !important}.ml-sm-n3,.mx-sm-n3{margin-left:-1rem !important}.m-sm-n4{margin:-1.5rem !important}.mt-sm-n4,.my-sm-n4{margin-top:-1.5rem !important}.mr-sm-n4,.mx-sm-n4{margin-right:-1.5rem !important}.mb-sm-n4,.my-sm-n4{margin-bottom:-1.5rem !important}.ml-sm-n4,.mx-sm-n4{margin-left:-1.5rem !important}.m-sm-n5{margin:-3rem !important}.mt-sm-n5,.my-sm-n5{margin-top:-3rem !important}.mr-sm-n5,.mx-sm-n5{margin-right:-3rem !important}.mb-sm-n5,.my-sm-n5{margin-bottom:-3rem !important}.ml-sm-n5,.mx-sm-n5{margin-left:-3rem !important}.m-sm-auto{margin:auto !important}.mt-sm-auto,.my-sm-auto{margin-top:auto !important}.mr-sm-auto,.mx-sm-auto{margin-right:auto !important}.mb-sm-auto,.my-sm-auto{margin-bottom:auto !important}.ml-sm-auto,.mx-sm-auto{margin-left:auto !important}}@media(min-width: 768px){.m-md-0{margin:0 !important}.mt-md-0,.my-md-0{margin-top:0 !important}.mr-md-0,.mx-md-0{margin-right:0 !important}.mb-md-0,.my-md-0{margin-bottom:0 !important}.ml-md-0,.mx-md-0{margin-left:0 !important}.m-md-1{margin:.25rem !important}.mt-md-1,.my-md-1{margin-top:.25rem !important}.mr-md-1,.mx-md-1{margin-right:.25rem !important}.mb-md-1,.my-md-1{margin-bottom:.25rem !important}.ml-md-1,.mx-md-1{margin-left:.25rem !important}.m-md-2{margin:.5rem !important}.mt-md-2,.my-md-2{margin-top:.5rem !important}.mr-md-2,.mx-md-2{margin-right:.5rem !important}.mb-md-2,.my-md-2{margin-bottom:.5rem !important}.ml-md-2,.mx-md-2{margin-left:.5rem !important}.m-md-3{margin:1rem !important}.mt-md-3,.my-md-3{margin-top:1rem !important}.mr-md-3,.mx-md-3{margin-right:1rem !important}.mb-md-3,.my-md-3{margin-bottom:1rem !important}.ml-md-3,.mx-md-3{margin-left:1rem !important}.m-md-4{margin:1.5rem !important}.mt-md-4,.my-md-4{margin-top:1.5rem !important}.mr-md-4,.mx-md-4{margin-right:1.5rem !important}.mb-md-4,.my-md-4{margin-bottom:1.5rem !important}.ml-md-4,.mx-md-4{margin-left:1.5rem !important}.m-md-5{margin:3rem !important}.mt-md-5,.my-md-5{margin-top:3rem !important}.mr-md-5,.mx-md-5{margin-right:3rem !important}.mb-md-5,.my-md-5{margin-bottom:3rem !important}.ml-md-5,.mx-md-5{margin-left:3rem !important}.p-md-0{padding:0 !important}.pt-md-0,.py-md-0{padding-top:0 !important}.pr-md-0,.px-md-0{padding-right:0 !important}.pb-md-0,.py-md-0{padding-bottom:0 !important}.pl-md-0,.px-md-0{padding-left:0 !important}.p-md-1{padding:.25rem !important}.pt-md-1,.py-md-1{padding-top:.25rem !important}.pr-md-1,.px-md-1{padding-right:.25rem !important}.pb-md-1,.py-md-1{padding-bottom:.25rem !important}.pl-md-1,.px-md-1{padding-left:.25rem !important}.p-md-2{padding:.5rem !important}.pt-md-2,.py-md-2{padding-top:.5rem !important}.pr-md-2,.px-md-2{padding-right:.5rem !important}.pb-md-2,.py-md-2{padding-bottom:.5rem !important}.pl-md-2,.px-md-2{padding-left:.5rem !important}.p-md-3{padding:1rem !important}.pt-md-3,.py-md-3{padding-top:1rem !important}.pr-md-3,.px-md-3{padding-right:1rem !important}.pb-md-3,.py-md-3{padding-bottom:1rem !important}.pl-md-3,.px-md-3{padding-left:1rem !important}.p-md-4{padding:1.5rem !important}.pt-md-4,.py-md-4{padding-top:1.5rem !important}.pr-md-4,.px-md-4{padding-right:1.5rem !important}.pb-md-4,.py-md-4{padding-bottom:1.5rem !important}.pl-md-4,.px-md-4{padding-left:1.5rem !important}.p-md-5{padding:3rem !important}.pt-md-5,.py-md-5{padding-top:3rem !important}.pr-md-5,.px-md-5{padding-right:3rem !important}.pb-md-5,.py-md-5{padding-bottom:3rem !important}.pl-md-5,.px-md-5{padding-left:3rem !important}.m-md-n1{margin:-0.25rem !important}.mt-md-n1,.my-md-n1{margin-top:-0.25rem !important}.mr-md-n1,.mx-md-n1{margin-right:-0.25rem !important}.mb-md-n1,.my-md-n1{margin-bottom:-0.25rem !important}.ml-md-n1,.mx-md-n1{margin-left:-0.25rem !important}.m-md-n2{margin:-0.5rem !important}.mt-md-n2,.my-md-n2{margin-top:-0.5rem !important}.mr-md-n2,.mx-md-n2{margin-right:-0.5rem !important}.mb-md-n2,.my-md-n2{margin-bottom:-0.5rem !important}.ml-md-n2,.mx-md-n2{margin-left:-0.5rem !important}.m-md-n3{margin:-1rem !important}.mt-md-n3,.my-md-n3{margin-top:-1rem !important}.mr-md-n3,.mx-md-n3{margin-right:-1rem !important}.mb-md-n3,.my-md-n3{margin-bottom:-1rem !important}.ml-md-n3,.mx-md-n3{margin-left:-1rem !important}.m-md-n4{margin:-1.5rem !important}.mt-md-n4,.my-md-n4{margin-top:-1.5rem !important}.mr-md-n4,.mx-md-n4{margin-right:-1.5rem !important}.mb-md-n4,.my-md-n4{margin-bottom:-1.5rem !important}.ml-md-n4,.mx-md-n4{margin-left:-1.5rem !important}.m-md-n5{margin:-3rem !important}.mt-md-n5,.my-md-n5{margin-top:-3rem !important}.mr-md-n5,.mx-md-n5{margin-right:-3rem !important}.mb-md-n5,.my-md-n5{margin-bottom:-3rem !important}.ml-md-n5,.mx-md-n5{margin-left:-3rem !important}.m-md-auto{margin:auto !important}.mt-md-auto,.my-md-auto{margin-top:auto !important}.mr-md-auto,.mx-md-auto{margin-right:auto !important}.mb-md-auto,.my-md-auto{margin-bottom:auto !important}.ml-md-auto,.mx-md-auto{margin-left:auto !important}}@media(min-width: 992px){.m-lg-0{margin:0 !important}.mt-lg-0,.my-lg-0{margin-top:0 !important}.mr-lg-0,.mx-lg-0{margin-right:0 !important}.mb-lg-0,.my-lg-0{margin-bottom:0 !important}.ml-lg-0,.mx-lg-0{margin-left:0 !important}.m-lg-1{margin:.25rem !important}.mt-lg-1,.my-lg-1{margin-top:.25rem !important}.mr-lg-1,.mx-lg-1{margin-right:.25rem !important}.mb-lg-1,.my-lg-1{margin-bottom:.25rem !important}.ml-lg-1,.mx-lg-1{margin-left:.25rem !important}.m-lg-2{margin:.5rem !important}.mt-lg-2,.my-lg-2{margin-top:.5rem !important}.mr-lg-2,.mx-lg-2{margin-right:.5rem !important}.mb-lg-2,.my-lg-2{margin-bottom:.5rem !important}.ml-lg-2,.mx-lg-2{margin-left:.5rem !important}.m-lg-3{margin:1rem !important}.mt-lg-3,.my-lg-3{margin-top:1rem !important}.mr-lg-3,.mx-lg-3{margin-right:1rem !important}.mb-lg-3,.my-lg-3{margin-bottom:1rem !important}.ml-lg-3,.mx-lg-3{margin-left:1rem !important}.m-lg-4{margin:1.5rem !important}.mt-lg-4,.my-lg-4{margin-top:1.5rem !important}.mr-lg-4,.mx-lg-4{margin-right:1.5rem !important}.mb-lg-4,.my-lg-4{margin-bottom:1.5rem !important}.ml-lg-4,.mx-lg-4{margin-left:1.5rem !important}.m-lg-5{margin:3rem !important}.mt-lg-5,.my-lg-5{margin-top:3rem !important}.mr-lg-5,.mx-lg-5{margin-right:3rem !important}.mb-lg-5,.my-lg-5{margin-bottom:3rem !important}.ml-lg-5,.mx-lg-5{margin-left:3rem !important}.p-lg-0{padding:0 !important}.pt-lg-0,.py-lg-0{padding-top:0 !important}.pr-lg-0,.px-lg-0{padding-right:0 !important}.pb-lg-0,.py-lg-0{padding-bottom:0 !important}.pl-lg-0,.px-lg-0{padding-left:0 !important}.p-lg-1{padding:.25rem !important}.pt-lg-1,.py-lg-1{padding-top:.25rem !important}.pr-lg-1,.px-lg-1{padding-right:.25rem !important}.pb-lg-1,.py-lg-1{padding-bottom:.25rem !important}.pl-lg-1,.px-lg-1{padding-left:.25rem !important}.p-lg-2{padding:.5rem !important}.pt-lg-2,.py-lg-2{padding-top:.5rem !important}.pr-lg-2,.px-lg-2{padding-right:.5rem !important}.pb-lg-2,.py-lg-2{padding-bottom:.5rem !important}.pl-lg-2,.px-lg-2{padding-left:.5rem !important}.p-lg-3{padding:1rem !important}.pt-lg-3,.py-lg-3{padding-top:1rem !important}.pr-lg-3,.px-lg-3{padding-right:1rem !important}.pb-lg-3,.py-lg-3{padding-bottom:1rem !important}.pl-lg-3,.px-lg-3{padding-left:1rem !important}.p-lg-4{padding:1.5rem !important}.pt-lg-4,.py-lg-4{padding-top:1.5rem !important}.pr-lg-4,.px-lg-4{padding-right:1.5rem !important}.pb-lg-4,.py-lg-4{padding-bottom:1.5rem !important}.pl-lg-4,.px-lg-4{padding-left:1.5rem !important}.p-lg-5{padding:3rem !important}.pt-lg-5,.py-lg-5{padding-top:3rem !important}.pr-lg-5,.px-lg-5{padding-right:3rem !important}.pb-lg-5,.py-lg-5{padding-bottom:3rem !important}.pl-lg-5,.px-lg-5{padding-left:3rem !important}.m-lg-n1{margin:-0.25rem !important}.mt-lg-n1,.my-lg-n1{margin-top:-0.25rem !important}.mr-lg-n1,.mx-lg-n1{margin-right:-0.25rem !important}.mb-lg-n1,.my-lg-n1{margin-bottom:-0.25rem !important}.ml-lg-n1,.mx-lg-n1{margin-left:-0.25rem !important}.m-lg-n2{margin:-0.5rem !important}.mt-lg-n2,.my-lg-n2{margin-top:-0.5rem !important}.mr-lg-n2,.mx-lg-n2{margin-right:-0.5rem !important}.mb-lg-n2,.my-lg-n2{margin-bottom:-0.5rem !important}.ml-lg-n2,.mx-lg-n2{margin-left:-0.5rem !important}.m-lg-n3{margin:-1rem !important}.mt-lg-n3,.my-lg-n3{margin-top:-1rem !important}.mr-lg-n3,.mx-lg-n3{margin-right:-1rem !important}.mb-lg-n3,.my-lg-n3{margin-bottom:-1rem !important}.ml-lg-n3,.mx-lg-n3{margin-left:-1rem !important}.m-lg-n4{margin:-1.5rem !important}.mt-lg-n4,.my-lg-n4{margin-top:-1.5rem !important}.mr-lg-n4,.mx-lg-n4{margin-right:-1.5rem !important}.mb-lg-n4,.my-lg-n4{margin-bottom:-1.5rem !important}.ml-lg-n4,.mx-lg-n4{margin-left:-1.5rem !important}.m-lg-n5{margin:-3rem !important}.mt-lg-n5,.my-lg-n5{margin-top:-3rem !important}.mr-lg-n5,.mx-lg-n5{margin-right:-3rem !important}.mb-lg-n5,.my-lg-n5{margin-bottom:-3rem !important}.ml-lg-n5,.mx-lg-n5{margin-left:-3rem !important}.m-lg-auto{margin:auto !important}.mt-lg-auto,.my-lg-auto{margin-top:auto !important}.mr-lg-auto,.mx-lg-auto{margin-right:auto !important}.mb-lg-auto,.my-lg-auto{margin-bottom:auto !important}.ml-lg-auto,.mx-lg-auto{margin-left:auto !important}}@media(min-width: 1200px){.m-xl-0{margin:0 !important}.mt-xl-0,.my-xl-0{margin-top:0 !important}.mr-xl-0,.mx-xl-0{margin-right:0 !important}.mb-xl-0,.my-xl-0{margin-bottom:0 !important}.ml-xl-0,.mx-xl-0{margin-left:0 !important}.m-xl-1{margin:.25rem !important}.mt-xl-1,.my-xl-1{margin-top:.25rem !important}.mr-xl-1,.mx-xl-1{margin-right:.25rem !important}.mb-xl-1,.my-xl-1{margin-bottom:.25rem !important}.ml-xl-1,.mx-xl-1{margin-left:.25rem !important}.m-xl-2{margin:.5rem !important}.mt-xl-2,.my-xl-2{margin-top:.5rem !important}.mr-xl-2,.mx-xl-2{margin-right:.5rem !important}.mb-xl-2,.my-xl-2{margin-bottom:.5rem !important}.ml-xl-2,.mx-xl-2{margin-left:.5rem !important}.m-xl-3{margin:1rem !important}.mt-xl-3,.my-xl-3{margin-top:1rem !important}.mr-xl-3,.mx-xl-3{margin-right:1rem !important}.mb-xl-3,.my-xl-3{margin-bottom:1rem !important}.ml-xl-3,.mx-xl-3{margin-left:1rem !important}.m-xl-4{margin:1.5rem !important}.mt-xl-4,.my-xl-4{margin-top:1.5rem !important}.mr-xl-4,.mx-xl-4{margin-right:1.5rem !important}.mb-xl-4,.my-xl-4{margin-bottom:1.5rem !important}.ml-xl-4,.mx-xl-4{margin-left:1.5rem !important}.m-xl-5{margin:3rem !important}.mt-xl-5,.my-xl-5{margin-top:3rem !important}.mr-xl-5,.mx-xl-5{margin-right:3rem !important}.mb-xl-5,.my-xl-5{margin-bottom:3rem !important}.ml-xl-5,.mx-xl-5{margin-left:3rem !important}.p-xl-0{padding:0 !important}.pt-xl-0,.py-xl-0{padding-top:0 !important}.pr-xl-0,.px-xl-0{padding-right:0 !important}.pb-xl-0,.py-xl-0{padding-bottom:0 !important}.pl-xl-0,.px-xl-0{padding-left:0 !important}.p-xl-1{padding:.25rem !important}.pt-xl-1,.py-xl-1{padding-top:.25rem !important}.pr-xl-1,.px-xl-1{padding-right:.25rem !important}.pb-xl-1,.py-xl-1{padding-bottom:.25rem !important}.pl-xl-1,.px-xl-1{padding-left:.25rem !important}.p-xl-2{padding:.5rem !important}.pt-xl-2,.py-xl-2{padding-top:.5rem !important}.pr-xl-2,.px-xl-2{padding-right:.5rem !important}.pb-xl-2,.py-xl-2{padding-bottom:.5rem !important}.pl-xl-2,.px-xl-2{padding-left:.5rem !important}.p-xl-3{padding:1rem !important}.pt-xl-3,.py-xl-3{padding-top:1rem !important}.pr-xl-3,.px-xl-3{padding-right:1rem !important}.pb-xl-3,.py-xl-3{padding-bottom:1rem !important}.pl-xl-3,.px-xl-3{padding-left:1rem !important}.p-xl-4{padding:1.5rem !important}.pt-xl-4,.py-xl-4{padding-top:1.5rem !important}.pr-xl-4,.px-xl-4{padding-right:1.5rem !important}.pb-xl-4,.py-xl-4{padding-bottom:1.5rem !important}.pl-xl-4,.px-xl-4{padding-left:1.5rem !important}.p-xl-5{padding:3rem !important}.pt-xl-5,.py-xl-5{padding-top:3rem !important}.pr-xl-5,.px-xl-5{padding-right:3rem !important}.pb-xl-5,.py-xl-5{padding-bottom:3rem !important}.pl-xl-5,.px-xl-5{padding-left:3rem !important}.m-xl-n1{margin:-0.25rem !important}.mt-xl-n1,.my-xl-n1{margin-top:-0.25rem !important}.mr-xl-n1,.mx-xl-n1{margin-right:-0.25rem !important}.mb-xl-n1,.my-xl-n1{margin-bottom:-0.25rem !important}.ml-xl-n1,.mx-xl-n1{margin-left:-0.25rem !important}.m-xl-n2{margin:-0.5rem !important}.mt-xl-n2,.my-xl-n2{margin-top:-0.5rem !important}.mr-xl-n2,.mx-xl-n2{margin-right:-0.5rem !important}.mb-xl-n2,.my-xl-n2{margin-bottom:-0.5rem !important}.ml-xl-n2,.mx-xl-n2{margin-left:-0.5rem !important}.m-xl-n3{margin:-1rem !important}.mt-xl-n3,.my-xl-n3{margin-top:-1rem !important}.mr-xl-n3,.mx-xl-n3{margin-right:-1rem !important}.mb-xl-n3,.my-xl-n3{margin-bottom:-1rem !important}.ml-xl-n3,.mx-xl-n3{margin-left:-1rem !important}.m-xl-n4{margin:-1.5rem !important}.mt-xl-n4,.my-xl-n4{margin-top:-1.5rem !important}.mr-xl-n4,.mx-xl-n4{margin-right:-1.5rem !important}.mb-xl-n4,.my-xl-n4{margin-bottom:-1.5rem !important}.ml-xl-n4,.mx-xl-n4{margin-left:-1.5rem !important}.m-xl-n5{margin:-3rem !important}.mt-xl-n5,.my-xl-n5{margin-top:-3rem !important}.mr-xl-n5,.mx-xl-n5{margin-right:-3rem !important}.mb-xl-n5,.my-xl-n5{margin-bottom:-3rem !important}.ml-xl-n5,.mx-xl-n5{margin-left:-3rem !important}.m-xl-auto{margin:auto !important}.mt-xl-auto,.my-xl-auto{margin-top:auto !important}.mr-xl-auto,.mx-xl-auto{margin-right:auto !important}.mb-xl-auto,.my-xl-auto{margin-bottom:auto !important}.ml-xl-auto,.mx-xl-auto{margin-left:auto !important}}.stretched-link::after{position:absolute;top:0;right:0;bottom:0;left:0;z-index:1;pointer-events:auto;content:"";background-color:rgba(0,0,0,0)}.text-monospace{font-family:SFMono-Regular,Menlo,Monaco,Consolas,"Liberation Mono","Courier New",monospace !important}.text-justify{text-align:justify !important}.text-wrap{white-space:normal !important}.text-nowrap{white-space:nowrap !important}.text-truncate{overflow:hidden;text-overflow:ellipsis;white-space:nowrap}.text-left{text-align:left !important}.text-right{text-align:right !important}.text-center{text-align:center !important}@media(min-width: 576px){.text-sm-left{text-align:left !important}.text-sm-right{text-align:right !important}.text-sm-center{text-align:center !important}}@media(min-width: 768px){.text-md-left{text-align:left !important}.text-md-right{text-align:right !important}.text-md-center{text-align:center !important}}@media(min-width: 992px){.text-lg-left{text-align:left !important}.text-lg-right{text-align:right !important}.text-lg-center{text-align:center !important}}@media(min-width: 1200px){.text-xl-left{text-align:left !important}.text-xl-right{text-align:right !important}.text-xl-center{text-align:center !important}}.text-lowercase{text-transform:lowercase !important}.text-uppercase{text-transform:uppercase !important}.text-capitalize{text-transform:capitalize !important}.font-weight-light{font-weight:300 !important}.font-weight-lighter{font-weight:lighter !important}.font-weight-normal{font-weight:400 !important}.font-weight-bold{font-weight:700 !important}.font-weight-bolder{font-weight:bolder !important}.font-italic{font-style:italic !important}.text-white{color:#fff !important}.text-primary{color:#00a99d !important}a.text-primary:hover,a.text-primary:focus{color:#005d56 !important}.text-secondary{color:#6c757d !important}a.text-secondary:hover,a.text-secondary:focus{color:#494f54 !important}.text-success{color:#28a745 !important}a.text-success:hover,a.text-success:focus{color:#19692c !important}.text-info{color:#17a2b8 !important}a.text-info:hover,a.text-info:focus{color:#0f6674 !important}.text-warning{color:#ffc107 !important}a.text-warning:hover,a.text-warning:focus{color:#ba8b00 !important}.text-danger{color:#dc3545 !important}a.text-danger:hover,a.text-danger:focus{color:#a71d2a !important}.text-light{color:#f8f9fa !important}a.text-light:hover,a.text-light:focus{color:#cbd3da !important}.text-dark{color:#343a40 !important}a.text-dark:hover,a.text-dark:focus{color:#121416 !important}.text-body{color:#212529 !important}.text-muted{color:#6c757d !important}.text-black-50{color:rgba(0,0,0,.5) !important}.text-white-50{color:rgba(255,255,255,.5) !important}.text-hide{font:0/0 a;color:transparent;text-shadow:none;background-color:transparent;border:0}.text-decoration-none{text-decoration:none !important}.text-break{word-break:break-word !important;word-wrap:break-word !important}.text-reset{color:inherit !important}.visible{visibility:visible !important}.invisible{visibility:hidden !important}@media print{*,*::before,*::after{text-shadow:none !important;box-shadow:none !important}a:not(.btn){text-decoration:underline}abbr[title]::after{content:" (" attr(title) ")"}pre{white-space:pre-wrap !important}pre,blockquote{border:1px solid #adb5bd;page-break-inside:avoid}thead{display:table-header-group}tr,img{page-break-inside:avoid}p,h2,h3{orphans:3;widows:3}h2,h3{page-break-after:avoid}@page{size:a3}body{min-width:992px !important}.container{min-width:992px !important}.navbar{display:none}.badge{border:1px solid #000}.table{border-collapse:collapse !important}.table td,.table th{background-color:#fff !important}.table-bordered th,.table-bordered td{border:1px solid #dee2e6 !important}.table-dark{color:inherit}.table-dark th,.table-dark td,.table-dark thead th,.table-dark tbody+tbody{border-color:#dee2e6}.table .thead-dark th{color:inherit;border-color:#dee2e6}}/*!\n * Font Awesome Free 5.15.2 by @fontawesome - https://fontawesome.com\n * License - https://fontawesome.com/license/free (Icons: CC BY 4.0, Fonts: SIL OFL 1.1, Code: MIT License)\n */.fa,.fas,.far,.fal,.fad,.fab{-moz-osx-font-smoothing:grayscale;-webkit-font-smoothing:antialiased;display:inline-block;font-style:normal;font-variant:normal;text-rendering:auto;line-height:1}.fa-lg{font-size:1.3333333333em;line-height:.75em;vertical-align:-0.0667em}.fa-xs{font-size:.75em}.fa-sm{font-size:.875em}.fa-1x{font-size:1em}.fa-2x{font-size:2em}.fa-3x{font-size:3em}.fa-4x{font-size:4em}.fa-5x{font-size:5em}.fa-6x{font-size:6em}.fa-7x{font-size:7em}.fa-8x{font-size:8em}.fa-9x{font-size:9em}.fa-10x{font-size:10em}.fa-fw{text-align:center;width:1.25em}.fa-ul{list-style-type:none;margin-left:2.5em;padding-left:0}.fa-ul>li{position:relative}.fa-li{left:-2em;position:absolute;text-align:center;width:2em;line-height:inherit}.fa-border{border:solid .08em #eee;border-radius:.1em;padding:.2em .25em .15em}.fa-pull-left{float:left}.fa-pull-right{float:right}.fa.fa-pull-left,.fas.fa-pull-left,.far.fa-pull-left,.fal.fa-pull-left,.fab.fa-pull-left{margin-right:.3em}.fa.fa-pull-right,.fas.fa-pull-right,.far.fa-pull-right,.fal.fa-pull-right,.fab.fa-pull-right{margin-left:.3em}.fa-spin{animation:fa-spin 2s infinite linear}.fa-pulse{animation:fa-spin 1s infinite steps(8)}@keyframes fa-spin{0%{transform:rotate(0deg)}100%{transform:rotate(360deg)}}.fa-rotate-90{-ms-filter:"progid:DXImageTransform.Microsoft.BasicImage(rotation=1)";transform:rotate(90deg)}.fa-rotate-180{-ms-filter:"progid:DXImageTransform.Microsoft.BasicImage(rotation=2)";transform:rotate(180deg)}.fa-rotate-270{-ms-filter:"progid:DXImageTransform.Microsoft.BasicImage(rotation=3)";transform:rotate(270deg)}.fa-flip-horizontal{-ms-filter:"progid:DXImageTransform.Microsoft.BasicImage(rotation=0, mirror=1)";transform:scale(-1, 1)}.fa-flip-vertical{-ms-filter:"progid:DXImageTransform.Microsoft.BasicImage(rotation=2, mirror=1)";transform:scale(1, -1)}.fa-flip-both,.fa-flip-horizontal.fa-flip-vertical{-ms-filter:"progid:DXImageTransform.Microsoft.BasicImage(rotation=2, mirror=1)";transform:scale(-1, -1)}:root .fa-rotate-90,:root .fa-rotate-180,:root .fa-rotate-270,:root .fa-flip-horizontal,:root .fa-flip-vertical,:root .fa-flip-both{filter:none}.fa-stack{display:inline-block;height:2em;line-height:2em;position:relative;vertical-align:middle;width:2.5em}.fa-stack-1x,.fa-stack-2x{left:0;position:absolute;text-align:center;width:100%}.fa-stack-1x{line-height:inherit}.fa-stack-2x{font-size:2em}.fa-inverse{color:#fff}.fa-500px:before{content:""}.fa-accessible-icon:before{content:""}.fa-accusoft:before{content:""}.fa-acquisitions-incorporated:before{content:""}.fa-ad:before{content:""}.fa-address-book:before{content:""}.fa-address-card:before{content:""}.fa-adjust:before{content:""}.fa-adn:before{content:""}.fa-adversal:before{content:""}.fa-affiliatetheme:before{content:""}.fa-air-freshener:before{content:""}.fa-airbnb:before{content:""}.fa-algolia:before{content:""}.fa-align-center:before{content:""}.fa-align-justify:before{content:""}.fa-align-left:before{content:""}.fa-align-right:before{content:""}.fa-alipay:before{content:""}.fa-allergies:before{content:""}.fa-amazon:before{content:""}.fa-amazon-pay:before{content:""}.fa-ambulance:before{content:""}.fa-american-sign-language-interpreting:before{content:""}.fa-amilia:before{content:""}.fa-anchor:before{content:""}.fa-android:before{content:""}.fa-angellist:before{content:""}.fa-angle-double-down:before{content:""}.fa-angle-double-left:before{content:""}.fa-angle-double-right:before{content:""}.fa-angle-double-up:before{content:""}.fa-angle-down:before{content:""}.fa-angle-left:before{content:""}.fa-angle-right:before{content:""}.fa-angle-up:before{content:""}.fa-angry:before{content:""}.fa-angrycreative:before{content:""}.fa-angular:before{content:""}.fa-ankh:before{content:""}.fa-app-store:before{content:""}.fa-app-store-ios:before{content:""}.fa-apper:before{content:""}.fa-apple:before{content:""}.fa-apple-alt:before{content:""}.fa-apple-pay:before{content:""}.fa-archive:before{content:""}.fa-archway:before{content:""}.fa-arrow-alt-circle-down:before{content:""}.fa-arrow-alt-circle-left:before{content:""}.fa-arrow-alt-circle-right:before{content:""}.fa-arrow-alt-circle-up:before{content:""}.fa-arrow-circle-down:before{content:""}.fa-arrow-circle-left:before{content:""}.fa-arrow-circle-right:before{content:""}.fa-arrow-circle-up:before{content:""}.fa-arrow-down:before{content:""}.fa-arrow-left:before{content:""}.fa-arrow-right:before{content:""}.fa-arrow-up:before{content:""}.fa-arrows-alt:before{content:""}.fa-arrows-alt-h:before{content:""}.fa-arrows-alt-v:before{content:""}.fa-artstation:before{content:""}.fa-assistive-listening-systems:before{content:""}.fa-asterisk:before{content:""}.fa-asymmetrik:before{content:""}.fa-at:before{content:""}.fa-atlas:before{content:""}.fa-atlassian:before{content:""}.fa-atom:before{content:""}.fa-audible:before{content:""}.fa-audio-description:before{content:""}.fa-autoprefixer:before{content:""}.fa-avianex:before{content:""}.fa-aviato:before{content:""}.fa-award:before{content:""}.fa-aws:before{content:""}.fa-baby:before{content:""}.fa-baby-carriage:before{content:""}.fa-backspace:before{content:""}.fa-backward:before{content:""}.fa-bacon:before{content:""}.fa-bacteria:before{content:""}.fa-bacterium:before{content:""}.fa-bahai:before{content:""}.fa-balance-scale:before{content:""}.fa-balance-scale-left:before{content:""}.fa-balance-scale-right:before{content:""}.fa-ban:before{content:""}.fa-band-aid:before{content:""}.fa-bandcamp:before{content:""}.fa-barcode:before{content:""}.fa-bars:before{content:""}.fa-baseball-ball:before{content:""}.fa-basketball-ball:before{content:""}.fa-bath:before{content:""}.fa-battery-empty:before{content:""}.fa-battery-full:before{content:""}.fa-battery-half:before{content:""}.fa-battery-quarter:before{content:""}.fa-battery-three-quarters:before{content:""}.fa-battle-net:before{content:""}.fa-bed:before{content:""}.fa-beer:before{content:""}.fa-behance:before{content:""}.fa-behance-square:before{content:""}.fa-bell:before{content:""}.fa-bell-slash:before{content:""}.fa-bezier-curve:before{content:""}.fa-bible:before{content:""}.fa-bicycle:before{content:""}.fa-biking:before{content:""}.fa-bimobject:before{content:""}.fa-binoculars:before{content:""}.fa-biohazard:before{content:""}.fa-birthday-cake:before{content:""}.fa-bitbucket:before{content:""}.fa-bitcoin:before{content:""}.fa-bity:before{content:""}.fa-black-tie:before{content:""}.fa-blackberry:before{content:""}.fa-blender:before{content:""}.fa-blender-phone:before{content:""}.fa-blind:before{content:""}.fa-blog:before{content:""}.fa-blogger:before{content:""}.fa-blogger-b:before{content:""}.fa-bluetooth:before{content:""}.fa-bluetooth-b:before{content:""}.fa-bold:before{content:""}.fa-bolt:before{content:""}.fa-bomb:before{content:""}.fa-bone:before{content:""}.fa-bong:before{content:""}.fa-book:before{content:""}.fa-book-dead:before{content:""}.fa-book-medical:before{content:""}.fa-book-open:before{content:""}.fa-book-reader:before{content:""}.fa-bookmark:before{content:""}.fa-bootstrap:before{content:""}.fa-border-all:before{content:""}.fa-border-none:before{content:""}.fa-border-style:before{content:""}.fa-bowling-ball:before{content:""}.fa-box:before{content:""}.fa-box-open:before{content:""}.fa-box-tissue:before{content:""}.fa-boxes:before{content:""}.fa-braille:before{content:""}.fa-brain:before{content:""}.fa-bread-slice:before{content:""}.fa-briefcase:before{content:""}.fa-briefcase-medical:before{content:""}.fa-broadcast-tower:before{content:""}.fa-broom:before{content:""}.fa-brush:before{content:""}.fa-btc:before{content:""}.fa-buffer:before{content:""}.fa-bug:before{content:""}.fa-building:before{content:""}.fa-bullhorn:before{content:""}.fa-bullseye:before{content:""}.fa-burn:before{content:""}.fa-buromobelexperte:before{content:""}.fa-bus:before{content:""}.fa-bus-alt:before{content:""}.fa-business-time:before{content:""}.fa-buy-n-large:before{content:""}.fa-buysellads:before{content:""}.fa-calculator:before{content:""}.fa-calendar:before{content:""}.fa-calendar-alt:before{content:""}.fa-calendar-check:before{content:""}.fa-calendar-day:before{content:""}.fa-calendar-minus:before{content:""}.fa-calendar-plus:before{content:""}.fa-calendar-times:before{content:""}.fa-calendar-week:before{content:""}.fa-camera:before{content:""}.fa-camera-retro:before{content:""}.fa-campground:before{content:""}.fa-canadian-maple-leaf:before{content:""}.fa-candy-cane:before{content:""}.fa-cannabis:before{content:""}.fa-capsules:before{content:""}.fa-car:before{content:""}.fa-car-alt:before{content:""}.fa-car-battery:before{content:""}.fa-car-crash:before{content:""}.fa-car-side:before{content:""}.fa-caravan:before{content:""}.fa-caret-down:before{content:""}.fa-caret-left:before{content:""}.fa-caret-right:before{content:""}.fa-caret-square-down:before{content:""}.fa-caret-square-left:before{content:""}.fa-caret-square-right:before{content:""}.fa-caret-square-up:before{content:""}.fa-caret-up:before{content:""}.fa-carrot:before{content:""}.fa-cart-arrow-down:before{content:""}.fa-cart-plus:before{content:""}.fa-cash-register:before{content:""}.fa-cat:before{content:""}.fa-cc-amazon-pay:before{content:""}.fa-cc-amex:before{content:""}.fa-cc-apple-pay:before{content:""}.fa-cc-diners-club:before{content:""}.fa-cc-discover:before{content:""}.fa-cc-jcb:before{content:""}.fa-cc-mastercard:before{content:""}.fa-cc-paypal:before{content:""}.fa-cc-stripe:before{content:""}.fa-cc-visa:before{content:""}.fa-centercode:before{content:""}.fa-centos:before{content:""}.fa-certificate:before{content:""}.fa-chair:before{content:""}.fa-chalkboard:before{content:""}.fa-chalkboard-teacher:before{content:""}.fa-charging-station:before{content:""}.fa-chart-area:before{content:""}.fa-chart-bar:before{content:""}.fa-chart-line:before{content:""}.fa-chart-pie:before{content:""}.fa-check:before{content:""}.fa-check-circle:before{content:""}.fa-check-double:before{content:""}.fa-check-square:before{content:""}.fa-cheese:before{content:""}.fa-chess:before{content:""}.fa-chess-bishop:before{content:""}.fa-chess-board:before{content:""}.fa-chess-king:before{content:""}.fa-chess-knight:before{content:""}.fa-chess-pawn:before{content:""}.fa-chess-queen:before{content:""}.fa-chess-rook:before{content:""}.fa-chevron-circle-down:before{content:""}.fa-chevron-circle-left:before{content:""}.fa-chevron-circle-right:before{content:""}.fa-chevron-circle-up:before{content:""}.fa-chevron-down:before{content:""}.fa-chevron-left:before{content:""}.fa-chevron-right:before{content:""}.fa-chevron-up:before{content:""}.fa-child:before{content:""}.fa-chrome:before{content:""}.fa-chromecast:before{content:""}.fa-church:before{content:""}.fa-circle:before{content:""}.fa-circle-notch:before{content:""}.fa-city:before{content:""}.fa-clinic-medical:before{content:""}.fa-clipboard:before{content:""}.fa-clipboard-check:before{content:""}.fa-clipboard-list:before{content:""}.fa-clock:before{content:""}.fa-clone:before{content:""}.fa-closed-captioning:before{content:""}.fa-cloud:before{content:""}.fa-cloud-download-alt:before{content:""}.fa-cloud-meatball:before{content:""}.fa-cloud-moon:before{content:""}.fa-cloud-moon-rain:before{content:""}.fa-cloud-rain:before{content:""}.fa-cloud-showers-heavy:before{content:""}.fa-cloud-sun:before{content:""}.fa-cloud-sun-rain:before{content:""}.fa-cloud-upload-alt:before{content:""}.fa-cloudflare:before{content:""}.fa-cloudscale:before{content:""}.fa-cloudsmith:before{content:""}.fa-cloudversify:before{content:""}.fa-cocktail:before{content:""}.fa-code:before{content:""}.fa-code-branch:before{content:""}.fa-codepen:before{content:""}.fa-codiepie:before{content:""}.fa-coffee:before{content:""}.fa-cog:before{content:""}.fa-cogs:before{content:""}.fa-coins:before{content:""}.fa-columns:before{content:""}.fa-comment:before{content:""}.fa-comment-alt:before{content:""}.fa-comment-dollar:before{content:""}.fa-comment-dots:before{content:""}.fa-comment-medical:before{content:""}.fa-comment-slash:before{content:""}.fa-comments:before{content:""}.fa-comments-dollar:before{content:""}.fa-compact-disc:before{content:""}.fa-compass:before{content:""}.fa-compress:before{content:""}.fa-compress-alt:before{content:""}.fa-compress-arrows-alt:before{content:""}.fa-concierge-bell:before{content:""}.fa-confluence:before{content:""}.fa-connectdevelop:before{content:""}.fa-contao:before{content:""}.fa-cookie:before{content:""}.fa-cookie-bite:before{content:""}.fa-copy:before{content:""}.fa-copyright:before{content:""}.fa-cotton-bureau:before{content:""}.fa-couch:before{content:""}.fa-cpanel:before{content:""}.fa-creative-commons:before{content:""}.fa-creative-commons-by:before{content:""}.fa-creative-commons-nc:before{content:""}.fa-creative-commons-nc-eu:before{content:""}.fa-creative-commons-nc-jp:before{content:""}.fa-creative-commons-nd:before{content:""}.fa-creative-commons-pd:before{content:""}.fa-creative-commons-pd-alt:before{content:""}.fa-creative-commons-remix:before{content:""}.fa-creative-commons-sa:before{content:""}.fa-creative-commons-sampling:before{content:""}.fa-creative-commons-sampling-plus:before{content:""}.fa-creative-commons-share:before{content:""}.fa-creative-commons-zero:before{content:""}.fa-credit-card:before{content:""}.fa-critical-role:before{content:""}.fa-crop:before{content:""}.fa-crop-alt:before{content:""}.fa-cross:before{content:""}.fa-crosshairs:before{content:""}.fa-crow:before{content:""}.fa-crown:before{content:""}.fa-crutch:before{content:""}.fa-css3:before{content:""}.fa-css3-alt:before{content:""}.fa-cube:before{content:""}.fa-cubes:before{content:""}.fa-cut:before{content:""}.fa-cuttlefish:before{content:""}.fa-d-and-d:before{content:""}.fa-d-and-d-beyond:before{content:""}.fa-dailymotion:before{content:""}.fa-dashcube:before{content:""}.fa-database:before{content:""}.fa-deaf:before{content:""}.fa-deezer:before{content:""}.fa-delicious:before{content:""}.fa-democrat:before{content:""}.fa-deploydog:before{content:""}.fa-deskpro:before{content:""}.fa-desktop:before{content:""}.fa-dev:before{content:""}.fa-deviantart:before{content:""}.fa-dharmachakra:before{content:""}.fa-dhl:before{content:""}.fa-diagnoses:before{content:""}.fa-diaspora:before{content:""}.fa-dice:before{content:""}.fa-dice-d20:before{content:""}.fa-dice-d6:before{content:""}.fa-dice-five:before{content:""}.fa-dice-four:before{content:""}.fa-dice-one:before{content:""}.fa-dice-six:before{content:""}.fa-dice-three:before{content:""}.fa-dice-two:before{content:""}.fa-digg:before{content:""}.fa-digital-ocean:before{content:""}.fa-digital-tachograph:before{content:""}.fa-directions:before{content:""}.fa-discord:before{content:""}.fa-discourse:before{content:""}.fa-disease:before{content:""}.fa-divide:before{content:""}.fa-dizzy:before{content:""}.fa-dna:before{content:""}.fa-dochub:before{content:""}.fa-docker:before{content:""}.fa-dog:before{content:""}.fa-dollar-sign:before{content:""}.fa-dolly:before{content:""}.fa-dolly-flatbed:before{content:""}.fa-donate:before{content:""}.fa-door-closed:before{content:""}.fa-door-open:before{content:""}.fa-dot-circle:before{content:""}.fa-dove:before{content:""}.fa-download:before{content:""}.fa-draft2digital:before{content:""}.fa-drafting-compass:before{content:""}.fa-dragon:before{content:""}.fa-draw-polygon:before{content:""}.fa-dribbble:before{content:""}.fa-dribbble-square:before{content:""}.fa-dropbox:before{content:""}.fa-drum:before{content:""}.fa-drum-steelpan:before{content:""}.fa-drumstick-bite:before{content:""}.fa-drupal:before{content:""}.fa-dumbbell:before{content:""}.fa-dumpster:before{content:""}.fa-dumpster-fire:before{content:""}.fa-dungeon:before{content:""}.fa-dyalog:before{content:""}.fa-earlybirds:before{content:""}.fa-ebay:before{content:""}.fa-edge:before{content:""}.fa-edge-legacy:before{content:""}.fa-edit:before{content:""}.fa-egg:before{content:""}.fa-eject:before{content:""}.fa-elementor:before{content:""}.fa-ellipsis-h:before{content:""}.fa-ellipsis-v:before{content:""}.fa-ello:before{content:""}.fa-ember:before{content:""}.fa-empire:before{content:""}.fa-envelope:before{content:""}.fa-envelope-open:before{content:""}.fa-envelope-open-text:before{content:""}.fa-envelope-square:before{content:""}.fa-envira:before{content:""}.fa-equals:before{content:""}.fa-eraser:before{content:""}.fa-erlang:before{content:""}.fa-ethereum:before{content:""}.fa-ethernet:before{content:""}.fa-etsy:before{content:""}.fa-euro-sign:before{content:""}.fa-evernote:before{content:""}.fa-exchange-alt:before{content:""}.fa-exclamation:before{content:""}.fa-exclamation-circle:before{content:""}.fa-exclamation-triangle:before{content:""}.fa-expand:before{content:""}.fa-expand-alt:before{content:""}.fa-expand-arrows-alt:before{content:""}.fa-expeditedssl:before{content:""}.fa-external-link-alt:before{content:""}.fa-external-link-square-alt:before{content:""}.fa-eye:before{content:""}.fa-eye-dropper:before{content:""}.fa-eye-slash:before{content:""}.fa-facebook:before{content:""}.fa-facebook-f:before{content:""}.fa-facebook-messenger:before{content:""}.fa-facebook-square:before{content:""}.fa-fan:before{content:""}.fa-fantasy-flight-games:before{content:""}.fa-fast-backward:before{content:""}.fa-fast-forward:before{content:""}.fa-faucet:before{content:""}.fa-fax:before{content:""}.fa-feather:before{content:""}.fa-feather-alt:before{content:""}.fa-fedex:before{content:""}.fa-fedora:before{content:""}.fa-female:before{content:""}.fa-fighter-jet:before{content:""}.fa-figma:before{content:""}.fa-file:before{content:""}.fa-file-alt:before{content:""}.fa-file-archive:before{content:""}.fa-file-audio:before{content:""}.fa-file-code:before{content:""}.fa-file-contract:before{content:""}.fa-file-csv:before{content:""}.fa-file-download:before{content:""}.fa-file-excel:before{content:""}.fa-file-export:before{content:""}.fa-file-image:before{content:""}.fa-file-import:before{content:""}.fa-file-invoice:before{content:""}.fa-file-invoice-dollar:before{content:""}.fa-file-medical:before{content:""}.fa-file-medical-alt:before{content:""}.fa-file-pdf:before{content:""}.fa-file-powerpoint:before{content:""}.fa-file-prescription:before{content:""}.fa-file-signature:before{content:""}.fa-file-upload:before{content:""}.fa-file-video:before{content:""}.fa-file-word:before{content:""}.fa-fill:before{content:""}.fa-fill-drip:before{content:""}.fa-film:before{content:""}.fa-filter:before{content:""}.fa-fingerprint:before{content:""}.fa-fire:before{content:""}.fa-fire-alt:before{content:""}.fa-fire-extinguisher:before{content:""}.fa-firefox:before{content:""}.fa-firefox-browser:before{content:""}.fa-first-aid:before{content:""}.fa-first-order:before{content:""}.fa-first-order-alt:before{content:""}.fa-firstdraft:before{content:""}.fa-fish:before{content:""}.fa-fist-raised:before{content:""}.fa-flag:before{content:""}.fa-flag-checkered:before{content:""}.fa-flag-usa:before{content:""}.fa-flask:before{content:""}.fa-flickr:before{content:""}.fa-flipboard:before{content:""}.fa-flushed:before{content:""}.fa-fly:before{content:""}.fa-folder:before{content:""}.fa-folder-minus:before{content:""}.fa-folder-open:before{content:""}.fa-folder-plus:before{content:""}.fa-font:before{content:""}.fa-font-awesome:before{content:""}.fa-font-awesome-alt:before{content:""}.fa-font-awesome-flag:before{content:""}.fa-font-awesome-logo-full:before{content:""}.fa-fonticons:before{content:""}.fa-fonticons-fi:before{content:""}.fa-football-ball:before{content:""}.fa-fort-awesome:before{content:""}.fa-fort-awesome-alt:before{content:""}.fa-forumbee:before{content:""}.fa-forward:before{content:""}.fa-foursquare:before{content:""}.fa-free-code-camp:before{content:""}.fa-freebsd:before{content:""}.fa-frog:before{content:""}.fa-frown:before{content:""}.fa-frown-open:before{content:""}.fa-fulcrum:before{content:""}.fa-funnel-dollar:before{content:""}.fa-futbol:before{content:""}.fa-galactic-republic:before{content:""}.fa-galactic-senate:before{content:""}.fa-gamepad:before{content:""}.fa-gas-pump:before{content:""}.fa-gavel:before{content:""}.fa-gem:before{content:""}.fa-genderless:before{content:""}.fa-get-pocket:before{content:""}.fa-gg:before{content:""}.fa-gg-circle:before{content:""}.fa-ghost:before{content:""}.fa-gift:before{content:""}.fa-gifts:before{content:""}.fa-git:before{content:""}.fa-git-alt:before{content:""}.fa-git-square:before{content:""}.fa-github:before{content:""}.fa-github-alt:before{content:""}.fa-github-square:before{content:""}.fa-gitkraken:before{content:""}.fa-gitlab:before{content:""}.fa-gitter:before{content:""}.fa-glass-cheers:before{content:""}.fa-glass-martini:before{content:""}.fa-glass-martini-alt:before{content:""}.fa-glass-whiskey:before{content:""}.fa-glasses:before{content:""}.fa-glide:before{content:""}.fa-glide-g:before{content:""}.fa-globe:before{content:""}.fa-globe-africa:before{content:""}.fa-globe-americas:before{content:""}.fa-globe-asia:before{content:""}.fa-globe-europe:before{content:""}.fa-gofore:before{content:""}.fa-golf-ball:before{content:""}.fa-goodreads:before{content:""}.fa-goodreads-g:before{content:""}.fa-google:before{content:""}.fa-google-drive:before{content:""}.fa-google-pay:before{content:""}.fa-google-play:before{content:""}.fa-google-plus:before{content:""}.fa-google-plus-g:before{content:""}.fa-google-plus-square:before{content:""}.fa-google-wallet:before{content:""}.fa-gopuram:before{content:""}.fa-graduation-cap:before{content:""}.fa-gratipay:before{content:""}.fa-grav:before{content:""}.fa-greater-than:before{content:""}.fa-greater-than-equal:before{content:""}.fa-grimace:before{content:""}.fa-grin:before{content:""}.fa-grin-alt:before{content:""}.fa-grin-beam:before{content:""}.fa-grin-beam-sweat:before{content:""}.fa-grin-hearts:before{content:""}.fa-grin-squint:before{content:""}.fa-grin-squint-tears:before{content:""}.fa-grin-stars:before{content:""}.fa-grin-tears:before{content:""}.fa-grin-tongue:before{content:""}.fa-grin-tongue-squint:before{content:""}.fa-grin-tongue-wink:before{content:""}.fa-grin-wink:before{content:""}.fa-grip-horizontal:before{content:""}.fa-grip-lines:before{content:""}.fa-grip-lines-vertical:before{content:""}.fa-grip-vertical:before{content:""}.fa-gripfire:before{content:""}.fa-grunt:before{content:""}.fa-guilded:before{content:""}.fa-guitar:before{content:""}.fa-gulp:before{content:""}.fa-h-square:before{content:""}.fa-hacker-news:before{content:""}.fa-hacker-news-square:before{content:""}.fa-hackerrank:before{content:""}.fa-hamburger:before{content:""}.fa-hammer:before{content:""}.fa-hamsa:before{content:""}.fa-hand-holding:before{content:""}.fa-hand-holding-heart:before{content:""}.fa-hand-holding-medical:before{content:""}.fa-hand-holding-usd:before{content:""}.fa-hand-holding-water:before{content:""}.fa-hand-lizard:before{content:""}.fa-hand-middle-finger:before{content:""}.fa-hand-paper:before{content:""}.fa-hand-peace:before{content:""}.fa-hand-point-down:before{content:""}.fa-hand-point-left:before{content:""}.fa-hand-point-right:before{content:""}.fa-hand-point-up:before{content:""}.fa-hand-pointer:before{content:""}.fa-hand-rock:before{content:""}.fa-hand-scissors:before{content:""}.fa-hand-sparkles:before{content:""}.fa-hand-spock:before{content:""}.fa-hands:before{content:""}.fa-hands-helping:before{content:""}.fa-hands-wash:before{content:""}.fa-handshake:before{content:""}.fa-handshake-alt-slash:before{content:""}.fa-handshake-slash:before{content:""}.fa-hanukiah:before{content:""}.fa-hard-hat:before{content:""}.fa-hashtag:before{content:""}.fa-hat-cowboy:before{content:""}.fa-hat-cowboy-side:before{content:""}.fa-hat-wizard:before{content:""}.fa-hdd:before{content:""}.fa-head-side-cough:before{content:""}.fa-head-side-cough-slash:before{content:""}.fa-head-side-mask:before{content:""}.fa-head-side-virus:before{content:""}.fa-heading:before{content:""}.fa-headphones:before{content:""}.fa-headphones-alt:before{content:""}.fa-headset:before{content:""}.fa-heart:before{content:""}.fa-heart-broken:before{content:""}.fa-heartbeat:before{content:""}.fa-helicopter:before{content:""}.fa-highlighter:before{content:""}.fa-hiking:before{content:""}.fa-hippo:before{content:""}.fa-hips:before{content:""}.fa-hire-a-helper:before{content:""}.fa-history:before{content:""}.fa-hive:before{content:""}.fa-hockey-puck:before{content:""}.fa-holly-berry:before{content:""}.fa-home:before{content:""}.fa-hooli:before{content:""}.fa-hornbill:before{content:""}.fa-horse:before{content:""}.fa-horse-head:before{content:""}.fa-hospital:before{content:""}.fa-hospital-alt:before{content:""}.fa-hospital-symbol:before{content:""}.fa-hospital-user:before{content:""}.fa-hot-tub:before{content:""}.fa-hotdog:before{content:""}.fa-hotel:before{content:""}.fa-hotjar:before{content:""}.fa-hourglass:before{content:""}.fa-hourglass-end:before{content:""}.fa-hourglass-half:before{content:""}.fa-hourglass-start:before{content:""}.fa-house-damage:before{content:""}.fa-house-user:before{content:""}.fa-houzz:before{content:""}.fa-hryvnia:before{content:""}.fa-html5:before{content:""}.fa-hubspot:before{content:""}.fa-i-cursor:before{content:""}.fa-ice-cream:before{content:""}.fa-icicles:before{content:""}.fa-icons:before{content:""}.fa-id-badge:before{content:""}.fa-id-card:before{content:""}.fa-id-card-alt:before{content:""}.fa-ideal:before{content:""}.fa-igloo:before{content:""}.fa-image:before{content:""}.fa-images:before{content:""}.fa-imdb:before{content:""}.fa-inbox:before{content:""}.fa-indent:before{content:""}.fa-industry:before{content:""}.fa-infinity:before{content:""}.fa-info:before{content:""}.fa-info-circle:before{content:""}.fa-innosoft:before{content:""}.fa-instagram:before{content:""}.fa-instagram-square:before{content:""}.fa-instalod:before{content:""}.fa-intercom:before{content:""}.fa-internet-explorer:before{content:""}.fa-invision:before{content:""}.fa-ioxhost:before{content:""}.fa-italic:before{content:""}.fa-itch-io:before{content:""}.fa-itunes:before{content:""}.fa-itunes-note:before{content:""}.fa-java:before{content:""}.fa-jedi:before{content:""}.fa-jedi-order:before{content:""}.fa-jenkins:before{content:""}.fa-jira:before{content:""}.fa-joget:before{content:""}.fa-joint:before{content:""}.fa-joomla:before{content:""}.fa-journal-whills:before{content:""}.fa-js:before{content:""}.fa-js-square:before{content:""}.fa-jsfiddle:before{content:""}.fa-kaaba:before{content:""}.fa-kaggle:before{content:""}.fa-key:before{content:""}.fa-keybase:before{content:""}.fa-keyboard:before{content:""}.fa-keycdn:before{content:""}.fa-khanda:before{content:""}.fa-kickstarter:before{content:""}.fa-kickstarter-k:before{content:""}.fa-kiss:before{content:""}.fa-kiss-beam:before{content:""}.fa-kiss-wink-heart:before{content:""}.fa-kiwi-bird:before{content:""}.fa-korvue:before{content:""}.fa-landmark:before{content:""}.fa-language:before{content:""}.fa-laptop:before{content:""}.fa-laptop-code:before{content:""}.fa-laptop-house:before{content:""}.fa-laptop-medical:before{content:""}.fa-laravel:before{content:""}.fa-lastfm:before{content:""}.fa-lastfm-square:before{content:""}.fa-laugh:before{content:""}.fa-laugh-beam:before{content:""}.fa-laugh-squint:before{content:""}.fa-laugh-wink:before{content:""}.fa-layer-group:before{content:""}.fa-leaf:before{content:""}.fa-leanpub:before{content:""}.fa-lemon:before{content:""}.fa-less:before{content:""}.fa-less-than:before{content:""}.fa-less-than-equal:before{content:""}.fa-level-down-alt:before{content:""}.fa-level-up-alt:before{content:""}.fa-life-ring:before{content:""}.fa-lightbulb:before{content:""}.fa-line:before{content:""}.fa-link:before{content:""}.fa-linkedin:before{content:""}.fa-linkedin-in:before{content:""}.fa-linode:before{content:""}.fa-linux:before{content:""}.fa-lira-sign:before{content:""}.fa-list:before{content:""}.fa-list-alt:before{content:""}.fa-list-ol:before{content:""}.fa-list-ul:before{content:""}.fa-location-arrow:before{content:""}.fa-lock:before{content:""}.fa-lock-open:before{content:""}.fa-long-arrow-alt-down:before{content:""}.fa-long-arrow-alt-left:before{content:""}.fa-long-arrow-alt-right:before{content:""}.fa-long-arrow-alt-up:before{content:""}.fa-low-vision:before{content:""}.fa-luggage-cart:before{content:""}.fa-lungs:before{content:""}.fa-lungs-virus:before{content:""}.fa-lyft:before{content:""}.fa-magento:before{content:""}.fa-magic:before{content:""}.fa-magnet:before{content:""}.fa-mail-bulk:before{content:""}.fa-mailchimp:before{content:""}.fa-male:before{content:""}.fa-mandalorian:before{content:""}.fa-map:before{content:""}.fa-map-marked:before{content:""}.fa-map-marked-alt:before{content:""}.fa-map-marker:before{content:""}.fa-map-marker-alt:before{content:""}.fa-map-pin:before{content:""}.fa-map-signs:before{content:""}.fa-markdown:before{content:""}.fa-marker:before{content:""}.fa-mars:before{content:""}.fa-mars-double:before{content:""}.fa-mars-stroke:before{content:""}.fa-mars-stroke-h:before{content:""}.fa-mars-stroke-v:before{content:""}.fa-mask:before{content:""}.fa-mastodon:before{content:""}.fa-maxcdn:before{content:""}.fa-mdb:before{content:""}.fa-medal:before{content:""}.fa-medapps:before{content:""}.fa-medium:before{content:""}.fa-medium-m:before{content:""}.fa-medkit:before{content:""}.fa-medrt:before{content:""}.fa-meetup:before{content:""}.fa-megaport:before{content:""}.fa-meh:before{content:""}.fa-meh-blank:before{content:""}.fa-meh-rolling-eyes:before{content:""}.fa-memory:before{content:""}.fa-mendeley:before{content:""}.fa-menorah:before{content:""}.fa-mercury:before{content:""}.fa-meteor:before{content:""}.fa-microblog:before{content:""}.fa-microchip:before{content:""}.fa-microphone:before{content:""}.fa-microphone-alt:before{content:""}.fa-microphone-alt-slash:before{content:""}.fa-microphone-slash:before{content:""}.fa-microscope:before{content:""}.fa-microsoft:before{content:""}.fa-minus:before{content:""}.fa-minus-circle:before{content:""}.fa-minus-square:before{content:""}.fa-mitten:before{content:""}.fa-mix:before{content:""}.fa-mixcloud:before{content:""}.fa-mixer:before{content:""}.fa-mizuni:before{content:""}.fa-mobile:before{content:""}.fa-mobile-alt:before{content:""}.fa-modx:before{content:""}.fa-monero:before{content:""}.fa-money-bill:before{content:""}.fa-money-bill-alt:before{content:""}.fa-money-bill-wave:before{content:""}.fa-money-bill-wave-alt:before{content:""}.fa-money-check:before{content:""}.fa-money-check-alt:before{content:""}.fa-monument:before{content:""}.fa-moon:before{content:""}.fa-mortar-pestle:before{content:""}.fa-mosque:before{content:""}.fa-motorcycle:before{content:""}.fa-mountain:before{content:""}.fa-mouse:before{content:""}.fa-mouse-pointer:before{content:""}.fa-mug-hot:before{content:""}.fa-music:before{content:""}.fa-napster:before{content:""}.fa-neos:before{content:""}.fa-network-wired:before{content:""}.fa-neuter:before{content:""}.fa-newspaper:before{content:""}.fa-nimblr:before{content:""}.fa-node:before{content:""}.fa-node-js:before{content:""}.fa-not-equal:before{content:""}.fa-notes-medical:before{content:""}.fa-npm:before{content:""}.fa-ns8:before{content:""}.fa-nutritionix:before{content:""}.fa-object-group:before{content:""}.fa-object-ungroup:before{content:""}.fa-octopus-deploy:before{content:""}.fa-odnoklassniki:before{content:""}.fa-odnoklassniki-square:before{content:""}.fa-oil-can:before{content:""}.fa-old-republic:before{content:""}.fa-om:before{content:""}.fa-opencart:before{content:""}.fa-openid:before{content:""}.fa-opera:before{content:""}.fa-optin-monster:before{content:""}.fa-orcid:before{content:""}.fa-osi:before{content:""}.fa-otter:before{content:""}.fa-outdent:before{content:""}.fa-page4:before{content:""}.fa-pagelines:before{content:""}.fa-pager:before{content:""}.fa-paint-brush:before{content:""}.fa-paint-roller:before{content:""}.fa-palette:before{content:""}.fa-palfed:before{content:""}.fa-pallet:before{content:""}.fa-paper-plane:before{content:""}.fa-paperclip:before{content:""}.fa-parachute-box:before{content:""}.fa-paragraph:before{content:""}.fa-parking:before{content:""}.fa-passport:before{content:""}.fa-pastafarianism:before{content:""}.fa-paste:before{content:""}.fa-patreon:before{content:""}.fa-pause:before{content:""}.fa-pause-circle:before{content:""}.fa-paw:before{content:""}.fa-paypal:before{content:""}.fa-peace:before{content:""}.fa-pen:before{content:""}.fa-pen-alt:before{content:""}.fa-pen-fancy:before{content:""}.fa-pen-nib:before{content:""}.fa-pen-square:before{content:""}.fa-pencil-alt:before{content:""}.fa-pencil-ruler:before{content:""}.fa-penny-arcade:before{content:""}.fa-people-arrows:before{content:""}.fa-people-carry:before{content:""}.fa-pepper-hot:before{content:""}.fa-perbyte:before{content:""}.fa-percent:before{content:""}.fa-percentage:before{content:""}.fa-periscope:before{content:""}.fa-person-booth:before{content:""}.fa-phabricator:before{content:""}.fa-phoenix-framework:before{content:""}.fa-phoenix-squadron:before{content:""}.fa-phone:before{content:""}.fa-phone-alt:before{content:""}.fa-phone-slash:before{content:""}.fa-phone-square:before{content:""}.fa-phone-square-alt:before{content:""}.fa-phone-volume:before{content:""}.fa-photo-video:before{content:""}.fa-php:before{content:""}.fa-pied-piper:before{content:""}.fa-pied-piper-alt:before{content:""}.fa-pied-piper-hat:before{content:""}.fa-pied-piper-pp:before{content:""}.fa-pied-piper-square:before{content:""}.fa-piggy-bank:before{content:""}.fa-pills:before{content:""}.fa-pinterest:before{content:""}.fa-pinterest-p:before{content:""}.fa-pinterest-square:before{content:""}.fa-pizza-slice:before{content:""}.fa-place-of-worship:before{content:""}.fa-plane:before{content:""}.fa-plane-arrival:before{content:""}.fa-plane-departure:before{content:""}.fa-plane-slash:before{content:""}.fa-play:before{content:""}.fa-play-circle:before{content:""}.fa-playstation:before{content:""}.fa-plug:before{content:""}.fa-plus:before{content:""}.fa-plus-circle:before{content:""}.fa-plus-square:before{content:""}.fa-podcast:before{content:""}.fa-poll:before{content:""}.fa-poll-h:before{content:""}.fa-poo:before{content:""}.fa-poo-storm:before{content:""}.fa-poop:before{content:""}.fa-portrait:before{content:""}.fa-pound-sign:before{content:""}.fa-power-off:before{content:""}.fa-pray:before{content:""}.fa-praying-hands:before{content:""}.fa-prescription:before{content:""}.fa-prescription-bottle:before{content:""}.fa-prescription-bottle-alt:before{content:""}.fa-print:before{content:""}.fa-procedures:before{content:""}.fa-product-hunt:before{content:""}.fa-project-diagram:before{content:""}.fa-pump-medical:before{content:""}.fa-pump-soap:before{content:""}.fa-pushed:before{content:""}.fa-puzzle-piece:before{content:""}.fa-python:before{content:""}.fa-qq:before{content:""}.fa-qrcode:before{content:""}.fa-question:before{content:""}.fa-question-circle:before{content:""}.fa-quidditch:before{content:""}.fa-quinscape:before{content:""}.fa-quora:before{content:""}.fa-quote-left:before{content:""}.fa-quote-right:before{content:""}.fa-quran:before{content:""}.fa-r-project:before{content:""}.fa-radiation:before{content:""}.fa-radiation-alt:before{content:""}.fa-rainbow:before{content:""}.fa-random:before{content:""}.fa-raspberry-pi:before{content:""}.fa-ravelry:before{content:""}.fa-react:before{content:""}.fa-reacteurope:before{content:""}.fa-readme:before{content:""}.fa-rebel:before{content:""}.fa-receipt:before{content:""}.fa-record-vinyl:before{content:""}.fa-recycle:before{content:""}.fa-red-river:before{content:""}.fa-reddit:before{content:""}.fa-reddit-alien:before{content:""}.fa-reddit-square:before{content:""}.fa-redhat:before{content:""}.fa-redo:before{content:""}.fa-redo-alt:before{content:""}.fa-registered:before{content:""}.fa-remove-format:before{content:""}.fa-renren:before{content:""}.fa-reply:before{content:""}.fa-reply-all:before{content:""}.fa-replyd:before{content:""}.fa-republican:before{content:""}.fa-researchgate:before{content:""}.fa-resolving:before{content:""}.fa-restroom:before{content:""}.fa-retweet:before{content:""}.fa-rev:before{content:""}.fa-ribbon:before{content:""}.fa-ring:before{content:""}.fa-road:before{content:""}.fa-robot:before{content:""}.fa-rocket:before{content:""}.fa-rocketchat:before{content:""}.fa-rockrms:before{content:""}.fa-route:before{content:""}.fa-rss:before{content:""}.fa-rss-square:before{content:""}.fa-ruble-sign:before{content:""}.fa-ruler:before{content:""}.fa-ruler-combined:before{content:""}.fa-ruler-horizontal:before{content:""}.fa-ruler-vertical:before{content:""}.fa-running:before{content:""}.fa-rupee-sign:before{content:""}.fa-rust:before{content:""}.fa-sad-cry:before{content:""}.fa-sad-tear:before{content:""}.fa-safari:before{content:""}.fa-salesforce:before{content:""}.fa-sass:before{content:""}.fa-satellite:before{content:""}.fa-satellite-dish:before{content:""}.fa-save:before{content:""}.fa-schlix:before{content:""}.fa-school:before{content:""}.fa-screwdriver:before{content:""}.fa-scribd:before{content:""}.fa-scroll:before{content:""}.fa-sd-card:before{content:""}.fa-search:before{content:""}.fa-search-dollar:before{content:""}.fa-search-location:before{content:""}.fa-search-minus:before{content:""}.fa-search-plus:before{content:""}.fa-searchengin:before{content:""}.fa-seedling:before{content:""}.fa-sellcast:before{content:""}.fa-sellsy:before{content:""}.fa-server:before{content:""}.fa-servicestack:before{content:""}.fa-shapes:before{content:""}.fa-share:before{content:""}.fa-share-alt:before{content:""}.fa-share-alt-square:before{content:""}.fa-share-square:before{content:""}.fa-shekel-sign:before{content:""}.fa-shield-alt:before{content:""}.fa-shield-virus:before{content:""}.fa-ship:before{content:""}.fa-shipping-fast:before{content:""}.fa-shirtsinbulk:before{content:""}.fa-shoe-prints:before{content:""}.fa-shopify:before{content:""}.fa-shopping-bag:before{content:""}.fa-shopping-basket:before{content:""}.fa-shopping-cart:before{content:""}.fa-shopware:before{content:""}.fa-shower:before{content:""}.fa-shuttle-van:before{content:""}.fa-sign:before{content:""}.fa-sign-in-alt:before{content:""}.fa-sign-language:before{content:""}.fa-sign-out-alt:before{content:""}.fa-signal:before{content:""}.fa-signature:before{content:""}.fa-sim-card:before{content:""}.fa-simplybuilt:before{content:""}.fa-sink:before{content:""}.fa-sistrix:before{content:""}.fa-sitemap:before{content:""}.fa-sith:before{content:""}.fa-skating:before{content:""}.fa-sketch:before{content:""}.fa-skiing:before{content:""}.fa-skiing-nordic:before{content:""}.fa-skull:before{content:""}.fa-skull-crossbones:before{content:""}.fa-skyatlas:before{content:""}.fa-skype:before{content:""}.fa-slack:before{content:""}.fa-slack-hash:before{content:""}.fa-slash:before{content:""}.fa-sleigh:before{content:""}.fa-sliders-h:before{content:""}.fa-slideshare:before{content:""}.fa-smile:before{content:""}.fa-smile-beam:before{content:""}.fa-smile-wink:before{content:""}.fa-smog:before{content:""}.fa-smoking:before{content:""}.fa-smoking-ban:before{content:""}.fa-sms:before{content:""}.fa-snapchat:before{content:""}.fa-snapchat-ghost:before{content:""}.fa-snapchat-square:before{content:""}.fa-snowboarding:before{content:""}.fa-snowflake:before{content:""}.fa-snowman:before{content:""}.fa-snowplow:before{content:""}.fa-soap:before{content:""}.fa-socks:before{content:""}.fa-solar-panel:before{content:""}.fa-sort:before{content:""}.fa-sort-alpha-down:before{content:""}.fa-sort-alpha-down-alt:before{content:""}.fa-sort-alpha-up:before{content:""}.fa-sort-alpha-up-alt:before{content:""}.fa-sort-amount-down:before{content:""}.fa-sort-amount-down-alt:before{content:""}.fa-sort-amount-up:before{content:""}.fa-sort-amount-up-alt:before{content:""}.fa-sort-down:before{content:""}.fa-sort-numeric-down:before{content:""}.fa-sort-numeric-down-alt:before{content:""}.fa-sort-numeric-up:before{content:""}.fa-sort-numeric-up-alt:before{content:""}.fa-sort-up:before{content:""}.fa-soundcloud:before{content:""}.fa-sourcetree:before{content:""}.fa-spa:before{content:""}.fa-space-shuttle:before{content:""}.fa-speakap:before{content:""}.fa-speaker-deck:before{content:""}.fa-spell-check:before{content:""}.fa-spider:before{content:""}.fa-spinner:before{content:""}.fa-splotch:before{content:""}.fa-spotify:before{content:""}.fa-spray-can:before{content:""}.fa-square:before{content:""}.fa-square-full:before{content:""}.fa-square-root-alt:before{content:""}.fa-squarespace:before{content:""}.fa-stack-exchange:before{content:""}.fa-stack-overflow:before{content:""}.fa-stackpath:before{content:""}.fa-stamp:before{content:""}.fa-star:before{content:""}.fa-star-and-crescent:before{content:""}.fa-star-half:before{content:""}.fa-star-half-alt:before{content:""}.fa-star-of-david:before{content:""}.fa-star-of-life:before{content:""}.fa-staylinked:before{content:""}.fa-steam:before{content:""}.fa-steam-square:before{content:""}.fa-steam-symbol:before{content:""}.fa-step-backward:before{content:""}.fa-step-forward:before{content:""}.fa-stethoscope:before{content:""}.fa-sticker-mule:before{content:""}.fa-sticky-note:before{content:""}.fa-stop:before{content:""}.fa-stop-circle:before{content:""}.fa-stopwatch:before{content:""}.fa-stopwatch-20:before{content:""}.fa-store:before{content:""}.fa-store-alt:before{content:""}.fa-store-alt-slash:before{content:""}.fa-store-slash:before{content:""}.fa-strava:before{content:""}.fa-stream:before{content:""}.fa-street-view:before{content:""}.fa-strikethrough:before{content:""}.fa-stripe:before{content:""}.fa-stripe-s:before{content:""}.fa-stroopwafel:before{content:""}.fa-studiovinari:before{content:""}.fa-stumbleupon:before{content:""}.fa-stumbleupon-circle:before{content:""}.fa-subscript:before{content:""}.fa-subway:before{content:""}.fa-suitcase:before{content:""}.fa-suitcase-rolling:before{content:""}.fa-sun:before{content:""}.fa-superpowers:before{content:""}.fa-superscript:before{content:""}.fa-supple:before{content:""}.fa-surprise:before{content:""}.fa-suse:before{content:""}.fa-swatchbook:before{content:""}.fa-swift:before{content:""}.fa-swimmer:before{content:""}.fa-swimming-pool:before{content:""}.fa-symfony:before{content:""}.fa-synagogue:before{content:""}.fa-sync:before{content:""}.fa-sync-alt:before{content:""}.fa-syringe:before{content:""}.fa-table:before{content:""}.fa-table-tennis:before{content:""}.fa-tablet:before{content:""}.fa-tablet-alt:before{content:""}.fa-tablets:before{content:""}.fa-tachometer-alt:before{content:""}.fa-tag:before{content:""}.fa-tags:before{content:""}.fa-tape:before{content:""}.fa-tasks:before{content:""}.fa-taxi:before{content:""}.fa-teamspeak:before{content:""}.fa-teeth:before{content:""}.fa-teeth-open:before{content:""}.fa-telegram:before{content:""}.fa-telegram-plane:before{content:""}.fa-temperature-high:before{content:""}.fa-temperature-low:before{content:""}.fa-tencent-weibo:before{content:""}.fa-tenge:before{content:""}.fa-terminal:before{content:""}.fa-text-height:before{content:""}.fa-text-width:before{content:""}.fa-th:before{content:""}.fa-th-large:before{content:""}.fa-th-list:before{content:""}.fa-the-red-yeti:before{content:""}.fa-theater-masks:before{content:""}.fa-themeco:before{content:""}.fa-themeisle:before{content:""}.fa-thermometer:before{content:""}.fa-thermometer-empty:before{content:""}.fa-thermometer-full:before{content:""}.fa-thermometer-half:before{content:""}.fa-thermometer-quarter:before{content:""}.fa-thermometer-three-quarters:before{content:""}.fa-think-peaks:before{content:""}.fa-thumbs-down:before{content:""}.fa-thumbs-up:before{content:""}.fa-thumbtack:before{content:""}.fa-ticket-alt:before{content:""}.fa-tiktok:before{content:""}.fa-times:before{content:""}.fa-times-circle:before{content:""}.fa-tint:before{content:""}.fa-tint-slash:before{content:""}.fa-tired:before{content:""}.fa-toggle-off:before{content:""}.fa-toggle-on:before{content:""}.fa-toilet:before{content:""}.fa-toilet-paper:before{content:""}.fa-toilet-paper-slash:before{content:""}.fa-toolbox:before{content:""}.fa-tools:before{content:""}.fa-tooth:before{content:""}.fa-torah:before{content:""}.fa-torii-gate:before{content:""}.fa-tractor:before{content:""}.fa-trade-federation:before{content:""}.fa-trademark:before{content:""}.fa-traffic-light:before{content:""}.fa-trailer:before{content:""}.fa-train:before{content:""}.fa-tram:before{content:""}.fa-transgender:before{content:""}.fa-transgender-alt:before{content:""}.fa-trash:before{content:""}.fa-trash-alt:before{content:""}.fa-trash-restore:before{content:""}.fa-trash-restore-alt:before{content:""}.fa-tree:before{content:""}.fa-trello:before{content:""}.fa-tripadvisor:before{content:""}.fa-trophy:before{content:""}.fa-truck:before{content:""}.fa-truck-loading:before{content:""}.fa-truck-monster:before{content:""}.fa-truck-moving:before{content:""}.fa-truck-pickup:before{content:""}.fa-tshirt:before{content:""}.fa-tty:before{content:""}.fa-tumblr:before{content:""}.fa-tumblr-square:before{content:""}.fa-tv:before{content:""}.fa-twitch:before{content:""}.fa-twitter:before{content:""}.fa-twitter-square:before{content:""}.fa-typo3:before{content:""}.fa-uber:before{content:""}.fa-ubuntu:before{content:""}.fa-uikit:before{content:""}.fa-umbraco:before{content:""}.fa-umbrella:before{content:""}.fa-umbrella-beach:before{content:""}.fa-uncharted:before{content:""}.fa-underline:before{content:""}.fa-undo:before{content:""}.fa-undo-alt:before{content:""}.fa-uniregistry:before{content:""}.fa-unity:before{content:""}.fa-universal-access:before{content:""}.fa-university:before{content:""}.fa-unlink:before{content:""}.fa-unlock:before{content:""}.fa-unlock-alt:before{content:""}.fa-unsplash:before{content:""}.fa-untappd:before{content:""}.fa-upload:before{content:""}.fa-ups:before{content:""}.fa-usb:before{content:""}.fa-user:before{content:""}.fa-user-alt:before{content:""}.fa-user-alt-slash:before{content:""}.fa-user-astronaut:before{content:""}.fa-user-check:before{content:""}.fa-user-circle:before{content:""}.fa-user-clock:before{content:""}.fa-user-cog:before{content:""}.fa-user-edit:before{content:""}.fa-user-friends:before{content:""}.fa-user-graduate:before{content:""}.fa-user-injured:before{content:""}.fa-user-lock:before{content:""}.fa-user-md:before{content:""}.fa-user-minus:before{content:""}.fa-user-ninja:before{content:""}.fa-user-nurse:before{content:""}.fa-user-plus:before{content:""}.fa-user-secret:before{content:""}.fa-user-shield:before{content:""}.fa-user-slash:before{content:""}.fa-user-tag:before{content:""}.fa-user-tie:before{content:""}.fa-user-times:before{content:""}.fa-users:before{content:""}.fa-users-cog:before{content:""}.fa-users-slash:before{content:""}.fa-usps:before{content:""}.fa-ussunnah:before{content:""}.fa-utensil-spoon:before{content:""}.fa-utensils:before{content:""}.fa-vaadin:before{content:""}.fa-vector-square:before{content:""}.fa-venus:before{content:""}.fa-venus-double:before{content:""}.fa-venus-mars:before{content:""}.fa-vest:before{content:""}.fa-vest-patches:before{content:""}.fa-viacoin:before{content:""}.fa-viadeo:before{content:""}.fa-viadeo-square:before{content:""}.fa-vial:before{content:""}.fa-vials:before{content:""}.fa-viber:before{content:""}.fa-video:before{content:""}.fa-video-slash:before{content:""}.fa-vihara:before{content:""}.fa-vimeo:before{content:""}.fa-vimeo-square:before{content:""}.fa-vimeo-v:before{content:""}.fa-vine:before{content:""}.fa-virus:before{content:""}.fa-virus-slash:before{content:""}.fa-viruses:before{content:""}.fa-vk:before{content:""}.fa-vnv:before{content:""}.fa-voicemail:before{content:""}.fa-volleyball-ball:before{content:""}.fa-volume-down:before{content:""}.fa-volume-mute:before{content:""}.fa-volume-off:before{content:""}.fa-volume-up:before{content:""}.fa-vote-yea:before{content:""}.fa-vr-cardboard:before{content:""}.fa-vuejs:before{content:""}.fa-walking:before{content:""}.fa-wallet:before{content:""}.fa-warehouse:before{content:""}.fa-watchman-monitoring:before{content:""}.fa-water:before{content:""}.fa-wave-square:before{content:""}.fa-waze:before{content:""}.fa-weebly:before{content:""}.fa-weibo:before{content:""}.fa-weight:before{content:""}.fa-weight-hanging:before{content:""}.fa-weixin:before{content:""}.fa-whatsapp:before{content:""}.fa-whatsapp-square:before{content:""}.fa-wheelchair:before{content:""}.fa-whmcs:before{content:""}.fa-wifi:before{content:""}.fa-wikipedia-w:before{content:""}.fa-wind:before{content:""}.fa-window-close:before{content:""}.fa-window-maximize:before{content:""}.fa-window-minimize:before{content:""}.fa-window-restore:before{content:""}.fa-windows:before{content:""}.fa-wine-bottle:before{content:""}.fa-wine-glass:before{content:""}.fa-wine-glass-alt:before{content:""}.fa-wix:before{content:""}.fa-wizards-of-the-coast:before{content:""}.fa-wodu:before{content:""}.fa-wolf-pack-battalion:before{content:""}.fa-won-sign:before{content:""}.fa-wordpress:before{content:""}.fa-wordpress-simple:before{content:""}.fa-wpbeginner:before{content:""}.fa-wpexplorer:before{content:""}.fa-wpforms:before{content:""}.fa-wpressr:before{content:""}.fa-wrench:before{content:""}.fa-x-ray:before{content:""}.fa-xbox:before{content:""}.fa-xing:before{content:""}.fa-xing-square:before{content:""}.fa-y-combinator:before{content:""}.fa-yahoo:before{content:""}.fa-yammer:before{content:""}.fa-yandex:before{content:""}.fa-yandex-international:before{content:""}.fa-yarn:before{content:""}.fa-yelp:before{content:""}.fa-yen-sign:before{content:""}.fa-yin-yang:before{content:""}.fa-yoast:before{content:""}.fa-youtube:before{content:""}.fa-youtube-square:before{content:""}.fa-zhihu:before{content:""}.sr-only{border:0;clip:rect(0, 0, 0, 0);height:1px;margin:-1px;overflow:hidden;padding:0;position:absolute;width:1px}.sr-only-focusable:active,.sr-only-focusable:focus{clip:auto;height:auto;margin:0;overflow:visible;position:static;width:auto}/*!\n * Font Awesome Free 5.15.2 by @fontawesome - https://fontawesome.com\n * License - https://fontawesome.com/license/free (Icons: CC BY 4.0, Fonts: SIL OFL 1.1, Code: MIT License)\n */.fa.fa-glass:before{content:""}.fa.fa-meetup{font-family:"Font Awesome 5 Brands";font-weight:400}.fa.fa-star-o{font-family:"Font Awesome 5 Free";font-weight:400}.fa.fa-star-o:before{content:""}.fa.fa-remove:before{content:""}.fa.fa-close:before{content:""}.fa.fa-gear:before{content:""}.fa.fa-trash-o{font-family:"Font Awesome 5 Free";font-weight:400}.fa.fa-trash-o:before{content:""}.fa.fa-file-o{font-family:"Font Awesome 5 Free";font-weight:400}.fa.fa-file-o:before{content:""}.fa.fa-clock-o{font-family:"Font Awesome 5 Free";font-weight:400}.fa.fa-clock-o:before{content:""}.fa.fa-arrow-circle-o-down{font-family:"Font Awesome 5 Free";font-weight:400}.fa.fa-arrow-circle-o-down:before{content:""}.fa.fa-arrow-circle-o-up{font-family:"Font Awesome 5 Free";font-weight:400}.fa.fa-arrow-circle-o-up:before{content:""}.fa.fa-play-circle-o{font-family:"Font Awesome 5 Free";font-weight:400}.fa.fa-play-circle-o:before{content:""}.fa.fa-repeat:before{content:""}.fa.fa-rotate-right:before{content:""}.fa.fa-refresh:before{content:""}.fa.fa-list-alt{font-family:"Font Awesome 5 Free";font-weight:400}.fa.fa-dedent:before{content:""}.fa.fa-video-camera:before{content:""}.fa.fa-picture-o{font-family:"Font Awesome 5 Free";font-weight:400}.fa.fa-picture-o:before{content:""}.fa.fa-photo{font-family:"Font Awesome 5 Free";font-weight:400}.fa.fa-photo:before{content:""}.fa.fa-image{font-family:"Font Awesome 5 Free";font-weight:400}.fa.fa-image:before{content:""}.fa.fa-pencil:before{content:""}.fa.fa-map-marker:before{content:""}.fa.fa-pencil-square-o{font-family:"Font Awesome 5 Free";font-weight:400}.fa.fa-pencil-square-o:before{content:""}.fa.fa-share-square-o{font-family:"Font Awesome 5 Free";font-weight:400}.fa.fa-share-square-o:before{content:""}.fa.fa-check-square-o{font-family:"Font Awesome 5 Free";font-weight:400}.fa.fa-check-square-o:before{content:""}.fa.fa-arrows:before{content:""}.fa.fa-times-circle-o{font-family:"Font Awesome 5 Free";font-weight:400}.fa.fa-times-circle-o:before{content:""}.fa.fa-check-circle-o{font-family:"Font Awesome 5 Free";font-weight:400}.fa.fa-check-circle-o:before{content:""}.fa.fa-mail-forward:before{content:""}.fa.fa-expand:before{content:""}.fa.fa-compress:before{content:""}.fa.fa-eye{font-family:"Font Awesome 5 Free";font-weight:400}.fa.fa-eye-slash{font-family:"Font Awesome 5 Free";font-weight:400}.fa.fa-warning:before{content:""}.fa.fa-calendar:before{content:""}.fa.fa-arrows-v:before{content:""}.fa.fa-arrows-h:before{content:""}.fa.fa-bar-chart{font-family:"Font Awesome 5 Free";font-weight:400}.fa.fa-bar-chart:before{content:""}.fa.fa-bar-chart-o{font-family:"Font Awesome 5 Free";font-weight:400}.fa.fa-bar-chart-o:before{content:""}.fa.fa-twitter-square{font-family:"Font Awesome 5 Brands";font-weight:400}.fa.fa-facebook-square{font-family:"Font Awesome 5 Brands";font-weight:400}.fa.fa-gears:before{content:""}.fa.fa-thumbs-o-up{font-family:"Font Awesome 5 Free";font-weight:400}.fa.fa-thumbs-o-up:before{content:""}.fa.fa-thumbs-o-down{font-family:"Font Awesome 5 Free";font-weight:400}.fa.fa-thumbs-o-down:before{content:""}.fa.fa-heart-o{font-family:"Font Awesome 5 Free";font-weight:400}.fa.fa-heart-o:before{content:""}.fa.fa-sign-out:before{content:""}.fa.fa-linkedin-square{font-family:"Font Awesome 5 Brands";font-weight:400}.fa.fa-linkedin-square:before{content:""}.fa.fa-thumb-tack:before{content:""}.fa.fa-external-link:before{content:""}.fa.fa-sign-in:before{content:""}.fa.fa-github-square{font-family:"Font Awesome 5 Brands";font-weight:400}.fa.fa-lemon-o{font-family:"Font Awesome 5 Free";font-weight:400}.fa.fa-lemon-o:before{content:""}.fa.fa-square-o{font-family:"Font Awesome 5 Free";font-weight:400}.fa.fa-square-o:before{content:""}.fa.fa-bookmark-o{font-family:"Font Awesome 5 Free";font-weight:400}.fa.fa-bookmark-o:before{content:""}.fa.fa-twitter{font-family:"Font Awesome 5 Brands";font-weight:400}.fa.fa-facebook{font-family:"Font Awesome 5 Brands";font-weight:400}.fa.fa-facebook:before{content:""}.fa.fa-facebook-f{font-family:"Font Awesome 5 Brands";font-weight:400}.fa.fa-facebook-f:before{content:""}.fa.fa-github{font-family:"Font Awesome 5 Brands";font-weight:400}.fa.fa-credit-card{font-family:"Font Awesome 5 Free";font-weight:400}.fa.fa-feed:before{content:""}.fa.fa-hdd-o{font-family:"Font Awesome 5 Free";font-weight:400}.fa.fa-hdd-o:before{content:""}.fa.fa-hand-o-right{font-family:"Font Awesome 5 Free";font-weight:400}.fa.fa-hand-o-right:before{content:""}.fa.fa-hand-o-left{font-family:"Font Awesome 5 Free";font-weight:400}.fa.fa-hand-o-left:before{content:""}.fa.fa-hand-o-up{font-family:"Font Awesome 5 Free";font-weight:400}.fa.fa-hand-o-up:before{content:""}.fa.fa-hand-o-down{font-family:"Font Awesome 5 Free";font-weight:400}.fa.fa-hand-o-down:before{content:""}.fa.fa-arrows-alt:before{content:""}.fa.fa-group:before{content:""}.fa.fa-chain:before{content:""}.fa.fa-scissors:before{content:""}.fa.fa-files-o{font-family:"Font Awesome 5 Free";font-weight:400}.fa.fa-files-o:before{content:""}.fa.fa-floppy-o{font-family:"Font Awesome 5 Free";font-weight:400}.fa.fa-floppy-o:before{content:""}.fa.fa-navicon:before{content:""}.fa.fa-reorder:before{content:""}.fa.fa-pinterest{font-family:"Font Awesome 5 Brands";font-weight:400}.fa.fa-pinterest-square{font-family:"Font Awesome 5 Brands";font-weight:400}.fa.fa-google-plus-square{font-family:"Font Awesome 5 Brands";font-weight:400}.fa.fa-google-plus{font-family:"Font Awesome 5 Brands";font-weight:400}.fa.fa-google-plus:before{content:""}.fa.fa-money{font-family:"Font Awesome 5 Free";font-weight:400}.fa.fa-money:before{content:""}.fa.fa-unsorted:before{content:""}.fa.fa-sort-desc:before{content:""}.fa.fa-sort-asc:before{content:""}.fa.fa-linkedin{font-family:"Font Awesome 5 Brands";font-weight:400}.fa.fa-linkedin:before{content:""}.fa.fa-rotate-left:before{content:""}.fa.fa-legal:before{content:""}.fa.fa-tachometer:before{content:""}.fa.fa-dashboard:before{content:""}.fa.fa-comment-o{font-family:"Font Awesome 5 Free";font-weight:400}.fa.fa-comment-o:before{content:""}.fa.fa-comments-o{font-family:"Font Awesome 5 Free";font-weight:400}.fa.fa-comments-o:before{content:""}.fa.fa-flash:before{content:""}.fa.fa-clipboard{font-family:"Font Awesome 5 Free";font-weight:400}.fa.fa-paste{font-family:"Font Awesome 5 Free";font-weight:400}.fa.fa-paste:before{content:""}.fa.fa-lightbulb-o{font-family:"Font Awesome 5 Free";font-weight:400}.fa.fa-lightbulb-o:before{content:""}.fa.fa-exchange:before{content:""}.fa.fa-cloud-download:before{content:""}.fa.fa-cloud-upload:before{content:""}.fa.fa-bell-o{font-family:"Font Awesome 5 Free";font-weight:400}.fa.fa-bell-o:before{content:""}.fa.fa-cutlery:before{content:""}.fa.fa-file-text-o{font-family:"Font Awesome 5 Free";font-weight:400}.fa.fa-file-text-o:before{content:""}.fa.fa-building-o{font-family:"Font Awesome 5 Free";font-weight:400}.fa.fa-building-o:before{content:""}.fa.fa-hospital-o{font-family:"Font Awesome 5 Free";font-weight:400}.fa.fa-hospital-o:before{content:""}.fa.fa-tablet:before{content:""}.fa.fa-mobile:before{content:""}.fa.fa-mobile-phone:before{content:""}.fa.fa-circle-o{font-family:"Font Awesome 5 Free";font-weight:400}.fa.fa-circle-o:before{content:""}.fa.fa-mail-reply:before{content:""}.fa.fa-github-alt{font-family:"Font Awesome 5 Brands";font-weight:400}.fa.fa-folder-o{font-family:"Font Awesome 5 Free";font-weight:400}.fa.fa-folder-o:before{content:""}.fa.fa-folder-open-o{font-family:"Font Awesome 5 Free";font-weight:400}.fa.fa-folder-open-o:before{content:""}.fa.fa-smile-o{font-family:"Font Awesome 5 Free";font-weight:400}.fa.fa-smile-o:before{content:""}.fa.fa-frown-o{font-family:"Font Awesome 5 Free";font-weight:400}.fa.fa-frown-o:before{content:""}.fa.fa-meh-o{font-family:"Font Awesome 5 Free";font-weight:400}.fa.fa-meh-o:before{content:""}.fa.fa-keyboard-o{font-family:"Font Awesome 5 Free";font-weight:400}.fa.fa-keyboard-o:before{content:""}.fa.fa-flag-o{font-family:"Font Awesome 5 Free";font-weight:400}.fa.fa-flag-o:before{content:""}.fa.fa-mail-reply-all:before{content:""}.fa.fa-star-half-o{font-family:"Font Awesome 5 Free";font-weight:400}.fa.fa-star-half-o:before{content:""}.fa.fa-star-half-empty{font-family:"Font Awesome 5 Free";font-weight:400}.fa.fa-star-half-empty:before{content:""}.fa.fa-star-half-full{font-family:"Font Awesome 5 Free";font-weight:400}.fa.fa-star-half-full:before{content:""}.fa.fa-code-fork:before{content:""}.fa.fa-chain-broken:before{content:""}.fa.fa-shield:before{content:""}.fa.fa-calendar-o{font-family:"Font Awesome 5 Free";font-weight:400}.fa.fa-calendar-o:before{content:""}.fa.fa-maxcdn{font-family:"Font Awesome 5 Brands";font-weight:400}.fa.fa-html5{font-family:"Font Awesome 5 Brands";font-weight:400}.fa.fa-css3{font-family:"Font Awesome 5 Brands";font-weight:400}.fa.fa-ticket:before{content:""}.fa.fa-minus-square-o{font-family:"Font Awesome 5 Free";font-weight:400}.fa.fa-minus-square-o:before{content:""}.fa.fa-level-up:before{content:""}.fa.fa-level-down:before{content:""}.fa.fa-pencil-square:before{content:""}.fa.fa-external-link-square:before{content:""}.fa.fa-compass{font-family:"Font Awesome 5 Free";font-weight:400}.fa.fa-caret-square-o-down{font-family:"Font Awesome 5 Free";font-weight:400}.fa.fa-caret-square-o-down:before{content:""}.fa.fa-toggle-down{font-family:"Font Awesome 5 Free";font-weight:400}.fa.fa-toggle-down:before{content:""}.fa.fa-caret-square-o-up{font-family:"Font Awesome 5 Free";font-weight:400}.fa.fa-caret-square-o-up:before{content:""}.fa.fa-toggle-up{font-family:"Font Awesome 5 Free";font-weight:400}.fa.fa-toggle-up:before{content:""}.fa.fa-caret-square-o-right{font-family:"Font Awesome 5 Free";font-weight:400}.fa.fa-caret-square-o-right:before{content:""}.fa.fa-toggle-right{font-family:"Font Awesome 5 Free";font-weight:400}.fa.fa-toggle-right:before{content:""}.fa.fa-eur:before{content:""}.fa.fa-euro:before{content:""}.fa.fa-gbp:before{content:""}.fa.fa-usd:before{content:""}.fa.fa-dollar:before{content:""}.fa.fa-inr:before{content:""}.fa.fa-rupee:before{content:""}.fa.fa-jpy:before{content:""}.fa.fa-cny:before{content:""}.fa.fa-rmb:before{content:""}.fa.fa-yen:before{content:""}.fa.fa-rub:before{content:""}.fa.fa-ruble:before{content:""}.fa.fa-rouble:before{content:""}.fa.fa-krw:before{content:""}.fa.fa-won:before{content:""}.fa.fa-btc{font-family:"Font Awesome 5 Brands";font-weight:400}.fa.fa-bitcoin{font-family:"Font Awesome 5 Brands";font-weight:400}.fa.fa-bitcoin:before{content:""}.fa.fa-file-text:before{content:""}.fa.fa-sort-alpha-asc:before{content:""}.fa.fa-sort-alpha-desc:before{content:""}.fa.fa-sort-amount-asc:before{content:""}.fa.fa-sort-amount-desc:before{content:""}.fa.fa-sort-numeric-asc:before{content:""}.fa.fa-sort-numeric-desc:before{content:""}.fa.fa-youtube-square{font-family:"Font Awesome 5 Brands";font-weight:400}.fa.fa-youtube{font-family:"Font Awesome 5 Brands";font-weight:400}.fa.fa-xing{font-family:"Font Awesome 5 Brands";font-weight:400}.fa.fa-xing-square{font-family:"Font Awesome 5 Brands";font-weight:400}.fa.fa-youtube-play{font-family:"Font Awesome 5 Brands";font-weight:400}.fa.fa-youtube-play:before{content:""}.fa.fa-dropbox{font-family:"Font Awesome 5 Brands";font-weight:400}.fa.fa-stack-overflow{font-family:"Font Awesome 5 Brands";font-weight:400}.fa.fa-instagram{font-family:"Font Awesome 5 Brands";font-weight:400}.fa.fa-flickr{font-family:"Font Awesome 5 Brands";font-weight:400}.fa.fa-adn{font-family:"Font Awesome 5 Brands";font-weight:400}.fa.fa-bitbucket{font-family:"Font Awesome 5 Brands";font-weight:400}.fa.fa-bitbucket-square{font-family:"Font Awesome 5 Brands";font-weight:400}.fa.fa-bitbucket-square:before{content:""}.fa.fa-tumblr{font-family:"Font Awesome 5 Brands";font-weight:400}.fa.fa-tumblr-square{font-family:"Font Awesome 5 Brands";font-weight:400}.fa.fa-long-arrow-down:before{content:""}.fa.fa-long-arrow-up:before{content:""}.fa.fa-long-arrow-left:before{content:""}.fa.fa-long-arrow-right:before{content:""}.fa.fa-apple{font-family:"Font Awesome 5 Brands";font-weight:400}.fa.fa-windows{font-family:"Font Awesome 5 Brands";font-weight:400}.fa.fa-android{font-family:"Font Awesome 5 Brands";font-weight:400}.fa.fa-linux{font-family:"Font Awesome 5 Brands";font-weight:400}.fa.fa-dribbble{font-family:"Font Awesome 5 Brands";font-weight:400}.fa.fa-skype{font-family:"Font Awesome 5 Brands";font-weight:400}.fa.fa-foursquare{font-family:"Font Awesome 5 Brands";font-weight:400}.fa.fa-trello{font-family:"Font Awesome 5 Brands";font-weight:400}.fa.fa-gratipay{font-family:"Font Awesome 5 Brands";font-weight:400}.fa.fa-gittip{font-family:"Font Awesome 5 Brands";font-weight:400}.fa.fa-gittip:before{content:""}.fa.fa-sun-o{font-family:"Font Awesome 5 Free";font-weight:400}.fa.fa-sun-o:before{content:""}.fa.fa-moon-o{font-family:"Font Awesome 5 Free";font-weight:400}.fa.fa-moon-o:before{content:""}.fa.fa-vk{font-family:"Font Awesome 5 Brands";font-weight:400}.fa.fa-weibo{font-family:"Font Awesome 5 Brands";font-weight:400}.fa.fa-renren{font-family:"Font Awesome 5 Brands";font-weight:400}.fa.fa-pagelines{font-family:"Font Awesome 5 Brands";font-weight:400}.fa.fa-stack-exchange{font-family:"Font Awesome 5 Brands";font-weight:400}.fa.fa-arrow-circle-o-right{font-family:"Font Awesome 5 Free";font-weight:400}.fa.fa-arrow-circle-o-right:before{content:""}.fa.fa-arrow-circle-o-left{font-family:"Font Awesome 5 Free";font-weight:400}.fa.fa-arrow-circle-o-left:before{content:""}.fa.fa-caret-square-o-left{font-family:"Font Awesome 5 Free";font-weight:400}.fa.fa-caret-square-o-left:before{content:""}.fa.fa-toggle-left{font-family:"Font Awesome 5 Free";font-weight:400}.fa.fa-toggle-left:before{content:""}.fa.fa-dot-circle-o{font-family:"Font Awesome 5 Free";font-weight:400}.fa.fa-dot-circle-o:before{content:""}.fa.fa-vimeo-square{font-family:"Font Awesome 5 Brands";font-weight:400}.fa.fa-try:before{content:""}.fa.fa-turkish-lira:before{content:""}.fa.fa-plus-square-o{font-family:"Font Awesome 5 Free";font-weight:400}.fa.fa-plus-square-o:before{content:""}.fa.fa-slack{font-family:"Font Awesome 5 Brands";font-weight:400}.fa.fa-wordpress{font-family:"Font Awesome 5 Brands";font-weight:400}.fa.fa-openid{font-family:"Font Awesome 5 Brands";font-weight:400}.fa.fa-institution:before{content:""}.fa.fa-bank:before{content:""}.fa.fa-mortar-board:before{content:""}.fa.fa-yahoo{font-family:"Font Awesome 5 Brands";font-weight:400}.fa.fa-google{font-family:"Font Awesome 5 Brands";font-weight:400}.fa.fa-reddit{font-family:"Font Awesome 5 Brands";font-weight:400}.fa.fa-reddit-square{font-family:"Font Awesome 5 Brands";font-weight:400}.fa.fa-stumbleupon-circle{font-family:"Font Awesome 5 Brands";font-weight:400}.fa.fa-stumbleupon{font-family:"Font Awesome 5 Brands";font-weight:400}.fa.fa-delicious{font-family:"Font Awesome 5 Brands";font-weight:400}.fa.fa-digg{font-family:"Font Awesome 5 Brands";font-weight:400}.fa.fa-pied-piper-pp{font-family:"Font Awesome 5 Brands";font-weight:400}.fa.fa-pied-piper-alt{font-family:"Font Awesome 5 Brands";font-weight:400}.fa.fa-drupal{font-family:"Font Awesome 5 Brands";font-weight:400}.fa.fa-joomla{font-family:"Font Awesome 5 Brands";font-weight:400}.fa.fa-spoon:before{content:""}.fa.fa-behance{font-family:"Font Awesome 5 Brands";font-weight:400}.fa.fa-behance-square{font-family:"Font Awesome 5 Brands";font-weight:400}.fa.fa-steam{font-family:"Font Awesome 5 Brands";font-weight:400}.fa.fa-steam-square{font-family:"Font Awesome 5 Brands";font-weight:400}.fa.fa-automobile:before{content:""}.fa.fa-envelope-o{font-family:"Font Awesome 5 Free";font-weight:400}.fa.fa-envelope-o:before{content:""}.fa.fa-spotify{font-family:"Font Awesome 5 Brands";font-weight:400}.fa.fa-deviantart{font-family:"Font Awesome 5 Brands";font-weight:400}.fa.fa-soundcloud{font-family:"Font Awesome 5 Brands";font-weight:400}.fa.fa-file-pdf-o{font-family:"Font Awesome 5 Free";font-weight:400}.fa.fa-file-pdf-o:before{content:""}.fa.fa-file-word-o{font-family:"Font Awesome 5 Free";font-weight:400}.fa.fa-file-word-o:before{content:""}.fa.fa-file-excel-o{font-family:"Font Awesome 5 Free";font-weight:400}.fa.fa-file-excel-o:before{content:""}.fa.fa-file-powerpoint-o{font-family:"Font Awesome 5 Free";font-weight:400}.fa.fa-file-powerpoint-o:before{content:""}.fa.fa-file-image-o{font-family:"Font Awesome 5 Free";font-weight:400}.fa.fa-file-image-o:before{content:""}.fa.fa-file-photo-o{font-family:"Font Awesome 5 Free";font-weight:400}.fa.fa-file-photo-o:before{content:""}.fa.fa-file-picture-o{font-family:"Font Awesome 5 Free";font-weight:400}.fa.fa-file-picture-o:before{content:""}.fa.fa-file-archive-o{font-family:"Font Awesome 5 Free";font-weight:400}.fa.fa-file-archive-o:before{content:""}.fa.fa-file-zip-o{font-family:"Font Awesome 5 Free";font-weight:400}.fa.fa-file-zip-o:before{content:""}.fa.fa-file-audio-o{font-family:"Font Awesome 5 Free";font-weight:400}.fa.fa-file-audio-o:before{content:""}.fa.fa-file-sound-o{font-family:"Font Awesome 5 Free";font-weight:400}.fa.fa-file-sound-o:before{content:""}.fa.fa-file-video-o{font-family:"Font Awesome 5 Free";font-weight:400}.fa.fa-file-video-o:before{content:""}.fa.fa-file-movie-o{font-family:"Font Awesome 5 Free";font-weight:400}.fa.fa-file-movie-o:before{content:""}.fa.fa-file-code-o{font-family:"Font Awesome 5 Free";font-weight:400}.fa.fa-file-code-o:before{content:""}.fa.fa-vine{font-family:"Font Awesome 5 Brands";font-weight:400}.fa.fa-codepen{font-family:"Font Awesome 5 Brands";font-weight:400}.fa.fa-jsfiddle{font-family:"Font Awesome 5 Brands";font-weight:400}.fa.fa-life-ring{font-family:"Font Awesome 5 Free";font-weight:400}.fa.fa-life-bouy{font-family:"Font Awesome 5 Free";font-weight:400}.fa.fa-life-bouy:before{content:""}.fa.fa-life-buoy{font-family:"Font Awesome 5 Free";font-weight:400}.fa.fa-life-buoy:before{content:""}.fa.fa-life-saver{font-family:"Font Awesome 5 Free";font-weight:400}.fa.fa-life-saver:before{content:""}.fa.fa-support{font-family:"Font Awesome 5 Free";font-weight:400}.fa.fa-support:before{content:""}.fa.fa-circle-o-notch:before{content:""}.fa.fa-rebel{font-family:"Font Awesome 5 Brands";font-weight:400}.fa.fa-ra{font-family:"Font Awesome 5 Brands";font-weight:400}.fa.fa-ra:before{content:""}.fa.fa-resistance{font-family:"Font Awesome 5 Brands";font-weight:400}.fa.fa-resistance:before{content:""}.fa.fa-empire{font-family:"Font Awesome 5 Brands";font-weight:400}.fa.fa-ge{font-family:"Font Awesome 5 Brands";font-weight:400}.fa.fa-ge:before{content:""}.fa.fa-git-square{font-family:"Font Awesome 5 Brands";font-weight:400}.fa.fa-git{font-family:"Font Awesome 5 Brands";font-weight:400}.fa.fa-hacker-news{font-family:"Font Awesome 5 Brands";font-weight:400}.fa.fa-y-combinator-square{font-family:"Font Awesome 5 Brands";font-weight:400}.fa.fa-y-combinator-square:before{content:""}.fa.fa-yc-square{font-family:"Font Awesome 5 Brands";font-weight:400}.fa.fa-yc-square:before{content:""}.fa.fa-tencent-weibo{font-family:"Font Awesome 5 Brands";font-weight:400}.fa.fa-qq{font-family:"Font Awesome 5 Brands";font-weight:400}.fa.fa-weixin{font-family:"Font Awesome 5 Brands";font-weight:400}.fa.fa-wechat{font-family:"Font Awesome 5 Brands";font-weight:400}.fa.fa-wechat:before{content:""}.fa.fa-send:before{content:""}.fa.fa-paper-plane-o{font-family:"Font Awesome 5 Free";font-weight:400}.fa.fa-paper-plane-o:before{content:""}.fa.fa-send-o{font-family:"Font Awesome 5 Free";font-weight:400}.fa.fa-send-o:before{content:""}.fa.fa-circle-thin{font-family:"Font Awesome 5 Free";font-weight:400}.fa.fa-circle-thin:before{content:""}.fa.fa-header:before{content:""}.fa.fa-sliders:before{content:""}.fa.fa-futbol-o{font-family:"Font Awesome 5 Free";font-weight:400}.fa.fa-futbol-o:before{content:""}.fa.fa-soccer-ball-o{font-family:"Font Awesome 5 Free";font-weight:400}.fa.fa-soccer-ball-o:before{content:""}.fa.fa-slideshare{font-family:"Font Awesome 5 Brands";font-weight:400}.fa.fa-twitch{font-family:"Font Awesome 5 Brands";font-weight:400}.fa.fa-yelp{font-family:"Font Awesome 5 Brands";font-weight:400}.fa.fa-newspaper-o{font-family:"Font Awesome 5 Free";font-weight:400}.fa.fa-newspaper-o:before{content:""}.fa.fa-paypal{font-family:"Font Awesome 5 Brands";font-weight:400}.fa.fa-google-wallet{font-family:"Font Awesome 5 Brands";font-weight:400}.fa.fa-cc-visa{font-family:"Font Awesome 5 Brands";font-weight:400}.fa.fa-cc-mastercard{font-family:"Font Awesome 5 Brands";font-weight:400}.fa.fa-cc-discover{font-family:"Font Awesome 5 Brands";font-weight:400}.fa.fa-cc-amex{font-family:"Font Awesome 5 Brands";font-weight:400}.fa.fa-cc-paypal{font-family:"Font Awesome 5 Brands";font-weight:400}.fa.fa-cc-stripe{font-family:"Font Awesome 5 Brands";font-weight:400}.fa.fa-bell-slash-o{font-family:"Font Awesome 5 Free";font-weight:400}.fa.fa-bell-slash-o:before{content:""}.fa.fa-trash:before{content:""}.fa.fa-copyright{font-family:"Font Awesome 5 Free";font-weight:400}.fa.fa-eyedropper:before{content:""}.fa.fa-area-chart:before{content:""}.fa.fa-pie-chart:before{content:""}.fa.fa-line-chart:before{content:""}.fa.fa-lastfm{font-family:"Font Awesome 5 Brands";font-weight:400}.fa.fa-lastfm-square{font-family:"Font Awesome 5 Brands";font-weight:400}.fa.fa-ioxhost{font-family:"Font Awesome 5 Brands";font-weight:400}.fa.fa-angellist{font-family:"Font Awesome 5 Brands";font-weight:400}.fa.fa-cc{font-family:"Font Awesome 5 Free";font-weight:400}.fa.fa-cc:before{content:""}.fa.fa-ils:before{content:""}.fa.fa-shekel:before{content:""}.fa.fa-sheqel:before{content:""}.fa.fa-meanpath{font-family:"Font Awesome 5 Brands";font-weight:400}.fa.fa-meanpath:before{content:""}.fa.fa-buysellads{font-family:"Font Awesome 5 Brands";font-weight:400}.fa.fa-connectdevelop{font-family:"Font Awesome 5 Brands";font-weight:400}.fa.fa-dashcube{font-family:"Font Awesome 5 Brands";font-weight:400}.fa.fa-forumbee{font-family:"Font Awesome 5 Brands";font-weight:400}.fa.fa-leanpub{font-family:"Font Awesome 5 Brands";font-weight:400}.fa.fa-sellsy{font-family:"Font Awesome 5 Brands";font-weight:400}.fa.fa-shirtsinbulk{font-family:"Font Awesome 5 Brands";font-weight:400}.fa.fa-simplybuilt{font-family:"Font Awesome 5 Brands";font-weight:400}.fa.fa-skyatlas{font-family:"Font Awesome 5 Brands";font-weight:400}.fa.fa-diamond{font-family:"Font Awesome 5 Free";font-weight:400}.fa.fa-diamond:before{content:""}.fa.fa-intersex:before{content:""}.fa.fa-facebook-official{font-family:"Font Awesome 5 Brands";font-weight:400}.fa.fa-facebook-official:before{content:""}.fa.fa-pinterest-p{font-family:"Font Awesome 5 Brands";font-weight:400}.fa.fa-whatsapp{font-family:"Font Awesome 5 Brands";font-weight:400}.fa.fa-hotel:before{content:""}.fa.fa-viacoin{font-family:"Font Awesome 5 Brands";font-weight:400}.fa.fa-medium{font-family:"Font Awesome 5 Brands";font-weight:400}.fa.fa-y-combinator{font-family:"Font Awesome 5 Brands";font-weight:400}.fa.fa-yc{font-family:"Font Awesome 5 Brands";font-weight:400}.fa.fa-yc:before{content:""}.fa.fa-optin-monster{font-family:"Font Awesome 5 Brands";font-weight:400}.fa.fa-opencart{font-family:"Font Awesome 5 Brands";font-weight:400}.fa.fa-expeditedssl{font-family:"Font Awesome 5 Brands";font-weight:400}.fa.fa-battery-4:before{content:""}.fa.fa-battery:before{content:""}.fa.fa-battery-3:before{content:""}.fa.fa-battery-2:before{content:""}.fa.fa-battery-1:before{content:""}.fa.fa-battery-0:before{content:""}.fa.fa-object-group{font-family:"Font Awesome 5 Free";font-weight:400}.fa.fa-object-ungroup{font-family:"Font Awesome 5 Free";font-weight:400}.fa.fa-sticky-note-o{font-family:"Font Awesome 5 Free";font-weight:400}.fa.fa-sticky-note-o:before{content:""}.fa.fa-cc-jcb{font-family:"Font Awesome 5 Brands";font-weight:400}.fa.fa-cc-diners-club{font-family:"Font Awesome 5 Brands";font-weight:400}.fa.fa-clone{font-family:"Font Awesome 5 Free";font-weight:400}.fa.fa-hourglass-o{font-family:"Font Awesome 5 Free";font-weight:400}.fa.fa-hourglass-o:before{content:""}.fa.fa-hourglass-1:before{content:""}.fa.fa-hourglass-2:before{content:""}.fa.fa-hourglass-3:before{content:""}.fa.fa-hand-rock-o{font-family:"Font Awesome 5 Free";font-weight:400}.fa.fa-hand-rock-o:before{content:""}.fa.fa-hand-grab-o{font-family:"Font Awesome 5 Free";font-weight:400}.fa.fa-hand-grab-o:before{content:""}.fa.fa-hand-paper-o{font-family:"Font Awesome 5 Free";font-weight:400}.fa.fa-hand-paper-o:before{content:""}.fa.fa-hand-stop-o{font-family:"Font Awesome 5 Free";font-weight:400}.fa.fa-hand-stop-o:before{content:""}.fa.fa-hand-scissors-o{font-family:"Font Awesome 5 Free";font-weight:400}.fa.fa-hand-scissors-o:before{content:""}.fa.fa-hand-lizard-o{font-family:"Font Awesome 5 Free";font-weight:400}.fa.fa-hand-lizard-o:before{content:""}.fa.fa-hand-spock-o{font-family:"Font Awesome 5 Free";font-weight:400}.fa.fa-hand-spock-o:before{content:""}.fa.fa-hand-pointer-o{font-family:"Font Awesome 5 Free";font-weight:400}.fa.fa-hand-pointer-o:before{content:""}.fa.fa-hand-peace-o{font-family:"Font Awesome 5 Free";font-weight:400}.fa.fa-hand-peace-o:before{content:""}.fa.fa-registered{font-family:"Font Awesome 5 Free";font-weight:400}.fa.fa-creative-commons{font-family:"Font Awesome 5 Brands";font-weight:400}.fa.fa-gg{font-family:"Font Awesome 5 Brands";font-weight:400}.fa.fa-gg-circle{font-family:"Font Awesome 5 Brands";font-weight:400}.fa.fa-tripadvisor{font-family:"Font Awesome 5 Brands";font-weight:400}.fa.fa-odnoklassniki{font-family:"Font Awesome 5 Brands";font-weight:400}.fa.fa-odnoklassniki-square{font-family:"Font Awesome 5 Brands";font-weight:400}.fa.fa-get-pocket{font-family:"Font Awesome 5 Brands";font-weight:400}.fa.fa-wikipedia-w{font-family:"Font Awesome 5 Brands";font-weight:400}.fa.fa-safari{font-family:"Font Awesome 5 Brands";font-weight:400}.fa.fa-chrome{font-family:"Font Awesome 5 Brands";font-weight:400}.fa.fa-firefox{font-family:"Font Awesome 5 Brands";font-weight:400}.fa.fa-opera{font-family:"Font Awesome 5 Brands";font-weight:400}.fa.fa-internet-explorer{font-family:"Font Awesome 5 Brands";font-weight:400}.fa.fa-television:before{content:""}.fa.fa-contao{font-family:"Font Awesome 5 Brands";font-weight:400}.fa.fa-500px{font-family:"Font Awesome 5 Brands";font-weight:400}.fa.fa-amazon{font-family:"Font Awesome 5 Brands";font-weight:400}.fa.fa-calendar-plus-o{font-family:"Font Awesome 5 Free";font-weight:400}.fa.fa-calendar-plus-o:before{content:""}.fa.fa-calendar-minus-o{font-family:"Font Awesome 5 Free";font-weight:400}.fa.fa-calendar-minus-o:before{content:""}.fa.fa-calendar-times-o{font-family:"Font Awesome 5 Free";font-weight:400}.fa.fa-calendar-times-o:before{content:""}.fa.fa-calendar-check-o{font-family:"Font Awesome 5 Free";font-weight:400}.fa.fa-calendar-check-o:before{content:""}.fa.fa-map-o{font-family:"Font Awesome 5 Free";font-weight:400}.fa.fa-map-o:before{content:""}.fa.fa-commenting:before{content:""}.fa.fa-commenting-o{font-family:"Font Awesome 5 Free";font-weight:400}.fa.fa-commenting-o:before{content:""}.fa.fa-houzz{font-family:"Font Awesome 5 Brands";font-weight:400}.fa.fa-vimeo{font-family:"Font Awesome 5 Brands";font-weight:400}.fa.fa-vimeo:before{content:""}.fa.fa-black-tie{font-family:"Font Awesome 5 Brands";font-weight:400}.fa.fa-fonticons{font-family:"Font Awesome 5 Brands";font-weight:400}.fa.fa-reddit-alien{font-family:"Font Awesome 5 Brands";font-weight:400}.fa.fa-edge{font-family:"Font Awesome 5 Brands";font-weight:400}.fa.fa-credit-card-alt:before{content:""}.fa.fa-codiepie{font-family:"Font Awesome 5 Brands";font-weight:400}.fa.fa-modx{font-family:"Font Awesome 5 Brands";font-weight:400}.fa.fa-fort-awesome{font-family:"Font Awesome 5 Brands";font-weight:400}.fa.fa-usb{font-family:"Font Awesome 5 Brands";font-weight:400}.fa.fa-product-hunt{font-family:"Font Awesome 5 Brands";font-weight:400}.fa.fa-mixcloud{font-family:"Font Awesome 5 Brands";font-weight:400}.fa.fa-scribd{font-family:"Font Awesome 5 Brands";font-weight:400}.fa.fa-pause-circle-o{font-family:"Font Awesome 5 Free";font-weight:400}.fa.fa-pause-circle-o:before{content:""}.fa.fa-stop-circle-o{font-family:"Font Awesome 5 Free";font-weight:400}.fa.fa-stop-circle-o:before{content:""}.fa.fa-bluetooth{font-family:"Font Awesome 5 Brands";font-weight:400}.fa.fa-bluetooth-b{font-family:"Font Awesome 5 Brands";font-weight:400}.fa.fa-gitlab{font-family:"Font Awesome 5 Brands";font-weight:400}.fa.fa-wpbeginner{font-family:"Font Awesome 5 Brands";font-weight:400}.fa.fa-wpforms{font-family:"Font Awesome 5 Brands";font-weight:400}.fa.fa-envira{font-family:"Font Awesome 5 Brands";font-weight:400}.fa.fa-wheelchair-alt{font-family:"Font Awesome 5 Brands";font-weight:400}.fa.fa-wheelchair-alt:before{content:""}.fa.fa-question-circle-o{font-family:"Font Awesome 5 Free";font-weight:400}.fa.fa-question-circle-o:before{content:""}.fa.fa-volume-control-phone:before{content:""}.fa.fa-asl-interpreting:before{content:""}.fa.fa-deafness:before{content:""}.fa.fa-hard-of-hearing:before{content:""}.fa.fa-glide{font-family:"Font Awesome 5 Brands";font-weight:400}.fa.fa-glide-g{font-family:"Font Awesome 5 Brands";font-weight:400}.fa.fa-signing:before{content:""}.fa.fa-viadeo{font-family:"Font Awesome 5 Brands";font-weight:400}.fa.fa-viadeo-square{font-family:"Font Awesome 5 Brands";font-weight:400}.fa.fa-snapchat{font-family:"Font Awesome 5 Brands";font-weight:400}.fa.fa-snapchat-ghost{font-family:"Font Awesome 5 Brands";font-weight:400}.fa.fa-snapchat-square{font-family:"Font Awesome 5 Brands";font-weight:400}.fa.fa-pied-piper{font-family:"Font Awesome 5 Brands";font-weight:400}.fa.fa-first-order{font-family:"Font Awesome 5 Brands";font-weight:400}.fa.fa-yoast{font-family:"Font Awesome 5 Brands";font-weight:400}.fa.fa-themeisle{font-family:"Font Awesome 5 Brands";font-weight:400}.fa.fa-google-plus-official{font-family:"Font Awesome 5 Brands";font-weight:400}.fa.fa-google-plus-official:before{content:""}.fa.fa-google-plus-circle{font-family:"Font Awesome 5 Brands";font-weight:400}.fa.fa-google-plus-circle:before{content:""}.fa.fa-font-awesome{font-family:"Font Awesome 5 Brands";font-weight:400}.fa.fa-fa{font-family:"Font Awesome 5 Brands";font-weight:400}.fa.fa-fa:before{content:""}.fa.fa-handshake-o{font-family:"Font Awesome 5 Free";font-weight:400}.fa.fa-handshake-o:before{content:""}.fa.fa-envelope-open-o{font-family:"Font Awesome 5 Free";font-weight:400}.fa.fa-envelope-open-o:before{content:""}.fa.fa-linode{font-family:"Font Awesome 5 Brands";font-weight:400}.fa.fa-address-book-o{font-family:"Font Awesome 5 Free";font-weight:400}.fa.fa-address-book-o:before{content:""}.fa.fa-vcard:before{content:""}.fa.fa-address-card-o{font-family:"Font Awesome 5 Free";font-weight:400}.fa.fa-address-card-o:before{content:""}.fa.fa-vcard-o{font-family:"Font Awesome 5 Free";font-weight:400}.fa.fa-vcard-o:before{content:""}.fa.fa-user-circle-o{font-family:"Font Awesome 5 Free";font-weight:400}.fa.fa-user-circle-o:before{content:""}.fa.fa-user-o{font-family:"Font Awesome 5 Free";font-weight:400}.fa.fa-user-o:before{content:""}.fa.fa-id-badge{font-family:"Font Awesome 5 Free";font-weight:400}.fa.fa-drivers-license:before{content:""}.fa.fa-id-card-o{font-family:"Font Awesome 5 Free";font-weight:400}.fa.fa-id-card-o:before{content:""}.fa.fa-drivers-license-o{font-family:"Font Awesome 5 Free";font-weight:400}.fa.fa-drivers-license-o:before{content:""}.fa.fa-quora{font-family:"Font Awesome 5 Brands";font-weight:400}.fa.fa-free-code-camp{font-family:"Font Awesome 5 Brands";font-weight:400}.fa.fa-telegram{font-family:"Font Awesome 5 Brands";font-weight:400}.fa.fa-thermometer-4:before{content:""}.fa.fa-thermometer:before{content:""}.fa.fa-thermometer-3:before{content:""}.fa.fa-thermometer-2:before{content:""}.fa.fa-thermometer-1:before{content:""}.fa.fa-thermometer-0:before{content:""}.fa.fa-bathtub:before{content:""}.fa.fa-s15:before{content:""}.fa.fa-window-maximize{font-family:"Font Awesome 5 Free";font-weight:400}.fa.fa-window-restore{font-family:"Font Awesome 5 Free";font-weight:400}.fa.fa-times-rectangle:before{content:""}.fa.fa-window-close-o{font-family:"Font Awesome 5 Free";font-weight:400}.fa.fa-window-close-o:before{content:""}.fa.fa-times-rectangle-o{font-family:"Font Awesome 5 Free";font-weight:400}.fa.fa-times-rectangle-o:before{content:""}.fa.fa-bandcamp{font-family:"Font Awesome 5 Brands";font-weight:400}.fa.fa-grav{font-family:"Font Awesome 5 Brands";font-weight:400}.fa.fa-etsy{font-family:"Font Awesome 5 Brands";font-weight:400}.fa.fa-imdb{font-family:"Font Awesome 5 Brands";font-weight:400}.fa.fa-ravelry{font-family:"Font Awesome 5 Brands";font-weight:400}.fa.fa-eercast{font-family:"Font Awesome 5 Brands";font-weight:400}.fa.fa-eercast:before{content:""}.fa.fa-snowflake-o{font-family:"Font Awesome 5 Free";font-weight:400}.fa.fa-snowflake-o:before{content:""}.fa.fa-superpowers{font-family:"Font Awesome 5 Brands";font-weight:400}.fa.fa-wpexplorer{font-family:"Font Awesome 5 Brands";font-weight:400}.fa.fa-cab:before{content:""}/*!\n * Bootstrap v4.4.1 (https://getbootstrap.com/)\n * Copyright 2011-2019 The Bootstrap Authors\n * Copyright 2011-2019 Twitter, Inc.\n * Licensed under MIT (https://github.com/twbs/bootstrap/blob/master/LICENSE)\n */:root{--blue: #007bff;--indigo: #6610f2;--purple: #6f42c1;--pink: #e83e8c;--red: #dc3545;--orange: #fd7e14;--yellow: #ffc107;--green: #28a745;--teal: #20c997;--cyan: #17a2b8;--white: #fff;--gray: #6c757d;--gray-dark: #343a40;--primary: #00a99d;--secondary: #6c757d;--success: #28a745;--info: #17a2b8;--warning: #ffc107;--danger: #dc3545;--light: #f8f9fa;--dark: #343a40;--breakpoint-xs: 0;--breakpoint-sm: 576px;--breakpoint-md: 768px;--breakpoint-lg: 992px;--breakpoint-xl: 1200px;--font-family-sans-serif: -apple-system, BlinkMacSystemFont, "Segoe UI", Roboto, "Helvetica Neue", Arial, "Noto Sans", "Liberation Sans", sans-serif, "Apple Color Emoji", "Segoe UI Emoji", "Segoe UI Symbol", "Noto Color Emoji";--font-family-monospace: SFMono-Regular, Menlo, Monaco, Consolas, "Liberation Mono", "Courier New", monospace}*,*::before,*::after{box-sizing:border-box}html{font-family:sans-serif;line-height:1.15;-webkit-text-size-adjust:100%;-webkit-tap-highlight-color:rgba(0,0,0,0)}article,aside,figcaption,figure,footer,header,hgroup,main,nav,section{display:block}body{margin:0;font-family:-apple-system,BlinkMacSystemFont,"Segoe UI",Roboto,"Helvetica Neue",Arial,"Noto Sans","Liberation Sans",sans-serif,"Apple Color Emoji","Segoe UI Emoji","Segoe UI Symbol","Noto Color Emoji";font-size:1rem;font-weight:400;line-height:1.5;color:#212529;text-align:left;background-color:#fff}[tabindex="-1"]:focus:not(:focus-visible){outline:0 !important}hr{box-sizing:content-box;height:0;overflow:visible}h1,h2,h3,h4,h5,h6{margin-top:0;margin-bottom:.5rem}p{margin-top:0;margin-bottom:1rem}abbr[title],abbr[data-original-title]{text-decoration:underline;text-decoration:underline dotted;cursor:help;border-bottom:0;text-decoration-skip-ink:none}address{margin-bottom:1rem;font-style:normal;line-height:inherit}ol,ul,dl{margin-top:0;margin-bottom:1rem}ol ol,ul ul,ol ul,ul ol{margin-bottom:0}dt{font-weight:700}dd{margin-bottom:.5rem;margin-left:0}blockquote{margin:0 0 1rem}b,strong{font-weight:bolder}small{font-size:80%}sub,sup{position:relative;font-size:75%;line-height:0;vertical-align:baseline}sub{bottom:-0.25em}sup{top:-0.5em}a{color:#00a99d;text-decoration:none;background-color:transparent}a:hover{color:#005d56;text-decoration:underline}a:not([href]){color:inherit;text-decoration:none}a:not([href]):hover{color:inherit;text-decoration:none}pre,code,kbd,samp{font-family:SFMono-Regular,Menlo,Monaco,Consolas,"Liberation Mono","Courier New",monospace;font-size:1em}pre{margin-top:0;margin-bottom:1rem;overflow:auto}figure{margin:0 0 1rem}img{vertical-align:middle;border-style:none}svg{overflow:hidden;vertical-align:middle}table{border-collapse:collapse}caption{padding-top:.75rem;padding-bottom:.75rem;color:#6c757d;text-align:left;caption-side:bottom}th{text-align:inherit}label{display:inline-block;margin-bottom:.5rem}button{border-radius:0}button:focus{outline:1px dotted;outline:5px auto -webkit-focus-ring-color}input,button,select,optgroup,textarea{margin:0;font-family:inherit;font-size:inherit;line-height:inherit}button,input{overflow:visible}button,select{text-transform:none}select{word-wrap:normal}button,[type=button],[type=reset],[type=submit]{-webkit-appearance:button}button:not(:disabled),[type=button]:not(:disabled),[type=reset]:not(:disabled),[type=submit]:not(:disabled){cursor:pointer}button::-moz-focus-inner,[type=button]::-moz-focus-inner,[type=reset]::-moz-focus-inner,[type=submit]::-moz-focus-inner{padding:0;border-style:none}input[type=radio],input[type=checkbox]{box-sizing:border-box;padding:0}input[type=date],input[type=time],input[type=datetime-local],input[type=month]{-webkit-appearance:listbox}textarea{overflow:auto;resize:vertical}fieldset{min-width:0;padding:0;margin:0;border:0}legend{display:block;width:100%;max-width:100%;padding:0;margin-bottom:.5rem;font-size:1.5rem;line-height:inherit;color:inherit;white-space:normal}progress{vertical-align:baseline}[type=number]::-webkit-inner-spin-button,[type=number]::-webkit-outer-spin-button{height:auto}[type=search]{outline-offset:-2px;-webkit-appearance:none}[type=search]::-webkit-search-decoration{-webkit-appearance:none}::-webkit-file-upload-button{font:inherit;-webkit-appearance:button}output{display:inline-block}summary{display:list-item;cursor:pointer}template{display:none}[hidden]{display:none !important}h1,h2,h3,h4,h5,h6,.h1,.h2,.h3,.h4,.h5,.h6{margin-bottom:.5rem;font-weight:500;line-height:1.2}h1,.h1{font-size:2.5rem}h2,.h2{font-size:2rem}h3,.h3{font-size:1.75rem}h4,.h4{font-size:1.5rem}h5,.h5{font-size:1.25rem}h6,.h6{font-size:1rem}.lead{font-size:1.25rem;font-weight:300}.display-1{font-size:6rem;font-weight:300;line-height:1.2}.display-2{font-size:5.5rem;font-weight:300;line-height:1.2}.display-3{font-size:4.5rem;font-weight:300;line-height:1.2}.display-4{font-size:3.5rem;font-weight:300;line-height:1.2}hr{margin-top:1rem;margin-bottom:1rem;border:0;border-top:1px solid rgba(0,0,0,.1)}small,.small{font-size:80%;font-weight:400}mark,.mark{padding:.2em;background-color:#fcf8e3}.list-unstyled{padding-left:0;list-style:none}.list-inline{padding-left:0;list-style:none}.list-inline-item{display:inline-block}.list-inline-item:not(:last-child){margin-right:.5rem}.initialism{font-size:90%;text-transform:uppercase}.blockquote{margin-bottom:1rem;font-size:1.25rem}.blockquote-footer{display:block;font-size:80%;color:#6c757d}.blockquote-footer::before{content:"— "}.img-fluid{max-width:100%;height:auto}.img-thumbnail{padding:.25rem;background-color:#fff;border:1px solid #dee2e6;border-radius:.25rem;max-width:100%;height:auto}.figure{display:inline-block}.figure-img{margin-bottom:.5rem;line-height:1}.figure-caption{font-size:90%;color:#6c757d}code{font-size:87.5%;color:#e83e8c;word-wrap:break-word}a>code{color:inherit}kbd{padding:.2rem .4rem;font-size:87.5%;color:#fff;background-color:#212529;border-radius:.2rem}kbd kbd{padding:0;font-size:100%;font-weight:700}pre{display:block;font-size:87.5%;color:#212529}pre code{font-size:inherit;color:inherit;word-break:normal}.pre-scrollable{max-height:340px;overflow-y:scroll}.container{width:100%;padding-right:15px;padding-left:15px;margin-right:auto;margin-left:auto}@media(min-width: 576px){.container{max-width:540px}}@media(min-width: 768px){.container{max-width:720px}}@media(min-width: 992px){.container{max-width:960px}}@media(min-width: 1200px){.container{max-width:1140px}}.container-fluid,.container-sm,.container-md,.container-lg,.container-xl{width:100%;padding-right:15px;padding-left:15px;margin-right:auto;margin-left:auto}@media(min-width: 576px){.container,.container-sm{max-width:540px}}@media(min-width: 768px){.container,.container-sm,.container-md{max-width:720px}}@media(min-width: 992px){.container,.container-sm,.container-md,.container-lg{max-width:960px}}@media(min-width: 1200px){.container,.container-sm,.container-md,.container-lg,.container-xl{max-width:1140px}}.row{display:flex;flex-wrap:wrap;margin-right:-15px;margin-left:-15px}.no-gutters{margin-right:0;margin-left:0}.no-gutters>.col,.no-gutters>[class*=col-]{padding-right:0;padding-left:0}.col-1,.col-2,.col-3,.col-4,.col-5,.col-6,.col-7,.col-8,.col-9,.col-10,.col-11,.col-12,.col,.col-auto,.col-sm-1,.col-sm-2,.col-sm-3,.col-sm-4,.col-sm-5,.col-sm-6,.col-sm-7,.col-sm-8,.col-sm-9,.col-sm-10,.col-sm-11,.col-sm-12,.col-sm,.col-sm-auto,.col-md-1,.col-md-2,.col-md-3,.col-md-4,.col-md-5,.col-md-6,.col-md-7,.col-md-8,.col-md-9,.col-md-10,.col-md-11,.col-md-12,.col-md,.col-md-auto,.col-lg-1,.col-lg-2,.col-lg-3,.col-lg-4,.col-lg-5,.col-lg-6,.col-lg-7,.col-lg-8,.col-lg-9,.col-lg-10,.col-lg-11,.col-lg-12,.col-lg,.col-lg-auto,.col-xl-1,.col-xl-2,.col-xl-3,.col-xl-4,.col-xl-5,.col-xl-6,.col-xl-7,.col-xl-8,.col-xl-9,.col-xl-10,.col-xl-11,.col-xl-12,.col-xl,.col-xl-auto{position:relative;width:100%;padding-right:15px;padding-left:15px}.col{flex-basis:0;flex-grow:1;max-width:100%}.row-cols-1>*{flex:0 0 100%;max-width:100%}.row-cols-2>*{flex:0 0 50%;max-width:50%}.row-cols-3>*{flex:0 0 33.3333333333%;max-width:33.3333333333%}.row-cols-4>*{flex:0 0 25%;max-width:25%}.row-cols-5>*{flex:0 0 20%;max-width:20%}.row-cols-6>*{flex:0 0 16.6666666667%;max-width:16.6666666667%}.col-auto{flex:0 0 auto;width:auto;max-width:100%}.col-1{flex:0 0 8.3333333333%;max-width:8.3333333333%}.col-2{flex:0 0 16.6666666667%;max-width:16.6666666667%}.col-3{flex:0 0 25%;max-width:25%}.col-4{flex:0 0 33.3333333333%;max-width:33.3333333333%}.col-5{flex:0 0 41.6666666667%;max-width:41.6666666667%}.col-6{flex:0 0 50%;max-width:50%}.col-7{flex:0 0 58.3333333333%;max-width:58.3333333333%}.col-8{flex:0 0 66.6666666667%;max-width:66.6666666667%}.col-9{flex:0 0 75%;max-width:75%}.col-10{flex:0 0 83.3333333333%;max-width:83.3333333333%}.col-11{flex:0 0 91.6666666667%;max-width:91.6666666667%}.col-12{flex:0 0 100%;max-width:100%}.order-first{order:-1}.order-last{order:13}.order-0{order:0}.order-1{order:1}.order-2{order:2}.order-3{order:3}.order-4{order:4}.order-5{order:5}.order-6{order:6}.order-7{order:7}.order-8{order:8}.order-9{order:9}.order-10{order:10}.order-11{order:11}.order-12{order:12}.offset-1{margin-left:8.3333333333%}.offset-2{margin-left:16.6666666667%}.offset-3{margin-left:25%}.offset-4{margin-left:33.3333333333%}.offset-5{margin-left:41.6666666667%}.offset-6{margin-left:50%}.offset-7{margin-left:58.3333333333%}.offset-8{margin-left:66.6666666667%}.offset-9{margin-left:75%}.offset-10{margin-left:83.3333333333%}.offset-11{margin-left:91.6666666667%}@media(min-width: 576px){.col-sm{flex-basis:0;flex-grow:1;max-width:100%}.row-cols-sm-1>*{flex:0 0 100%;max-width:100%}.row-cols-sm-2>*{flex:0 0 50%;max-width:50%}.row-cols-sm-3>*{flex:0 0 33.3333333333%;max-width:33.3333333333%}.row-cols-sm-4>*{flex:0 0 25%;max-width:25%}.row-cols-sm-5>*{flex:0 0 20%;max-width:20%}.row-cols-sm-6>*{flex:0 0 16.6666666667%;max-width:16.6666666667%}.col-sm-auto{flex:0 0 auto;width:auto;max-width:100%}.col-sm-1{flex:0 0 8.3333333333%;max-width:8.3333333333%}.col-sm-2{flex:0 0 16.6666666667%;max-width:16.6666666667%}.col-sm-3{flex:0 0 25%;max-width:25%}.col-sm-4{flex:0 0 33.3333333333%;max-width:33.3333333333%}.col-sm-5{flex:0 0 41.6666666667%;max-width:41.6666666667%}.col-sm-6{flex:0 0 50%;max-width:50%}.col-sm-7{flex:0 0 58.3333333333%;max-width:58.3333333333%}.col-sm-8{flex:0 0 66.6666666667%;max-width:66.6666666667%}.col-sm-9{flex:0 0 75%;max-width:75%}.col-sm-10{flex:0 0 83.3333333333%;max-width:83.3333333333%}.col-sm-11{flex:0 0 91.6666666667%;max-width:91.6666666667%}.col-sm-12{flex:0 0 100%;max-width:100%}.order-sm-first{order:-1}.order-sm-last{order:13}.order-sm-0{order:0}.order-sm-1{order:1}.order-sm-2{order:2}.order-sm-3{order:3}.order-sm-4{order:4}.order-sm-5{order:5}.order-sm-6{order:6}.order-sm-7{order:7}.order-sm-8{order:8}.order-sm-9{order:9}.order-sm-10{order:10}.order-sm-11{order:11}.order-sm-12{order:12}.offset-sm-0{margin-left:0}.offset-sm-1{margin-left:8.3333333333%}.offset-sm-2{margin-left:16.6666666667%}.offset-sm-3{margin-left:25%}.offset-sm-4{margin-left:33.3333333333%}.offset-sm-5{margin-left:41.6666666667%}.offset-sm-6{margin-left:50%}.offset-sm-7{margin-left:58.3333333333%}.offset-sm-8{margin-left:66.6666666667%}.offset-sm-9{margin-left:75%}.offset-sm-10{margin-left:83.3333333333%}.offset-sm-11{margin-left:91.6666666667%}}@media(min-width: 768px){.col-md{flex-basis:0;flex-grow:1;max-width:100%}.row-cols-md-1>*{flex:0 0 100%;max-width:100%}.row-cols-md-2>*{flex:0 0 50%;max-width:50%}.row-cols-md-3>*{flex:0 0 33.3333333333%;max-width:33.3333333333%}.row-cols-md-4>*{flex:0 0 25%;max-width:25%}.row-cols-md-5>*{flex:0 0 20%;max-width:20%}.row-cols-md-6>*{flex:0 0 16.6666666667%;max-width:16.6666666667%}.col-md-auto{flex:0 0 auto;width:auto;max-width:100%}.col-md-1{flex:0 0 8.3333333333%;max-width:8.3333333333%}.col-md-2{flex:0 0 16.6666666667%;max-width:16.6666666667%}.col-md-3{flex:0 0 25%;max-width:25%}.col-md-4{flex:0 0 33.3333333333%;max-width:33.3333333333%}.col-md-5{flex:0 0 41.6666666667%;max-width:41.6666666667%}.col-md-6{flex:0 0 50%;max-width:50%}.col-md-7{flex:0 0 58.3333333333%;max-width:58.3333333333%}.col-md-8{flex:0 0 66.6666666667%;max-width:66.6666666667%}.col-md-9{flex:0 0 75%;max-width:75%}.col-md-10{flex:0 0 83.3333333333%;max-width:83.3333333333%}.col-md-11{flex:0 0 91.6666666667%;max-width:91.6666666667%}.col-md-12{flex:0 0 100%;max-width:100%}.order-md-first{order:-1}.order-md-last{order:13}.order-md-0{order:0}.order-md-1{order:1}.order-md-2{order:2}.order-md-3{order:3}.order-md-4{order:4}.order-md-5{order:5}.order-md-6{order:6}.order-md-7{order:7}.order-md-8{order:8}.order-md-9{order:9}.order-md-10{order:10}.order-md-11{order:11}.order-md-12{order:12}.offset-md-0{margin-left:0}.offset-md-1{margin-left:8.3333333333%}.offset-md-2{margin-left:16.6666666667%}.offset-md-3{margin-left:25%}.offset-md-4{margin-left:33.3333333333%}.offset-md-5{margin-left:41.6666666667%}.offset-md-6{margin-left:50%}.offset-md-7{margin-left:58.3333333333%}.offset-md-8{margin-left:66.6666666667%}.offset-md-9{margin-left:75%}.offset-md-10{margin-left:83.3333333333%}.offset-md-11{margin-left:91.6666666667%}}@media(min-width: 992px){.col-lg{flex-basis:0;flex-grow:1;max-width:100%}.row-cols-lg-1>*{flex:0 0 100%;max-width:100%}.row-cols-lg-2>*{flex:0 0 50%;max-width:50%}.row-cols-lg-3>*{flex:0 0 33.3333333333%;max-width:33.3333333333%}.row-cols-lg-4>*{flex:0 0 25%;max-width:25%}.row-cols-lg-5>*{flex:0 0 20%;max-width:20%}.row-cols-lg-6>*{flex:0 0 16.6666666667%;max-width:16.6666666667%}.col-lg-auto{flex:0 0 auto;width:auto;max-width:100%}.col-lg-1{flex:0 0 8.3333333333%;max-width:8.3333333333%}.col-lg-2{flex:0 0 16.6666666667%;max-width:16.6666666667%}.col-lg-3{flex:0 0 25%;max-width:25%}.col-lg-4{flex:0 0 33.3333333333%;max-width:33.3333333333%}.col-lg-5{flex:0 0 41.6666666667%;max-width:41.6666666667%}.col-lg-6{flex:0 0 50%;max-width:50%}.col-lg-7{flex:0 0 58.3333333333%;max-width:58.3333333333%}.col-lg-8{flex:0 0 66.6666666667%;max-width:66.6666666667%}.col-lg-9{flex:0 0 75%;max-width:75%}.col-lg-10{flex:0 0 83.3333333333%;max-width:83.3333333333%}.col-lg-11{flex:0 0 91.6666666667%;max-width:91.6666666667%}.col-lg-12{flex:0 0 100%;max-width:100%}.order-lg-first{order:-1}.order-lg-last{order:13}.order-lg-0{order:0}.order-lg-1{order:1}.order-lg-2{order:2}.order-lg-3{order:3}.order-lg-4{order:4}.order-lg-5{order:5}.order-lg-6{order:6}.order-lg-7{order:7}.order-lg-8{order:8}.order-lg-9{order:9}.order-lg-10{order:10}.order-lg-11{order:11}.order-lg-12{order:12}.offset-lg-0{margin-left:0}.offset-lg-1{margin-left:8.3333333333%}.offset-lg-2{margin-left:16.6666666667%}.offset-lg-3{margin-left:25%}.offset-lg-4{margin-left:33.3333333333%}.offset-lg-5{margin-left:41.6666666667%}.offset-lg-6{margin-left:50%}.offset-lg-7{margin-left:58.3333333333%}.offset-lg-8{margin-left:66.6666666667%}.offset-lg-9{margin-left:75%}.offset-lg-10{margin-left:83.3333333333%}.offset-lg-11{margin-left:91.6666666667%}}@media(min-width: 1200px){.col-xl{flex-basis:0;flex-grow:1;max-width:100%}.row-cols-xl-1>*{flex:0 0 100%;max-width:100%}.row-cols-xl-2>*{flex:0 0 50%;max-width:50%}.row-cols-xl-3>*{flex:0 0 33.3333333333%;max-width:33.3333333333%}.row-cols-xl-4>*{flex:0 0 25%;max-width:25%}.row-cols-xl-5>*{flex:0 0 20%;max-width:20%}.row-cols-xl-6>*{flex:0 0 16.6666666667%;max-width:16.6666666667%}.col-xl-auto{flex:0 0 auto;width:auto;max-width:100%}.col-xl-1{flex:0 0 8.3333333333%;max-width:8.3333333333%}.col-xl-2{flex:0 0 16.6666666667%;max-width:16.6666666667%}.col-xl-3{flex:0 0 25%;max-width:25%}.col-xl-4{flex:0 0 33.3333333333%;max-width:33.3333333333%}.col-xl-5{flex:0 0 41.6666666667%;max-width:41.6666666667%}.col-xl-6{flex:0 0 50%;max-width:50%}.col-xl-7{flex:0 0 58.3333333333%;max-width:58.3333333333%}.col-xl-8{flex:0 0 66.6666666667%;max-width:66.6666666667%}.col-xl-9{flex:0 0 75%;max-width:75%}.col-xl-10{flex:0 0 83.3333333333%;max-width:83.3333333333%}.col-xl-11{flex:0 0 91.6666666667%;max-width:91.6666666667%}.col-xl-12{flex:0 0 100%;max-width:100%}.order-xl-first{order:-1}.order-xl-last{order:13}.order-xl-0{order:0}.order-xl-1{order:1}.order-xl-2{order:2}.order-xl-3{order:3}.order-xl-4{order:4}.order-xl-5{order:5}.order-xl-6{order:6}.order-xl-7{order:7}.order-xl-8{order:8}.order-xl-9{order:9}.order-xl-10{order:10}.order-xl-11{order:11}.order-xl-12{order:12}.offset-xl-0{margin-left:0}.offset-xl-1{margin-left:8.3333333333%}.offset-xl-2{margin-left:16.6666666667%}.offset-xl-3{margin-left:25%}.offset-xl-4{margin-left:33.3333333333%}.offset-xl-5{margin-left:41.6666666667%}.offset-xl-6{margin-left:50%}.offset-xl-7{margin-left:58.3333333333%}.offset-xl-8{margin-left:66.6666666667%}.offset-xl-9{margin-left:75%}.offset-xl-10{margin-left:83.3333333333%}.offset-xl-11{margin-left:91.6666666667%}}.table{width:100%;margin-bottom:1rem;color:#212529}.table th,.table td{padding:.75rem;vertical-align:top;border-top:1px solid #dee2e6}.table thead th{vertical-align:bottom;border-bottom:2px solid #dee2e6}.table tbody+tbody{border-top:2px solid #dee2e6}.table-sm th,.table-sm td{padding:.3rem}.table-bordered{border:1px solid #dee2e6}.table-bordered th,.table-bordered td{border:1px solid #dee2e6}.table-bordered thead th,.table-bordered thead td{border-bottom-width:2px}.table-borderless th,.table-borderless td,.table-borderless thead th,.table-borderless tbody+tbody{border:0}.table-striped tbody tr:nth-of-type(odd){background-color:rgba(0,0,0,.05)}.table-hover tbody tr:hover{color:#212529;background-color:rgba(0,0,0,.075)}.table-primary,.table-primary>th,.table-primary>td{background-color:#b8e7e4}.table-primary th,.table-primary td,.table-primary thead th,.table-primary tbody+tbody{border-color:#7ad2cc}.table-hover .table-primary:hover{background-color:#a5e1dd}.table-hover .table-primary:hover>td,.table-hover .table-primary:hover>th{background-color:#a5e1dd}.table-secondary,.table-secondary>th,.table-secondary>td{background-color:#d6d8db}.table-secondary th,.table-secondary td,.table-secondary thead th,.table-secondary tbody+tbody{border-color:#b3b7bb}.table-hover .table-secondary:hover{background-color:#c8cbcf}.table-hover .table-secondary:hover>td,.table-hover .table-secondary:hover>th{background-color:#c8cbcf}.table-success,.table-success>th,.table-success>td{background-color:#c3e6cb}.table-success th,.table-success td,.table-success thead th,.table-success tbody+tbody{border-color:#8fd19e}.table-hover .table-success:hover{background-color:#b1dfbb}.table-hover .table-success:hover>td,.table-hover .table-success:hover>th{background-color:#b1dfbb}.table-info,.table-info>th,.table-info>td{background-color:#bee5eb}.table-info th,.table-info td,.table-info thead th,.table-info tbody+tbody{border-color:#86cfda}.table-hover .table-info:hover{background-color:#abdde5}.table-hover .table-info:hover>td,.table-hover .table-info:hover>th{background-color:#abdde5}.table-warning,.table-warning>th,.table-warning>td{background-color:#ffeeba}.table-warning th,.table-warning td,.table-warning thead th,.table-warning tbody+tbody{border-color:#ffdf7e}.table-hover .table-warning:hover{background-color:#ffe8a1}.table-hover .table-warning:hover>td,.table-hover .table-warning:hover>th{background-color:#ffe8a1}.table-danger,.table-danger>th,.table-danger>td{background-color:#f5c6cb}.table-danger th,.table-danger td,.table-danger thead th,.table-danger tbody+tbody{border-color:#ed969e}.table-hover .table-danger:hover{background-color:#f1b0b7}.table-hover .table-danger:hover>td,.table-hover .table-danger:hover>th{background-color:#f1b0b7}.table-light,.table-light>th,.table-light>td{background-color:#fdfdfe}.table-light th,.table-light td,.table-light thead th,.table-light tbody+tbody{border-color:#fbfcfc}.table-hover .table-light:hover{background-color:#ececf6}.table-hover .table-light:hover>td,.table-hover .table-light:hover>th{background-color:#ececf6}.table-dark,.table-dark>th,.table-dark>td{background-color:#c6c8ca}.table-dark th,.table-dark td,.table-dark thead th,.table-dark tbody+tbody{border-color:#95999c}.table-hover .table-dark:hover{background-color:#b9bbbe}.table-hover .table-dark:hover>td,.table-hover .table-dark:hover>th{background-color:#b9bbbe}.table-active,.table-active>th,.table-active>td{background-color:rgba(0,0,0,.075)}.table-hover .table-active:hover{background-color:rgba(0,0,0,.075)}.table-hover .table-active:hover>td,.table-hover .table-active:hover>th{background-color:rgba(0,0,0,.075)}.table .thead-dark th{color:#fff;background-color:#343a40;border-color:#454d55}.table .thead-light th{color:#495057;background-color:#e9ecef;border-color:#dee2e6}.table-dark{color:#fff;background-color:#343a40}.table-dark th,.table-dark td,.table-dark thead th{border-color:#454d55}.table-dark.table-bordered{border:0}.table-dark.table-striped tbody tr:nth-of-type(odd){background-color:rgba(255,255,255,.05)}.table-dark.table-hover tbody tr:hover{color:#fff;background-color:rgba(255,255,255,.075)}@media(max-width: 575.98px){.table-responsive-sm{display:block;width:100%;overflow-x:auto;-webkit-overflow-scrolling:touch}.table-responsive-sm>.table-bordered{border:0}}@media(max-width: 767.98px){.table-responsive-md{display:block;width:100%;overflow-x:auto;-webkit-overflow-scrolling:touch}.table-responsive-md>.table-bordered{border:0}}@media(max-width: 991.98px){.table-responsive-lg{display:block;width:100%;overflow-x:auto;-webkit-overflow-scrolling:touch}.table-responsive-lg>.table-bordered{border:0}}@media(max-width: 1199.98px){.table-responsive-xl{display:block;width:100%;overflow-x:auto;-webkit-overflow-scrolling:touch}.table-responsive-xl>.table-bordered{border:0}}.table-responsive{display:block;width:100%;overflow-x:auto;-webkit-overflow-scrolling:touch}.table-responsive>.table-bordered{border:0}.form-control{display:block;width:100%;height:calc(1.5em + 0.75rem + 2px);padding:.375rem .75rem;font-size:1rem;font-weight:400;line-height:1.5;color:#495057;background-color:#fff;background-clip:padding-box;border:1px solid #ced4da;border-radius:.25rem;transition:border-color .15s ease-in-out,box-shadow .15s ease-in-out}@media(prefers-reduced-motion: reduce){.form-control{transition:none}}.form-control::-ms-expand{background-color:transparent;border:0}.form-control:-moz-focusring{color:transparent;text-shadow:0 0 0 #495057}.form-control:focus{color:#495057;background-color:#fff;border-color:#2afff0;outline:0;box-shadow:0 0 0 .2rem rgba(0,169,157,.25)}.form-control::placeholder{color:#6c757d;opacity:1}.form-control:disabled,.form-control[readonly]{background-color:#e9ecef;opacity:1}select.form-control:focus::-ms-value{color:#495057;background-color:#fff}.form-control-file,.form-control-range{display:block;width:100%}.col-form-label{padding-top:calc(0.375rem + 1px);padding-bottom:calc(0.375rem + 1px);margin-bottom:0;font-size:inherit;line-height:1.5}.col-form-label-lg{padding-top:calc(0.5rem + 1px);padding-bottom:calc(0.5rem + 1px);font-size:1.25rem;line-height:1.5}.col-form-label-sm{padding-top:calc(0.25rem + 1px);padding-bottom:calc(0.25rem + 1px);font-size:0.875rem;line-height:1.5}.form-control-plaintext{display:block;width:100%;padding:.375rem 0;margin-bottom:0;font-size:1rem;line-height:1.5;color:#212529;background-color:transparent;border:solid transparent;border-width:1px 0}.form-control-plaintext.form-control-sm,.form-control-plaintext.form-control-lg{padding-right:0;padding-left:0}.form-control-sm{height:calc(1.5em + 0.5rem + 2px);padding:.25rem .5rem;font-size:0.875rem;line-height:1.5;border-radius:.2rem}.form-control-lg{height:calc(1.5em + 1rem + 2px);padding:.5rem 1rem;font-size:1.25rem;line-height:1.5;border-radius:.3rem}select.form-control[size],select.form-control[multiple]{height:auto}textarea.form-control{height:auto}.form-group{margin-bottom:1rem}.form-text{display:block;margin-top:.25rem}.form-row{display:flex;flex-wrap:wrap;margin-right:-5px;margin-left:-5px}.form-row>.col,.form-row>[class*=col-]{padding-right:5px;padding-left:5px}.form-check{position:relative;display:block;padding-left:1.25rem}.form-check-input{position:absolute;margin-top:.3rem;margin-left:-1.25rem}.form-check-input[disabled]~.form-check-label,.form-check-input:disabled~.form-check-label{color:#6c757d}.form-check-label{margin-bottom:0}.form-check-inline{display:inline-flex;align-items:center;padding-left:0;margin-right:.75rem}.form-check-inline .form-check-input{position:static;margin-top:0;margin-right:.3125rem;margin-left:0}.valid-feedback{display:none;width:100%;margin-top:.25rem;font-size:80%;color:#28a745}.valid-tooltip{position:absolute;top:100%;z-index:5;display:none;max-width:100%;padding:.25rem .5rem;margin-top:.1rem;font-size:0.875rem;line-height:1.5;color:#fff;background-color:rgba(40,167,69,.9);border-radius:.25rem}.was-validated :valid~.valid-feedback,.was-validated :valid~.valid-tooltip,.is-valid~.valid-feedback,.is-valid~.valid-tooltip{display:block}.was-validated .form-control:valid,.form-control.is-valid{border-color:#28a745;padding-right:calc(1.5em + 0.75rem);background-image:url%28"data:image/svg+xml,%3csvg xmlns=\'http://www.w3.org/2000/svg\' width=\'8\' height=\'8\' viewBox=\'0 0 8 8\'%3e%3cpath fill=\'%2328a745\' d=\'M2.3 6.73L.6 4.53c-.4-1.04.46-1.4 1.1-.8l1.1 1.4 3.4-3.8c.6-.63 1.6-.27 1.2.7l-4 4.6c-.43.5-.8.4-1.1.1z\'/%3e%3c/svg%3e"%29;background-repeat:no-repeat;background-position:right calc(0.375em + 0.1875rem) center;background-size:calc(0.75em + 0.375rem) calc(0.75em + 0.375rem)}.was-validated .form-control:valid:focus,.form-control.is-valid:focus{border-color:#28a745;box-shadow:0 0 0 .2rem rgba(40,167,69,.25)}.was-validated textarea.form-control:valid,textarea.form-control.is-valid{padding-right:calc(1.5em + 0.75rem);background-position:top calc(0.375em + 0.1875rem) right calc(0.375em + 0.1875rem)}.was-validated .custom-select:valid,.custom-select.is-valid{border-color:#28a745;padding-right:calc(0.75em + 2.3125rem);background:url("data:image/svg+xml,%3csvg xmlns=\'http://www.w3.org/2000/svg\' width=\'4\' height=\'5\' viewBox=\'0 0 4 5\'%3e%3cpath fill=\'%23343a40\' d=\'M2 0L0 2h4zm0 5L0 3h4z\'/%3e%3c/svg%3e") right .75rem center/8px 10px no-repeat,url%28"data:image/svg+xml,%3csvg xmlns=\'http://www.w3.org/2000/svg\' width=\'8\' height=\'8\' viewBox=\'0 0 8 8\'%3e%3cpath fill=\'%2328a745\' d=\'M2.3 6.73L.6 4.53c-.4-1.04.46-1.4 1.1-.8l1.1 1.4 3.4-3.8c.6-.63 1.6-.27 1.2.7l-4 4.6c-.43.5-.8.4-1.1.1z\'/%3e%3c/svg%3e"%29 #fff no-repeat center right 1.75rem/calc(0.75em + 0.375rem) calc(0.75em + 0.375rem)}.was-validated .custom-select:valid:focus,.custom-select.is-valid:focus{border-color:#28a745;box-shadow:0 0 0 .2rem rgba(40,167,69,.25)}.was-validated .form-check-input:valid~.form-check-label,.form-check-input.is-valid~.form-check-label{color:#28a745}.was-validated .form-check-input:valid~.valid-feedback,.was-validated .form-check-input:valid~.valid-tooltip,.form-check-input.is-valid~.valid-feedback,.form-check-input.is-valid~.valid-tooltip{display:block}.was-validated .custom-control-input:valid~.custom-control-label,.custom-control-input.is-valid~.custom-control-label{color:#28a745}.was-validated .custom-control-input:valid~.custom-control-label::before,.custom-control-input.is-valid~.custom-control-label::before{border-color:#28a745}.was-validated .custom-control-input:valid:checked~.custom-control-label::before,.custom-control-input.is-valid:checked~.custom-control-label::before{border-color:#34ce57;background-color:#34ce57}.was-validated .custom-control-input:valid:focus~.custom-control-label::before,.custom-control-input.is-valid:focus~.custom-control-label::before{box-shadow:0 0 0 .2rem rgba(40,167,69,.25)}.was-validated .custom-control-input:valid:focus:not(:checked)~.custom-control-label::before,.custom-control-input.is-valid:focus:not(:checked)~.custom-control-label::before{border-color:#28a745}.was-validated .custom-file-input:valid~.custom-file-label,.custom-file-input.is-valid~.custom-file-label{border-color:#28a745}.was-validated .custom-file-input:valid:focus~.custom-file-label,.custom-file-input.is-valid:focus~.custom-file-label{border-color:#28a745;box-shadow:0 0 0 .2rem rgba(40,167,69,.25)}.invalid-feedback{display:none;width:100%;margin-top:.25rem;font-size:80%;color:#dc3545}.invalid-tooltip{position:absolute;top:100%;z-index:5;display:none;max-width:100%;padding:.25rem .5rem;margin-top:.1rem;font-size:0.875rem;line-height:1.5;color:#fff;background-color:rgba(220,53,69,.9);border-radius:.25rem}.was-validated :invalid~.invalid-feedback,.was-validated :invalid~.invalid-tooltip,.is-invalid~.invalid-feedback,.is-invalid~.invalid-tooltip{display:block}.was-validated .form-control:invalid,.form-control.is-invalid{border-color:#dc3545;padding-right:calc(1.5em + 0.75rem);background-image:url%28"data:image/svg+xml,%3csvg xmlns=\'http://www.w3.org/2000/svg\' width=\'12\' height=\'12\' fill=\'none\' stroke=\'%23dc3545\' viewBox=\'0 0 12 12\'%3e%3ccircle cx=\'6\' cy=\'6\' r=\'4.5\'/%3e%3cpath stroke-linejoin=\'round\' d=\'M5.8 3.6h.4L6 6.5z\'/%3e%3ccircle cx=\'6\' cy=\'8.2\' r=\'.6\' fill=\'%23dc3545\' stroke=\'none\'/%3e%3c/svg%3e"%29;background-repeat:no-repeat;background-position:right calc(0.375em + 0.1875rem) center;background-size:calc(0.75em + 0.375rem) calc(0.75em + 0.375rem)}.was-validated .form-control:invalid:focus,.form-control.is-invalid:focus{border-color:#dc3545;box-shadow:0 0 0 .2rem rgba(220,53,69,.25)}.was-validated textarea.form-control:invalid,textarea.form-control.is-invalid{padding-right:calc(1.5em + 0.75rem);background-position:top calc(0.375em + 0.1875rem) right calc(0.375em + 0.1875rem)}.was-validated .custom-select:invalid,.custom-select.is-invalid{border-color:#dc3545;padding-right:calc(0.75em + 2.3125rem);background:url("data:image/svg+xml,%3csvg xmlns=\'http://www.w3.org/2000/svg\' width=\'4\' height=\'5\' viewBox=\'0 0 4 5\'%3e%3cpath fill=\'%23343a40\' d=\'M2 0L0 2h4zm0 5L0 3h4z\'/%3e%3c/svg%3e") right .75rem center/8px 10px no-repeat,url%28"data:image/svg+xml,%3csvg xmlns=\'http://www.w3.org/2000/svg\' width=\'12\' height=\'12\' fill=\'none\' stroke=\'%23dc3545\' viewBox=\'0 0 12 12\'%3e%3ccircle cx=\'6\' cy=\'6\' r=\'4.5\'/%3e%3cpath stroke-linejoin=\'round\' d=\'M5.8 3.6h.4L6 6.5z\'/%3e%3ccircle cx=\'6\' cy=\'8.2\' r=\'.6\' fill=\'%23dc3545\' stroke=\'none\'/%3e%3c/svg%3e"%29 #fff no-repeat center right 1.75rem/calc(0.75em + 0.375rem) calc(0.75em + 0.375rem)}.was-validated .custom-select:invalid:focus,.custom-select.is-invalid:focus{border-color:#dc3545;box-shadow:0 0 0 .2rem rgba(220,53,69,.25)}.was-validated .form-check-input:invalid~.form-check-label,.form-check-input.is-invalid~.form-check-label{color:#dc3545}.was-validated .form-check-input:invalid~.invalid-feedback,.was-validated .form-check-input:invalid~.invalid-tooltip,.form-check-input.is-invalid~.invalid-feedback,.form-check-input.is-invalid~.invalid-tooltip{display:block}.was-validated .custom-control-input:invalid~.custom-control-label,.custom-control-input.is-invalid~.custom-control-label{color:#dc3545}.was-validated .custom-control-input:invalid~.custom-control-label::before,.custom-control-input.is-invalid~.custom-control-label::before{border-color:#dc3545}.was-validated .custom-control-input:invalid:checked~.custom-control-label::before,.custom-control-input.is-invalid:checked~.custom-control-label::before{border-color:#e4606d;background-color:#e4606d}.was-validated .custom-control-input:invalid:focus~.custom-control-label::before,.custom-control-input.is-invalid:focus~.custom-control-label::before{box-shadow:0 0 0 .2rem rgba(220,53,69,.25)}.was-validated .custom-control-input:invalid:focus:not(:checked)~.custom-control-label::before,.custom-control-input.is-invalid:focus:not(:checked)~.custom-control-label::before{border-color:#dc3545}.was-validated .custom-file-input:invalid~.custom-file-label,.custom-file-input.is-invalid~.custom-file-label{border-color:#dc3545}.was-validated .custom-file-input:invalid:focus~.custom-file-label,.custom-file-input.is-invalid:focus~.custom-file-label{border-color:#dc3545;box-shadow:0 0 0 .2rem rgba(220,53,69,.25)}.form-inline{display:flex;flex-flow:row wrap;align-items:center}.form-inline .form-check{width:100%}@media(min-width: 576px){.form-inline label{display:flex;align-items:center;justify-content:center;margin-bottom:0}.form-inline .form-group{display:flex;flex:0 0 auto;flex-flow:row wrap;align-items:center;margin-bottom:0}.form-inline .form-control{display:inline-block;width:auto;vertical-align:middle}.form-inline .form-control-plaintext{display:inline-block}.form-inline .input-group,.form-inline .custom-select{width:auto}.form-inline .form-check{display:flex;align-items:center;justify-content:center;width:auto;padding-left:0}.form-inline .form-check-input{position:relative;flex-shrink:0;margin-top:0;margin-right:.25rem;margin-left:0}.form-inline .custom-control{align-items:center;justify-content:center}.form-inline .custom-control-label{margin-bottom:0}}.btn{display:inline-block;font-weight:400;color:#212529;text-align:center;vertical-align:middle;cursor:pointer;user-select:none;background-color:transparent;border:1px solid transparent;padding:.375rem .75rem;font-size:1rem;line-height:1.5;border-radius:.25rem;transition:color .15s ease-in-out,background-color .15s ease-in-out,border-color .15s ease-in-out,box-shadow .15s ease-in-out}@media(prefers-reduced-motion: reduce){.btn{transition:none}}.btn:hover{color:#212529;text-decoration:none}.btn:focus,.btn.focus{outline:0;box-shadow:0 0 0 .2rem rgba(0,169,157,.25)}.btn.disabled,.btn:disabled{opacity:.65}a.btn.disabled,fieldset:disabled a.btn{pointer-events:none}.btn-primary{color:#fff;background-color:#00a99d;border-color:#00a99d}.btn-primary:hover{color:#fff;background-color:#008379;border-color:#00766e}.btn-primary:focus,.btn-primary.focus{color:#fff;background-color:#008379;border-color:#00766e;box-shadow:0 0 0 .2rem rgba(38,182,172,.5)}.btn-primary.disabled,.btn-primary:disabled{color:#fff;background-color:#00a99d;border-color:#00a99d}.btn-primary:not(:disabled):not(.disabled):active,.btn-primary:not(:disabled):not(.disabled).active,.show>.btn-primary.dropdown-toggle{color:#fff;background-color:#00766e;border-color:#006962}.btn-primary:not(:disabled):not(.disabled):active:focus,.btn-primary:not(:disabled):not(.disabled).active:focus,.show>.btn-primary.dropdown-toggle:focus{box-shadow:0 0 0 .2rem rgba(38,182,172,.5)}.btn-secondary{color:#fff;background-color:#6c757d;border-color:#6c757d}.btn-secondary:hover{color:#fff;background-color:#5a6268;border-color:#545b62}.btn-secondary:focus,.btn-secondary.focus{color:#fff;background-color:#5a6268;border-color:#545b62;box-shadow:0 0 0 .2rem rgba(130,138,145,.5)}.btn-secondary.disabled,.btn-secondary:disabled{color:#fff;background-color:#6c757d;border-color:#6c757d}.btn-secondary:not(:disabled):not(.disabled):active,.btn-secondary:not(:disabled):not(.disabled).active,.show>.btn-secondary.dropdown-toggle{color:#fff;background-color:#545b62;border-color:#4e555b}.btn-secondary:not(:disabled):not(.disabled):active:focus,.btn-secondary:not(:disabled):not(.disabled).active:focus,.show>.btn-secondary.dropdown-toggle:focus{box-shadow:0 0 0 .2rem rgba(130,138,145,.5)}.btn-success{color:#fff;background-color:#28a745;border-color:#28a745}.btn-success:hover{color:#fff;background-color:#218838;border-color:#1e7e34}.btn-success:focus,.btn-success.focus{color:#fff;background-color:#218838;border-color:#1e7e34;box-shadow:0 0 0 .2rem rgba(72,180,97,.5)}.btn-success.disabled,.btn-success:disabled{color:#fff;background-color:#28a745;border-color:#28a745}.btn-success:not(:disabled):not(.disabled):active,.btn-success:not(:disabled):not(.disabled).active,.show>.btn-success.dropdown-toggle{color:#fff;background-color:#1e7e34;border-color:#1c7430}.btn-success:not(:disabled):not(.disabled):active:focus,.btn-success:not(:disabled):not(.disabled).active:focus,.show>.btn-success.dropdown-toggle:focus{box-shadow:0 0 0 .2rem rgba(72,180,97,.5)}.btn-info{color:#fff;background-color:#17a2b8;border-color:#17a2b8}.btn-info:hover{color:#fff;background-color:#138496;border-color:#117a8b}.btn-info:focus,.btn-info.focus{color:#fff;background-color:#138496;border-color:#117a8b;box-shadow:0 0 0 .2rem rgba(58,176,195,.5)}.btn-info.disabled,.btn-info:disabled{color:#fff;background-color:#17a2b8;border-color:#17a2b8}.btn-info:not(:disabled):not(.disabled):active,.btn-info:not(:disabled):not(.disabled).active,.show>.btn-info.dropdown-toggle{color:#fff;background-color:#117a8b;border-color:#10707f}.btn-info:not(:disabled):not(.disabled):active:focus,.btn-info:not(:disabled):not(.disabled).active:focus,.show>.btn-info.dropdown-toggle:focus{box-shadow:0 0 0 .2rem rgba(58,176,195,.5)}.btn-warning{color:#212529;background-color:#ffc107;border-color:#ffc107}.btn-warning:hover{color:#212529;background-color:#e0a800;border-color:#d39e00}.btn-warning:focus,.btn-warning.focus{color:#212529;background-color:#e0a800;border-color:#d39e00;box-shadow:0 0 0 .2rem rgba(222,170,12,.5)}.btn-warning.disabled,.btn-warning:disabled{color:#212529;background-color:#ffc107;border-color:#ffc107}.btn-warning:not(:disabled):not(.disabled):active,.btn-warning:not(:disabled):not(.disabled).active,.show>.btn-warning.dropdown-toggle{color:#212529;background-color:#d39e00;border-color:#c69500}.btn-warning:not(:disabled):not(.disabled):active:focus,.btn-warning:not(:disabled):not(.disabled).active:focus,.show>.btn-warning.dropdown-toggle:focus{box-shadow:0 0 0 .2rem rgba(222,170,12,.5)}.btn-danger{color:#fff;background-color:#dc3545;border-color:#dc3545}.btn-danger:hover{color:#fff;background-color:#c82333;border-color:#bd2130}.btn-danger:focus,.btn-danger.focus{color:#fff;background-color:#c82333;border-color:#bd2130;box-shadow:0 0 0 .2rem rgba(225,83,97,.5)}.btn-danger.disabled,.btn-danger:disabled{color:#fff;background-color:#dc3545;border-color:#dc3545}.btn-danger:not(:disabled):not(.disabled):active,.btn-danger:not(:disabled):not(.disabled).active,.show>.btn-danger.dropdown-toggle{color:#fff;background-color:#bd2130;border-color:#b21f2d}.btn-danger:not(:disabled):not(.disabled):active:focus,.btn-danger:not(:disabled):not(.disabled).active:focus,.show>.btn-danger.dropdown-toggle:focus{box-shadow:0 0 0 .2rem rgba(225,83,97,.5)}.btn-light{color:#212529;background-color:#f8f9fa;border-color:#f8f9fa}.btn-light:hover{color:#212529;background-color:#e2e6ea;border-color:#dae0e5}.btn-light:focus,.btn-light.focus{color:#212529;background-color:#e2e6ea;border-color:#dae0e5;box-shadow:0 0 0 .2rem rgba(216,217,219,.5)}.btn-light.disabled,.btn-light:disabled{color:#212529;background-color:#f8f9fa;border-color:#f8f9fa}.btn-light:not(:disabled):not(.disabled):active,.btn-light:not(:disabled):not(.disabled).active,.show>.btn-light.dropdown-toggle{color:#212529;background-color:#dae0e5;border-color:#d3d9df}.btn-light:not(:disabled):not(.disabled):active:focus,.btn-light:not(:disabled):not(.disabled).active:focus,.show>.btn-light.dropdown-toggle:focus{box-shadow:0 0 0 .2rem rgba(216,217,219,.5)}.btn-dark{color:#fff;background-color:#343a40;border-color:#343a40}.btn-dark:hover{color:#fff;background-color:#23272b;border-color:#1d2124}.btn-dark:focus,.btn-dark.focus{color:#fff;background-color:#23272b;border-color:#1d2124;box-shadow:0 0 0 .2rem rgba(82,88,93,.5)}.btn-dark.disabled,.btn-dark:disabled{color:#fff;background-color:#343a40;border-color:#343a40}.btn-dark:not(:disabled):not(.disabled):active,.btn-dark:not(:disabled):not(.disabled).active,.show>.btn-dark.dropdown-toggle{color:#fff;background-color:#1d2124;border-color:#171a1d}.btn-dark:not(:disabled):not(.disabled):active:focus,.btn-dark:not(:disabled):not(.disabled).active:focus,.show>.btn-dark.dropdown-toggle:focus{box-shadow:0 0 0 .2rem rgba(82,88,93,.5)}.btn-outline-primary{color:#00a99d;border-color:#00a99d}.btn-outline-primary:hover{color:#fff;background-color:#00a99d;border-color:#00a99d}.btn-outline-primary:focus,.btn-outline-primary.focus{box-shadow:0 0 0 .2rem rgba(0,169,157,.5)}.btn-outline-primary.disabled,.btn-outline-primary:disabled{color:#00a99d;background-color:transparent}.btn-outline-primary:not(:disabled):not(.disabled):active,.btn-outline-primary:not(:disabled):not(.disabled).active,.show>.btn-outline-primary.dropdown-toggle{color:#fff;background-color:#00a99d;border-color:#00a99d}.btn-outline-primary:not(:disabled):not(.disabled):active:focus,.btn-outline-primary:not(:disabled):not(.disabled).active:focus,.show>.btn-outline-primary.dropdown-toggle:focus{box-shadow:0 0 0 .2rem rgba(0,169,157,.5)}.btn-outline-secondary{color:#6c757d;border-color:#6c757d}.btn-outline-secondary:hover{color:#fff;background-color:#6c757d;border-color:#6c757d}.btn-outline-secondary:focus,.btn-outline-secondary.focus{box-shadow:0 0 0 .2rem rgba(108,117,125,.5)}.btn-outline-secondary.disabled,.btn-outline-secondary:disabled{color:#6c757d;background-color:transparent}.btn-outline-secondary:not(:disabled):not(.disabled):active,.btn-outline-secondary:not(:disabled):not(.disabled).active,.show>.btn-outline-secondary.dropdown-toggle{color:#fff;background-color:#6c757d;border-color:#6c757d}.btn-outline-secondary:not(:disabled):not(.disabled):active:focus,.btn-outline-secondary:not(:disabled):not(.disabled).active:focus,.show>.btn-outline-secondary.dropdown-toggle:focus{box-shadow:0 0 0 .2rem rgba(108,117,125,.5)}.btn-outline-success{color:#28a745;border-color:#28a745}.btn-outline-success:hover{color:#fff;background-color:#28a745;border-color:#28a745}.btn-outline-success:focus,.btn-outline-success.focus{box-shadow:0 0 0 .2rem rgba(40,167,69,.5)}.btn-outline-success.disabled,.btn-outline-success:disabled{color:#28a745;background-color:transparent}.btn-outline-success:not(:disabled):not(.disabled):active,.btn-outline-success:not(:disabled):not(.disabled).active,.show>.btn-outline-success.dropdown-toggle{color:#fff;background-color:#28a745;border-color:#28a745}.btn-outline-success:not(:disabled):not(.disabled):active:focus,.btn-outline-success:not(:disabled):not(.disabled).active:focus,.show>.btn-outline-success.dropdown-toggle:focus{box-shadow:0 0 0 .2rem rgba(40,167,69,.5)}.btn-outline-info{color:#17a2b8;border-color:#17a2b8}.btn-outline-info:hover{color:#fff;background-color:#17a2b8;border-color:#17a2b8}.btn-outline-info:focus,.btn-outline-info.focus{box-shadow:0 0 0 .2rem rgba(23,162,184,.5)}.btn-outline-info.disabled,.btn-outline-info:disabled{color:#17a2b8;background-color:transparent}.btn-outline-info:not(:disabled):not(.disabled):active,.btn-outline-info:not(:disabled):not(.disabled).active,.show>.btn-outline-info.dropdown-toggle{color:#fff;background-color:#17a2b8;border-color:#17a2b8}.btn-outline-info:not(:disabled):not(.disabled):active:focus,.btn-outline-info:not(:disabled):not(.disabled).active:focus,.show>.btn-outline-info.dropdown-toggle:focus{box-shadow:0 0 0 .2rem rgba(23,162,184,.5)}.btn-outline-warning{color:#ffc107;border-color:#ffc107}.btn-outline-warning:hover{color:#212529;background-color:#ffc107;border-color:#ffc107}.btn-outline-warning:focus,.btn-outline-warning.focus{box-shadow:0 0 0 .2rem rgba(255,193,7,.5)}.btn-outline-warning.disabled,.btn-outline-warning:disabled{color:#ffc107;background-color:transparent}.btn-outline-warning:not(:disabled):not(.disabled):active,.btn-outline-warning:not(:disabled):not(.disabled).active,.show>.btn-outline-warning.dropdown-toggle{color:#212529;background-color:#ffc107;border-color:#ffc107}.btn-outline-warning:not(:disabled):not(.disabled):active:focus,.btn-outline-warning:not(:disabled):not(.disabled).active:focus,.show>.btn-outline-warning.dropdown-toggle:focus{box-shadow:0 0 0 .2rem rgba(255,193,7,.5)}.btn-outline-danger{color:#dc3545;border-color:#dc3545}.btn-outline-danger:hover{color:#fff;background-color:#dc3545;border-color:#dc3545}.btn-outline-danger:focus,.btn-outline-danger.focus{box-shadow:0 0 0 .2rem rgba(220,53,69,.5)}.btn-outline-danger.disabled,.btn-outline-danger:disabled{color:#dc3545;background-color:transparent}.btn-outline-danger:not(:disabled):not(.disabled):active,.btn-outline-danger:not(:disabled):not(.disabled).active,.show>.btn-outline-danger.dropdown-toggle{color:#fff;background-color:#dc3545;border-color:#dc3545}.btn-outline-danger:not(:disabled):not(.disabled):active:focus,.btn-outline-danger:not(:disabled):not(.disabled).active:focus,.show>.btn-outline-danger.dropdown-toggle:focus{box-shadow:0 0 0 .2rem rgba(220,53,69,.5)}.btn-outline-light{color:#f8f9fa;border-color:#f8f9fa}.btn-outline-light:hover{color:#212529;background-color:#f8f9fa;border-color:#f8f9fa}.btn-outline-light:focus,.btn-outline-light.focus{box-shadow:0 0 0 .2rem rgba(248,249,250,.5)}.btn-outline-light.disabled,.btn-outline-light:disabled{color:#f8f9fa;background-color:transparent}.btn-outline-light:not(:disabled):not(.disabled):active,.btn-outline-light:not(:disabled):not(.disabled).active,.show>.btn-outline-light.dropdown-toggle{color:#212529;background-color:#f8f9fa;border-color:#f8f9fa}.btn-outline-light:not(:disabled):not(.disabled):active:focus,.btn-outline-light:not(:disabled):not(.disabled).active:focus,.show>.btn-outline-light.dropdown-toggle:focus{box-shadow:0 0 0 .2rem rgba(248,249,250,.5)}.btn-outline-dark{color:#343a40;border-color:#343a40}.btn-outline-dark:hover{color:#fff;background-color:#343a40;border-color:#343a40}.btn-outline-dark:focus,.btn-outline-dark.focus{box-shadow:0 0 0 .2rem rgba(52,58,64,.5)}.btn-outline-dark.disabled,.btn-outline-dark:disabled{color:#343a40;background-color:transparent}.btn-outline-dark:not(:disabled):not(.disabled):active,.btn-outline-dark:not(:disabled):not(.disabled).active,.show>.btn-outline-dark.dropdown-toggle{color:#fff;background-color:#343a40;border-color:#343a40}.btn-outline-dark:not(:disabled):not(.disabled):active:focus,.btn-outline-dark:not(:disabled):not(.disabled).active:focus,.show>.btn-outline-dark.dropdown-toggle:focus{box-shadow:0 0 0 .2rem rgba(52,58,64,.5)}.btn-link{font-weight:400;color:#00a99d;text-decoration:none}.btn-link:hover{color:#005d56;text-decoration:underline}.btn-link:focus,.btn-link.focus{text-decoration:underline;box-shadow:none}.btn-link:disabled,.btn-link.disabled{color:#6c757d;pointer-events:none}.btn-lg,.btn-group-lg>.btn{padding:.5rem 1rem;font-size:1.25rem;line-height:1.5;border-radius:.3rem}.btn-sm,.btn-group-sm>.btn{padding:.25rem .5rem;font-size:0.875rem;line-height:1.5;border-radius:.2rem}.btn-block{display:block;width:100%}.btn-block+.btn-block{margin-top:.5rem}input[type=submit].btn-block,input[type=reset].btn-block,input[type=button].btn-block{width:100%}.fade{transition:opacity .15s linear}@media(prefers-reduced-motion: reduce){.fade{transition:none}}.fade:not(.show){opacity:0}.collapse:not(.show){display:none}.collapsing{position:relative;height:0;overflow:hidden;transition:height .35s ease}@media(prefers-reduced-motion: reduce){.collapsing{transition:none}}.dropup,.dropright,.dropdown,.dropleft{position:relative}.dropdown-toggle{white-space:nowrap}.dropdown-toggle::after{display:inline-block;margin-left:.255em;vertical-align:.255em;content:"";border-top:.3em solid;border-right:.3em solid transparent;border-bottom:0;border-left:.3em solid transparent}.dropdown-toggle:empty::after{margin-left:0}.dropdown-menu{position:absolute;top:100%;left:0;z-index:1000;display:none;float:left;min-width:10rem;padding:.5rem 0;margin:.125rem 0 0;font-size:1rem;color:#212529;text-align:left;list-style:none;background-color:#fff;background-clip:padding-box;border:1px solid rgba(0,0,0,.15);border-radius:.25rem}.dropdown-menu-left{right:auto;left:0}.dropdown-menu-right{right:0;left:auto}@media(min-width: 576px){.dropdown-menu-sm-left{right:auto;left:0}.dropdown-menu-sm-right{right:0;left:auto}}@media(min-width: 768px){.dropdown-menu-md-left{right:auto;left:0}.dropdown-menu-md-right{right:0;left:auto}}@media(min-width: 992px){.dropdown-menu-lg-left{right:auto;left:0}.dropdown-menu-lg-right{right:0;left:auto}}@media(min-width: 1200px){.dropdown-menu-xl-left{right:auto;left:0}.dropdown-menu-xl-right{right:0;left:auto}}.dropup .dropdown-menu{top:auto;bottom:100%;margin-top:0;margin-bottom:.125rem}.dropup .dropdown-toggle::after{display:inline-block;margin-left:.255em;vertical-align:.255em;content:"";border-top:0;border-right:.3em solid transparent;border-bottom:.3em solid;border-left:.3em solid transparent}.dropup .dropdown-toggle:empty::after{margin-left:0}.dropright .dropdown-menu{top:0;right:auto;left:100%;margin-top:0;margin-left:.125rem}.dropright .dropdown-toggle::after{display:inline-block;margin-left:.255em;vertical-align:.255em;content:"";border-top:.3em solid transparent;border-right:0;border-bottom:.3em solid transparent;border-left:.3em solid}.dropright .dropdown-toggle:empty::after{margin-left:0}.dropright .dropdown-toggle::after{vertical-align:0}.dropleft .dropdown-menu{top:0;right:100%;left:auto;margin-top:0;margin-right:.125rem}.dropleft .dropdown-toggle::after{display:inline-block;margin-left:.255em;vertical-align:.255em;content:""}.dropleft .dropdown-toggle::after{display:none}.dropleft .dropdown-toggle::before{display:inline-block;margin-right:.255em;vertical-align:.255em;content:"";border-top:.3em solid transparent;border-right:.3em solid;border-bottom:.3em solid transparent}.dropleft .dropdown-toggle:empty::after{margin-left:0}.dropleft .dropdown-toggle::before{vertical-align:0}.dropdown-menu[x-placement^=top],.dropdown-menu[x-placement^=right],.dropdown-menu[x-placement^=bottom],.dropdown-menu[x-placement^=left]{right:auto;bottom:auto}.dropdown-divider{height:0;margin:.5rem 0;overflow:hidden;border-top:1px solid #e9ecef}.dropdown-item{display:block;width:100%;padding:.25rem 1.5rem;clear:both;font-weight:400;color:#212529;text-align:inherit;white-space:nowrap;background-color:transparent;border:0}.dropdown-item:hover,.dropdown-item:focus{color:#16181b;text-decoration:none;background-color:#e9ecef}.dropdown-item.active,.dropdown-item:active{color:#fff;text-decoration:none;background-color:#00a99d}.dropdown-item.disabled,.dropdown-item:disabled{color:#adb5bd;pointer-events:none;background-color:transparent}.dropdown-menu.show{display:block}.dropdown-header{display:block;padding:.5rem 1.5rem;margin-bottom:0;font-size:0.875rem;color:#6c757d;white-space:nowrap}.dropdown-item-text{display:block;padding:.25rem 1.5rem;color:#212529}.btn-group,.btn-group-vertical{position:relative;display:inline-flex;vertical-align:middle}.btn-group>.btn,.btn-group-vertical>.btn{position:relative;flex:1 1 auto}.btn-group>.btn:hover,.btn-group-vertical>.btn:hover{z-index:1}.btn-group>.btn:focus,.btn-group>.btn:active,.btn-group>.btn.active,.btn-group-vertical>.btn:focus,.btn-group-vertical>.btn:active,.btn-group-vertical>.btn.active{z-index:1}.btn-toolbar{display:flex;flex-wrap:wrap;justify-content:flex-start}.btn-toolbar .input-group{width:auto}.btn-group>.btn:not(:first-child),.btn-group>.btn-group:not(:first-child){margin-left:-1px}.btn-group>.btn:not(:last-child):not(.dropdown-toggle),.btn-group>.btn-group:not(:last-child)>.btn{border-top-right-radius:0;border-bottom-right-radius:0}.btn-group>.btn:not(:first-child),.btn-group>.btn-group:not(:first-child)>.btn{border-top-left-radius:0;border-bottom-left-radius:0}.dropdown-toggle-split{padding-right:.5625rem;padding-left:.5625rem}.dropdown-toggle-split::after,.dropup .dropdown-toggle-split::after,.dropright .dropdown-toggle-split::after{margin-left:0}.dropleft .dropdown-toggle-split::before{margin-right:0}.btn-sm+.dropdown-toggle-split,.btn-group-sm>.btn+.dropdown-toggle-split{padding-right:.375rem;padding-left:.375rem}.btn-lg+.dropdown-toggle-split,.btn-group-lg>.btn+.dropdown-toggle-split{padding-right:.75rem;padding-left:.75rem}.btn-group-vertical{flex-direction:column;align-items:flex-start;justify-content:center}.btn-group-vertical>.btn,.btn-group-vertical>.btn-group{width:100%}.btn-group-vertical>.btn:not(:first-child),.btn-group-vertical>.btn-group:not(:first-child){margin-top:-1px}.btn-group-vertical>.btn:not(:last-child):not(.dropdown-toggle),.btn-group-vertical>.btn-group:not(:last-child)>.btn{border-bottom-right-radius:0;border-bottom-left-radius:0}.btn-group-vertical>.btn:not(:first-child),.btn-group-vertical>.btn-group:not(:first-child)>.btn{border-top-left-radius:0;border-top-right-radius:0}.btn-group-toggle>.btn,.btn-group-toggle>.btn-group>.btn{margin-bottom:0}.btn-group-toggle>.btn input[type=radio],.btn-group-toggle>.btn input[type=checkbox],.btn-group-toggle>.btn-group>.btn input[type=radio],.btn-group-toggle>.btn-group>.btn input[type=checkbox]{position:absolute;clip:rect(0, 0, 0, 0);pointer-events:none}.input-group{position:relative;display:flex;flex-wrap:wrap;align-items:stretch;width:100%}.input-group>.form-control,.input-group>.form-control-plaintext,.input-group>.custom-select,.input-group>.custom-file{position:relative;flex:1 1 0%;min-width:0;margin-bottom:0}.input-group>.form-control+.form-control,.input-group>.form-control+.custom-select,.input-group>.form-control+.custom-file,.input-group>.form-control-plaintext+.form-control,.input-group>.form-control-plaintext+.custom-select,.input-group>.form-control-plaintext+.custom-file,.input-group>.custom-select+.form-control,.input-group>.custom-select+.custom-select,.input-group>.custom-select+.custom-file,.input-group>.custom-file+.form-control,.input-group>.custom-file+.custom-select,.input-group>.custom-file+.custom-file{margin-left:-1px}.input-group>.form-control:focus,.input-group>.custom-select:focus,.input-group>.custom-file .custom-file-input:focus~.custom-file-label{z-index:3}.input-group>.custom-file .custom-file-input:focus{z-index:4}.input-group>.form-control:not(:last-child),.input-group>.custom-select:not(:last-child){border-top-right-radius:0;border-bottom-right-radius:0}.input-group>.form-control:not(:first-child),.input-group>.custom-select:not(:first-child){border-top-left-radius:0;border-bottom-left-radius:0}.input-group>.custom-file{display:flex;align-items:center}.input-group>.custom-file:not(:last-child) .custom-file-label,.input-group>.custom-file:not(:last-child) .custom-file-label::after{border-top-right-radius:0;border-bottom-right-radius:0}.input-group>.custom-file:not(:first-child) .custom-file-label{border-top-left-radius:0;border-bottom-left-radius:0}.input-group-prepend,.input-group-append{display:flex}.input-group-prepend .btn,.input-group-append .btn{position:relative;z-index:2}.input-group-prepend .btn:focus,.input-group-append .btn:focus{z-index:3}.input-group-prepend .btn+.btn,.input-group-prepend .btn+.input-group-text,.input-group-prepend .input-group-text+.input-group-text,.input-group-prepend .input-group-text+.btn,.input-group-append .btn+.btn,.input-group-append .btn+.input-group-text,.input-group-append .input-group-text+.input-group-text,.input-group-append .input-group-text+.btn{margin-left:-1px}.input-group-prepend{margin-right:-1px}.input-group-append{margin-left:-1px}.input-group-text{display:flex;align-items:center;padding:.375rem .75rem;margin-bottom:0;font-size:1rem;font-weight:400;line-height:1.5;color:#495057;text-align:center;white-space:nowrap;background-color:#e9ecef;border:1px solid #ced4da;border-radius:.25rem}.input-group-text input[type=radio],.input-group-text input[type=checkbox]{margin-top:0}.input-group-lg>.form-control:not(textarea),.input-group-lg>.custom-select{height:calc(1.5em + 1rem + 2px)}.input-group-lg>.form-control,.input-group-lg>.custom-select,.input-group-lg>.input-group-prepend>.input-group-text,.input-group-lg>.input-group-append>.input-group-text,.input-group-lg>.input-group-prepend>.btn,.input-group-lg>.input-group-append>.btn{padding:.5rem 1rem;font-size:1.25rem;line-height:1.5;border-radius:.3rem}.input-group-sm>.form-control:not(textarea),.input-group-sm>.custom-select{height:calc(1.5em + 0.5rem + 2px)}.input-group-sm>.form-control,.input-group-sm>.custom-select,.input-group-sm>.input-group-prepend>.input-group-text,.input-group-sm>.input-group-append>.input-group-text,.input-group-sm>.input-group-prepend>.btn,.input-group-sm>.input-group-append>.btn{padding:.25rem .5rem;font-size:0.875rem;line-height:1.5;border-radius:.2rem}.input-group-lg>.custom-select,.input-group-sm>.custom-select{padding-right:1.75rem}.input-group>.input-group-prepend>.btn,.input-group>.input-group-prepend>.input-group-text,.input-group>.input-group-append:not(:last-child)>.btn,.input-group>.input-group-append:not(:last-child)>.input-group-text,.input-group>.input-group-append:last-child>.btn:not(:last-child):not(.dropdown-toggle),.input-group>.input-group-append:last-child>.input-group-text:not(:last-child){border-top-right-radius:0;border-bottom-right-radius:0}.input-group>.input-group-append>.btn,.input-group>.input-group-append>.input-group-text,.input-group>.input-group-prepend:not(:first-child)>.btn,.input-group>.input-group-prepend:not(:first-child)>.input-group-text,.input-group>.input-group-prepend:first-child>.btn:not(:first-child),.input-group>.input-group-prepend:first-child>.input-group-text:not(:first-child){border-top-left-radius:0;border-bottom-left-radius:0}.custom-control{position:relative;display:block;min-height:1.5rem;padding-left:1.5rem}.custom-control-inline{display:inline-flex;margin-right:1rem}.custom-control-input{position:absolute;left:0;z-index:-1;width:1rem;height:1.25rem;opacity:0}.custom-control-input:checked~.custom-control-label::before{color:#fff;border-color:#00a99d;background-color:#00a99d}.custom-control-input:focus~.custom-control-label::before{box-shadow:0 0 0 .2rem rgba(0,169,157,.25)}.custom-control-input:focus:not(:checked)~.custom-control-label::before{border-color:#2afff0}.custom-control-input:not(:disabled):active~.custom-control-label::before{color:#fff;background-color:#5dfff3;border-color:#5dfff3}.custom-control-input[disabled]~.custom-control-label,.custom-control-input:disabled~.custom-control-label{color:#6c757d}.custom-control-input[disabled]~.custom-control-label::before,.custom-control-input:disabled~.custom-control-label::before{background-color:#e9ecef}.custom-control-label{position:relative;margin-bottom:0;vertical-align:top}.custom-control-label::before{position:absolute;top:.25rem;left:-1.5rem;display:block;width:1rem;height:1rem;pointer-events:none;content:"";background-color:#fff;border:#adb5bd solid 1px}.custom-control-label::after{position:absolute;top:.25rem;left:-1.5rem;display:block;width:1rem;height:1rem;content:"";background:no-repeat 50%/50% 50%}.custom-checkbox .custom-control-label::before{border-radius:.25rem}.custom-checkbox .custom-control-input:checked~.custom-control-label::after{background-image:url%28"data:image/svg+xml,%3csvg xmlns=\'http://www.w3.org/2000/svg\' width=\'8\' height=\'8\' viewBox=\'0 0 8 8\'%3e%3cpath fill=\'%23fff\' d=\'M6.564.75l-3.59 3.612-1.538-1.55L0 4.26l2.974 2.99L8 2.193z\'/%3e%3c/svg%3e"%29}.custom-checkbox .custom-control-input:indeterminate~.custom-control-label::before{border-color:#00a99d;background-color:#00a99d}.custom-checkbox .custom-control-input:indeterminate~.custom-control-label::after{background-image:url%28"data:image/svg+xml,%3csvg xmlns=\'http://www.w3.org/2000/svg\' width=\'4\' height=\'4\' viewBox=\'0 0 4 4\'%3e%3cpath stroke=\'%23fff\' d=\'M0 2h4\'/%3e%3c/svg%3e"%29}.custom-checkbox .custom-control-input:disabled:checked~.custom-control-label::before{background-color:rgba(0,169,157,.5)}.custom-checkbox .custom-control-input:disabled:indeterminate~.custom-control-label::before{background-color:rgba(0,169,157,.5)}.custom-radio .custom-control-label::before{border-radius:50%}.custom-radio .custom-control-input:checked~.custom-control-label::after{background-image:url%28"data:image/svg+xml,%3csvg xmlns=\'http://www.w3.org/2000/svg\' width=\'12\' height=\'12\' viewBox=\'-4 -4 8 8\'%3e%3ccircle r=\'3\' fill=\'%23fff\'/%3e%3c/svg%3e"%29}.custom-radio .custom-control-input:disabled:checked~.custom-control-label::before{background-color:rgba(0,169,157,.5)}.custom-switch{padding-left:2.25rem}.custom-switch .custom-control-label::before{left:-2.25rem;width:1.75rem;pointer-events:all;border-radius:.5rem}.custom-switch .custom-control-label::after{top:calc(0.25rem + 2px);left:calc(-2.25rem + 2px);width:calc(1rem - 4px);height:calc(1rem - 4px);background-color:#adb5bd;border-radius:.5rem;transition:transform .15s ease-in-out,background-color .15s ease-in-out,border-color .15s ease-in-out,box-shadow .15s ease-in-out}@media(prefers-reduced-motion: reduce){.custom-switch .custom-control-label::after{transition:none}}.custom-switch .custom-control-input:checked~.custom-control-label::after{background-color:#fff;transform:translateX(0.75rem)}.custom-switch .custom-control-input:disabled:checked~.custom-control-label::before{background-color:rgba(0,169,157,.5)}.custom-select{display:inline-block;width:100%;height:calc(1.5em + 0.75rem + 2px);padding:.375rem 1.75rem .375rem .75rem;font-size:1rem;font-weight:400;line-height:1.5;color:#495057;vertical-align:middle;background:#fff url("data:image/svg+xml,%3csvg xmlns=\'http://www.w3.org/2000/svg\' width=\'4\' height=\'5\' viewBox=\'0 0 4 5\'%3e%3cpath fill=\'%23343a40\' d=\'M2 0L0 2h4zm0 5L0 3h4z\'/%3e%3c/svg%3e") right .75rem center/8px 10px no-repeat;border:1px solid #ced4da;border-radius:.25rem;appearance:none}.custom-select:focus{border-color:#2afff0;outline:0;box-shadow:0 0 0 .2rem rgba(0,169,157,.25)}.custom-select:focus::-ms-value{color:#495057;background-color:#fff}.custom-select[multiple],.custom-select[size]:not([size="1"]){height:auto;padding-right:.75rem;background-image:none}.custom-select:disabled{color:#6c757d;background-color:#e9ecef}.custom-select::-ms-expand{display:none}.custom-select:-moz-focusring{color:transparent;text-shadow:0 0 0 #495057}.custom-select-sm{height:calc(1.5em + 0.5rem + 2px);padding-top:.25rem;padding-bottom:.25rem;padding-left:.5rem;font-size:0.875rem}.custom-select-lg{height:calc(1.5em + 1rem + 2px);padding-top:.5rem;padding-bottom:.5rem;padding-left:1rem;font-size:1.25rem}.custom-file{position:relative;display:inline-block;width:100%;height:calc(1.5em + 0.75rem + 2px);margin-bottom:0}.custom-file-input{position:relative;z-index:2;width:100%;height:calc(1.5em + 0.75rem + 2px);margin:0;opacity:0}.custom-file-input:focus~.custom-file-label{border-color:#2afff0;box-shadow:0 0 0 .2rem rgba(0,169,157,.25)}.custom-file-input[disabled]~.custom-file-label,.custom-file-input:disabled~.custom-file-label{background-color:#e9ecef}.custom-file-input:lang(en)~.custom-file-label::after{content:"Browse"}.custom-file-input~.custom-file-label[data-browse]::after{content:attr(data-browse)}.custom-file-label{position:absolute;top:0;right:0;left:0;z-index:1;height:calc(1.5em + 0.75rem + 2px);padding:.375rem .75rem;font-weight:400;line-height:1.5;color:#495057;background-color:#fff;border:1px solid #ced4da;border-radius:.25rem}.custom-file-label::after{position:absolute;top:0;right:0;bottom:0;z-index:3;display:block;height:calc(1.5em + 0.75rem);padding:.375rem .75rem;line-height:1.5;color:#495057;content:"Browse";background-color:#e9ecef;border-left:inherit;border-radius:0 .25rem .25rem 0}.custom-range{width:100%;height:1.4rem;padding:0;background-color:transparent;appearance:none}.custom-range:focus{outline:none}.custom-range:focus::-webkit-slider-thumb{box-shadow:0 0 0 1px #fff,0 0 0 .2rem rgba(0,169,157,.25)}.custom-range:focus::-moz-range-thumb{box-shadow:0 0 0 1px #fff,0 0 0 .2rem rgba(0,169,157,.25)}.custom-range:focus::-ms-thumb{box-shadow:0 0 0 1px #fff,0 0 0 .2rem rgba(0,169,157,.25)}.custom-range::-moz-focus-outer{border:0}.custom-range::-webkit-slider-thumb{width:1rem;height:1rem;margin-top:-0.25rem;background-color:#00a99d;border:0;border-radius:1rem;transition:background-color .15s ease-in-out,border-color .15s ease-in-out,box-shadow .15s ease-in-out;appearance:none}@media(prefers-reduced-motion: reduce){.custom-range::-webkit-slider-thumb{transition:none}}.custom-range::-webkit-slider-thumb:active{background-color:#5dfff3}.custom-range::-webkit-slider-runnable-track{width:100%;height:.5rem;color:transparent;cursor:pointer;background-color:#dee2e6;border-color:transparent;border-radius:1rem}.custom-range::-moz-range-thumb{width:1rem;height:1rem;background-color:#00a99d;border:0;border-radius:1rem;transition:background-color .15s ease-in-out,border-color .15s ease-in-out,box-shadow .15s ease-in-out;appearance:none}@media(prefers-reduced-motion: reduce){.custom-range::-moz-range-thumb{transition:none}}.custom-range::-moz-range-thumb:active{background-color:#5dfff3}.custom-range::-moz-range-track{width:100%;height:.5rem;color:transparent;cursor:pointer;background-color:#dee2e6;border-color:transparent;border-radius:1rem}.custom-range::-ms-thumb{width:1rem;height:1rem;margin-top:0;margin-right:.2rem;margin-left:.2rem;background-color:#00a99d;border:0;border-radius:1rem;transition:background-color .15s ease-in-out,border-color .15s ease-in-out,box-shadow .15s ease-in-out;appearance:none}@media(prefers-reduced-motion: reduce){.custom-range::-ms-thumb{transition:none}}.custom-range::-ms-thumb:active{background-color:#5dfff3}.custom-range::-ms-track{width:100%;height:.5rem;color:transparent;cursor:pointer;background-color:transparent;border-color:transparent;border-width:.5rem}.custom-range::-ms-fill-lower{background-color:#dee2e6;border-radius:1rem}.custom-range::-ms-fill-upper{margin-right:15px;background-color:#dee2e6;border-radius:1rem}.custom-range:disabled::-webkit-slider-thumb{background-color:#adb5bd}.custom-range:disabled::-webkit-slider-runnable-track{cursor:default}.custom-range:disabled::-moz-range-thumb{background-color:#adb5bd}.custom-range:disabled::-moz-range-track{cursor:default}.custom-range:disabled::-ms-thumb{background-color:#adb5bd}.custom-control-label::before,.custom-file-label,.custom-select{transition:background-color .15s ease-in-out,border-color .15s ease-in-out,box-shadow .15s ease-in-out}@media(prefers-reduced-motion: reduce){.custom-control-label::before,.custom-file-label,.custom-select{transition:none}}.nav{display:flex;flex-wrap:wrap;padding-left:0;margin-bottom:0;list-style:none}.nav-link{display:block;padding:.5rem 1rem}.nav-link:hover,.nav-link:focus{text-decoration:none}.nav-link.disabled{color:#6c757d;pointer-events:none;cursor:default}.nav-tabs{border-bottom:1px solid #dee2e6}.nav-tabs .nav-item{margin-bottom:-1px}.nav-tabs .nav-link{border:1px solid transparent;border-top-left-radius:.25rem;border-top-right-radius:.25rem}.nav-tabs .nav-link:hover,.nav-tabs .nav-link:focus{border-color:#e9ecef #e9ecef #dee2e6}.nav-tabs .nav-link.disabled{color:#6c757d;background-color:transparent;border-color:transparent}.nav-tabs .nav-link.active,.nav-tabs .nav-item.show .nav-link{color:#495057;background-color:#fff;border-color:#dee2e6 #dee2e6 #fff}.nav-tabs .dropdown-menu{margin-top:-1px;border-top-left-radius:0;border-top-right-radius:0}.nav-pills .nav-link{border-radius:.25rem}.nav-pills .nav-link.active,.nav-pills .show>.nav-link{color:#fff;background-color:#00a99d}.nav-fill .nav-item{flex:1 1 auto;text-align:center}.nav-justified .nav-item{flex-basis:0;flex-grow:1;text-align:center}.tab-content>.tab-pane{display:none}.tab-content>.active{display:block}.navbar{position:relative;display:flex;flex-wrap:wrap;align-items:center;justify-content:space-between;padding:.5rem 1rem}.navbar .container,.navbar .container-fluid,.navbar .container-sm,.navbar .container-md,.navbar .container-lg,.navbar .container-xl{display:flex;flex-wrap:wrap;align-items:center;justify-content:space-between}.navbar-brand{display:inline-block;padding-top:.3125rem;padding-bottom:.3125rem;margin-right:1rem;font-size:1.25rem;line-height:inherit;white-space:nowrap}.navbar-brand:hover,.navbar-brand:focus{text-decoration:none}.navbar-nav{display:flex;flex-direction:column;padding-left:0;margin-bottom:0;list-style:none}.navbar-nav .nav-link{padding-right:0;padding-left:0}.navbar-nav .dropdown-menu{position:static;float:none}.navbar-text{display:inline-block;padding-top:.5rem;padding-bottom:.5rem}.navbar-collapse{flex-basis:100%;flex-grow:1;align-items:center}.navbar-toggler{padding:.25rem .75rem;font-size:1.25rem;line-height:1;background-color:transparent;border:1px solid transparent;border-radius:.25rem}.navbar-toggler:hover,.navbar-toggler:focus{text-decoration:none}.navbar-toggler-icon{display:inline-block;width:1.5em;height:1.5em;vertical-align:middle;content:"";background:no-repeat center center;background-size:100% 100%}@media(max-width: 575.98px){.navbar-expand-sm>.container,.navbar-expand-sm>.container-fluid,.navbar-expand-sm>.container-sm,.navbar-expand-sm>.container-md,.navbar-expand-sm>.container-lg,.navbar-expand-sm>.container-xl{padding-right:0;padding-left:0}}@media(min-width: 576px){.navbar-expand-sm{flex-flow:row nowrap;justify-content:flex-start}.navbar-expand-sm .navbar-nav{flex-direction:row}.navbar-expand-sm .navbar-nav .dropdown-menu{position:absolute}.navbar-expand-sm .navbar-nav .nav-link{padding-right:.5rem;padding-left:.5rem}.navbar-expand-sm>.container,.navbar-expand-sm>.container-fluid,.navbar-expand-sm>.container-sm,.navbar-expand-sm>.container-md,.navbar-expand-sm>.container-lg,.navbar-expand-sm>.container-xl{flex-wrap:nowrap}.navbar-expand-sm .navbar-collapse{display:flex !important;flex-basis:auto}.navbar-expand-sm .navbar-toggler{display:none}}@media(max-width: 767.98px){.navbar-expand-md>.container,.navbar-expand-md>.container-fluid,.navbar-expand-md>.container-sm,.navbar-expand-md>.container-md,.navbar-expand-md>.container-lg,.navbar-expand-md>.container-xl{padding-right:0;padding-left:0}}@media(min-width: 768px){.navbar-expand-md{flex-flow:row nowrap;justify-content:flex-start}.navbar-expand-md .navbar-nav{flex-direction:row}.navbar-expand-md .navbar-nav .dropdown-menu{position:absolute}.navbar-expand-md .navbar-nav .nav-link{padding-right:.5rem;padding-left:.5rem}.navbar-expand-md>.container,.navbar-expand-md>.container-fluid,.navbar-expand-md>.container-sm,.navbar-expand-md>.container-md,.navbar-expand-md>.container-lg,.navbar-expand-md>.container-xl{flex-wrap:nowrap}.navbar-expand-md .navbar-collapse{display:flex !important;flex-basis:auto}.navbar-expand-md .navbar-toggler{display:none}}@media(max-width: 991.98px){.navbar-expand-lg>.container,.navbar-expand-lg>.container-fluid,.navbar-expand-lg>.container-sm,.navbar-expand-lg>.container-md,.navbar-expand-lg>.container-lg,.navbar-expand-lg>.container-xl{padding-right:0;padding-left:0}}@media(min-width: 992px){.navbar-expand-lg{flex-flow:row nowrap;justify-content:flex-start}.navbar-expand-lg .navbar-nav{flex-direction:row}.navbar-expand-lg .navbar-nav .dropdown-menu{position:absolute}.navbar-expand-lg .navbar-nav .nav-link{padding-right:.5rem;padding-left:.5rem}.navbar-expand-lg>.container,.navbar-expand-lg>.container-fluid,.navbar-expand-lg>.container-sm,.navbar-expand-lg>.container-md,.navbar-expand-lg>.container-lg,.navbar-expand-lg>.container-xl{flex-wrap:nowrap}.navbar-expand-lg .navbar-collapse{display:flex !important;flex-basis:auto}.navbar-expand-lg .navbar-toggler{display:none}}@media(max-width: 1199.98px){.navbar-expand-xl>.container,.navbar-expand-xl>.container-fluid,.navbar-expand-xl>.container-sm,.navbar-expand-xl>.container-md,.navbar-expand-xl>.container-lg,.navbar-expand-xl>.container-xl{padding-right:0;padding-left:0}}@media(min-width: 1200px){.navbar-expand-xl{flex-flow:row nowrap;justify-content:flex-start}.navbar-expand-xl .navbar-nav{flex-direction:row}.navbar-expand-xl .navbar-nav .dropdown-menu{position:absolute}.navbar-expand-xl .navbar-nav .nav-link{padding-right:.5rem;padding-left:.5rem}.navbar-expand-xl>.container,.navbar-expand-xl>.container-fluid,.navbar-expand-xl>.container-sm,.navbar-expand-xl>.container-md,.navbar-expand-xl>.container-lg,.navbar-expand-xl>.container-xl{flex-wrap:nowrap}.navbar-expand-xl .navbar-collapse{display:flex !important;flex-basis:auto}.navbar-expand-xl .navbar-toggler{display:none}}.navbar-expand{flex-flow:row nowrap;justify-content:flex-start}.navbar-expand>.container,.navbar-expand>.container-fluid,.navbar-expand>.container-sm,.navbar-expand>.container-md,.navbar-expand>.container-lg,.navbar-expand>.container-xl{padding-right:0;padding-left:0}.navbar-expand .navbar-nav{flex-direction:row}.navbar-expand .navbar-nav .dropdown-menu{position:absolute}.navbar-expand .navbar-nav .nav-link{padding-right:.5rem;padding-left:.5rem}.navbar-expand>.container,.navbar-expand>.container-fluid,.navbar-expand>.container-sm,.navbar-expand>.container-md,.navbar-expand>.container-lg,.navbar-expand>.container-xl{flex-wrap:nowrap}.navbar-expand .navbar-collapse{display:flex !important;flex-basis:auto}.navbar-expand .navbar-toggler{display:none}.navbar-light .navbar-brand{color:rgba(0,0,0,.9)}.navbar-light .navbar-brand:hover,.navbar-light .navbar-brand:focus{color:rgba(0,0,0,.9)}.navbar-light .navbar-nav .nav-link{color:rgba(0,0,0,.5)}.navbar-light .navbar-nav .nav-link:hover,.navbar-light .navbar-nav .nav-link:focus{color:rgba(0,0,0,.7)}.navbar-light .navbar-nav .nav-link.disabled{color:rgba(0,0,0,.3)}.navbar-light .navbar-nav .show>.nav-link,.navbar-light .navbar-nav .active>.nav-link,.navbar-light .navbar-nav .nav-link.show,.navbar-light .navbar-nav .nav-link.active{color:rgba(0,0,0,.9)}.navbar-light .navbar-toggler{color:rgba(0,0,0,.5);border-color:rgba(0,0,0,.1)}.navbar-light .navbar-toggler-icon{background-image:url%28"data:image/svg+xml,%3csvg xmlns=\'http://www.w3.org/2000/svg\' width=\'30\' height=\'30\' viewBox=\'0 0 30 30\'%3e%3cpath stroke=\'rgba%280, 0, 0, 0.5%29\' stroke-linecap=\'round\' stroke-miterlimit=\'10\' stroke-width=\'2\' d=\'M4 7h22M4 15h22M4 23h22\'/%3e%3c/svg%3e"%29}.navbar-light .navbar-text{color:rgba(0,0,0,.5)}.navbar-light .navbar-text a{color:rgba(0,0,0,.9)}.navbar-light .navbar-text a:hover,.navbar-light .navbar-text a:focus{color:rgba(0,0,0,.9)}.navbar-dark .navbar-brand{color:#fff}.navbar-dark .navbar-brand:hover,.navbar-dark .navbar-brand:focus{color:#fff}.navbar-dark .navbar-nav .nav-link{color:rgba(255,255,255,.5)}.navbar-dark .navbar-nav .nav-link:hover,.navbar-dark .navbar-nav .nav-link:focus{color:rgba(255,255,255,.75)}.navbar-dark .navbar-nav .nav-link.disabled{color:rgba(255,255,255,.25)}.navbar-dark .navbar-nav .show>.nav-link,.navbar-dark .navbar-nav .active>.nav-link,.navbar-dark .navbar-nav .nav-link.show,.navbar-dark .navbar-nav .nav-link.active{color:#fff}.navbar-dark .navbar-toggler{color:rgba(255,255,255,.5);border-color:rgba(255,255,255,.1)}.navbar-dark .navbar-toggler-icon{background-image:url%28"data:image/svg+xml,%3csvg xmlns=\'http://www.w3.org/2000/svg\' width=\'30\' height=\'30\' viewBox=\'0 0 30 30\'%3e%3cpath stroke=\'rgba%28255, 255, 255, 0.5%29\' stroke-linecap=\'round\' stroke-miterlimit=\'10\' stroke-width=\'2\' d=\'M4 7h22M4 15h22M4 23h22\'/%3e%3c/svg%3e"%29}.navbar-dark .navbar-text{color:rgba(255,255,255,.5)}.navbar-dark .navbar-text a{color:#fff}.navbar-dark .navbar-text a:hover,.navbar-dark .navbar-text a:focus{color:#fff}.card{position:relative;display:flex;flex-direction:column;min-width:0;word-wrap:break-word;background-color:#fff;background-clip:border-box;border:1px solid rgba(0,0,0,.125);border-radius:.25rem}.card>hr{margin-right:0;margin-left:0}.card>.list-group:first-child .list-group-item:first-child{border-top-left-radius:.25rem;border-top-right-radius:.25rem}.card>.list-group:last-child .list-group-item:last-child{border-bottom-right-radius:.25rem;border-bottom-left-radius:.25rem}.card-body{flex:1 1 auto;min-height:1px;padding:1.25rem}.card-title{margin-bottom:.75rem}.card-subtitle{margin-top:-0.375rem;margin-bottom:0}.card-text:last-child{margin-bottom:0}.card-link:hover{text-decoration:none}.card-link+.card-link{margin-left:1.25rem}.card-header{padding:.75rem 1.25rem;margin-bottom:0;background-color:rgba(0,0,0,.03);border-bottom:1px solid rgba(0,0,0,.125)}.card-header:first-child{border-radius:calc(0.25rem - 1px) calc(0.25rem - 1px) 0 0}.card-header+.list-group .list-group-item:first-child{border-top:0}.card-footer{padding:.75rem 1.25rem;background-color:rgba(0,0,0,.03);border-top:1px solid rgba(0,0,0,.125)}.card-footer:last-child{border-radius:0 0 calc(0.25rem - 1px) calc(0.25rem - 1px)}.card-header-tabs{margin-right:-0.625rem;margin-bottom:-0.75rem;margin-left:-0.625rem;border-bottom:0}.card-header-pills{margin-right:-0.625rem;margin-left:-0.625rem}.card-img-overlay{position:absolute;top:0;right:0;bottom:0;left:0;padding:1.25rem}.card-img,.card-img-top,.card-img-bottom{flex-shrink:0;width:100%}.card-img,.card-img-top{border-top-left-radius:calc(0.25rem - 1px);border-top-right-radius:calc(0.25rem - 1px)}.card-img,.card-img-bottom{border-bottom-right-radius:calc(0.25rem - 1px);border-bottom-left-radius:calc(0.25rem - 1px)}.card-deck .card{margin-bottom:15px}@media(min-width: 576px){.card-deck{display:flex;flex-flow:row wrap;margin-right:-15px;margin-left:-15px}.card-deck .card{flex:1 0 0%;margin-right:15px;margin-bottom:0;margin-left:15px}}.card-group>.card{margin-bottom:15px}@media(min-width: 576px){.card-group{display:flex;flex-flow:row wrap}.card-group>.card{flex:1 0 0%;margin-bottom:0}.card-group>.card+.card{margin-left:0;border-left:0}.card-group>.card:not(:last-child){border-top-right-radius:0;border-bottom-right-radius:0}.card-group>.card:not(:last-child) .card-img-top,.card-group>.card:not(:last-child) .card-header{border-top-right-radius:0}.card-group>.card:not(:last-child) .card-img-bottom,.card-group>.card:not(:last-child) .card-footer{border-bottom-right-radius:0}.card-group>.card:not(:first-child){border-top-left-radius:0;border-bottom-left-radius:0}.card-group>.card:not(:first-child) .card-img-top,.card-group>.card:not(:first-child) .card-header{border-top-left-radius:0}.card-group>.card:not(:first-child) .card-img-bottom,.card-group>.card:not(:first-child) .card-footer{border-bottom-left-radius:0}}.card-columns .card{margin-bottom:.75rem}@media(min-width: 576px){.card-columns{column-count:3;column-gap:1.25rem;orphans:1;widows:1}.card-columns .card{display:inline-block;width:100%}}.accordion>.card{overflow:hidden}.accordion>.card:not(:last-of-type){border-bottom:0;border-bottom-right-radius:0;border-bottom-left-radius:0}.accordion>.card:not(:first-of-type){border-top-left-radius:0;border-top-right-radius:0}.accordion>.card>.card-header{border-radius:0;margin-bottom:-1px}.breadcrumb{display:flex;flex-wrap:wrap;padding:.75rem 1rem;margin-bottom:1rem;list-style:none;background-color:#e9ecef;border-radius:.25rem}.breadcrumb-item+.breadcrumb-item{padding-left:.5rem}.breadcrumb-item+.breadcrumb-item::before{display:inline-block;padding-right:.5rem;color:#6c757d;content:"/"}.breadcrumb-item+.breadcrumb-item:hover::before{text-decoration:underline}.breadcrumb-item+.breadcrumb-item:hover::before{text-decoration:none}.breadcrumb-item.active{color:#6c757d}.pagination{display:flex;padding-left:0;list-style:none;border-radius:.25rem}.page-link{position:relative;display:block;padding:.5rem .75rem;margin-left:-1px;line-height:1.25;color:#00a99d;background-color:#fff;border:1px solid #dee2e6}.page-link:hover{z-index:2;color:#005d56;text-decoration:none;background-color:#e9ecef;border-color:#dee2e6}.page-link:focus{z-index:3;outline:0;box-shadow:0 0 0 .2rem rgba(0,169,157,.25)}.page-item:first-child .page-link{margin-left:0;border-top-left-radius:.25rem;border-bottom-left-radius:.25rem}.page-item:last-child .page-link{border-top-right-radius:.25rem;border-bottom-right-radius:.25rem}.page-item.active .page-link{z-index:3;color:#fff;background-color:#00a99d;border-color:#00a99d}.page-item.disabled .page-link{color:#6c757d;pointer-events:none;cursor:auto;background-color:#fff;border-color:#dee2e6}.pagination-lg .page-link{padding:.75rem 1.5rem;font-size:1.25rem;line-height:1.5}.pagination-lg .page-item:first-child .page-link{border-top-left-radius:.3rem;border-bottom-left-radius:.3rem}.pagination-lg .page-item:last-child .page-link{border-top-right-radius:.3rem;border-bottom-right-radius:.3rem}.pagination-sm .page-link{padding:.25rem .5rem;font-size:0.875rem;line-height:1.5}.pagination-sm .page-item:first-child .page-link{border-top-left-radius:.2rem;border-bottom-left-radius:.2rem}.pagination-sm .page-item:last-child .page-link{border-top-right-radius:.2rem;border-bottom-right-radius:.2rem}.badge{display:inline-block;padding:.25em .4em;font-size:75%;font-weight:700;line-height:1;text-align:center;white-space:nowrap;vertical-align:baseline;border-radius:.25rem;transition:color .15s ease-in-out,background-color .15s ease-in-out,border-color .15s ease-in-out,box-shadow .15s ease-in-out}@media(prefers-reduced-motion: reduce){.badge{transition:none}}a.badge:hover,a.badge:focus{text-decoration:none}.badge:empty{display:none}.btn .badge{position:relative;top:-1px}.badge-pill{padding-right:.6em;padding-left:.6em;border-radius:10rem}.badge-primary{color:#fff;background-color:#00a99d}a.badge-primary:hover,a.badge-primary:focus{color:#fff;background-color:#00766e}a.badge-primary:focus,a.badge-primary.focus{outline:0;box-shadow:0 0 0 .2rem rgba(0,169,157,.5)}.badge-secondary{color:#fff;background-color:#6c757d}a.badge-secondary:hover,a.badge-secondary:focus{color:#fff;background-color:#545b62}a.badge-secondary:focus,a.badge-secondary.focus{outline:0;box-shadow:0 0 0 .2rem rgba(108,117,125,.5)}.badge-success{color:#fff;background-color:#28a745}a.badge-success:hover,a.badge-success:focus{color:#fff;background-color:#1e7e34}a.badge-success:focus,a.badge-success.focus{outline:0;box-shadow:0 0 0 .2rem rgba(40,167,69,.5)}.badge-info{color:#fff;background-color:#17a2b8}a.badge-info:hover,a.badge-info:focus{color:#fff;background-color:#117a8b}a.badge-info:focus,a.badge-info.focus{outline:0;box-shadow:0 0 0 .2rem rgba(23,162,184,.5)}.badge-warning{color:#212529;background-color:#ffc107}a.badge-warning:hover,a.badge-warning:focus{color:#212529;background-color:#d39e00}a.badge-warning:focus,a.badge-warning.focus{outline:0;box-shadow:0 0 0 .2rem rgba(255,193,7,.5)}.badge-danger{color:#fff;background-color:#dc3545}a.badge-danger:hover,a.badge-danger:focus{color:#fff;background-color:#bd2130}a.badge-danger:focus,a.badge-danger.focus{outline:0;box-shadow:0 0 0 .2rem rgba(220,53,69,.5)}.badge-light{color:#212529;background-color:#f8f9fa}a.badge-light:hover,a.badge-light:focus{color:#212529;background-color:#dae0e5}a.badge-light:focus,a.badge-light.focus{outline:0;box-shadow:0 0 0 .2rem rgba(248,249,250,.5)}.badge-dark{color:#fff;background-color:#343a40}a.badge-dark:hover,a.badge-dark:focus{color:#fff;background-color:#1d2124}a.badge-dark:focus,a.badge-dark.focus{outline:0;box-shadow:0 0 0 .2rem rgba(52,58,64,.5)}.jumbotron{padding:2rem 1rem;margin-bottom:2rem;background-color:#e9ecef;border-radius:.3rem}@media(min-width: 576px){.jumbotron{padding:4rem 2rem}}.jumbotron-fluid{padding-right:0;padding-left:0;border-radius:0}.alert{position:relative;padding:.75rem 1.25rem;margin-bottom:1rem;border:1px solid transparent;border-radius:.25rem}.alert-heading{color:inherit}.alert-link{font-weight:700}.alert-dismissible{padding-right:4rem}.alert-dismissible .close{position:absolute;top:0;right:0;padding:.75rem 1.25rem;color:inherit}.alert-primary{color:#005852;background-color:#cceeeb;border-color:#b8e7e4}.alert-primary hr{border-top-color:#a5e1dd}.alert-primary .alert-link{color:#002522}.alert-secondary{color:#383d41;background-color:#e2e3e5;border-color:#d6d8db}.alert-secondary hr{border-top-color:#c8cbcf}.alert-secondary .alert-link{color:#202326}.alert-success{color:#155724;background-color:#d4edda;border-color:#c3e6cb}.alert-success hr{border-top-color:#b1dfbb}.alert-success .alert-link{color:#0b2e13}.alert-info{color:#0c5460;background-color:#d1ecf1;border-color:#bee5eb}.alert-info hr{border-top-color:#abdde5}.alert-info .alert-link{color:#062c33}.alert-warning{color:#856404;background-color:#fff3cd;border-color:#ffeeba}.alert-warning hr{border-top-color:#ffe8a1}.alert-warning .alert-link{color:#533f03}.alert-danger{color:#721c24;background-color:#f8d7da;border-color:#f5c6cb}.alert-danger hr{border-top-color:#f1b0b7}.alert-danger .alert-link{color:#491217}.alert-light{color:#818182;background-color:#fefefe;border-color:#fdfdfe}.alert-light hr{border-top-color:#ececf6}.alert-light .alert-link{color:#686868}.alert-dark{color:#1b1e21;background-color:#d6d8d9;border-color:#c6c8ca}.alert-dark hr{border-top-color:#b9bbbe}.alert-dark .alert-link{color:#040505}@keyframes progress-bar-stripes{from{background-position:1rem 0}to{background-position:0 0}}.progress{display:flex;height:1rem;overflow:hidden;font-size:0.75rem;background-color:#e9ecef;border-radius:.25rem}.progress-bar{display:flex;flex-direction:column;justify-content:center;overflow:hidden;color:#fff;text-align:center;white-space:nowrap;background-color:#00a99d;transition:width .6s ease}@media(prefers-reduced-motion: reduce){.progress-bar{transition:none}}.progress-bar-striped{background-image:linear-gradient(45deg, rgba(255, 255, 255, 0.15) 25%, transparent 25%, transparent 50%, rgba(255, 255, 255, 0.15) 50%, rgba(255, 255, 255, 0.15) 75%, transparent 75%, transparent);background-size:1rem 1rem}.progress-bar-animated{animation:progress-bar-stripes 1s linear infinite}@media(prefers-reduced-motion: reduce){.progress-bar-animated{animation:none}}.media{display:flex;align-items:flex-start}.media-body{flex:1}.list-group{display:flex;flex-direction:column;padding-left:0;margin-bottom:0}.list-group-item-action{width:100%;color:#495057;text-align:inherit}.list-group-item-action:hover,.list-group-item-action:focus{z-index:1;color:#495057;text-decoration:none;background-color:#f8f9fa}.list-group-item-action:active{color:#212529;background-color:#e9ecef}.list-group-item{position:relative;display:block;padding:.75rem 1.25rem;background-color:#fff;border:1px solid rgba(0,0,0,.125)}.list-group-item:first-child{border-top-left-radius:.25rem;border-top-right-radius:.25rem}.list-group-item:last-child{border-bottom-right-radius:.25rem;border-bottom-left-radius:.25rem}.list-group-item.disabled,.list-group-item:disabled{color:#6c757d;pointer-events:none;background-color:#fff}.list-group-item.active{z-index:2;color:#fff;background-color:#00a99d;border-color:#00a99d}.list-group-item+.list-group-item{border-top-width:0}.list-group-item+.list-group-item.active{margin-top:-1px;border-top-width:1px}.list-group-horizontal{flex-direction:row}.list-group-horizontal .list-group-item:first-child{border-bottom-left-radius:.25rem;border-top-right-radius:0}.list-group-horizontal .list-group-item:last-child{border-top-right-radius:.25rem;border-bottom-left-radius:0}.list-group-horizontal .list-group-item.active{margin-top:0}.list-group-horizontal .list-group-item+.list-group-item{border-top-width:1px;border-left-width:0}.list-group-horizontal .list-group-item+.list-group-item.active{margin-left:-1px;border-left-width:1px}@media(min-width: 576px){.list-group-horizontal-sm{flex-direction:row}.list-group-horizontal-sm .list-group-item:first-child{border-bottom-left-radius:.25rem;border-top-right-radius:0}.list-group-horizontal-sm .list-group-item:last-child{border-top-right-radius:.25rem;border-bottom-left-radius:0}.list-group-horizontal-sm .list-group-item.active{margin-top:0}.list-group-horizontal-sm .list-group-item+.list-group-item{border-top-width:1px;border-left-width:0}.list-group-horizontal-sm .list-group-item+.list-group-item.active{margin-left:-1px;border-left-width:1px}}@media(min-width: 768px){.list-group-horizontal-md{flex-direction:row}.list-group-horizontal-md .list-group-item:first-child{border-bottom-left-radius:.25rem;border-top-right-radius:0}.list-group-horizontal-md .list-group-item:last-child{border-top-right-radius:.25rem;border-bottom-left-radius:0}.list-group-horizontal-md .list-group-item.active{margin-top:0}.list-group-horizontal-md .list-group-item+.list-group-item{border-top-width:1px;border-left-width:0}.list-group-horizontal-md .list-group-item+.list-group-item.active{margin-left:-1px;border-left-width:1px}}@media(min-width: 992px){.list-group-horizontal-lg{flex-direction:row}.list-group-horizontal-lg .list-group-item:first-child{border-bottom-left-radius:.25rem;border-top-right-radius:0}.list-group-horizontal-lg .list-group-item:last-child{border-top-right-radius:.25rem;border-bottom-left-radius:0}.list-group-horizontal-lg .list-group-item.active{margin-top:0}.list-group-horizontal-lg .list-group-item+.list-group-item{border-top-width:1px;border-left-width:0}.list-group-horizontal-lg .list-group-item+.list-group-item.active{margin-left:-1px;border-left-width:1px}}@media(min-width: 1200px){.list-group-horizontal-xl{flex-direction:row}.list-group-horizontal-xl .list-group-item:first-child{border-bottom-left-radius:.25rem;border-top-right-radius:0}.list-group-horizontal-xl .list-group-item:last-child{border-top-right-radius:.25rem;border-bottom-left-radius:0}.list-group-horizontal-xl .list-group-item.active{margin-top:0}.list-group-horizontal-xl .list-group-item+.list-group-item{border-top-width:1px;border-left-width:0}.list-group-horizontal-xl .list-group-item+.list-group-item.active{margin-left:-1px;border-left-width:1px}}.list-group-flush .list-group-item{border-right-width:0;border-left-width:0;border-radius:0}.list-group-flush .list-group-item:first-child{border-top-width:0}.list-group-flush:last-child .list-group-item:last-child{border-bottom-width:0}.list-group-item-primary{color:#005852;background-color:#b8e7e4}.list-group-item-primary.list-group-item-action:hover,.list-group-item-primary.list-group-item-action:focus{color:#005852;background-color:#a5e1dd}.list-group-item-primary.list-group-item-action.active{color:#fff;background-color:#005852;border-color:#005852}.list-group-item-secondary{color:#383d41;background-color:#d6d8db}.list-group-item-secondary.list-group-item-action:hover,.list-group-item-secondary.list-group-item-action:focus{color:#383d41;background-color:#c8cbcf}.list-group-item-secondary.list-group-item-action.active{color:#fff;background-color:#383d41;border-color:#383d41}.list-group-item-success{color:#155724;background-color:#c3e6cb}.list-group-item-success.list-group-item-action:hover,.list-group-item-success.list-group-item-action:focus{color:#155724;background-color:#b1dfbb}.list-group-item-success.list-group-item-action.active{color:#fff;background-color:#155724;border-color:#155724}.list-group-item-info{color:#0c5460;background-color:#bee5eb}.list-group-item-info.list-group-item-action:hover,.list-group-item-info.list-group-item-action:focus{color:#0c5460;background-color:#abdde5}.list-group-item-info.list-group-item-action.active{color:#fff;background-color:#0c5460;border-color:#0c5460}.list-group-item-warning{color:#856404;background-color:#ffeeba}.list-group-item-warning.list-group-item-action:hover,.list-group-item-warning.list-group-item-action:focus{color:#856404;background-color:#ffe8a1}.list-group-item-warning.list-group-item-action.active{color:#fff;background-color:#856404;border-color:#856404}.list-group-item-danger{color:#721c24;background-color:#f5c6cb}.list-group-item-danger.list-group-item-action:hover,.list-group-item-danger.list-group-item-action:focus{color:#721c24;background-color:#f1b0b7}.list-group-item-danger.list-group-item-action.active{color:#fff;background-color:#721c24;border-color:#721c24}.list-group-item-light{color:#818182;background-color:#fdfdfe}.list-group-item-light.list-group-item-action:hover,.list-group-item-light.list-group-item-action:focus{color:#818182;background-color:#ececf6}.list-group-item-light.list-group-item-action.active{color:#fff;background-color:#818182;border-color:#818182}.list-group-item-dark{color:#1b1e21;background-color:#c6c8ca}.list-group-item-dark.list-group-item-action:hover,.list-group-item-dark.list-group-item-action:focus{color:#1b1e21;background-color:#b9bbbe}.list-group-item-dark.list-group-item-action.active{color:#fff;background-color:#1b1e21;border-color:#1b1e21}.close{float:right;font-size:1.5rem;font-weight:700;line-height:1;color:#000;text-shadow:0 1px 0 #fff;opacity:.5}.close:hover{color:#000;text-decoration:none}.close:not(:disabled):not(.disabled):hover,.close:not(:disabled):not(.disabled):focus{opacity:.75}button.close{padding:0;background-color:transparent;border:0;appearance:none}a.close.disabled{pointer-events:none}.toast{max-width:350px;overflow:hidden;font-size:0.875rem;background-color:rgba(255,255,255,.85);background-clip:padding-box;border:1px solid rgba(0,0,0,.1);box-shadow:0 .25rem .75rem rgba(0,0,0,.1);backdrop-filter:blur(10px);opacity:0;border-radius:.25rem}.toast:not(:last-child){margin-bottom:.75rem}.toast.showing{opacity:1}.toast.show{display:block;opacity:1}.toast.hide{display:none}.toast-header{display:flex;align-items:center;padding:.25rem .75rem;color:#6c757d;background-color:rgba(255,255,255,.85);background-clip:padding-box;border-bottom:1px solid rgba(0,0,0,.05)}.toast-body{padding:.75rem}.modal-open{overflow:hidden}.modal-open .modal{overflow-x:hidden;overflow-y:auto}.modal{position:fixed;top:0;left:0;z-index:1050;display:none;width:100%;height:100%;overflow:hidden;outline:0}.modal-dialog{position:relative;width:auto;margin:.5rem;pointer-events:none}.modal.fade .modal-dialog{transition:transform .3s ease-out;transform:translate(0, -50px)}@media(prefers-reduced-motion: reduce){.modal.fade .modal-dialog{transition:none}}.modal.show .modal-dialog{transform:none}.modal.modal-static .modal-dialog{transform:scale(1.02)}.modal-dialog-scrollable{display:flex;max-height:calc(100% - 1rem)}.modal-dialog-scrollable .modal-content{max-height:calc(100vh - 1rem);overflow:hidden}.modal-dialog-scrollable .modal-header,.modal-dialog-scrollable .modal-footer{flex-shrink:0}.modal-dialog-scrollable .modal-body{overflow-y:auto}.modal-dialog-centered{display:flex;align-items:center;min-height:calc(100% - 1rem)}.modal-dialog-centered::before{display:block;height:calc(100vh - 1rem);content:""}.modal-dialog-centered.modal-dialog-scrollable{flex-direction:column;justify-content:center;height:100%}.modal-dialog-centered.modal-dialog-scrollable .modal-content{max-height:none}.modal-dialog-centered.modal-dialog-scrollable::before{content:none}.modal-content{position:relative;display:flex;flex-direction:column;width:100%;pointer-events:auto;background-color:#fff;background-clip:padding-box;border:1px solid rgba(0,0,0,.2);border-radius:.3rem;outline:0}.modal-backdrop{position:fixed;top:0;left:0;z-index:1040;width:100vw;height:100vh;background-color:#000}.modal-backdrop.fade{opacity:0}.modal-backdrop.show{opacity:.5}.modal-header{display:flex;align-items:flex-start;justify-content:space-between;padding:1rem 1rem;border-bottom:1px solid #dee2e6;border-top-left-radius:calc(0.3rem - 1px);border-top-right-radius:calc(0.3rem - 1px)}.modal-header .close{padding:1rem 1rem;margin:-1rem -1rem -1rem auto}.modal-title{margin-bottom:0;line-height:1.5}.modal-body{position:relative;flex:1 1 auto;padding:1rem}.modal-footer{display:flex;flex-wrap:wrap;align-items:center;justify-content:flex-end;padding:.75rem;border-top:1px solid #dee2e6;border-bottom-right-radius:calc(0.3rem - 1px);border-bottom-left-radius:calc(0.3rem - 1px)}.modal-footer>*{margin:.25rem}.modal-scrollbar-measure{position:absolute;top:-9999px;width:50px;height:50px;overflow:scroll}@media(min-width: 576px){.modal-dialog{max-width:500px;margin:1.75rem auto}.modal-dialog-scrollable{max-height:calc(100% - 3.5rem)}.modal-dialog-scrollable .modal-content{max-height:calc(100vh - 3.5rem)}.modal-dialog-centered{min-height:calc(100% - 3.5rem)}.modal-dialog-centered::before{height:calc(100vh - 3.5rem)}.modal-sm{max-width:300px}}@media(min-width: 992px){.modal-lg,.modal-xl{max-width:800px}}@media(min-width: 1200px){.modal-xl{max-width:1140px}}.tooltip{position:absolute;z-index:1070;display:block;margin:0;font-family:-apple-system,BlinkMacSystemFont,"Segoe UI",Roboto,"Helvetica Neue",Arial,"Noto Sans","Liberation Sans",sans-serif,"Apple Color Emoji","Segoe UI Emoji","Segoe UI Symbol","Noto Color Emoji";font-style:normal;font-weight:400;line-height:1.5;text-align:left;text-align:start;text-decoration:none;text-shadow:none;text-transform:none;letter-spacing:normal;word-break:normal;word-spacing:normal;white-space:normal;line-break:auto;font-size:0.875rem;word-wrap:break-word;opacity:0}.tooltip.show{opacity:.9}.tooltip .arrow{position:absolute;display:block;width:.8rem;height:.4rem}.tooltip .arrow::before{position:absolute;content:"";border-color:transparent;border-style:solid}.bs-tooltip-top,.bs-tooltip-auto[x-placement^=top]{padding:.4rem 0}.bs-tooltip-top .arrow,.bs-tooltip-auto[x-placement^=top] .arrow{bottom:0}.bs-tooltip-top .arrow::before,.bs-tooltip-auto[x-placement^=top] .arrow::before{top:0;border-width:.4rem .4rem 0;border-top-color:#000}.bs-tooltip-right,.bs-tooltip-auto[x-placement^=right]{padding:0 .4rem}.bs-tooltip-right .arrow,.bs-tooltip-auto[x-placement^=right] .arrow{left:0;width:.4rem;height:.8rem}.bs-tooltip-right .arrow::before,.bs-tooltip-auto[x-placement^=right] .arrow::before{right:0;border-width:.4rem .4rem .4rem 0;border-right-color:#000}.bs-tooltip-bottom,.bs-tooltip-auto[x-placement^=bottom]{padding:.4rem 0}.bs-tooltip-bottom .arrow,.bs-tooltip-auto[x-placement^=bottom] .arrow{top:0}.bs-tooltip-bottom .arrow::before,.bs-tooltip-auto[x-placement^=bottom] .arrow::before{bottom:0;border-width:0 .4rem .4rem;border-bottom-color:#000}.bs-tooltip-left,.bs-tooltip-auto[x-placement^=left]{padding:0 .4rem}.bs-tooltip-left .arrow,.bs-tooltip-auto[x-placement^=left] .arrow{right:0;width:.4rem;height:.8rem}.bs-tooltip-left .arrow::before,.bs-tooltip-auto[x-placement^=left] .arrow::before{left:0;border-width:.4rem 0 .4rem .4rem;border-left-color:#000}.tooltip-inner{max-width:200px;padding:.25rem .5rem;color:#fff;text-align:center;background-color:#000;border-radius:.25rem}.popover{position:absolute;top:0;left:0;z-index:1060;display:block;max-width:276px;font-family:-apple-system,BlinkMacSystemFont,"Segoe UI",Roboto,"Helvetica Neue",Arial,"Noto Sans","Liberation Sans",sans-serif,"Apple Color Emoji","Segoe UI Emoji","Segoe UI Symbol","Noto Color Emoji";font-style:normal;font-weight:400;line-height:1.5;text-align:left;text-align:start;text-decoration:none;text-shadow:none;text-transform:none;letter-spacing:normal;word-break:normal;word-spacing:normal;white-space:normal;line-break:auto;font-size:0.875rem;word-wrap:break-word;background-color:#fff;background-clip:padding-box;border:1px solid rgba(0,0,0,.2);border-radius:.3rem}.popover .arrow{position:absolute;display:block;width:1rem;height:.5rem;margin:0 .3rem}.popover .arrow::before,.popover .arrow::after{position:absolute;display:block;content:"";border-color:transparent;border-style:solid}.bs-popover-top,.bs-popover-auto[x-placement^=top]{margin-bottom:.5rem}.bs-popover-top>.arrow,.bs-popover-auto[x-placement^=top]>.arrow{bottom:calc(-0.5rem - 1px)}.bs-popover-top>.arrow::before,.bs-popover-auto[x-placement^=top]>.arrow::before{bottom:0;border-width:.5rem .5rem 0;border-top-color:rgba(0,0,0,.25)}.bs-popover-top>.arrow::after,.bs-popover-auto[x-placement^=top]>.arrow::after{bottom:1px;border-width:.5rem .5rem 0;border-top-color:#fff}.bs-popover-right,.bs-popover-auto[x-placement^=right]{margin-left:.5rem}.bs-popover-right>.arrow,.bs-popover-auto[x-placement^=right]>.arrow{left:calc(-0.5rem - 1px);width:.5rem;height:1rem;margin:.3rem 0}.bs-popover-right>.arrow::before,.bs-popover-auto[x-placement^=right]>.arrow::before{left:0;border-width:.5rem .5rem .5rem 0;border-right-color:rgba(0,0,0,.25)}.bs-popover-right>.arrow::after,.bs-popover-auto[x-placement^=right]>.arrow::after{left:1px;border-width:.5rem .5rem .5rem 0;border-right-color:#fff}.bs-popover-bottom,.bs-popover-auto[x-placement^=bottom]{margin-top:.5rem}.bs-popover-bottom>.arrow,.bs-popover-auto[x-placement^=bottom]>.arrow{top:calc(-0.5rem - 1px)}.bs-popover-bottom>.arrow::before,.bs-popover-auto[x-placement^=bottom]>.arrow::before{top:0;border-width:0 .5rem .5rem .5rem;border-bottom-color:rgba(0,0,0,.25)}.bs-popover-bottom>.arrow::after,.bs-popover-auto[x-placement^=bottom]>.arrow::after{top:1px;border-width:0 .5rem .5rem .5rem;border-bottom-color:#fff}.bs-popover-bottom .popover-header::before,.bs-popover-auto[x-placement^=bottom] .popover-header::before{position:absolute;top:0;left:50%;display:block;width:1rem;margin-left:-0.5rem;content:"";border-bottom:1px solid #f7f7f7}.bs-popover-left,.bs-popover-auto[x-placement^=left]{margin-right:.5rem}.bs-popover-left>.arrow,.bs-popover-auto[x-placement^=left]>.arrow{right:calc(-0.5rem - 1px);width:.5rem;height:1rem;margin:.3rem 0}.bs-popover-left>.arrow::before,.bs-popover-auto[x-placement^=left]>.arrow::before{right:0;border-width:.5rem 0 .5rem .5rem;border-left-color:rgba(0,0,0,.25)}.bs-popover-left>.arrow::after,.bs-popover-auto[x-placement^=left]>.arrow::after{right:1px;border-width:.5rem 0 .5rem .5rem;border-left-color:#fff}.popover-header{padding:.5rem .75rem;margin-bottom:0;font-size:1rem;background-color:#f7f7f7;border-bottom:1px solid #ebebeb;border-top-left-radius:calc(0.3rem - 1px);border-top-right-radius:calc(0.3rem - 1px)}.popover-header:empty{display:none}.popover-body{padding:.5rem .75rem;color:#212529}.carousel{position:relative}.carousel.pointer-event{touch-action:pan-y}.carousel-inner{position:relative;width:100%;overflow:hidden}.carousel-inner::after{display:block;clear:both;content:""}.carousel-item{position:relative;display:none;float:left;width:100%;margin-right:-100%;backface-visibility:hidden;transition:transform .6s ease-in-out}@media(prefers-reduced-motion: reduce){.carousel-item{transition:none}}.carousel-item.active,.carousel-item-next,.carousel-item-prev{display:block}.carousel-item-next:not(.carousel-item-left),.active.carousel-item-right{transform:translateX(100%)}.carousel-item-prev:not(.carousel-item-right),.active.carousel-item-left{transform:translateX(-100%)}.carousel-fade .carousel-item{opacity:0;transition-property:opacity;transform:none}.carousel-fade .carousel-item.active,.carousel-fade .carousel-item-next.carousel-item-left,.carousel-fade .carousel-item-prev.carousel-item-right{z-index:1;opacity:1}.carousel-fade .active.carousel-item-left,.carousel-fade .active.carousel-item-right{z-index:0;opacity:0;transition:opacity 0s .6s}@media(prefers-reduced-motion: reduce){.carousel-fade .active.carousel-item-left,.carousel-fade .active.carousel-item-right{transition:none}}.carousel-control-prev,.carousel-control-next{position:absolute;top:0;bottom:0;z-index:1;display:flex;align-items:center;justify-content:center;width:15%;color:#fff;text-align:center;opacity:.5;transition:opacity .15s ease}@media(prefers-reduced-motion: reduce){.carousel-control-prev,.carousel-control-next{transition:none}}.carousel-control-prev:hover,.carousel-control-prev:focus,.carousel-control-next:hover,.carousel-control-next:focus{color:#fff;text-decoration:none;outline:0;opacity:.9}.carousel-control-prev{left:0}.carousel-control-next{right:0}.carousel-control-prev-icon,.carousel-control-next-icon{display:inline-block;width:20px;height:20px;background:no-repeat 50%/100% 100%}.carousel-control-prev-icon{background-image:url%28"data:image/svg+xml,%3csvg xmlns=\'http://www.w3.org/2000/svg\' fill=\'%23fff\' width=\'8\' height=\'8\' viewBox=\'0 0 8 8\'%3e%3cpath d=\'M5.25 0l-4 4 4 4 1.5-1.5L4.25 4l2.5-2.5L5.25 0z\'/%3e%3c/svg%3e"%29}.carousel-control-next-icon{background-image:url%28"data:image/svg+xml,%3csvg xmlns=\'http://www.w3.org/2000/svg\' fill=\'%23fff\' width=\'8\' height=\'8\' viewBox=\'0 0 8 8\'%3e%3cpath d=\'M2.75 0l-1.5 1.5L3.75 4l-2.5 2.5L2.75 8l4-4-4-4z\'/%3e%3c/svg%3e"%29}.carousel-indicators{position:absolute;right:0;bottom:0;left:0;z-index:15;display:flex;justify-content:center;padding-left:0;margin-right:15%;margin-left:15%;list-style:none}.carousel-indicators li{box-sizing:content-box;flex:0 1 auto;width:30px;height:3px;margin-right:3px;margin-left:3px;text-indent:-999px;cursor:pointer;background-color:#fff;background-clip:padding-box;border-top:10px solid transparent;border-bottom:10px solid transparent;opacity:.5;transition:opacity .6s ease}@media(prefers-reduced-motion: reduce){.carousel-indicators li{transition:none}}.carousel-indicators .active{opacity:1}.carousel-caption{position:absolute;right:15%;bottom:20px;left:15%;z-index:10;padding-top:20px;padding-bottom:20px;color:#fff;text-align:center}@keyframes spinner-border{to{transform:rotate(360deg)}}.spinner-border{display:inline-block;width:2rem;height:2rem;vertical-align:text-bottom;border:.25em solid currentColor;border-right-color:transparent;border-radius:50%;animation:spinner-border .75s linear infinite}.spinner-border-sm{width:1rem;height:1rem;border-width:.2em}@keyframes spinner-grow{0%{transform:scale(0)}50%{opacity:1}}.spinner-grow{display:inline-block;width:2rem;height:2rem;vertical-align:text-bottom;background-color:currentColor;border-radius:50%;opacity:0;animation:spinner-grow .75s linear infinite}.spinner-grow-sm{width:1rem;height:1rem}.align-baseline{vertical-align:baseline !important}.align-top{vertical-align:top !important}.align-middle{vertical-align:middle !important}.align-bottom{vertical-align:bottom !important}.align-text-bottom{vertical-align:text-bottom !important}.align-text-top{vertical-align:text-top !important}.bg-primary{background-color:#00a99d !important}a.bg-primary:hover,a.bg-primary:focus,button.bg-primary:hover,button.bg-primary:focus{background-color:#00766e !important}.bg-secondary{background-color:#6c757d !important}a.bg-secondary:hover,a.bg-secondary:focus,button.bg-secondary:hover,button.bg-secondary:focus{background-color:#545b62 !important}.bg-success{background-color:#28a745 !important}a.bg-success:hover,a.bg-success:focus,button.bg-success:hover,button.bg-success:focus{background-color:#1e7e34 !important}.bg-info{background-color:#17a2b8 !important}a.bg-info:hover,a.bg-info:focus,button.bg-info:hover,button.bg-info:focus{background-color:#117a8b !important}.bg-warning{background-color:#ffc107 !important}a.bg-warning:hover,a.bg-warning:focus,button.bg-warning:hover,button.bg-warning:focus{background-color:#d39e00 !important}.bg-danger{background-color:#dc3545 !important}a.bg-danger:hover,a.bg-danger:focus,button.bg-danger:hover,button.bg-danger:focus{background-color:#bd2130 !important}.bg-light{background-color:#f8f9fa !important}a.bg-light:hover,a.bg-light:focus,button.bg-light:hover,button.bg-light:focus{background-color:#dae0e5 !important}.bg-dark{background-color:#343a40 !important}a.bg-dark:hover,a.bg-dark:focus,button.bg-dark:hover,button.bg-dark:focus{background-color:#1d2124 !important}.bg-white{background-color:#fff !important}.bg-transparent{background-color:transparent !important}.border{border:1px solid #dee2e6 !important}.border-top{border-top:1px solid #dee2e6 !important}.border-right{border-right:1px solid #dee2e6 !important}.border-bottom{border-bottom:1px solid #dee2e6 !important}.border-left{border-left:1px solid #dee2e6 !important}.border-0{border:0 !important}.border-top-0{border-top:0 !important}.border-right-0{border-right:0 !important}.border-bottom-0{border-bottom:0 !important}.border-left-0{border-left:0 !important}.border-primary{border-color:#00a99d !important}.border-secondary{border-color:#6c757d !important}.border-success{border-color:#28a745 !important}.border-info{border-color:#17a2b8 !important}.border-warning{border-color:#ffc107 !important}.border-danger{border-color:#dc3545 !important}.border-light{border-color:#f8f9fa !important}.border-dark{border-color:#343a40 !important}.border-white{border-color:#fff !important}.rounded-sm{border-radius:.2rem !important}.rounded{border-radius:.25rem !important}.rounded-top{border-top-left-radius:.25rem !important;border-top-right-radius:.25rem !important}.rounded-right{border-top-right-radius:.25rem !important;border-bottom-right-radius:.25rem !important}.rounded-bottom{border-bottom-right-radius:.25rem !important;border-bottom-left-radius:.25rem !important}.rounded-left{border-top-left-radius:.25rem !important;border-bottom-left-radius:.25rem !important}.rounded-lg{border-radius:.3rem !important}.rounded-circle{border-radius:50% !important}.rounded-pill{border-radius:50rem !important}.rounded-0{border-radius:0 !important}.clearfix::after{display:block;clear:both;content:""}.d-none{display:none !important}.d-inline{display:inline !important}.d-inline-block{display:inline-block !important}.d-block{display:block !important}.d-table{display:table !important}.d-table-row{display:table-row !important}.d-table-cell{display:table-cell !important}.d-flex{display:flex !important}.d-inline-flex{display:inline-flex !important}@media(min-width: 576px){.d-sm-none{display:none !important}.d-sm-inline{display:inline !important}.d-sm-inline-block{display:inline-block !important}.d-sm-block{display:block !important}.d-sm-table{display:table !important}.d-sm-table-row{display:table-row !important}.d-sm-table-cell{display:table-cell !important}.d-sm-flex{display:flex !important}.d-sm-inline-flex{display:inline-flex !important}}@media(min-width: 768px){.d-md-none{display:none !important}.d-md-inline{display:inline !important}.d-md-inline-block{display:inline-block !important}.d-md-block{display:block !important}.d-md-table{display:table !important}.d-md-table-row{display:table-row !important}.d-md-table-cell{display:table-cell !important}.d-md-flex{display:flex !important}.d-md-inline-flex{display:inline-flex !important}}@media(min-width: 992px){.d-lg-none{display:none !important}.d-lg-inline{display:inline !important}.d-lg-inline-block{display:inline-block !important}.d-lg-block{display:block !important}.d-lg-table{display:table !important}.d-lg-table-row{display:table-row !important}.d-lg-table-cell{display:table-cell !important}.d-lg-flex{display:flex !important}.d-lg-inline-flex{display:inline-flex !important}}@media(min-width: 1200px){.d-xl-none{display:none !important}.d-xl-inline{display:inline !important}.d-xl-inline-block{display:inline-block !important}.d-xl-block{display:block !important}.d-xl-table{display:table !important}.d-xl-table-row{display:table-row !important}.d-xl-table-cell{display:table-cell !important}.d-xl-flex{display:flex !important}.d-xl-inline-flex{display:inline-flex !important}}@media print{.d-print-none{display:none !important}.d-print-inline{display:inline !important}.d-print-inline-block{display:inline-block !important}.d-print-block{display:block !important}.d-print-table{display:table !important}.d-print-table-row{display:table-row !important}.d-print-table-cell{display:table-cell !important}.d-print-flex{display:flex !important}.d-print-inline-flex{display:inline-flex !important}}.embed-responsive{position:relative;display:block;width:100%;padding:0;overflow:hidden}.embed-responsive::before{display:block;content:""}.embed-responsive .embed-responsive-item,.embed-responsive iframe,.embed-responsive embed,.embed-responsive object,.embed-responsive video{position:absolute;top:0;bottom:0;left:0;width:100%;height:100%;border:0}.embed-responsive-21by9::before{padding-top:42.8571428571%}.embed-responsive-16by9::before{padding-top:56.25%}.embed-responsive-4by3::before{padding-top:75%}.embed-responsive-1by1::before{padding-top:100%}.embed-responsive-21by9::before{padding-top:42.8571428571%}.embed-responsive-16by9::before{padding-top:56.25%}.embed-responsive-4by3::before{padding-top:75%}.embed-responsive-1by1::before{padding-top:100%}.flex-row{flex-direction:row !important}.flex-column{flex-direction:column !important}.flex-row-reverse{flex-direction:row-reverse !important}.flex-column-reverse{flex-direction:column-reverse !important}.flex-wrap{flex-wrap:wrap !important}.flex-nowrap{flex-wrap:nowrap !important}.flex-wrap-reverse{flex-wrap:wrap-reverse !important}.flex-fill{flex:1 1 auto !important}.flex-grow-0{flex-grow:0 !important}.flex-grow-1{flex-grow:1 !important}.flex-shrink-0{flex-shrink:0 !important}.flex-shrink-1{flex-shrink:1 !important}.justify-content-start{justify-content:flex-start !important}.justify-content-end{justify-content:flex-end !important}.justify-content-center{justify-content:center !important}.justify-content-between{justify-content:space-between !important}.justify-content-around{justify-content:space-around !important}.align-items-start{align-items:flex-start !important}.align-items-end{align-items:flex-end !important}.align-items-center{align-items:center !important}.align-items-baseline{align-items:baseline !important}.align-items-stretch{align-items:stretch !important}.align-content-start{align-content:flex-start !important}.align-content-end{align-content:flex-end !important}.align-content-center{align-content:center !important}.align-content-between{align-content:space-between !important}.align-content-around{align-content:space-around !important}.align-content-stretch{align-content:stretch !important}.align-self-auto{align-self:auto !important}.align-self-start{align-self:flex-start !important}.align-self-end{align-self:flex-end !important}.align-self-center{align-self:center !important}.align-self-baseline{align-self:baseline !important}.align-self-stretch{align-self:stretch !important}@media(min-width: 576px){.flex-sm-row{flex-direction:row !important}.flex-sm-column{flex-direction:column !important}.flex-sm-row-reverse{flex-direction:row-reverse !important}.flex-sm-column-reverse{flex-direction:column-reverse !important}.flex-sm-wrap{flex-wrap:wrap !important}.flex-sm-nowrap{flex-wrap:nowrap !important}.flex-sm-wrap-reverse{flex-wrap:wrap-reverse !important}.flex-sm-fill{flex:1 1 auto !important}.flex-sm-grow-0{flex-grow:0 !important}.flex-sm-grow-1{flex-grow:1 !important}.flex-sm-shrink-0{flex-shrink:0 !important}.flex-sm-shrink-1{flex-shrink:1 !important}.justify-content-sm-start{justify-content:flex-start !important}.justify-content-sm-end{justify-content:flex-end !important}.justify-content-sm-center{justify-content:center !important}.justify-content-sm-between{justify-content:space-between !important}.justify-content-sm-around{justify-content:space-around !important}.align-items-sm-start{align-items:flex-start !important}.align-items-sm-end{align-items:flex-end !important}.align-items-sm-center{align-items:center !important}.align-items-sm-baseline{align-items:baseline !important}.align-items-sm-stretch{align-items:stretch !important}.align-content-sm-start{align-content:flex-start !important}.align-content-sm-end{align-content:flex-end !important}.align-content-sm-center{align-content:center !important}.align-content-sm-between{align-content:space-between !important}.align-content-sm-around{align-content:space-around !important}.align-content-sm-stretch{align-content:stretch !important}.align-self-sm-auto{align-self:auto !important}.align-self-sm-start{align-self:flex-start !important}.align-self-sm-end{align-self:flex-end !important}.align-self-sm-center{align-self:center !important}.align-self-sm-baseline{align-self:baseline !important}.align-self-sm-stretch{align-self:stretch !important}}@media(min-width: 768px){.flex-md-row{flex-direction:row !important}.flex-md-column{flex-direction:column !important}.flex-md-row-reverse{flex-direction:row-reverse !important}.flex-md-column-reverse{flex-direction:column-reverse !important}.flex-md-wrap{flex-wrap:wrap !important}.flex-md-nowrap{flex-wrap:nowrap !important}.flex-md-wrap-reverse{flex-wrap:wrap-reverse !important}.flex-md-fill{flex:1 1 auto !important}.flex-md-grow-0{flex-grow:0 !important}.flex-md-grow-1{flex-grow:1 !important}.flex-md-shrink-0{flex-shrink:0 !important}.flex-md-shrink-1{flex-shrink:1 !important}.justify-content-md-start{justify-content:flex-start !important}.justify-content-md-end{justify-content:flex-end !important}.justify-content-md-center{justify-content:center !important}.justify-content-md-between{justify-content:space-between !important}.justify-content-md-around{justify-content:space-around !important}.align-items-md-start{align-items:flex-start !important}.align-items-md-end{align-items:flex-end !important}.align-items-md-center{align-items:center !important}.align-items-md-baseline{align-items:baseline !important}.align-items-md-stretch{align-items:stretch !important}.align-content-md-start{align-content:flex-start !important}.align-content-md-end{align-content:flex-end !important}.align-content-md-center{align-content:center !important}.align-content-md-between{align-content:space-between !important}.align-content-md-around{align-content:space-around !important}.align-content-md-stretch{align-content:stretch !important}.align-self-md-auto{align-self:auto !important}.align-self-md-start{align-self:flex-start !important}.align-self-md-end{align-self:flex-end !important}.align-self-md-center{align-self:center !important}.align-self-md-baseline{align-self:baseline !important}.align-self-md-stretch{align-self:stretch !important}}@media(min-width: 992px){.flex-lg-row{flex-direction:row !important}.flex-lg-column{flex-direction:column !important}.flex-lg-row-reverse{flex-direction:row-reverse !important}.flex-lg-column-reverse{flex-direction:column-reverse !important}.flex-lg-wrap{flex-wrap:wrap !important}.flex-lg-nowrap{flex-wrap:nowrap !important}.flex-lg-wrap-reverse{flex-wrap:wrap-reverse !important}.flex-lg-fill{flex:1 1 auto !important}.flex-lg-grow-0{flex-grow:0 !important}.flex-lg-grow-1{flex-grow:1 !important}.flex-lg-shrink-0{flex-shrink:0 !important}.flex-lg-shrink-1{flex-shrink:1 !important}.justify-content-lg-start{justify-content:flex-start !important}.justify-content-lg-end{justify-content:flex-end !important}.justify-content-lg-center{justify-content:center !important}.justify-content-lg-between{justify-content:space-between !important}.justify-content-lg-around{justify-content:space-around !important}.align-items-lg-start{align-items:flex-start !important}.align-items-lg-end{align-items:flex-end !important}.align-items-lg-center{align-items:center !important}.align-items-lg-baseline{align-items:baseline !important}.align-items-lg-stretch{align-items:stretch !important}.align-content-lg-start{align-content:flex-start !important}.align-content-lg-end{align-content:flex-end !important}.align-content-lg-center{align-content:center !important}.align-content-lg-between{align-content:space-between !important}.align-content-lg-around{align-content:space-around !important}.align-content-lg-stretch{align-content:stretch !important}.align-self-lg-auto{align-self:auto !important}.align-self-lg-start{align-self:flex-start !important}.align-self-lg-end{align-self:flex-end !important}.align-self-lg-center{align-self:center !important}.align-self-lg-baseline{align-self:baseline !important}.align-self-lg-stretch{align-self:stretch !important}}@media(min-width: 1200px){.flex-xl-row{flex-direction:row !important}.flex-xl-column{flex-direction:column !important}.flex-xl-row-reverse{flex-direction:row-reverse !important}.flex-xl-column-reverse{flex-direction:column-reverse !important}.flex-xl-wrap{flex-wrap:wrap !important}.flex-xl-nowrap{flex-wrap:nowrap !important}.flex-xl-wrap-reverse{flex-wrap:wrap-reverse !important}.flex-xl-fill{flex:1 1 auto !important}.flex-xl-grow-0{flex-grow:0 !important}.flex-xl-grow-1{flex-grow:1 !important}.flex-xl-shrink-0{flex-shrink:0 !important}.flex-xl-shrink-1{flex-shrink:1 !important}.justify-content-xl-start{justify-content:flex-start !important}.justify-content-xl-end{justify-content:flex-end !important}.justify-content-xl-center{justify-content:center !important}.justify-content-xl-between{justify-content:space-between !important}.justify-content-xl-around{justify-content:space-around !important}.align-items-xl-start{align-items:flex-start !important}.align-items-xl-end{align-items:flex-end !important}.align-items-xl-center{align-items:center !important}.align-items-xl-baseline{align-items:baseline !important}.align-items-xl-stretch{align-items:stretch !important}.align-content-xl-start{align-content:flex-start !important}.align-content-xl-end{align-content:flex-end !important}.align-content-xl-center{align-content:center !important}.align-content-xl-between{align-content:space-between !important}.align-content-xl-around{align-content:space-around !important}.align-content-xl-stretch{align-content:stretch !important}.align-self-xl-auto{align-self:auto !important}.align-self-xl-start{align-self:flex-start !important}.align-self-xl-end{align-self:flex-end !important}.align-self-xl-center{align-self:center !important}.align-self-xl-baseline{align-self:baseline !important}.align-self-xl-stretch{align-self:stretch !important}}.float-left{float:left !important}.float-right{float:right !important}.float-none{float:none !important}@media(min-width: 576px){.float-sm-left{float:left !important}.float-sm-right{float:right !important}.float-sm-none{float:none !important}}@media(min-width: 768px){.float-md-left{float:left !important}.float-md-right{float:right !important}.float-md-none{float:none !important}}@media(min-width: 992px){.float-lg-left{float:left !important}.float-lg-right{float:right !important}.float-lg-none{float:none !important}}@media(min-width: 1200px){.float-xl-left{float:left !important}.float-xl-right{float:right !important}.float-xl-none{float:none !important}}.overflow-auto{overflow:auto !important}.overflow-hidden{overflow:hidden !important}.position-static{position:static !important}.position-relative{position:relative !important}.position-absolute{position:absolute !important}.position-fixed{position:fixed !important}.position-sticky{position:sticky !important}.fixed-top{position:fixed;top:0;right:0;left:0;z-index:1030}.fixed-bottom{position:fixed;right:0;bottom:0;left:0;z-index:1030}@supports(position: sticky){.sticky-top{position:sticky;top:0;z-index:1020}}.sr-only{position:absolute;width:1px;height:1px;padding:0;margin:-1px;overflow:hidden;clip:rect(0, 0, 0, 0);white-space:nowrap;border:0}.sr-only-focusable:active,.sr-only-focusable:focus{position:static;width:auto;height:auto;overflow:visible;clip:auto;white-space:normal}.shadow-sm{box-shadow:0 .125rem .25rem rgba(0,0,0,.075) !important}.shadow{box-shadow:0 .5rem 1rem rgba(0,0,0,.15) !important}.shadow-lg{box-shadow:0 1rem 3rem rgba(0,0,0,.175) !important}.shadow-none{box-shadow:none !important}.w-25{width:25% !important}.w-50{width:50% !important}.w-75{width:75% !important}.w-100{width:100% !important}.w-auto{width:auto !important}.h-25{height:25% !important}.h-50{height:50% !important}.h-75{height:75% !important}.h-100{height:100% !important}.h-auto{height:auto !important}.mw-100{max-width:100% !important}.mh-100{max-height:100% !important}.min-vw-100{min-width:100vw !important}.min-vh-100{min-height:100vh !important}.vw-100{width:100vw !important}.vh-100{height:100vh !important}.stretched-link::after{position:absolute;top:0;right:0;bottom:0;left:0;z-index:1;pointer-events:auto;content:"";background-color:rgba(0,0,0,0)}.m-0{margin:0 !important}.mt-0,.my-0{margin-top:0 !important}.mr-0,.mx-0{margin-right:0 !important}.mb-0,.my-0{margin-bottom:0 !important}.ml-0,.mx-0{margin-left:0 !important}.m-1{margin:.25rem !important}.mt-1,.my-1{margin-top:.25rem !important}.mr-1,.mx-1{margin-right:.25rem !important}.mb-1,.my-1{margin-bottom:.25rem !important}.ml-1,.mx-1{margin-left:.25rem !important}.m-2{margin:.5rem !important}.mt-2,.my-2{margin-top:.5rem !important}.mr-2,.mx-2{margin-right:.5rem !important}.mb-2,.my-2{margin-bottom:.5rem !important}.ml-2,.mx-2{margin-left:.5rem !important}.m-3{margin:1rem !important}.mt-3,.my-3{margin-top:1rem !important}.mr-3,.mx-3{margin-right:1rem !important}.mb-3,.my-3{margin-bottom:1rem !important}.ml-3,.mx-3{margin-left:1rem !important}.m-4{margin:1.5rem !important}.mt-4,.my-4{margin-top:1.5rem !important}.mr-4,.mx-4{margin-right:1.5rem !important}.mb-4,.my-4{margin-bottom:1.5rem !important}.ml-4,.mx-4{margin-left:1.5rem !important}.m-5{margin:3rem !important}.mt-5,.my-5{margin-top:3rem !important}.mr-5,.mx-5{margin-right:3rem !important}.mb-5,.my-5{margin-bottom:3rem !important}.ml-5,.mx-5{margin-left:3rem !important}.p-0{padding:0 !important}.pt-0,.py-0{padding-top:0 !important}.pr-0,.px-0{padding-right:0 !important}.pb-0,.py-0{padding-bottom:0 !important}.pl-0,.px-0{padding-left:0 !important}.p-1{padding:.25rem !important}.pt-1,.py-1{padding-top:.25rem !important}.pr-1,.px-1{padding-right:.25rem !important}.pb-1,.py-1{padding-bottom:.25rem !important}.pl-1,.px-1{padding-left:.25rem !important}.p-2{padding:.5rem !important}.pt-2,.py-2{padding-top:.5rem !important}.pr-2,.px-2{padding-right:.5rem !important}.pb-2,.py-2{padding-bottom:.5rem !important}.pl-2,.px-2{padding-left:.5rem !important}.p-3{padding:1rem !important}.pt-3,.py-3{padding-top:1rem !important}.pr-3,.px-3{padding-right:1rem !important}.pb-3,.py-3{padding-bottom:1rem !important}.pl-3,.px-3{padding-left:1rem !important}.p-4{padding:1.5rem !important}.pt-4,.py-4{padding-top:1.5rem !important}.pr-4,.px-4{padding-right:1.5rem !important}.pb-4,.py-4{padding-bottom:1.5rem !important}.pl-4,.px-4{padding-left:1.5rem !important}.p-5{padding:3rem !important}.pt-5,.py-5{padding-top:3rem !important}.pr-5,.px-5{padding-right:3rem !important}.pb-5,.py-5{padding-bottom:3rem !important}.pl-5,.px-5{padding-left:3rem !important}.m-n1{margin:-0.25rem !important}.mt-n1,.my-n1{margin-top:-0.25rem !important}.mr-n1,.mx-n1{margin-right:-0.25rem !important}.mb-n1,.my-n1{margin-bottom:-0.25rem !important}.ml-n1,.mx-n1{margin-left:-0.25rem !important}.m-n2{margin:-0.5rem !important}.mt-n2,.my-n2{margin-top:-0.5rem !important}.mr-n2,.mx-n2{margin-right:-0.5rem !important}.mb-n2,.my-n2{margin-bottom:-0.5rem !important}.ml-n2,.mx-n2{margin-left:-0.5rem !important}.m-n3{margin:-1rem !important}.mt-n3,.my-n3{margin-top:-1rem !important}.mr-n3,.mx-n3{margin-right:-1rem !important}.mb-n3,.my-n3{margin-bottom:-1rem !important}.ml-n3,.mx-n3{margin-left:-1rem !important}.m-n4{margin:-1.5rem !important}.mt-n4,.my-n4{margin-top:-1.5rem !important}.mr-n4,.mx-n4{margin-right:-1.5rem !important}.mb-n4,.my-n4{margin-bottom:-1.5rem !important}.ml-n4,.mx-n4{margin-left:-1.5rem !important}.m-n5{margin:-3rem !important}.mt-n5,.my-n5{margin-top:-3rem !important}.mr-n5,.mx-n5{margin-right:-3rem !important}.mb-n5,.my-n5{margin-bottom:-3rem !important}.ml-n5,.mx-n5{margin-left:-3rem !important}.m-auto{margin:auto !important}.mt-auto,.my-auto{margin-top:auto !important}.mr-auto,.mx-auto{margin-right:auto !important}.mb-auto,.my-auto{margin-bottom:auto !important}.ml-auto,.mx-auto{margin-left:auto !important}@media(min-width: 576px){.m-sm-0{margin:0 !important}.mt-sm-0,.my-sm-0{margin-top:0 !important}.mr-sm-0,.mx-sm-0{margin-right:0 !important}.mb-sm-0,.my-sm-0{margin-bottom:0 !important}.ml-sm-0,.mx-sm-0{margin-left:0 !important}.m-sm-1{margin:.25rem !important}.mt-sm-1,.my-sm-1{margin-top:.25rem !important}.mr-sm-1,.mx-sm-1{margin-right:.25rem !important}.mb-sm-1,.my-sm-1{margin-bottom:.25rem !important}.ml-sm-1,.mx-sm-1{margin-left:.25rem !important}.m-sm-2{margin:.5rem !important}.mt-sm-2,.my-sm-2{margin-top:.5rem !important}.mr-sm-2,.mx-sm-2{margin-right:.5rem !important}.mb-sm-2,.my-sm-2{margin-bottom:.5rem !important}.ml-sm-2,.mx-sm-2{margin-left:.5rem !important}.m-sm-3{margin:1rem !important}.mt-sm-3,.my-sm-3{margin-top:1rem !important}.mr-sm-3,.mx-sm-3{margin-right:1rem !important}.mb-sm-3,.my-sm-3{margin-bottom:1rem !important}.ml-sm-3,.mx-sm-3{margin-left:1rem !important}.m-sm-4{margin:1.5rem !important}.mt-sm-4,.my-sm-4{margin-top:1.5rem !important}.mr-sm-4,.mx-sm-4{margin-right:1.5rem !important}.mb-sm-4,.my-sm-4{margin-bottom:1.5rem !important}.ml-sm-4,.mx-sm-4{margin-left:1.5rem !important}.m-sm-5{margin:3rem !important}.mt-sm-5,.my-sm-5{margin-top:3rem !important}.mr-sm-5,.mx-sm-5{margin-right:3rem !important}.mb-sm-5,.my-sm-5{margin-bottom:3rem !important}.ml-sm-5,.mx-sm-5{margin-left:3rem !important}.p-sm-0{padding:0 !important}.pt-sm-0,.py-sm-0{padding-top:0 !important}.pr-sm-0,.px-sm-0{padding-right:0 !important}.pb-sm-0,.py-sm-0{padding-bottom:0 !important}.pl-sm-0,.px-sm-0{padding-left:0 !important}.p-sm-1{padding:.25rem !important}.pt-sm-1,.py-sm-1{padding-top:.25rem !important}.pr-sm-1,.px-sm-1{padding-right:.25rem !important}.pb-sm-1,.py-sm-1{padding-bottom:.25rem !important}.pl-sm-1,.px-sm-1{padding-left:.25rem !important}.p-sm-2{padding:.5rem !important}.pt-sm-2,.py-sm-2{padding-top:.5rem !important}.pr-sm-2,.px-sm-2{padding-right:.5rem !important}.pb-sm-2,.py-sm-2{padding-bottom:.5rem !important}.pl-sm-2,.px-sm-2{padding-left:.5rem !important}.p-sm-3{padding:1rem !important}.pt-sm-3,.py-sm-3{padding-top:1rem !important}.pr-sm-3,.px-sm-3{padding-right:1rem !important}.pb-sm-3,.py-sm-3{padding-bottom:1rem !important}.pl-sm-3,.px-sm-3{padding-left:1rem !important}.p-sm-4{padding:1.5rem !important}.pt-sm-4,.py-sm-4{padding-top:1.5rem !important}.pr-sm-4,.px-sm-4{padding-right:1.5rem !important}.pb-sm-4,.py-sm-4{padding-bottom:1.5rem !important}.pl-sm-4,.px-sm-4{padding-left:1.5rem !important}.p-sm-5{padding:3rem !important}.pt-sm-5,.py-sm-5{padding-top:3rem !important}.pr-sm-5,.px-sm-5{padding-right:3rem !important}.pb-sm-5,.py-sm-5{padding-bottom:3rem !important}.pl-sm-5,.px-sm-5{padding-left:3rem !important}.m-sm-n1{margin:-0.25rem !important}.mt-sm-n1,.my-sm-n1{margin-top:-0.25rem !important}.mr-sm-n1,.mx-sm-n1{margin-right:-0.25rem !important}.mb-sm-n1,.my-sm-n1{margin-bottom:-0.25rem !important}.ml-sm-n1,.mx-sm-n1{margin-left:-0.25rem !important}.m-sm-n2{margin:-0.5rem !important}.mt-sm-n2,.my-sm-n2{margin-top:-0.5rem !important}.mr-sm-n2,.mx-sm-n2{margin-right:-0.5rem !important}.mb-sm-n2,.my-sm-n2{margin-bottom:-0.5rem !important}.ml-sm-n2,.mx-sm-n2{margin-left:-0.5rem !important}.m-sm-n3{margin:-1rem !important}.mt-sm-n3,.my-sm-n3{margin-top:-1rem !important}.mr-sm-n3,.mx-sm-n3{margin-right:-1rem !important}.mb-sm-n3,.my-sm-n3{margin-bottom:-1rem !important}.ml-sm-n3,.mx-sm-n3{margin-left:-1rem !important}.m-sm-n4{margin:-1.5rem !important}.mt-sm-n4,.my-sm-n4{margin-top:-1.5rem !important}.mr-sm-n4,.mx-sm-n4{margin-right:-1.5rem !important}.mb-sm-n4,.my-sm-n4{margin-bottom:-1.5rem !important}.ml-sm-n4,.mx-sm-n4{margin-left:-1.5rem !important}.m-sm-n5{margin:-3rem !important}.mt-sm-n5,.my-sm-n5{margin-top:-3rem !important}.mr-sm-n5,.mx-sm-n5{margin-right:-3rem !important}.mb-sm-n5,.my-sm-n5{margin-bottom:-3rem !important}.ml-sm-n5,.mx-sm-n5{margin-left:-3rem !important}.m-sm-auto{margin:auto !important}.mt-sm-auto,.my-sm-auto{margin-top:auto !important}.mr-sm-auto,.mx-sm-auto{margin-right:auto !important}.mb-sm-auto,.my-sm-auto{margin-bottom:auto !important}.ml-sm-auto,.mx-sm-auto{margin-left:auto !important}}@media(min-width: 768px){.m-md-0{margin:0 !important}.mt-md-0,.my-md-0{margin-top:0 !important}.mr-md-0,.mx-md-0{margin-right:0 !important}.mb-md-0,.my-md-0{margin-bottom:0 !important}.ml-md-0,.mx-md-0{margin-left:0 !important}.m-md-1{margin:.25rem !important}.mt-md-1,.my-md-1{margin-top:.25rem !important}.mr-md-1,.mx-md-1{margin-right:.25rem !important}.mb-md-1,.my-md-1{margin-bottom:.25rem !important}.ml-md-1,.mx-md-1{margin-left:.25rem !important}.m-md-2{margin:.5rem !important}.mt-md-2,.my-md-2{margin-top:.5rem !important}.mr-md-2,.mx-md-2{margin-right:.5rem !important}.mb-md-2,.my-md-2{margin-bottom:.5rem !important}.ml-md-2,.mx-md-2{margin-left:.5rem !important}.m-md-3{margin:1rem !important}.mt-md-3,.my-md-3{margin-top:1rem !important}.mr-md-3,.mx-md-3{margin-right:1rem !important}.mb-md-3,.my-md-3{margin-bottom:1rem !important}.ml-md-3,.mx-md-3{margin-left:1rem !important}.m-md-4{margin:1.5rem !important}.mt-md-4,.my-md-4{margin-top:1.5rem !important}.mr-md-4,.mx-md-4{margin-right:1.5rem !important}.mb-md-4,.my-md-4{margin-bottom:1.5rem !important}.ml-md-4,.mx-md-4{margin-left:1.5rem !important}.m-md-5{margin:3rem !important}.mt-md-5,.my-md-5{margin-top:3rem !important}.mr-md-5,.mx-md-5{margin-right:3rem !important}.mb-md-5,.my-md-5{margin-bottom:3rem !important}.ml-md-5,.mx-md-5{margin-left:3rem !important}.p-md-0{padding:0 !important}.pt-md-0,.py-md-0{padding-top:0 !important}.pr-md-0,.px-md-0{padding-right:0 !important}.pb-md-0,.py-md-0{padding-bottom:0 !important}.pl-md-0,.px-md-0{padding-left:0 !important}.p-md-1{padding:.25rem !important}.pt-md-1,.py-md-1{padding-top:.25rem !important}.pr-md-1,.px-md-1{padding-right:.25rem !important}.pb-md-1,.py-md-1{padding-bottom:.25rem !important}.pl-md-1,.px-md-1{padding-left:.25rem !important}.p-md-2{padding:.5rem !important}.pt-md-2,.py-md-2{padding-top:.5rem !important}.pr-md-2,.px-md-2{padding-right:.5rem !important}.pb-md-2,.py-md-2{padding-bottom:.5rem !important}.pl-md-2,.px-md-2{padding-left:.5rem !important}.p-md-3{padding:1rem !important}.pt-md-3,.py-md-3{padding-top:1rem !important}.pr-md-3,.px-md-3{padding-right:1rem !important}.pb-md-3,.py-md-3{padding-bottom:1rem !important}.pl-md-3,.px-md-3{padding-left:1rem !important}.p-md-4{padding:1.5rem !important}.pt-md-4,.py-md-4{padding-top:1.5rem !important}.pr-md-4,.px-md-4{padding-right:1.5rem !important}.pb-md-4,.py-md-4{padding-bottom:1.5rem !important}.pl-md-4,.px-md-4{padding-left:1.5rem !important}.p-md-5{padding:3rem !important}.pt-md-5,.py-md-5{padding-top:3rem !important}.pr-md-5,.px-md-5{padding-right:3rem !important}.pb-md-5,.py-md-5{padding-bottom:3rem !important}.pl-md-5,.px-md-5{padding-left:3rem !important}.m-md-n1{margin:-0.25rem !important}.mt-md-n1,.my-md-n1{margin-top:-0.25rem !important}.mr-md-n1,.mx-md-n1{margin-right:-0.25rem !important}.mb-md-n1,.my-md-n1{margin-bottom:-0.25rem !important}.ml-md-n1,.mx-md-n1{margin-left:-0.25rem !important}.m-md-n2{margin:-0.5rem !important}.mt-md-n2,.my-md-n2{margin-top:-0.5rem !important}.mr-md-n2,.mx-md-n2{margin-right:-0.5rem !important}.mb-md-n2,.my-md-n2{margin-bottom:-0.5rem !important}.ml-md-n2,.mx-md-n2{margin-left:-0.5rem !important}.m-md-n3{margin:-1rem !important}.mt-md-n3,.my-md-n3{margin-top:-1rem !important}.mr-md-n3,.mx-md-n3{margin-right:-1rem !important}.mb-md-n3,.my-md-n3{margin-bottom:-1rem !important}.ml-md-n3,.mx-md-n3{margin-left:-1rem !important}.m-md-n4{margin:-1.5rem !important}.mt-md-n4,.my-md-n4{margin-top:-1.5rem !important}.mr-md-n4,.mx-md-n4{margin-right:-1.5rem !important}.mb-md-n4,.my-md-n4{margin-bottom:-1.5rem !important}.ml-md-n4,.mx-md-n4{margin-left:-1.5rem !important}.m-md-n5{margin:-3rem !important}.mt-md-n5,.my-md-n5{margin-top:-3rem !important}.mr-md-n5,.mx-md-n5{margin-right:-3rem !important}.mb-md-n5,.my-md-n5{margin-bottom:-3rem !important}.ml-md-n5,.mx-md-n5{margin-left:-3rem !important}.m-md-auto{margin:auto !important}.mt-md-auto,.my-md-auto{margin-top:auto !important}.mr-md-auto,.mx-md-auto{margin-right:auto !important}.mb-md-auto,.my-md-auto{margin-bottom:auto !important}.ml-md-auto,.mx-md-auto{margin-left:auto !important}}@media(min-width: 992px){.m-lg-0{margin:0 !important}.mt-lg-0,.my-lg-0{margin-top:0 !important}.mr-lg-0,.mx-lg-0{margin-right:0 !important}.mb-lg-0,.my-lg-0{margin-bottom:0 !important}.ml-lg-0,.mx-lg-0{margin-left:0 !important}.m-lg-1{margin:.25rem !important}.mt-lg-1,.my-lg-1{margin-top:.25rem !important}.mr-lg-1,.mx-lg-1{margin-right:.25rem !important}.mb-lg-1,.my-lg-1{margin-bottom:.25rem !important}.ml-lg-1,.mx-lg-1{margin-left:.25rem !important}.m-lg-2{margin:.5rem !important}.mt-lg-2,.my-lg-2{margin-top:.5rem !important}.mr-lg-2,.mx-lg-2{margin-right:.5rem !important}.mb-lg-2,.my-lg-2{margin-bottom:.5rem !important}.ml-lg-2,.mx-lg-2{margin-left:.5rem !important}.m-lg-3{margin:1rem !important}.mt-lg-3,.my-lg-3{margin-top:1rem !important}.mr-lg-3,.mx-lg-3{margin-right:1rem !important}.mb-lg-3,.my-lg-3{margin-bottom:1rem !important}.ml-lg-3,.mx-lg-3{margin-left:1rem !important}.m-lg-4{margin:1.5rem !important}.mt-lg-4,.my-lg-4{margin-top:1.5rem !important}.mr-lg-4,.mx-lg-4{margin-right:1.5rem !important}.mb-lg-4,.my-lg-4{margin-bottom:1.5rem !important}.ml-lg-4,.mx-lg-4{margin-left:1.5rem !important}.m-lg-5{margin:3rem !important}.mt-lg-5,.my-lg-5{margin-top:3rem !important}.mr-lg-5,.mx-lg-5{margin-right:3rem !important}.mb-lg-5,.my-lg-5{margin-bottom:3rem !important}.ml-lg-5,.mx-lg-5{margin-left:3rem !important}.p-lg-0{padding:0 !important}.pt-lg-0,.py-lg-0{padding-top:0 !important}.pr-lg-0,.px-lg-0{padding-right:0 !important}.pb-lg-0,.py-lg-0{padding-bottom:0 !important}.pl-lg-0,.px-lg-0{padding-left:0 !important}.p-lg-1{padding:.25rem !important}.pt-lg-1,.py-lg-1{padding-top:.25rem !important}.pr-lg-1,.px-lg-1{padding-right:.25rem !important}.pb-lg-1,.py-lg-1{padding-bottom:.25rem !important}.pl-lg-1,.px-lg-1{padding-left:.25rem !important}.p-lg-2{padding:.5rem !important}.pt-lg-2,.py-lg-2{padding-top:.5rem !important}.pr-lg-2,.px-lg-2{padding-right:.5rem !important}.pb-lg-2,.py-lg-2{padding-bottom:.5rem !important}.pl-lg-2,.px-lg-2{padding-left:.5rem !important}.p-lg-3{padding:1rem !important}.pt-lg-3,.py-lg-3{padding-top:1rem !important}.pr-lg-3,.px-lg-3{padding-right:1rem !important}.pb-lg-3,.py-lg-3{padding-bottom:1rem !important}.pl-lg-3,.px-lg-3{padding-left:1rem !important}.p-lg-4{padding:1.5rem !important}.pt-lg-4,.py-lg-4{padding-top:1.5rem !important}.pr-lg-4,.px-lg-4{padding-right:1.5rem !important}.pb-lg-4,.py-lg-4{padding-bottom:1.5rem !important}.pl-lg-4,.px-lg-4{padding-left:1.5rem !important}.p-lg-5{padding:3rem !important}.pt-lg-5,.py-lg-5{padding-top:3rem !important}.pr-lg-5,.px-lg-5{padding-right:3rem !important}.pb-lg-5,.py-lg-5{padding-bottom:3rem !important}.pl-lg-5,.px-lg-5{padding-left:3rem !important}.m-lg-n1{margin:-0.25rem !important}.mt-lg-n1,.my-lg-n1{margin-top:-0.25rem !important}.mr-lg-n1,.mx-lg-n1{margin-right:-0.25rem !important}.mb-lg-n1,.my-lg-n1{margin-bottom:-0.25rem !important}.ml-lg-n1,.mx-lg-n1{margin-left:-0.25rem !important}.m-lg-n2{margin:-0.5rem !important}.mt-lg-n2,.my-lg-n2{margin-top:-0.5rem !important}.mr-lg-n2,.mx-lg-n2{margin-right:-0.5rem !important}.mb-lg-n2,.my-lg-n2{margin-bottom:-0.5rem !important}.ml-lg-n2,.mx-lg-n2{margin-left:-0.5rem !important}.m-lg-n3{margin:-1rem !important}.mt-lg-n3,.my-lg-n3{margin-top:-1rem !important}.mr-lg-n3,.mx-lg-n3{margin-right:-1rem !important}.mb-lg-n3,.my-lg-n3{margin-bottom:-1rem !important}.ml-lg-n3,.mx-lg-n3{margin-left:-1rem !important}.m-lg-n4{margin:-1.5rem !important}.mt-lg-n4,.my-lg-n4{margin-top:-1.5rem !important}.mr-lg-n4,.mx-lg-n4{margin-right:-1.5rem !important}.mb-lg-n4,.my-lg-n4{margin-bottom:-1.5rem !important}.ml-lg-n4,.mx-lg-n4{margin-left:-1.5rem !important}.m-lg-n5{margin:-3rem !important}.mt-lg-n5,.my-lg-n5{margin-top:-3rem !important}.mr-lg-n5,.mx-lg-n5{margin-right:-3rem !important}.mb-lg-n5,.my-lg-n5{margin-bottom:-3rem !important}.ml-lg-n5,.mx-lg-n5{margin-left:-3rem !important}.m-lg-auto{margin:auto !important}.mt-lg-auto,.my-lg-auto{margin-top:auto !important}.mr-lg-auto,.mx-lg-auto{margin-right:auto !important}.mb-lg-auto,.my-lg-auto{margin-bottom:auto !important}.ml-lg-auto,.mx-lg-auto{margin-left:auto !important}}@media(min-width: 1200px){.m-xl-0{margin:0 !important}.mt-xl-0,.my-xl-0{margin-top:0 !important}.mr-xl-0,.mx-xl-0{margin-right:0 !important}.mb-xl-0,.my-xl-0{margin-bottom:0 !important}.ml-xl-0,.mx-xl-0{margin-left:0 !important}.m-xl-1{margin:.25rem !important}.mt-xl-1,.my-xl-1{margin-top:.25rem !important}.mr-xl-1,.mx-xl-1{margin-right:.25rem !important}.mb-xl-1,.my-xl-1{margin-bottom:.25rem !important}.ml-xl-1,.mx-xl-1{margin-left:.25rem !important}.m-xl-2{margin:.5rem !important}.mt-xl-2,.my-xl-2{margin-top:.5rem !important}.mr-xl-2,.mx-xl-2{margin-right:.5rem !important}.mb-xl-2,.my-xl-2{margin-bottom:.5rem !important}.ml-xl-2,.mx-xl-2{margin-left:.5rem !important}.m-xl-3{margin:1rem !important}.mt-xl-3,.my-xl-3{margin-top:1rem !important}.mr-xl-3,.mx-xl-3{margin-right:1rem !important}.mb-xl-3,.my-xl-3{margin-bottom:1rem !important}.ml-xl-3,.mx-xl-3{margin-left:1rem !important}.m-xl-4{margin:1.5rem !important}.mt-xl-4,.my-xl-4{margin-top:1.5rem !important}.mr-xl-4,.mx-xl-4{margin-right:1.5rem !important}.mb-xl-4,.my-xl-4{margin-bottom:1.5rem !important}.ml-xl-4,.mx-xl-4{margin-left:1.5rem !important}.m-xl-5{margin:3rem !important}.mt-xl-5,.my-xl-5{margin-top:3rem !important}.mr-xl-5,.mx-xl-5{margin-right:3rem !important}.mb-xl-5,.my-xl-5{margin-bottom:3rem !important}.ml-xl-5,.mx-xl-5{margin-left:3rem !important}.p-xl-0{padding:0 !important}.pt-xl-0,.py-xl-0{padding-top:0 !important}.pr-xl-0,.px-xl-0{padding-right:0 !important}.pb-xl-0,.py-xl-0{padding-bottom:0 !important}.pl-xl-0,.px-xl-0{padding-left:0 !important}.p-xl-1{padding:.25rem !important}.pt-xl-1,.py-xl-1{padding-top:.25rem !important}.pr-xl-1,.px-xl-1{padding-right:.25rem !important}.pb-xl-1,.py-xl-1{padding-bottom:.25rem !important}.pl-xl-1,.px-xl-1{padding-left:.25rem !important}.p-xl-2{padding:.5rem !important}.pt-xl-2,.py-xl-2{padding-top:.5rem !important}.pr-xl-2,.px-xl-2{padding-right:.5rem !important}.pb-xl-2,.py-xl-2{padding-bottom:.5rem !important}.pl-xl-2,.px-xl-2{padding-left:.5rem !important}.p-xl-3{padding:1rem !important}.pt-xl-3,.py-xl-3{padding-top:1rem !important}.pr-xl-3,.px-xl-3{padding-right:1rem !important}.pb-xl-3,.py-xl-3{padding-bottom:1rem !important}.pl-xl-3,.px-xl-3{padding-left:1rem !important}.p-xl-4{padding:1.5rem !important}.pt-xl-4,.py-xl-4{padding-top:1.5rem !important}.pr-xl-4,.px-xl-4{padding-right:1.5rem !important}.pb-xl-4,.py-xl-4{padding-bottom:1.5rem !important}.pl-xl-4,.px-xl-4{padding-left:1.5rem !important}.p-xl-5{padding:3rem !important}.pt-xl-5,.py-xl-5{padding-top:3rem !important}.pr-xl-5,.px-xl-5{padding-right:3rem !important}.pb-xl-5,.py-xl-5{padding-bottom:3rem !important}.pl-xl-5,.px-xl-5{padding-left:3rem !important}.m-xl-n1{margin:-0.25rem !important}.mt-xl-n1,.my-xl-n1{margin-top:-0.25rem !important}.mr-xl-n1,.mx-xl-n1{margin-right:-0.25rem !important}.mb-xl-n1,.my-xl-n1{margin-bottom:-0.25rem !important}.ml-xl-n1,.mx-xl-n1{margin-left:-0.25rem !important}.m-xl-n2{margin:-0.5rem !important}.mt-xl-n2,.my-xl-n2{margin-top:-0.5rem !important}.mr-xl-n2,.mx-xl-n2{margin-right:-0.5rem !important}.mb-xl-n2,.my-xl-n2{margin-bottom:-0.5rem !important}.ml-xl-n2,.mx-xl-n2{margin-left:-0.5rem !important}.m-xl-n3{margin:-1rem !important}.mt-xl-n3,.my-xl-n3{margin-top:-1rem !important}.mr-xl-n3,.mx-xl-n3{margin-right:-1rem !important}.mb-xl-n3,.my-xl-n3{margin-bottom:-1rem !important}.ml-xl-n3,.mx-xl-n3{margin-left:-1rem !important}.m-xl-n4{margin:-1.5rem !important}.mt-xl-n4,.my-xl-n4{margin-top:-1.5rem !important}.mr-xl-n4,.mx-xl-n4{margin-right:-1.5rem !important}.mb-xl-n4,.my-xl-n4{margin-bottom:-1.5rem !important}.ml-xl-n4,.mx-xl-n4{margin-left:-1.5rem !important}.m-xl-n5{margin:-3rem !important}.mt-xl-n5,.my-xl-n5{margin-top:-3rem !important}.mr-xl-n5,.mx-xl-n5{margin-right:-3rem !important}.mb-xl-n5,.my-xl-n5{margin-bottom:-3rem !important}.ml-xl-n5,.mx-xl-n5{margin-left:-3rem !important}.m-xl-auto{margin:auto !important}.mt-xl-auto,.my-xl-auto{margin-top:auto !important}.mr-xl-auto,.mx-xl-auto{margin-right:auto !important}.mb-xl-auto,.my-xl-auto{margin-bottom:auto !important}.ml-xl-auto,.mx-xl-auto{margin-left:auto !important}}.text-monospace{font-family:SFMono-Regular,Menlo,Monaco,Consolas,"Liberation Mono","Courier New",monospace !important}.text-justify{text-align:justify !important}.text-wrap{white-space:normal !important}.text-nowrap{white-space:nowrap !important}.text-truncate{overflow:hidden;text-overflow:ellipsis;white-space:nowrap}.text-left{text-align:left !important}.text-right{text-align:right !important}.text-center{text-align:center !important}@media(min-width: 576px){.text-sm-left{text-align:left !important}.text-sm-right{text-align:right !important}.text-sm-center{text-align:center !important}}@media(min-width: 768px){.text-md-left{text-align:left !important}.text-md-right{text-align:right !important}.text-md-center{text-align:center !important}}@media(min-width: 992px){.text-lg-left{text-align:left !important}.text-lg-right{text-align:right !important}.text-lg-center{text-align:center !important}}@media(min-width: 1200px){.text-xl-left{text-align:left !important}.text-xl-right{text-align:right !important}.text-xl-center{text-align:center !important}}.text-lowercase{text-transform:lowercase !important}.text-uppercase{text-transform:uppercase !important}.text-capitalize{text-transform:capitalize !important}.font-weight-light{font-weight:300 !important}.font-weight-lighter{font-weight:lighter !important}.font-weight-normal{font-weight:400 !important}.font-weight-bold{font-weight:700 !important}.font-weight-bolder{font-weight:bolder !important}.font-italic{font-style:italic !important}.text-white{color:#fff !important}.text-primary{color:#00a99d !important}a.text-primary:hover,a.text-primary:focus{color:#005d56 !important}.text-secondary{color:#6c757d !important}a.text-secondary:hover,a.text-secondary:focus{color:#494f54 !important}.text-success{color:#28a745 !important}a.text-success:hover,a.text-success:focus{color:#19692c !important}.text-info{color:#17a2b8 !important}a.text-info:hover,a.text-info:focus{color:#0f6674 !important}.text-warning{color:#ffc107 !important}a.text-warning:hover,a.text-warning:focus{color:#ba8b00 !important}.text-danger{color:#dc3545 !important}a.text-danger:hover,a.text-danger:focus{color:#a71d2a !important}.text-light{color:#f8f9fa !important}a.text-light:hover,a.text-light:focus{color:#cbd3da !important}.text-dark{color:#343a40 !important}a.text-dark:hover,a.text-dark:focus{color:#121416 !important}.text-body{color:#212529 !important}.text-muted{color:#6c757d !important}.text-black-50{color:rgba(0,0,0,.5) !important}.text-white-50{color:rgba(255,255,255,.5) !important}.text-hide{font:0/0 a;color:transparent;text-shadow:none;background-color:transparent;border:0}.text-decoration-none{text-decoration:none !important}.text-break{word-break:break-word !important;overflow-wrap:break-word !important}.text-reset{color:inherit !important}.visible{visibility:visible !important}.invisible{visibility:hidden !important}@media print{*,*::before,*::after{text-shadow:none !important;box-shadow:none !important}a:not(.btn){text-decoration:underline}abbr[title]::after{content:" (" attr(title) ")"}pre{white-space:pre-wrap !important}pre,blockquote{border:1px solid #adb5bd;page-break-inside:avoid}thead{display:table-header-group}tr,img{page-break-inside:avoid}p,h2,h3{orphans:3;widows:3}h2,h3{page-break-after:avoid}@page{size:a3}body{min-width:992px !important}.container{min-width:992px !important}.navbar{display:none}.badge{border:1px solid #000}.table{border-collapse:collapse !important}.table td,.table th{background-color:#fff !important}.table-bordered th,.table-bordered td{border:1px solid #dee2e6 !important}.table-dark{color:inherit}.table-dark th,.table-dark td,.table-dark thead th,.table-dark tbody+tbody{border-color:#dee2e6}.table .thead-dark th{color:inherit;border-color:#dee2e6}}div.alignmentjs-container{float:left;padding:0;margin:0}svg#alignmentjs-labels>text{font-family:Courier;font-size:14px}.alert-info{position:relative;font-family:"Montserrat",sans-serif;border:none;color:#156057;word-spacing:1.5px}.alert-dismissable,.alert-dismissible{padding:20px 100px 20px 27px}.alert-dismissable .close{position:absolute;top:0px;right:0px;color:#fff;font-weight:400;font-size:28px;padding:0px 18px 3px 18px;background-color:#00a99d;opacity:1}.alert-dismissable .close:hover{opacity:.85}.alert>p+p{margin:12px 0px 5px 0px}.btn{border-radius:0;padding:.375rem .75rem;font-family:"Montserrat",sans-serif;height:38px;letter-spacing:.25px}.btn .caret{margin:0 5px}.btn-group-justified{border:1px solid #ccc;box-sizing:border-box}.btn-primary{border:none;background-color:#00a99d}.btn-primary:focus,.btn-primary.focus{color:#fff;background-color:#049b8c}.btn-primary:hover{color:#fff;background-color:#048276;-webkit-transition:ease-in-out 125ms;transition:125ms}.btn-primary:active,.btn-primary.active,.open>.dropdown-toggle.btn-primary{background-color:#00a99d}.btn-primary:active,.btn-primary.active,.btn-secondary{color:#333;background-color:#fff;border-color:#ccc}.open>.dropdown-toggle.btn-primary{color:#fff;background-color:#00766e;border-color:#11aa9b}a:hover{-webkit-transition:ease-in-out 125ms;-moz-transition:ease-in-out 125ms;-o-transition:ease-in-out 125ms;transition:ease-in-out 125ms}p{margin:0 0 5px 0}.glyphicon{right:5px;line-height:1.1}small{font-size:80%}html,body{font-size:14px;line-height:1.5;margin:0px 0px 50px 0px;padding:0}.navbar{display:flex;justify-content:space-between;border-bottom:2px solid #00a99d}.nav-button:active,.nav-button:focus,.nav-button:hover{background-color:#fff;border:1px solid #028e80;color:#049b8c}#hyphy-logo{width:250px;height:50px}toolbar-dropdown::before{content:"";height:68px;display:inline-block;vertical-align:middle}.dropup,.dropdown{position:relative;float:right}.dropdown-menu{position:absolute;top:75%;left:0;z-index:1000;display:none;float:left;padding:0;min-width:160px;margin:2px 0 0;list-style:none;font-size:14px;text-align:left;background-color:#fff;border:1px solid #ccc;border:3px solid #00a99d;border-radius:0px;-webkit-box-shadow:0 6px 12px rgba(0,0,0,.175);box-shadow:0 6px 12px rgba(0,0,0,.175);background-clip:padding-box}.dropdown-menu>li>a{display:block;padding:8px 20px;clear:both;font-family:"Montserrat",sans-serif;font-size:13.5px;font-weight:500;line-height:1.5;color:#444;white-space:nowrap}.dropdown-menu>li:hover,.dropdown-menu>li:hover{text-decoration:none;color:#fff;background-color:#00a99d}.main-result{margin-bottom:0px}#results,#summary-tab{margin-top:0}.summary-method-name{font-size:19px;font-weight:400}#slac-table .col-md-12 p{margin:10px 23px}.input-group-addon{padding:6px 12px;border:none;text-align:center}table{margin-top:0;margin-bottom:20px}.site-table{margin:10px 0px 0px 0px}.fade-site-table{cursor:pointer}@media(min-width: 1200px){.fade-site-table{table-layout:fixed}}.table>thead>tr>th{border-bottom:2px solid #a1c6b3;text-align:center}.table>caption+thead>tr:first-child>th,.table>colgroup+thead>tr:first-child>th,.table>thead:first-child>tr:first-child>th,.table>caption+thead>tr:first-child>td,.table>colgroup+thead>tr:first-child>td,.table>thead:first-child>tr:first-child>td{color:#00a99d}table.dm-table>thead>tr>th,table.dm-table>thead>th{text-align:center}.table>thead>tr>th,.table>tbody>tr>th,.table>tfoot>tr>th,.table>thead>tr>td,.table>tbody>tr>td,.table>tfoot>tr>td,.table-condensed>tfoot>tr>td{font-family:"Noto Sans",sans-serif;text-align:center;vertical-align:middle}.table>thead>tr>td.info,.table>tbody>tr>td.info,.table>tfoot>tr>td.info,.table>thead>tr>th.info,.table>tbody>tr>th.info,.table>tfoot>tr>th.info,.table>thead>tr.info>td,.table>tbody>tr.info>td,.table>tfoot>tr.info>td,.table>thead>tr.info>th,.table>tbody>tr.info>th,.table>tfoot>tr.info>th{background-color:#eaf4f2}.table-condensed>tbody>tr>th{padding:15px 9px;border-top:none}th,td{font-weight:bold;text-align:center;padding:15px 9px}.panel{box-shadow:none;border:none;margin-bottom:40px}.panel-title{font-family:"Montserrat",sans-serif}.panel-default>.panel-heading,.panel-body{text-align:center;border:none}.panel-heading{padding:20px 15px;background-color:#eaf4f2}.panel-heading>.btn-group>.btn{height:28px}.clear-padding{padding:0}.popover{font-family:"Montserrat",sans-serif;font-size:13.5px;letter-spacing:.25px;color:#444;border:3px solid #00a99d;border-radius:0;-webkit-box-shadow:0 6px 12px rgba(0,0,0,.45);box-shadow:0 6px 12px rgba(0,0,0,.45)}.popover-title{padding:12px 14px 0px 14px;margin:0;font-family:"Montserrat",sans-serif;font-size:15.5px;font-weight:700;background-color:#fff;color:#038c7f;letter-spacing:.25px}.popover-title:after{content:"";display:block;width:100%;height:3px;margin:9px auto 0px auto;background-color:#91cec2}.popover-content ul{list-style:none;padding:0}.popover-content ul li{font-family:"Noto Sans",sans-serif;font-size:13.5px}.popover.bottom>.arrow{top:-16px;left:50%;margin-left:-11px;border-top-width:3px;border-bottom-color:#00a99d}.popover.bottom>.arrow:after{top:1px;margin-left:-10px;content:" ";border-top-width:3px;border-bottom-color:#fff}.form-group label{font-family:"Noto Sans",sans-serif;font-size:15px;margin:10px 0 5px 0;color:#00a99d}#input-info{padding:5px 20px 5px 23px}.divider{padding:0px 12px}.blue-badge,.white-badge,.red-badge{border-radius:12px;color:#fff;font-size:11px;padding:2px 8px 3px 8px;font-family:"Montserrat",sans-serif;font-weight:500;letter-spacing:.25px}.blue-badge{background-color:#3d4699}.white-badge{background-color:#fff;color:#333;border:1px solid #ccc}.red-badge{background-color:#cc4242}svg g>text{font-family:"Noto Sans" sans-serif;font-size:13px;font-weight:600}.form-control{display:inline-block;height:38px;border-width:2px;width:100px}.form-group{margin-bottom:0}.hyphy-busted-site-table{margin-top:0px}#prop-chart{width:75%}.dm-table-header{margin-top:30px;margin-bottom:0;padding:12px 10px 13px 5px;border-bottom:2px solid}input[type=range]{width:100%}.list-group-item-heading{margin-top:68px}.table-responsive{order:none;overflow-x:visible}table table{margin-bottom:0}tr.highlight{background-color:#bfbfbf}.fa-rotate-45{-webkit-transform:rotate(45deg);-moz-transform:rotate(45deg);-ms-transform:rotate(45deg);-o-transform:rotate(45deg);transform:rotate(45deg)}.fa-rotate-135{-webkit-transform:rotate(135deg);-moz-transform:rotate(135deg);-ms-transform:rotate(135deg);-o-transform:rotate(135deg);transform:rotate(135deg)}.hyphy-omega-line-reference{stroke:#000;stroke-width:6px;opacity:1}.hyphy-omega-bar line,.hyphy-omega-bar path{fill:none;stroke:#000;shape-rendering:crispEdges}.hyphy-omega-line{stroke:#000;stroke-width:6px}.hyphy-axis path,.hyphy-axis line{fill:none;stroke:#000;shape-rendering:crispEdges}.hyphy-axis-label{font:18px sans-serif}.hyphy-axis text{font:12px sans-serif}.hyphy-weight-label{font:14px sans-serif;text-anchor:middle}.hyphy-neutral-line{stroke-dasharray:2,10;stroke:#000;stroke-width:1px}.hyphy-displacement-line{fill:none;stroke:#000;opacity:.7;stroke-width:1.5px}#hyphy-chart-image{display:none}#hyphy-chart-canvas{display:none}.hyphy-busted-axis path,.hyphy-busted-axis line{fill:none;stroke:#000;shape-rendering:crispEdges}.hyphy-busted-area{fill:#4682b4}.hyphy-busted-site-table{margin-top:20px}.hyphy-busted-btn-export{margin:5px}.hyphy-busted-dist-export{margin-left:5px}.hyphy-blue{color:#1f77b4}.hyphy-red{color:#d62728}.hyphy-navy{color:#001f3f}.hyphy-omega-chart-btn{color:#000;padding-left:15px;padding-right:15px;padding-top:5px;padding-bottom:5px}.fixed{position:fixed}.nav-pills a{color:#00a99d}.nav-pills a:hover{color:#7ec6b3}.nav-pills>li.active>a{color:#fff;background-color:#00a99d}.nav-pills>li.active>a:hover{background-color:#7ec6b3}#summary-method-name{color:#00a99d;font-family:"Montserrat"}.popover{color:#111}table.dm-table>thead>tr>th,table.dm-table>thead>th{font-family:"Noto Sans",sans-serif;color:#00a99d}.hyphy-highlight{color:#00a99d}div.main-result a{color:#00a99d}#summary-pmid{color:#00a99d}#input-info{padding:0px 20px 0px 23px;font-family:"Montserrat",sans-serif;font-weight:500}div#input-info span{font-family:"Montserrat";font-weight:700}div#input-info p{font-family:"Montserrat";font-weight:600;font-size:14px}.main-result{border-color:#00a99d;padding:10px 10px 10px 10px}div.main-result p{font-family:"Noto Sans";color:#545454;font-size:17px}div.main-result hr{margin-top:10px;margin-bottom:10px}tbody>tr{background-color:#dddedf}p.description{font-family:"Noto Sans";line-height:1.5;font-size:15px;color:#545454;padding:0px 20px;margin:0px 0px 20px 0px}.v-align{float:none;display:inline-block;vertical-align:middle}.popover{max-width:100%}.bs-docs-sidebar .nav{min-width:185px}.nav-tabs{border-bottom:none}.bs-docs-sidebar .nav>li>a{display:block;font-family:"Montserrat",sans-serif;font-size:15px;font-weight:500;color:#777;letter-spacing:.75px;border-right:3px solid #ccc;padding:10px 0px 10px 34px;border-radius:0}.nav-pills .nav-link.active,.nav-pills .show>.nav-link,.nav-link:hover{color:#00a99d;border-color:#00a99d;background-color:#fff;border-top:none;border-bottom:none;border-left:none}.bs-docs-sidebar .nav>li>a:hover{color:#00a99d;border-right-color:#aaa}.bs-docs-sidebar .nav>.active>ul.nav{display:block}.bs-docs-section{margin-bottom:60px}.bs-docs-example{margin-top:20px;margin-bottom:20px}.hyphy-row{margin-top:20px}.axis path,.axis line{fill:none;stroke:#000;shape-rendering:crispEdges}g{font:10px sans-serif}.dot1{stroke:#000}.dot2{stroke:#000}.dot3{stroke:#000}.dot4{stroke:#000}.dot5{stroke:#000}.dot6{stroke:#000}.dot7{stroke:#000}.dot8{stroke:#000}.dot9{stroke:#000}.dot10{stroke:#000}.dot11{stroke:#000}.dot12{stroke:#000}.dot13{stroke:#000}.dot14{stroke:#000}.dot15{stroke:#000}.dot16{stroke:#000}.dot17{stroke:#000}.dot18{stroke:#000}.dot19{stroke:#000}.dot20{stroke:#000}circle.property1{fill:#fff;stroke:#000}line.sep_line{fill:#fff;stroke:#000;stroke-width:2}circle.property2{fill:#fff;stroke:red}circle.property3{fill:#fff;stroke:#ff7400}circle.property4{fill:#fff;stroke:#099}circle.property5{fill:#fff;stroke:#0c0}.rule line{stroke:#eee;shape-rendering:crispEdges}.rule line.axis{stroke:#000}.grid .tick{stroke:#d3d3d3;opacity:.7}.grid path{stroke-width:0}.exch_rate{fill:#fff;stroke:#ccc}div.dc-chart{float:left}.dc-chart rect.bar{stroke:none;cursor:pointer}.dc-chart rect.bar:hover{fill-opacity:.5}.dc-chart rect.stack1{stroke:none;fill:red}.dc-chart rect.stack2{stroke:none;fill:green}.dc-chart rect.deselected{stroke:none;fill:#ccc}.dc-chart .pie-slice{fill:#fff;font-size:12px;cursor:pointer}.dc-chart .pie-slice :hover{fill-opacity:.8}.dc-chart .selected path{stroke-width:3;stroke:#ccc;fill-opacity:1}.dc-chart .deselected path{stroke:none;fill-opacity:.5;fill:#ccc}.axis line,.dc-chart .axis path{fill:none;stroke:#000;shape-rendering:crispEdges}.dc-chart .axis text{font:10px sans-serif}.dc-chart .grid-line,.dc-chart .grid-line line{fill:none;stroke:#ccc;opacity:.5;shape-rendering:crispEdges}.dc-chart .brush rect.background{z-index:-999}.dc-chart .brush rect.extent{fill:#4682b4;fill-opacity:.125}.dc-chart .brush .resize path{fill:#eee;stroke:#666}.dc-chart path.line{fill:none;stroke-width:1.5px}.dc-chart circle.dot{stroke:none}.dc-chart g.dc-tooltip path{fill:none;stroke:gray;stroke-opacity:.8}.dc-chart path.area{fill-opacity:.3;stroke:none}.dc-chart .node{font-size:.7em;cursor:pointer}.dc-chart .node :hover{fill-opacity:.8}.dc-chart .selected circle{stroke-width:3;stroke:#ccc;fill-opacity:1}.dc-chart .deselected circle{stroke:none;fill-opacity:.5;fill:#ccc}.dc-chart .bubble{stroke:none;fill-opacity:.6}.dc-data-count{float:right;margin-top:15px;margin-right:15px}.dc-data-count .filter-count,.dc-data-count .total-count{color:#3182bd;font-weight:700}.dc-chart g.state{cursor:pointer}.dc-chart g.state :hover{fill-opacity:.8}.dc-chart g.state path{stroke:#fff}.dc-chart g.deselected path{fill:gray}.dc-chart g.deselected text{display:none}.dc-chart g.county path{stroke:#fff;fill:none}.dc-chart g.debug rect{fill:blue;fill-opacity:.2}.dc-chart g.row rect{fill-opacity:.8;cursor:pointer}.dc-chart g.row rect:hover{fill-opacity:.6}.dc-chart g.row text{fill:#fff;font-size:12px;cursor:pointer}.dc-chart .highlight,.dc-chart path.highlight{fill-opacity:1;stroke-opacity:1}.dc-legend{font-size:11px}.dc-legend-item{cursor:pointer}.dc-chart g.axis text{-webkit-user-select:none;-moz-user-select:none;-ms-user-select:none;-o-user-select:none;user-select:none;pointer-events:none}.dc-chart path.highlight{stroke-width:3}.dc-chart .fadeout{fill-opacity:.2;stroke-opacity:.2}.dc-chart path.dc-symbol{fill-opacity:.5;stroke-opacity:.5}.dc-hard .number-display{float:none}.tree-selection-brush .extent{fill-opacity:.05;stroke:#fff;shape-rendering:crispEdges}.tree-scale-bar text{font:sans-serif}.tree-scale-bar line,.tree-scale-bar path{fill:none;stroke:#000;shape-rendering:crispEdges}.node circle,.node ellipse,.node rect{fill:#4682b4;stroke:#000;stroke-width:.5px}.internal-node circle,.internal-node ellipse,.internal-node rect{fill:#ccc;stroke:#000;stroke-width:.5px}.node{font:10px sans-serif}.node-selected{fill:red}.node-collapsed circle,.node-collapsed ellipse,.node-collapsed rect{fill:#000}.node-tagged{fill:blue}.branch{fill:none;stroke:#999;stroke-width:2px}.clade{fill:#1f77b4;stroke:#444;stroke-width:2px;opacity:.5}.branch-selected{stroke:red;stroke-width:3px}.branch-tagged{stroke:blue;stroke-dasharray:10,5;stroke-width:2px}.branch-tracer{stroke:#bbb;stroke-dasharray:3,4;stroke-width:1px}.branch-multiple{stroke-dasharray:5,5,1,5;stroke-width:3px}.branch:hover{stroke-width:10px}.internal-node circle:hover,.internal-node ellipse:hover,.internal-node rect:hover{fill:#000;stroke:#ccc}.tree-widget{text-align:center}.summary-method-name{font-family:"Montserrat",sans-serif;font-size:21px;color:#00a99d}.results-summary{font-family:"Montserrat",sans-serif;font-size:48px;color:#00a99d;font-weight:600;line-height:1}.nav-stacked{padding-top:45px}a:hover{transition:ease-in-out 175ms}li.active>a:hover{border-right:3px solid #00a99d}#tree-tab{margin-bottom:20px}#summary-tab{margin-top:-20px}.nav-button{display:inline-block;padding:0px 24px;height:38px;margin-left:7px;background-color:transparent;font-family:"Montserrat",sans-serif;font-size:13.5px;font-weight:500;line-height:38px;vertical-align:middle;letter-spacing:.25px;color:#00a99d;box-sizing:border-box;border:1px solid #27c1a7}.nav-button:active,.nav-button:focus,.nav-button:hover{background-color:#00a99d;text-decoration:none;color:#fff;border:1px solid transparent}.divider{font-size:21px;padding:0px 10px;color:#91cec2;vertical-align:sub}.brush .extent{stroke:#fff;fill-opacity:.125;shape-rendering:crispEdges}.modal .modal-dialog{width:850px;max-width:850px}.fa-bar-chart:hover{color:#00a99d}.positive-selection-row{color:#fff;background-color:#00a99d;text-align:center}.negative-selection-row{color:#fff;background-color:#333;text-align:center}#tree_tooltip{color:#fff;display:block;position:fixed;padding:20px;max-width:200px;background-color:rgba(0,0,0,.85);-webkit-border-radius:12px;-moz-border-radius:12px;border-radius:12px;display:none}.hyphy-export-dropdown{float:left}.custom-file-upload{border:1px solid #ccc;display:inline-block;padding:6px 12px;cursor:pointer}.input-group-btn,.btn-group-toggle{display:inline-block}.btn-sm,.btn-group-sm>.btn{font-size:1rem}.hyphy-vision-datamonkey-table{overflow-x:scroll}.btn-export-csv-round{background-color:#00a99d;color:#fff;letter-spacing:1px;text-align:center;line-height:20px;font-family:"Noto Sans",sans-serif;font-size:15px;font-weight:600;border:2px solid #00a99d;border-radius:40px;transition:all .3s ease 0s}.btn-export-chart-png{background-color:#00a99d;color:#fff;letter-spacing:1px;text-align:center;line-height:20px;font-family:"Noto Sans",sans-serif;font-size:10px;font-weight:600;border:2px solid #fff}.btn-export-chart-svg{background-color:#00a99d;color:#fff;letter-spacing:1px;text-align:center;line-height:20px;font-family:"Noto Sans",sans-serif;font-size:10px;font-weight:600;border:2px solid #fff}.gard{font-family:"Source Serif Pro",Iowan Old Style,Apple Garamond,Palatino Linotype,Times New Roman,"Droid Serif",Times,serif,Apple Color Emoji,Segoe UI Emoji,Segoe UI Symbol}.gard td{background:#fff;background-color:#fff !important}/*!\n * Font Awesome Free 5.15.2 by @fontawesome - https://fontawesome.com\n * License - https://fontawesome.com/license/free (Icons: CC BY 4.0, Fonts: SIL OFL 1.1, Code: MIT License)\n */.fa,.fas,.far,.fal,.fad,.fab{-moz-osx-font-smoothing:grayscale;-webkit-font-smoothing:antialiased;display:inline-block;font-style:normal;font-variant:normal;text-rendering:auto;line-height:1}.fa-lg{font-size:1.3333333333em;line-height:.75em;vertical-align:-0.0667em}.fa-xs{font-size:.75em}.fa-sm{font-size:.875em}.fa-1x{font-size:1em}.fa-2x{font-size:2em}.fa-3x{font-size:3em}.fa-4x{font-size:4em}.fa-5x{font-size:5em}.fa-6x{font-size:6em}.fa-7x{font-size:7em}.fa-8x{font-size:8em}.fa-9x{font-size:9em}.fa-10x{font-size:10em}.fa-fw{text-align:center;width:1.25em}.fa-ul{list-style-type:none;margin-left:2.5em;padding-left:0}.fa-ul>li{position:relative}.fa-li{left:-2em;position:absolute;text-align:center;width:2em;line-height:inherit}.fa-border{border:solid .08em #eee;border-radius:.1em;padding:.2em .25em .15em}.fa-pull-left{float:left}.fa-pull-right{float:right}.fa.fa-pull-left,.fas.fa-pull-left,.far.fa-pull-left,.fal.fa-pull-left,.fab.fa-pull-left{margin-right:.3em}.fa.fa-pull-right,.fas.fa-pull-right,.far.fa-pull-right,.fal.fa-pull-right,.fab.fa-pull-right{margin-left:.3em}.fa-spin{animation:fa-spin 2s infinite linear}.fa-pulse{animation:fa-spin 1s infinite steps(8)}@keyframes fa-spin{0%{transform:rotate(0deg)}100%{transform:rotate(360deg)}}.fa-rotate-90{-ms-filter:"progid:DXImageTransform.Microsoft.BasicImage(rotation=1)";transform:rotate(90deg)}.fa-rotate-180{-ms-filter:"progid:DXImageTransform.Microsoft.BasicImage(rotation=2)";transform:rotate(180deg)}.fa-rotate-270{-ms-filter:"progid:DXImageTransform.Microsoft.BasicImage(rotation=3)";transform:rotate(270deg)}.fa-flip-horizontal{-ms-filter:"progid:DXImageTransform.Microsoft.BasicImage(rotation=0, mirror=1)";transform:scale(-1, 1)}.fa-flip-vertical{-ms-filter:"progid:DXImageTransform.Microsoft.BasicImage(rotation=2, mirror=1)";transform:scale(1, -1)}.fa-flip-both,.fa-flip-horizontal.fa-flip-vertical{-ms-filter:"progid:DXImageTransform.Microsoft.BasicImage(rotation=2, mirror=1)";transform:scale(-1, -1)}:root .fa-rotate-90,:root .fa-rotate-180,:root .fa-rotate-270,:root .fa-flip-horizontal,:root .fa-flip-vertical,:root .fa-flip-both{filter:none}.fa-stack{display:inline-block;height:2em;line-height:2em;position:relative;vertical-align:middle;width:2.5em}.fa-stack-1x,.fa-stack-2x{left:0;position:absolute;text-align:center;width:100%}.fa-stack-1x{line-height:inherit}.fa-stack-2x{font-size:2em}.fa-inverse{color:#fff}.fa-500px:before{content:""}.fa-accessible-icon:before{content:""}.fa-accusof
```
