## supplementary data file 3 for "Repeated truncation of a modular antimicrobial peptide gene for neural context": moment_timeline.bcb1cafa923482f4826e32741fe16a98.js

```
(window.__twttrll=window.__twttrll||[]).push([[0],{168:function(e,t,r){var i=r(39),n=r(170),o=r(7);(i=Object.create(i)).build=o(i.build,null,n),e.exports=i},169:function(e,t,r){var i=r(42),n=r(37),o=r(38),s=r(0),a=r(7),c=r(33),u=r(5),l=r(174);e.exports=function(e){e.params({partner:{fallback:a(c.val,c,"partner")}}),e.define("scribeItems",function(){return{}}),e.define("scribeNamespace",function(){return{client:"tfw"}}),e.define("scribeData",function(){return{widget_origin:o.rootDocumentLocation(),widget_frame:o.isFramed()&&o.currentDocumentLocation(),widget_partner:this.params.partner,widget_site_screen_name:l(c.val("site")),widget_site_user_id:u.asNumber(c.val("site:id")),widget_creator_screen_name:l(c.val("creator")),widget_creator_user_id:u.asNumber(c.val("creator:id"))}}),e.define("scribe",function(e,t,r){e=s.aug(this.scribeNamespace(),e||{}),t=s.aug(this.scribeData(),t||{}),i.scribe(e,t,!1,r)}),e.define("scribeInteraction",function(e,t,r){var i=n.extractTermsFromDOM(e.target);i.action=e.type,"url"===i.element&&(i.element=n.clickEventElement(e.target)),this.scribe(i,t,r)})}},170:function(e,t,r){var i=r(40),n=r(0),o=r(171);function s(){i.apply(this,arguments),this.Widget=this.Component}s.prototype=Object.create(i.prototype),n.aug(s.prototype,{factory:o,build:function(){return i.prototype.build.apply(this,arguments)},selectors:function(e){var t=this.Widget.prototype.selectors;e=e||{},this.Widget.prototype.selectors=n.aug({},e,t)}}),e.exports=s},171:function(e,t,r){var i=r(6),n=r(35),o=r(41),s=r(0),a=r(7),c=r(172),u="twitter-widget-";e.exports=function(){var e=o();function t(t,r){e.apply(this,arguments),this.id=u+c(),this.sandbox=r}return t.prototype=Object.create(e.prototype),s.aug(t.prototype,{selectors:{},hydrate:function(){return i.resolve()},prepForInsertion:function(){},render:function(){return i.resolve()},show:function(){return i.resolve()},resize:function(){return i.resolve()},select:function(e,t){return 1===arguments.length&&(t=e,e=this.el),e?(t=this.selectors[t]||t,s.toRealArray(e.querySelectorAll(t))):[]},selectOne:function(){return this.select.apply(this,arguments)[0]},selectLast:function(){return this.select.apply(this,arguments).pop()},on:function(e,t,r){var i,o=this.el;this.el&&(e=(e||"").split(/\s+/),2===arguments.length?r=t:i=t,i=this.selectors[i]||i,r=a(r,this),e.forEach(i?function(e){n.delegate(o,e,i,r)}:function(e){o.addEventListener(e,r,!1)}))}}),t}},172:function(e,t){var r=0;e.exports=function(){return String(r++)}},173:function(e,t,r){var i=r(5),n=r(0);e.exports=function(e){e.define("widgetDataAttributes",function(){return{}}),e.define("setDataAttributes",function(){var e=this.sandbox.sandboxEl;n.forIn(this.widgetDataAttributes(),function(t,r){i.hasValue(r)&&e.setAttribute("data-"+t,r)})}),e.after("render",function(){this.setDataAttributes()})}},174:function(e,t){e.exports=function(e){return e&&"@"===e[0]?e.substr(1):e}},176:function(e,t,r){var i=r(73);e.exports=function(e){return e?(e=Array.isArray(e)?e:[e]).reduce(function(e,t){var r=t.getAttribute("data-tweet-id"),n=t.getAttribute("data-rendered-tweet-id")||r;return t.getAttribute("data-tweet-item-type")===i.QUOTE_TWEET.toString()?e[r]={item_type:i.QUOTE_TWEET}:r===n?e[n]={item_type:i.TWEET}:r&&(e[n]={item_type:i.RETWEET,target_type:i.TWEET,target_id:r}),e},{}):{}}},177:function(e,t,r){var i=r(71),n=r(180),o=r(7),s=r(72);function a(e,t,r,o){var a,c;return r=function(e){return"dark"===e?"dark":"light"}(r),a=i.isRtlLang(t)?"rtl":"ltr",c=[e,o?n.holdback_css:n.css,r,a,"css"].join("."),s.resourceBaseUrl+(o?"/holdback":"")+"/css/"+c}e.exports={dmButton:function(){return s.resourceBaseUrl+"/css/"+["dm_button",n.css,"css"].join(".")},tweet:o(a,null,"tweet"),timeline:o(a,null,"timeline"),video:o(a,null,"video"),moment:o(a,null,"moment"),grid:o(a,null,"grid"),periscopeOnAir:function(){return s.resourceBaseUrl+"/css/"+["periscope_on_air",n.css,"css"].join(".")}}},178:function(e,t,r){var i=r(34),n=r(168),o=r(176);e.exports=n.couple(r(169),function(e){e.selectors({tweetIdInfo:".js-tweetIdInfo",quotedTweetInfo:'[data-tweet-item-type="23"]'}),e.define("scribeClickInteraction",function(e,t){var r=i.closest(this.selectors.tweetIdInfo,t,this.el),n=r&&r.querySelector(this.selectors.quotedTweetInfo);this.scribeInteraction(e,function(e,t){var r;if(e)return r=o(t?[e,t]:[e]),{item_ids:Object.keys(r),item_details:r}}(r,n))}),e.after("render",function(){this.on("click","A",this.scribeClickInteraction),this.on("click","BUTTON",this.scribeClickInteraction)})})},179:function(e,t,r){var i=r(4),n=r(5),o=i.createElement("div");e.exports=function(e){return n.isNumber(e)&&(e+="px"),o.style.width="",o.style.width=e,o.style.width||null}},180:function(e){e.exports={css:"32f7f89e2e680ebfe3f4cfefb27966ae",holdback_css:"ec70a6a3eabf66ea7567c1f46719df0f"}},181:function(e,t,r){var i=r(183),n=r(36),o=r(3),s=r(6),a="data-url-ref-attrs-injected";e.exports=function(e){var t={};e.define("injectRefUrlParams",function(e){return o.isTwitterURL(e.href)?e.getAttribute(a)?s.resolve():(t={twcamp:this.params.productName,twterm:this.params.id,twcon:e.getAttribute("data-twcon")},n.getActiveExperimentDataString().then(function(r){e.setAttribute(a,!0),t.twgr=r,e.href=i(e.href,t)}.bind(this)).catch(function(){e.setAttribute(a,!0),e.href=i(e.href,t)}.bind(this))):s.resolve()}),e.after("render",function(){this.on("click","A",function(e,t){this.injectRefUrlParams(t)})})}},183:function(e,t,r){var i=r(11),n=r(38),o="^",s="|",a="twsrc",c="twterm",u="twcamp",l="twgr",d="twcon";function h(e,t){return e+o+t}e.exports=function(e,t){var r=[h(a,"tfw")];return t&&(r=r.concat(function(e){var t=[];return e.twcamp&&t.push(h(u,e.twcamp)),e.twterm&&t.push(h(c,e.twterm)),e.twgr&&t.push(h(l,e.twgr)),e.twcon&&t.push(h(d,e.twcon)),t}(t))),function(e,t){return i.url(e,{ref_src:t,ref_url:n.rootDocumentLocation()})}(e,function(e){return e.reduce(function(e,t){return e+s+t})}(r))}},184:function(e,t,r){var i=r(37),n=r(168),o=r(186);e.exports=n.couple(r(169),function(e){e.selectors({inViewportMarker:".js-inViewportScribingTarget"}),e.define("scribeInViewportSeen",function(e,t){var r=i.extractTermsFromDOM(e);r.action="seen",this.scribe(r,t)}),e.after("show",function(){var e=this.selectors.inViewportMarker;this.select(e).forEach(function(e){e&&o.inViewportOnce(e,this.sandbox.sandboxEl,function(){this.scribeInViewportSeen(e,this.scribeItems())}.bind(this))},this)})})},185:function(e,t,r){var i=r(1),n=r(0),o=r(43),s=r(11),a=r(78),c=r(21),u=r(211),l=r(10),d="https://pbs.twimg.com/cards/player-placeholder",h=/max-width:\s*([\d.]+px)/,f=/top:\s*(-?[\d.]+%)/,p=/left:\s*(-?[\d.]+%)/,m=/padding-bottom:\s*([\d.]+%)/,b={64:"tiny",120:"120x120",240:"240x240",360:"360x360",680:"small",900:"900x900",1200:"medium",2048:"large",4096:"4096x4096"},g=Object.keys(b).sort(function(e,t){return e-t}),v=2;function w(e,t){e.getAttribute("data-image")===d?e.src=d+".png":e.getAttribute("data-image")?x(e,t):I(e,t)}function y(e,t,r){var i,n,o,s,a;if(n=C(e),o=t.split(",").map(function(e){return new function(e){var t=e.split(" ");this.url=decodeURIComponent(t[0].trim()),this.width=+t[1].replace(/w$/,"").trim()}(e.trim())}),r)for(a=0;a<o.length;a++)o[a].url===r&&(i=o[a]);return s=o.reduce(function(e,t){return t.width<e.width&&t.width>=n?t:e},o[0]),i&&i.width>s.width?i:s}function x(e,t){var r,i,n,o,c,u,l;i=(r=s.decodeURL(e.src).name)&&a(g,function(e){if(b[e]===r)return e}),n=function(e){return{width:parseInt(e.getAttribute("width")),height:parseInt(e.getAttribute("height"))||1}}(e),i>=(c=((l=n).height>l.width?C(t)*n.height/n.width:C(t))||680)||(o=e.getAttribute("data-image"),u=a(g,function(e){if(e>=c)return e})||4096,e.src=s.url(o,{format:e.getAttribute("data-image-format")||"jpg",name:b[u]}))}function C(e){return i.devicePixelRatio?e*Math.min(i.devicePixelRatio,v):e}function I(e,t){var r,i=e.getAttribute("data-srcset"),n=e.src;i&&(r=y(t,i,n),e.src=r.url)}function E(e,t,r){e&&(n.toRealArray(e.querySelectorAll(".NaturalImage-image")).forEach(function(e){r(function(){w(e,t)})}),n.toRealArray(e.querySelectorAll(".CroppedImage-image")).forEach(function(e){r(function(){w(e,t/2)})}),n.toRealArray(e.querySelectorAll("img.autosized-media")).forEach(function(e){r(function(){w(e,t),e.removeAttribute("width"),e.removeAttribute("height")})}))}function _(e,t,r,i){e&&n.toRealArray(e.querySelectorAll("iframe.autosized-media, .wvp-player-container")).forEach(function(e){var n=k(e.getAttribute("data-width"),e.getAttribute("data-height"),u.effectiveWidth(e.parentElement)||t,r);i(function(){e.setAttribute("width",n.width),e.setAttribute("height",n.height),l.present(e,"wvp-player-container")?(e.style.width=n.width,e.style.height=n.height):(e.width=n.width,e.height=n.height)})})}function k(e,t,r,i,n,o){return r=r||e,i=i||t,n=n||0,o=o||0,e>r&&(t*=r/e,e=r),t>i&&(e*=i/t,t=i),e<n&&(t*=n/e,e=n),t<o&&(e*=o/t,t=o),{width:Math.floor(e),height:Math.floor(t)}}function A(e,t,r,i){n.toRealArray(e.querySelectorAll(t)).forEach(function(e){var t=e.getAttribute("style")||e.getAttribute("data-style"),n=i.test(t)&&RegExp.$1;n&&(e.setAttribute("data-csp-fix",!0),e.style[r]=n)})}e.exports={scaleDimensions:k,retinize:function(e,t){t=void 0!==t?!!t:c.retina(),n.toRealArray(e.getElementsByTagName("IMG")).forEach(function(e){var r=e.getAttribute("data-src-1x")||e.getAttribute("src"),i=e.getAttribute("data-src-2x");t&&i?e.src=i:r&&(e.src=r)})},setSrcForImgs:E,sizeIframes:_,constrainMedia:function(e,t,r,i){E(e,t,i),_(e,t,r,i)},fixMediaCardLayout:function(e){o.inlineStyle()||(A(e,".MediaCard-widthConstraint","maxWidth",h),A(e,".MediaCard-mediaContainer","paddingBottom",m),A(e,".CroppedImage-image","top",f),A(e,".CroppedImage-image","left",p))},__setSrcFromSet:I,__setSrcFromImage:x,__setImageSrc:w}},186:function(e,t,r){var i=r(187),n=r(44),o=r(188),s=r(1),a=r(21),c=function(e){return(s.requestIdleCallback||s.requestAnimationFrame||function(e){e()})(e)},u=function(){this.observers=[]};u.prototype._register=function(e,t,r){var n,u=this;return a.hasIntersectionObserverSupport()?((n=new s.IntersectionObserver(function(e){e.forEach(function(e){e.intersectionRatio>=1&&(c(r),u._unregister(n))})},{threshold:1})).observe(e),n):(n={update:function(o,s){i(e,{viewportWidth:o,viewportHeight:s,sandboxEl:t})&&(r(),u._unregister(n))}},this.observers.push(n),1===this.observers.length&&(this.unlisten=o.addScrollListener(this._onViewportChange.bind(this))),this._onViewportChange(),n)},u.prototype._unregister=function(e){var t;a.hasIntersectionObserverSupport()&&e instanceof s.IntersectionObserver?e.disconnect():(t=this.observers.indexOf(e))>-1&&(this.observers.splice(t,1),0===this.observers.length&&this.unlisten&&this.unlisten())},u.prototype._onViewportChange=function(){n(c(function(){this._notify(o.getWidth(),o.getHeight())}.bind(this)),50,this)},u.prototype._notify=function(e,t){this.observers.forEach(function(r){r.update(e,t)})},u.prototype.inViewportOnce=function(e,t,r){return this._register(e,t,r)},e.exports=new u},187:function(e,t,r){var i=r(4),n=r(1);e.exports=function(e,t){var r,o,s,a;return o=(t=t||{}).viewportWidth||n.innerWidth,r=t.viewportHeight||n.innerHeight,s=e.getBoundingClientRect(),e.ownerDocument!==i&&t.sandboxEl&&(a=t.sandboxEl.getBoundingClientRect(),s={top:s.top+a.top,bottom:s.bottom+a.top,left:s.left+a.left,right:s.right+a.left}),s.top>=0&&s.left>=0&&s.bottom<=r&&s.right<=o}},188:function(e,t,r){var i=r(1),n={_addListener:function(e,t){var r=function(){t()};return i.addEventListener(e,r),function(){i.removeEventListener(e,r)}},addScrollListener:function(e){return this._addListener("scroll",e)},getHeight:function(){return i.innerHeight},getWidth:function(){return i.innerWidth}};e.exports=n},192:function(e,t,r){var i=r(18),n=r(185),o=375;e.exports=function(e){e.after("prepForInsertion",function(e){n.sizeIframes(e,this.sandbox.width,o,i.sync)}),e.after("resize",function(){n.sizeIframes(this.el,this.sandbox.width,o,i.write)})}},193:function(e,t,r){var i=r(43),n=(r(12),r(0)),o=/^([a-zA-Z-]+):\s*(.+)$/;function s(e){var t=(e.getAttribute("data-style")||"").split(";").reduce(function(e,t){var r,i,n;return o.test(t.trim())&&(r=RegExp.$1,i=RegExp.$2,e[(n=r,n.replace(/-(.)/g,function(e,t){return t.toUpperCase()}))]=i),e},{});0!==Object.keys(t).length&&(e.setAttribute("data-csp-fix","true"),n.forIn(t,function(t,r){e.style[t]=r}))}e.exports=function(e){e.selectors({cspForcedStyle:".js-cspForcedStyle"}),e.after("prepForInsertion",function(e){i.inlineStyle()||this.select(e,"cspForcedStyle").forEach(s)})}},194:function(e,t,r){var i=r(185);e.exports=function(e){e.after("prepForInsertion",function(e){i.retinize(e)})}},195:function(e,t,r){var i=r(18),n=r(185);e.exports=function(e){e.after("prepForInsertion",function(e){n.setSrcForImgs(e,this.sandbox.width,i.sync)}),e.after("resize",function(){n.setSrcForImgs(this.el,this.sandbox.width,i.write)})}},196:function(e,t){var r="data-iframe-title";e.exports=function(e){e.after("render",function(){var e=this.el.getAttribute(r);e&&this.sandbox.setTitle&&this.sandbox.setTitle(e)})}},197:function(e,t,r){var i=r(6),n=r(5),o="env-bp-",s=o+"min";function a(e){return e.every(n.isInt)}function c(e){var t=e.map(function(e){return{size:+e,className:o+e}}).sort(function(e,t){return e.size-t.size});return t.unshift({size:0,className:s}),t}e.exports=function(e){e.params({breakpoints:{required:!0,validate:a,transform:c}}),e.define("getClassForWidth",function(e){var t,r,i;for(r=this.params.breakpoints.length-1;r>=0;r--)if(e>(i=this.params.breakpoints[r]).size){t=i.className;break}return t}),e.after("initialize",function(){this.allBreakpoints=this.params.breakpoints.map(function(e){return e.className})}),e.define("recalculateBreakpoints",function(){var e=this.getClassForWidth(this.sandbox.width);return e&&this.sandbox.hasRootClass(e)?i.resolve():i.all([this.sandbox.removeRootClass(this.allBreakpoints),this.sandbox.addRootClass(e)])}),e.after("render",function(){return this.recalculateBreakpoints()}),e.after("resize",function(){return this.recalculateBreakpoints()})}},198:function(e,t,r){var i=r(2),n=r(77),o=r(213),s=null;function a(e,t){var r,i;if(i={scribeContext:(t=t||{}).scribeContext||{client:"tfw"},languageCode:t.languageCode,hideControls:t.hideControls||!1,addTwitterBranding:t.addBranding||!1,widgetOrigin:t.widgetOrigin,borderRadius:t.borderRadius,autoPlay:t.autoPlay},(r=n(e,".wvp-player-container")).length>0)return s&&o.setBaseUrl(s),{element:r[0],options:i}}e.exports={insertForTweet:function(e,t,r){var n,s=new i,c=a(e,r);if(c)return(n=o.createPlayerForTweet(c.element,t,c.options))?(s.resolve(n),s.promise):s.reject(new Error("unable to create tweet video player"))},insertForEvent:function(e,t,r){var n,s=new i,c=a(e,r);return c?((n=o.createPlayerForLiveVideo(c.element,t,c.options)).on("ready",function(){n.playPreview(),s.resolve(n)}),s.promise):s.reject(new Error("unable to initialize event video player"))},remove:function(e){var t=e.querySelector(".wvp-player-container"),r=t&&o.findPlayerForElement(t);if(r)return r.teardown()},find:function(e){return o.findPlayerForElement(e)}}},199:function(e,t,r){var i=r(34),n=r(214),o=r(168),s=r(36),a=r(37),c=r(0),u=r(6),l="data-click-to-open-target";e.exports=o.couple(r(169),function(e){e.selectors({clickToOpen:".js-clickToOpenTarget"}),e.define("shouldOpenTarget",function(e){var t=i.closest("A",e.target,this.el),r=i.closest("BUTTON",e.target,this.el),n=this.sandbox.hasSelectedText();return!t&&!r&&!n}),e.define("openTarget",function(e,t){var r=t&&t.getAttribute(l),i={twcamp:this.params.productName,twterm:this.params.id,twcon:t.getAttribute("data-twcon")};return r?s.getActiveExperimentDataString().then(function(t){i.twgr=t,n(r,i),this.scribeOpenClick(e)}.bind(this)).catch(function(){n(r,i),this.scribeOpenClick(e)}.bind(this)):u.resolve()}),e.define("attemptToOpenTarget",function(e,t){return this.shouldOpenTarget(e)?this.openTarget(e,t):u.resolve()}),e.define("scribeOpenClick",function(e){var t=a.extractTermsFromDOM(e.target),r={associations:a.formatTweetAssociation(t)},i=c.aug({},{element:"chrome",action:"click"},t);this.scribe(i,r)}),e.after("render",function(){this.on("click","clickToOpen",this.attemptToOpenTarget)})})},200:function(e,t,r){var i=r(34),n=r(33),o=r(74),s=r(38),a=r(11),c=r(3),u=r(7),l="data-url-params-injected";e.exports=function(e){e.params({productName:{required:!0},dataSource:{required:!1},related:{required:!1},partner:{fallback:u(n.val,n,"partner")}}),e.selectors({timeline:".timeline",tweetIdInfo:".js-tweetIdInfo"}),e.define("injectWebIntentParams",function(e){var t=i.closest(this.selectors.timeline,e,this.el),r=i.closest(this.selectors.tweetIdInfo,e,this.el);e.getAttribute(l)||(e.setAttribute(l,!0),e.href=a.url(e.href,{tw_w:this.params.dataSource&&this.params.dataSource.id,tw_i:r&&r.getAttribute("data-tweet-id"),tw_p:this.params.productName,related:this.params.related,partner:this.params.partner,query:t&&t.getAttribute("data-search-query"),profile_id:t&&t.getAttribute("data-profile-id"),original_referer:s.rootDocumentLocation()}))}),e.after("render",function(){this.on("click","A",function(e,t){c.isIntentURL(t.href)&&(this.injectWebIntentParams(t),o.open(t.href,this.sandbox.sandboxEl,e))})})}},201:function(e,t,r){var i=r(21);e.exports=function(e){e.before("render",function(){i.ios()&&this.sandbox.addRootClass("env-ios"),i.ie9()&&this.sandbox.addRootClass("ie9"),i.touch()&&this.sandbox.addRootClass("is-touch")})}},202:function(e,t,r){var i=r(168),n=r(215),o=1;e.exports=i.couple(r(169),function(e){var t={action:"dimensions"},r=new n(o);e.after("show",function(){var e;r.nextBoolean()&&(e={widget_width:this.sandbox.width,widget_height:this.sandbox.height},this.scribe(t,e))})})},203:function(e,t,r){var i=r(0);e.exports=function(e){var t={transparent:!1,hideBorder:!1,hideHeader:!1,hideFooter:!1,hideScrollBar:!1};return e=e||"",i.contains(e,"transparent")&&(t.transparent=!0),i.contains(e,"noborders")&&(t.hideBorder=!0),i.contains(e,"noheader")&&(t.hideHeader=!0),i.contains(e,"nofooter")&&(t.hideFooter=!0),i.contains(e,"noscrollbar")&&(t.hideScrollBar=!0),t}},204:function(e,t,r){var i=r(10),n=r(34),o=r(4),s=r(0),a="is-openedAbove",c="is-openedBelow";e.exports=function(e){e.selectors({shareMenuOpener:".js-showShareMenu",shareMenu:".timeline-ShareMenu",shareMenuTimelineHeader:".timeline-Header",shareMenuTimelineFooter:".timeline-Footer"}),e.define("getHeaderHeight",function(){var e=this.selectOne("shareMenuTimelineHeader");return e?e.getBoundingClientRect().height:0}),e.define("getFooterHeight",function(){var e=this.selectOne("shareMenuTimelineFooter");return e?e.getBoundingClientRect().height:0}),e.define("getShareMenuPositionClass",function(e){var t=e.getBoundingClientRect(),r=t.top-this.getHeaderHeight();return this.sandbox.height-t.bottom-this.getFooterHeight()<r?a:c}),e.after("render",function(){this.on("click","shareMenuOpener",function(e,t){var r,a=this,c=n.closest(this.selectors.shareMenu,e.target,this.el);function u(){i.remove(c,r),a.el.removeEventListener("click",u,!1),o.removeEventListener("click",u,!1)}e.preventDefault(),c&&(r=this.getShareMenuPositionClass(t),i.add(c,r),s.async(function(){a.el.addEventListener("click",u,!1),o.addEventListener("click",u,!1)}))})})}},211:function(e,t){e.exports={effectiveWidth:function e(t){return t&&1===t.nodeType?t.offsetWidth||e(t.parentNode):0}}},213:function(e,t,r){var i,n;n=this,void 0===(i=function(){return n.TwitterVideoPlayer=function(){var e="https://twitter.com",t=/^https?:\/\/([a-zA-Z0-9]+\.)*twitter.com(:\d+)?$/,r={suppressScribing:!1,squareCorners:!1,hideControls:!1,addTwitterBranding:!1},i=0,n={};function o(e){if(e&&e.data&&e.data.params&&e.data.params[0]){var t=e.data.params[0],r=e.data.id;if(t&&t.context&&"TwitterVideoPlayer"===t.context){var i=t.playerId;delete t.playerId,delete t.context;var o=n[i];o&&o.processMessage(e.data.method,t,r)}}}function s(t,r,s,a,c){var u=t.ownerDocument,l=u.defaultView;l.addEventListener("message",o),this.playerId=i++;var d={embed_source:"clientlib",player_id:this.playerId,rpc_init:1,autoplay:a.autoPlay};if(this.scribeParams={},this.scribeParams.suppressScribing=a&&a.suppressScribing,!this.scribeParams.suppressScribing){if(!a.scribeContext)throw"video_player: Missing scribe context";if(!a.scribeContext.client)throw"video_player: Scribe context missing client property";this.scribeParams.client=a.scribeContext.client,this.scribeParams.page=a.scribeContext.page,this.scribeParams.section=a.scribeContext.section,this.scribeParams.component=a.scribeContext.component}this.scribeParams.debugScribe=a&&a.scribeContext&&a.scribeContext.debugScribing,this.scribeParams.scribeUrl=a&&a.scribeContext&&a.scribeContext.scribeUrl,this.promotedLogParams=a.promotedContext,this.adRequestCallback=a.adRequestCallback,a.languageCode&&(d.language_code=a.languageCode),"tfw"===this.scribeParams.client&&(d.use_syndication_guest_id=!0),a.autoPlay&&(d.autoplay=1);var h=function(e,t,r){var i=Object.keys(r).filter(function(e){return null!=r[e]}).map(function(e){var t=r[e];return encodeURIComponent(e)+"="+encodeURIComponent(t)}).join("&");return i&&(i="?"+i),e+t+i}(e,r,d);return this.videoIframe=document.createElement("iframe"),this.videoIframe.setAttribute("src",h),this.videoIframe.setAttribute("allowfullscreen",""),this.videoIframe.setAttribute("allow","autoplay; fullscreen"),this.videoIframe.setAttribute("id",s),this.videoIframe.setAttribute("style","width: 100%; height: 100%; position: absolute; top: 0; left: 0;"),this.domElement=t,this.domElement.appendChild(this.videoIframe),n[this.playerId]=this,this.eventCallbacks={},this.emitEvent=function(e,t){var r=this.eventCallbacks[e];void 0!==r&&r.forEach(function(e){e.apply(this.playerInterface,[t])}.bind(this))},this.jsonRpc=function(e){var t=this.videoIframe.contentWindow;e.jsonrpc="2.0",t&&t.postMessage&&t.postMessage(JSON.stringify(e),"*")},this.jsonRpcCall=function(e,t){this.jsonRpc({method:e,params:t})},this.jsonRpcResult=function(e,t){this.jsonRpc({result:e,id:t})},this.processMessage=function(e,t,r){switch(e){case"requestPlayerConfig":this.jsonRpcResult({scribeParams:this.scribeParams,promotedLogParams:this.promotedLogParams,squareCorners:a.squareCorners,borderRadius:a.borderRadius,hideControls:a.hideControls,embedded:a.addTwitterBranding,widgetOrigin:a.widgetOrigin,ignoreFineGrainGeoblocking:a.ignoreFineGrainGeoblocking},r);break;case"videoPlayerAdStart":this.emitEvent("adStart",t);break;case"videoPlayerAdEnd":this.emitEvent("adEnd",t);break;case"videoPlayerPlay":this.emitEvent("play",t);break;case"videoPlayerPause":this.emitEvent("pause",t);break;case"videoPlayerMute":this.emitEvent("mute",t);break;case"videoPlayerUnmute":this.emitEvent("unmute",t);break;case"videoPlayerPlaybackComplete":this.emitEvent("playbackComplete",t);break;case"videoPlayerReady":this.emitEvent("ready",t);break;case"videoView":this.emitEvent("view",t);break;case"debugLoggingEvent":this.emitEvent("logged",t);break;case"requestDynamicAd":"function"==typeof this.adRequestCallback?this.jsonRpcResult(this.adRequestCallback(),r):this.jsonRpcResult({},r);break;case"videoPlayerError":t&&"NO_COOKIES_ERROR"===t.error_category?this.emitEvent("noCookiesError",t):t&&"GEOBLOCK_ERROR"===t.error_category&&this.emitEvent("geoblockError",t)}},this.playerInterface={on:function(e,t){return void 0===this.eventCallbacks[e]&&(this.eventCallbacks[e]=[]),this.eventCallbacks[e].push(t),this.playerInterface}.bind(this),off:function(e,t){if(void 0===t)delete this.eventCallbacks[e];else{var r=this.eventCallbacks[e];if(void 0!==r){var i=r.indexOf(t);i>-1&&r.splice(i,1)}}return this.playerInterface}.bind(this),play:function(){return this.jsonRpcCall("play"),this.playerInterface}.bind(this),pause:function(){return this.jsonRpcCall("pause"),this.playerInterface}.bind(this),mute:function(){return this.jsonRpcCall("mute"),this.playerInterface}.bind(this),unmute:function(){return this.jsonRpcCall("unmute"),this.playerInterface}.bind(this),playPreview:function(){return this.jsonRpcCall("autoPlayPreview"),this.playerInterface}.bind(this),pausePreview:function(){return this.jsonRpcCall("autoPlayPreviewStop"),this.playerInterface}.bind(this),updatePosition:function(e){return this.jsonRpcCall("updatePosition",[e]),this.playerInterface}.bind(this),updateLayoutBreakpoint:function(e){return this.jsonRpcCall("updateLayoutBreakpoint",[e]),this.playerInterface}.bind(this),enterFullScreen:function(){return this.jsonRpcCall("enterFullScreen"),this.playerInterface}.bind(this),exitFullScreen:function(){return this.jsonRpcCall("exitFullScreen"),this.playerInterface}.bind(this),teardown:function(){this.eventCallbacks={},t.removeChild(this.videoIframe),this.videoIframe=void 0,delete n[this.playerId]}.bind(this)},this.playerInterface}return{setBaseUrl:function(r){t.test(r)?e=r:window.console.error("newBaseUrl "+r+" not allowed")},createPlayerForTweet:function(e,t,i){var n="/i/videos/tweet/"+t,o="player_tweet_"+t;return new s(e,n,o,i||r)},createPlayerForDm:function(e,t,i){var n="/i/videos/dm/"+t,o="player_dm_"+t;return new s(e,n,o,i||r)},createPlayerForLiveVideo:function(e,t,i){var n="/i/videos/live_video/"+t,o="player_live_video_"+t;return new s(e,n,o,i||r)},findPlayerForElement:function(e){for(var t in n)if(n.hasOwnProperty(t)){var r=n[t];if(r&&r.domElement===e)return r.playerInterface}return null}}}()}.call(t,r,t,e))||(e.exports=i)},214:function(e,t,r){var i=r(1),n=r(183),o=r(3);e.exports=function(e,t){o.isTwitterURL(e)&&(e=n(e,t)),i.open(e)}},215:function(e,t){function r(e){this.percentage=e}r.prototype.nextBoolean=function(){return 100*Math.random()<this.percentage},e.exports=r}}]);
```
