## supplementary data file 3 for "Repeated truncation of a modular antimicrobial peptide gene for neural context": saved_resource.html

 

Twitter Ads info and privacy

### Tweets by ‎@hyphy\_software

Load new Tweets

1. HyPhy Retweeted

   Spyros Lytras
   @SpyrosLytras

   Replying to @SpyrosLytras

   We used an array of methods implemented in @hyphy\_software to search for site-, branch- and ORF-specific selection in the phylogenetic clade SARS-CoV-2 emerged from (we refer to as the 'nCoV' clade) 9/18

   Mar 13, 2021

   - ### Share on

     - Twitter
     - Facebook
     - LinkedIn
     - Tumblr
2. HyPhy Retweeted

   Sergei Pond
   @sergeilkp

   Rapid app development is a must. Excited to start moving http://Datamonkey.org  visualization development to embedded @observablehq notebooks. The new GARD analysis result page is one such example. Focus on added value, don't waste time with "plumbing" http://datamonkey.org/gard/6048d68555c66b34799c6f0c …

   Mar 10, 2021

   - ### Share on

     - Twitter
     - Facebook
     - LinkedIn
     - Tumblr
3. HyPhy Retweeted

   Alison L Hill
   @alison\_l\_hill

   Join us for the next Virus Dynamics & Evolution meeting! Bringing together modeling/computational/phylogenetic work on HIV, SARS-CoV-2, HCV, HBC, flu, dengue, CMV, Ebola, and any + all other human viruses. Abstracts due Wed March 17th. More info : https://cpd.ucsd.edu/hivdynamics/index.html … (1/3)

   Mar 9, 2021

   - ### Share on

     - Twitter
     - Facebook
     - LinkedIn
     - Tumblr
4. HyPhy Retweeted

   Sergei Pond
   @sergeilkp

   In 2020, about 10% of all the submissions to http://datamonkey.org  were CoV related. Pushed by @GISAID data volumes we significantly improved @hyphy\_software performance on larger data sets => Datamonkey users can now analyze larger datasets as well http://blog.datamonkey.org/2021/01/07/maximum-limits/ …

   Jan 17, 2021

   - ### Share on

     - Twitter
     - Facebook
     - LinkedIn
     - Tumblr
5. HyPhy Retweeted

   Sergei Pond
   @sergeilkp

   One very challenging aspect of viral evolution is that it is rarely, if ever, predictable. We can be pretty certain that changes will arise, but where, when, and what effect they may have? For example, could we have guessed what amino-acids could arise at S/501 in #SARSCoV2 ?

   Jan 16, 2021

   - ### Share on

     - Twitter
     - Facebook
     - LinkedIn
     - Tumblr
6. HyPhy Retweeted

   Sergei Pond
   @sergeilkp

   Update of #SARSCoV2 selection analyses with data up to 12/31/2020 from @GISAID at https://observablehq.com/@spond/revised-sars-cov-2-analytics-page …  
     
   There are 21 new sites with some signal for selection added since the previous analysis, including S/179, S/1117, S/1228 (https://observablehq.com/@spond/sars\_cov\_2\_sites?sites=nsp2/465,nsp2/489,nsp2/510,nsp3/338,nsp3/1269,nsp4/461,3C/71,nsp6/142,nsp6/149,exonuclease/37,exonuclease/255,endornase/112,S/179,S/1117,S/1228,N/373,N/389,ORF3a/22,ORF3a/107,ORF3a/199,ORF7a/14 …)

   ## Evolutionary annotation of B.1.1.7 SARS-CoV-2/COVID-19 genomes enabled by data from

   Evolutionary annotation of SARS-CoV-2/COVID-19 genomes enabled by data from There are codons in the SARS-CoV-2 genome with available annotation evolutionary attributes and sites selected for display....

   observablehq.com

   Jan 9, 2021

   - ### Share on

     - Twitter
     - Facebook
     - LinkedIn
     - Tumblr
7. HyPhy
   @hyphy\_software

   Effective immediately, and reflecting recent improvements in both architecture and HyPhy software improvements, http://datamonkey.org  has been updated with new maximum sequence and site limits. Please see http://blog.datamonkey.org/2021/01/07/maximum-limits/ … for details.

   Jan 8, 2021

   - ### Share on

     - Twitter
     - Facebook
     - LinkedIn
     - Tumblr
8. HyPhy Retweeted

   Sergei Pond
   @sergeilkp

   The M1 @apple chip is impressive. In my hands (@hyphy\_software, now with code to use ARM NEON) is running faster on the M1 than on the i9 Intel from the most recent Intel MB Pro, despite only having 128bit SIMD (so 2 doubles/cycle) vs 256bit AVX in Intel (4 dbl / cycle)

   Jan 3, 2021

   - ### Share on

     - Twitter
     - Facebook
     - LinkedIn
     - Tumblr
9. HyPhy Retweeted

   Sergei Pond
   @sergeilkp

   Redesigned #SARSCoV2 selection analysis up to early December sequences (>200,000) using @GISAID data can be found at https://observablehq.com/@spond/revised-sars-cov-2-analytics-page …  
     
   Recent analyses show an uptick in selection on the virus including S/501 S/484 and other sites reported from the UK and South Africa.

   ## Natural selection analysis of global SARS-CoV-2/COVID-19 enabled by data from

   Natural selection analysis of SARS-CoV-2/COVID-19 enabled by data from Analysis details Which positions in the SARS-CoV-2 genome may be subject to positive selection, i.e., involved in adaptation, or...

   observablehq.com

   Dec 22, 2020

   - ### Share on

     - Twitter
     - Facebook
     - LinkedIn
     - Tumblr
10. HyPhy Retweeted

    Sergei Pond
    @sergeilkp

    Emergence and rapid spread of a new severe acute respiratory syndrome-related coronavirus 2 (SARS-CoV-2) lineage with multiple spike mutations in South Africa https://www.krisp.org.za/publications.php?pubid=315 …   
      
    Lots of new selection signal in SARS-CoV-2 in the last 1-2 months.

    Dec 21, 2020

    - ### Share on

      - Twitter
      - Facebook
      - LinkedIn
      - Tumblr
11. HyPhy Retweeted

    Sergei Pond
    @sergeilkp

    #SARSCoV2 selection analysis update with up to ~80,000 sequences from @GISAID at http://covid19.datamonkey.org  . Two few positions in Spike (S5, S769), low frequency but consistently occurring are now in the Top 5.

    Aug 27, 2020

    - ### Share on

      - Twitter
      - Facebook
      - LinkedIn
      - Tumblr
12. HyPhy Retweeted

    Steven Weaver
    @stvnwvr

    I've started working on a comprehensive benchmarking notebook for HyPhy and started with commonly used methods. If there are any specific benchmarks you'd like to see, let me know and I'll include if reasonable. https://observablehq.com/@stevenweaver/hyphy-benchmarks-and-profiling …

    ## HyPhy Benchmarks and Profiling

    The following are benchmarks from a SARS-CoV-2 spike coding region dataset with number of sequences 250 and number of sites 3822. Commands were run using the HYPHYMPI target starting from 8 process...

    observablehq.com

    Aug 24, 2020

    - ### Share on

      - Twitter
      - Facebook
      - LinkedIn
      - Tumblr
13. HyPhy
    @hyphy\_software

    Datamonkey now been released with a new API! Submit jobs from your pipeline scripts or from anywhere you else you may find convenient. Documentation can be found here -- https://github.com/kjlevitz/datamonkey-js/wiki/API …

    Aug 10, 2020

    - ### Share on

      - Twitter
      - Facebook
      - LinkedIn
      - Tumblr
14. HyPhy Retweeted

    Sergei Pond
    @sergeilkp

    As @BallouxFrancois also pointed out, S 614 (talked about a lot) and RdRp 323 (not talked about much at all) are strongly linked (LD R2 > 0.99), and both show about the same extent of evidence for selection: both arose early on and stayed "on" https://twitter.com/OscarMacLean1/status/1289159229371068417

    Jul 31, 2020

    - ### Share on

      - Twitter
      - Facebook
      - LinkedIn
      - Tumblr
15. HyPhy Retweeted

    Oscar MacLean
    @OscarMacLean1

    In our updated manuscript we investigated and compared the viral evolutionary history which preceded and followed the emergence of #covid19 in humans. Thanks to @spyroslytas @sergeilkp @robertsonlab @stevenweaver @lemeylab @maciekboni @joshwentviral for a great collaboration

    Jul 31, 2020

    - ### Share on

      - Twitter
      - Facebook
      - LinkedIn
      - Tumblr
16. HyPhy Retweeted

    Robertson lab
    @robertson\_lab

    Update to our pre-print online with new analysis, 'Natural selection in the evolution of SARS-CoV-2 in bats, not humans, created a highly capable human pathogen'; the evidence points to the significant evolution that created #SARSCoV2 occurring in bats: https://doi.org/10.1101/2020.05.28.122366 …

    Jul 31, 2020

    - ### Share on

      - Twitter
      - Facebook
      - LinkedIn
      - Tumblr
17. HyPhy
    @hyphy\_software

    Interested in gene-wide episodic diversifying selection? Check out our latest how-to video on using BUSTED[S] on http://datamonkey.org : https://www.youtube.com/watch?v=FRcJjYIcnY8 …

    YouTube
    ‎@**YouTube**

    Jul 16, 2020

    - ### Share on

      - Twitter
      - Facebook
      - LinkedIn
      - Tumblr
18. HyPhy Retweeted

    Sergei Pond
    @sergeilkp

    @GISAID #SARS\_CoV\_2 Natural selection analysis now based on over 65,000 sequences. https://observablehq.com/@spond/natural-selection-analysis-of-sars-cov-2-covid-19 …  
    Still mostly neutral, with "original" sites, like RdRp 323 (which is in perfect linkage with S 614), topping the list.

    ## Notice: this page has been deprecated in favor of a fully redesigned and updated version 2

    Notice: this page has been deprecated in favor of a fully redesigned and updated version 2 Natural selection analysis of SARS-CoV-2/COVID-19 enabled by data from Last update: September 7th, 2020...

    observablehq.com

    Jul 15, 2020

    - ### Share on

      - Twitter
      - Facebook
      - LinkedIn
      - Tumblr
19. HyPhy Retweeted

    Sergei Pond
    @sergeilkp

    SARS-CoV-2 came from dogs? Not so much...https://academic.oup.com/mbe/article/doi/10.1093/molbev/msaa178/5870838 …

    ## Viral CpG Deficiency Provides No Evidence That Dogs Were Intermediate Hosts for SARS-CoV-2

    Abstract. Due to the scope and impact of the COVID-19 pandemic there exists a strong desire to understand where the SARS-CoV-2 virus came from and how it jumped

    academic.oup.com

    Jul 13, 2020

    - ### Share on

      - Twitter
      - Facebook
      - LinkedIn
      - Tumblr
20. HyPhy
    @hyphy\_software

    New video alert! Have you been enjoying our DataMonkey videos but want to run more than 500 sequences at once or use a method not yet implemented there? Check out our tutorial on how to install HyPhy via the line: https://youtu.be/fgNrPbOTpxE 

    YouTube
    ‎@**YouTube**

    Jun 17, 2020

    - ### Share on

      - Twitter
      - Facebook
      - LinkedIn
      - Tumblr

Load more Tweets

There are no more Tweets in this timeline.

Embed
View on Twitter
