## supplementary data file 3 for "Repeated truncation of a modular antimicrobial peptide gene for neural context": timeline.28ecda9667eeb8e1b18898b99fee6c31.js

```
(window.__twttrll=window.__twttrll||[]).push([[7],{175:function(e,t,i){var r=i(0);e.exports=function(e){return r.isType("string",e)}},182:function(e,t){var i=/^(dark|light)$/;e.exports=function(e){return i.test(e)}},189:function(e,t,i){var r=i(179),n=i(7),s=i(71),o=i(175),a=i(182),d=i(33),l=i(5),c={dataSource:{required:!0},id:{validate:o},lang:{required:!0,transform:s.matchLanguage,fallback:"en"},isPreconfigured:{required:!0,fallback:!1},width:{validate:r,transform:r},height:{validate:r,transform:r},theme:{fallback:[n(d.val,d,"widgets:theme")],validate:a},tweetLimit:{transform:l.asInt},partner:{fallback:n(d.val,d,"partner")},staticContent:{required:!1,transform:l.asBoolean},screenName:{required:!1},showReplies:{required:!1},userId:{required:!1}};e.exports=c},190:function(e){e.exports={INITIAL:1,NEWER:2,OLDER:3}},205:function(e,t,i){var r=i(190);e.exports=function(e,t,i){switch(e.cursors=e.cursors||{},e.pollInterval=t.pollInterval,i){case r.INITIAL:e.cursors.min=t.minCursorPosition,e.cursors.max=t.maxCursorPosition;break;case r.NEWER:e.cursors.max=t.maxCursorPosition||e.cursors.max;break;case r.OLDER:e.cursors.min=t.minCursorPosition||e.cursors.min}}},220:function(e,t,i){var r=i(177),n=i(176),s=i(168),o=i(18),a=i(6),d=i(189),l=i(5),c=i(205),u=i(0),h=i(73),f=i(190),m="180px",p="100%",w="200px",v="520px",g="600px",b=1;e.exports=s.couple(i(169),i(173),function(e){e.params(d),e.selectors({header:".timeline-Header",footer:".timeline-Footer",viewport:".timeline-Viewport",tweetList:".timeline-TweetList",tweetsInStream:".timeline-Tweet"}),e.around("scribeNamespace",function(e){return u.aug(e(),{page:"timeline"})}),e.around("scribeData",function(e){var t=this.params.dataSource.id;return u.aug(e(),{widget_id:l.isNumber(t)?t:void 0,widget_data_source:t,query:this.el&&this.el.getAttribute("data-search-query"),profile_id:this.el&&this.el.getAttribute("data-profile-id")})}),e.around("widgetDataAttributes",function(e){return u.aug({"widget-id":this.params.dataSource.id,"user-id":this.el&&this.el.getAttribute("data-profile-id"),"search-query":this.el&&this.el.getAttribute("data-search-query")},e())}),e.define("updateViewportHeight",function(){var e,t=this.sandbox,i=this.selectOne("header"),r=this.selectOne("footer"),n=this.selectOne("viewport");return o.read(function(){e=t.height-2*b,e-=i?i.offsetHeight:0,e-=r?r.offsetHeight:0}),o.write(function(){n.style.height=e+"px"})}),e.define("adjustWidgetSize",function(){return this.isFullyExpanded?this.sandbox.matchHeightToContent():this.updateViewportHeight()}),e.define("scribeImpressionsForInitialTweetSet",function(){var e=n(this.select("tweetsInStream")),t=Object.keys(e),i=t.length?"results":"no_results",r=this.el.getAttribute("data-collection-id");r&&(t.push(r),e[r]={item_type:h.CUSTOM_TIMELINE}),this.scribe({component:"timeline",element:"initial",action:i},{item_ids:t,item_details:e})}),e.override("initialize",function(){this.params.width||(this.params.width=this.params.isPreconfigured?v:p),this.isStaticTimeline=this.params.staticContent||this.params.tweetLimit>0,this.params.theme=this.params.theme||"light",this.isFullyExpanded=this.isStaticTimeline||!this.params.isPreconfigured&&!this.params.height,this.isFullyExpanded||this.params.height||(this.params.height=g)}),e.override("hydrate",function(){var e=this;return this.params.dataSource.fetch().then(function(t){e.html=t.html,c(e,t,f.INITIAL)})}),e.override("render",function(){var e,t=this;return this.el=this.sandbox.htmlToElement(this.html),this.el?(this.el.lang=this.params.lang,this.isFullyExpanded&&this.sandbox.addRootClass("var-fully-expanded"),this.isStaticTimeline&&this.sandbox.addRootClass("var-static"),e=r.timeline(this.params.lang,this.params.theme),a.all([this.sandbox.appendStyleSheet(e),this.sandbox.styleSelf({display:"inline-block",maxWidth:p,width:this.params.width,minWidth:m,marginTop:0,marginBottom:0})]).then(function(){return t.prepForInsertion(t.el),t.sandbox.injectWidgetEl(t.el)})):a.reject(new Error("unable to render"))}),e.override("show",function(){var e=this.sandbox,t=this;return this.sandbox.makeVisible().then(function(){return e.styleSelf({minHeight:t.isStaticTimeline?void 0:w,height:t.params.height})}).then(function(){return t.adjustWidgetSize()}).then(function(){return o.read(function(){t.scribeImpressionsForInitialTweetSet()})})}),e.last("resize",function(){return this.adjustWidgetSize()})})},221:function(e,t,i){var r=i(18),n=i(203);e.exports=function(e){e.params({chrome:{transform:n,fallback:""}}),e.selectors({streamContainer:".timeline-Viewport",tweetStream:".timeline-TweetList"}),e.before("render",function(){this.params.chrome.transparent&&this.sandbox.addRootClass("var-chromeless"),this.params.chrome.hideBorder&&this.sandbox.addRootClass("var-borderless"),this.params.chrome.hideHeader&&this.sandbox.addRootClass("var-headerless"),this.params.chrome.hideFooter&&this.sandbox.addRootClass("var-footerless")}),e.after("render",function(){if(this.params.chrome.hideScrollBar)return this.hideScrollBar()}),e.after("resize",function(){if(this.params.chrome.hideScrollBar)return this.hideScrollBar()}),e.define("hideScrollBar",function(){var e=this.selectOne("streamContainer"),t=this.selectOne("tweetStream");return r.defer(function(){var i,r;e.style.width="",i=e.offsetWidth-t.offsetWidth,r=e.offsetWidth+i,e.style.width=r+"px"})})}},222:function(e,t){e.exports=function(e){e.params({ariaLive:{fallback:""}}),e.selectors({newTweetsNotifier:".new-tweets-bar"}),e.after("render",function(){var e=this.selectOne("newTweetsNotifier");"assertive"===this.params.ariaLive&&e&&e.setAttribute("aria-live","assertive")})}},223:function(e,t,i){var r=i(74),n=i(3),s=i(5);e.exports=function(e){e.selectors({followButton:".follow-button"}),e.define("handleFollowButtonClick",function(e,t){var i=n.intentForFollowURL(t.href);s.asBoolean(t.getAttribute("data-age-gate"))||r.open(i,e)}),e.after("render",function(){this.on("click","followButton",function(e,t){this.handleFollowButtonClick(e,t)})})}},224:function(e,t,i){var r=i(34),n=i(10);e.exports=function(e){e.selectors({mediaCard:".MediaCard",mediaCardNsfwDismissalTarget:".MediaCard-dismissNsfw"}),e.define("dismissNsfwWarning",function(e,t){var i=r.closest(this.selectors.mediaCard,t,this.el);i&&n.remove(i,"is-nsfw")}),e.after("render",function(){this.on("click","mediaCardNsfwDismissalTarget",this.dismissNsfwWarning)})}},225:function(e,t,i){var r,n,s=i(10),o=i(31),a=i(168),d=i(34),l=i(71),c=i(18),u=i(11),h=i(198),f=i(6),m={autoplay:"1"},p="js-forceRedraw";e.exports=a.couple(i(226),function(e){function t(e){var t=e.createElement("div");return t.className="MediaCard-mediaAsset",t}e.params({lang:{required:!0,transform:l.matchLanguage,fallback:"en"},videoPlayerBorderRadius:{fallback:{}},videoPlayerBranding:{fallback:!0}}),e.selectors({mediaAsset:".MediaCard-mediaAsset",cardInterstitial:".js-cardPlayerInterstitial",wvpInterstitial:".js-playableMediaInterstitial",sourceIdInfo:".js-tweetIdInfo"}),e.define("videoPlayerOptions",function(){var e=(this.scribeData()||{}).widget_origin,t=this.scribeNamespace()||{};return{addBranding:this.params.videoPlayerBranding,borderRadius:this.params.videoPlayerBorderRadius,languageCode:this.params.lang,widgetOrigin:e,autoPlay:!0,scribeContext:{page:t.page,client:t.client}}}),e.define("replaceInterstitialWithMedia",function(e,t){return f.all([this.restoreLastMediaInterstitial(),c.write(function(){r=e,n=e.parentNode,e.parentNode.replaceChild(t,e)})])}),e.define("restoreLastMediaInterstitial",function(){var e;return r&&n?(e=n.firstElementChild,h.remove(e),c.write(function(){n.replaceChild(r,e)})):f.resolve()}),e.define("playWebVideoPlayerMediaAsset",function(e,t){var i,r=d.closest(this.selectors.sourceIdInfo,t,this.el),n=r.getAttribute("data-tweet-id"),s=h.insertForTweet;return n||(n=r.getAttribute("data-event-id"),s=h.insertForEvent),n?(e.preventDefault(),i=this.selectOne(r,this.selectors.wvpInterstitial),this.getConsent(r,i).then(function(){this.displayWebVideoPlayerMediaAsset(r,n,s)}.bind(this))):f.reject(new Error("No source id found for player"))}),e.define("displayWebVideoPlayerMediaAsset",function(e,i,r){var n=this.selectOne(e,this.selectors.mediaAsset),s=t(this.sandbox),o=this.sandbox.createElement("div"),a=this.videoPlayerOptions();return o.className="wvp-player-container",s.appendChild(o),this.replaceInterstitialWithMedia(n,s).then(function(){return r.call(this,s,i,a)})}),e.define("displayIframeMediaAsset",function(e,i){var r,n,a,l=d.closest(this.selectors.mediaAsset,i,this.el),h=d.closest(this.selectors.cardInterstitial,i,this.el),w=h.getAttribute("data-player-src"),v=h.getAttribute("data-player-width"),g=h.getAttribute("data-player-height"),b=h.getAttribute("data-player-title");return w?(e.preventDefault(),a=w,w=u.url(a,m),r=t(this.sandbox),(n=o({src:w,allowfullscreen:"true",width:v,height:g,title:b||""})).className="FilledIframe",r.appendChild(n),this.replaceInterstitialWithMedia(l,r).then(function(){n.focus(),c.write(function(){s.add(r,p),s.add(n,p)})})):f.reject(new Error("No Player frame source"))}),e.after("render",function(){var e=this.selectOne(this.selectors.wvpInterstitial);e&&s.remove(e,"u-hidden"),this.on("click","cardInterstitial",this.displayIframeMediaAsset),this.on("click","wvpInterstitial",this.playWebVideoPlayerMediaAsset)})})},226:function(e,t,i){var r=i(10),n=i(227),s=i(2),o=i(20),a=i(18),d=i(24),l=i(6),c=i(36);e.exports=function(e){e.selectors({cookieConsentButton:".js-cookieConsentButton",interstitial:".js-interstitial"}),e.define("getConsent",function(e,t){var i=this.selectOne(e,this.selectors.interstitial);return i?c.shouldObtainCookieConsent().catch(function(){return l.resolve(!0)}).then(function(e){var r,n;return e?(r=new s,n=function(){this.handleCookieConsentClick(t,i),r.resolve()}.bind(this),a.write(function(){this.scribe({component:"cookie_consent",action:"show"}),this.showConsentInterstitial(i,t),this.attachConsentListener(i,n)},this),r.promise):l.resolve()}.bind(this)):l.resolve()}),e.define("attachConsentListener",function(e,t){var i=this.selectOne(e,this.selectors.cookieConsentButton);i&&i.addEventListener("click",t,{once:!0})}),e.define("showConsentInterstitial",function(e,t){r.remove(e,"u-hidden"),t&&r.add(t,"is-backgrounded")}),e.define("hideConsentInterstitial",function(e,t){r.add(e,"u-hidden"),t&&r.remove(t,"is-backgrounded")}),e.define("setCookieConsentCookie",function(){return n.request(o.cookieConsent()).catch(function(e){throw new Error("CORS request failed: "+e)})}),e.define("handleCookieConsentClick",function(e,t){return d.allSettled([a.write(this.hideConsentInterstitial.bind(this,t,e)),this.setCookieConsentCookie()])})}},227:function(e,t,i){var r=i(2),n=i(6),s=i(21),o=i(25),a=i(11),d=i(0),l=i(1),c={method:"GET",params:{},headers:{},credentials:"include",isSuccess:function(e){return e>=200&&e<300}},u={JSON:"application/json",TEXT:"text/plain"},h={NO_XHR:new Error("No suitable XHR implementation available."),REQUEST_FAILED:new Error("XHR request failed."),REQUEST_ABORTED:new Error("XHR request aborted."),REQUEST_TIMED_OUT:new Error("XHR request timed out."),NETWORK_ERROR:new Error("Network error.")};e.exports={request:function(e,t){var i,f;return t=d.aug({},c,t||{}),i=a.url(e,t.params),(f=l.fetch)?f(i,t).catch(function(){return n.reject(h.NETWORK_ERROR)}).then(function(e){if(t.isSuccess(e.status))return e.text().then(function(t){var i=e.headers.get("content-type");return t&&d.contains(i,u.JSON)?o.parse(t):t});throw new Error("Request failed with status: "+e.status)}):function(e,t){var i,a=new r,c=s.ie9(),f=c?l.XDomainRequest:l.XMLHttpRequest;if(!f)return n.reject(h.NO_XHR);function m(){var e=c?i.contentType:i.getResponseHeader("content-type"),r=d.contains(e,u.JSON)?function(e){return e?o.parse(e):e}(i.responseText):i.responseText;c||t.isSuccess(i.status)?a.resolve(r):c||0!==i.status?a.reject(r):a.reject(h.NETWORK_ERROR)}return(i=new f).onreadystatechange=function(){4===i.readyState&&m()},i.onload=m,i.onerror=function(){a.reject(h.REQUEST_FAILED)},i.onabort=function(){a.reject(h.REQUEST_ABORTED)},i.ontimeout=function(){a.reject(h.REQUEST_TIMED_OUT)},i.open(t.method,e),"include"===t.credentials&&(i.withCredentials=!0),i.setRequestHeader&&d.forIn(t.headers,function(e){i.setRequestHeader(e,t.headers[e])}),i.send(),a.promise}(i,t)},mimeTypes:u,errors:h}},228:function(e,t,i){var r=i(6),n=i(168);e.exports=n.couple(i(229),function(e){e.override("resizeSandboxDueToCardChange",function(){return this.isFullyExpanded?this.sandbox.matchHeightToContent():r.resolve()})})},229:function(e,t,i){var r=i(10),n=i(18),s=i(47),o=i(21),a=i(6),d=i(0),l=i(168),c=50,u="data-card-breakpoints",h="data-theme",f="is-loaded",m="is-constrainedByMaxWidth";e.exports=l.couple(i(169),function(e){e.selectors({prerenderedCard:".PrerenderedCard",periscopeVideo:".PlayerCard--video",rootCardEl:".TwitterCard .CardContent > *:first-child"}),e.define("scribeCardShown",function(e){this.scribe({component:"card",action:"shown"},{items:[{card_name:e.getAttribute("data-card-name")}]},2)}),e.define("resizeSandboxDueToCardChange",function(){return this.sandbox.matchHeightToContent()}),e.define("markCardElAsLoaded",function(e){var t=this,i=!1;function s(){i&&t.resizeSandboxDueToCardChange()}return this.select(e,"img").forEach(function(e){e.addEventListener("load",s,!1)}),this.scribeCardShown(e),n.write(function(){r.add(e,f)}).then(function(){i=!0,t.resizeSandboxDueToCardChange()})}),e.define("updateCardWidthConstraints",function(){var e=this;return a.all(this.select("prerenderedCard").map(function(t){var i=e.selectOne(t,"rootCardEl");return n.defer(function(){var e,n=0;o.ios()?(r.remove(t,m),n=s(t.parentElement).width,t.style.maxWidth=n+"px"):n=s(t.parentElement).width,e=function(e){var t,i="";for(t=Math.floor(e/c);t>0;t--)i+="w"+t*c+" ";return i}(n),i.setAttribute(u,e),r.add(t,m)}).then(function(){return e.resizeSandboxDueToCardChange()})}))}),e.define("setCardTheme",function(e){var t=this.selectOne(e,"rootCardEl");this.params.theme&&t.setAttribute(h,this.params.theme)}),e.after("prepForInsertion",function(e){var t=this,i=this.select(e,"prerenderedCard").reduce(function(e,t){var i=t.getAttribute("data-css");return i&&(e[i]=e[i]||[],e[i].push(t)),e},{});d.forIn(i,function(e,i){t.sandbox.prependStyleSheet(e).then(function(){i.forEach(function(e){t.setCardTheme(e),t.markCardElAsLoaded(e)})})})}),e.after("show",function(){var e;return o.anyIE()&&(e=this.selectOne("periscopeVideo"))&&(e.style.display="none"),this.updateCardWidthConstraints()}),e.after("resize",function(){return this.updateCardWidthConstraints()})})},230:function(e,t,i){var r=i(2),n=i(6),s=i(10),o=i(18),a=i(1),d=i(8),l=i(0),c=i(168),u=i(176),h=i(231),f=i(205),m=i(23),p=i(232),w=i(190),v=50,g=5e3,b=500,C="is-atEndOfTimeline";e.exports=c.couple(i(169),function(e){e.params({dataSource:{required:!0},isPreviewTimeline:{required:!1,fallback:!1}}),e.selectors({timelineTweet:".timeline-Tweet",viewport:".timeline-Viewport",tweetList:".timeline-TweetList",tweetsInStream:".timeline-Tweet",newTweetsNotifier:".new-tweets-bar",loadMore:".timeline-LoadMore",loadMoreButton:".timeline-LoadMore-prompt"}),e.define("gcTweetsSync",function(){var e="custom"===this.el.getAttribute("data-timeline-type"),t=this.selectOne("tweetList");if(e)return n.resolve();h(t,v)}),e.define("scribeImpressionsForDynamicTweetSet",function(e,t){var i=l.toRealArray(e.querySelectorAll(this.selectors.timelineTweet)),r=u(i),n=Object.keys(r),s=t?"newer":"older",o=t?p.CLIENT_SIDE_APP:p.CLIENT_SIDE_USER;this.scribe({component:"timeline",element:s,action:"results"},{item_ids:n,item_details:r,event_initiator:o})}),e.define("fetchTweets",function(e,t){var i=this,r=function(e,t,i){var r={};return e=e||{},i&&e.max?r.minPosition=e.max:!i&&e.min?r.maxPosition=e.min:i?r.sinceId=t:r.maxId=t,r}(this.cursors,e,t);return this.params.dataSource.poll(r,t).then(function(r){var n,s,o=i.sandbox.createFragment(),a=i.sandbox.createElement("ol"),d=t?w.NEWER:w.OLDER;return f(i,r,d),a.innerHTML=r.html,(n=a.firstElementChild)&&(s=i.selectOne(n,"timelineTweet")),s&&"LI"===n.tagName?(s.getAttribute("data-tweet-id")===e&&a.removeChild(n),i.scribeImpressionsForDynamicTweetSet(a,t),i.prepForInsertion(a),l.toRealArray(a.children).forEach(function(e){o.appendChild(e)}),o):o},function(e){return"404"===e?i.pollInterval=null:"503"===e&&(i.pollInterval*=1.5),n.reject(e)})}),e.define("loadOldTweets",function(){var e=this,t=this.selectLast("tweetsInStream"),i=t&&t.getAttribute("data-tweet-id");return i?this.fetchTweets(i,!1).then(function(t){var i=e.selectOne("tweetList"),r=e.selectOne("loadMore");return o.write(function(){t.childNodes.length>0?i.appendChild(t):s.add(r,C)})}):n.reject(new Error("unable to load more"))}),e.after("loadOldTweets",function(){return m.trigger("timelineUpdated",{target:this.sandbox.sandboxEl,region:"older"}),this.resize()}),e.define("loadNewTweets",function(){var e=this,t=this.selectOne("tweetsInStream"),i=t&&t.getAttribute("data-tweet-id");return i?this.fetchTweets(i,!0).then(function(t){var i,r,n=e.selectOne("viewport"),s=e.selectOne("tweetList");if(0!==t.childNodes.length)return o.read(function(){i=n.scrollTop,r=n.scrollHeight}),o.defer(function(){var o;s.insertBefore(t,s.firstElementChild),o=i+n.scrollHeight-r,i>40||e.mouseIsOverWidget?(n.scrollTop=o,e.showNewTweetsNotifier()):(n.scrollTop=0,e.gcTweetsSync())})}):n.reject(new Error("unable to load new tweets"))}),e.after("loadNewTweets",function(){return m.trigger("timelineUpdated",{target:this.sandbox.sandboxEl,region:"newer"}),this.resize()}),e.define("showNewTweetsNotifier",function(){var e=this,t=this.selectOne("newTweetsNotifier"),i=t&&t.firstElementChild;return a.setTimeout(function(){e.hideNewTweetsNotifier()},g),o.write(function(){t.removeChild(i),t.appendChild(i),s.add(t,"is-displayed")}),o.defer(function(){s.add(t,"is-opaque")})}),e.define("hideNewTweetsNotifier",function(e){var t=new r,i=this.selectOne("newTweetsNotifier");return!(e=e||{}).force&&this.mouseIsOverNewTweetsNotifier?(t.resolve(),t.promise):(o.write(function(){s.remove(i,"is-opaque")}),a.setTimeout(function(){o.write(function(){s.remove(i,"is-displayed")}).then(t.resolve,t.reject)},b),t.promise)}),e.define("scrollToTopOfViewport",function(){var e=this.selectOne("viewport");return o.write(function(){e.scrollTop=0,e.focus()})}),e.define("schedulePolling",function(){var e=this,t=d.get("timeline.pollInterval");function i(){e.isPollInProgress=!1}!function r(){var n=t||e.pollInterval;n&&a.setTimeout(function(){e.isPollInProgress||(e.isPollInProgress=!0,e.loadNewTweets(e.sandbox).then(i,i)),r()},n)}()}),e.after("initialize",function(){this.isPollInProgress=!1,this.mouseIsOverWidget=!1,this.mouseIsOverNewTweetsNotifier=!1,this.cursors={},this.pollInterval=1e4}),e.after("render",function(){this.isStaticTimeline||this.params.isPreviewTimeline||(this.select("timelineTweet").length>0&&this.schedulePolling(),this.on("mouseover",function(){this.mouseIsOverWidget=!0}),this.on("mouseout",function(){this.mouseIsOverWidget=!1}),this.on("mouseover","newTweetsNotifier",function(){this.mouseIsOverNewTweetsNotifier=!0}),this.on("mouseout","newTweetsNotifier",function(){this.mouseIsOverNewTweetsNotifier=!1}),this.on("click","newTweetsNotifier",function(){this.scrollToTopOfViewport(),this.hideNewTweetsNotifier({force:!0})}),this.on("click","loadMoreButton",function(){this.loadOldTweets()}))})})},231:function(e,t){e.exports=function(e,t){if(e)for(;e.children[t];)e.removeChild(e.children[t])}},232:function(e){e.exports={CLIENT_SIDE_USER:0,CLIENT_SIDE_APP:2}},233:function(e,t,i){var r=i(33),n=i(234),s=i(7),o=".customisable-border";e.exports=function(e){e.params({borderColor:{fallback:[s(r.val,r,"widgets:border-color")],validate:n}}),e.after("render",function(){var e=this.params.borderColor;e&&this.sandbox.appendCss(function(e){return o+"{border-color:"+e+";}"}(e))})}},234:function(e,t){var i=/^#(?:[a-f\d]{3}){1,2}$/i;e.exports=function(e){return i.test(e)}},235:function(e,t,i){var r=i(34),n=i(45);e.exports=function(e){e.after("render",function(){var e,t=this.sandbox.sandboxEl,i=t.tagName;if(n(t,"td "+i))return e=r.closest("td",t),this.sandbox.styleSelf({maxWidth:e.clientWidth+"px"})})}},88:function(e,t,i){var r=i(168);e.exports=r.build([i(220),i(221),i(201),i(178),i(184),i(181),i(222),i(194),i(195),i(192),i(193),i(223),i(224),i(225),i(228),i(230),i(204),i(200),i(199),i(233),i(196),i(197),i(202),i(235)],{pageForAudienceImpression:"timeline",productName:"embeddedtimeline",breakpoints:[330,430,550,660,820,970]})}}]);
```
