## supplementary data file 3 for "Repeated truncation of a modular antimicrobial peptide gene for neural context": vendors_datamonkey.js

```
(window.webpackJsonp=window.webpackJsonp||[]).push([[1],[function(t,e,n){(function(t){t.exports=function(){"use strict";var e,r;function i(){return e.apply(null,arguments)}function a(t){return t instanceof Array||"[object Array]"===Object.prototype.toString.call(t)}function o(t){return null!=t&&"[object Object]"===Object.prototype.toString.call(t)}function u(t,e){return Object.prototype.hasOwnProperty.call(t,e)}function s(t){if(Object.getOwnPropertyNames)return 0===Object.getOwnPropertyNames(t).length;var e;for(e in t)if(u(t,e))return!1;return!0}function c(t){return void 0===t}function l(t){return"number"==typeof t||"[object Number]"===Object.prototype.toString.call(t)}function f(t){return t instanceof Date||"[object Date]"===Object.prototype.toString.call(t)}function d(t,e){var n,r=[];for(n=0;n<t.length;++n)r.push(e(t[n],n));return r}function h(t,e){for(var n in e)u(e,n)&&(t[n]=e[n]);return u(e,"toString")&&(t.toString=e.toString),u(e,"valueOf")&&(t.valueOf=e.valueOf),t}function p(t,e,n,r){return Ee(t,e,n,r,!0).utc()}function m(t){return null==t._pf&&(t._pf={empty:!1,unusedTokens:[],unusedInput:[],overflow:-2,charsLeftOver:0,nullInput:!1,invalidEra:null,invalidMonth:null,invalidFormat:!1,userInvalidated:!1,iso:!1,parsedDateParts:[],era:null,meridiem:null,rfc2822:!1,weekdayMismatch:!1}),t._pf}function v(t){if(null==t._isValid){var e=m(t),n=r.call(e.parsedDateParts,(function(t){return null!=t})),i=!isNaN(t._d.getTime())&&e.overflow<0&&!e.empty&&!e.invalidEra&&!e.invalidMonth&&!e.invalidWeekday&&!e.weekdayMismatch&&!e.nullInput&&!e.invalidFormat&&!e.userInvalidated&&(!e.meridiem||e.meridiem&&n);if(t._strict&&(i=i&&0===e.charsLeftOver&&0===e.unusedTokens.length&&void 0===e.bigHour),null!=Object.isFrozen&&Object.isFrozen(t))return i;t._isValid=i}return t._isValid}function g(t){var e=p(NaN);return null!=t?h(m(e),t):m(e).userInvalidated=!0,e}r=Array.prototype.some?Array.prototype.some:function(t){var e,n=Object(this),r=n.length>>>0;for(e=0;e<r;e++)if(e in n&&t.call(this,n[e],e,n))return!0;return!1};var _=i.momentProperties=[],y=!1;function b(t,e){var n,r,i;if(c(e._isAMomentObject)||(t._isAMomentObject=e._isAMomentObject),c(e._i)||(t._i=e._i),c(e._f)||(t._f=e._f),c(e._l)||(t._l=e._l),c(e._strict)||(t._strict=e._strict),c(e._tzm)||(t._tzm=e._tzm),c(e._isUTC)||(t._isUTC=e._isUTC),c(e._offset)||(t._offset=e._offset),c(e._pf)||(t._pf=m(e)),c(e._locale)||(t._locale=e._locale),_.length>0)for(n=0;n<_.length;n++)c(i=e[r=_[n]])||(t[r]=i);return t}function w(t){b(this,t),this._d=new Date(null!=t._d?t._d.getTime():NaN),this.isValid()||(this._d=new Date(NaN)),!1===y&&(y=!0,i.updateOffset(this),y=!1)}function x(t){return t instanceof w||null!=t&&null!=t._isAMomentObject}function k(t){!1===i.suppressDeprecationWarnings&&"undefined"!=typeof console&&console.warn&&console.warn("Deprecation warning: "+t)}function M(t,e){var n=!0;return h((function(){if(null!=i.deprecationHandler&&i.deprecationHandler(null,t),n){var r,a,o,s=[];for(a=0;a<arguments.length;a++){if(r="","object"==typeof arguments[a]){for(o in r+="\n["+a+"] ",arguments[0])u(arguments[0],o)&&(r+=o+": "+arguments[0][o]+", ");r=r.slice(0,-2)}else r=arguments[a];s.push(r)}k(t+"\nArguments: "+Array.prototype.slice.call(s).join("")+"\n"+(new Error).stack),n=!1}return e.apply(this,arguments)}),e)}var E,S={};function T(t,e){null!=i.deprecationHandler&&i.deprecationHandler(t,e),S[t]||(k(e),S[t]=!0)}function j(t){return"undefined"!=typeof Function&&t instanceof Function||"[object Function]"===Object.prototype.toString.call(t)}function O(t,e){var n,r=h({},t);for(n in e)u(e,n)&&(o(t[n])&&o(e[n])?(r[n]={},h(r[n],t[n]),h(r[n],e[n])):null!=e[n]?r[n]=e[n]:delete r[n]);for(n in t)u(t,n)&&!u(e,n)&&o(t[n])&&(r[n]=h({},r[n]));return r}function A(t){null!=t&&this.set(t)}function L(t,e,n){var r=""+Math.abs(t),i=e-r.length;return(t>=0?n?"+":"":"-")+Math.pow(10,Math.max(0,i)).toString().substr(1)+r}i.suppressDeprecationWarnings=!1,i.deprecationHandler=null,E=Object.keys?Object.keys:function(t){var e,n=[];for(e in t)u(t,e)&&n.push(e);return n};var C=/(\[[^\[]*\])|(\\)?([Hh]mm(ss)?|Mo|MM?M?M?|Do|DDDo|DD?D?D?|ddd?d?|do?|w[o|w]?|W[o|W]?|Qo?|N{1,5}|YYYYYY|YYYYY|YYYY|YY|y{2,4}|yo?|gg(ggg?)?|GG(GGG?)?|e|E|a|A|hh?|HH?|kk?|mm?|ss?|S{1,9}|x|X|zz?|ZZ?|.)/g,D=/(\[[^\[]*\])|(\\)?(LTS|LT|LL?L?L?|l{1,4})/g,N={},P={};function R(t,e,n,r){var i=r;"string"==typeof r&&(i=function(){return this[r]()}),t&&(P[t]=i),e&&(P[e[0]]=function(){return L(i.apply(this,arguments),e[1],e[2])}),n&&(P[n]=function(){return this.localeData().ordinal(i.apply(this,arguments),t)})}function Y(t,e){return t.isValid()?(e=I(e,t.localeData()),N[e]=N[e]||function(t){var e,n,r,i=t.match(C);for(e=0,n=i.length;e<n;e++)P[i[e]]?i[e]=P[i[e]]:i[e]=(r=i[e]).match(/\[[\s\S]/)?r.replace(/^\[|\]$/g,""):r.replace(/\\/g,"");return function(e){var r,a="";for(r=0;r<n;r++)a+=j(i[r])?i[r].call(e,t):i[r];return a}}(e),N[e](t)):t.localeData().invalidDate()}function I(t,e){var n=5;function r(t){return e.longDateFormat(t)||t}for(D.lastIndex=0;n>=0&&D.test(t);)t=t.replace(D,r),D.lastIndex=0,n-=1;return t}var z={};function F(t,e){var n=t.toLowerCase();z[n]=z[n+"s"]=z[e]=t}function H(t){return"string"==typeof t?z[t]||z[t.toLowerCase()]:void 0}function B(t){var e,n,r={};for(n in t)u(t,n)&&(e=H(n))&&(r[e]=t[n]);return r}var q={};function U(t,e){q[t]=e}function W(t){return t%4==0&&t%100!=0||t%400==0}function G(t){return t<0?Math.ceil(t)||0:Math.floor(t)}function V(t){var e=+t,n=0;return 0!==e&&isFinite(e)&&(n=G(e)),n}function $(t,e){return function(n){return null!=n?(J(this,t,n),i.updateOffset(this,e),this):X(this,t)}}function X(t,e){return t.isValid()?t._d["get"+(t._isUTC?"UTC":"")+e]():NaN}function J(t,e,n){t.isValid()&&!isNaN(n)&&("FullYear"===e&&W(t.year())&&1===t.month()&&29===t.date()?(n=V(n),t._d["set"+(t._isUTC?"UTC":"")+e](n,t.month(),xt(n,t.month()))):t._d["set"+(t._isUTC?"UTC":"")+e](n))}var Z,K=/\d/,Q=/\d\d/,tt=/\d{3}/,et=/\d{4}/,nt=/[+-]?\d{6}/,rt=/\d\d?/,it=/\d\d\d\d?/,at=/\d\d\d\d\d\d?/,ot=/\d{1,3}/,ut=/\d{1,4}/,st=/[+-]?\d{1,6}/,ct=/\d+/,lt=/[+-]?\d+/,ft=/Z|[+-]\d\d:?\d\d/gi,dt=/Z|[+-]\d\d(?::?\d\d)?/gi,ht=/[0-9]{0,256}['a-z\u00A0-\u05FF\u0700-\uD7FF\uF900-\uFDCF\uFDF0-\uFF07\uFF10-\uFFEF]{1,256}|[\u0600-\u06FF\/]{1,256}(\s*?[\u0600-\u06FF]{1,256}){1,2}/i;function pt(t,e,n){Z[t]=j(e)?e:function(t,r){return t&&n?n:e}}function mt(t,e){return u(Z,t)?Z[t](e._strict,e._locale):new RegExp(vt(t.replace("\\","").replace(/\\(\[)|\\(\])|\[([^\]\[]*)\]|\\(.)/g,(function(t,e,n,r,i){return e||n||r||i}))))}function vt(t){return t.replace(/[-\/\\^$*+?.()|[\]{}]/g,"\\$&")}Z={};var gt,_t={};function yt(t,e){var n,r=e;for("string"==typeof t&&(t=[t]),l(e)&&(r=function(t,n){n[e]=V(t)}),n=0;n<t.length;n++)_t[t[n]]=r}function bt(t,e){yt(t,(function(t,n,r,i){r._w=r._w||{},e(t,r._w,r,i)}))}function wt(t,e,n){null!=e&&u(_t,t)&&_t[t](e,n._a,n,t)}function xt(t,e){if(isNaN(t)||isNaN(e))return NaN;var n,r=(e%(n=12)+n)%n;return t+=(e-r)/12,1===r?W(t)?29:28:31-r%7%2}gt=Array.prototype.indexOf?Array.prototype.indexOf:function(t){var e;for(e=0;e<this.length;++e)if(this[e]===t)return e;return-1},R("M",["MM",2],"Mo",(function(){return this.month()+1})),R("MMM",0,0,(function(t){return this.localeData().monthsShort(this,t)})),R("MMMM",0,0,(function(t){return this.localeData().months(this,t)})),F("month","M"),U("month",8),pt("M",rt),pt("MM",rt,Q),pt("MMM",(function(t,e){return e.monthsShortRegex(t)})),pt("MMMM",(function(t,e){return e.monthsRegex(t)})),yt(["M","MM"],(function(t,e){e[1]=V(t)-1})),yt(["MMM","MMMM"],(function(t,e,n,r){var i=n._locale.monthsParse(t,r,n._strict);null!=i?e[1]=i:m(n).invalidMonth=t}));var kt="January_February_March_April_May_June_July_August_September_October_November_December".split("_"),Mt="Jan_Feb_Mar_Apr_May_Jun_Jul_Aug_Sep_Oct_Nov_Dec".split("_"),Et=/D[oD]?(\[[^\[\]]*\]|\s)+MMMM?/,St=ht,Tt=ht;function jt(t,e,n){var r,i,a,o=t.toLocaleLowerCase();if(!this._monthsParse)for(this._monthsParse=[],this._longMonthsParse=[],this._shortMonthsParse=[],r=0;r<12;++r)a=p([2e3,r]),this._shortMonthsParse[r]=this.monthsShort(a,"").toLocaleLowerCase(),this._longMonthsParse[r]=this.months(a,"").toLocaleLowerCase();return n?"MMM"===e?-1!==(i=gt.call(this._shortMonthsParse,o))?i:null:-1!==(i=gt.call(this._longMonthsParse,o))?i:null:"MMM"===e?-1!==(i=gt.call(this._shortMonthsParse,o))||-1!==(i=gt.call(this._longMonthsParse,o))?i:null:-1!==(i=gt.call(this._longMonthsParse,o))||-1!==(i=gt.call(this._shortMonthsParse,o))?i:null}function Ot(t,e){var n;if(!t.isValid())return t;if("string"==typeof e)if(/^\d+$/.test(e))e=V(e);else if(!l(e=t.localeData().monthsParse(e)))return t;return n=Math.min(t.date(),xt(t.year(),e)),t._d["set"+(t._isUTC?"UTC":"")+"Month"](e,n),t}function At(t){return null!=t?(Ot(this,t),i.updateOffset(this,!0),this):X(this,"Month")}function Lt(){function t(t,e){return e.length-t.length}var e,n,r=[],i=[],a=[];for(e=0;e<12;e++)n=p([2e3,e]),r.push(this.monthsShort(n,"")),i.push(this.months(n,"")),a.push(this.months(n,"")),a.push(this.monthsShort(n,""));for(r.sort(t),i.sort(t),a.sort(t),e=0;e<12;e++)r[e]=vt(r[e]),i[e]=vt(i[e]);for(e=0;e<24;e++)a[e]=vt(a[e]);this._monthsRegex=new RegExp("^("+a.join("|")+")","i"),this._monthsShortRegex=this._monthsRegex,this._monthsStrictRegex=new RegExp("^("+i.join("|")+")","i"),this._monthsShortStrictRegex=new RegExp("^("+r.join("|")+")","i")}function Ct(t){return W(t)?366:365}R("Y",0,0,(function(){var t=this.year();return t<=9999?L(t,4):"+"+t})),R(0,["YY",2],0,(function(){return this.year()%100})),R(0,["YYYY",4],0,"year"),R(0,["YYYYY",5],0,"year"),R(0,["YYYYYY",6,!0],0,"year"),F("year","y"),U("year",1),pt("Y",lt),pt("YY",rt,Q),pt("YYYY",ut,et),pt("YYYYY",st,nt),pt("YYYYYY",st,nt),yt(["YYYYY","YYYYYY"],0),yt("YYYY",(function(t,e){e[0]=2===t.length?i.parseTwoDigitYear(t):V(t)})),yt("YY",(function(t,e){e[0]=i.parseTwoDigitYear(t)})),yt("Y",(function(t,e){e[0]=parseInt(t,10)})),i.parseTwoDigitYear=function(t){return V(t)+(V(t)>68?1900:2e3)};var Dt=$("FullYear",!0);function Nt(t,e,n,r,i,a,o){var u;return t<100&&t>=0?(u=new Date(t+400,e,n,r,i,a,o),isFinite(u.getFullYear())&&u.setFullYear(t)):u=new Date(t,e,n,r,i,a,o),u}function Pt(t){var e,n;return t<100&&t>=0?((n=Array.prototype.slice.call(arguments))[0]=t+400,e=new Date(Date.UTC.apply(null,n)),isFinite(e.getUTCFullYear())&&e.setUTCFullYear(t)):e=new Date(Date.UTC.apply(null,arguments)),e}function Rt(t,e,n){var r=7+e-n;return-(7+Pt(t,0,r).getUTCDay()-e)%7+r-1}function Yt(t,e,n,r,i){var a,o,u=1+7*(e-1)+(7+n-r)%7+Rt(t,r,i);return u<=0?o=Ct(a=t-1)+u:u>Ct(t)?(a=t+1,o=u-Ct(t)):(a=t,o=u),{year:a,dayOfYear:o}}function It(t,e,n){var r,i,a=Rt(t.year(),e,n),o=Math.floor((t.dayOfYear()-a-1)/7)+1;return o<1?r=o+zt(i=t.year()-1,e,n):o>zt(t.year(),e,n)?(r=o-zt(t.year(),e,n),i=t.year()+1):(i=t.year(),r=o),{week:r,year:i}}function zt(t,e,n){var r=Rt(t,e,n),i=Rt(t+1,e,n);return(Ct(t)-r+i)/7}function Ft(t,e){return t.slice(e,7).concat(t.slice(0,e))}R("w",["ww",2],"wo","week"),R("W",["WW",2],"Wo","isoWeek"),F("week","w"),F("isoWeek","W"),U("week",5),U("isoWeek",5),pt("w",rt),pt("ww",rt,Q),pt("W",rt),pt("WW",rt,Q),bt(["w","ww","W","WW"],(function(t,e,n,r){e[r.substr(0,1)]=V(t)})),R("d",0,"do","day"),R("dd",0,0,(function(t){return this.localeData().weekdaysMin(this,t)})),R("ddd",0,0,(function(t){return this.localeData().weekdaysShort(this,t)})),R("dddd",0,0,(function(t){return this.localeData().weekdays(this,t)})),R("e",0,0,"weekday"),R("E",0,0,"isoWeekday"),F("day","d"),F("weekday","e"),F("isoWeekday","E"),U("day",11),U("weekday",11),U("isoWeekday",11),pt("d",rt),pt("e",rt),pt("E",rt),pt("dd",(function(t,e){return e.weekdaysMinRegex(t)})),pt("ddd",(function(t,e){return e.weekdaysShortRegex(t)})),pt("dddd",(function(t,e){return e.weekdaysRegex(t)})),bt(["dd","ddd","dddd"],(function(t,e,n,r){var i=n._locale.weekdaysParse(t,r,n._strict);null!=i?e.d=i:m(n).invalidWeekday=t})),bt(["d","e","E"],(function(t,e,n,r){e[r]=V(t)}));var Ht="Sunday_Monday_Tuesday_Wednesday_Thursday_Friday_Saturday".split("_"),Bt="Sun_Mon_Tue_Wed_Thu_Fri_Sat".split("_"),qt="Su_Mo_Tu_We_Th_Fr_Sa".split("_"),Ut=ht,Wt=ht,Gt=ht;function Vt(t,e,n){var r,i,a,o=t.toLocaleLowerCase();if(!this._weekdaysParse)for(this._weekdaysParse=[],this._shortWeekdaysParse=[],this._minWeekdaysParse=[],r=0;r<7;++r)a=p([2e3,1]).day(r),this._minWeekdaysParse[r]=this.weekdaysMin(a,"").toLocaleLowerCase(),this._shortWeekdaysParse[r]=this.weekdaysShort(a,"").toLocaleLowerCase(),this._weekdaysParse[r]=this.weekdays(a,"").toLocaleLowerCase();return n?"dddd"===e?-1!==(i=gt.call(this._weekdaysParse,o))?i:null:"ddd"===e?-1!==(i=gt.call(this._shortWeekdaysParse,o))?i:null:-1!==(i=gt.call(this._minWeekdaysParse,o))?i:null:"dddd"===e?-1!==(i=gt.call(this._weekdaysParse,o))||-1!==(i=gt.call(this._shortWeekdaysParse,o))||-1!==(i=gt.call(this._minWeekdaysParse,o))?i:null:"ddd"===e?-1!==(i=gt.call(this._shortWeekdaysParse,o))||-1!==(i=gt.call(this._weekdaysParse,o))||-1!==(i=gt.call(this._minWeekdaysParse,o))?i:null:-1!==(i=gt.call(this._minWeekdaysParse,o))||-1!==(i=gt.call(this._weekdaysParse,o))||-1!==(i=gt.call(this._shortWeekdaysParse,o))?i:null}function $t(){function t(t,e){return e.length-t.length}var e,n,r,i,a,o=[],u=[],s=[],c=[];for(e=0;e<7;e++)n=p([2e3,1]).day(e),r=vt(this.weekdaysMin(n,"")),i=vt(this.weekdaysShort(n,"")),a=vt(this.weekdays(n,"")),o.push(r),u.push(i),s.push(a),c.push(r),c.push(i),c.push(a);o.sort(t),u.sort(t),s.sort(t),c.sort(t),this._weekdaysRegex=new RegExp("^("+c.join("|")+")","i"),this._weekdaysShortRegex=this._weekdaysRegex,this._weekdaysMinRegex=this._weekdaysRegex,this._weekdaysStrictRegex=new RegExp("^("+s.join("|")+")","i"),this._weekdaysShortStrictRegex=new RegExp("^("+u.join("|")+")","i"),this._weekdaysMinStrictRegex=new RegExp("^("+o.join("|")+")","i")}function Xt(){return this.hours()%12||12}function Jt(t,e){R(t,0,0,(function(){return this.localeData().meridiem(this.hours(),this.minutes(),e)}))}function Zt(t,e){return e._meridiemParse}R("H",["HH",2],0,"hour"),R("h",["hh",2],0,Xt),R("k",["kk",2],0,(function(){return this.hours()||24})),R("hmm",0,0,(function(){return""+Xt.apply(this)+L(this.minutes(),2)})),R("hmmss",0,0,(function(){return""+Xt.apply(this)+L(this.minutes(),2)+L(this.seconds(),2)})),R("Hmm",0,0,(function(){return""+this.hours()+L(this.minutes(),2)})),R("Hmmss",0,0,(function(){return""+this.hours()+L(this.minutes(),2)+L(this.seconds(),2)})),Jt("a",!0),Jt("A",!1),F("hour","h"),U("hour",13),pt("a",Zt),pt("A",Zt),pt("H",rt),pt("h",rt),pt("k",rt),pt("HH",rt,Q),pt("hh",rt,Q),pt("kk",rt,Q),pt("hmm",it),pt("hmmss",at),pt("Hmm",it),pt("Hmmss",at),yt(["H","HH"],3),yt(["k","kk"],(function(t,e,n){var r=V(t);e[3]=24===r?0:r})),yt(["a","A"],(function(t,e,n){n._isPm=n._locale.isPM(t),n._meridiem=t})),yt(["h","hh"],(function(t,e,n){e[3]=V(t),m(n).bigHour=!0})),yt("hmm",(function(t,e,n){var r=t.length-2;e[3]=V(t.substr(0,r)),e[4]=V(t.substr(r)),m(n).bigHour=!0})),yt("hmmss",(function(t,e,n){var r=t.length-4,i=t.length-2;e[3]=V(t.substr(0,r)),e[4]=V(t.substr(r,2)),e[5]=V(t.substr(i)),m(n).bigHour=!0})),yt("Hmm",(function(t,e,n){var r=t.length-2;e[3]=V(t.substr(0,r)),e[4]=V(t.substr(r))})),yt("Hmmss",(function(t,e,n){var r=t.length-4,i=t.length-2;e[3]=V(t.substr(0,r)),e[4]=V(t.substr(r,2)),e[5]=V(t.substr(i))}));var Kt,Qt=$("Hours",!0),te={calendar:{sameDay:"[Today at] LT",nextDay:"[Tomorrow at] LT",nextWeek:"dddd [at] LT",lastDay:"[Yesterday at] LT",lastWeek:"[Last] dddd [at] LT",sameElse:"L"},longDateFormat:{LTS:"h:mm:ss A",LT:"h:mm A",L:"MM/DD/YYYY",LL:"MMMM D, YYYY",LLL:"MMMM D, YYYY h:mm A",LLLL:"dddd, MMMM D, YYYY h:mm A"},invalidDate:"Invalid date",ordinal:"%d",dayOfMonthOrdinalParse:/\d{1,2}/,relativeTime:{future:"in %s",past:"%s ago",s:"a few seconds",ss:"%d seconds",m:"a minute",mm:"%d minutes",h:"an hour",hh:"%d hours",d:"a day",dd:"%d days",w:"a week",ww:"%d weeks",M:"a month",MM:"%d months",y:"a year",yy:"%d years"},months:kt,monthsShort:Mt,week:{dow:0,doy:6},weekdays:Ht,weekdaysMin:qt,weekdaysShort:Bt,meridiemParse:/[ap]\.?m?\.?/i},ee={},ne={};function re(t,e){var n,r=Math.min(t.length,e.length);for(n=0;n<r;n+=1)if(t[n]!==e[n])return n;return r}function ie(t){return t?t.toLowerCase().replace("_","-"):t}function ae(e){var r=null;if(void 0===ee[e]&&void 0!==t&&t&&t.exports)try{r=Kt._abbr,n(717)("./"+e),oe(r)}catch(t){ee[e]=null}return ee[e]}function oe(t,e){var n;return t&&((n=c(e)?se(t):ue(t,e))?Kt=n:"undefined"!=typeof console&&console.warn&&console.warn("Locale "+t+" not found. Did you forget to load it?")),Kt._abbr}function ue(t,e){if(null!==e){var n,r=te;if(e.abbr=t,null!=ee[t])T("defineLocaleOverride","use moment.updateLocale(localeName, config) to change an existing locale. moment.defineLocale(localeName, config) should only be used for creating a new locale See http://momentjs.com/guides/#/warnings/define-locale/ for more info."),r=ee[t]._config;else if(null!=e.parentLocale)if(null!=ee[e.parentLocale])r=ee[e.parentLocale]._config;else{if(null==(n=ae(e.parentLocale)))return ne[e.parentLocale]||(ne[e.parentLocale]=[]),ne[e.parentLocale].push({name:t,config:e}),null;r=n._config}return ee[t]=new A(O(r,e)),ne[t]&&ne[t].forEach((function(t){ue(t.name,t.config)})),oe(t),ee[t]}return delete ee[t],null}function se(t){var e;if(t&&t._locale&&t._locale._abbr&&(t=t._locale._abbr),!t)return Kt;if(!a(t)){if(e=ae(t))return e;t=[t]}return function(t){for(var e,n,r,i,a=0;a<t.length;){for(e=(i=ie(t[a]).split("-")).length,n=(n=ie(t[a+1]))?n.split("-"):null;e>0;){if(r=ae(i.slice(0,e).join("-")))return r;if(n&&n.length>=e&&re(i,n)>=e-1)break;e--}a++}return Kt}(t)}function ce(t){var e,n=t._a;return n&&-2===m(t).overflow&&(e=n[1]<0||n[1]>11?1:n[2]<1||n[2]>xt(n[0],n[1])?2:n[3]<0||n[3]>24||24===n[3]&&(0!==n[4]||0!==n[5]||0!==n[6])?3:n[4]<0||n[4]>59?4:n[5]<0||n[5]>59?5:n[6]<0||n[6]>999?6:-1,m(t)._overflowDayOfYear&&(e<0||e>2)&&(e=2),m(t)._overflowWeeks&&-1===e&&(e=7),m(t)._overflowWeekday&&-1===e&&(e=8),m(t).overflow=e),t}var le=/^\s*((?:[+-]\d{6}|\d{4})-(?:\d\d-\d\d|W\d\d-\d|W\d\d|\d\d\d|\d\d))(?:(T| )(\d\d(?::\d\d(?::\d\d(?:[.,]\d+)?)?)?)([+-]\d\d(?::?\d\d)?|\s*Z)?)?$/,fe=/^\s*((?:[+-]\d{6}|\d{4})(?:\d\d\d\d|W\d\d\d|W\d\d|\d\d\d|\d\d|))(?:(T| )(\d\d(?:\d\d(?:\d\d(?:[.,]\d+)?)?)?)([+-]\d\d(?::?\d\d)?|\s*Z)?)?$/,de=/Z|[+-]\d\d(?::?\d\d)?/,he=[["YYYYYY-MM-DD",/[+-]\d{6}-\d\d-\d\d/],["YYYY-MM-DD",/\d{4}-\d\d-\d\d/],["GGGG-[W]WW-E",/\d{4}-W\d\d-\d/],["GGGG-[W]WW",/\d{4}-W\d\d/,!1],["YYYY-DDD",/\d{4}-\d{3}/],["YYYY-MM",/\d{4}-\d\d/,!1],["YYYYYYMMDD",/[+-]\d{10}/],["YYYYMMDD",/\d{8}/],["GGGG[W]WWE",/\d{4}W\d{3}/],["GGGG[W]WW",/\d{4}W\d{2}/,!1],["YYYYDDD",/\d{7}/],["YYYYMM",/\d{6}/,!1],["YYYY",/\d{4}/,!1]],pe=[["HH:mm:ss.SSSS",/\d\d:\d\d:\d\d\.\d+/],["HH:mm:ss,SSSS",/\d\d:\d\d:\d\d,\d+/],["HH:mm:ss",/\d\d:\d\d:\d\d/],["HH:mm",/\d\d:\d\d/],["HHmmss.SSSS",/\d\d\d\d\d\d\.\d+/],["HHmmss,SSSS",/\d\d\d\d\d\d,\d+/],["HHmmss",/\d\d\d\d\d\d/],["HHmm",/\d\d\d\d/],["HH",/\d\d/]],me=/^\/?Date\((-?\d+)/i,ve=/^(?:(Mon|Tue|Wed|Thu|Fri|Sat|Sun),?\s)?(\d{1,2})\s(Jan|Feb|Mar|Apr|May|Jun|Jul|Aug|Sep|Oct|Nov|Dec)\s(\d{2,4})\s(\d\d):(\d\d)(?::(\d\d))?\s(?:(UT|GMT|[ECMP][SD]T)|([Zz])|([+-]\d{4}))$/,ge={UT:0,GMT:0,EDT:-240,EST:-300,CDT:-300,CST:-360,MDT:-360,MST:-420,PDT:-420,PST:-480};function _e(t){var e,n,r,i,a,o,u=t._i,s=le.exec(u)||fe.exec(u);if(s){for(m(t).iso=!0,e=0,n=he.length;e<n;e++)if(he[e][1].exec(s[1])){i=he[e][0],r=!1!==he[e][2];break}if(null==i)return void(t._isValid=!1);if(s[3]){for(e=0,n=pe.length;e<n;e++)if(pe[e][1].exec(s[3])){a=(s[2]||" ")+pe[e][0];break}if(null==a)return void(t._isValid=!1)}if(!r&&null!=a)return void(t._isValid=!1);if(s[4]){if(!de.exec(s[4]))return void(t._isValid=!1);o="Z"}t._f=i+(a||"")+(o||""),ke(t)}else t._isValid=!1}function ye(t){var e=parseInt(t,10);return e<=49?2e3+e:e<=999?1900+e:e}function be(t){var e,n,r,i,a,o,u,s,c=ve.exec(t._i.replace(/\([^)]*\)|[\n\t]/g," ").replace(/(\s\s+)/g," ").replace(/^\s\s*/,"").replace(/\s\s*$/,""));if(c){if(n=c[4],r=c[3],i=c[2],a=c[5],o=c[6],u=c[7],s=[ye(n),Mt.indexOf(r),parseInt(i,10),parseInt(a,10),parseInt(o,10)],u&&s.push(parseInt(u,10)),e=s,!function(t,e,n){return!t||Bt.indexOf(t)===new Date(e[0],e[1],e[2]).getDay()||(m(n).weekdayMismatch=!0,n._isValid=!1,!1)}(c[1],e,t))return;t._a=e,t._tzm=function(t,e,n){if(t)return ge[t];if(e)return 0;var r=parseInt(n,10),i=r%100;return(r-i)/100*60+i}(c[8],c[9],c[10]),t._d=Pt.apply(null,t._a),t._d.setUTCMinutes(t._d.getUTCMinutes()-t._tzm),m(t).rfc2822=!0}else t._isValid=!1}function we(t,e,n){return null!=t?t:null!=e?e:n}function xe(t){var e,n,r,a,o,u=[];if(!t._d){for(r=function(t){var e=new Date(i.now());return t._useUTC?[e.getUTCFullYear(),e.getUTCMonth(),e.getUTCDate()]:[e.getFullYear(),e.getMonth(),e.getDate()]}(t),t._w&&null==t._a[2]&&null==t._a[1]&&function(t){var e,n,r,i,a,o,u,s,c;null!=(e=t._w).GG||null!=e.W||null!=e.E?(a=1,o=4,n=we(e.GG,t._a[0],It(Se(),1,4).year),r=we(e.W,1),((i=we(e.E,1))<1||i>7)&&(s=!0)):(a=t._locale._week.dow,o=t._locale._week.doy,c=It(Se(),a,o),n=we(e.gg,t._a[0],c.year),r=we(e.w,c.week),null!=e.d?((i=e.d)<0||i>6)&&(s=!0):null!=e.e?(i=e.e+a,(e.e<0||e.e>6)&&(s=!0)):i=a),r<1||r>zt(n,a,o)?m(t)._overflowWeeks=!0:null!=s?m(t)._overflowWeekday=!0:(u=Yt(n,r,i,a,o),t._a[0]=u.year,t._dayOfYear=u.dayOfYear)}(t),null!=t._dayOfYear&&(o=we(t._a[0],r[0]),(t._dayOfYear>Ct(o)||0===t._dayOfYear)&&(m(t)._overflowDayOfYear=!0),n=Pt(o,0,t._dayOfYear),t._a[1]=n.getUTCMonth(),t._a[2]=n.getUTCDate()),e=0;e<3&&null==t._a[e];++e)t._a[e]=u[e]=r[e];for(;e<7;e++)t._a[e]=u[e]=null==t._a[e]?2===e?1:0:t._a[e];24===t._a[3]&&0===t._a[4]&&0===t._a[5]&&0===t._a[6]&&(t._nextDay=!0,t._a[3]=0),t._d=(t._useUTC?Pt:Nt).apply(null,u),a=t._useUTC?t._d.getUTCDay():t._d.getDay(),null!=t._tzm&&t._d.setUTCMinutes(t._d.getUTCMinutes()-t._tzm),t._nextDay&&(t._a[3]=24),t._w&&void 0!==t._w.d&&t._w.d!==a&&(m(t).weekdayMismatch=!0)}}function ke(t){if(t._f!==i.ISO_8601)if(t._f!==i.RFC_2822){t._a=[],m(t).empty=!0;var e,n,r,a,o,u,s=""+t._i,c=s.length,l=0;for(r=I(t._f,t._locale).match(C)||[],e=0;e<r.length;e++)a=r[e],(n=(s.match(mt(a,t))||[])[0])&&((o=s.substr(0,s.indexOf(n))).length>0&&m(t).unusedInput.push(o),s=s.slice(s.indexOf(n)+n.length),l+=n.length),P[a]?(n?m(t).empty=!1:m(t).unusedTokens.push(a),wt(a,n,t)):t._strict&&!n&&m(t).unusedTokens.push(a);m(t).charsLeftOver=c-l,s.length>0&&m(t).unusedInput.push(s),t._a[3]<=12&&!0===m(t).bigHour&&t._a[3]>0&&(m(t).bigHour=void 0),m(t).parsedDateParts=t._a.slice(0),m(t).meridiem=t._meridiem,t._a[3]=function(t,e,n){var r;return null==n?e:null!=t.meridiemHour?t.meridiemHour(e,n):null!=t.isPM?((r=t.isPM(n))&&e<12&&(e+=12),r||12!==e||(e=0),e):e}(t._locale,t._a[3],t._meridiem),null!==(u=m(t).era)&&(t._a[0]=t._locale.erasConvertYear(u,t._a[0])),xe(t),ce(t)}else be(t);else _e(t)}function Me(t){var e=t._i,n=t._f;return t._locale=t._locale||se(t._l),null===e||void 0===n&&""===e?g({nullInput:!0}):("string"==typeof e&&(t._i=e=t._locale.preparse(e)),x(e)?new w(ce(e)):(f(e)?t._d=e:a(n)?function(t){var e,n,r,i,a,o,u=!1;if(0===t._f.length)return m(t).invalidFormat=!0,void(t._d=new Date(NaN));for(i=0;i<t._f.length;i++)a=0,o=!1,e=b({},t),null!=t._useUTC&&(e._useUTC=t._useUTC),e._f=t._f[i],ke(e),v(e)&&(o=!0),a+=m(e).charsLeftOver,a+=10*m(e).unusedTokens.length,m(e).score=a,u?a<r&&(r=a,n=e):(null==r||a<r||o)&&(r=a,n=e,o&&(u=!0));h(t,n||e)}(t):n?ke(t):function(t){var e=t._i;c(e)?t._d=new Date(i.now()):f(e)?t._d=new Date(e.valueOf()):"string"==typeof e?function(t){var e=me.exec(t._i);null===e?(_e(t),!1===t._isValid&&(delete t._isValid,be(t),!1===t._isValid&&(delete t._isValid,t._strict?t._isValid=!1:i.createFromInputFallback(t)))):t._d=new Date(+e[1])}(t):a(e)?(t._a=d(e.slice(0),(function(t){return parseInt(t,10)})),xe(t)):o(e)?function(t){if(!t._d){var e=B(t._i),n=void 0===e.day?e.date:e.day;t._a=d([e.year,e.month,n,e.hour,e.minute,e.second,e.millisecond],(function(t){return t&&parseInt(t,10)})),xe(t)}}(t):l(e)?t._d=new Date(e):i.createFromInputFallback(t)}(t),v(t)||(t._d=null),t))}function Ee(t,e,n,r,i){var u,c={};return!0!==e&&!1!==e||(r=e,e=void 0),!0!==n&&!1!==n||(r=n,n=void 0),(o(t)&&s(t)||a(t)&&0===t.length)&&(t=void 0),c._isAMomentObject=!0,c._useUTC=c._isUTC=i,c._l=n,c._i=t,c._f=e,c._strict=r,(u=new w(ce(Me(c))))._nextDay&&(u.add(1,"d"),u._nextDay=void 0),u}function Se(t,e,n,r){return Ee(t,e,n,r,!1)}i.createFromInputFallback=M("value provided is not in a recognized RFC2822 or ISO format. moment construction falls back to js Date(), which is not reliable across all browsers and versions. Non RFC2822/ISO date formats are discouraged. Please refer to http://momentjs.com/guides/#/warnings/js-date/ for more info.",(function(t){t._d=new Date(t._i+(t._useUTC?" UTC":""))})),i.ISO_8601=function(){},i.RFC_2822=function(){};var Te=M("moment().min is deprecated, use moment.max instead. http://momentjs.com/guides/#/warnings/min-max/",(function(){var t=Se.apply(null,arguments);return this.isValid()&&t.isValid()?t<this?this:t:g()})),je=M("moment().max is deprecated, use moment.min instead. http://momentjs.com/guides/#/warnings/min-max/",(function(){var t=Se.apply(null,arguments);return this.isValid()&&t.isValid()?t>this?this:t:g()}));function Oe(t,e){var n,r;if(1===e.length&&a(e[0])&&(e=e[0]),!e.length)return Se();for(n=e[0],r=1;r<e.length;++r)e[r].isValid()&&!e[r][t](n)||(n=e[r]);return n}var Ae=["year","quarter","month","week","day","hour","minute","second","millisecond"];function Le(t){var e=B(t),n=e.year||0,r=e.quarter||0,i=e.month||0,a=e.week||e.isoWeek||0,o=e.day||0,s=e.hour||0,c=e.minute||0,l=e.second||0,f=e.millisecond||0;this._isValid=function(t){var e,n,r=!1;for(e in t)if(u(t,e)&&(-1===gt.call(Ae,e)||null!=t[e]&&isNaN(t[e])))return!1;for(n=0;n<Ae.length;++n)if(t[Ae[n]]){if(r)return!1;parseFloat(t[Ae[n]])!==V(t[Ae[n]])&&(r=!0)}return!0}(e),this._milliseconds=+f+1e3*l+6e4*c+1e3*s*60*60,this._days=+o+7*a,this._months=+i+3*r+12*n,this._data={},this._locale=se(),this._bubble()}function Ce(t){return t instanceof Le}function De(t){return t<0?-1*Math.round(-1*t):Math.round(t)}function Ne(t,e){R(t,0,0,(function(){var t=this.utcOffset(),n="+";return t<0&&(t=-t,n="-"),n+L(~~(t/60),2)+e+L(~~t%60,2)}))}Ne("Z",":"),Ne("ZZ",""),pt("Z",dt),pt("ZZ",dt),yt(["Z","ZZ"],(function(t,e,n){n._useUTC=!0,n._tzm=Re(dt,t)}));var Pe=/([\+\-]|\d\d)/gi;function Re(t,e){var n,r,i=(e||"").match(t);return null===i?null:0===(r=60*(n=((i[i.length-1]||[])+"").match(Pe)||["-",0,0])[1]+V(n[2]))?0:"+"===n[0]?r:-r}function Ye(t,e){var n,r;return e._isUTC?(n=e.clone(),r=(x(t)||f(t)?t.valueOf():Se(t).valueOf())-n.valueOf(),n._d.setTime(n._d.valueOf()+r),i.updateOffset(n,!1),n):Se(t).local()}function Ie(t){return-Math.round(t._d.getTimezoneOffset())}function ze(){return!!this.isValid()&&this._isUTC&&0===this._offset}i.updateOffset=function(){};var Fe=/^(-|\+)?(?:(\d*)[. ])?(\d+):(\d+)(?::(\d+)(\.\d*)?)?$/,He=/^(-|\+)?P(?:([-+]?[0-9,.]*)Y)?(?:([-+]?[0-9,.]*)M)?(?:([-+]?[0-9,.]*)W)?(?:([-+]?[0-9,.]*)D)?(?:T(?:([-+]?[0-9,.]*)H)?(?:([-+]?[0-9,.]*)M)?(?:([-+]?[0-9,.]*)S)?)?$/;function Be(t,e){var n,r,i,a,o,s,c=t,f=null;return Ce(t)?c={ms:t._milliseconds,d:t._days,M:t._months}:l(t)||!isNaN(+t)?(c={},e?c[e]=+t:c.milliseconds=+t):(f=Fe.exec(t))?(n="-"===f[1]?-1:1,c={y:0,d:V(f[2])*n,h:V(f[3])*n,m:V(f[4])*n,s:V(f[5])*n,ms:V(De(1e3*f[6]))*n}):(f=He.exec(t))?(n="-"===f[1]?-1:1,c={y:qe(f[2],n),M:qe(f[3],n),w:qe(f[4],n),d:qe(f[5],n),h:qe(f[6],n),m:qe(f[7],n),s:qe(f[8],n)}):null==c?c={}:"object"==typeof c&&("from"in c||"to"in c)&&(a=Se(c.from),o=Se(c.to),i=a.isValid()&&o.isValid()?(o=Ye(o,a),a.isBefore(o)?s=Ue(a,o):((s=Ue(o,a)).milliseconds=-s.milliseconds,s.months=-s.months),s):{milliseconds:0,months:0},(c={}).ms=i.milliseconds,c.M=i.months),r=new Le(c),Ce(t)&&u(t,"_locale")&&(r._locale=t._locale),Ce(t)&&u(t,"_isValid")&&(r._isValid=t._isValid),r}function qe(t,e){var n=t&&parseFloat(t.replace(",","."));return(isNaN(n)?0:n)*e}function Ue(t,e){var n={};return n.months=e.month()-t.month()+12*(e.year()-t.year()),t.clone().add(n.months,"M").isAfter(e)&&--n.months,n.milliseconds=+e-+t.clone().add(n.months,"M"),n}function We(t,e){return function(n,r){var i;return null===r||isNaN(+r)||(T(e,"moment()."+e+"(period, number) is deprecated. Please use moment()."+e+"(number, period). See http://momentjs.com/guides/#/warnings/add-inverted-param/ for more info."),i=n,n=r,r=i),Ge(this,Be(n,r),t),this}}function Ge(t,e,n,r){var a=e._milliseconds,o=De(e._days),u=De(e._months);t.isValid()&&(r=null==r||r,u&&Ot(t,X(t,"Month")+u*n),o&&J(t,"Date",X(t,"Date")+o*n),a&&t._d.setTime(t._d.valueOf()+a*n),r&&i.updateOffset(t,o||u))}Be.fn=Le.prototype,Be.invalid=function(){return Be(NaN)};var Ve=We(1,"add"),$e=We(-1,"subtract");function Xe(t){return"string"==typeof t||t instanceof String}function Je(t){return x(t)||f(t)||Xe(t)||l(t)||function(t){var e=a(t),n=!1;return e&&(n=0===t.filter((function(e){return!l(e)&&Xe(t)})).length),e&&n}(t)||function(t){var e,n,r=o(t)&&!s(t),i=!1,a=["years","year","y","months","month","M","days","day","d","dates","date","D","hours","hour","h","minutes","minute","m","seconds","second","s","milliseconds","millisecond","ms"];for(e=0;e<a.length;e+=1)n=a[e],i=i||u(t,n);return r&&i}(t)||null==t}function Ze(t){var e,n=o(t)&&!s(t),r=!1,i=["sameDay","nextDay","lastDay","nextWeek","lastWeek","sameElse"];for(e=0;e<i.length;e+=1)r=r||u(t,i[e]);return n&&r}function Ke(t,e){if(t.date()<e.date())return-Ke(e,t);var n=12*(e.year()-t.year())+(e.month()-t.month()),r=t.clone().add(n,"months");return-(n+(e-r<0?(e-r)/(r-t.clone().add(n-1,"months")):(e-r)/(t.clone().add(n+1,"months")-r)))||0}function Qe(t){var e;return void 0===t?this._locale._abbr:(null!=(e=se(t))&&(this._locale=e),this)}i.defaultFormat="YYYY-MM-DDTHH:mm:ssZ",i.defaultFormatUtc="YYYY-MM-DDTHH:mm:ss[Z]";var tn=M("moment().lang() is deprecated. Instead, use moment().localeData() to get the language configuration. Use moment().locale() to change languages.",(function(t){return void 0===t?this.localeData():this.locale(t)}));function en(){return this._locale}function nn(t,e){return(t%e+e)%e}function rn(t,e,n){return t<100&&t>=0?new Date(t+400,e,n)-126227808e5:new Date(t,e,n).valueOf()}function an(t,e,n){return t<100&&t>=0?Date.UTC(t+400,e,n)-126227808e5:Date.UTC(t,e,n)}function on(t,e){return e.erasAbbrRegex(t)}function un(){var t,e,n=[],r=[],i=[],a=[],o=this.eras();for(t=0,e=o.length;t<e;++t)r.push(vt(o[t].name)),n.push(vt(o[t].abbr)),i.push(vt(o[t].narrow)),a.push(vt(o[t].name)),a.push(vt(o[t].abbr)),a.push(vt(o[t].narrow));this._erasRegex=new RegExp("^("+a.join("|")+")","i"),this._erasNameRegex=new RegExp("^("+r.join("|")+")","i"),this._erasAbbrRegex=new RegExp("^("+n.join("|")+")","i"),this._erasNarrowRegex=new RegExp("^("+i.join("|")+")","i")}function sn(t,e){R(0,[t,t.length],0,e)}function cn(t,e,n,r,i){var a;return null==t?It(this,r,i).year:(e>(a=zt(t,r,i))&&(e=a),ln.call(this,t,e,n,r,i))}function ln(t,e,n,r,i){var a=Yt(t,e,n,r,i),o=Pt(a.year,0,a.dayOfYear);return this.year(o.getUTCFullYear()),this.month(o.getUTCMonth()),this.date(o.getUTCDate()),this}R("N",0,0,"eraAbbr"),R("NN",0,0,"eraAbbr"),R("NNN",0,0,"eraAbbr"),R("NNNN",0,0,"eraName"),R("NNNNN",0,0,"eraNarrow"),R("y",["y",1],"yo","eraYear"),R("y",["yy",2],0,"eraYear"),R("y",["yyy",3],0,"eraYear"),R("y",["yyyy",4],0,"eraYear"),pt("N",on),pt("NN",on),pt("NNN",on),pt("NNNN",(function(t,e){return e.erasNameRegex(t)})),pt("NNNNN",(function(t,e){return e.erasNarrowRegex(t)})),yt(["N","NN","NNN","NNNN","NNNNN"],(function(t,e,n,r){var i=n._locale.erasParse(t,r,n._strict);i?m(n).era=i:m(n).invalidEra=t})),pt("y",ct),pt("yy",ct),pt("yyy",ct),pt("yyyy",ct),pt("yo",(function(t,e){return e._eraYearOrdinalRegex||ct})),yt(["y","yy","yyy","yyyy"],0),yt(["yo"],(function(t,e,n,r){var i;n._locale._eraYearOrdinalRegex&&(i=t.match(n._locale._eraYearOrdinalRegex)),n._locale.eraYearOrdinalParse?e[0]=n._locale.eraYearOrdinalParse(t,i):e[0]=parseInt(t,10)})),R(0,["gg",2],0,(function(){return this.weekYear()%100})),R(0,["GG",2],0,(function(){return this.isoWeekYear()%100})),sn("gggg","weekYear"),sn("ggggg","weekYear"),sn("GGGG","isoWeekYear"),sn("GGGGG","isoWeekYear"),F("weekYear","gg"),F("isoWeekYear","GG"),U("weekYear",1),U("isoWeekYear",1),pt("G",lt),pt("g",lt),pt("GG",rt,Q),pt("gg",rt,Q),pt("GGGG",ut,et),pt("gggg",ut,et),pt("GGGGG",st,nt),pt("ggggg",st,nt),bt(["gggg","ggggg","GGGG","GGGGG"],(function(t,e,n,r){e[r.substr(0,2)]=V(t)})),bt(["gg","GG"],(function(t,e,n,r){e[r]=i.parseTwoDigitYear(t)})),R("Q",0,"Qo","quarter"),F("quarter","Q"),U("quarter",7),pt("Q",K),yt("Q",(function(t,e){e[1]=3*(V(t)-1)})),R("D",["DD",2],"Do","date"),F("date","D"),U("date",9),pt("D",rt),pt("DD",rt,Q),pt("Do",(function(t,e){return t?e._dayOfMonthOrdinalParse||e._ordinalParse:e._dayOfMonthOrdinalParseLenient})),yt(["D","DD"],2),yt("Do",(function(t,e){e[2]=V(t.match(rt)[0])}));var fn=$("Date",!0);R("DDD",["DDDD",3],"DDDo","dayOfYear"),F("dayOfYear","DDD"),U("dayOfYear",4),pt("DDD",ot),pt("DDDD",tt),yt(["DDD","DDDD"],(function(t,e,n){n._dayOfYear=V(t)})),R("m",["mm",2],0,"minute"),F("minute","m"),U("minute",14),pt("m",rt),pt("mm",rt,Q),yt(["m","mm"],4);var dn=$("Minutes",!1);R("s",["ss",2],0,"second"),F("second","s"),U("second",15),pt("s",rt),pt("ss",rt,Q),yt(["s","ss"],5);var hn,pn,mn=$("Seconds",!1);for(R("S",0,0,(function(){return~~(this.millisecond()/100)})),R(0,["SS",2],0,(function(){return~~(this.millisecond()/10)})),R(0,["SSS",3],0,"millisecond"),R(0,["SSSS",4],0,(function(){return 10*this.millisecond()})),R(0,["SSSSS",5],0,(function(){return 100*this.millisecond()})),R(0,["SSSSSS",6],0,(function(){return 1e3*this.millisecond()})),R(0,["SSSSSSS",7],0,(function(){return 1e4*this.millisecond()})),R(0,["SSSSSSSS",8],0,(function(){return 1e5*this.millisecond()})),R(0,["SSSSSSSSS",9],0,(function(){return 1e6*this.millisecond()})),F("millisecond","ms"),U("millisecond",16),pt("S",ot,K),pt("SS",ot,Q),pt("SSS",ot,tt),hn="SSSS";hn.length<=9;hn+="S")pt(hn,ct);function vn(t,e){e[6]=V(1e3*("0."+t))}for(hn="S";hn.length<=9;hn+="S")yt(hn,vn);pn=$("Milliseconds",!1),R("z",0,0,"zoneAbbr"),R("zz",0,0,"zoneName");var gn=w.prototype;function _n(t){return t}gn.add=Ve,gn.calendar=function(t,e){1===arguments.length&&(arguments[0]?Je(arguments[0])?(t=arguments[0],e=void 0):Ze(arguments[0])&&(e=arguments[0],t=void 0):(t=void 0,e=void 0));var n=t||Se(),r=Ye(n,this).startOf("day"),a=i.calendarFormat(this,r)||"sameElse",o=e&&(j(e[a])?e[a].call(this,n):e[a]);return this.format(o||this.localeData().calendar(a,this,Se(n)))},gn.clone=function(){return new w(this)},gn.diff=function(t,e,n){var r,i,a;if(!this.isValid())return NaN;if(!(r=Ye(t,this)).isValid())return NaN;switch(i=6e4*(r.utcOffset()-this.utcOffset()),e=H(e)){case"year":a=Ke(this,r)/12;break;case"month":a=Ke(this,r);break;case"quarter":a=Ke(this,r)/3;break;case"second":a=(this-r)/1e3;break;case"minute":a=(this-r)/6e4;break;case"hour":a=(this-r)/36e5;break;case"day":a=(this-r-i)/864e5;break;case"week":a=(this-r-i)/6048e5;break;default:a=this-r}return n?a:G(a)},gn.endOf=function(t){var e,n;if(void 0===(t=H(t))||"millisecond"===t||!this.isValid())return this;switch(n=this._isUTC?an:rn,t){case"year":e=n(this.year()+1,0,1)-1;break;case"quarter":e=n(this.year(),this.month()-this.month()%3+3,1)-1;break;case"month":e=n(this.year(),this.month()+1,1)-1;break;case"week":e=n(this.year(),this.month(),this.date()-this.weekday()+7)-1;break;case"isoWeek":e=n(this.year(),this.month(),this.date()-(this.isoWeekday()-1)+7)-1;break;case"day":case"date":e=n(this.year(),this.month(),this.date()+1)-1;break;case"hour":e=this._d.valueOf(),e+=36e5-nn(e+(this._isUTC?0:6e4*this.utcOffset()),36e5)-1;break;case"minute":e=this._d.valueOf(),e+=6e4-nn(e,6e4)-1;break;case"second":e=this._d.valueOf(),e+=1e3-nn(e,1e3)-1}return this._d.setTime(e),i.updateOffset(this,!0),this},gn.format=function(t){t||(t=this.isUtc()?i.defaultFormatUtc:i.defaultFormat);var e=Y(this,t);return this.localeData().postformat(e)},gn.from=function(t,e){return this.isValid()&&(x(t)&&t.isValid()||Se(t).isValid())?Be({to:this,from:t}).locale(this.locale()).humanize(!e):this.localeData().invalidDate()},gn.fromNow=function(t){return this.from(Se(),t)},gn.to=function(t,e){return this.isValid()&&(x(t)&&t.isValid()||Se(t).isValid())?Be({from:this,to:t}).locale(this.locale()).humanize(!e):this.localeData().invalidDate()},gn.toNow=function(t){return this.to(Se(),t)},gn.get=function(t){return j(this[t=H(t)])?this[t]():this},gn.invalidAt=function(){return m(this).overflow},gn.isAfter=function(t,e){var n=x(t)?t:Se(t);return!(!this.isValid()||!n.isValid())&&("millisecond"===(e=H(e)||"millisecond")?this.valueOf()>n.valueOf():n.valueOf()<this.clone().startOf(e).valueOf())},gn.isBefore=function(t,e){var n=x(t)?t:Se(t);return!(!this.isValid()||!n.isValid())&&("millisecond"===(e=H(e)||"millisecond")?this.valueOf()<n.valueOf():this.clone().endOf(e).valueOf()<n.valueOf())},gn.isBetween=function(t,e,n,r){var i=x(t)?t:Se(t),a=x(e)?e:Se(e);return!!(this.isValid()&&i.isValid()&&a.isValid())&&(("("===(r=r||"()")[0]?this.isAfter(i,n):!this.isBefore(i,n))&&(")"===r[1]?this.isBefore(a,n):!this.isAfter(a,n)))},gn.isSame=function(t,e){var n,r=x(t)?t:Se(t);return!(!this.isValid()||!r.isValid())&&("millisecond"===(e=H(e)||"millisecond")?this.valueOf()===r.valueOf():(n=r.valueOf(),this.clone().startOf(e).valueOf()<=n&&n<=this.clone().endOf(e).valueOf()))},gn.isSameOrAfter=function(t,e){return this.isSame(t,e)||this.isAfter(t,e)},gn.isSameOrBefore=function(t,e){return this.isSame(t,e)||this.isBefore(t,e)},gn.isValid=function(){return v(this)},gn.lang=tn,gn.locale=Qe,gn.localeData=en,gn.max=je,gn.min=Te,gn.parsingFlags=function(){return h({},m(this))},gn.set=function(t,e){if("object"==typeof t){var n,r=function(t){var e,n=[];for(e in t)u(t,e)&&n.push({unit:e,priority:q[e]});return n.sort((function(t,e){return t.priority-e.priority})),n}(t=B(t));for(n=0;n<r.length;n++)this[r[n].unit](t[r[n].unit])}else if(j(this[t=H(t)]))return this[t](e);return this},gn.startOf=function(t){var e,n;if(void 0===(t=H(t))||"millisecond"===t||!this.isValid())return this;switch(n=this._isUTC?an:rn,t){case"year":e=n(this.year(),0,1);break;case"quarter":e=n(this.year(),this.month()-this.month()%3,1);break;case"month":e=n(this.year(),this.month(),1);break;case"week":e=n(this.year(),this.month(),this.date()-this.weekday());break;case"isoWeek":e=n(this.year(),this.month(),this.date()-(this.isoWeekday()-1));break;case"day":case"date":e=n(this.year(),this.month(),this.date());break;case"hour":e=this._d.valueOf(),e-=nn(e+(this._isUTC?0:6e4*this.utcOffset()),36e5);break;case"minute":e=this._d.valueOf(),e-=nn(e,6e4);break;case"second":e=this._d.valueOf(),e-=nn(e,1e3)}return this._d.setTime(e),i.updateOffset(this,!0),this},gn.subtract=$e,gn.toArray=function(){var t=this;return[t.year(),t.month(),t.date(),t.hour(),t.minute(),t.second(),t.millisecond()]},gn.toObject=function(){var t=this;return{years:t.year(),months:t.month(),date:t.date(),hours:t.hours(),minutes:t.minutes(),seconds:t.seconds(),milliseconds:t.milliseconds()}},gn.toDate=function(){return new Date(this.valueOf())},gn.toISOString=function(t){if(!this.isValid())return null;var e=!0!==t,n=e?this.clone().utc():this;return n.year()<0||n.year()>9999?Y(n,e?"YYYYYY-MM-DD[T]HH:mm:ss.SSS[Z]":"YYYYYY-MM-DD[T]HH:mm:ss.SSSZ"):j(Date.prototype.toISOString)?e?this.toDate().toISOString():new Date(this.valueOf()+60*this.utcOffset()*1e3).toISOString().replace("Z",Y(n,"Z")):Y(n,e?"YYYY-MM-DD[T]HH:mm:ss.SSS[Z]":"YYYY-MM-DD[T]HH:mm:ss.SSSZ")},gn.inspect=function(){if(!this.isValid())return"moment.invalid(/* "+this._i+" */)";var t,e,n,r="moment",i="";return this.isLocal()||(r=0===this.utcOffset()?"moment.utc":"moment.parseZone",i="Z"),t="["+r+'("]',e=0<=this.year()&&this.year()<=9999?"YYYY":"YYYYYY",n=i+'[")]',this.format(t+e+"-MM-DD[T]HH:mm:ss.SSS"+n)},"undefined"!=typeof Symbol&&null!=Symbol.for&&(gn[Symbol.for("nodejs.util.inspect.custom")]=function(){return"Moment<"+this.format()+">"}),gn.toJSON=function(){return this.isValid()?this.toISOString():null},gn.toString=function(){return this.clone().locale("en").format("ddd MMM DD YYYY HH:mm:ss [GMT]ZZ")},gn.unix=function(){return Math.floor(this.valueOf()/1e3)},gn.valueOf=function(){return this._d.valueOf()-6e4*(this._offset||0)},gn.creationData=function(){return{input:this._i,format:this._f,locale:this._locale,isUTC:this._isUTC,strict:this._strict}},gn.eraName=function(){var t,e,n,r=this.localeData().eras();for(t=0,e=r.length;t<e;++t){if(n=this.clone().startOf("day").valueOf(),r[t].since<=n&&n<=r[t].until)return r[t].name;if(r[t].until<=n&&n<=r[t].since)return r[t].name}return""},gn.eraNarrow=function(){var t,e,n,r=this.localeData().eras();for(t=0,e=r.length;t<e;++t){if(n=this.clone().startOf("day").valueOf(),r[t].since<=n&&n<=r[t].until)return r[t].narrow;if(r[t].until<=n&&n<=r[t].since)return r[t].narrow}return""},gn.eraAbbr=function(){var t,e,n,r=this.localeData().eras();for(t=0,e=r.length;t<e;++t){if(n=this.clone().startOf("day").valueOf(),r[t].since<=n&&n<=r[t].until)return r[t].abbr;if(r[t].until<=n&&n<=r[t].since)return r[t].abbr}return""},gn.eraYear=function(){var t,e,n,r,a=this.localeData().eras();for(t=0,e=a.length;t<e;++t)if(n=a[t].since<=a[t].until?1:-1,r=this.clone().startOf("day").valueOf(),a[t].since<=r&&r<=a[t].until||a[t].until<=r&&r<=a[t].since)return(this.year()-i(a[t].since).year())*n+a[t].offset;return this.year()},gn.year=Dt,gn.isLeapYear=function(){return W(this.year())},gn.weekYear=function(t){return cn.call(this,t,this.week(),this.weekday(),this.localeData()._week.dow,this.localeData()._week.doy)},gn.isoWeekYear=function(t){return cn.call(this,t,this.isoWeek(),this.isoWeekday(),1,4)},gn.quarter=gn.quarters=function(t){return null==t?Math.ceil((this.month()+1)/3):this.month(3*(t-1)+this.month()%3)},gn.month=At,gn.daysInMonth=function(){return xt(this.year(),this.month())},gn.week=gn.weeks=function(t){var e=this.localeData().week(this);return null==t?e:this.add(7*(t-e),"d")},gn.isoWeek=gn.isoWeeks=function(t){var e=It(this,1,4).week;return null==t?e:this.add(7*(t-e),"d")},gn.weeksInYear=function(){var t=this.localeData()._week;return zt(this.year(),t.dow,t.doy)},gn.weeksInWeekYear=function(){var t=this.localeData()._week;return zt(this.weekYear(),t.dow,t.doy)},gn.isoWeeksInYear=function(){return zt(this.year(),1,4)},gn.isoWeeksInISOWeekYear=function(){return zt(this.isoWeekYear(),1,4)},gn.date=fn,gn.day=gn.days=function(t){if(!this.isValid())return null!=t?this:NaN;var e=this._isUTC?this._d.getUTCDay():this._d.getDay();return null!=t?(t=function(t,e){return"string"!=typeof t?t:isNaN(t)?"number"==typeof(t=e.weekdaysParse(t))?t:null:parseInt(t,10)}(t,this.localeData()),this.add(t-e,"d")):e},gn.weekday=function(t){if(!this.isValid())return null!=t?this:NaN;var e=(this.day()+7-this.localeData()._week.dow)%7;return null==t?e:this.add(t-e,"d")},gn.isoWeekday=function(t){if(!this.isValid())return null!=t?this:NaN;if(null!=t){var e=function(t,e){return"string"==typeof t?e.weekdaysParse(t)%7||7:isNaN(t)?null:t}(t,this.localeData());return this.day(this.day()%7?e:e-7)}return this.day()||7},gn.dayOfYear=function(t){var e=Math.round((this.clone().startOf("day")-this.clone().startOf("year"))/864e5)+1;return null==t?e:this.add(t-e,"d")},gn.hour=gn.hours=Qt,gn.minute=gn.minutes=dn,gn.second=gn.seconds=mn,gn.millisecond=gn.milliseconds=pn,gn.utcOffset=function(t,e,n){var r,a=this._offset||0;if(!this.isValid())return null!=t?this:NaN;if(null!=t){if("string"==typeof t){if(null===(t=Re(dt,t)))return this}else Math.abs(t)<16&&!n&&(t*=60);return!this._isUTC&&e&&(r=Ie(this)),this._offset=t,this._isUTC=!0,null!=r&&this.add(r,"m"),a!==t&&(!e||this._changeInProgress?Ge(this,Be(t-a,"m"),1,!1):this._changeInProgress||(this._changeInProgress=!0,i.updateOffset(this,!0),this._changeInProgress=null)),this}return this._isUTC?a:Ie(this)},gn.utc=function(t){return this.utcOffset(0,t)},gn.local=function(t){return this._isUTC&&(this.utcOffset(0,t),this._isUTC=!1,t&&this.subtract(Ie(this),"m")),this},gn.parseZone=function(){if(null!=this._tzm)this.utcOffset(this._tzm,!1,!0);else if("string"==typeof this._i){var t=Re(ft,this._i);null!=t?this.utcOffset(t):this.utcOffset(0,!0)}return this},gn.hasAlignedHourOffset=function(t){return!!this.isValid()&&(t=t?Se(t).utcOffset():0,(this.utcOffset()-t)%60==0)},gn.isDST=function(){return this.utcOffset()>this.clone().month(0).utcOffset()||this.utcOffset()>this.clone().month(5).utcOffset()},gn.isLocal=function(){return!!this.isValid()&&!this._isUTC},gn.isUtcOffset=function(){return!!this.isValid()&&this._isUTC},gn.isUtc=ze,gn.isUTC=ze,gn.zoneAbbr=function(){return this._isUTC?"UTC":""},gn.zoneName=function(){return this._isUTC?"Coordinated Universal Time":""},gn.dates=M("dates accessor is deprecated. Use date instead.",fn),gn.months=M("months accessor is deprecated. Use month instead",At),gn.years=M("years accessor is deprecated. Use year instead",Dt),gn.zone=M("moment().zone is deprecated, use moment().utcOffset instead. http://momentjs.com/guides/#/warnings/zone/",(function(t,e){return null!=t?("string"!=typeof t&&(t=-t),this.utcOffset(t,e),this):-this.utcOffset()})),gn.isDSTShifted=M("isDSTShifted is deprecated. See http://momentjs.com/guides/#/warnings/dst-shifted/ for more information",(function(){if(!c(this._isDSTShifted))return this._isDSTShifted;var t,e={};return b(e,this),(e=Me(e))._a?(t=e._isUTC?p(e._a):Se(e._a),this._isDSTShifted=this.isValid()&&function(t,e,n){var r,i=Math.min(t.length,e.length),a=Math.abs(t.length-e.length),o=0;for(r=0;r<i;r++)(n&&t[r]!==e[r]||!n&&V(t[r])!==V(e[r]))&&o++;return o+a}(e._a,t.toArray())>0):this._isDSTShifted=!1,this._isDSTShifted}));var yn=A.prototype;function bn(t,e,n,r){var i=se(),a=p().set(r,e);return i[n](a,t)}function wn(t,e,n){if(l(t)&&(e=t,t=void 0),t=t||"",null!=e)return bn(t,e,n,"month");var r,i=[];for(r=0;r<12;r++)i[r]=bn(t,r,n,"month");return i}function xn(t,e,n,r){"boolean"==typeof t?(l(e)&&(n=e,e=void 0),e=e||""):(n=e=t,t=!1,l(e)&&(n=e,e=void 0),e=e||"");var i,a=se(),o=t?a._week.dow:0,u=[];if(null!=n)return bn(e,(n+o)%7,r,"day");for(i=0;i<7;i++)u[i]=bn(e,(i+o)%7,r,"day");return u}yn.calendar=function(t,e,n){var r=this._calendar[t]||this._calendar.sameElse;return j(r)?r.call(e,n):r},yn.longDateFormat=function(t){var e=this._longDateFormat[t],n=this._longDateFormat[t.toUpperCase()];return e||!n?e:(this._longDateFormat[t]=n.match(C).map((function(t){return"MMMM"===t||"MM"===t||"DD"===t||"dddd"===t?t.slice(1):t})).join(""),this._longDateFormat[t])},yn.invalidDate=function(){return this._invalidDate},yn.ordinal=function(t){return this._ordinal.replace("%d",t)},yn.preparse=_n,yn.postformat=_n,yn.relativeTime=function(t,e,n,r){var i=this._relativeTime[n];return j(i)?i(t,e,n,r):i.replace(/%d/i,t)},yn.pastFuture=function(t,e){var n=this._relativeTime[t>0?"future":"past"];return j(n)?n(e):n.replace(/%s/i,e)},yn.set=function(t){var e,n;for(n in t)u(t,n)&&(j(e=t[n])?this[n]=e:this["_"+n]=e);this._config=t,this._dayOfMonthOrdinalParseLenient=new RegExp((this._dayOfMonthOrdinalParse.source||this._ordinalParse.source)+"|"+/\d{1,2}/.source)},yn.eras=function(t,e){var n,r,a,o=this._eras||se("en")._eras;for(n=0,r=o.length;n<r;++n){switch(typeof o[n].since){case"string":a=i(o[n].since).startOf("day"),o[n].since=a.valueOf()}switch(typeof o[n].until){case"undefined":o[n].until=1/0;break;case"string":a=i(o[n].until).startOf("day").valueOf(),o[n].until=a.valueOf()}}return o},yn.erasParse=function(t,e,n){var r,i,a,o,u,s=this.eras();for(t=t.toUpperCase(),r=0,i=s.length;r<i;++r)if(a=s[r].name.toUpperCase(),o=s[r].abbr.toUpperCase(),u=s[r].narrow.toUpperCase(),n)switch(e){case"N":case"NN":case"NNN":if(o===t)return s[r];break;case"NNNN":if(a===t)return s[r];break;case"NNNNN":if(u===t)return s[r]}else if([a,o,u].indexOf(t)>=0)return s[r]},yn.erasConvertYear=function(t,e){var n=t.since<=t.until?1:-1;return void 0===e?i(t.since).year():i(t.since).year()+(e-t.offset)*n},yn.erasAbbrRegex=function(t){return u(this,"_erasAbbrRegex")||un.call(this),t?this._erasAbbrRegex:this._erasRegex},yn.erasNameRegex=function(t){return u(this,"_erasNameRegex")||un.call(this),t?this._erasNameRegex:this._erasRegex},yn.erasNarrowRegex=function(t){return u(this,"_erasNarrowRegex")||un.call(this),t?this._erasNarrowRegex:this._erasRegex},yn.months=function(t,e){return t?a(this._months)?this._months[t.month()]:this._months[(this._months.isFormat||Et).test(e)?"format":"standalone"][t.month()]:a(this._months)?this._months:this._months.standalone},yn.monthsShort=function(t,e){return t?a(this._monthsShort)?this._monthsShort[t.month()]:this._monthsShort[Et.test(e)?"format":"standalone"][t.month()]:a(this._monthsShort)?this._monthsShort:this._monthsShort.standalone},yn.monthsParse=function(t,e,n){var r,i,a;if(this._monthsParseExact)return jt.call(this,t,e,n);for(this._monthsParse||(this._monthsParse=[],this._longMonthsParse=[],this._shortMonthsParse=[]),r=0;r<12;r++){if(i=p([2e3,r]),n&&!this._longMonthsParse[r]&&(this._longMonthsParse[r]=new RegExp("^"+this.months(i,"").replace(".","")+"$","i"),this._shortMonthsParse[r]=new RegExp("^"+this.monthsShort(i,"").replace(".","")+"$","i")),n||this._monthsParse[r]||(a="^"+this.months(i,"")+"|^"+this.monthsShort(i,""),this._monthsParse[r]=new RegExp(a.replace(".",""),"i")),n&&"MMMM"===e&&this._longMonthsParse[r].test(t))return r;if(n&&"MMM"===e&&this._shortMonthsParse[r].test(t))return r;if(!n&&this._monthsParse[r].test(t))return r}},yn.monthsRegex=function(t){return this._monthsParseExact?(u(this,"_monthsRegex")||Lt.call(this),t?this._monthsStrictRegex:this._monthsRegex):(u(this,"_monthsRegex")||(this._monthsRegex=Tt),this._monthsStrictRegex&&t?this._monthsStrictRegex:this._monthsRegex)},yn.monthsShortRegex=function(t){return this._monthsParseExact?(u(this,"_monthsRegex")||Lt.call(this),t?this._monthsShortStrictRegex:this._monthsShortRegex):(u(this,"_monthsShortRegex")||(this._monthsShortRegex=St),this._monthsShortStrictRegex&&t?this._monthsShortStrictRegex:this._monthsShortRegex)},yn.week=function(t){return It(t,this._week.dow,this._week.doy).week},yn.firstDayOfYear=function(){return this._week.doy},yn.firstDayOfWeek=function(){return this._week.dow},yn.weekdays=function(t,e){var n=a(this._weekdays)?this._weekdays:this._weekdays[t&&!0!==t&&this._weekdays.isFormat.test(e)?"format":"standalone"];return!0===t?Ft(n,this._week.dow):t?n[t.day()]:n},yn.weekdaysMin=function(t){return!0===t?Ft(this._weekdaysMin,this._week.dow):t?this._weekdaysMin[t.day()]:this._weekdaysMin},yn.weekdaysShort=function(t){return!0===t?Ft(this._weekdaysShort,this._week.dow):t?this._weekdaysShort[t.day()]:this._weekdaysShort},yn.weekdaysParse=function(t,e,n){var r,i,a;if(this._weekdaysParseExact)return Vt.call(this,t,e,n);for(this._weekdaysParse||(this._weekdaysParse=[],this._minWeekdaysParse=[],this._shortWeekdaysParse=[],this._fullWeekdaysParse=[]),r=0;r<7;r++){if(i=p([2e3,1]).day(r),n&&!this._fullWeekdaysParse[r]&&(this._fullWeekdaysParse[r]=new RegExp("^"+this.weekdays(i,"").replace(".","\\.?")+"$","i"),this._shortWeekdaysParse[r]=new RegExp("^"+this.weekdaysShort(i,"").replace(".","\\.?")+"$","i"),this._minWeekdaysParse[r]=new RegExp("^"+this.weekdaysMin(i,"").replace(".","\\.?")+"$","i")),this._weekdaysParse[r]||(a="^"+this.weekdays(i,"")+"|^"+this.weekdaysShort(i,"")+"|^"+this.weekdaysMin(i,""),this._weekdaysParse[r]=new RegExp(a.replace(".",""),"i")),n&&"dddd"===e&&this._fullWeekdaysParse[r].test(t))return r;if(n&&"ddd"===e&&this._shortWeekdaysParse[r].test(t))return r;if(n&&"dd"===e&&this._minWeekdaysParse[r].test(t))return r;if(!n&&this._weekdaysParse[r].test(t))return r}},yn.weekdaysRegex=function(t){return this._weekdaysParseExact?(u(this,"_weekdaysRegex")||$t.call(this),t?this._weekdaysStrictRegex:this._weekdaysRegex):(u(this,"_weekdaysRegex")||(this._weekdaysRegex=Ut),this._weekdaysStrictRegex&&t?this._weekdaysStrictRegex:this._weekdaysRegex)},yn.weekdaysShortRegex=function(t){return this._weekdaysParseExact?(u(this,"_weekdaysRegex")||$t.call(this),t?this._weekdaysShortStrictRegex:this._weekdaysShortRegex):(u(this,"_weekdaysShortRegex")||(this._weekdaysShortRegex=Wt),this._weekdaysShortStrictRegex&&t?this._weekdaysShortStrictRegex:this._weekdaysShortRegex)},yn.weekdaysMinRegex=function(t){return this._weekdaysParseExact?(u(this,"_weekdaysRegex")||$t.call(this),t?this._weekdaysMinStrictRegex:this._weekdaysMinRegex):(u(this,"_weekdaysMinRegex")||(this._weekdaysMinRegex=Gt),this._weekdaysMinStrictRegex&&t?this._weekdaysMinStrictRegex:this._weekdaysMinRegex)},yn.isPM=function(t){return"p"===(t+"").toLowerCase().charAt(0)},yn.meridiem=function(t,e,n){return t>11?n?"pm":"PM":n?"am":"AM"},oe("en",{eras:[{since:"0001-01-01",until:1/0,offset:1,name:"Anno Domini",narrow:"AD",abbr:"AD"},{since:"0000-12-31",until:-1/0,offset:1,name:"Before Christ",narrow:"BC",abbr:"BC"}],dayOfMonthOrdinalParse:/\d{1,2}(th|st|nd|rd)/,ordinal:function(t){var e=t%10;return t+(1===V(t%100/10)?"th":1===e?"st":2===e?"nd":3===e?"rd":"th")}}),i.lang=M("moment.lang is deprecated. Use moment.locale instead.",oe),i.langData=M("moment.langData is deprecated. Use moment.localeData instead.",se);var kn=Math.abs;function Mn(t,e,n,r){var i=Be(e,n);return t._milliseconds+=r*i._milliseconds,t._days+=r*i._days,t._months+=r*i._months,t._bubble()}function En(t){return t<0?Math.floor(t):Math.ceil(t)}function Sn(t){return 4800*t/146097}function Tn(t){return 146097*t/4800}function jn(t){return function(){return this.as(t)}}var On=jn("ms"),An=jn("s"),Ln=jn("m"),Cn=jn("h"),Dn=jn("d"),Nn=jn("w"),Pn=jn("M"),Rn=jn("Q"),Yn=jn("y");function In(t){return function(){return this.isValid()?this._data[t]:NaN}}var zn=In("milliseconds"),Fn=In("seconds"),Hn=In("minutes"),Bn=In("hours"),qn=In("days"),Un=In("months"),Wn=In("years"),Gn=Math.round,Vn={ss:44,s:45,m:45,h:22,d:26,w:null,M:11};function $n(t,e,n,r,i){return i.relativeTime(e||1,!!n,t,r)}var Xn=Math.abs;function Jn(t){return(t>0)-(t<0)||+t}function Zn(){if(!this.isValid())return this.localeData().invalidDate();var t,e,n,r,i,a,o,u,s=Xn(this._milliseconds)/1e3,c=Xn(this._days),l=Xn(this._months),f=this.asSeconds();return f?(t=G(s/60),e=G(t/60),s%=60,t%=60,n=G(l/12),l%=12,r=s?s.toFixed(3).replace(/\.?0+$/,""):"",i=f<0?"-":"",a=Jn(this._months)!==Jn(f)?"-":"",o=Jn(this._days)!==Jn(f)?"-":"",u=Jn(this._milliseconds)!==Jn(f)?"-":"",i+"P"+(n?a+n+"Y":"")+(l?a+l+"M":"")+(c?o+c+"D":"")+(e||t||s?"T":"")+(e?u+e+"H":"")+(t?u+t+"M":"")+(s?u+r+"S":"")):"P0D"}var Kn=Le.prototype;return Kn.isValid=function(){return this._isValid},Kn.abs=function(){var t=this._data;return this._milliseconds=kn(this._milliseconds),this._days=kn(this._days),this._months=kn(this._months),t.milliseconds=kn(t.milliseconds),t.seconds=kn(t.seconds),t.minutes=kn(t.minutes),t.hours=kn(t.hours),t.months=kn(t.months),t.years=kn(t.years),this},Kn.add=function(t,e){return Mn(this,t,e,1)},Kn.subtract=function(t,e){return Mn(this,t,e,-1)},Kn.as=function(t){if(!this.isValid())return NaN;var e,n,r=this._milliseconds;if("month"===(t=H(t))||"quarter"===t||"year"===t)switch(e=this._days+r/864e5,n=this._months+Sn(e),t){case"month":return n;case"quarter":return n/3;case"year":return n/12}else switch(e=this._days+Math.round(Tn(this._months)),t){case"week":return e/7+r/6048e5;case"day":return e+r/864e5;case"hour":return 24*e+r/36e5;case"minute":return 1440*e+r/6e4;case"second":return 86400*e+r/1e3;case"millisecond":return Math.floor(864e5*e)+r;default:throw new Error("Unknown unit "+t)}},Kn.asMilliseconds=On,Kn.asSeconds=An,Kn.asMinutes=Ln,Kn.asHours=Cn,Kn.asDays=Dn,Kn.asWeeks=Nn,Kn.asMonths=Pn,Kn.asQuarters=Rn,Kn.asYears=Yn,Kn.valueOf=function(){return this.isValid()?this._milliseconds+864e5*this._days+this._months%12*2592e6+31536e6*V(this._months/12):NaN},Kn._bubble=function(){var t,e,n,r,i,a=this._milliseconds,o=this._days,u=this._months,s=this._data;return a>=0&&o>=0&&u>=0||a<=0&&o<=0&&u<=0||(a+=864e5*En(Tn(u)+o),o=0,u=0),s.milliseconds=a%1e3,t=G(a/1e3),s.seconds=t%60,e=G(t/60),s.minutes=e%60,n=G(e/60),s.hours=n%24,o+=G(n/24),i=G(Sn(o)),u+=i,o-=En(Tn(i)),r=G(u/12),u%=12,s.days=o,s.months=u,s.years=r,this},Kn.clone=function(){return Be(this)},Kn.get=function(t){return t=H(t),this.isValid()?this[t+"s"]():NaN},Kn.milliseconds=zn,Kn.seconds=Fn,Kn.minutes=Hn,Kn.hours=Bn,Kn.days=qn,Kn.weeks=function(){return G(this.days()/7)},Kn.months=Un,Kn.years=Wn,Kn.humanize=function(t,e){if(!this.isValid())return this.localeData().invalidDate();var n,r,i=!1,a=Vn;return"object"==typeof t&&(e=t,t=!1),"boolean"==typeof t&&(i=t),"object"==typeof e&&(a=Object.assign({},Vn,e),null!=e.s&&null==e.ss&&(a.ss=e.s-1)),n=this.localeData(),r=function(t,e,n,r){var i=Be(t).abs(),a=Gn(i.as("s")),o=Gn(i.as("m")),u=Gn(i.as("h")),s=Gn(i.as("d")),c=Gn(i.as("M")),l=Gn(i.as("w")),f=Gn(i.as("y")),d=a<=n.ss&&["s",a]||a<n.s&&["ss",a]||o<=1&&["m"]||o<n.m&&["mm",o]||u<=1&&["h"]||u<n.h&&["hh",u]||s<=1&&["d"]||s<n.d&&["dd",s];return null!=n.w&&(d=d||l<=1&&["w"]||l<n.w&&["ww",l]),(d=d||c<=1&&["M"]||c<n.M&&["MM",c]||f<=1&&["y"]||["yy",f])[2]=e,d[3]=+t>0,d[4]=r,$n.apply(null,d)}(this,!i,a,n),i&&(r=n.pastFuture(+this,r)),n.postformat(r)},Kn.toISOString=Zn,Kn.toString=Zn,Kn.toJSON=Zn,Kn.locale=Qe,Kn.localeData=en,Kn.toIsoString=M("toIsoString() is deprecated. Please use toISOString() instead (notice the capitals)",Zn),Kn.lang=tn,R("X",0,0,"unix"),R("x",0,0,"valueOf"),pt("x",lt),pt("X",/[+-]?\d+(\.\d{1,3})?/),yt("X",(function(t,e,n){n._d=new Date(1e3*parseFloat(t))})),yt("x",(function(t,e,n){n._d=new Date(V(t))})),
//! moment.js
i.version="2.29.1",e=Se,i.fn=gn,i.min=function(){var t=[].slice.call(arguments,0);return Oe("isBefore",t)},i.max=function(){var t=[].slice.call(arguments,0);return Oe("isAfter",t)},i.now=function(){return Date.now?Date.now():+new Date},i.utc=p,i.unix=function(t){return Se(1e3*t)},i.months=function(t,e){return wn(t,e,"months")},i.isDate=f,i.locale=oe,i.invalid=g,i.duration=Be,i.isMoment=x,i.weekdays=function(t,e,n){return xn(t,e,n,"weekdays")},i.parseZone=function(){return Se.apply(null,arguments).parseZone()},i.localeData=se,i.isDuration=Ce,i.monthsShort=function(t,e){return wn(t,e,"monthsShort")},i.weekdaysMin=function(t,e,n){return xn(t,e,n,"weekdaysMin")},i.defineLocale=ue,i.updateLocale=function(t,e){if(null!=e){var n,r,i=te;null!=ee[t]&&null!=ee[t].parentLocale?ee[t].set(O(ee[t]._config,e)):(null!=(r=ae(t))&&(i=r._config),e=O(i,e),null==r&&(e.abbr=t),(n=new A(e)).parentLocale=ee[t],ee[t]=n),oe(t)}else null!=ee[t]&&(null!=ee[t].parentLocale?(ee[t]=ee[t].parentLocale,t===oe()&&oe(t)):null!=ee[t]&&delete ee[t]);return ee[t]},i.locales=function(){return E(ee)},i.weekdaysShort=function(t,e,n){return xn(t,e,n,"weekdaysShort")},i.normalizeUnits=H,i.relativeTimeRounding=function(t){return void 0===t?Gn:"function"==typeof t&&(Gn=t,!0)},i.relativeTimeThreshold=function(t,e){return void 0!==Vn[t]&&(void 0===e?Vn[t]:(Vn[t]=e,"s"===t&&(Vn.ss=e-1),!0))},i.calendarFormat=function(t,e){var n=t.diff(e,"days",!0);return n<-6?"sameElse":n<-1?"lastWeek":n<0?"lastDay":n<1?"sameDay":n<2?"nextDay":n<7?"nextWeek":"sameElse"},i.prototype=gn,i.HTML5_FMT={DATETIME_LOCAL:"YYYY-MM-DDTHH:mm",DATETIME_LOCAL_SECONDS:"YYYY-MM-DDTHH:mm:ss",DATETIME_LOCAL_MS:"YYYY-MM-DDTHH:mm:ss.SSS",DATE:"YYYY-MM-DD",TIME:"HH:mm",TIME_SECONDS:"HH:mm:ss",TIME_MS:"HH:mm:ss.SSS",WEEK:"GGGG-[W]WW",MONTH:"YYYY-MM"},i}()}).call(this,n(27)(t))},function(t,e,n){"use strict";t.exports=n(512)},function(t,e,n){"use strict";n.d(e,"b",(function(){return s})),n.d(e,"d",(function(){return o})),n.d(e,"c",(function(){return u})),n.d(e,"a",(function(){return r})),n.d(e,"e",(function(){return i})),n.d(e,"f",(function(){return f})),n.d(e,"g",(function(){return d})),n.d(e,"h",(function(){return m})),n.d(e,"i",(function(){return v})),n.d(e,"j",(function(){return A})),n.d(e,"w",(function(){return C})),n.d(e,"x",(function(){return D})),n.d(e,"y",(function(){return O})),n.d(e,"k",(function(){return N})),n.d(e,"l",(function(){return P})),n.d(e,"m",(function(){return R})),n.d(e,"n",(function(){return Y})),n.d(e,"o",(function(){return I})),n.d(e,"p",(function(){return c})),n.d(e,"q",(function(){return z})),n.d(e,"r",(function(){return L})),n.d(e,"s",(function(){return x})),n.d(e,"t",(function(){return F})),n.d(e,"u",(function(){return H})),n.d(e,"v",(function(){return B})),n.d(e,"B",(function(){return S})),n.d(e,"z",(function(){return T})),n.d(e,"A",(function(){return j})),n.d(e,"C",(function(){return q})),n.d(e,"D",(function(){return p})),n.d(e,"E",(function(){return W}));var r=function(t,e){return t<e?-1:t>e?1:t>=e?0:NaN},i=function(t){var e;return 1===t.length&&(e=t,t=function(t,n){return r(e(t),n)}),{left:function(e,n,r,i){for(null==r&&(r=0),null==i&&(i=e.length);r<i;){var a=r+i>>>1;t(e[a],n)<0?r=a+1:i=a}return r},right:function(e,n,r,i){for(null==r&&(r=0),null==i&&(i=e.length);r<i;){var a=r+i>>>1;t(e[a],n)>0?i=a:r=a+1}return r}}};var a=i(r),o=a.right,u=a.left,s=o,c=function(t,e){null==e&&(e=l);for(var n=0,r=t.length-1,i=t[0],a=new Array(r<0?0:r);n<r;)a[n]=e(i,i=t[++n]);return a};function l(t,e){return[t,e]}var f=function(t,e,n){var r,i,a,o,u=t.length,s=e.length,c=new Array(u*s);for(null==n&&(n=l),r=a=0;r<u;++r)for(o=t[r],i=0;i<s;++i,++a)c[a]=n(o,e[i]);return c},d=function(t,e){return e<t?-1:e>t?1:e>=t?0:NaN},h=function(t){return null===t?NaN:+t},p=function(t,e){var n,r,i=t.length,a=0,o=-1,u=0,s=0;if(null==e)for(;++o<i;)isNaN(n=h(t[o]))||(s+=(r=n-u)*(n-(u+=r/++a)));else for(;++o<i;)isNaN(n=h(e(t[o],o,t)))||(s+=(r=n-u)*(n-(u+=r/++a)));if(a>1)return s/(a-1)},m=function(t,e){var n=p(t,e);return n?Math.sqrt(n):n},v=function(t,e){var n,r,i,a=t.length,o=-1;if(null==e){for(;++o<a;)if(null!=(n=t[o])&&n>=n)for(r=i=n;++o<a;)null!=(n=t[o])&&(r>n&&(r=n),i<n&&(i=n))}else for(;++o<a;)if(null!=(n=e(t[o],o,t))&&n>=n)for(r=i=n;++o<a;)null!=(n=e(t[o],o,t))&&(r>n&&(r=n),i<n&&(i=n));return[r,i]},g=Array.prototype,_=g.slice,y=g.map,b=function(t){return function(){return t}},w=function(t){return t},x=function(t,e,n){t=+t,e=+e,n=(i=arguments.length)<2?(e=t,t=0,1):i<3?1:+n;for(var r=-1,i=0|Math.max(0,Math.ceil((e-t)/n)),a=new Array(i);++r<i;)a[r]=t+r*n;return a},k=Math.sqrt(50),M=Math.sqrt(10),E=Math.sqrt(2),S=function(t,e,n){var r,i,a,o,u=-1;if(n=+n,(t=+t)===(e=+e)&&n>0)return[t];if((r=e<t)&&(i=t,t=e,e=i),0===(o=T(t,e,n))||!isFinite(o))return[];if(o>0)for(t=Math.ceil(t/o),e=Math.floor(e/o),a=new Array(i=Math.ceil(e-t+1));++u<i;)a[u]=(t+u)*o;else for(t=Math.floor(t*o),e=Math.ceil(e*o),a=new Array(i=Math.ceil(t-e+1));++u<i;)a[u]=(t-u)/o;return r&&a.reverse(),a};function T(t,e,n){var r=(e-t)/Math.max(0,n),i=Math.floor(Math.log(r)/Math.LN10),a=r/Math.pow(10,i);return i>=0?(a>=k?10:a>=M?5:a>=E?2:1)*Math.pow(10,i):-Math.pow(10,-i)/(a>=k?10:a>=M?5:a>=E?2:1)}function j(t,e,n){var r=Math.abs(e-t)/Math.max(0,n),i=Math.pow(10,Math.floor(Math.log(r)/Math.LN10)),a=r/i;return a>=k?i*=10:a>=M?i*=5:a>=E&&(i*=2),e<t?-i:i}var O=function(t){return Math.ceil(Math.log(t.length)/Math.LN2)+1},A=function(){var t=w,e=v,n=O;function r(r){var i,a,o=r.length,u=new Array(o);for(i=0;i<o;++i)u[i]=t(r[i],i,r);var c=e(u),l=c[0],f=c[1],d=n(u,l,f);Array.isArray(d)||(d=j(l,f,d),d=x(Math.ceil(l/d)*d,f,d));for(var h=d.length;d[0]<=l;)d.shift(),--h;for(;d[h-1]>f;)d.pop(),--h;var p,m=new Array(h+1);for(i=0;i<=h;++i)(p=m[i]=[]).x0=i>0?d[i-1]:l,p.x1=i<h?d[i]:f;for(i=0;i<o;++i)l<=(a=u[i])&&a<=f&&m[s(d,a,0,h)].push(r[i]);return m}return r.value=function(e){return arguments.length?(t="function"==typeof e?e:b(e),r):t},r.domain=function(t){return arguments.length?(e="function"==typeof t?t:b([t[0],t[1]]),r):e},r.thresholds=function(t){return arguments.length?(n="function"==typeof t?t:Array.isArray(t)?b(_.call(t)):b(t),r):n},r},L=function(t,e,n){if(null==n&&(n=h),r=t.length){if((e=+e)<=0||r<2)return+n(t[0],0,t);if(e>=1)return+n(t[r-1],r-1,t);var r,i=(r-1)*e,a=Math.floor(i),o=+n(t[a],a,t);return o+(+n(t[a+1],a+1,t)-o)*(i-a)}},C=function(t,e,n){return t=y.call(t,h).sort(r),Math.ceil((n-e)/(2*(L(t,.75)-L(t,.25))*Math.pow(t.length,-1/3)))},D=function(t,e,n){return Math.ceil((n-e)/(3.5*m(t)*Math.pow(t.length,-1/3)))},N=function(t,e){var n,r,i=t.length,a=-1;if(null==e){for(;++a<i;)if(null!=(n=t[a])&&n>=n)for(r=n;++a<i;)null!=(n=t[a])&&n>r&&(r=n)}else for(;++a<i;)if(null!=(n=e(t[a],a,t))&&n>=n)for(r=n;++a<i;)null!=(n=e(t[a],a,t))&&n>r&&(r=n);return r},P=function(t,e){var n,r=t.length,i=r,a=-1,o=0;if(null==e)for(;++a<r;)isNaN(n=h(t[a]))?--i:o+=n;else for(;++a<r;)isNaN(n=h(e(t[a],a,t)))?--i:o+=n;if(i)return o/i},R=function(t,e){var n,i=t.length,a=-1,o=[];if(null==e)for(;++a<i;)isNaN(n=h(t[a]))||o.push(n);else for(;++a<i;)isNaN(n=h(e(t[a],a,t)))||o.push(n);return L(o.sort(r),.5)},Y=function(t){for(var e,n,r,i=t.length,a=-1,o=0;++a<i;)o+=t[a].length;for(n=new Array(o);--i>=0;)for(e=(r=t[i]).length;--e>=0;)n[--o]=r[e];return n},I=function(t,e){var n,r,i=t.length,a=-1;if(null==e){for(;++a<i;)if(null!=(n=t[a])&&n>=n)for(r=n;++a<i;)null!=(n=t[a])&&r>n&&(r=n)}else for(;++a<i;)if(null!=(n=e(t[a],a,t))&&n>=n)for(r=n;++a<i;)null!=(n=e(t[a],a,t))&&r>n&&(r=n);return r},z=function(t,e){for(var n=e.length,r=new Array(n);n--;)r[n]=t[e[n]];return r},F=function(t,e){if(n=t.length){var n,i,a=0,o=0,u=t[o];for(null==e&&(e=r);++a<n;)(e(i=t[a],u)<0||0!==e(u,u))&&(u=i,o=a);return 0===e(u,u)?o:void 0}},H=function(t,e,n){for(var r,i,a=(null==n?t.length:n)-(e=null==e?0:+e);a;)i=Math.random()*a--|0,r=t[a+e],t[a+e]=t[i+e],t[i+e]=r;return t},B=function(t,e){var n,r=t.length,i=-1,a=0;if(null==e)for(;++i<r;)(n=+t[i])&&(a+=n);else for(;++i<r;)(n=+e(t[i],i,t))&&(a+=n);return a},q=function(t){if(!(i=t.length))return[];for(var e=-1,n=I(t,U),r=new Array(n);++e<n;)for(var i,a=-1,o=r[e]=new Array(i);++a<i;)o[a]=t[a][e];return r};function U(t){return t.length}var W=function(){return q(arguments)}},function(t,e,n){"use strict";e.a=function(t){for(var e=t.length/6|0,n=new Array(e),r=0;r<e;)n[r]="#"+t.slice(6*r,6*++r);return n}},function(t,e,n){"use strict";(function(t){n.d(e,"e",(function(){return r})),n.d(e,"p",(function(){return i})),n.d(e,"a",(function(){return a})),n.d(e,"c",(function(){return o})),n.d(e,"d",(function(){return u})),n.d(e,"o",(function(){return s})),n.d(e,"q",(function(){return c})),n.d(e,"t",(function(){return l})),n.d(e,"i",(function(){return f})),n.d(e,"r",(function(){return d})),n.d(e,"s",(function(){return h})),n.d(e,"k",(function(){return p})),n.d(e,"m",(function(){return m})),n.d(e,"j",(function(){return v})),n.d(e,"l",(function(){return g})),n.d(e,"g",(function(){return _})),n.d(e,"f",(function(){return y})),n.d(e,"h",(function(){return b})),n.d(e,"n",(function(){return w})),n.d(e,"b",(function(){return x}));var r="1.12.0",i="object"==typeof self&&self.self===self&&self||"object"==typeof t&&t.global===t&&t||Function("return this")()||{},a=Array.prototype,o=Object.prototype,u="undefined"!=typeof Symbol?Symbol.prototype:null,s=a.push,c=a.slice,l=o.toString,f=o.hasOwnProperty,d="undefined"!=typeof ArrayBuffer,h="undefined"!=typeof DataView,p=Array.isArray,m=Object.keys,v=Object.create,g=d&&ArrayBuffer.isView,_=isNaN,y=isFinite,b=!{toString:null}.propertyIsEnumerable("toString"),w=["valueOf","isPrototypeOf","toString","propertyIsEnumerable","hasOwnProperty","toLocaleString"],x=Math.pow(2,53)-1}).call(this,n(6))},function(t,e,n){"use strict";e.a=function(t){return function(){return t}}},function(t,e){var n;n=function(){return this}();try{n=n||new Function("return this")()}catch(t){"object"==typeof window&&(n=window)}t.exports=n},function(t,e,n){"use strict";n.d(e,"a",(function(){return a}));var r=new Date,i=new Date;function a(t,e,n,o){function u(e){return t(e=0===arguments.length?new Date:new Date(+e)),e}return u.floor=function(e){return t(e=new Date(+e)),e},u.ceil=function(n){return t(n=new Date(n-1)),e(n,1),t(n),n},u.round=function(t){var e=u(t),n=u.ceil(t);return t-e<n-t?e:n},u.offset=function(t,n){return e(t=new Date(+t),null==n?1:Math.floor(n)),t},u.range=function(n,r,i){var a,o=[];if(n=u.ceil(n),i=null==i?1:Math.floor(i),!(n<r&&i>0))return o;do{o.push(a=new Date(+n)),e(n,i),t(n)}while(a<n&&n<r);return o},u.filter=function(n){return a((function(e){if(e>=e)for(;t(e),!n(e);)e.setTime(e-1)}),(function(t,r){if(t>=t)if(r<0)for(;++r<=0;)for(;e(t,-1),!n(t););else for(;--r>=0;)for(;e(t,1),!n(t););}))},n&&(u.count=function(e,a){return r.setTime(+e),i.setTime(+a),t(r),t(i),Math.floor(n(r,i))},u.every=function(t){return t=Math.floor(t),isFinite(t)&&t>0?t>1?u.filter(o?function(e){return o(e)%t==0}:function(e){return u.count(0,e)%t==0}):u:null}),u}},function(t,e,n){(function(e){t.exports=e.jQuery=n(471)}).call(this,n(6))},function(t,e,n){"use strict";n.d(e,"d",(function(){return r})),n.d(e,"c",(function(){return i})),n.d(e,"b",(function(){return a})),n.d(e,"a",(function(){return o})),n.d(e,"e",(function(){return u}));var r=1e3,i=6e4,a=36e5,o=864e5,u=6048e5},function(t,e,n){"use strict";n.d(e,"c",(function(){return a})),n.d(e,"b",(function(){return o})),n.d(e,"a",(function(){return u}));var r=n(107);function i(t,e){return function(n){return t+n*e}}function a(t,e){var n=e-t;return n?i(t,n>180||n<-180?n-360*Math.round(n/360):n):Object(r.a)(isNaN(t)?e:t)}function o(t){return 1==(t=+t)?u:function(e,n){return n-e?function(t,e,n){return t=Math.pow(t,n),e=Math.pow(e,n)-t,n=1/n,function(r){return Math.pow(t+r*e,n)}}(e,n,t):Object(r.a)(isNaN(e)?n:e)}}function u(t,e){var n=e-t;return n?i(t,n):Object(r.a)(isNaN(t)?e:t)}},function(t,e){var n=t.exports={version:"2.6.11"};"number"==typeof __e&&(__e=n)},function(t,e,n){"use strict";n.d(e,"g",(function(){return o})),n.d(e,"c",(function(){return u})),n.d(e,"k",(function(){return s})),n.d(e,"m",(function(){return c})),n.d(e,"i",(function(){return l})),n.d(e,"a",(function(){return f})),n.d(e,"e",(function(){return d})),n.d(e,"h",(function(){return h})),n.d(e,"d",(function(){return p})),n.d(e,"l",(function(){return m})),n.d(e,"n",(function(){return v})),n.d(e,"j",(function(){return g})),n.d(e,"b",(function(){return _})),n.d(e,"f",(function(){return y}));var r=n(7),i=n(9);function a(t){return Object(r.a)((function(e){e.setDate(e.getDate()-(e.getDay()+7-t)%7),e.setHours(0,0,0,0)}),(function(t,e){t.setDate(t.getDate()+7*e)}),(function(t,e){return(e-t-(e.getTimezoneOffset()-t.getTimezoneOffset())*i.c)/i.e}))}var o=a(0),u=a(1),s=a(2),c=a(3),l=a(4),f=a(5),d=a(6),h=o.range,p=u.range,m=s.range,v=c.range,g=l.range,_=f.range,y=d.range},function(t,e,n){"use strict";n.d(e,"g",(function(){return o})),n.d(e,"c",(function(){return u})),n.d(e,"k",(function(){return s})),n.d(e,"m",(function(){return c})),n.d(e,"i",(function(){return l})),n.d(e,"a",(function(){return f})),n.d(e,"e",(function(){return d})),n.d(e,"h",(function(){return h})),n.d(e,"d",(function(){return p})),n.d(e,"l",(function(){return m})),n.d(e,"n",(function(){return v})),n.d(e,"j",(function(){return g})),n.d(e,"b",(function(){return _})),n.d(e,"f",(function(){return y}));var r=n(7),i=n(9);function a(t){return Object(r.a)((function(e){e.setUTCDate(e.getUTCDate()-(e.getUTCDay()+7-t)%7),e.setUTCHours(0,0,0,0)}),(function(t,e){t.setUTCDate(t.getUTCDate()+7*e)}),(function(t,e){return(e-t)/i.e}))}var o=a(0),u=a(1),s=a(2),c=a(3),l=a(4),f=a(5),d=a(6),h=o.range,p=u.range,m=s.range,v=c.range,g=l.range,_=f.range,y=d.range},function(t,e,n){"use strict";n.d(e,"a",(function(){return i})),n.d(e,"d",(function(){return a})),n.d(e,"c",(function(){return o})),n.d(e,"e",(function(){return b})),n.d(e,"h",(function(){return k})),n.d(e,"g",(function(){return M})),n.d(e,"b",(function(){return E})),n.d(e,"f",(function(){return L}));var r=n(26);function i(){}var a=.7,o=1/a,u="\\s*([+-]?\\d+)\\s*",s="\\s*([+-]?\\d*\\.?\\d+(?:[eE][+-]?\\d+)?)\\s*",c="\\s*([+-]?\\d*\\.?\\d+(?:[eE][+-]?\\d+)?)%\\s*",l=/^#([0-9a-f]{3,8})$/,f=new RegExp("^rgb\\("+[u,u,u]+"\\)$"),d=new RegExp("^rgb\\("+[c,c,c]+"\\)$"),h=new RegExp("^rgba\\("+[u,u,u,s]+"\\)$"),p=new RegExp("^rgba\\("+[c,c,c,s]+"\\)$"),m=new RegExp("^hsl\\("+[s,c,c]+"\\)$"),v=new RegExp("^hsla\\("+[s,c,c,s]+"\\)$"),g={aliceblue:15792383,antiquewhite:16444375,aqua:65535,aquamarine:8388564,azure:15794175,beige:16119260,bisque:16770244,black:0,blanchedalmond:16772045,blue:255,blueviolet:9055202,brown:10824234,burlywood:14596231,cadetblue:6266528,chartreuse:8388352,chocolate:13789470,coral:16744272,cornflowerblue:6591981,cornsilk:16775388,crimson:14423100,cyan:65535,darkblue:139,darkcyan:35723,darkgoldenrod:12092939,darkgray:11119017,darkgreen:25600,darkgrey:11119017,darkkhaki:12433259,darkmagenta:9109643,darkolivegreen:5597999,darkorange:16747520,darkorchid:10040012,darkred:9109504,darksalmon:15308410,darkseagreen:9419919,darkslateblue:4734347,darkslategray:3100495,darkslategrey:3100495,darkturquoise:52945,darkviolet:9699539,deeppink:16716947,deepskyblue:49151,dimgray:6908265,dimgrey:6908265,dodgerblue:2003199,firebrick:11674146,floralwhite:16775920,forestgreen:2263842,fuchsia:16711935,gainsboro:14474460,ghostwhite:16316671,gold:16766720,goldenrod:14329120,gray:8421504,green:32768,greenyellow:11403055,grey:8421504,honeydew:15794160,hotpink:16738740,indianred:13458524,indigo:4915330,ivory:16777200,khaki:15787660,lavender:15132410,lavenderblush:16773365,lawngreen:8190976,lemonchiffon:16775885,lightblue:11393254,lightcoral:15761536,lightcyan:14745599,lightgoldenrodyellow:16448210,lightgray:13882323,lightgreen:9498256,lightgrey:13882323,lightpink:16758465,lightsalmon:16752762,lightseagreen:2142890,lightskyblue:8900346,lightslategray:7833753,lightslategrey:7833753,lightsteelblue:11584734,lightyellow:16777184,lime:65280,limegreen:3329330,linen:16445670,magenta:16711935,maroon:8388608,mediumaquamarine:6737322,mediumblue:205,mediumorchid:12211667,mediumpurple:9662683,mediumseagreen:3978097,mediumslateblue:8087790,mediumspringgreen:64154,mediumturquoise:4772300,mediumvioletred:13047173,midnightblue:1644912,mintcream:16121850,mistyrose:16770273,moccasin:16770229,navajowhite:16768685,navy:128,oldlace:16643558,olive:8421376,olivedrab:7048739,orange:16753920,orangered:16729344,orchid:14315734,palegoldenrod:15657130,palegreen:10025880,paleturquoise:11529966,palevioletred:14381203,papayawhip:16773077,peachpuff:16767673,peru:13468991,pink:16761035,plum:14524637,powderblue:11591910,purple:8388736,rebeccapurple:6697881,red:16711680,rosybrown:12357519,royalblue:4286945,saddlebrown:9127187,salmon:16416882,sandybrown:16032864,seagreen:3050327,seashell:16774638,sienna:10506797,silver:12632256,skyblue:8900331,slateblue:6970061,slategray:7372944,slategrey:7372944,snow:16775930,springgreen:65407,steelblue:4620980,tan:13808780,teal:32896,thistle:14204888,tomato:16737095,turquoise:4251856,violet:15631086,wheat:16113331,white:16777215,whitesmoke:16119285,yellow:16776960,yellowgreen:10145074};function _(){return this.rgb().formatHex()}function y(){return this.rgb().formatRgb()}function b(t){var e,n;return t=(t+"").trim().toLowerCase(),(e=l.exec(t))?(n=e[1].length,e=parseInt(e[1],16),6===n?w(e):3===n?new E(e>>8&15|e>>4&240,e>>4&15|240&e,(15&e)<<4|15&e,1):8===n?x(e>>24&255,e>>16&255,e>>8&255,(255&e)/255):4===n?x(e>>12&15|e>>8&240,e>>8&15|e>>4&240,e>>4&15|240&e,((15&e)<<4|15&e)/255):null):(e=f.exec(t))?new E(e[1],e[2],e[3],1):(e=d.exec(t))?new E(255*e[1]/100,255*e[2]/100,255*e[3]/100,1):(e=h.exec(t))?x(e[1],e[2],e[3],e[4]):(e=p.exec(t))?x(255*e[1]/100,255*e[2]/100,255*e[3]/100,e[4]):(e=m.exec(t))?O(e[1],e[2]/100,e[3]/100,1):(e=v.exec(t))?O(e[1],e[2]/100,e[3]/100,e[4]):g.hasOwnProperty(t)?w(g[t]):"transparent"===t?new E(NaN,NaN,NaN,0):null}function w(t){return new E(t>>16&255,t>>8&255,255&t,1)}function x(t,e,n,r){return r<=0&&(t=e=n=NaN),new E(t,e,n,r)}function k(t){return t instanceof i||(t=b(t)),t?new E((t=t.rgb()).r,t.g,t.b,t.opacity):new E}function M(t,e,n,r){return 1===arguments.length?k(t):new E(t,e,n,null==r?1:r)}function E(t,e,n,r){this.r=+t,this.g=+e,this.b=+n,this.opacity=+r}function S(){return"#"+j(this.r)+j(this.g)+j(this.b)}function T(){var t=this.opacity;return(1===(t=isNaN(t)?1:Math.max(0,Math.min(1,t)))?"rgb(":"rgba(")+Math.max(0,Math.min(255,Math.round(this.r)||0))+", "+Math.max(0,Math.min(255,Math.round(this.g)||0))+", "+Math.max(0,Math.min(255,Math.round(this.b)||0))+(1===t?")":", "+t+")")}function j(t){return((t=Math.max(0,Math.min(255,Math.round(t)||0)))<16?"0":"")+t.toString(16)}function O(t,e,n,r){return r<=0?t=e=n=NaN:n<=0||n>=1?t=e=NaN:e<=0&&(t=NaN),new C(t,e,n,r)}function A(t){if(t instanceof C)return new C(t.h,t.s,t.l,t.opacity);if(t instanceof i||(t=b(t)),!t)return new C;if(t instanceof C)return t;var e=(t=t.rgb()).r/255,n=t.g/255,r=t.b/255,a=Math.min(e,n,r),o=Math.max(e,n,r),u=NaN,s=o-a,c=(o+a)/2;return s?(u=e===o?(n-r)/s+6*(n<r):n===o?(r-e)/s+2:(e-n)/s+4,s/=c<.5?o+a:2-o-a,u*=60):s=c>0&&c<1?0:u,new C(u,s,c,t.opacity)}function L(t,e,n,r){return 1===arguments.length?A(t):new C(t,e,n,null==r?1:r)}function C(t,e,n,r){this.h=+t,this.s=+e,this.l=+n,this.opacity=+r}function D(t,e,n){return 255*(t<60?e+(n-e)*t/60:t<180?n:t<240?e+(n-e)*(240-t)/60:e)}Object(r.a)(i,b,{copy:function(t){return Object.assign(new this.constructor,this,t)},displayable:function(){return this.rgb().displayable()},hex:_,formatHex:_,formatHsl:function(){return A(this).formatHsl()},formatRgb:y,toString:y}),Object(r.a)(E,M,Object(r.b)(i,{brighter:function(t){return t=null==t?o:Math.pow(o,t),new E(this.r*t,this.g*t,this.b*t,this.opacity)},darker:function(t){return t=null==t?a:Math.pow(a,t),new E(this.r*t,this.g*t,this.b*t,this.opacity)},rgb:function(){return this},displayable:function(){return-.5<=this.r&&this.r<255.5&&-.5<=this.g&&this.g<255.5&&-.5<=this.b&&this.b<255.5&&0<=this.opacity&&this.opacity<=1},hex:S,formatHex:S,formatRgb:T,toString:T})),Object(r.a)(C,L,Object(r.b)(i,{brighter:function(t){return t=null==t?o:Math.pow(o,t),new C(this.h,this.s,this.l*t,this.opacity)},darker:function(t){return t=null==t?a:Math.pow(a,t),new C(this.h,this.s,this.l*t,this.opacity)},rgb:function(){var t=this.h%360+360*(this.h<0),e=isNaN(t)||isNaN(this.s)?0:this.s,n=this.l,r=n+(n<.5?n:1-n)*e,i=2*n-r;return new E(D(t>=240?t-240:t+120,i,r),D(t,i,r),D(t<120?t+240:t-120,i,r),this.opacity)},displayable:function(){return(0<=this.s&&this.s<=1||isNaN(this.s))&&0<=this.l&&this.l<=1&&0<=this.opacity&&this.opacity<=1},formatHsl:function(){var t=this.opacity;return(1===(t=isNaN(t)?1:Math.max(0,Math.min(1,t)))?"hsl(":"hsla(")+(this.h||0)+", "+100*(this.s||0)+"%, "+100*(this.l||0)+"%"+(1===t?")":", "+t+")")}}))},function(t,e,n){"use strict";!function t(){if("undefined"!=typeof __REACT_DEVTOOLS_GLOBAL_HOOK__&&"function"==typeof __REACT_DEVTOOLS_GLOBAL_HOOK__.checkDCE){0;try{__REACT_DEVTOOLS_GLOBAL_HOOK__.checkDCE(t)}catch(t){console.error(t)}}}(),t.exports=n(516)},function(t,e,n){"use strict";n.d(e,"b",(function(){return a})),n.d(e,"c",(function(){return r.a})),n.d(e,"f",(function(){return u})),n.d(e,"g",(function(){return c.a})),n.d(e,"h",(function(){return l.a})),n.d(e,"i",(function(){return f.a})),n.d(e,"j",(function(){return d.a})),n.d(e,"a",(function(){return h.a})),n.d(e,"k",(function(){return i.a})),n.d(e,"l",(function(){return m})),n.d(e,"m",(function(){return p.b})),n.d(e,"n",(function(){return v.a})),n.d(e,"o",(function(){return g.a})),n.d(e,"p",(function(){return _.b})),n.d(e,"q",(function(){return y.a})),n.d(e,"r",(function(){return w})),n.d(e,"s",(function(){return x.a})),n.d(e,"e",(function(){return k.c})),n.d(e,"d",(function(){return k.a}));var r=n(59),i=n(106),a=function(t){return Object(i.a)(Object(r.a)(t).call(document.documentElement))},o=0;function u(){return new s}function s(){this._="@"+(++o).toString(36)}s.prototype=u.prototype={constructor:s,get:function(t){for(var e=this._;!(e in t);)if(!(t=t.parentNode))return;return t[e]},set:function(t,e){return t[this._]=e},remove:function(t){return this._ in t&&delete t[this._]},toString:function(){return this._}};var c=n(161),l=n(255),f=n(95),d=n(62),h=n(60),p=n(44),m=function(t){return"string"==typeof t?new p.a([document.querySelectorAll(t)],[document.documentElement]):new p.a([null==t?[]:t],p.c)},v=n(96),g=n(162),_=n(163),y=n(254),b=n(105),w=function(t,e){null==e&&(e=Object(b.a)().touches);for(var n=0,r=e?e.length:0,i=new Array(r);n<r;++n)i[n]=Object(h.a)(t,e[n]);return i},x=n(97),k=n(42)},function(t,e,n){var r=n(18),i=n(11),a=n(84),o=n(67),u=n(69),s=function(t,e,n){var c,l,f,d=t&s.F,h=t&s.G,p=t&s.S,m=t&s.P,v=t&s.B,g=t&s.W,_=h?i:i[e]||(i[e]={}),y=_.prototype,b=h?r:p?r[e]:(r[e]||{}).prototype;for(c in h&&(n=e),n)(l=!d&&b&&void 0!==b[c])&&u(_,c)||(f=l?b[c]:n[c],_[c]=h&&"function"!=typeof b[c]?n[c]:v&&l?a(f,r):g&&b[c]==f?function(t){var e=function(e,n,r){if(this instanceof t){switch(arguments.length){case 0:return new t;case 1:return new t(e);case 2:return new t(e,n)}return new t(e,n,r)}return t.apply(this,arguments)};return e.prototype=t.prototype,e}(f):m&&"function"==typeof f?a(Function.call,f):f,m&&((_.virtual||(_.virtual={}))[c]=f,t&s.R&&y&&!y[c]&&o(y,c,f)))};s.F=1,s.G=2,s.S=4,s.P=8,s.B=16,s.W=32,s.U=64,s.R=128,t.exports=s},function(t,e){var n=t.exports="undefined"!=typeof window&&window.Math==Math?window:"undefined"!=typeof self&&self.Math==Math?self:Function("return this")();"number"==typeof __g&&(__g=n)},function(t,e,n){"use strict";n.r(e),n.d(e,"default",(function(){return wn})),n.d(e,"VERSION",(function(){return i.e})),n.d(e,"restArguments",(function(){return a})),n.d(e,"isObject",(function(){return o})),n.d(e,"isNull",(function(){return u})),n.d(e,"isUndefined",(function(){return s})),n.d(e,"isBoolean",(function(){return c})),n.d(e,"isElement",(function(){return l})),n.d(e,"isString",(function(){return d})),n.d(e,"isNumber",(function(){return h})),n.d(e,"isDate",(function(){return p})),n.d(e,"isRegExp",(function(){return m})),n.d(e,"isError",(function(){return v})),n.d(e,"isSymbol",(function(){return g})),n.d(e,"isArrayBuffer",(function(){return _})),n.d(e,"isDataView",(function(){return S})),n.d(e,"isArray",(function(){return T})),n.d(e,"isFunction",(function(){return w})),n.d(e,"isArguments",(function(){return A})),n.d(e,"isFinite",(function(){return L})),n.d(e,"isNaN",(function(){return C})),n.d(e,"isTypedArray",(function(){return z})),n.d(e,"isEmpty",(function(){return q})),n.d(e,"isMatch",(function(){return U})),n.d(e,"isEqual",(function(){return $})),n.d(e,"isMap",(function(){return rt})),n.d(e,"isWeakMap",(function(){return it})),n.d(e,"isSet",(function(){return at})),n.d(e,"isWeakSet",(function(){return ot})),n.d(e,"keys",(function(){return B})),n.d(e,"allKeys",(function(){return X})),n.d(e,"values",(function(){return ut})),n.d(e,"pairs",(function(){return st})),n.d(e,"invert",(function(){return ct})),n.d(e,"functions",(function(){return lt})),n.d(e,"methods",(function(){return lt})),n.d(e,"extend",(function(){return dt})),n.d(e,"extendOwn",(function(){return ht})),n.d(e,"assign",(function(){return ht})),n.d(e,"defaults",(function(){return pt})),n.d(e,"create",(function(){return vt})),n.d(e,"clone",(function(){return gt})),n.d(e,"tap",(function(){return _t})),n.d(e,"get",(function(){return xt})),n.d(e,"has",(function(){return kt})),n.d(e,"mapObject",(function(){return Lt})),n.d(e,"identity",(function(){return Mt})),n.d(e,"constant",(function(){return D})),n.d(e,"noop",(function(){return Ct})),n.d(e,"toPath",(function(){return yt})),n.d(e,"property",(function(){return St})),n.d(e,"propertyOf",(function(){return Dt})),n.d(e,"matcher",(function(){return Et})),n.d(e,"matches",(function(){return Et})),n.d(e,"times",(function(){return Nt})),n.d(e,"random",(function(){return Pt})),n.d(e,"now",(function(){return Rt})),n.d(e,"escape",(function(){return zt})),n.d(e,"unescape",(function(){return Ft})),n.d(e,"templateSettings",(function(){return Ht})),n.d(e,"template",(function(){return Gt})),n.d(e,"result",(function(){return Vt})),n.d(e,"uniqueId",(function(){return Xt})),n.d(e,"chain",(function(){return Jt})),n.d(e,"iteratee",(function(){return Ot})),n.d(e,"partial",(function(){return Qt})),n.d(e,"bind",(function(){return te})),n.d(e,"bindAll",(function(){return re})),n.d(e,"memoize",(function(){return ie})),n.d(e,"delay",(function(){return ae})),n.d(e,"defer",(function(){return oe})),n.d(e,"throttle",(function(){return ue})),n.d(e,"debounce",(function(){return se})),n.d(e,"wrap",(function(){return ce})),n.d(e,"negate",(function(){return le})),n.d(e,"compose",(function(){return fe})),n.d(e,"after",(function(){return de})),n.d(e,"before",(function(){return he})),n.d(e,"once",(function(){return pe})),n.d(e,"findKey",(function(){return me})),n.d(e,"findIndex",(function(){return ge})),n.d(e,"findLastIndex",(function(){return _e})),n.d(e,"sortedIndex",(function(){return ye})),n.d(e,"indexOf",(function(){return we})),n.d(e,"lastIndexOf",(function(){return xe})),n.d(e,"find",(function(){return ke})),n.d(e,"detect",(function(){return ke})),n.d(e,"findWhere",(function(){return Me})),n.d(e,"each",(function(){return Ee})),n.d(e,"forEach",(function(){return Ee})),n.d(e,"map",(function(){return Se})),n.d(e,"collect",(function(){return Se})),n.d(e,"reduce",(function(){return je})),n.d(e,"foldl",(function(){return je})),n.d(e,"inject",(function(){return je})),n.d(e,"reduceRight",(function(){return Oe})),n.d(e,"foldr",(function(){return Oe})),n.d(e,"filter",(function(){return Ae})),n.d(e,"select",(function(){return Ae})),n.d(e,"reject",(function(){return Le})),n.d(e,"every",(function(){return Ce})),n.d(e,"all",(function(){return Ce})),n.d(e,"some",(function(){return De})),n.d(e,"any",(function(){return De})),n.d(e,"contains",(function(){return Ne})),n.d(e,"includes",(function(){return Ne})),n.d(e,"include",(function(){return Ne})),n.d(e,"invoke",(function(){return Pe})),n.d(e,"pluck",(function(){return Re})),n.d(e,"where",(function(){return Ye})),n.d(e,"max",(function(){return Ie})),n.d(e,"min",(function(){return ze})),n.d(e,"shuffle",(function(){return He})),n.d(e,"sample",(function(){return Fe})),n.d(e,"sortBy",(function(){return Be})),n.d(e,"groupBy",(function(){return Ue})),n.d(e,"indexBy",(function(){return We})),n.d(e,"countBy",(function(){return Ge})),n.d(e,"partition",(function(){return Ve})),n.d(e,"toArray",(function(){return Xe})),n.d(e,"size",(function(){return Je})),n.d(e,"pick",(function(){return Ke})),n.d(e,"omit",(function(){return Qe})),n.d(e,"first",(function(){return en})),n.d(e,"head",(function(){return en})),n.d(e,"take",(function(){return en})),n.d(e,"initial",(function(){return tn})),n.d(e,"last",(function(){return rn})),n.d(e,"rest",(function(){return nn})),n.d(e,"tail",(function(){return nn})),n.d(e,"drop",(function(){return nn})),n.d(e,"compact",(function(){return an})),n.d(e,"flatten",(function(){return on})),n.d(e,"without",(function(){return sn})),n.d(e,"uniq",(function(){return cn})),n.d(e,"unique",(function(){return cn})),n.d(e,"union",(function(){return ln})),n.d(e,"intersection",(function(){return fn})),n.d(e,"difference",(function(){return un})),n.d(e,"unzip",(function(){return dn})),n.d(e,"transpose",(function(){return dn})),n.d(e,"zip",(function(){return hn})),n.d(e,"object",(function(){return pn})),n.d(e,"range",(function(){return mn})),n.d(e,"chunk",(function(){return vn})),n.d(e,"mixin",(function(){return _n}));var r={};n.r(r),n.d(r,"VERSION",(function(){return i.e})),n.d(r,"restArguments",(function(){return a})),n.d(r,"isObject",(function(){return o})),n.d(r,"isNull",(function(){return u})),n.d(r,"isUndefined",(function(){return s})),n.d(r,"isBoolean",(function(){return c})),n.d(r,"isElement",(function(){return l})),n.d(r,"isString",(function(){return d})),n.d(r,"isNumber",(function(){return h})),n.d(r,"isDate",(function(){return p})),n.d(r,"isRegExp",(function(){return m})),n.d(r,"isError",(function(){return v})),n.d(r,"isSymbol",(function(){return g})),n.d(r,"isArrayBuffer",(function(){return _})),n.d(r,"isDataView",(function(){return S})),n.d(r,"isArray",(function(){return T})),n.d(r,"isFunction",(function(){return w})),n.d(r,"isArguments",(function(){return A})),n.d(r,"isFinite",(function(){return L})),n.d(r,"isNaN",(function(){return C})),n.d(r,"isTypedArray",(function(){return z})),n.d(r,"isEmpty",(function(){return q})),n.d(r,"isMatch",(function(){return U})),n.d(r,"isEqual",(function(){return $})),n.d(r,"isMap",(function(){return rt})),n.d(r,"isWeakMap",(function(){return it})),n.d(r,"isSet",(function(){return at})),n.d(r,"isWeakSet",(function(){return ot})),n.d(r,"keys",(function(){return B})),n.d(r,"allKeys",(function(){return X})),n.d(r,"values",(function(){return ut})),n.d(r,"pairs",(function(){return st})),n.d(r,"invert",(function(){return ct})),n.d(r,"functions",(function(){return lt})),n.d(r,"methods",(function(){return lt})),n.d(r,"extend",(function(){return dt})),n.d(r,"extendOwn",(function(){return ht})),n.d(r,"assign",(function(){return ht})),n.d(r,"defaults",(function(){return pt})),n.d(r,"create",(function(){return vt})),n.d(r,"clone",(function(){return gt})),n.d(r,"tap",(function(){return _t})),n.d(r,"get",(function(){return xt})),n.d(r,"has",(function(){return kt})),n.d(r,"mapObject",(function(){return Lt})),n.d(r,"identity",(function(){return Mt})),n.d(r,"constant",(function(){return D})),n.d(r,"noop",(function(){return Ct})),n.d(r,"toPath",(function(){return yt})),n.d(r,"property",(function(){return St})),n.d(r,"propertyOf",(function(){return Dt})),n.d(r,"matcher",(function(){return Et})),n.d(r,"matches",(function(){return Et})),n.d(r,"times",(function(){return Nt})),n.d(r,"random",(function(){return Pt})),n.d(r,"now",(function(){return Rt})),n.d(r,"escape",(function(){return zt})),n.d(r,"unescape",(function(){return Ft})),n.d(r,"templateSettings",(function(){return Ht})),n.d(r,"template",(function(){return Gt})),n.d(r,"result",(function(){return Vt})),n.d(r,"uniqueId",(function(){return Xt})),n.d(r,"chain",(function(){return Jt})),n.d(r,"iteratee",(function(){return Ot})),n.d(r,"partial",(function(){return Qt})),n.d(r,"bind",(function(){return te})),n.d(r,"bindAll",(function(){return re})),n.d(r,"memoize",(function(){return ie})),n.d(r,"delay",(function(){return ae})),n.d(r,"defer",(function(){return oe})),n.d(r,"throttle",(function(){return ue})),n.d(r,"debounce",(function(){return se})),n.d(r,"wrap",(function(){return ce})),n.d(r,"negate",(function(){return le})),n.d(r,"compose",(function(){return fe})),n.d(r,"after",(function(){return de})),n.d(r,"before",(function(){return he})),n.d(r,"once",(function(){return pe})),n.d(r,"findKey",(function(){return me})),n.d(r,"findIndex",(function(){return ge})),n.d(r,"findLastIndex",(function(){return _e})),n.d(r,"sortedIndex",(function(){return ye})),n.d(r,"indexOf",(function(){return we})),n.d(r,"lastIndexOf",(function(){return xe})),n.d(r,"find",(function(){return ke})),n.d(r,"detect",(function(){return ke})),n.d(r,"findWhere",(function(){return Me})),n.d(r,"each",(function(){return Ee})),n.d(r,"forEach",(function(){return Ee})),n.d(r,"map",(function(){return Se})),n.d(r,"collect",(function(){return Se})),n.d(r,"reduce",(function(){return je})),n.d(r,"foldl",(function(){return je})),n.d(r,"inject",(function(){return je})),n.d(r,"reduceRight",(function(){return Oe})),n.d(r,"foldr",(function(){return Oe})),n.d(r,"filter",(function(){return Ae})),n.d(r,"select",(function(){return Ae})),n.d(r,"reject",(function(){return Le})),n.d(r,"every",(function(){return Ce})),n.d(r,"all",(function(){return Ce})),n.d(r,"some",(function(){return De})),n.d(r,"any",(function(){return De})),n.d(r,"contains",(function(){return Ne})),n.d(r,"includes",(function(){return Ne})),n.d(r,"include",(function(){return Ne})),n.d(r,"invoke",(function(){return Pe})),n.d(r,"pluck",(function(){return Re})),n.d(r,"where",(function(){return Ye})),n.d(r,"max",(function(){return Ie})),n.d(r,"min",(function(){return ze})),n.d(r,"shuffle",(function(){return He})),n.d(r,"sample",(function(){return Fe})),n.d(r,"sortBy",(function(){return Be})),n.d(r,"groupBy",(function(){return Ue})),n.d(r,"indexBy",(function(){return We})),n.d(r,"countBy",(function(){return Ge})),n.d(r,"partition",(function(){return Ve})),n.d(r,"toArray",(function(){return Xe})),n.d(r,"size",(function(){return Je})),n.d(r,"pick",(function(){return Ke})),n.d(r,"omit",(function(){return Qe})),n.d(r,"first",(function(){return en})),n.d(r,"head",(function(){return en})),n.d(r,"take",(function(){return en})),n.d(r,"initial",(function(){return tn})),n.d(r,"last",(function(){return rn})),n.d(r,"rest",(function(){return nn})),n.d(r,"tail",(function(){return nn})),n.d(r,"drop",(function(){return nn})),n.d(r,"compact",(function(){return an})),n.d(r,"flatten",(function(){return on})),n.d(r,"without",(function(){return sn})),n.d(r,"uniq",(function(){return cn})),n.d(r,"unique",(function(){return cn})),n.d(r,"union",(function(){return ln})),n.d(r,"intersection",(function(){return fn})),n.d(r,"difference",(function(){return un})),n.d(r,"unzip",(function(){return dn})),n.d(r,"transpose",(function(){return dn})),n.d(r,"zip",(function(){return hn})),n.d(r,"object",(function(){return pn})),n.d(r,"range",(function(){return mn})),n.d(r,"chunk",(function(){return vn})),n.d(r,"mixin",(function(){return _n})),n.d(r,"default",(function(){return yn}));var i=n(4);function a(t,e){return e=null==e?t.length-1:+e,function(){for(var n=Math.max(arguments.length-e,0),r=Array(n),i=0;i<n;i++)r[i]=arguments[i+e];switch(e){case 0:return t.call(this,r);case 1:return t.call(this,arguments[0],r);case 2:return t.call(this,arguments[0],arguments[1],r)}var a=Array(e+1);for(i=0;i<e;i++)a[i]=arguments[i];return a[e]=r,t.apply(this,a)}}function o(t){var e=typeof t;return"function"===e||"object"===e&&!!t}function u(t){return null===t}function s(t){return void 0===t}function c(t){return!0===t||!1===t||"[object Boolean]"===i.t.call(t)}function l(t){return!(!t||1!==t.nodeType)}function f(t){var e="[object "+t+"]";return function(t){return i.t.call(t)===e}}var d=f("String"),h=f("Number"),p=f("Date"),m=f("RegExp"),v=f("Error"),g=f("Symbol"),_=f("ArrayBuffer"),y=f("Function"),b=i.p.document&&i.p.document.childNodes;"object"!=typeof Int8Array&&"function"!=typeof b&&(y=function(t){return"function"==typeof t||!1});var w=y,x=f("Object"),k=i.s&&x(new DataView(new ArrayBuffer(8))),M="undefined"!=typeof Map&&x(new Map),E=f("DataView");var S=k?function(t){return null!=t&&w(t.getInt8)&&_(t.buffer)}:E,T=i.k||f("Array");function j(t,e){return null!=t&&i.i.call(t,e)}var O=f("Arguments");!function(){O(arguments)||(O=function(t){return j(t,"callee")})}();var A=O;function L(t){return!g(t)&&Object(i.f)(t)&&!isNaN(parseFloat(t))}function C(t){return h(t)&&Object(i.g)(t)}function D(t){return function(){return t}}function N(t){return function(e){var n=t(e);return"number"==typeof n&&n>=0&&n<=i.b}}function P(t){return function(e){return null==e?void 0:e[t]}}var R=P("byteLength"),Y=N(R),I=/\[object ((I|Ui)nt(8|16|32)|Float(32|64)|Uint8Clamped|Big(I|Ui)nt64)Array\]/;var z=i.r?function(t){return i.l?Object(i.l)(t)&&!S(t):Y(t)&&I.test(i.t.call(t))}:D(!1),F=P("length");function H(t,e){e=function(t){for(var e={},n=t.length,r=0;r<n;++r)e[t[r]]=!0;return{contains:function(t){return e[t]},push:function(n){return e[n]=!0,t.push(n)}}}(e);var n=i.n.length,r=t.constructor,a=w(r)&&r.prototype||i.c,o="constructor";for(j(t,o)&&!e.contains(o)&&e.push(o);n--;)(o=i.n[n])in t&&t[o]!==a[o]&&!e.contains(o)&&e.push(o)}function B(t){if(!o(t))return[];if(i.m)return Object(i.m)(t);var e=[];for(var n in t)j(t,n)&&e.push(n);return i.h&&H(t,e),e}function q(t){if(null==t)return!0;var e=F(t);return"number"==typeof e&&(T(t)||d(t)||A(t))?0===e:0===F(B(t))}function U(t,e){var n=B(e),r=n.length;if(null==t)return!r;for(var i=Object(t),a=0;a<r;a++){var o=n[a];if(e[o]!==i[o]||!(o in i))return!1}return!0}function W(t){return t instanceof W?t:this instanceof W?void(this._wrapped=t):new W(t)}function G(t){return new Uint8Array(t.buffer||t,t.byteOffset||0,R(t))}W.VERSION=i.e,W.prototype.value=function(){return this._wrapped},W.prototype.valueOf=W.prototype.toJSON=W.prototype.value,W.prototype.toString=function(){return String(this._wrapped)};function V(t,e,n,r){if(t===e)return 0!==t||1/t==1/e;if(null==t||null==e)return!1;if(t!=t)return e!=e;var a=typeof t;return("function"===a||"object"===a||"object"==typeof e)&&function t(e,n,r,a){e instanceof W&&(e=e._wrapped);n instanceof W&&(n=n._wrapped);var o=i.t.call(e);if(o!==i.t.call(n))return!1;if(k&&"[object Object]"==o&&S(e)){if(!S(n))return!1;o="[object DataView]"}switch(o){case"[object RegExp]":case"[object String]":return""+e==""+n;case"[object Number]":return+e!=+e?+n!=+n:0==+e?1/+e==1/n:+e==+n;case"[object Date]":case"[object Boolean]":return+e==+n;case"[object Symbol]":return i.d.valueOf.call(e)===i.d.valueOf.call(n);case"[object ArrayBuffer]":case"[object DataView]":return t(G(e),G(n),r,a)}var u="[object Array]"===o;if(!u&&z(e)){if(R(e)!==R(n))return!1;if(e.buffer===n.buffer&&e.byteOffset===n.byteOffset)return!0;u=!0}if(!u){if("object"!=typeof e||"object"!=typeof n)return!1;var s=e.constructor,c=n.constructor;if(s!==c&&!(w(s)&&s instanceof s&&w(c)&&c instanceof c)&&"constructor"in e&&"constructor"in n)return!1}a=a||[];var l=(r=r||[]).length;for(;l--;)if(r[l]===e)return a[l]===n;if(r.push(e),a.push(n),u){if((l=e.length)!==n.length)return!1;for(;l--;)if(!V(e[l],n[l],r,a))return!1}else{var f,d=B(e);if(l=d.length,B(n).length!==l)return!1;for(;l--;)if(f=d[l],!j(n,f)||!V(e[f],n[f],r,a))return!1}return r.pop(),a.pop(),!0}(t,e,n,r)}function $(t,e){return V(t,e)}function X(t){if(!o(t))return[];var e=[];for(var n in t)e.push(n);return i.h&&H(t,e),e}function J(t){var e=F(t);return function(n){if(null==n)return!1;var r=X(n);if(F(r))return!1;for(var i=0;i<e;i++)if(!w(n[t[i]]))return!1;return t!==et||!w(n[Z])}}var Z="forEach",K=["clear","delete"],Q=["get","has","set"],tt=K.concat(Z,Q),et=K.concat(Q),nt=["add"].concat(K,Z,"has"),rt=M?J(tt):f("Map"),it=M?J(et):f("WeakMap"),at=M?J(nt):f("Set"),ot=f("WeakSet");function ut(t){for(var e=B(t),n=e.length,r=Array(n),i=0;i<n;i++)r[i]=t[e[i]];return r}function st(t){for(var e=B(t),n=e.length,r=Array(n),i=0;i<n;i++)r[i]=[e[i],t[e[i]]];return r}function ct(t){for(var e={},n=B(t),r=0,i=n.length;r<i;r++)e[t[n[r]]]=n[r];return e}function lt(t){var e=[];for(var n in t)w(t[n])&&e.push(n);return e.sort()}function ft(t,e){return function(n){var r=arguments.length;if(e&&(n=Object(n)),r<2||null==n)return n;for(var i=1;i<r;i++)for(var a=arguments[i],o=t(a),u=o.length,s=0;s<u;s++){var c=o[s];e&&void 0!==n[c]||(n[c]=a[c])}return n}}var dt=ft(X),ht=ft(B),pt=ft(X,!0);function mt(t){if(!o(t))return{};if(i.j)return Object(i.j)(t);var e=function(){};e.prototype=t;var n=new e;return e.prototype=null,n}function vt(t,e){var n=mt(t);return e&&ht(n,e),n}function gt(t){return o(t)?T(t)?t.slice():dt({},t):t}function _t(t,e){return e(t),t}function yt(t){return T(t)?t:[t]}function bt(t){return W.toPath(t)}function wt(t,e){for(var n=e.length,r=0;r<n;r++){if(null==t)return;t=t[e[r]]}return n?t:void 0}function xt(t,e,n){var r=wt(t,bt(e));return s(r)?n:r}function kt(t,e){for(var n=(e=bt(e)).length,r=0;r<n;r++){var i=e[r];if(!j(t,i))return!1;t=t[i]}return!!n}function Mt(t){return t}function Et(t){return t=ht({},t),function(e){return U(e,t)}}function St(t){return t=bt(t),function(e){return wt(e,t)}}function Tt(t,e,n){if(void 0===e)return t;switch(null==n?3:n){case 1:return function(n){return t.call(e,n)};case 3:return function(n,r,i){return t.call(e,n,r,i)};case 4:return function(n,r,i,a){return t.call(e,n,r,i,a)}}return function(){return t.apply(e,arguments)}}function jt(t,e,n){return null==t?Mt:w(t)?Tt(t,e,n):o(t)&&!T(t)?Et(t):St(t)}function Ot(t,e){return jt(t,e,1/0)}function At(t,e,n){return W.iteratee!==Ot?W.iteratee(t,e):jt(t,e,n)}function Lt(t,e,n){e=At(e,n);for(var r=B(t),i=r.length,a={},o=0;o<i;o++){var u=r[o];a[u]=e(t[u],u,t)}return a}function Ct(){}function Dt(t){return null==t?Ct:function(e){return xt(t,e)}}function Nt(t,e,n){var r=Array(Math.max(0,t));e=Tt(e,n,1);for(var i=0;i<t;i++)r[i]=e(i);return r}function Pt(t,e){return null==e&&(e=t,t=0),t+Math.floor(Math.random()*(e-t+1))}W.toPath=yt,W.iteratee=Ot;var Rt=Date.now||function(){return(new Date).getTime()};function Yt(t){var e=function(e){return t[e]},n="(?:"+B(t).join("|")+")",r=RegExp(n),i=RegExp(n,"g");return function(t){return t=null==t?"":""+t,r.test(t)?t.replace(i,e):t}}var It={"&":"&amp;","<":"&lt;",">":"&gt;",'"':"&quot;","'":"&#x27;","`":"&#x60;"},zt=Yt(It),Ft=Yt(ct(It)),Ht=W.templateSettings={evaluate:/<%([\s\S]+?)%>/g,interpolate:/<%=([\s\S]+?)%>/g,escape:/<%-([\s\S]+?)%>/g},Bt=/(.)^/,qt={"'":"'","\\":"\\","\r":"r","\n":"n","\u2028":"u2028","\u2029":"u2029"},Ut=/\\|'|\r|\n|\u2028|\u2029/g;function Wt(t){return"\\"+qt[t]}function Gt(t,e,n){!e&&n&&(e=n),e=pt({},e,W.templateSettings);var r,i=RegExp([(e.escape||Bt).source,(e.interpolate||Bt).source,(e.evaluate||Bt).source].join("|")+"|$","g"),a=0,o="__p+='";t.replace(i,(function(e,n,r,i,u){return o+=t.slice(a,u).replace(Ut,Wt),a=u+e.length,n?o+="'+\n((__t=("+n+"))==null?'':_.escape(__t))+\n'":r?o+="'+\n((__t=("+r+"))==null?'':__t)+\n'":i&&(o+="';\n"+i+"\n__p+='"),e})),o+="';\n",e.variable||(o="with(obj||{}){\n"+o+"}\n"),o="var __t,__p='',__j=Array.prototype.join,print=function(){__p+=__j.call(arguments,'');};\n"+o+"return __p;\n";try{r=new Function(e.variable||"obj","_",o)}catch(t){throw t.source=o,t}var u=function(t){return r.call(this,t,W)},s=e.variable||"obj";return u.source="function("+s+"){\n"+o+"}",u}function Vt(t,e,n){var r=(e=bt(e)).length;if(!r)return w(n)?n.call(t):n;for(var i=0;i<r;i++){var a=null==t?void 0:t[e[i]];void 0===a&&(a=n,i=r),t=w(a)?a.call(t):a}return t}var $t=0;function Xt(t){var e=++$t+"";return t?t+e:e}function Jt(t){var e=W(t);return e._chain=!0,e}function Zt(t,e,n,r,i){if(!(r instanceof e))return t.apply(n,i);var a=mt(t.prototype),u=t.apply(a,i);return o(u)?u:a}var Kt=a((function(t,e){var n=Kt.placeholder,r=function(){for(var i=0,a=e.length,o=Array(a),u=0;u<a;u++)o[u]=e[u]===n?arguments[i++]:e[u];for(;i<arguments.length;)o.push(arguments[i++]);return Zt(t,r,this,this,o)};return r}));Kt.placeholder=W;var Qt=Kt,te=a((function(t,e,n){if(!w(t))throw new TypeError("Bind must be called on a function");var r=a((function(i){return Zt(t,r,e,this,n.concat(i))}));return r})),ee=N(F);function ne(t,e,n,r){if(r=r||[],e||0===e){if(e<=0)return r.concat(t)}else e=1/0;for(var i=r.length,a=0,o=F(t);a<o;a++){var u=t[a];if(ee(u)&&(T(u)||A(u)))if(e>1)ne(u,e-1,n,r),i=r.length;else for(var s=0,c=u.length;s<c;)r[i++]=u[s++];else n||(r[i++]=u)}return r}var re=a((function(t,e){var n=(e=ne(e,!1,!1)).length;if(n<1)throw new Error("bindAll must be passed function names");for(;n--;){var r=e[n];t[r]=te(t[r],t)}return t}));function ie(t,e){var n=function(r){var i=n.cache,a=""+(e?e.apply(this,arguments):r);return j(i,a)||(i[a]=t.apply(this,arguments)),i[a]};return n.cache={},n}var ae=a((function(t,e,n){return setTimeout((function(){return t.apply(null,n)}),e)})),oe=Qt(ae,W,1);function ue(t,e,n){var r,i,a,o,u=0;n||(n={});var s=function(){u=!1===n.leading?0:Rt(),r=null,o=t.apply(i,a),r||(i=a=null)},c=function(){var c=Rt();u||!1!==n.leading||(u=c);var l=e-(c-u);return i=this,a=arguments,l<=0||l>e?(r&&(clearTimeout(r),r=null),u=c,o=t.apply(i,a),r||(i=a=null)):r||!1===n.trailing||(r=setTimeout(s,l)),o};return c.cancel=function(){clearTimeout(r),u=0,r=i=a=null},c}function se(t,e,n){var r,i,o=function(e,n){r=null,n&&(i=t.apply(e,n))},u=a((function(a){if(r&&clearTimeout(r),n){var u=!r;r=setTimeout(o,e),u&&(i=t.apply(this,a))}else r=ae(o,e,this,a);return i}));return u.cancel=function(){clearTimeout(r),r=null},u}function ce(t,e){return Qt(e,t)}function le(t){return function(){return!t.apply(this,arguments)}}function fe(){var t=arguments,e=t.length-1;return function(){for(var n=e,r=t[e].apply(this,arguments);n--;)r=t[n].call(this,r);return r}}function de(t,e){return function(){if(--t<1)return e.apply(this,arguments)}}function he(t,e){var n;return function(){return--t>0&&(n=e.apply(this,arguments)),t<=1&&(e=null),n}}var pe=Qt(he,2);function me(t,e,n){e=At(e,n);for(var r,i=B(t),a=0,o=i.length;a<o;a++)if(e(t[r=i[a]],r,t))return r}function ve(t){return function(e,n,r){n=At(n,r);for(var i=F(e),a=t>0?0:i-1;a>=0&&a<i;a+=t)if(n(e[a],a,e))return a;return-1}}var ge=ve(1),_e=ve(-1);function ye(t,e,n,r){for(var i=(n=At(n,r,1))(e),a=0,o=F(t);a<o;){var u=Math.floor((a+o)/2);n(t[u])<i?a=u+1:o=u}return a}function be(t,e,n){return function(r,a,o){var u=0,s=F(r);if("number"==typeof o)t>0?u=o>=0?o:Math.max(o+s,u):s=o>=0?Math.min(o+1,s):o+s+1;else if(n&&o&&s)return r[o=n(r,a)]===a?o:-1;if(a!=a)return(o=e(i.q.call(r,u,s),C))>=0?o+u:-1;for(o=t>0?u:s-1;o>=0&&o<s;o+=t)if(r[o]===a)return o;return-1}}var we=be(1,ge,ye),xe=be(-1,_e);function ke(t,e,n){var r=(ee(t)?ge:me)(t,e,n);if(void 0!==r&&-1!==r)return t[r]}function Me(t,e){return ke(t,Et(e))}function Ee(t,e,n){var r,i;if(e=Tt(e,n),ee(t))for(r=0,i=t.length;r<i;r++)e(t[r],r,t);else{var a=B(t);for(r=0,i=a.length;r<i;r++)e(t[a[r]],a[r],t)}return t}function Se(t,e,n){e=At(e,n);for(var r=!ee(t)&&B(t),i=(r||t).length,a=Array(i),o=0;o<i;o++){var u=r?r[o]:o;a[o]=e(t[u],u,t)}return a}function Te(t){var e=function(e,n,r,i){var a=!ee(e)&&B(e),o=(a||e).length,u=t>0?0:o-1;for(i||(r=e[a?a[u]:u],u+=t);u>=0&&u<o;u+=t){var s=a?a[u]:u;r=n(r,e[s],s,e)}return r};return function(t,n,r,i){var a=arguments.length>=3;return e(t,Tt(n,i,4),r,a)}}var je=Te(1),Oe=Te(-1);function Ae(t,e,n){var r=[];return e=At(e,n),Ee(t,(function(t,n,i){e(t,n,i)&&r.push(t)})),r}function Le(t,e,n){return Ae(t,le(At(e)),n)}function Ce(t,e,n){e=At(e,n);for(var r=!ee(t)&&B(t),i=(r||t).length,a=0;a<i;a++){var o=r?r[a]:a;if(!e(t[o],o,t))return!1}return!0}function De(t,e,n){e=At(e,n);for(var r=!ee(t)&&B(t),i=(r||t).length,a=0;a<i;a++){var o=r?r[a]:a;if(e(t[o],o,t))return!0}return!1}function Ne(t,e,n,r){return ee(t)||(t=ut(t)),("number"!=typeof n||r)&&(n=0),we(t,e,n)>=0}var Pe=a((function(t,e,n){var r,i;return w(e)?i=e:(e=bt(e),r=e.slice(0,-1),e=e[e.length-1]),Se(t,(function(t){var a=i;if(!a){if(r&&r.length&&(t=wt(t,r)),null==t)return;a=t[e]}return null==a?a:a.apply(t,n)}))}));function Re(t,e){return Se(t,St(e))}function Ye(t,e){return Ae(t,Et(e))}function Ie(t,e,n){var r,i,a=-1/0,o=-1/0;if(null==e||"number"==typeof e&&"object"!=typeof t[0]&&null!=t)for(var u=0,s=(t=ee(t)?t:ut(t)).length;u<s;u++)null!=(r=t[u])&&r>a&&(a=r);else e=At(e,n),Ee(t,(function(t,n,r){((i=e(t,n,r))>o||i===-1/0&&a===-1/0)&&(a=t,o=i)}));return a}function ze(t,e,n){var r,i,a=1/0,o=1/0;if(null==e||"number"==typeof e&&"object"!=typeof t[0]&&null!=t)for(var u=0,s=(t=ee(t)?t:ut(t)).length;u<s;u++)null!=(r=t[u])&&r<a&&(a=r);else e=At(e,n),Ee(t,(function(t,n,r){((i=e(t,n,r))<o||i===1/0&&a===1/0)&&(a=t,o=i)}));return a}function Fe(t,e,n){if(null==e||n)return ee(t)||(t=ut(t)),t[Pt(t.length-1)];var r=ee(t)?gt(t):ut(t),i=F(r);e=Math.max(Math.min(e,i),0);for(var a=i-1,o=0;o<e;o++){var u=Pt(o,a),s=r[o];r[o]=r[u],r[u]=s}return r.slice(0,e)}function He(t){return Fe(t,1/0)}function Be(t,e,n){var r=0;return e=At(e,n),Re(Se(t,(function(t,n,i){return{value:t,index:r++,criteria:e(t,n,i)}})).sort((function(t,e){var n=t.criteria,r=e.criteria;if(n!==r){if(n>r||void 0===n)return 1;if(n<r||void 0===r)return-1}return t.index-e.index})),"value")}function qe(t,e){return function(n,r,i){var a=e?[[],[]]:{};return r=At(r,i),Ee(n,(function(e,i){var o=r(e,i,n);t(a,e,o)})),a}}var Ue=qe((function(t,e,n){j(t,n)?t[n].push(e):t[n]=[e]})),We=qe((function(t,e,n){t[n]=e})),Ge=qe((function(t,e,n){j(t,n)?t[n]++:t[n]=1})),Ve=qe((function(t,e,n){t[n?0:1].push(e)}),!0),$e=/[^\ud800-\udfff]|[\ud800-\udbff][\udc00-\udfff]|[\ud800-\udfff]/g;function Xe(t){return t?T(t)?i.q.call(t):d(t)?t.match($e):ee(t)?Se(t,Mt):ut(t):[]}function Je(t){return null==t?0:ee(t)?t.length:B(t).length}function Ze(t,e,n){return e in n}var Ke=a((function(t,e){var n={},r=e[0];if(null==t)return n;w(r)?(e.length>1&&(r=Tt(r,e[1])),e=X(t)):(r=Ze,e=ne(e,!1,!1),t=Object(t));for(var i=0,a=e.length;i<a;i++){var o=e[i],u=t[o];r(u,o,t)&&(n[o]=u)}return n})),Qe=a((function(t,e){var n,r=e[0];return w(r)?(r=le(r),e.length>1&&(n=e[1])):(e=Se(ne(e,!1,!1),String),r=function(t,n){return!Ne(e,n)}),Ke(t,r,n)}));function tn(t,e,n){return i.q.call(t,0,Math.max(0,t.length-(null==e||n?1:e)))}function en(t,e,n){return null==t||t.length<1?null==e||n?void 0:[]:null==e||n?t[0]:tn(t,t.length-e)}function nn(t,e,n){return i.q.call(t,null==e||n?1:e)}function rn(t,e,n){return null==t||t.length<1?null==e||n?void 0:[]:null==e||n?t[t.length-1]:nn(t,Math.max(0,t.length-e))}function an(t){return Ae(t,Boolean)}function on(t,e){return ne(t,e,!1)}var un=a((function(t,e){return e=ne(e,!0,!0),Ae(t,(function(t){return!Ne(e,t)}))})),sn=a((function(t,e){return un(t,e)}));function cn(t,e,n,r){c(e)||(r=n,n=e,e=!1),null!=n&&(n=At(n,r));for(var i=[],a=[],o=0,u=F(t);o<u;o++){var s=t[o],l=n?n(s,o,t):s;e&&!n?(o&&a===l||i.push(s),a=l):n?Ne(a,l)||(a.push(l),i.push(s)):Ne(i,s)||i.push(s)}return i}var ln=a((function(t){return cn(ne(t,!0,!0))}));function fn(t){for(var e=[],n=arguments.length,r=0,i=F(t);r<i;r++){var a=t[r];if(!Ne(e,a)){var o;for(o=1;o<n&&Ne(arguments[o],a);o++);o===n&&e.push(a)}}return e}function dn(t){for(var e=t&&Ie(t,F).length||0,n=Array(e),r=0;r<e;r++)n[r]=Re(t,r);return n}var hn=a(dn);function pn(t,e){for(var n={},r=0,i=F(t);r<i;r++)e?n[t[r]]=e[r]:n[t[r][0]]=t[r][1];return n}function mn(t,e,n){null==e&&(e=t||0,t=0),n||(n=e<t?-1:1);for(var r=Math.max(Math.ceil((e-t)/n),0),i=Array(r),a=0;a<r;a++,t+=n)i[a]=t;return i}function vn(t,e){if(null==e||e<1)return[];for(var n=[],r=0,a=t.length;r<a;)n.push(i.q.call(t,r,r+=e));return n}function gn(t,e){return t._chain?W(e).chain():e}function _n(t){return Ee(lt(t),(function(e){var n=W[e]=t[e];W.prototype[e]=function(){var t=[this._wrapped];return i.o.apply(t,arguments),gn(this,n.apply(W,t))}})),W}Ee(["pop","push","reverse","shift","sort","splice","unshift"],(function(t){var e=i.a[t];W.prototype[t]=function(){var n=this._wrapped;return null!=n&&(e.apply(n,arguments),"shift"!==t&&"splice"!==t||0!==n.length||delete n[0]),gn(this,n)}})),Ee(["concat","join","slice"],(function(t){var e=i.a[t];W.prototype[t]=function(){var t=this._wrapped;return null!=t&&(t=e.apply(t,arguments)),gn(this,t)}}));var yn=W,bn=_n(r);bn._=bn;var wn=bn},function(t,e,n){"use strict";var r=n(14);n.d(e,"a",(function(){return r.e})),n.d(e,"h",(function(){return r.g})),n.d(e,"e",(function(){return r.f}));var i=n(111);n.d(e,"f",(function(){return i.a})),n.d(e,"d",(function(){return i.c})),n.d(e,"g",(function(){return i.d})),n.d(e,"c",(function(){return i.b}));var a=n(171);n.d(e,"b",(function(){return a.a}))},function(t,e,n){t.exports=n(535)},function(t,e,n){var r=n(197)("wks"),i=n(137),a=n(18).Symbol,o="function"==typeof a;(t.exports=function(t){return r[t]||(r[t]=o&&a[t]||(o?a:i)("Symbol."+t))}).store=r},function(t,e,n){"use strict";n.d(e,"a",(function(){return r.a})),n.d(e,"b",(function(){return i.a})),n.d(e,"c",(function(){return a.b})),n.d(e,"d",(function(){return o.a})),n.d(e,"g",(function(){return u.a})),n.d(e,"h",(function(){return s})),n.d(e,"m",(function(){return l})),n.d(e,"o",(function(){return f.a})),n.d(e,"p",(function(){return d.a})),n.d(e,"q",(function(){return h.a})),n.d(e,"u",(function(){return p.a})),n.d(e,"v",(function(){return m.a})),n.d(e,"w",(function(){return v.a})),n.d(e,"x",(function(){return v.b})),n.d(e,"y",(function(){return g.a})),n.d(e,"r",(function(){return _.a})),n.d(e,"s",(function(){return _.b})),n.d(e,"t",(function(){return _.c})),n.d(e,"k",(function(){return w})),n.d(e,"l",(function(){return x})),n.d(e,"n",(function(){return M})),n.d(e,"i",(function(){return S})),n.d(e,"j",(function(){return T})),n.d(e,"e",(function(){return j.b})),n.d(e,"f",(function(){return j.a})),n.d(e,"z",(function(){return O})),n.d(e,"A",(function(){return A}));var r=n(98),i=n(165),a=n(94),o=n(159),u=n(164),s=function(t){var e=t.length;return function(n){return t[Math.max(0,Math.min(e-1,Math.floor(n*e)))]}},c=n(10),l=function(t,e){var n=Object(c.c)(+t,+e);return function(t){var e=n(t);return e-360*Math.floor(e/360)}},f=n(25),d=n(63),h=n(166),p=n(259),m=n(160),v=n(227),g=n(260),_=n(78),y=n(14);function b(t){return function(e,n){var r=t((e=Object(y.f)(e)).h,(n=Object(y.f)(n)).h),i=Object(c.a)(e.s,n.s),a=Object(c.a)(e.l,n.l),o=Object(c.a)(e.opacity,n.opacity);return function(t){return e.h=r(t),e.s=i(t),e.l=a(t),e.opacity=o(t),e+""}}}var w=b(c.c),x=b(c.a),k=n(111);function M(t,e){var n=Object(c.a)((t=Object(k.a)(t)).l,(e=Object(k.a)(e)).l),r=Object(c.a)(t.a,e.a),i=Object(c.a)(t.b,e.b),a=Object(c.a)(t.opacity,e.opacity);return function(e){return t.l=n(e),t.a=r(e),t.b=i(e),t.opacity=a(e),t+""}}function E(t){return function(e,n){var r=t((e=Object(k.c)(e)).h,(n=Object(k.c)(n)).h),i=Object(c.a)(e.c,n.c),a=Object(c.a)(e.l,n.l),o=Object(c.a)(e.opacity,n.opacity);return function(t){return e.h=r(t),e.c=i(t),e.l=a(t),e.opacity=o(t),e+""}}}var S=E(c.c),T=E(c.a),j=n(179);function O(t,e){for(var n=0,r=e.length-1,i=e[0],a=new Array(r<0?0:r);n<r;)a[n]=t(i,i=e[++n]);return function(t){var e=Math.max(0,Math.min(r-1,Math.floor(t*=r)));return a[e](t-e)}}var A=function(t,e){for(var n=new Array(e),r=0;r<e;++r)n[r]=t(r/(e-1));return n}},function(t,e,n){"use strict";n.d(e,"d",(function(){return o})),n.d(e,"e",(function(){return p})),n.d(e,"c",(function(){return a})),n.d(e,"b",(function(){return m})),n.d(e,"f",(function(){return v})),n.d(e,"a",(function(){return g}));function r(){}function i(t,e){var n=new r;if(t instanceof r)t.each((function(t,e){n.set(e,t)}));else if(Array.isArray(t)){var i,a=-1,o=t.length;if(null==e)for(;++a<o;)n.set(a,t[a]);else for(;++a<o;)n.set(e(i=t[a],a,t),i)}else if(t)for(var u in t)n.set(u,t[u]);return n}r.prototype=i.prototype={constructor:r,has:function(t){return"$"+t in this},get:function(t){return this["$"+t]},set:function(t,e){return this["$"+t]=e,this},remove:function(t){var e="$"+t;return e in this&&delete this[e]},clear:function(){for(var t in this)"$"===t[0]&&delete this[t]},keys:function(){var t=[];for(var e in this)"$"===e[0]&&t.push(e.slice(1));return t},values:function(){var t=[];for(var e in this)"$"===e[0]&&t.push(this[e]);return t},entries:function(){var t=[];for(var e in this)"$"===e[0]&&t.push({key:e.slice(1),value:this[e]});return t},size:function(){var t=0;for(var e in this)"$"===e[0]&&++t;return t},empty:function(){for(var t in this)if("$"===t[0])return!1;return!0},each:function(t){for(var e in this)"$"===e[0]&&t(this[e],e.slice(1),this)}};var a=i,o=function(){var t,e,n,r=[],i=[];function o(n,i,u,s){if(i>=r.length)return null!=t&&n.sort(t),null!=e?e(n):n;for(var c,l,f,d=-1,h=n.length,p=r[i++],m=a(),v=u();++d<h;)(f=m.get(c=p(l=n[d])+""))?f.push(l):m.set(c,[l]);return m.each((function(t,e){s(v,e,o(t,i,u,s))})),v}return n={object:function(t){return o(t,0,u,s)},map:function(t){return o(t,0,c,l)},entries:function(t){return function t(n,a){if(++a>r.length)return n;var o,u=i[a-1];return null!=e&&a>=r.length?o=n.entries():(o=[],n.each((function(e,n){o.push({key:n,values:t(e,a)})}))),null!=u?o.sort((function(t,e){return u(t.key,e.key)})):o}(o(t,0,c,l),0)},key:function(t){return r.push(t),n},sortKeys:function(t){return i[r.length-1]=t,n},sortValues:function(e){return t=e,n},rollup:function(t){return e=t,n}}};function u(){return{}}function s(t,e,n){t[e]=n}function c(){return a()}function l(t,e,n){t.set(e,n)}function f(){}var d=a.prototype;function h(t,e){var n=new f;if(t instanceof f)t.each((function(t){n.add(t)}));else if(t){var r=-1,i=t.length;if(null==e)for(;++r<i;)n.add(t[r]);else for(;++r<i;)n.add(e(t[r],r,t))}return n}f.prototype=h.prototype={constructor:f,has:d.has,add:function(t){return this["$"+(t+="")]=t,this},remove:d.remove,clear:d.clear,values:d.keys,size:d.size,empty:d.empty,each:d.each};var p=h,m=function(t){var e=[];for(var n in t)e.push(n);return e},v=function(t){var e=[];for(var n in t)e.push(t[n]);return e},g=function(t){var e=[];for(var n in t)e.push({key:n,value:t[n]});return e}},function(t,e,n){"use strict";e.a=function(t,e){return t=+t,e=+e,function(n){return t*(1-n)+e*n}}},function(t,e,n){"use strict";function r(t,e){var n=Object.create(t.prototype);for(var r in e)n[r]=e[r];return n}n.d(e,"b",(function(){return r})),e.a=function(t,e,n){t.prototype=e.prototype=n,n.constructor=t}},function(t,e){t.exports=function(t){return t.webpackPolyfill||(t.deprecate=function(){},t.paths=[],t.children||(t.children=[]),Object.defineProperty(t,"loaded",{enumerable:!0,get:function(){return t.l}}),Object.defineProperty(t,"id",{enumerable:!0,get:function(){return t.i}}),t.webpackPolyfill=1),t}},function(t,e,n){"use strict";(function(t){
/*!
 * The buffer module from node.js, for the browser.
 *
 * @author   Feross Aboukhadijeh <http://feross.org>
 * @license  MIT
 */
var r=n(597),i=n(598),a=n(599);function o(){return s.TYPED_ARRAY_SUPPORT?2147483647:1073741823}function u(t,e){if(o()<e)throw new RangeError("Invalid typed array length");return s.TYPED_ARRAY_SUPPORT?(t=new Uint8Array(e)).__proto__=s.prototype:(null===t&&(t=new s(e)),t.length=e),t}function s(t,e,n){if(!(s.TYPED_ARRAY_SUPPORT||this instanceof s))return new s(t,e,n);if("number"==typeof t){if("string"==typeof e)throw new Error("If encoding is specified then the first argument must be a string");return f(this,t)}return c(this,t,e,n)}function c(t,e,n,r){if("number"==typeof e)throw new TypeError('"value" argument must not be a number');return"undefined"!=typeof ArrayBuffer&&e instanceof ArrayBuffer?function(t,e,n,r){if(e.byteLength,n<0||e.byteLength<n)throw new RangeError("'offset' is out of bounds");if(e.byteLength<n+(r||0))throw new RangeError("'length' is out of bounds");e=void 0===n&&void 0===r?new Uint8Array(e):void 0===r?new Uint8Array(e,n):new Uint8Array(e,n,r);s.TYPED_ARRAY_SUPPORT?(t=e).__proto__=s.prototype:t=d(t,e);return t}(t,e,n,r):"string"==typeof e?function(t,e,n){"string"==typeof n&&""!==n||(n="utf8");if(!s.isEncoding(n))throw new TypeError('"encoding" must be a valid string encoding');var r=0|p(e,n),i=(t=u(t,r)).write(e,n);i!==r&&(t=t.slice(0,i));return t}(t,e,n):function(t,e){if(s.isBuffer(e)){var n=0|h(e.length);return 0===(t=u(t,n)).length||e.copy(t,0,0,n),t}if(e){if("undefined"!=typeof ArrayBuffer&&e.buffer instanceof ArrayBuffer||"length"in e)return"number"!=typeof e.length||(r=e.length)!=r?u(t,0):d(t,e);if("Buffer"===e.type&&a(e.data))return d(t,e.data)}var r;throw new TypeError("First argument must be a string, Buffer, ArrayBuffer, Array, or array-like object.")}(t,e)}function l(t){if("number"!=typeof t)throw new TypeError('"size" argument must be a number');if(t<0)throw new RangeError('"size" argument must not be negative')}function f(t,e){if(l(e),t=u(t,e<0?0:0|h(e)),!s.TYPED_ARRAY_SUPPORT)for(var n=0;n<e;++n)t[n]=0;return t}function d(t,e){var n=e.length<0?0:0|h(e.length);t=u(t,n);for(var r=0;r<n;r+=1)t[r]=255&e[r];return t}function h(t){if(t>=o())throw new RangeError("Attempt to allocate Buffer larger than maximum size: 0x"+o().toString(16)+" bytes");return 0|t}function p(t,e){if(s.isBuffer(t))return t.length;if("undefined"!=typeof ArrayBuffer&&"function"==typeof ArrayBuffer.isView&&(ArrayBuffer.isView(t)||t instanceof ArrayBuffer))return t.byteLength;"string"!=typeof t&&(t=""+t);var n=t.length;if(0===n)return 0;for(var r=!1;;)switch(e){case"ascii":case"latin1":case"binary":return n;case"utf8":case"utf-8":case void 0:return F(t).length;case"ucs2":case"ucs-2":case"utf16le":case"utf-16le":return 2*n;case"hex":return n>>>1;case"base64":return H(t).length;default:if(r)return F(t).length;e=(""+e).toLowerCase(),r=!0}}function m(t,e,n){var r=!1;if((void 0===e||e<0)&&(e=0),e>this.length)return"";if((void 0===n||n>this.length)&&(n=this.length),n<=0)return"";if((n>>>=0)<=(e>>>=0))return"";for(t||(t="utf8");;)switch(t){case"hex":return O(this,e,n);case"utf8":case"utf-8":return S(this,e,n);case"ascii":return T(this,e,n);case"latin1":case"binary":return j(this,e,n);case"base64":return E(this,e,n);case"ucs2":case"ucs-2":case"utf16le":case"utf-16le":return A(this,e,n);default:if(r)throw new TypeError("Unknown encoding: "+t);t=(t+"").toLowerCase(),r=!0}}function v(t,e,n){var r=t[e];t[e]=t[n],t[n]=r}function g(t,e,n,r,i){if(0===t.length)return-1;if("string"==typeof n?(r=n,n=0):n>2147483647?n=2147483647:n<-2147483648&&(n=-2147483648),n=+n,isNaN(n)&&(n=i?0:t.length-1),n<0&&(n=t.length+n),n>=t.length){if(i)return-1;n=t.length-1}else if(n<0){if(!i)return-1;n=0}if("string"==typeof e&&(e=s.from(e,r)),s.isBuffer(e))return 0===e.length?-1:_(t,e,n,r,i);if("number"==typeof e)return e&=255,s.TYPED_ARRAY_SUPPORT&&"function"==typeof Uint8Array.prototype.indexOf?i?Uint8Array.prototype.indexOf.call(t,e,n):Uint8Array.prototype.lastIndexOf.call(t,e,n):_(t,[e],n,r,i);throw new TypeError("val must be string, number or Buffer")}function _(t,e,n,r,i){var a,o=1,u=t.length,s=e.length;if(void 0!==r&&("ucs2"===(r=String(r).toLowerCase())||"ucs-2"===r||"utf16le"===r||"utf-16le"===r)){if(t.length<2||e.length<2)return-1;o=2,u/=2,s/=2,n/=2}function c(t,e){return 1===o?t[e]:t.readUInt16BE(e*o)}if(i){var l=-1;for(a=n;a<u;a++)if(c(t,a)===c(e,-1===l?0:a-l)){if(-1===l&&(l=a),a-l+1===s)return l*o}else-1!==l&&(a-=a-l),l=-1}else for(n+s>u&&(n=u-s),a=n;a>=0;a--){for(var f=!0,d=0;d<s;d++)if(c(t,a+d)!==c(e,d)){f=!1;break}if(f)return a}return-1}function y(t,e,n,r){n=Number(n)||0;var i=t.length-n;r?(r=Number(r))>i&&(r=i):r=i;var a=e.length;if(a%2!=0)throw new TypeError("Invalid hex string");r>a/2&&(r=a/2);for(var o=0;o<r;++o){var u=parseInt(e.substr(2*o,2),16);if(isNaN(u))return o;t[n+o]=u}return o}function b(t,e,n,r){return B(F(e,t.length-n),t,n,r)}function w(t,e,n,r){return B(function(t){for(var e=[],n=0;n<t.length;++n)e.push(255&t.charCodeAt(n));return e}(e),t,n,r)}function x(t,e,n,r){return w(t,e,n,r)}function k(t,e,n,r){return B(H(e),t,n,r)}function M(t,e,n,r){return B(function(t,e){for(var n,r,i,a=[],o=0;o<t.length&&!((e-=2)<0);++o)n=t.charCodeAt(o),r=n>>8,i=n%256,a.push(i),a.push(r);return a}(e,t.length-n),t,n,r)}function E(t,e,n){return 0===e&&n===t.length?r.fromByteArray(t):r.fromByteArray(t.slice(e,n))}function S(t,e,n){n=Math.min(t.length,n);for(var r=[],i=e;i<n;){var a,o,u,s,c=t[i],l=null,f=c>239?4:c>223?3:c>191?2:1;if(i+f<=n)switch(f){case 1:c<128&&(l=c);break;case 2:128==(192&(a=t[i+1]))&&(s=(31&c)<<6|63&a)>127&&(l=s);break;case 3:a=t[i+1],o=t[i+2],128==(192&a)&&128==(192&o)&&(s=(15&c)<<12|(63&a)<<6|63&o)>2047&&(s<55296||s>57343)&&(l=s);break;case 4:a=t[i+1],o=t[i+2],u=t[i+3],128==(192&a)&&128==(192&o)&&128==(192&u)&&(s=(15&c)<<18|(63&a)<<12|(63&o)<<6|63&u)>65535&&s<1114112&&(l=s)}null===l?(l=65533,f=1):l>65535&&(l-=65536,r.push(l>>>10&1023|55296),l=56320|1023&l),r.push(l),i+=f}return function(t){var e=t.length;if(e<=4096)return String.fromCharCode.apply(String,t);var n="",r=0;for(;r<e;)n+=String.fromCharCode.apply(String,t.slice(r,r+=4096));return n}(r)}e.Buffer=s,e.SlowBuffer=function(t){+t!=t&&(t=0);return s.alloc(+t)},e.INSPECT_MAX_BYTES=50,s.TYPED_ARRAY_SUPPORT=void 0!==t.TYPED_ARRAY_SUPPORT?t.TYPED_ARRAY_SUPPORT:function(){try{var t=new Uint8Array(1);return t.__proto__={__proto__:Uint8Array.prototype,foo:function(){return 42}},42===t.foo()&&"function"==typeof t.subarray&&0===t.subarray(1,1).byteLength}catch(t){return!1}}(),e.kMaxLength=o(),s.poolSize=8192,s._augment=function(t){return t.__proto__=s.prototype,t},s.from=function(t,e,n){return c(null,t,e,n)},s.TYPED_ARRAY_SUPPORT&&(s.prototype.__proto__=Uint8Array.prototype,s.__proto__=Uint8Array,"undefined"!=typeof Symbol&&Symbol.species&&s[Symbol.species]===s&&Object.defineProperty(s,Symbol.species,{value:null,configurable:!0})),s.alloc=function(t,e,n){return function(t,e,n,r){return l(e),e<=0?u(t,e):void 0!==n?"string"==typeof r?u(t,e).fill(n,r):u(t,e).fill(n):u(t,e)}(null,t,e,n)},s.allocUnsafe=function(t){return f(null,t)},s.allocUnsafeSlow=function(t){return f(null,t)},s.isBuffer=function(t){return!(null==t||!t._isBuffer)},s.compare=function(t,e){if(!s.isBuffer(t)||!s.isBuffer(e))throw new TypeError("Arguments must be Buffers");if(t===e)return 0;for(var n=t.length,r=e.length,i=0,a=Math.min(n,r);i<a;++i)if(t[i]!==e[i]){n=t[i],r=e[i];break}return n<r?-1:r<n?1:0},s.isEncoding=function(t){switch(String(t).toLowerCase()){case"hex":case"utf8":case"utf-8":case"ascii":case"latin1":case"binary":case"base64":case"ucs2":case"ucs-2":case"utf16le":case"utf-16le":return!0;default:return!1}},s.concat=function(t,e){if(!a(t))throw new TypeError('"list" argument must be an Array of Buffers');if(0===t.length)return s.alloc(0);var n;if(void 0===e)for(e=0,n=0;n<t.length;++n)e+=t[n].length;var r=s.allocUnsafe(e),i=0;for(n=0;n<t.length;++n){var o=t[n];if(!s.isBuffer(o))throw new TypeError('"list" argument must be an Array of Buffers');o.copy(r,i),i+=o.length}return r},s.byteLength=p,s.prototype._isBuffer=!0,s.prototype.swap16=function(){var t=this.length;if(t%2!=0)throw new RangeError("Buffer size must be a multiple of 16-bits");for(var e=0;e<t;e+=2)v(this,e,e+1);return this},s.prototype.swap32=function(){var t=this.length;if(t%4!=0)throw new RangeError("Buffer size must be a multiple of 32-bits");for(var e=0;e<t;e+=4)v(this,e,e+3),v(this,e+1,e+2);return this},s.prototype.swap64=function(){var t=this.length;if(t%8!=0)throw new RangeError("Buffer size must be a multiple of 64-bits");for(var e=0;e<t;e+=8)v(this,e,e+7),v(this,e+1,e+6),v(this,e+2,e+5),v(this,e+3,e+4);return this},s.prototype.toString=function(){var t=0|this.length;return 0===t?"":0===arguments.length?S(this,0,t):m.apply(this,arguments)},s.prototype.equals=function(t){if(!s.isBuffer(t))throw new TypeError("Argument must be a Buffer");return this===t||0===s.compare(this,t)},s.prototype.inspect=function(){var t="",n=e.INSPECT_MAX_BYTES;return this.length>0&&(t=this.toString("hex",0,n).match(/.{2}/g).join(" "),this.length>n&&(t+=" ... ")),"<Buffer "+t+">"},s.prototype.compare=function(t,e,n,r,i){if(!s.isBuffer(t))throw new TypeError("Argument must be a Buffer");if(void 0===e&&(e=0),void 0===n&&(n=t?t.length:0),void 0===r&&(r=0),void 0===i&&(i=this.length),e<0||n>t.length||r<0||i>this.length)throw new RangeError("out of range index");if(r>=i&&e>=n)return 0;if(r>=i)return-1;if(e>=n)return 1;if(this===t)return 0;for(var a=(i>>>=0)-(r>>>=0),o=(n>>>=0)-(e>>>=0),u=Math.min(a,o),c=this.slice(r,i),l=t.slice(e,n),f=0;f<u;++f)if(c[f]!==l[f]){a=c[f],o=l[f];break}return a<o?-1:o<a?1:0},s.prototype.includes=function(t,e,n){return-1!==this.indexOf(t,e,n)},s.prototype.indexOf=function(t,e,n){return g(this,t,e,n,!0)},s.prototype.lastIndexOf=function(t,e,n){return g(this,t,e,n,!1)},s.prototype.write=function(t,e,n,r){if(void 0===e)r="utf8",n=this.length,e=0;else if(void 0===n&&"string"==typeof e)r=e,n=this.length,e=0;else{if(!isFinite(e))throw new Error("Buffer.write(string, encoding, offset[, length]) is no longer supported");e|=0,isFinite(n)?(n|=0,void 0===r&&(r="utf8")):(r=n,n=void 0)}var i=this.length-e;if((void 0===n||n>i)&&(n=i),t.length>0&&(n<0||e<0)||e>this.length)throw new RangeError("Attempt to write outside buffer bounds");r||(r="utf8");for(var a=!1;;)switch(r){case"hex":return y(this,t,e,n);case"utf8":case"utf-8":return b(this,t,e,n);case"ascii":return w(this,t,e,n);case"latin1":case"binary":return x(this,t,e,n);case"base64":return k(this,t,e,n);case"ucs2":case"ucs-2":case"utf16le":case"utf-16le":return M(this,t,e,n);default:if(a)throw new TypeError("Unknown encoding: "+r);r=(""+r).toLowerCase(),a=!0}},s.prototype.toJSON=function(){return{type:"Buffer",data:Array.prototype.slice.call(this._arr||this,0)}};function T(t,e,n){var r="";n=Math.min(t.length,n);for(var i=e;i<n;++i)r+=String.fromCharCode(127&t[i]);return r}function j(t,e,n){var r="";n=Math.min(t.length,n);for(var i=e;i<n;++i)r+=String.fromCharCode(t[i]);return r}function O(t,e,n){var r=t.length;(!e||e<0)&&(e=0),(!n||n<0||n>r)&&(n=r);for(var i="",a=e;a<n;++a)i+=z(t[a]);return i}function A(t,e,n){for(var r=t.slice(e,n),i="",a=0;a<r.length;a+=2)i+=String.fromCharCode(r[a]+256*r[a+1]);return i}function L(t,e,n){if(t%1!=0||t<0)throw new RangeError("offset is not uint");if(t+e>n)throw new RangeError("Trying to access beyond buffer length")}function C(t,e,n,r,i,a){if(!s.isBuffer(t))throw new TypeError('"buffer" argument must be a Buffer instance');if(e>i||e<a)throw new RangeError('"value" argument is out of bounds');if(n+r>t.length)throw new RangeError("Index out of range")}function D(t,e,n,r){e<0&&(e=65535+e+1);for(var i=0,a=Math.min(t.length-n,2);i<a;++i)t[n+i]=(e&255<<8*(r?i:1-i))>>>8*(r?i:1-i)}function N(t,e,n,r){e<0&&(e=4294967295+e+1);for(var i=0,a=Math.min(t.length-n,4);i<a;++i)t[n+i]=e>>>8*(r?i:3-i)&255}function P(t,e,n,r,i,a){if(n+r>t.length)throw new RangeError("Index out of range");if(n<0)throw new RangeError("Index out of range")}function R(t,e,n,r,a){return a||P(t,0,n,4),i.write(t,e,n,r,23,4),n+4}function Y(t,e,n,r,a){return a||P(t,0,n,8),i.write(t,e,n,r,52,8),n+8}s.prototype.slice=function(t,e){var n,r=this.length;if((t=~~t)<0?(t+=r)<0&&(t=0):t>r&&(t=r),(e=void 0===e?r:~~e)<0?(e+=r)<0&&(e=0):e>r&&(e=r),e<t&&(e=t),s.TYPED_ARRAY_SUPPORT)(n=this.subarray(t,e)).__proto__=s.prototype;else{var i=e-t;n=new s(i,void 0);for(var a=0;a<i;++a)n[a]=this[a+t]}return n},s.prototype.readUIntLE=function(t,e,n){t|=0,e|=0,n||L(t,e,this.length);for(var r=this[t],i=1,a=0;++a<e&&(i*=256);)r+=this[t+a]*i;return r},s.prototype.readUIntBE=function(t,e,n){t|=0,e|=0,n||L(t,e,this.length);for(var r=this[t+--e],i=1;e>0&&(i*=256);)r+=this[t+--e]*i;return r},s.prototype.readUInt8=function(t,e){return e||L(t,1,this.length),this[t]},s.prototype.readUInt16LE=function(t,e){return e||L(t,2,this.length),this[t]|this[t+1]<<8},s.prototype.readUInt16BE=function(t,e){return e||L(t,2,this.length),this[t]<<8|this[t+1]},s.prototype.readUInt32LE=function(t,e){return e||L(t,4,this.length),(this[t]|this[t+1]<<8|this[t+2]<<16)+16777216*this[t+3]},s.prototype.readUInt32BE=function(t,e){return e||L(t,4,this.length),16777216*this[t]+(this[t+1]<<16|this[t+2]<<8|this[t+3])},s.prototype.readIntLE=function(t,e,n){t|=0,e|=0,n||L(t,e,this.length);for(var r=this[t],i=1,a=0;++a<e&&(i*=256);)r+=this[t+a]*i;return r>=(i*=128)&&(r-=Math.pow(2,8*e)),r},s.prototype.readIntBE=function(t,e,n){t|=0,e|=0,n||L(t,e,this.length);for(var r=e,i=1,a=this[t+--r];r>0&&(i*=256);)a+=this[t+--r]*i;return a>=(i*=128)&&(a-=Math.pow(2,8*e)),a},s.prototype.readInt8=function(t,e){return e||L(t,1,this.length),128&this[t]?-1*(255-this[t]+1):this[t]},s.prototype.readInt16LE=function(t,e){e||L(t,2,this.length);var n=this[t]|this[t+1]<<8;return 32768&n?4294901760|n:n},s.prototype.readInt16BE=function(t,e){e||L(t,2,this.length);var n=this[t+1]|this[t]<<8;return 32768&n?4294901760|n:n},s.prototype.readInt32LE=function(t,e){return e||L(t,4,this.length),this[t]|this[t+1]<<8|this[t+2]<<16|this[t+3]<<24},s.prototype.readInt32BE=function(t,e){return e||L(t,4,this.length),this[t]<<24|this[t+1]<<16|this[t+2]<<8|this[t+3]},s.prototype.readFloatLE=function(t,e){return e||L(t,4,this.length),i.read(this,t,!0,23,4)},s.prototype.readFloatBE=function(t,e){return e||L(t,4,this.length),i.read(this,t,!1,23,4)},s.prototype.readDoubleLE=function(t,e){return e||L(t,8,this.length),i.read(this,t,!0,52,8)},s.prototype.readDoubleBE=function(t,e){return e||L(t,8,this.length),i.read(this,t,!1,52,8)},s.prototype.writeUIntLE=function(t,e,n,r){(t=+t,e|=0,n|=0,r)||C(this,t,e,n,Math.pow(2,8*n)-1,0);var i=1,a=0;for(this[e]=255&t;++a<n&&(i*=256);)this[e+a]=t/i&255;return e+n},s.prototype.writeUIntBE=function(t,e,n,r){(t=+t,e|=0,n|=0,r)||C(this,t,e,n,Math.pow(2,8*n)-1,0);var i=n-1,a=1;for(this[e+i]=255&t;--i>=0&&(a*=256);)this[e+i]=t/a&255;return e+n},s.prototype.writeUInt8=function(t,e,n){return t=+t,e|=0,n||C(this,t,e,1,255,0),s.TYPED_ARRAY_SUPPORT||(t=Math.floor(t)),this[e]=255&t,e+1},s.prototype.writeUInt16LE=function(t,e,n){return t=+t,e|=0,n||C(this,t,e,2,65535,0),s.TYPED_ARRAY_SUPPORT?(this[e]=255&t,this[e+1]=t>>>8):D(this,t,e,!0),e+2},s.prototype.writeUInt16BE=function(t,e,n){return t=+t,e|=0,n||C(this,t,e,2,65535,0),s.TYPED_ARRAY_SUPPORT?(this[e]=t>>>8,this[e+1]=255&t):D(this,t,e,!1),e+2},s.prototype.writeUInt32LE=function(t,e,n){return t=+t,e|=0,n||C(this,t,e,4,4294967295,0),s.TYPED_ARRAY_SUPPORT?(this[e+3]=t>>>24,this[e+2]=t>>>16,this[e+1]=t>>>8,this[e]=255&t):N(this,t,e,!0),e+4},s.prototype.writeUInt32BE=function(t,e,n){return t=+t,e|=0,n||C(this,t,e,4,4294967295,0),s.TYPED_ARRAY_SUPPORT?(this[e]=t>>>24,this[e+1]=t>>>16,this[e+2]=t>>>8,this[e+3]=255&t):N(this,t,e,!1),e+4},s.prototype.writeIntLE=function(t,e,n,r){if(t=+t,e|=0,!r){var i=Math.pow(2,8*n-1);C(this,t,e,n,i-1,-i)}var a=0,o=1,u=0;for(this[e]=255&t;++a<n&&(o*=256);)t<0&&0===u&&0!==this[e+a-1]&&(u=1),this[e+a]=(t/o>>0)-u&255;return e+n},s.prototype.writeIntBE=function(t,e,n,r){if(t=+t,e|=0,!r){var i=Math.pow(2,8*n-1);C(this,t,e,n,i-1,-i)}var a=n-1,o=1,u=0;for(this[e+a]=255&t;--a>=0&&(o*=256);)t<0&&0===u&&0!==this[e+a+1]&&(u=1),this[e+a]=(t/o>>0)-u&255;return e+n},s.prototype.writeInt8=function(t,e,n){return t=+t,e|=0,n||C(this,t,e,1,127,-128),s.TYPED_ARRAY_SUPPORT||(t=Math.floor(t)),t<0&&(t=255+t+1),this[e]=255&t,e+1},s.prototype.writeInt16LE=function(t,e,n){return t=+t,e|=0,n||C(this,t,e,2,32767,-32768),s.TYPED_ARRAY_SUPPORT?(this[e]=255&t,this[e+1]=t>>>8):D(this,t,e,!0),e+2},s.prototype.writeInt16BE=function(t,e,n){return t=+t,e|=0,n||C(this,t,e,2,32767,-32768),s.TYPED_ARRAY_SUPPORT?(this[e]=t>>>8,this[e+1]=255&t):D(this,t,e,!1),e+2},s.prototype.writeInt32LE=function(t,e,n){return t=+t,e|=0,n||C(this,t,e,4,2147483647,-2147483648),s.TYPED_ARRAY_SUPPORT?(this[e]=255&t,this[e+1]=t>>>8,this[e+2]=t>>>16,this[e+3]=t>>>24):N(this,t,e,!0),e+4},s.prototype.writeInt32BE=function(t,e,n){return t=+t,e|=0,n||C(this,t,e,4,2147483647,-2147483648),t<0&&(t=4294967295+t+1),s.TYPED_ARRAY_SUPPORT?(this[e]=t>>>24,this[e+1]=t>>>16,this[e+2]=t>>>8,this[e+3]=255&t):N(this,t,e,!1),e+4},s.prototype.writeFloatLE=function(t,e,n){return R(this,t,e,!0,n)},s.prototype.writeFloatBE=function(t,e,n){return R(this,t,e,!1,n)},s.prototype.writeDoubleLE=function(t,e,n){return Y(this,t,e,!0,n)},s.prototype.writeDoubleBE=function(t,e,n){return Y(this,t,e,!1,n)},s.prototype.copy=function(t,e,n,r){if(n||(n=0),r||0===r||(r=this.length),e>=t.length&&(e=t.length),e||(e=0),r>0&&r<n&&(r=n),r===n)return 0;if(0===t.length||0===this.length)return 0;if(e<0)throw new RangeError("targetStart out of bounds");if(n<0||n>=this.length)throw new RangeError("sourceStart out of bounds");if(r<0)throw new RangeError("sourceEnd out of bounds");r>this.length&&(r=this.length),t.length-e<r-n&&(r=t.length-e+n);var i,a=r-n;if(this===t&&n<e&&e<r)for(i=a-1;i>=0;--i)t[i+e]=this[i+n];else if(a<1e3||!s.TYPED_ARRAY_SUPPORT)for(i=0;i<a;++i)t[i+e]=this[i+n];else Uint8Array.prototype.set.call(t,this.subarray(n,n+a),e);return a},s.prototype.fill=function(t,e,n,r){if("string"==typeof t){if("string"==typeof e?(r=e,e=0,n=this.length):"string"==typeof n&&(r=n,n=this.length),1===t.length){var i=t.charCodeAt(0);i<256&&(t=i)}if(void 0!==r&&"string"!=typeof r)throw new TypeError("encoding must be a string");if("string"==typeof r&&!s.isEncoding(r))throw new TypeError("Unknown encoding: "+r)}else"number"==typeof t&&(t&=255);if(e<0||this.length<e||this.length<n)throw new RangeError("Out of range index");if(n<=e)return this;var a;if(e>>>=0,n=void 0===n?this.length:n>>>0,t||(t=0),"number"==typeof t)for(a=e;a<n;++a)this[a]=t;else{var o=s.isBuffer(t)?t:F(new s(t,r).toString()),u=o.length;for(a=0;a<n-e;++a)this[a+e]=o[a%u]}return this};var I=/[^+\/0-9A-Za-z-_]/g;function z(t){return t<16?"0"+t.toString(16):t.toString(16)}function F(t,e){var n;e=e||1/0;for(var r=t.length,i=null,a=[],o=0;o<r;++o){if((n=t.charCodeAt(o))>55295&&n<57344){if(!i){if(n>56319){(e-=3)>-1&&a.push(239,191,189);continue}if(o+1===r){(e-=3)>-1&&a.push(239,191,189);continue}i=n;continue}if(n<56320){(e-=3)>-1&&a.push(239,191,189),i=n;continue}n=65536+(i-55296<<10|n-56320)}else i&&(e-=3)>-1&&a.push(239,191,189);if(i=null,n<128){if((e-=1)<0)break;a.push(n)}else if(n<2048){if((e-=2)<0)break;a.push(n>>6|192,63&n|128)}else if(n<65536){if((e-=3)<0)break;a.push(n>>12|224,n>>6&63|128,63&n|128)}else{if(!(n<1114112))throw new Error("Invalid code point");if((e-=4)<0)break;a.push(n>>18|240,n>>12&63|128,n>>6&63|128,63&n|128)}}return a}function H(t){return r.toByteArray(function(t){if((t=function(t){return t.trim?t.trim():t.replace(/^\s+|\s+$/g,"")}(t).replace(I,"")).length<2)return"";for(;t.length%4!=0;)t+="=";return t}(t))}function B(t,e,n,r){for(var i=0;i<r&&!(i+n>=e.length||i>=t.length);++i)e[i+n]=t[i];return i}}).call(this,n(6))},function(t,e){t.exports=function(){throw new Error("define cannot be used indirect")}},function(t,e,n){"use strict";t.exports=function(t){return i.test("number"==typeof t?r(t):t.charAt(0))};var r=String.fromCharCode,i=/\s/},function(t,e,n){var r=n(54);t.exports=function(t){if(!r(t))throw TypeError(t+" is not an object!");return t}},function(t,e,n){"use strict";n.d(e,"c",(function(){return K})),n.d(e,"a",(function(){return it})),n.d(e,"b",(function(){return h}));var r=n(44),i=n(252),a=n(49),o=n(253),u=Object(i.a)("start","end","cancel","interrupt"),s=[],c=function(t,e,n,r,i,c){var l=t.__transition;if(l){if(n in l)return}else t.__transition={};!function(t,e,n){var r,i=t.__transition;function u(a){var l,f,d,h;if(1!==n.state)return c();for(l in i)if((h=i[l]).name===n.name){if(3===h.state)return Object(o.a)(u);4===h.state?(h.state=6,h.timer.stop(),h.on.call("interrupt",t,t.__data__,h.index,h.group),delete i[l]):+l<e&&(h.state=6,h.timer.stop(),h.on.call("cancel",t,t.__data__,h.index,h.group),delete i[l])}if(Object(o.a)((function(){3===n.state&&(n.state=4,n.timer.restart(s,n.delay,n.time),s(a))})),n.state=2,n.on.call("start",t,t.__data__,n.index,n.group),2===n.state){for(n.state=3,r=new Array(d=n.tween.length),l=0,f=-1;l<d;++l)(h=n.tween[l].value.call(t,t.__data__,n.index,n.group))&&(r[++f]=h);r.length=f+1}}function s(e){for(var i=e<n.duration?n.ease.call(null,e/n.duration):(n.timer.restart(c),n.state=5,1),a=-1,o=r.length;++a<o;)r[a].call(t,i);5===n.state&&(n.on.call("end",t,t.__data__,n.index,n.group),c())}function c(){for(var r in n.state=6,n.timer.stop(),delete i[e],i)return;delete t.__transition}i[e]=n,n.timer=Object(a.c)((function(t){n.state=1,n.timer.restart(u,n.delay,n.time),n.delay<=t&&u(t-n.delay)}),0,n.time)}(t,n,{name:e,index:r,group:i,on:u,tween:s,time:c.time,delay:c.delay,duration:c.duration,ease:c.ease,timer:null,state:0})};function l(t,e){var n=d(t,e);if(n.state>0)throw new Error("too late; already scheduled");return n}function f(t,e){var n=d(t,e);if(n.state>3)throw new Error("too late; already running");return n}function d(t,e){var n=t.__transition;if(!n||!(n=n[e]))throw new Error("transition not found");return n}var h=function(t,e){var n,r,i,a=t.__transition,o=!0;if(a){for(i in e=null==e?null:e+"",a)(n=a[i]).name===e?(r=n.state>2&&n.state<5,n.state=6,n.timer.stop(),n.on.call(r?"interrupt":"cancel",t,t.__data__,n.index,n.group),delete a[i]):o=!1;o&&delete t.__transition}},p=n(227),m=n(95);function v(t,e){var n,r;return function(){var i=f(this,t),a=i.tween;if(a!==n)for(var o=0,u=(r=n=a).length;o<u;++o)if(r[o].name===e){(r=r.slice()).splice(o,1);break}i.tween=r}}function g(t,e,n){var r,i;if("function"!=typeof n)throw new Error;return function(){var a=f(this,t),o=a.tween;if(o!==r){i=(r=o).slice();for(var u={name:e,value:n},s=0,c=i.length;s<c;++s)if(i[s].name===e){i[s]=u;break}s===c&&i.push(u)}a.tween=i}}function _(t,e,n){var r=t._id;return t.each((function(){var t=f(this,r);(t.value||(t.value={}))[e]=n.apply(this,arguments)})),function(t){return d(t,r).value[e]}}var y=n(14),b=n(25),w=n(78),x=n(160),k=function(t,e){var n;return("number"==typeof e?b.a:e instanceof y.e?w.a:(n=Object(y.e)(e))?(e=n,w.a):x.a)(t,e)};function M(t){return function(){this.removeAttribute(t)}}function E(t){return function(){this.removeAttributeNS(t.space,t.local)}}function S(t,e,n){var r,i,a=n+"";return function(){var o=this.getAttribute(t);return o===a?null:o===r?i:i=e(r=o,n)}}function T(t,e,n){var r,i,a=n+"";return function(){var o=this.getAttributeNS(t.space,t.local);return o===a?null:o===r?i:i=e(r=o,n)}}function j(t,e,n){var r,i,a;return function(){var o,u,s=n(this);if(null!=s)return(o=this.getAttribute(t))===(u=s+"")?null:o===r&&u===i?a:(i=u,a=e(r=o,s));this.removeAttribute(t)}}function O(t,e,n){var r,i,a;return function(){var o,u,s=n(this);if(null!=s)return(o=this.getAttributeNS(t.space,t.local))===(u=s+"")?null:o===r&&u===i?a:(i=u,a=e(r=o,s));this.removeAttributeNS(t.space,t.local)}}function A(t,e){return function(n){this.setAttribute(t,e.call(this,n))}}function L(t,e){return function(n){this.setAttributeNS(t.space,t.local,e.call(this,n))}}function C(t,e){var n,r;function i(){var i=e.apply(this,arguments);return i!==r&&(n=(r=i)&&L(t,i)),n}return i._value=e,i}function D(t,e){var n,r;function i(){var i=e.apply(this,arguments);return i!==r&&(n=(r=i)&&A(t,i)),n}return i._value=e,i}function N(t,e){return function(){l(this,t).delay=+e.apply(this,arguments)}}function P(t,e){return e=+e,function(){l(this,t).delay=e}}function R(t,e){return function(){f(this,t).duration=+e.apply(this,arguments)}}function Y(t,e){return e=+e,function(){f(this,t).duration=e}}function I(t,e){if("function"!=typeof e)throw new Error;return function(){f(this,t).ease=e}}var z=n(161);function F(t,e,n){var r,i,a=function(t){return(t+"").trim().split(/^|\s+/).every((function(t){var e=t.indexOf(".");return e>=0&&(t=t.slice(0,e)),!t||"start"===t}))}(e)?l:f;return function(){var o=a(this,t),u=o.on;u!==r&&(i=(r=u).copy()).on(e,n),o.on=i}}var H=n(96),B=n(162),q=r.b.prototype.constructor,U=n(163);function W(t){return function(){this.style.removeProperty(t)}}function G(t,e,n){return function(r){this.style.setProperty(t,e.call(this,r),n)}}function V(t,e,n){var r,i;function a(){var a=e.apply(this,arguments);return a!==i&&(r=(i=a)&&G(t,a,n)),r}return a._value=e,a}function $(t){return function(e){this.textContent=t.call(this,e)}}function X(t){var e,n;function r(){var r=t.apply(this,arguments);return r!==n&&(e=(n=r)&&$(r)),e}return r._value=t,r}var J=0;function Z(t,e,n,r){this._groups=t,this._parents=e,this._name=n,this._id=r}function K(t){return Object(r.b)().transition(t)}function Q(){return++J}var tt=r.b.prototype;Z.prototype=K.prototype={constructor:Z,select:function(t){var e=this._name,n=this._id;"function"!=typeof t&&(t=Object(H.a)(t));for(var r=this._groups,i=r.length,a=new Array(i),o=0;o<i;++o)for(var u,s,l=r[o],f=l.length,h=a[o]=new Array(f),p=0;p<f;++p)(u=l[p])&&(s=t.call(u,u.__data__,p,l))&&("__data__"in u&&(s.__data__=u.__data__),h[p]=s,c(h[p],e,n,p,h,d(u,n)));return new Z(a,this._parents,e,n)},selectAll:function(t){var e=this._name,n=this._id;"function"!=typeof t&&(t=Object(B.a)(t));for(var r=this._groups,i=r.length,a=[],o=[],u=0;u<i;++u)for(var s,l=r[u],f=l.length,h=0;h<f;++h)if(s=l[h]){for(var p,m=t.call(s,s.__data__,h,l),v=d(s,n),g=0,_=m.length;g<_;++g)(p=m[g])&&c(p,e,n,g,m,v);a.push(m),o.push(s)}return new Z(a,o,e,n)},filter:function(t){"function"!=typeof t&&(t=Object(z.a)(t));for(var e=this._groups,n=e.length,r=new Array(n),i=0;i<n;++i)for(var a,o=e[i],u=o.length,s=r[i]=[],c=0;c<u;++c)(a=o[c])&&t.call(a,a.__data__,c,o)&&s.push(a);return new Z(r,this._parents,this._name,this._id)},merge:function(t){if(t._id!==this._id)throw new Error;for(var e=this._groups,n=t._groups,r=e.length,i=n.length,a=Math.min(r,i),o=new Array(r),u=0;u<a;++u)for(var s,c=e[u],l=n[u],f=c.length,d=o[u]=new Array(f),h=0;h<f;++h)(s=c[h]||l[h])&&(d[h]=s);for(;u<r;++u)o[u]=e[u];return new Z(o,this._parents,this._name,this._id)},selection:function(){return new q(this._groups,this._parents)},transition:function(){for(var t=this._name,e=this._id,n=Q(),r=this._groups,i=r.length,a=0;a<i;++a)for(var o,u=r[a],s=u.length,l=0;l<s;++l)if(o=u[l]){var f=d(o,e);c(o,t,n,l,u,{time:f.time+f.delay+f.duration,delay:0,duration:f.duration,ease:f.ease})}return new Z(r,this._parents,t,n)},call:tt.call,nodes:tt.nodes,node:tt.node,size:tt.size,empty:tt.empty,each:tt.each,on:function(t,e){var n=this._id;return arguments.length<2?d(this.node(),n).on.on(t):this.each(F(n,t,e))},attr:function(t,e){var n=Object(m.a)(t),r="transform"===n?p.b:k;return this.attrTween(t,"function"==typeof e?(n.local?O:j)(n,r,_(this,"attr."+t,e)):null==e?(n.local?E:M)(n):(n.local?T:S)(n,r,e))},attrTween:function(t,e){var n="attr."+t;if(arguments.length<2)return(n=this.tween(n))&&n._value;if(null==e)return this.tween(n,null);if("function"!=typeof e)throw new Error;var r=Object(m.a)(t);return this.tween(n,(r.local?C:D)(r,e))},style:function(t,e,n){var r="transform"==(t+="")?p.a:k;return null==e?this.styleTween(t,function(t,e){var n,r,i;return function(){var a=Object(U.b)(this,t),o=(this.style.removeProperty(t),Object(U.b)(this,t));return a===o?null:a===n&&o===r?i:i=e(n=a,r=o)}}(t,r)).on("end.style."+t,W(t)):"function"==typeof e?this.styleTween(t,function(t,e,n){var r,i,a;return function(){var o=Object(U.b)(this,t),u=n(this),s=u+"";return null==u&&(this.style.removeProperty(t),s=u=Object(U.b)(this,t)),o===s?null:o===r&&s===i?a:(i=s,a=e(r=o,u))}}(t,r,_(this,"style."+t,e))).each(function(t,e){var n,r,i,a,o="style."+e,u="end."+o;return function(){var s=f(this,t),c=s.on,l=null==s.value[o]?a||(a=W(e)):void 0;c===n&&i===l||(r=(n=c).copy()).on(u,i=l),s.on=r}}(this._id,t)):this.styleTween(t,function(t,e,n){var r,i,a=n+"";return function(){var o=Object(U.b)(this,t);return o===a?null:o===r?i:i=e(r=o,n)}}(t,r,e),n).on("end.style."+t,null)},styleTween:function(t,e,n){var r="style."+(t+="");if(arguments.length<2)return(r=this.tween(r))&&r._value;if(null==e)return this.tween(r,null);if("function"!=typeof e)throw new Error;return this.tween(r,V(t,e,null==n?"":n))},text:function(t){return this.tween("text","function"==typeof t?function(t){return function(){var e=t(this);this.textContent=null==e?"":e}}(_(this,"text",t)):function(t){return function(){this.textContent=t}}(null==t?"":t+""))},textTween:function(t){var e="text";if(arguments.length<1)return(e=this.tween(e))&&e._value;if(null==t)return this.tween(e,null);if("function"!=typeof t)throw new Error;return this.tween(e,X(t))},remove:function(){return this.on("end.remove",(t=this._id,function(){var e=this.parentNode;for(var n in this.__transition)if(+n!==t)return;e&&e.removeChild(this)}));var t},tween:function(t,e){var n=this._id;if(t+="",arguments.length<2){for(var r,i=d(this.node(),n).tween,a=0,o=i.length;a<o;++a)if((r=i[a]).name===t)return r.value;return null}return this.each((null==e?v:g)(n,t,e))},delay:function(t){var e=this._id;return arguments.length?this.each(("function"==typeof t?N:P)(e,t)):d(this.node(),e).delay},duration:function(t){var e=this._id;return arguments.length?this.each(("function"==typeof t?R:Y)(e,t)):d(this.node(),e).duration},ease:function(t){var e=this._id;return arguments.length?this.each(I(e,t)):d(this.node(),e).ease},end:function(){var t,e,n=this,r=n._id,i=n.size();return new Promise((function(a,o){var u={value:o},s={value:function(){0==--i&&a()}};n.each((function(){var n=f(this,r),i=n.on;i!==t&&((e=(t=i).copy())._.cancel.push(u),e._.interrupt.push(u),e._.end.push(s)),n.on=e}))}))}};var et={time:null,delay:0,duration:250,ease:n(110).b};function nt(t,e){for(var n;!(n=t.__transition)||!(n=n[e]);)if(!(t=t.parentNode))return et.time=Object(a.b)(),et;return n}r.b.prototype.interrupt=function(t){return this.each((function(){h(this,t)}))},r.b.prototype.transition=function(t){var e,n;t instanceof Z?(e=t._id,t=t._name):(e=Q(),(n=et).time=Object(a.b)(),t=null==t?null:t+"");for(var r=this._groups,i=r.length,o=0;o<i;++o)for(var u,s=r[o],l=s.length,f=0;f<l;++f)(u=s[f])&&c(u,t,e,f,s,n||nt(u,e));return new Z(r,this._parents,t,e)};var rt=[null],it=function(t,e){var n,r,i=t.__transition;if(i)for(r in e=null==e?null:e+"",i)if((n=i[r]).state>1&&n.name===e)return new Z([[t]],rt,e,+r);return null}},function(t,e,n){"use strict";n.d(e,"b",(function(){return i}));var r=n(42);function i(){r.c.stopImmediatePropagation()}e.a=function(){r.c.preventDefault(),r.c.stopImmediatePropagation()}},function(t,e,n){(function(e){t.exports=e.d3=n(509)}).call(this,n(6))},function(t,e,n){"use strict";var r=n(228);n.d(e,"a",(function(){return r.a}))},function(t,e){var n,r,i=t.exports={};function a(){throw new Error("setTimeout has not been defined")}function o(){throw new Error("clearTimeout has not been defined")}function u(t){if(n===setTimeout)return setTimeout(t,0);if((n===a||!n)&&setTimeout)return n=setTimeout,setTimeout(t,0);try{return n(t,0)}catch(e){try{return n.call(null,t,0)}catch(e){return n.call(this,t,0)}}}!function(){try{n="function"==typeof setTimeout?setTimeout:a}catch(t){n=a}try{r="function"==typeof clearTimeout?clearTimeout:o}catch(t){r=o}}();var s,c=[],l=!1,f=-1;function d(){l&&s&&(l=!1,s.length?c=s.concat(c):f=-1,c.length&&h())}function h(){if(!l){var t=u(d);l=!0;for(var e=c.length;e;){for(s=c,c=[];++f<e;)s&&s[f].run();f=-1,e=c.length}s=null,l=!1,function(t){if(r===clearTimeout)return clearTimeout(t);if((r===o||!r)&&clearTimeout)return r=clearTimeout,clearTimeout(t);try{r(t)}catch(e){try{return r.call(null,t)}catch(e){return r.call(this,t)}}}(t)}}function p(t,e){this.fun=t,this.array=e}function m(){}i.nextTick=function(t){var e=new Array(arguments.length-1);if(arguments.length>1)for(var n=1;n<arguments.length;n++)e[n-1]=arguments[n];c.push(new p(t,e)),1!==c.length||l||u(h)},p.prototype.run=function(){this.fun.apply(null,this.array)},i.title="browser",i.browser=!0,i.env={},i.argv=[],i.version="",i.versions={},i.on=m,i.addListener=m,i.once=m,i.off=m,i.removeListener=m,i.removeAllListeners=m,i.emit=m,i.prependListener=m,i.prependOnceListener=m,i.listeners=function(t){return[]},i.binding=function(t){throw new Error("process.binding is not supported")},i.cwd=function(){return"/"},i.chdir=function(t){throw new Error("process.chdir is not supported")},i.umask=function(){return 0}},function(t,e){t.exports=function(){for(var t={},e=0;e<arguments.length;e++){var r=arguments[e];for(var i in r)n.call(r,i)&&(t[i]=r[i])}return t};var n=Object.prototype.hasOwnProperty},function(t,e){t.exports=function(t){return t&&t.__esModule?t:{default:t}}},function(t,e,n){t.exports=!n(68)((function(){return 7!=Object.defineProperty({},"a",{get:function(){return 7}}).a}))},function(t,e,n){"use strict";n.r(e),n.d(e,"timeFormatDefaultLocale",(function(){return r.a})),n.d(e,"timeFormat",(function(){return r.b})),n.d(e,"timeParse",(function(){return r.c})),n.d(e,"utcFormat",(function(){return r.d})),n.d(e,"utcParse",(function(){return r.e})),n.d(e,"timeFormatLocale",(function(){return i.a})),n.d(e,"isoFormat",(function(){return a})),n.d(e,"isoParse",(function(){return o}));var r=n(41),i=n(167);var a=Date.prototype.toISOString?function(t){return t.toISOString()}:Object(r.d)("%Y-%m-%dT%H:%M:%S.%LZ");var o=+new Date("2000-01-01T00:00:00.000Z")?function(t){var e=new Date(t);return isNaN(e)?null:e}:Object(r.e)("%Y-%m-%dT%H:%M:%S.%LZ")},function(t,e,n){"use strict";n.d(e,"b",(function(){return i})),n.d(e,"c",(function(){return a})),n.d(e,"d",(function(){return o})),n.d(e,"e",(function(){return u})),n.d(e,"a",(function(){return c}));var r,i,a,o,u,s=n(167);function c(t){return r=Object(s.a)(t),i=r.format,a=r.parse,o=r.utcFormat,u=r.utcParse,r}c({dateTime:"%x, %X",date:"%-m/%-d/%Y",time:"%-I:%M:%S %p",periods:["AM","PM"],days:["Sunday","Monday","Tuesday","Wednesday","Thursday","Friday","Saturday"],shortDays:["Sun","Mon","Tue","Wed","Thu","Fri","Sat"],months:["January","February","March","April","May","June","July","August","September","October","November","December"],shortMonths:["Jan","Feb","Mar","Apr","May","Jun","Jul","Aug","Sep","Oct","Nov","Dec"]})},function(t,e,n){"use strict";n.d(e,"c",(function(){return i})),n.d(e,"a",(function(){return l}));var r={},i=null;"undefined"!=typeof document&&("onmouseenter"in document.documentElement||(r={mouseenter:"mouseover",mouseleave:"mouseout"}));function a(t,e,n){return t=o(t,e,n),function(e){var n=e.relatedTarget;n&&(n===this||8&n.compareDocumentPosition(this))||t.call(this,e)}}function o(t,e,n){return function(r){var a=i;i=r;try{t.call(this,this.__data__,e,n)}finally{i=a}}}function u(t){return t.trim().split(/^|\s+/).map((function(t){var e="",n=t.indexOf(".");return n>=0&&(e=t.slice(n+1),t=t.slice(0,n)),{type:t,name:e}}))}function s(t){return function(){var e=this.__on;if(e){for(var n,r=0,i=-1,a=e.length;r<a;++r)n=e[r],t.type&&n.type!==t.type||n.name!==t.name?e[++i]=n:this.removeEventListener(n.type,n.listener,n.capture);++i?e.length=i:delete this.__on}}}function c(t,e,n){var i=r.hasOwnProperty(t.type)?a:o;return function(r,a,o){var u,s=this.__on,c=i(e,a,o);if(s)for(var l=0,f=s.length;l<f;++l)if((u=s[l]).type===t.type&&u.name===t.name)return this.removeEventListener(u.type,u.listener,u.capture),this.addEventListener(u.type,u.listener=c,u.capture=n),void(u.value=e);this.addEventListener(t.type,c,n),u={type:t.type,name:t.name,value:e,listener:c,capture:n},s?s.push(u):this.__on=[u]}}function l(t,e,n,r){var a=i;t.sourceEvent=i,i=t;try{return e.apply(n,r)}finally{i=a}}e.b=function(t,e,n){var r,i,a=u(t+""),o=a.length;if(!(arguments.length<2)){for(l=e?c:s,null==n&&(n=!1),r=0;r<o;++r)this.each(l(a[r],e,n));return this}var l=this.node().__on;if(l)for(var f,d=0,h=l.length;d<h;++d)for(r=0,f=l[d];r<o;++r)if((i=a[r]).type===f.type&&i.name===f.name)return f.value}},function(t,e,n){"use strict";var r=n(79);e.a=function(t){return(t=Object(r.a)(Math.abs(t)))?t[1]:NaN}},function(t,e,n){"use strict";n.d(e,"c",(function(){return $})),n.d(e,"a",(function(){return X}));var r=n(96),i=n(162),a=n(161),o=function(t){return new Array(t.length)};function u(t,e){this.ownerDocument=t.ownerDocument,this.namespaceURI=t.namespaceURI,this._next=null,this._parent=t,this.__data__=e}u.prototype={constructor:u,appendChild:function(t){return this._parent.insertBefore(t,this._next)},insertBefore:function(t,e){return this._parent.insertBefore(t,e)},querySelector:function(t){return this._parent.querySelector(t)},querySelectorAll:function(t){return this._parent.querySelectorAll(t)}};function s(t,e,n,r,i,a){for(var o,s=0,c=e.length,l=a.length;s<l;++s)(o=e[s])?(o.__data__=a[s],r[s]=o):n[s]=new u(t,a[s]);for(;s<c;++s)(o=e[s])&&(i[s]=o)}function c(t,e,n,r,i,a,o){var s,c,l,f={},d=e.length,h=a.length,p=new Array(d);for(s=0;s<d;++s)(c=e[s])&&(p[s]=l="$"+o.call(c,c.__data__,s,e),l in f?i[s]=c:f[l]=c);for(s=0;s<h;++s)(c=f[l="$"+o.call(t,a[s],s,a)])?(r[s]=c,c.__data__=a[s],f[l]=null):n[s]=new u(t,a[s]);for(s=0;s<d;++s)(c=e[s])&&f[p[s]]===c&&(i[s]=c)}function l(t,e){return t<e?-1:t>e?1:t>=e?0:NaN}var f=n(95);function d(t){return function(){this.removeAttribute(t)}}function h(t){return function(){this.removeAttributeNS(t.space,t.local)}}function p(t,e){return function(){this.setAttribute(t,e)}}function m(t,e){return function(){this.setAttributeNS(t.space,t.local,e)}}function v(t,e){return function(){var n=e.apply(this,arguments);null==n?this.removeAttribute(t):this.setAttribute(t,n)}}function g(t,e){return function(){var n=e.apply(this,arguments);null==n?this.removeAttributeNS(t.space,t.local):this.setAttributeNS(t.space,t.local,n)}}var _=n(163);function y(t){return function(){delete this[t]}}function b(t,e){return function(){this[t]=e}}function w(t,e){return function(){var n=e.apply(this,arguments);null==n?delete this[t]:this[t]=n}}function x(t){return t.trim().split(/^|\s+/)}function k(t){return t.classList||new M(t)}function M(t){this._node=t,this._names=x(t.getAttribute("class")||"")}function E(t,e){for(var n=k(t),r=-1,i=e.length;++r<i;)n.add(e[r])}function S(t,e){for(var n=k(t),r=-1,i=e.length;++r<i;)n.remove(e[r])}function T(t){return function(){E(this,t)}}function j(t){return function(){S(this,t)}}function O(t,e){return function(){(e.apply(this,arguments)?E:S)(this,t)}}M.prototype={add:function(t){this._names.indexOf(t)<0&&(this._names.push(t),this._node.setAttribute("class",this._names.join(" ")))},remove:function(t){var e=this._names.indexOf(t);e>=0&&(this._names.splice(e,1),this._node.setAttribute("class",this._names.join(" ")))},contains:function(t){return this._names.indexOf(t)>=0}};function A(){this.textContent=""}function L(t){return function(){this.textContent=t}}function C(t){return function(){var e=t.apply(this,arguments);this.textContent=null==e?"":e}}function D(){this.innerHTML=""}function N(t){return function(){this.innerHTML=t}}function P(t){return function(){var e=t.apply(this,arguments);this.innerHTML=null==e?"":e}}function R(){this.nextSibling&&this.parentNode.appendChild(this)}function Y(){this.previousSibling&&this.parentNode.insertBefore(this,this.parentNode.firstChild)}var I=n(59);function z(){return null}function F(){var t=this.parentNode;t&&t.removeChild(this)}function H(){var t=this.cloneNode(!1),e=this.parentNode;return e?e.insertBefore(t,this.nextSibling):t}function B(){var t=this.cloneNode(!0),e=this.parentNode;return e?e.insertBefore(t,this.nextSibling):t}var q=n(42),U=n(97);function W(t,e,n){var r=Object(U.a)(t),i=r.CustomEvent;"function"==typeof i?i=new i(e,n):(i=r.document.createEvent("Event"),n?(i.initEvent(e,n.bubbles,n.cancelable),i.detail=n.detail):i.initEvent(e,!1,!1)),t.dispatchEvent(i)}function G(t,e){return function(){return W(this,t,e)}}function V(t,e){return function(){return W(this,t,e.apply(this,arguments))}}var $=[null];function X(t,e){this._groups=t,this._parents=e}function J(){return new X([[document.documentElement]],$)}X.prototype=J.prototype={constructor:X,select:function(t){"function"!=typeof t&&(t=Object(r.a)(t));for(var e=this._groups,n=e.length,i=new Array(n),a=0;a<n;++a)for(var o,u,s=e[a],c=s.length,l=i[a]=new Array(c),f=0;f<c;++f)(o=s[f])&&(u=t.call(o,o.__data__,f,s))&&("__data__"in o&&(u.__data__=o.__data__),l[f]=u);return new X(i,this._parents)},selectAll:function(t){"function"!=typeof t&&(t=Object(i.a)(t));for(var e=this._groups,n=e.length,r=[],a=[],o=0;o<n;++o)for(var u,s=e[o],c=s.length,l=0;l<c;++l)(u=s[l])&&(r.push(t.call(u,u.__data__,l,s)),a.push(u));return new X(r,a)},filter:function(t){"function"!=typeof t&&(t=Object(a.a)(t));for(var e=this._groups,n=e.length,r=new Array(n),i=0;i<n;++i)for(var o,u=e[i],s=u.length,c=r[i]=[],l=0;l<s;++l)(o=u[l])&&t.call(o,o.__data__,l,u)&&c.push(o);return new X(r,this._parents)},data:function(t,e){if(!t)return v=new Array(this.size()),d=-1,this.each((function(t){v[++d]=t})),v;var n,r=e?c:s,i=this._parents,a=this._groups;"function"!=typeof t&&(n=t,t=function(){return n});for(var o=a.length,u=new Array(o),l=new Array(o),f=new Array(o),d=0;d<o;++d){var h=i[d],p=a[d],m=p.length,v=t.call(h,h&&h.__data__,d,i),g=v.length,_=l[d]=new Array(g),y=u[d]=new Array(g);r(h,p,_,y,f[d]=new Array(m),v,e);for(var b,w,x=0,k=0;x<g;++x)if(b=_[x]){for(x>=k&&(k=x+1);!(w=y[k])&&++k<g;);b._next=w||null}}return(u=new X(u,i))._enter=l,u._exit=f,u},enter:function(){return new X(this._enter||this._groups.map(o),this._parents)},exit:function(){return new X(this._exit||this._groups.map(o),this._parents)},join:function(t,e,n){var r=this.enter(),i=this,a=this.exit();return r="function"==typeof t?t(r):r.append(t+""),null!=e&&(i=e(i)),null==n?a.remove():n(a),r&&i?r.merge(i).order():i},merge:function(t){for(var e=this._groups,n=t._groups,r=e.length,i=n.length,a=Math.min(r,i),o=new Array(r),u=0;u<a;++u)for(var s,c=e[u],l=n[u],f=c.length,d=o[u]=new Array(f),h=0;h<f;++h)(s=c[h]||l[h])&&(d[h]=s);for(;u<r;++u)o[u]=e[u];return new X(o,this._parents)},order:function(){for(var t=this._groups,e=-1,n=t.length;++e<n;)for(var r,i=t[e],a=i.length-1,o=i[a];--a>=0;)(r=i[a])&&(o&&4^r.compareDocumentPosition(o)&&o.parentNode.insertBefore(r,o),o=r);return this},sort:function(t){function e(e,n){return e&&n?t(e.__data__,n.__data__):!e-!n}t||(t=l);for(var n=this._groups,r=n.length,i=new Array(r),a=0;a<r;++a){for(var o,u=n[a],s=u.length,c=i[a]=new Array(s),f=0;f<s;++f)(o=u[f])&&(c[f]=o);c.sort(e)}return new X(i,this._parents).order()},call:function(){var t=arguments[0];return arguments[0]=this,t.apply(null,arguments),this},nodes:function(){var t=new Array(this.size()),e=-1;return this.each((function(){t[++e]=this})),t},node:function(){for(var t=this._groups,e=0,n=t.length;e<n;++e)for(var r=t[e],i=0,a=r.length;i<a;++i){var o=r[i];if(o)return o}return null},size:function(){var t=0;return this.each((function(){++t})),t},empty:function(){return!this.node()},each:function(t){for(var e=this._groups,n=0,r=e.length;n<r;++n)for(var i,a=e[n],o=0,u=a.length;o<u;++o)(i=a[o])&&t.call(i,i.__data__,o,a);return this},attr:function(t,e){var n=Object(f.a)(t);if(arguments.length<2){var r=this.node();return n.local?r.getAttributeNS(n.space,n.local):r.getAttribute(n)}return this.each((null==e?n.local?h:d:"function"==typeof e?n.local?g:v:n.local?m:p)(n,e))},style:_.a,property:function(t,e){return arguments.length>1?this.each((null==e?y:"function"==typeof e?w:b)(t,e)):this.node()[t]},classed:function(t,e){var n=x(t+"");if(arguments.length<2){for(var r=k(this.node()),i=-1,a=n.length;++i<a;)if(!r.contains(n[i]))return!1;return!0}return this.each(("function"==typeof e?O:e?T:j)(n,e))},text:function(t){return arguments.length?this.each(null==t?A:("function"==typeof t?C:L)(t)):this.node().textContent},html:function(t){return arguments.length?this.each(null==t?D:("function"==typeof t?P:N)(t)):this.node().innerHTML},raise:function(){return this.each(R)},lower:function(){return this.each(Y)},append:function(t){var e="function"==typeof t?t:Object(I.a)(t);return this.select((function(){return this.appendChild(e.apply(this,arguments))}))},insert:function(t,e){var n="function"==typeof t?t:Object(I.a)(t),i=null==e?z:"function"==typeof e?e:Object(r.a)(e);return this.select((function(){return this.insertBefore(n.apply(this,arguments),i.apply(this,arguments)||null)}))},remove:function(){return this.each(F)},clone:function(t){return this.select(t?B:H)},datum:function(t){return arguments.length?this.property("__data__",t):this.node().__data__},on:q.b,dispatch:function(t,e){return this.each(("function"==typeof e?V:G)(t,e))}};e.b=J},function(t,e,n){"use strict";var r=n(252);n.d(e,"a",(function(){return r.a}))},function(t,e,n){"use strict";var r=n(7);n.d(e,"g",(function(){return r.a}));var i=n(113);n.d(e,"h",(function(){return i.a})),n.d(e,"i",(function(){return i.b})),n.d(e,"L",(function(){return i.a})),n.d(e,"M",(function(){return i.b}));var a=n(112);n.d(e,"r",(function(){return a.a})),n.d(e,"s",(function(){return a.b})),n.d(e,"V",(function(){return a.a})),n.d(e,"W",(function(){return a.b}));var o=n(182);n.d(e,"j",(function(){return o.a})),n.d(e,"k",(function(){return o.b}));var u=n(181);n.d(e,"e",(function(){return u.a})),n.d(e,"f",(function(){return u.b}));var s=n(127);n.d(e,"a",(function(){return s.b})),n.d(e,"b",(function(){return s.a}));var c=n(12);n.d(e,"B",(function(){return c.g})),n.d(e,"C",(function(){return c.h})),n.d(e,"t",(function(){return c.g})),n.d(e,"u",(function(){return c.h})),n.d(e,"l",(function(){return c.c})),n.d(e,"m",(function(){return c.d})),n.d(e,"x",(function(){return c.k})),n.d(e,"y",(function(){return c.l})),n.d(e,"z",(function(){return c.m})),n.d(e,"A",(function(){return c.n})),n.d(e,"v",(function(){return c.i})),n.d(e,"w",(function(){return c.j})),n.d(e,"c",(function(){return c.a})),n.d(e,"d",(function(){return c.b})),n.d(e,"p",(function(){return c.e})),n.d(e,"q",(function(){return c.f}));var l=n(180);n.d(e,"n",(function(){return l.a})),n.d(e,"o",(function(){return l.b}));var f=n(64);n.d(e,"D",(function(){return f.a})),n.d(e,"E",(function(){return f.b}));var d=n(185);n.d(e,"N",(function(){return d.a})),n.d(e,"O",(function(){return d.b}));var h=n(184);n.d(e,"J",(function(){return h.a})),n.d(e,"K",(function(){return h.b}));var p=n(128);n.d(e,"F",(function(){return p.a})),n.d(e,"G",(function(){return p.b}));var m=n(13);n.d(e,"fb",(function(){return m.g})),n.d(e,"gb",(function(){return m.h})),n.d(e,"X",(function(){return m.g})),n.d(e,"Y",(function(){return m.h})),n.d(e,"P",(function(){return m.c})),n.d(e,"Q",(function(){return m.d})),n.d(e,"bb",(function(){return m.k})),n.d(e,"cb",(function(){return m.l})),n.d(e,"db",(function(){return m.m})),n.d(e,"eb",(function(){return m.n})),n.d(e,"Z",(function(){return m.i})),n.d(e,"ab",(function(){return m.j})),n.d(e,"H",(function(){return m.a})),n.d(e,"I",(function(){return m.b})),n.d(e,"T",(function(){return m.e})),n.d(e,"U",(function(){return m.f}));var v=n(183);n.d(e,"R",(function(){return v.a})),n.d(e,"S",(function(){return v.b}));var g=n(65);n.d(e,"hb",(function(){return g.a})),n.d(e,"ib",(function(){return g.b}))},function(t,e,n){var r=n(31),i=n(264),a=n(193),o=Object.defineProperty;e.f=n(39)?Object.defineProperty:function(t,e,n){if(r(t),e=a(e,!0),r(n),i)try{return o(t,e,n)}catch(t){}if("get"in n||"set"in n)throw TypeError("Accessors not supported!");return"value"in n&&(t[e]=n.value),t}},function(t,e,n){t.exports=n(513)()},function(t,e,n){"use strict";n.d(e,"b",(function(){return h})),n.d(e,"a",(function(){return m})),n.d(e,"c",(function(){return v})),n.d(e,"d",(function(){return g}));var r,i,a=0,o=0,u=0,s=0,c=0,l=0,f="object"==typeof performance&&performance.now?performance:Date,d="object"==typeof window&&window.requestAnimationFrame?window.requestAnimationFrame.bind(window):function(t){setTimeout(t,17)};function h(){return c||(d(p),c=f.now()+l)}function p(){c=0}function m(){this._call=this._time=this._next=null}function v(t,e,n){var r=new m;return r.restart(t,e,n),r}function g(){h(),++a;for(var t,e=r;e;)(t=c-e._time)>=0&&e._call.call(null,t),e=e._next;--a}function _(){c=(s=f.now())+l,a=o=0;try{g()}finally{a=0,function(){var t,e,n=r,a=1/0;for(;n;)n._call?(a>n._time&&(a=n._time),t=n,n=n._next):(e=n._next,n._next=null,n=t?t._next=e:r=e);i=t,b(a)}(),c=0}}function y(){var t=f.now(),e=t-s;e>1e3&&(l-=e,s=t)}function b(t){a||(o&&(o=clearTimeout(o)),t-c>24?(t<1/0&&(o=setTimeout(_,t-f.now()-l)),u&&(u=clearInterval(u))):(u||(s=f.now(),u=setInterval(y,1e3)),a=1,d(_)))}m.prototype=v.prototype={constructor:m,restart:function(t,e,n){if("function"!=typeof t)throw new TypeError("callback is not a function");n=(null==n?h():+n)+(null==e?0:+e),this._next||i===this||(i?i._next=this:r=this,i=this),this._call=t,this._time=n,b()},stop:function(){this._call&&(this._call=null,this._time=1/0,b())}}},function(t,e,n){"use strict";function r(t){return t[0]}function i(t){return t[1]}n.d(e,"a",(function(){return r})),n.d(e,"b",(function(){return i}))},function(t,e,n){"use strict";n.d(e,"f",(function(){return a})),n.d(e,"g",(function(){return o})),n.d(e,"a",(function(){return u})),n.d(e,"b",(function(){return s})),n.d(e,"d",(function(){return c})),n.d(e,"c",(function(){return l})),n.d(e,"e",(function(){return f}));var r=n(100),i=Object(r.a)(","),a=i.parse,o=i.parseRows,u=i.format,s=i.formatBody,c=i.formatRows,l=i.formatRow,f=i.formatValue},function(t,e,n){"use strict";n.d(e,"f",(function(){return a})),n.d(e,"g",(function(){return o})),n.d(e,"a",(function(){return u})),n.d(e,"b",(function(){return s})),n.d(e,"d",(function(){return c})),n.d(e,"c",(function(){return l})),n.d(e,"e",(function(){return f}));var r=n(100),i=Object(r.a)("\t"),a=i.parse,o=i.parseRows,u=i.format,s=i.formatBody,c=i.formatRows,l=i.formatRow,f=i.formatValue},function(t,e,n){"use strict";var r=n(134);n.d(e,"c",(function(){return r.a})),n.d(e,"b",(function(){return r.b})),n.d(e,"e",(function(){return r.c}));var i=n(169);n.d(e,"d",(function(){return i.a}));var a=n(101);n.d(e,"f",(function(){return a.b})),n.d(e,"a",(function(){return a.a}));var o=n(256);n.d(e,"g",(function(){return o.a}));var u=n(257);n.d(e,"h",(function(){return u.a}));var s=n(258);n.d(e,"i",(function(){return s.a}))},function(t,e){t.exports=function(t){return"object"==typeof t?null!==t:"function"==typeof t}},function(t,e,n){var r=n(268),i=n(136);t.exports=function(t){return r(i(t))}},function(t,e){t.exports=function(t){return t&&t.__esModule?t:{default:t}}},function(t,e,n){"use strict";var r="undefined"!=typeof Uint8Array&&"undefined"!=typeof Uint16Array&&"undefined"!=typeof Int32Array;function i(t,e){return Object.prototype.hasOwnProperty.call(t,e)}e.assign=function(t){for(var e=Array.prototype.slice.call(arguments,1);e.length;){var n=e.shift();if(n){if("object"!=typeof n)throw new TypeError(n+"must be non-object");for(var r in n)i(n,r)&&(t[r]=n[r])}}return t},e.shrinkBuf=function(t,e){return t.length===e?t:t.subarray?t.subarray(0,e):(t.length=e,t)};var a={arraySet:function(t,e,n,r,i){if(e.subarray&&t.subarray)t.set(e.subarray(n,n+r),i);else for(var a=0;a<r;a++)t[i+a]=e[n+a]},flattenChunks:function(t){var e,n,r,i,a,o;for(r=0,e=0,n=t.length;e<n;e++)r+=t[e].length;for(o=new Uint8Array(r),i=0,e=0,n=t.length;e<n;e++)a=t[e],o.set(a,i),i+=a.length;return o}},o={arraySet:function(t,e,n,r,i){for(var a=0;a<r;a++)t[i+a]=e[n+a]},flattenChunks:function(t){return[].concat.apply([],t)}};e.setTyped=function(t){t?(e.Buf8=Uint8Array,e.Buf16=Uint16Array,e.Buf32=Int32Array,e.assign(e,a)):(e.Buf8=Array,e.Buf16=Array,e.Buf32=Array,e.assign(e,o))},e.setTyped(r)},function(t,e,n){"use strict";var r=n(1),i=n(676);if(void 0===r)throw Error("create-react-class could not find the React object. If you are using script tags, make sure that React is being loaded before create-react-class.");var a=(new r.Component).updater;t.exports=i(r.Component,r.isValidElement,a)},function(t,e,n){"use strict";var r=n(95),i=n(62);function a(t){return function(){var e=this.ownerDocument,n=this.namespaceURI;return n===i.b&&e.documentElement.namespaceURI===i.b?e.createElement(t):e.createElementNS(n,t)}}function o(t){return function(){return this.ownerDocument.createElementNS(t.space,t.local)}}e.a=function(t){var e=Object(r.a)(t);return(e.local?o:a)(e)}},function(t,e,n){"use strict";e.a=function(t,e){var n=t.ownerSVGElement||t;if(n.createSVGPoint){var r=n.createSVGPoint();return r.x=e.clientX,r.y=e.clientY,[(r=r.matrixTransform(t.getScreenCTM().inverse())).x,r.y]}var i=t.getBoundingClientRect();return[e.clientX-i.left-t.clientLeft,e.clientY-i.top-t.clientTop]}},function(t,e,n){"use strict";function r(t){this._context=t}r.prototype={areaStart:function(){this._line=0},areaEnd:function(){this._line=NaN},lineStart:function(){this._point=0},lineEnd:function(){(this._line||0!==this._line&&1===this._point)&&this._context.closePath(),this._line=1-this._line},point:function(t,e){switch(t=+t,e=+e,this._point){case 0:this._point=1,this._line?this._context.lineTo(t,e):this._context.moveTo(t,e);break;case 1:this._point=2;default:this._context.lineTo(t,e)}}},e.a=function(t){return new r(t)}},function(t,e,n){"use strict";n.d(e,"b",(function(){return r}));var r="http://www.w3.org/1999/xhtml";e.a={svg:"http://www.w3.org/2000/svg",xhtml:r,xlink:"http://www.w3.org/1999/xlink",xml:"http://www.w3.org/XML/1998/namespace",xmlns:"http://www.w3.org/2000/xmlns/"}},function(t,e,n){"use strict";function r(t){return ArrayBuffer.isView(t)&&!(t instanceof DataView)}n.d(e,"b",(function(){return r})),e.a=function(t,e){e||(e=[]);var n,r=t?Math.min(e.length,t.length):0,i=e.slice();return function(a){for(n=0;n<r;++n)i[n]=t[n]*(1-a)+e[n]*a;return i}}},function(t,e,n){"use strict";n.d(e,"b",(function(){return a}));var r=n(7),i=Object(r.a)((function(t){t.setMonth(0,1),t.setHours(0,0,0,0)}),(function(t,e){t.setFullYear(t.getFullYear()+e)}),(function(t,e){return e.getFullYear()-t.getFullYear()}),(function(t){return t.getFullYear()}));i.every=function(t){return isFinite(t=Math.floor(t))&&t>0?Object(r.a)((function(e){e.setFullYear(Math.floor(e.getFullYear()/t)*t),e.setMonth(0,1),e.setHours(0,0,0,0)}),(function(e,n){e.setFullYear(e.getFullYear()+n*t)})):null},e.a=i;var a=i.range},function(t,e,n){"use strict";n.d(e,"b",(function(){return a}));var r=n(7),i=Object(r.a)((function(t){t.setUTCMonth(0,1),t.setUTCHours(0,0,0,0)}),(function(t,e){t.setUTCFullYear(t.getUTCFullYear()+e)}),(function(t,e){return e.getUTCFullYear()-t.getUTCFullYear()}),(function(t){return t.getUTCFullYear()}));i.every=function(t){return isFinite(t=Math.floor(t))&&t>0?Object(r.a)((function(e){e.setUTCFullYear(Math.floor(e.getUTCFullYear()/t)*t),e.setUTCMonth(0,1),e.setUTCHours(0,0,0,0)}),(function(e,n){e.setUTCFullYear(e.getUTCFullYear()+n*t)})):null},e.a=i;var a=i.range},function(t,e){(function(e){t.exports=e}).call(this,{})},function(t,e,n){var r=n(47),i=n(114);t.exports=n(39)?function(t,e,n){return r.f(t,e,i(1,n))}:function(t,e,n){return t[e]=n,t}},function(t,e){t.exports=function(t){try{return!!t()}catch(t){return!0}}},function(t,e){var n={}.hasOwnProperty;t.exports=function(t,e){return n.call(t,e)}},function(t,e,n){var r=n(21);t.exports=function(t,e,n){return e in t?r(t,e,{value:n,enumerable:!0,configurable:!0,writable:!0}):t[e]=n,t}},function(t,e){t.exports=function(t,e){if(!(t instanceof e))throw new TypeError("Cannot call a class as a function")}},function(t,e,n){var r=n(21);function i(t,e){for(var n=0;n<e.length;n++){var i=e[n];i.enumerable=i.enumerable||!1,i.configurable=!0,"value"in i&&(i.writable=!0),r(t,i.key,i)}}t.exports=function(t,e,n){return e&&i(t.prototype,e),n&&i(t,n),t}},function(t,e,n){t.exports=n(275)},function(t,e){function n(t,e,n,r,i,a,o){try{var u=t[a](o),s=u.value}catch(t){return void n(t)}u.done?e(s):Promise.resolve(s).then(r,i)}t.exports=function(t){return function(){var e=this,r=arguments;return new Promise((function(i,a){var o=t.apply(e,r);function u(t){n(o,i,a,u,s,"next",t)}function s(t){n(o,i,a,u,s,"throw",t)}u(void 0)}))}}},function(t,e){"function"==typeof Object.create?t.exports=function(t,e){e&&(t.super_=e,t.prototype=Object.create(e.prototype,{constructor:{value:t,enumerable:!1,writable:!0,configurable:!0}}))}:t.exports=function(t,e){if(e){t.super_=e;var n=function(){};n.prototype=e.prototype,t.prototype=new n,t.prototype.constructor=t}}},function(t,e,n){"use strict";var r=n(143),i=Object.keys||function(t){var e=[];for(var n in t)e.push(n);return e};t.exports=f;var a=Object.create(n(122));a.inherits=n(75);var o=n(306),u=n(213);a.inherits(f,o);for(var s=i(u.prototype),c=0;c<s.length;c++){var l=s[c];f.prototype[l]||(f.prototype[l]=u.prototype[l])}function f(t){if(!(this instanceof f))return new f(t);o.call(this,t),u.call(this,t),t&&!1===t.readable&&(this.readable=!1),t&&!1===t.writable&&(this.writable=!1),this.allowHalfOpen=!0,t&&!1===t.allowHalfOpen&&(this.allowHalfOpen=!1),this.once("end",d)}function d(){this.allowHalfOpen||this._writableState.ended||r.nextTick(h,this)}function h(t){t.end()}Object.defineProperty(f.prototype,"writableHighWaterMark",{enumerable:!1,get:function(){return this._writableState.highWaterMark}}),Object.defineProperty(f.prototype,"destroyed",{get:function(){return void 0!==this._readableState&&void 0!==this._writableState&&(this._readableState.destroyed&&this._writableState.destroyed)},set:function(t){void 0!==this._readableState&&void 0!==this._writableState&&(this._readableState.destroyed=t,this._writableState.destroyed=t)}}),f.prototype._destroy=function(t,e){this.push(null),this.end(),r.nextTick(e,t)}},function(t,e,n){"use strict";n.d(e,"b",(function(){return r.b})),n.d(e,"d",(function(){return r.c})),n.d(e,"e",(function(){return r.d})),n.d(e,"c",(function(){return i.a})),n.d(e,"a",(function(){return a}));var r=n(49),i=n(253),a=function(t,e,n){var i=new r.a,a=e;return null==e?(i.restart(t,e,n),i):(e=+e,n=null==n?Object(r.b)():+n,i.restart((function r(o){o+=a,i.restart(r,a+=e,n),t(o)}),e,n),i)}},function(t,e,n){"use strict";n.d(e,"b",(function(){return s})),n.d(e,"c",(function(){return c}));var r=n(14),i=n(94),a=n(159),o=n(10);function u(t){return function(e){var n,i,a=e.length,o=new Array(a),u=new Array(a),s=new Array(a);for(n=0;n<a;++n)i=Object(r.g)(e[n]),o[n]=i.r||0,u[n]=i.g||0,s[n]=i.b||0;return o=t(o),u=t(u),s=t(s),i.opacity=1,function(t){return i.r=o(t),i.g=u(t),i.b=s(t),i+""}}}e.a=function t(e){var n=Object(o.b)(e);function i(t,e){var i=n((t=Object(r.g)(t)).r,(e=Object(r.g)(e)).r),a=n(t.g,e.g),u=n(t.b,e.b),s=Object(o.a)(t.opacity,e.opacity);return function(e){return t.r=i(e),t.g=a(e),t.b=u(e),t.opacity=s(e),t+""}}return i.gamma=t,i}(1);var s=u(i.b),c=u(a.a)},function(t,e,n){"use strict";e.a=function(t,e){if((n=(t=e?t.toExponential(e-1):t.toExponential()).indexOf("e"))<0)return null;var n,r=t.slice(0,n);return[r.length>1?r[0]+r.slice(2):r,+t.slice(n+1)]}},function(t,e,n){"use strict";n.d(e,"a",(function(){return d})),n.d(e,"l",(function(){return p})),n.d(e,"g",(function(){return R})),n.d(e,"i",(function(){return P})),n.d(e,"j",(function(){return W})),n.d(e,"w",(function(){return X})),n.d(e,"k",(function(){return f})),n.d(e,"h",(function(){return l})),n.d(e,"m",(function(){return tt})),n.d(e,"v",(function(){return et})),n.d(e,"n",(function(){return nt})),n.d(e,"o",(function(){return rt})),n.d(e,"x",(function(){return it})),n.d(e,"y",(function(){return gt})),n.d(e,"z",(function(){return Mt})),n.d(e,"p",(function(){return Tt})),n.d(e,"q",(function(){return jt})),n.d(e,"r",(function(){return At})),n.d(e,"t",(function(){return Lt})),n.d(e,"u",(function(){return Ot})),n.d(e,"s",(function(){return Ct})),n.d(e,"b",(function(){return Nt})),n.d(e,"c",(function(){return Pt})),n.d(e,"d",(function(){return Yt})),n.d(e,"e",(function(){return It})),n.d(e,"f",(function(){return Rt})),n.d(e,"A",(function(){return D}));var r=n(2);function i(t,e){switch(arguments.length){case 0:break;case 1:this.range(t);break;default:this.range(e).domain(t)}return this}function a(t,e){switch(arguments.length){case 0:break;case 1:this.interpolator(t);break;default:this.interpolator(e).domain(t)}return this}var o=n(24),u=Array.prototype,s=u.map,c=u.slice,l={name:"implicit"};function f(){var t=Object(o.c)(),e=[],n=[],r=l;function a(i){var a=i+"",o=t.get(a);if(!o){if(r!==l)return r;t.set(a,o=e.push(i))}return n[(o-1)%n.length]}return a.domain=function(n){if(!arguments.length)return e.slice();e=[],t=Object(o.c)();for(var r,i,u=-1,s=n.length;++u<s;)t.has(i=(r=n[u])+"")||t.set(i,e.push(r));return a},a.range=function(t){return arguments.length?(n=c.call(t),a):n.slice()},a.unknown=function(t){return arguments.length?(r=t,a):r},a.copy=function(){return f(e,n).unknown(r)},i.apply(a,arguments),a}function d(){var t,e,n=f().unknown(void 0),a=n.domain,o=n.range,u=[0,1],s=!1,c=0,l=0,h=.5;function p(){var n=a().length,i=u[1]<u[0],f=u[i-0],d=u[1-i];t=(d-f)/Math.max(1,n-c+2*l),s&&(t=Math.floor(t)),f+=(d-f-t*(n-c))*h,e=t*(1-c),s&&(f=Math.round(f),e=Math.round(e));var p=Object(r.s)(n).map((function(e){return f+t*e}));return o(i?p.reverse():p)}return delete n.unknown,n.domain=function(t){return arguments.length?(a(t),p()):a()},n.range=function(t){return arguments.length?(u=[+t[0],+t[1]],p()):u.slice()},n.rangeRound=function(t){return u=[+t[0],+t[1]],s=!0,p()},n.bandwidth=function(){return e},n.step=function(){return t},n.round=function(t){return arguments.length?(s=!!t,p()):s},n.padding=function(t){return arguments.length?(c=Math.min(1,l=+t),p()):c},n.paddingInner=function(t){return arguments.length?(c=Math.min(1,t),p()):c},n.paddingOuter=function(t){return arguments.length?(l=+t,p()):l},n.align=function(t){return arguments.length?(h=Math.max(0,Math.min(1,t)),p()):h},n.copy=function(){return d(a(),u).round(s).paddingInner(c).paddingOuter(l).align(h)},i.apply(p(),arguments)}function h(t){var e=t.copy;return t.padding=t.paddingOuter,delete t.paddingInner,delete t.paddingOuter,t.copy=function(){return h(e())},t}function p(){return h(d.apply(null,arguments).paddingInner(1))}var m=n(98),v=n(25),g=n(259),_=function(t){return+t},y=[0,1];function b(t){return t}function w(t,e){return(e-=t=+t)?function(n){return(n-t)/e}:(n=isNaN(e)?NaN:.5,function(){return n});var n}function x(t){var e,n=t[0],r=t[t.length-1];return n>r&&(e=n,n=r,r=e),function(t){return Math.max(n,Math.min(r,t))}}function k(t,e,n){var r=t[0],i=t[1],a=e[0],o=e[1];return i<r?(r=w(i,r),a=n(o,a)):(r=w(r,i),a=n(a,o)),function(t){return a(r(t))}}function M(t,e,n){var i=Math.min(t.length,e.length)-1,a=new Array(i),o=new Array(i),u=-1;for(t[i]<t[0]&&(t=t.slice().reverse(),e=e.slice().reverse());++u<i;)a[u]=w(t[u],t[u+1]),o[u]=n(e[u],e[u+1]);return function(e){var n=Object(r.b)(t,e,1,i)-1;return o[n](a[n](e))}}function E(t,e){return e.domain(t.domain()).range(t.range()).interpolate(t.interpolate()).clamp(t.clamp()).unknown(t.unknown())}function S(){var t,e,n,r,i,a,o=y,u=y,l=m.a,f=b;function d(){return r=Math.min(o.length,u.length)>2?M:k,i=a=null,h}function h(e){return isNaN(e=+e)?n:(i||(i=r(o.map(t),u,l)))(t(f(e)))}return h.invert=function(n){return f(e((a||(a=r(u,o.map(t),v.a)))(n)))},h.domain=function(t){return arguments.length?(o=s.call(t,_),f===b||(f=x(o)),d()):o.slice()},h.range=function(t){return arguments.length?(u=c.call(t),d()):u.slice()},h.rangeRound=function(t){return u=c.call(t),l=g.a,d()},h.clamp=function(t){return arguments.length?(f=t?x(o):b,h):f!==b},h.interpolate=function(t){return arguments.length?(l=t,d()):l},h.unknown=function(t){return arguments.length?(n=t,h):n},function(n,r){return t=n,e=r,d()}}function T(t,e){return S()(t,e)}var j=n(101),O=n(257),A=n(134),L=n(258),C=n(256),D=function(t,e,n,i){var a,o=Object(r.A)(t,e,n);switch((i=Object(j.b)(null==i?",f":i)).type){case"s":var u=Math.max(Math.abs(t),Math.abs(e));return null!=i.precision||isNaN(a=Object(O.a)(o,u))||(i.precision=a),Object(A.c)(i,u);case"":case"e":case"g":case"p":case"r":null!=i.precision||isNaN(a=Object(L.a)(o,Math.max(Math.abs(t),Math.abs(e))))||(i.precision=a-("e"===i.type));break;case"f":case"%":null!=i.precision||isNaN(a=Object(C.a)(o))||(i.precision=a-2*("%"===i.type))}return Object(A.b)(i)};function N(t){var e=t.domain;return t.ticks=function(t){var n=e();return Object(r.B)(n[0],n[n.length-1],null==t?10:t)},t.tickFormat=function(t,n){var r=e();return D(r[0],r[r.length-1],null==t?10:t,n)},t.nice=function(n){null==n&&(n=10);var i,a=e(),o=0,u=a.length-1,s=a[o],c=a[u];return c<s&&(i=s,s=c,c=i,i=o,o=u,u=i),(i=Object(r.z)(s,c,n))>0?(s=Math.floor(s/i)*i,c=Math.ceil(c/i)*i,i=Object(r.z)(s,c,n)):i<0&&(s=Math.ceil(s*i)/i,c=Math.floor(c*i)/i,i=Object(r.z)(s,c,n)),i>0?(a[o]=Math.floor(s/i)*i,a[u]=Math.ceil(c/i)*i,e(a)):i<0&&(a[o]=Math.ceil(s*i)/i,a[u]=Math.floor(c*i)/i,e(a)),t},t}function P(){var t=T(b,b);return t.copy=function(){return E(t,P())},i.apply(t,arguments),N(t)}function R(t){var e;function n(t){return isNaN(t=+t)?e:t}return n.invert=n,n.domain=n.range=function(e){return arguments.length?(t=s.call(e,_),n):t.slice()},n.unknown=function(t){return arguments.length?(e=t,n):e},n.copy=function(){return R(t).unknown(e)},t=arguments.length?s.call(t,_):[0,1],N(n)}var Y=function(t,e){var n,r=0,i=(t=t.slice()).length-1,a=t[r],o=t[i];return o<a&&(n=r,r=i,i=n,n=a,a=o,o=n),t[r]=e.floor(a),t[i]=e.ceil(o),t};function I(t){return Math.log(t)}function z(t){return Math.exp(t)}function F(t){return-Math.log(-t)}function H(t){return-Math.exp(-t)}function B(t){return isFinite(t)?+("1e"+t):t<0?0:t}function q(t){return function(e){return-t(-e)}}function U(t){var e,n,i=t(I,z),a=i.domain,o=10;function u(){return e=function(t){return t===Math.E?Math.log:10===t&&Math.log10||2===t&&Math.log2||(t=Math.log(t),function(e){return Math.log(e)/t})}(o),n=function(t){return 10===t?B:t===Math.E?Math.exp:function(e){return Math.pow(t,e)}}(o),a()[0]<0?(e=q(e),n=q(n),t(F,H)):t(I,z),i}return i.base=function(t){return arguments.length?(o=+t,u()):o},i.domain=function(t){return arguments.length?(a(t),u()):a()},i.ticks=function(t){var i,u=a(),s=u[0],c=u[u.length-1];(i=c<s)&&(h=s,s=c,c=h);var l,f,d,h=e(s),p=e(c),m=null==t?10:+t,v=[];if(!(o%1)&&p-h<m){if(h=Math.round(h)-1,p=Math.round(p)+1,s>0){for(;h<p;++h)for(f=1,l=n(h);f<o;++f)if(!((d=l*f)<s)){if(d>c)break;v.push(d)}}else for(;h<p;++h)for(f=o-1,l=n(h);f>=1;--f)if(!((d=l*f)<s)){if(d>c)break;v.push(d)}}else v=Object(r.B)(h,p,Math.min(p-h,m)).map(n);return i?v.reverse():v},i.tickFormat=function(t,r){if(null==r&&(r=10===o?".0e":","),"function"!=typeof r&&(r=Object(A.b)(r)),t===1/0)return r;null==t&&(t=10);var a=Math.max(1,o*t/i.ticks().length);return function(t){var i=t/n(Math.round(e(t)));return i*o<o-.5&&(i*=o),i<=a?r(t):""}},i.nice=function(){return a(Y(a(),{floor:function(t){return n(Math.floor(e(t)))},ceil:function(t){return n(Math.ceil(e(t)))}}))},i}function W(){var t=U(S()).domain([1,10]);return t.copy=function(){return E(t,W()).base(t.base())},i.apply(t,arguments),t}function G(t){return function(e){return Math.sign(e)*Math.log1p(Math.abs(e/t))}}function V(t){return function(e){return Math.sign(e)*Math.expm1(Math.abs(e))*t}}function $(t){var e=1,n=t(G(e),V(e));return n.constant=function(n){return arguments.length?t(G(e=+n),V(e)):e},N(n)}function X(){var t=$(S());return t.copy=function(){return E(t,X()).constant(t.constant())},i.apply(t,arguments)}function J(t){return function(e){return e<0?-Math.pow(-e,t):Math.pow(e,t)}}function Z(t){return t<0?-Math.sqrt(-t):Math.sqrt(t)}function K(t){return t<0?-t*t:t*t}function Q(t){var e=t(b,b),n=1;function r(){return 1===n?t(b,b):.5===n?t(Z,K):t(J(n),J(1/n))}return e.exponent=function(t){return arguments.length?(n=+t,r()):n},N(e)}function tt(){var t=Q(S());return t.copy=function(){return E(t,tt()).exponent(t.exponent())},i.apply(t,arguments),t}function et(){return tt.apply(null,arguments).exponent(.5)}function nt(){var t,e=[],n=[],a=[];function o(){var t=0,i=Math.max(1,n.length);for(a=new Array(i-1);++t<i;)a[t-1]=Object(r.r)(e,t/i);return u}function u(e){return isNaN(e=+e)?t:n[Object(r.b)(a,e)]}return u.invertExtent=function(t){var r=n.indexOf(t);return r<0?[NaN,NaN]:[r>0?a[r-1]:e[0],r<a.length?a[r]:e[e.length-1]]},u.domain=function(t){if(!arguments.length)return e.slice();e=[];for(var n,i=0,a=t.length;i<a;++i)null==(n=t[i])||isNaN(n=+n)||e.push(n);return e.sort(r.a),o()},u.range=function(t){return arguments.length?(n=c.call(t),o()):n.slice()},u.unknown=function(e){return arguments.length?(t=e,u):t},u.quantiles=function(){return a.slice()},u.copy=function(){return nt().domain(e).range(n).unknown(t)},i.apply(u,arguments)}function rt(){var t,e=0,n=1,a=1,o=[.5],u=[0,1];function s(e){return e<=e?u[Object(r.b)(o,e,0,a)]:t}function l(){var t=-1;for(o=new Array(a);++t<a;)o[t]=((t+1)*n-(t-a)*e)/(a+1);return s}return s.domain=function(t){return arguments.length?(e=+t[0],n=+t[1],l()):[e,n]},s.range=function(t){return arguments.length?(a=(u=c.call(t)).length-1,l()):u.slice()},s.invertExtent=function(t){var r=u.indexOf(t);return r<0?[NaN,NaN]:r<1?[e,o[0]]:r>=a?[o[a-1],n]:[o[r-1],o[r]]},s.unknown=function(e){return arguments.length?(t=e,s):s},s.thresholds=function(){return o.slice()},s.copy=function(){return rt().domain([e,n]).range(u).unknown(t)},i.apply(N(s),arguments)}function it(){var t,e=[.5],n=[0,1],a=1;function o(i){return i<=i?n[Object(r.b)(e,i,0,a)]:t}return o.domain=function(t){return arguments.length?(e=c.call(t),a=Math.min(e.length,n.length-1),o):e.slice()},o.range=function(t){return arguments.length?(n=c.call(t),a=Math.min(e.length,n.length-1),o):n.slice()},o.invertExtent=function(t){var r=n.indexOf(t);return[e[r-1],e[r]]},o.unknown=function(e){return arguments.length?(t=e,o):t},o.copy=function(){return it().domain(e).range(n).unknown(t)},i.apply(o,arguments)}var at=n(64),ot=n(180),ut=n(12),st=n(127),ct=n(181),lt=n(182),ft=n(112),dt=n(113),ht=n(41);function pt(t){return new Date(t)}function mt(t){return t instanceof Date?+t:+new Date(+t)}function vt(t,e,n,i,a,o,u,c,l){var f=T(b,b),d=f.invert,h=f.domain,p=l(".%L"),m=l(":%S"),v=l("%I:%M"),g=l("%I %p"),_=l("%a %d"),y=l("%b %d"),w=l("%B"),x=l("%Y"),k=[[u,1,1e3],[u,5,5e3],[u,15,15e3],[u,30,3e4],[o,1,6e4],[o,5,3e5],[o,15,9e5],[o,30,18e5],[a,1,36e5],[a,3,108e5],[a,6,216e5],[a,12,432e5],[i,1,864e5],[i,2,1728e5],[n,1,6048e5],[e,1,2592e6],[e,3,7776e6],[t,1,31536e6]];function M(r){return(u(r)<r?p:o(r)<r?m:a(r)<r?v:i(r)<r?g:e(r)<r?n(r)<r?_:y:t(r)<r?w:x)(r)}function S(e,n,i,a){if(null==e&&(e=10),"number"==typeof e){var o=Math.abs(i-n)/e,u=Object(r.e)((function(t){return t[2]})).right(k,o);u===k.length?(a=Object(r.A)(n/31536e6,i/31536e6,e),e=t):u?(a=(u=k[o/k[u-1][2]<k[u][2]/o?u-1:u])[1],e=u[0]):(a=Math.max(Object(r.A)(n,i,e),1),e=c)}return null==a?e:e.every(a)}return f.invert=function(t){return new Date(d(t))},f.domain=function(t){return arguments.length?h(s.call(t,mt)):h().map(pt)},f.ticks=function(t,e){var n,r=h(),i=r[0],a=r[r.length-1],o=a<i;return o&&(n=i,i=a,a=n),n=(n=S(t,i,a,e))?n.range(i,a+1):[],o?n.reverse():n},f.tickFormat=function(t,e){return null==e?M:l(e)},f.nice=function(t,e){var n=h();return(t=S(t,n[0],n[n.length-1],e))?h(Y(n,t)):f},f.copy=function(){return E(f,vt(t,e,n,i,a,o,u,c,l))},f}var gt=function(){return i.apply(vt(at.a,ot.a,ut.g,st.b,ct.a,lt.a,ft.a,dt.a,ht.b).domain([new Date(2e3,0,1),new Date(2e3,0,2)]),arguments)},_t=n(65),yt=n(183),bt=n(13),wt=n(128),xt=n(184),kt=n(185),Mt=function(){return i.apply(vt(_t.a,yt.a,bt.g,wt.a,xt.a,kt.a,ft.a,dt.a,ht.d).domain([Date.UTC(2e3,0,1),Date.UTC(2e3,0,2)]),arguments)};function Et(){var t,e,n,r,i,a=0,o=1,u=b,s=!1;function c(e){return isNaN(e=+e)?i:u(0===n?.5:(e=(r(e)-t)*n,s?Math.max(0,Math.min(1,e)):e))}return c.domain=function(i){return arguments.length?(t=r(a=+i[0]),e=r(o=+i[1]),n=t===e?0:1/(e-t),c):[a,o]},c.clamp=function(t){return arguments.length?(s=!!t,c):s},c.interpolator=function(t){return arguments.length?(u=t,c):u},c.unknown=function(t){return arguments.length?(i=t,c):i},function(i){return r=i,t=i(a),e=i(o),n=t===e?0:1/(e-t),c}}function St(t,e){return e.domain(t.domain()).interpolator(t.interpolator()).clamp(t.clamp()).unknown(t.unknown())}function Tt(){var t=N(Et()(b));return t.copy=function(){return St(t,Tt())},a.apply(t,arguments)}function jt(){var t=U(Et()).domain([1,10]);return t.copy=function(){return St(t,jt()).base(t.base())},a.apply(t,arguments)}function Ot(){var t=$(Et());return t.copy=function(){return St(t,Ot()).constant(t.constant())},a.apply(t,arguments)}function At(){var t=Q(Et());return t.copy=function(){return St(t,At()).exponent(t.exponent())},a.apply(t,arguments)}function Lt(){return At.apply(null,arguments).exponent(.5)}function Ct(){var t=[],e=b;function n(n){if(!isNaN(n=+n))return e((Object(r.b)(t,n)-1)/(t.length-1))}return n.domain=function(e){if(!arguments.length)return t.slice();t=[];for(var i,a=0,o=e.length;a<o;++a)null==(i=e[a])||isNaN(i=+i)||t.push(i);return t.sort(r.a),n},n.interpolator=function(t){return arguments.length?(e=t,n):e},n.copy=function(){return Ct(e).domain(t)},a.apply(n,arguments)}function Dt(){var t,e,n,r,i,a,o,u=0,s=.5,c=1,l=b,f=!1;function d(t){return isNaN(t=+t)?o:(t=.5+((t=+a(t))-e)*(t<e?r:i),l(f?Math.max(0,Math.min(1,t)):t))}return d.domain=function(o){return arguments.length?(t=a(u=+o[0]),e=a(s=+o[1]),n=a(c=+o[2]),r=t===e?0:.5/(e-t),i=e===n?0:.5/(n-e),d):[u,s,c]},d.clamp=function(t){return arguments.length?(f=!!t,d):f},d.interpolator=function(t){return arguments.length?(l=t,d):l},d.unknown=function(t){return arguments.length?(o=t,d):o},function(o){return a=o,t=o(u),e=o(s),n=o(c),r=t===e?0:.5/(e-t),i=e===n?0:.5/(n-e),d}}function Nt(){var t=N(Dt()(b));return t.copy=function(){return St(t,Nt())},a.apply(t,arguments)}function Pt(){var t=U(Dt()).domain([.1,1,10]);return t.copy=function(){return St(t,Pt()).base(t.base())},a.apply(t,arguments)}function Rt(){var t=$(Dt());return t.copy=function(){return St(t,Rt()).constant(t.constant())},a.apply(t,arguments)}function Yt(){var t=Q(Dt());return t.copy=function(){return St(t,Yt()).exponent(t.exponent())},a.apply(t,arguments)}function It(){return Yt.apply(null,arguments).exponent(.5)}},function(t,e,n){(function(t){var r=n(487),i=n(488),a=n(489),o=n(490),u=n(491),s=n(492),c=navigator.userAgent.match(/Android/i),l=/PhantomJS/i.test(navigator.userAgent),f=c||l;e.protocol=3;var d=e.packets={open:0,close:1,ping:2,pong:3,message:4,upgrade:5,noop:6},h=r(d),p={type:"error",data:"parser error"},m=n(493);function v(t,e,n){for(var r=new Array(t.length),i=u(t.length,n),a=function(t,n,i){e(n,(function(e,n){r[t]=n,i(e,r)}))},o=0;o<t.length;o++)a(o,t[o],i)}e.encodePacket=function(n,r,i,a){"function"==typeof r&&(a=r,r=!1),"function"==typeof i&&(a=i,i=null);var o=void 0===n.data?void 0:n.data.buffer||n.data;if(t.ArrayBuffer&&o instanceof ArrayBuffer)return function(t,n,r){if(!n)return e.encodeBase64Packet(t,r);var i=t.data,a=new Uint8Array(i),o=new Uint8Array(1+i.byteLength);o[0]=d[t.type];for(var u=0;u<a.length;u++)o[u+1]=a[u];return r(o.buffer)}(n,r,a);if(m&&o instanceof t.Blob)return function(t,n,r){if(!n)return e.encodeBase64Packet(t,r);if(f)return function(t,n,r){if(!n)return e.encodeBase64Packet(t,r);var i=new FileReader;return i.onload=function(){t.data=i.result,e.encodePacket(t,n,!0,r)},i.readAsArrayBuffer(t.data)}(t,n,r);var i=new Uint8Array(1);i[0]=d[t.type];var a=new m([i.buffer,t.data]);return r(a)}(n,r,a);if(o&&o.base64)return function(t,n){var r="b"+e.packets[t.type]+t.data.data;return n(r)}(n,a);var u=d[n.type];return void 0!==n.data&&(u+=i?s.encode(String(n.data)):String(n.data)),a(""+u)},e.encodeBase64Packet=function(n,r){var i,a="b"+e.packets[n.type];if(m&&n.data instanceof t.Blob){var o=new FileReader;return o.onload=function(){var t=o.result.split(",")[1];r(a+t)},o.readAsDataURL(n.data)}try{i=String.fromCharCode.apply(null,new Uint8Array(n.data))}catch(t){for(var u=new Uint8Array(n.data),s=new Array(u.length),c=0;c<u.length;c++)s[c]=u[c];i=String.fromCharCode.apply(null,s)}return a+=t.btoa(i),r(a)},e.decodePacket=function(t,n,r){if("string"==typeof t||void 0===t){if("b"==t.charAt(0))return e.decodeBase64Packet(t.substr(1),n);if(r)try{t=s.decode(t)}catch(t){return p}var i=t.charAt(0);return Number(i)==i&&h[i]?t.length>1?{type:h[i],data:t.substring(1)}:{type:h[i]}:p}i=new Uint8Array(t)[0];var o=a(t,1);return m&&"blob"===n&&(o=new m([o])),{type:h[i],data:o}},e.decodeBase64Packet=function(e,n){var r=h[e.charAt(0)];if(!t.ArrayBuffer)return{type:r,data:{base64:!0,data:e.substr(1)}};var i=o.decode(e.substr(1));return"blob"===n&&m&&(i=new m([i])),{type:r,data:i}},e.encodePayload=function(t,n,r){"function"==typeof n&&(r=n,n=null);var a=i(t);if(n&&a)return m&&!f?e.encodePayloadAsBlob(t,r):e.encodePayloadAsArrayBuffer(t,r);if(!t.length)return r("0:");v(t,(function(t,r){e.encodePacket(t,!!a&&n,!0,(function(t){r(null,function(t){return t.length+":"+t}(t))}))}),(function(t,e){return r(e.join(""))}))},e.decodePayload=function(t,n,r){if("string"!=typeof t)return e.decodePayloadAsBinary(t,n,r);var i;if("function"==typeof n&&(r=n,n=null),""==t)return r(p,0,1);for(var a,o,u="",s=0,c=t.length;s<c;s++){var l=t.charAt(s);if(":"!=l)u+=l;else{if(""==u||u!=(a=Number(u)))return r(p,0,1);if(u!=(o=t.substr(s+1,a)).length)return r(p,0,1);if(o.length){if(i=e.decodePacket(o,n,!0),p.type==i.type&&p.data==i.data)return r(p,0,1);if(!1===r(i,s+a,c))return}s+=a,u=""}}return""!=u?r(p,0,1):void 0},e.encodePayloadAsArrayBuffer=function(t,n){if(!t.length)return n(new ArrayBuffer(0));v(t,(function(t,n){e.encodePacket(t,!0,!0,(function(t){return n(null,t)}))}),(function(t,e){var r=e.reduce((function(t,e){var n;return t+(n="string"==typeof e?e.length:e.byteLength).toString().length+n+2}),0),i=new Uint8Array(r),a=0;return e.forEach((function(t){var e="string"==typeof t,n=t;if(e){for(var r=new Uint8Array(t.length),o=0;o<t.length;o++)r[o]=t.charCodeAt(o);n=r.buffer}i[a++]=e?0:1;var u=n.byteLength.toString();for(o=0;o<u.length;o++)i[a++]=parseInt(u[o]);i[a++]=255;for(r=new Uint8Array(n),o=0;o<r.length;o++)i[a++]=r[o]})),n(i.buffer)}))},e.encodePayloadAsBlob=function(t,n){v(t,(function(t,n){e.encodePacket(t,!0,!0,(function(t){var e=new Uint8Array(1);if(e[0]=1,"string"==typeof t){for(var r=new Uint8Array(t.length),i=0;i<t.length;i++)r[i]=t.charCodeAt(i);t=r.buffer,e[0]=0}var a=(t instanceof ArrayBuffer?t.byteLength:t.size).toString(),o=new Uint8Array(a.length+1);for(i=0;i<a.length;i++)o[i]=parseInt(a[i]);if(o[a.length]=255,m){var u=new m([e.buffer,o.buffer,t]);n(null,u)}}))}),(function(t,e){return n(new m(e))}))},e.decodePayloadAsBinary=function(t,n,r){"function"==typeof n&&(r=n,n=null);for(var i=t,o=[],u=!1;i.byteLength>0;){for(var s=new Uint8Array(i),c=0===s[0],l="",f=1;255!=s[f];f++){if(l.length>310){u=!0;break}l+=s[f]}if(u)return r(p,0,1);i=a(i,2+l.length),l=parseInt(l);var d=a(i,0,l);if(c)try{d=String.fromCharCode.apply(null,new Uint8Array(d))}catch(t){var h=new Uint8Array(d);d="";for(f=0;f<h.length;f++)d+=String.fromCharCode(h[f])}o.push(d),i=a(i,l)}var m=o.length;o.forEach((function(t,i){r(e.decodePacket(t,n,!0),i,m)}))}}).call(this,n(6))},function(t,e,n){"use strict";var r=n(467);n.d(e,"a",(function(){return r.a}))},function(t,e,n){"use strict";var r;function i(t,e,n){return e in t?Object.defineProperty(t,e,{value:n,enumerable:!0,configurable:!0,writable:!0}):t[e]=n,t}Object.defineProperty(e,"__esModule",{value:!0}),e.amino_acid_text_color=e.amino_acid_colors=e.amino_acid_color=e.nucleotide_difference=e.nucleotide_text_color=e.nucleotide_colors=e.nucleotide_color=void 0;var a={a:"LightPink",g:"LemonChiffon",t:"LightBlue",c:"MediumPurple",y:"DarkGrey",r:"DarkGrey",w:"DarkGrey",s:"DarkGrey",k:"DarkGrey",m:"DarkGrey",d:"DarkGrey",v:"DarkGrey",h:"DarkGrey",b:"DarkGrey",n:"DarkGrey",x:"DarkGrey",A:"LightPink",G:"LemonChiffon",T:"LightBlue",C:"MediumPurple",Y:"DarkGrey",R:"DarkGrey",W:"DarkGrey",S:"DarkGrey",K:"DarkGrey",M:"DarkGrey",D:"DarkGrey",V:"DarkGrey",H:"DarkGrey",B:"DarkGrey",N:"DarkGrey",X:"DarkGrey","-":"lightgrey"},o={a:"Red",g:"GoldenRod",t:"Blue",c:"DarkMagenta",y:"black",r:"black",w:"black",s:"black",k:"black",m:"black",d:"black",v:"black",h:"black",b:"black",n:"black",x:"black",A:"Red",G:"GoldenRod",T:"Blue",C:"DarkMagenta",Y:"black",R:"black",W:"black",S:"black",K:"black",M:"black",D:"black",V:"black",H:"black",B:"black",N:"black",X:"black","-":"DarkGrey"},u=(i(r={"-":"Snow",a:"lightblue",c:"pink",d:"LightSteelBlue",e:"purple",f:"AntiqueWhite",g:"LightSalmon",h:"CadetBlue",i:"Crimson",k:"DarkCyan",l:"DarkKhaki",m:"steelblue",r:"DarkSeaGreen",p:"yellow",q:"lightgreen"},"r","orange"),i(r,"s","green"),i(r,"t","DeepSkyBlue"),i(r,"v","Gold"),i(r,"w","HotPink"),i(r,"x","black"),i(r,"y","IndianRed"),i(r,"A","lightblue"),i(r,"C","pink"),i(r,"D","LightSteelBlue"),i(r,"E","purple"),i(r,"F","AntiqueWhite"),i(r,"G","LightSalmon"),i(r,"H","CadetBlue"),i(r,"I","Crimson"),i(r,"K","DarkCyan"),i(r,"L","DarkKhaki"),i(r,"M","steelblue"),i(r,"N","DarkSeaGreen"),i(r,"P","yellow"),i(r,"Q","lightgreen"),i(r,"R","orange"),i(r,"S","green"),i(r,"T","DeepSkyBlue"),i(r,"V","Gold"),i(r,"W","HotPink"),i(r,"X","black"),i(r,"Y","IndianRed"),r);e.nucleotide_difference=function(t){var e=t.header,n=t.seq;return function(t,r,i){var o=a[t];return"-"==t?a["-"]:i==e?o:t==n[r-1]?"white":o}},e.amino_acid_text_color=function(t,e,n){return"X"!=t.toUpperCase()?"black":"white"},e.amino_acid_color=function(t,e,n){return u[t]},e.nucleotide_text_color=function(t,e,n){return o[t]},e.nucleotide_color=function(t,e,n){return a[t]},e.amino_acid_colors=u,e.nucleotide_colors=a},function(t,e,n){var r=n(135);t.exports=function(t,e,n){if(r(t),void 0===e)return t;switch(n){case 1:return function(n){return t.call(e,n)};case 2:return function(n,r){return t.call(e,n,r)};case 3:return function(n,r,i){return t.call(e,n,r,i)}}return function(){return t.apply(e,arguments)}}},function(t,e,n){var r=n(136);t.exports=function(t){return Object(r(t))}},function(t,e){t.exports={}},function(t,e,n){t.exports=n(574)},function(t,e){t.exports=function(t,e){if(!(t instanceof e))throw new TypeError("Cannot call a class as a function")}},function(t,e){function n(t,e){for(var n=0;n<e.length;n++){var r=e[n];r.enumerable=r.enumerable||!1,r.configurable=!0,"value"in r&&(r.writable=!0),Object.defineProperty(t,r.key,r)}}t.exports=function(t,e,r){return e&&n(t.prototype,e),r&&n(t,r),t}},function(t,e){(e=t.exports=function(t){return t.replace(/^\s*|\s*$/g,"")}).left=function(t){return t.replace(/^\s*/,"")},e.right=function(t){return t.replace(/\s*$/,"")}},function(t,e,n){"use strict";n.r(e),n.d(e,"version",(function(){return r})),n.d(e,"bisect",(function(){return i.b})),n.d(e,"bisectRight",(function(){return i.d})),n.d(e,"bisectLeft",(function(){return i.c})),n.d(e,"ascending",(function(){return i.a})),n.d(e,"bisector",(function(){return i.e})),n.d(e,"cross",(function(){return i.f})),n.d(e,"descending",(function(){return i.g})),n.d(e,"deviation",(function(){return i.h})),n.d(e,"extent",(function(){return i.i})),n.d(e,"histogram",(function(){return i.j})),n.d(e,"thresholdFreedmanDiaconis",(function(){return i.w})),n.d(e,"thresholdScott",(function(){return i.x})),n.d(e,"thresholdSturges",(function(){return i.y})),n.d(e,"max",(function(){return i.k})),n.d(e,"mean",(function(){return i.l})),n.d(e,"median",(function(){return i.m})),n.d(e,"merge",(function(){return i.n})),n.d(e,"min",(function(){return i.o})),n.d(e,"pairs",(function(){return i.p})),n.d(e,"permute",(function(){return i.q})),n.d(e,"quantile",(function(){return i.r})),n.d(e,"range",(function(){return i.s})),n.d(e,"scan",(function(){return i.t})),n.d(e,"shuffle",(function(){return i.u})),n.d(e,"sum",(function(){return i.v})),n.d(e,"ticks",(function(){return i.B})),n.d(e,"tickIncrement",(function(){return i.z})),n.d(e,"tickStep",(function(){return i.A})),n.d(e,"transpose",(function(){return i.C})),n.d(e,"variance",(function(){return i.D})),n.d(e,"zip",(function(){return i.E})),n.d(e,"axisTop",(function(){return a.d})),n.d(e,"axisRight",(function(){return a.c})),n.d(e,"axisBottom",(function(){return a.a})),n.d(e,"axisLeft",(function(){return a.b})),n.d(e,"brush",(function(){return o.a})),n.d(e,"brushX",(function(){return o.c})),n.d(e,"brushY",(function(){return o.d})),n.d(e,"brushSelection",(function(){return o.b})),n.d(e,"chord",(function(){return u.a})),n.d(e,"ribbon",(function(){return u.b})),n.d(e,"nest",(function(){return s.d})),n.d(e,"set",(function(){return s.e})),n.d(e,"map",(function(){return s.c})),n.d(e,"keys",(function(){return s.b})),n.d(e,"values",(function(){return s.f})),n.d(e,"entries",(function(){return s.a})),n.d(e,"color",(function(){return c.a})),n.d(e,"rgb",(function(){return c.h})),n.d(e,"hsl",(function(){return c.e})),n.d(e,"lab",(function(){return c.f})),n.d(e,"hcl",(function(){return c.d})),n.d(e,"lch",(function(){return c.g})),n.d(e,"gray",(function(){return c.c})),n.d(e,"cubehelix",(function(){return c.b})),n.d(e,"contours",(function(){return l.b})),n.d(e,"contourDensity",(function(){return l.a})),n.d(e,"dispatch",(function(){return f.a})),n.d(e,"drag",(function(){return d.a})),n.d(e,"dragDisable",(function(){return d.b})),n.d(e,"dragEnable",(function(){return d.c})),n.d(e,"dsvFormat",(function(){return h.i})),n.d(e,"csvParse",(function(){return h.g})),n.d(e,"csvParseRows",(function(){return h.h})),n.d(e,"csvFormat",(function(){return h.b})),n.d(e,"csvFormatBody",(function(){return h.c})),n.d(e,"csvFormatRows",(function(){return h.e})),n.d(e,"csvFormatRow",(function(){return h.d})),n.d(e,"csvFormatValue",(function(){return h.f})),n.d(e,"tsvParse",(function(){return h.o})),n.d(e,"tsvParseRows",(function(){return h.p})),n.d(e,"tsvFormat",(function(){return h.j})),n.d(e,"tsvFormatBody",(function(){return h.k})),n.d(e,"tsvFormatRows",(function(){return h.m})),n.d(e,"tsvFormatRow",(function(){return h.l})),n.d(e,"tsvFormatValue",(function(){return h.n})),n.d(e,"autoType",(function(){return h.a})),n.d(e,"easeLinear",(function(){return p.y})),n.d(e,"easeQuad",(function(){return p.D})),n.d(e,"easeQuadIn",(function(){return p.E})),n.d(e,"easeQuadOut",(function(){return p.G})),n.d(e,"easeQuadInOut",(function(){return p.F})),n.d(e,"easeCubic",(function(){return p.m})),n.d(e,"easeCubicIn",(function(){return p.n})),n.d(e,"easeCubicOut",(function(){return p.p})),n.d(e,"easeCubicInOut",(function(){return p.o})),n.d(e,"easePoly",(function(){return p.z})),n.d(e,"easePolyIn",(function(){return p.A})),n.d(e,"easePolyOut",(function(){return p.C})),n.d(e,"easePolyInOut",(function(){return p.B})),n.d(e,"easeSin",(function(){return p.H})),n.d(e,"easeSinIn",(function(){return p.I})),n.d(e,"easeSinOut",(function(){return p.K})),n.d(e,"easeSinInOut",(function(){return p.J})),n.d(e,"easeExp",(function(){return p.u})),n.d(e,"easeExpIn",(function(){return p.v})),n.d(e,"easeExpOut",(function(){return p.x})),n.d(e,"easeExpInOut",(function(){return p.w})),n.d(e,"easeCircle",(function(){return p.i})),n.d(e,"easeCircleIn",(function(){return p.j})),n.d(e,"easeCircleOut",(function(){return p.l})),n.d(e,"easeCircleInOut",(function(){return p.k})),n.d(e,"easeBounce",(function(){return p.e})),n.d(e,"easeBounceIn",(function(){return p.f})),n.d(e,"easeBounceOut",(function(){return p.h})),n.d(e,"easeBounceInOut",(function(){return p.g})),n.d(e,"easeBack",(function(){return p.a})),n.d(e,"easeBackIn",(function(){return p.b})),n.d(e,"easeBackOut",(function(){return p.d})),n.d(e,"easeBackInOut",(function(){return p.c})),n.d(e,"easeElastic",(function(){return p.q})),n.d(e,"easeElasticIn",(function(){return p.r})),n.d(e,"easeElasticOut",(function(){return p.t})),n.d(e,"easeElasticInOut",(function(){return p.s})),n.d(e,"blob",(function(){return m.a})),n.d(e,"buffer",(function(){return m.b})),n.d(e,"dsv",(function(){return m.d})),n.d(e,"csv",(function(){return m.c})),n.d(e,"tsv",(function(){return m.j})),n.d(e,"image",(function(){return m.f})),n.d(e,"json",(function(){return m.g})),n.d(e,"text",(function(){return m.i})),n.d(e,"xml",(function(){return m.k})),n.d(e,"html",(function(){return m.e})),n.d(e,"svg",(function(){return m.h})),n.d(e,"forceCenter",(function(){return v.a})),n.d(e,"forceCollide",(function(){return v.b})),n.d(e,"forceLink",(function(){return v.c})),n.d(e,"forceManyBody",(function(){return v.d})),n.d(e,"forceRadial",(function(){return v.e})),n.d(e,"forceSimulation",(function(){return v.f})),n.d(e,"forceX",(function(){return v.g})),n.d(e,"forceY",(function(){return v.h})),n.d(e,"formatDefaultLocale",(function(){return g.c})),n.d(e,"format",(function(){return g.b})),n.d(e,"formatPrefix",(function(){return g.e})),n.d(e,"formatLocale",(function(){return g.d})),n.d(e,"formatSpecifier",(function(){return g.f})),n.d(e,"FormatSpecifier",(function(){return g.a})),n.d(e,"precisionFixed",(function(){return g.g})),n.d(e,"precisionPrefix",(function(){return g.h})),n.d(e,"precisionRound",(function(){return g.i})),n.d(e,"geoArea",(function(){return _.c})),n.d(e,"geoBounds",(function(){return _.h})),n.d(e,"geoCentroid",(function(){return _.i})),n.d(e,"geoCircle",(function(){return _.j})),n.d(e,"geoClipAntimeridian",(function(){return _.k})),n.d(e,"geoClipCircle",(function(){return _.l})),n.d(e,"geoClipExtent",(function(){return _.m})),n.d(e,"geoClipRectangle",(function(){return _.n})),n.d(e,"geoContains",(function(){return _.u})),n.d(e,"geoDistance",(function(){return _.v})),n.d(e,"geoGraticule",(function(){return _.C})),n.d(e,"geoGraticule10",(function(){return _.D})),n.d(e,"geoInterpolate",(function(){return _.F})),n.d(e,"geoLength",(function(){return _.G})),n.d(e,"geoPath",(function(){return _.N})),n.d(e,"geoAlbers",(function(){return _.a})),n.d(e,"geoAlbersUsa",(function(){return _.b})),n.d(e,"geoAzimuthalEqualArea",(function(){return _.d})),n.d(e,"geoAzimuthalEqualAreaRaw",(function(){return _.e})),n.d(e,"geoAzimuthalEquidistant",(function(){return _.f})),n.d(e,"geoAzimuthalEquidistantRaw",(function(){return _.g})),n.d(e,"geoConicConformal",(function(){return _.o})),n.d(e,"geoConicConformalRaw",(function(){return _.p})),n.d(e,"geoConicEqualArea",(function(){return _.q})),n.d(e,"geoConicEqualAreaRaw",(function(){return _.r})),n.d(e,"geoConicEquidistant",(function(){return _.s})),n.d(e,"geoConicEquidistantRaw",(function(){return _.t})),n.d(e,"geoEqualEarth",(function(){return _.w})),n.d(e,"geoEqualEarthRaw",(function(){return _.x})),n.d(e,"geoEquirectangular",(function(){return _.y})),n.d(e,"geoEquirectangularRaw",(function(){return _.z})),n.d(e,"geoGnomonic",(function(){return _.A})),n.d(e,"geoGnomonicRaw",(function(){return _.B})),n.d(e,"geoIdentity",(function(){return _.E})),n.d(e,"geoProjection",(function(){return _.O})),n.d(e,"geoProjectionMutator",(function(){return _.P})),n.d(e,"geoMercator",(function(){return _.H})),n.d(e,"geoMercatorRaw",(function(){return _.I})),n.d(e,"geoNaturalEarth1",(function(){return _.J})),n.d(e,"geoNaturalEarth1Raw",(function(){return _.K})),n.d(e,"geoOrthographic",(function(){return _.L})),n.d(e,"geoOrthographicRaw",(function(){return _.M})),n.d(e,"geoStereographic",(function(){return _.R})),n.d(e,"geoStereographicRaw",(function(){return _.S})),n.d(e,"geoTransverseMercator",(function(){return _.V})),n.d(e,"geoTransverseMercatorRaw",(function(){return _.W})),n.d(e,"geoRotation",(function(){return _.Q})),n.d(e,"geoStream",(function(){return _.T})),n.d(e,"geoTransform",(function(){return _.U})),n.d(e,"cluster",(function(){return y.a})),n.d(e,"hierarchy",(function(){return y.b})),n.d(e,"pack",(function(){return y.c})),n.d(e,"packSiblings",(function(){return y.e})),n.d(e,"packEnclose",(function(){return y.d})),n.d(e,"partition",(function(){return y.f})),n.d(e,"stratify",(function(){return y.g})),n.d(e,"tree",(function(){return y.h})),n.d(e,"treemap",(function(){return y.i})),n.d(e,"treemapBinary",(function(){return y.j})),n.d(e,"treemapDice",(function(){return y.k})),n.d(e,"treemapSlice",(function(){return y.m})),n.d(e,"treemapSliceDice",(function(){return y.n})),n.d(e,"treemapSquarify",(function(){return y.o})),n.d(e,"treemapResquarify",(function(){return y.l})),n.d(e,"interpolate",(function(){return b.a})),n.d(e,"interpolateArray",(function(){return b.b})),n.d(e,"interpolateBasis",(function(){return b.c})),n.d(e,"interpolateBasisClosed",(function(){return b.d})),n.d(e,"interpolateDate",(function(){return b.g})),n.d(e,"interpolateDiscrete",(function(){return b.h})),n.d(e,"interpolateHue",(function(){return b.m})),n.d(e,"interpolateNumber",(function(){return b.o})),n.d(e,"interpolateNumberArray",(function(){return b.p})),n.d(e,"interpolateObject",(function(){return b.q})),n.d(e,"interpolateRound",(function(){return b.u})),n.d(e,"interpolateString",(function(){return b.v})),n.d(e,"interpolateTransformCss",(function(){return b.w})),n.d(e,"interpolateTransformSvg",(function(){return b.x})),n.d(e,"interpolateZoom",(function(){return b.y})),n.d(e,"interpolateRgb",(function(){return b.r})),n.d(e,"interpolateRgbBasis",(function(){return b.s})),n.d(e,"interpolateRgbBasisClosed",(function(){return b.t})),n.d(e,"interpolateHsl",(function(){return b.k})),n.d(e,"interpolateHslLong",(function(){return b.l})),n.d(e,"interpolateLab",(function(){return b.n})),n.d(e,"interpolateHcl",(function(){return b.i})),n.d(e,"interpolateHclLong",(function(){return b.j})),n.d(e,"interpolateCubehelix",(function(){return b.e})),n.d(e,"interpolateCubehelixLong",(function(){return b.f})),n.d(e,"piecewise",(function(){return b.z})),n.d(e,"quantize",(function(){return b.A})),n.d(e,"path",(function(){return w.a})),n.d(e,"polygonArea",(function(){return x.a})),n.d(e,"polygonCentroid",(function(){return x.b})),n.d(e,"polygonHull",(function(){return x.d})),n.d(e,"polygonContains",(function(){return x.c})),n.d(e,"polygonLength",(function(){return x.e})),n.d(e,"quadtree",(function(){return k.a})),n.d(e,"randomUniform",(function(){return M.f})),n.d(e,"randomNormal",(function(){return M.e})),n.d(e,"randomLogNormal",(function(){return M.d})),n.d(e,"randomBates",(function(){return M.a})),n.d(e,"randomIrwinHall",(function(){return M.c})),n.d(e,"randomExponential",(function(){return M.b})),n.d(e,"scaleBand",(function(){return C})),n.d(e,"scalePoint",(function(){return N})),n.d(e,"scaleIdentity",(function(){return nt})),n.d(e,"scaleLinear",(function(){return et})),n.d(e,"scaleLog",(function(){return ft})),n.d(e,"scaleSymlog",(function(){return mt})),n.d(e,"scaleOrdinal",(function(){return L})),n.d(e,"scaleImplicit",(function(){return A})),n.d(e,"scalePow",(function(){return bt})),n.d(e,"scaleSqrt",(function(){return wt})),n.d(e,"scaleQuantile",(function(){return xt})),n.d(e,"scaleQuantize",(function(){return kt})),n.d(e,"scaleThreshold",(function(){return Mt})),n.d(e,"scaleTime",(function(){return Yt})),n.d(e,"scaleUtc",(function(){return Ut})),n.d(e,"scaleSequential",(function(){return Vt})),n.d(e,"scaleSequentialLog",(function(){return $t})),n.d(e,"scaleSequentialPow",(function(){return Jt})),n.d(e,"scaleSequentialSqrt",(function(){return Zt})),n.d(e,"scaleSequentialSymlog",(function(){return Xt})),n.d(e,"scaleSequentialQuantile",(function(){return Kt})),n.d(e,"scaleDiverging",(function(){return te})),n.d(e,"scaleDivergingLog",(function(){return ee})),n.d(e,"scaleDivergingPow",(function(){return re})),n.d(e,"scaleDivergingSqrt",(function(){return ie})),n.d(e,"scaleDivergingSymlog",(function(){return ne})),n.d(e,"tickFormat",(function(){return Q})),n.d(e,"schemeCategory10",(function(){return ae.R})),n.d(e,"schemeAccent",(function(){return ae.M})),n.d(e,"schemeDark2",(function(){return ae.S})),n.d(e,"schemePaired",(function(){return ae.Z})),n.d(e,"schemePastel1",(function(){return ae.ab})),n.d(e,"schemePastel2",(function(){return ae.bb})),n.d(e,"schemeSet1",(function(){return ae.ob})),n.d(e,"schemeSet2",(function(){return ae.pb})),n.d(e,"schemeSet3",(function(){return ae.qb})),n.d(e,"schemeTableau10",(function(){return ae.sb})),n.d(e,"interpolateBrBG",(function(){return ae.b})),n.d(e,"schemeBrBG",(function(){return ae.O})),n.d(e,"interpolatePRGn",(function(){return ae.o})),n.d(e,"schemePRGn",(function(){return ae.Y})),n.d(e,"interpolatePiYG",(function(){return ae.p})),n.d(e,"schemePiYG",(function(){return ae.cb})),n.d(e,"interpolatePuOr",(function(){return ae.t})),n.d(e,"schemePuOr",(function(){return ae.fb})),n.d(e,"interpolateRdBu",(function(){return ae.x})),n.d(e,"schemeRdBu",(function(){return ae.ib})),n.d(e,"interpolateRdGy",(function(){return ae.y})),n.d(e,"schemeRdGy",(function(){return ae.jb})),n.d(e,"interpolateRdYlBu",(function(){return ae.A})),n.d(e,"schemeRdYlBu",(function(){return ae.lb})),n.d(e,"interpolateRdYlGn",(function(){return ae.B})),n.d(e,"schemeRdYlGn",(function(){return ae.mb})),n.d(e,"interpolateSpectral",(function(){return ae.E})),n.d(e,"schemeSpectral",(function(){return ae.rb})),n.d(e,"interpolateBuGn",(function(){return ae.c})),n.d(e,"schemeBuGn",(function(){return ae.P})),n.d(e,"interpolateBuPu",(function(){return ae.d})),n.d(e,"schemeBuPu",(function(){return ae.Q})),n.d(e,"interpolateGnBu",(function(){return ae.h})),n.d(e,"schemeGnBu",(function(){return ae.T})),n.d(e,"interpolateOrRd",(function(){return ae.m})),n.d(e,"schemeOrRd",(function(){return ae.W})),n.d(e,"interpolatePuBuGn",(function(){return ae.s})),n.d(e,"schemePuBuGn",(function(){return ae.eb})),n.d(e,"interpolatePuBu",(function(){return ae.r})),n.d(e,"schemePuBu",(function(){return ae.db})),n.d(e,"interpolatePuRd",(function(){return ae.u})),n.d(e,"schemePuRd",(function(){return ae.gb})),n.d(e,"interpolateRdPu",(function(){return ae.z})),n.d(e,"schemeRdPu",(function(){return ae.kb})),n.d(e,"interpolateYlGnBu",(function(){return ae.J})),n.d(e,"schemeYlGnBu",(function(){return ae.ub})),n.d(e,"interpolateYlGn",(function(){return ae.I})),n.d(e,"schemeYlGn",(function(){return ae.tb})),n.d(e,"interpolateYlOrBr",(function(){return ae.K})),n.d(e,"schemeYlOrBr",(function(){return ae.vb})),n.d(e,"interpolateYlOrRd",(function(){return ae.L})),n.d(e,"schemeYlOrRd",(function(){return ae.wb})),n.d(e,"interpolateBlues",(function(){return ae.a})),n.d(e,"schemeBlues",(function(){return ae.N})),n.d(e,"interpolateGreens",(function(){return ae.i})),n.d(e,"schemeGreens",(function(){return ae.U})),n.d(e,"interpolateGreys",(function(){return ae.j})),n.d(e,"schemeGreys",(function(){return ae.V})),n.d(e,"interpolatePurples",(function(){return ae.v})),n.d(e,"schemePurples",(function(){return ae.hb})),n.d(e,"interpolateReds",(function(){return ae.C})),n.d(e,"schemeReds",(function(){return ae.nb})),n.d(e,"interpolateOranges",(function(){return ae.n})),n.d(e,"schemeOranges",(function(){return ae.X})),n.d(e,"interpolateCividis",(function(){return ae.e})),n.d(e,"interpolateCubehelixDefault",(function(){return ae.g})),n.d(e,"interpolateRainbow",(function(){return ae.w})),n.d(e,"interpolateWarm",(function(){return ae.H})),n.d(e,"interpolateCool",(function(){return ae.f})),n.d(e,"interpolateSinebow",(function(){return ae.D})),n.d(e,"interpolateTurbo",(function(){return ae.F})),n.d(e,"interpolateViridis",(function(){return ae.G})),n.d(e,"interpolateMagma",(function(){return ae.l})),n.d(e,"interpolateInferno",(function(){return ae.k})),n.d(e,"interpolatePlasma",(function(){return ae.q})),n.d(e,"create",(function(){return oe.b})),n.d(e,"creator",(function(){return oe.c})),n.d(e,"local",(function(){return oe.f})),n.d(e,"matcher",(function(){return oe.g})),n.d(e,"mouse",(function(){return oe.h})),n.d(e,"namespace",(function(){return oe.i})),n.d(e,"namespaces",(function(){return oe.j})),n.d(e,"clientPoint",(function(){return oe.a})),n.d(e,"select",(function(){return oe.k})),n.d(e,"selectAll",(function(){return oe.l})),n.d(e,"selection",(function(){return oe.m})),n.d(e,"selector",(function(){return oe.n})),n.d(e,"selectorAll",(function(){return oe.o})),n.d(e,"style",(function(){return oe.p})),n.d(e,"touch",(function(){return oe.q})),n.d(e,"touches",(function(){return oe.r})),n.d(e,"window",(function(){return oe.s})),n.d(e,"event",(function(){return oe.e})),n.d(e,"customEvent",(function(){return oe.d})),n.d(e,"arc",(function(){return ue.a})),n.d(e,"area",(function(){return ue.b})),n.d(e,"line",(function(){return ue.v})),n.d(e,"pie",(function(){return ue.A})),n.d(e,"areaRadial",(function(){return ue.c})),n.d(e,"radialArea",(function(){return ue.C})),n.d(e,"lineRadial",(function(){return ue.w})),n.d(e,"radialLine",(function(){return ue.D})),n.d(e,"pointRadial",(function(){return ue.B})),n.d(e,"linkHorizontal",(function(){return ue.x})),n.d(e,"linkVertical",(function(){return ue.z})),n.d(e,"linkRadial",(function(){return ue.y})),n.d(e,"symbol",(function(){return ue.Q})),n.d(e,"symbols",(function(){return ue.Y})),n.d(e,"symbolCircle",(function(){return ue.R})),n.d(e,"symbolCross",(function(){return ue.S})),n.d(e,"symbolDiamond",(function(){return ue.T})),n.d(e,"symbolSquare",(function(){return ue.U})),n.d(e,"symbolStar",(function(){return ue.V})),n.d(e,"symbolTriangle",(function(){return ue.W})),n.d(e,"symbolWye",(function(){return ue.X})),n.d(e,"curveBasisClosed",(function(){return ue.e})),n.d(e,"curveBasisOpen",(function(){return ue.f})),n.d(e,"curveBasis",(function(){return ue.d})),n.d(e,"curveBundle",(function(){return ue.g})),n.d(e,"curveCardinalClosed",(function(){return ue.i})),n.d(e,"curveCardinalOpen",(function(){return ue.j})),n.d(e,"curveCardinal",(function(){return ue.h})),n.d(e,"curveCatmullRomClosed",(function(){return ue.l})),n.d(e,"curveCatmullRomOpen",(function(){return ue.m})),n.d(e,"curveCatmullRom",(function(){return ue.k})),n.d(e,"curveLinearClosed",(function(){return ue.o})),n.d(e,"curveLinear",(function(){return ue.n})),n.d(e,"curveMonotoneX",(function(){return ue.p})),n.d(e,"curveMonotoneY",(function(){return ue.q})),n.d(e,"curveNatural",(function(){return ue.r})),n.d(e,"curveStep",(function(){return ue.s})),n.d(e,"curveStepAfter",(function(){return ue.t})),n.d(e,"curveStepBefore",(function(){return ue.u})),n.d(e,"stack",(function(){return ue.E})),n.d(e,"stackOffsetExpand",(function(){return ue.G})),n.d(e,"stackOffsetDiverging",(function(){return ue.F})),n.d(e,"stackOffsetNone",(function(){return ue.H})),n.d(e,"stackOffsetSilhouette",(function(){return ue.I})),n.d(e,"stackOffsetWiggle",(function(){return ue.J})),n.d(e,"stackOrderAppearance",(function(){return ue.K})),n.d(e,"stackOrderAscending",(function(){return ue.L})),n.d(e,"stackOrderDescending",(function(){return ue.M})),n.d(e,"stackOrderInsideOut",(function(){return ue.N})),n.d(e,"stackOrderNone",(function(){return ue.O})),n.d(e,"stackOrderReverse",(function(){return ue.P})),n.d(e,"timeInterval",(function(){return se.g})),n.d(e,"timeMillisecond",(function(){return se.h})),n.d(e,"timeMilliseconds",(function(){return se.i})),n.d(e,"utcMillisecond",(function(){return se.L})),n.d(e,"utcMilliseconds",(function(){return se.M})),n.d(e,"timeSecond",(function(){return se.r})),n.d(e,"timeSeconds",(function(){return se.s})),n.d(e,"utcSecond",(function(){return se.V})),n.d(e,"utcSeconds",(function(){return se.W})),n.d(e,"timeMinute",(function(){return se.j})),n.d(e,"timeMinutes",(function(){return se.k})),n.d(e,"timeHour",(function(){return se.e})),n.d(e,"timeHours",(function(){return se.f})),n.d(e,"timeDay",(function(){return se.a})),n.d(e,"timeDays",(function(){return se.b})),n.d(e,"timeWeek",(function(){return se.B})),n.d(e,"timeWeeks",(function(){return se.C})),n.d(e,"timeSunday",(function(){return se.t})),n.d(e,"timeSundays",(function(){return se.u})),n.d(e,"timeMonday",(function(){return se.l})),n.d(e,"timeMondays",(function(){return se.m})),n.d(e,"timeTuesday",(function(){return se.x})),n.d(e,"timeTuesdays",(function(){return se.y})),n.d(e,"timeWednesday",(function(){return se.z})),n.d(e,"timeWednesdays",(function(){return se.A})),n.d(e,"timeThursday",(function(){return se.v})),n.d(e,"timeThursdays",(function(){return se.w})),n.d(e,"timeFriday",(function(){return se.c})),n.d(e,"timeFridays",(function(){return se.d})),n.d(e,"timeSaturday",(function(){return se.p})),n.d(e,"timeSaturdays",(function(){return se.q})),n.d(e,"timeMonth",(function(){return se.n})),n.d(e,"timeMonths",(function(){return se.o})),n.d(e,"timeYear",(function(){return se.D})),n.d(e,"timeYears",(function(){return se.E})),n.d(e,"utcMinute",(function(){return se.N})),n.d(e,"utcMinutes",(function(){return se.O})),n.d(e,"utcHour",(function(){return se.J})),n.d(e,"utcHours",(function(){return se.K})),n.d(e,"utcDay",(function(){return se.F})),n.d(e,"utcDays",(function(){return se.G})),n.d(e,"utcWeek",(function(){return se.fb})),n.d(e,"utcWeeks",(function(){return se.gb})),n.d(e,"utcSunday",(function(){return se.X})),n.d(e,"utcSundays",(function(){return se.Y})),n.d(e,"utcMonday",(function(){return se.P})),n.d(e,"utcMondays",(function(){return se.Q})),n.d(e,"utcTuesday",(function(){return se.bb})),n.d(e,"utcTuesdays",(function(){return se.cb})),n.d(e,"utcWednesday",(function(){return se.db})),n.d(e,"utcWednesdays",(function(){return se.eb})),n.d(e,"utcThursday",(function(){return se.Z})),n.d(e,"utcThursdays",(function(){return se.ab})),n.d(e,"utcFriday",(function(){return se.H})),n.d(e,"utcFridays",(function(){return se.I})),n.d(e,"utcSaturday",(function(){return se.T})),n.d(e,"utcSaturdays",(function(){return se.U})),n.d(e,"utcMonth",(function(){return se.R})),n.d(e,"utcMonths",(function(){return se.S})),n.d(e,"utcYear",(function(){return se.hb})),n.d(e,"utcYears",(function(){return se.ib})),n.d(e,"timeFormatDefaultLocale",(function(){return ce.timeFormatDefaultLocale})),n.d(e,"timeFormat",(function(){return ce.timeFormat})),n.d(e,"timeParse",(function(){return ce.timeParse})),n.d(e,"utcFormat",(function(){return ce.utcFormat})),n.d(e,"utcParse",(function(){return ce.utcParse})),n.d(e,"timeFormatLocale",(function(){return ce.timeFormatLocale})),n.d(e,"isoFormat",(function(){return ce.isoFormat})),n.d(e,"isoParse",(function(){return ce.isoParse})),n.d(e,"now",(function(){return le.b})),n.d(e,"timer",(function(){return le.d})),n.d(e,"timerFlush",(function(){return le.e})),n.d(e,"timeout",(function(){return le.c})),n.d(e,"interval",(function(){return le.a})),n.d(e,"transition",(function(){return fe.c})),n.d(e,"active",(function(){return fe.a})),n.d(e,"interrupt",(function(){return fe.b})),n.d(e,"voronoi",(function(){return de.a})),n.d(e,"zoom",(function(){return he.a})),n.d(e,"zoomTransform",(function(){return he.c})),n.d(e,"zoomIdentity",(function(){return he.b}));var r="5.16.0",i=n(2),a=n(157),o=n(156),u=n(154),s=n(24),c=n(20),l=n(149),f=n(45),d=n(92),h=n(126),p=n(125),m=n(151),v=n(148),g=n(53),_=n(146),y=n(147),b=n(23),w=n(35),x=n(153),k=n(82),M=n(152);function E(t,e){switch(arguments.length){case 0:break;case 1:this.range(t);break;default:this.range(e).domain(t)}return this}function S(t,e){switch(arguments.length){case 0:break;case 1:this.interpolator(t);break;default:this.interpolator(e).domain(t)}return this}var T=Array.prototype,j=T.map,O=T.slice,A={name:"implicit"};function L(){var t=Object(s.c)(),e=[],n=[],r=A;function i(i){var a=i+"",o=t.get(a);if(!o){if(r!==A)return r;t.set(a,o=e.push(i))}return n[(o-1)%n.length]}return i.domain=function(n){if(!arguments.length)return e.slice();e=[],t=Object(s.c)();for(var r,a,o=-1,u=n.length;++o<u;)t.has(a=(r=n[o])+"")||t.set(a,e.push(r));return i},i.range=function(t){return arguments.length?(n=O.call(t),i):n.slice()},i.unknown=function(t){return arguments.length?(r=t,i):r},i.copy=function(){return L(e,n).unknown(r)},E.apply(i,arguments),i}function C(){var t,e,n=L().unknown(void 0),r=n.domain,a=n.range,o=[0,1],u=!1,s=0,c=0,l=.5;function f(){var n=r().length,f=o[1]<o[0],d=o[f-0],h=o[1-f];t=(h-d)/Math.max(1,n-s+2*c),u&&(t=Math.floor(t)),d+=(h-d-t*(n-s))*l,e=t*(1-s),u&&(d=Math.round(d),e=Math.round(e));var p=Object(i.s)(n).map((function(e){return d+t*e}));return a(f?p.reverse():p)}return delete n.unknown,n.domain=function(t){return arguments.length?(r(t),f()):r()},n.range=function(t){return arguments.length?(o=[+t[0],+t[1]],f()):o.slice()},n.rangeRound=function(t){return o=[+t[0],+t[1]],u=!0,f()},n.bandwidth=function(){return e},n.step=function(){return t},n.round=function(t){return arguments.length?(u=!!t,f()):u},n.padding=function(t){return arguments.length?(s=Math.min(1,c=+t),f()):s},n.paddingInner=function(t){return arguments.length?(s=Math.min(1,t),f()):s},n.paddingOuter=function(t){return arguments.length?(c=+t,f()):c},n.align=function(t){return arguments.length?(l=Math.max(0,Math.min(1,t)),f()):l},n.copy=function(){return C(r(),o).round(u).paddingInner(s).paddingOuter(c).align(l)},E.apply(f(),arguments)}function D(t){var e=t.copy;return t.padding=t.paddingOuter,delete t.paddingInner,delete t.paddingOuter,t.copy=function(){return D(e())},t}function N(){return D(C.apply(null,arguments).paddingInner(1))}var P=n(98),R=n(25),Y=n(259),I=function(t){return+t},z=[0,1];function F(t){return t}function H(t,e){return(e-=t=+t)?function(n){return(n-t)/e}:(n=isNaN(e)?NaN:.5,function(){return n});var n}function B(t){var e,n=t[0],r=t[t.length-1];return n>r&&(e=n,n=r,r=e),function(t){return Math.max(n,Math.min(r,t))}}function q(t,e,n){var r=t[0],i=t[1],a=e[0],o=e[1];return i<r?(r=H(i,r),a=n(o,a)):(r=H(r,i),a=n(a,o)),function(t){return a(r(t))}}function U(t,e,n){var r=Math.min(t.length,e.length)-1,a=new Array(r),o=new Array(r),u=-1;for(t[r]<t[0]&&(t=t.slice().reverse(),e=e.slice().reverse());++u<r;)a[u]=H(t[u],t[u+1]),o[u]=n(e[u],e[u+1]);return function(e){var n=Object(i.b)(t,e,1,r)-1;return o[n](a[n](e))}}function W(t,e){return e.domain(t.domain()).range(t.range()).interpolate(t.interpolate()).clamp(t.clamp()).unknown(t.unknown())}function G(){var t,e,n,r,i,a,o=z,u=z,s=P.a,c=F;function l(){return r=Math.min(o.length,u.length)>2?U:q,i=a=null,f}function f(e){return isNaN(e=+e)?n:(i||(i=r(o.map(t),u,s)))(t(c(e)))}return f.invert=function(n){return c(e((a||(a=r(u,o.map(t),R.a)))(n)))},f.domain=function(t){return arguments.length?(o=j.call(t,I),c===F||(c=B(o)),l()):o.slice()},f.range=function(t){return arguments.length?(u=O.call(t),l()):u.slice()},f.rangeRound=function(t){return u=O.call(t),s=Y.a,l()},f.clamp=function(t){return arguments.length?(c=t?B(o):F,f):c!==F},f.interpolate=function(t){return arguments.length?(s=t,l()):s},f.unknown=function(t){return arguments.length?(n=t,f):n},function(n,r){return t=n,e=r,l()}}function V(t,e){return G()(t,e)}var $=n(101),X=n(257),J=n(134),Z=n(258),K=n(256),Q=function(t,e,n,r){var a,o=Object(i.A)(t,e,n);switch((r=Object($.b)(null==r?",f":r)).type){case"s":var u=Math.max(Math.abs(t),Math.abs(e));return null!=r.precision||isNaN(a=Object(X.a)(o,u))||(r.precision=a),Object(J.c)(r,u);case"":case"e":case"g":case"p":case"r":null!=r.precision||isNaN(a=Object(Z.a)(o,Math.max(Math.abs(t),Math.abs(e))))||(r.precision=a-("e"===r.type));break;case"f":case"%":null!=r.precision||isNaN(a=Object(K.a)(o))||(r.precision=a-2*("%"===r.type))}return Object(J.b)(r)};function tt(t){var e=t.domain;return t.ticks=function(t){var n=e();return Object(i.B)(n[0],n[n.length-1],null==t?10:t)},t.tickFormat=function(t,n){var r=e();return Q(r[0],r[r.length-1],null==t?10:t,n)},t.nice=function(n){null==n&&(n=10);var r,a=e(),o=0,u=a.length-1,s=a[o],c=a[u];return c<s&&(r=s,s=c,c=r,r=o,o=u,u=r),(r=Object(i.z)(s,c,n))>0?(s=Math.floor(s/r)*r,c=Math.ceil(c/r)*r,r=Object(i.z)(s,c,n)):r<0&&(s=Math.ceil(s*r)/r,c=Math.floor(c*r)/r,r=Object(i.z)(s,c,n)),r>0?(a[o]=Math.floor(s/r)*r,a[u]=Math.ceil(c/r)*r,e(a)):r<0&&(a[o]=Math.ceil(s*r)/r,a[u]=Math.floor(c*r)/r,e(a)),t},t}function et(){var t=V(F,F);return t.copy=function(){return W(t,et())},E.apply(t,arguments),tt(t)}function nt(t){var e;function n(t){return isNaN(t=+t)?e:t}return n.invert=n,n.domain=n.range=function(e){return arguments.length?(t=j.call(e,I),n):t.slice()},n.unknown=function(t){return arguments.length?(e=t,n):e},n.copy=function(){return nt(t).unknown(e)},t=arguments.length?j.call(t,I):[0,1],tt(n)}var rt=function(t,e){var n,r=0,i=(t=t.slice()).length-1,a=t[r],o=t[i];return o<a&&(n=r,r=i,i=n,n=a,a=o,o=n),t[r]=e.floor(a),t[i]=e.ceil(o),t};function it(t){return Math.log(t)}function at(t){return Math.exp(t)}function ot(t){return-Math.log(-t)}function ut(t){return-Math.exp(-t)}function st(t){return isFinite(t)?+("1e"+t):t<0?0:t}function ct(t){return function(e){return-t(-e)}}function lt(t){var e,n,r=t(it,at),a=r.domain,o=10;function u(){return e=function(t){return t===Math.E?Math.log:10===t&&Math.log10||2===t&&Math.log2||(t=Math.log(t),function(e){return Math.log(e)/t})}(o),n=function(t){return 10===t?st:t===Math.E?Math.exp:function(e){return Math.pow(t,e)}}(o),a()[0]<0?(e=ct(e),n=ct(n),t(ot,ut)):t(it,at),r}return r.base=function(t){return arguments.length?(o=+t,u()):o},r.domain=function(t){return arguments.length?(a(t),u()):a()},r.ticks=function(t){var r,u=a(),s=u[0],c=u[u.length-1];(r=c<s)&&(h=s,s=c,c=h);var l,f,d,h=e(s),p=e(c),m=null==t?10:+t,v=[];if(!(o%1)&&p-h<m){if(h=Math.round(h)-1,p=Math.round(p)+1,s>0){for(;h<p;++h)for(f=1,l=n(h);f<o;++f)if(!((d=l*f)<s)){if(d>c)break;v.push(d)}}else for(;h<p;++h)for(f=o-1,l=n(h);f>=1;--f)if(!((d=l*f)<s)){if(d>c)break;v.push(d)}}else v=Object(i.B)(h,p,Math.min(p-h,m)).map(n);return r?v.reverse():v},r.tickFormat=function(t,i){if(null==i&&(i=10===o?".0e":","),"function"!=typeof i&&(i=Object(J.b)(i)),t===1/0)return i;null==t&&(t=10);var a=Math.max(1,o*t/r.ticks().length);return function(t){var r=t/n(Math.round(e(t)));return r*o<o-.5&&(r*=o),r<=a?i(t):""}},r.nice=function(){return a(rt(a(),{floor:function(t){return n(Math.floor(e(t)))},ceil:function(t){return n(Math.ceil(e(t)))}}))},r}function ft(){var t=lt(G()).domain([1,10]);return t.copy=function(){return W(t,ft()).base(t.base())},E.apply(t,arguments),t}function dt(t){return function(e){return Math.sign(e)*Math.log1p(Math.abs(e/t))}}function ht(t){return function(e){return Math.sign(e)*Math.expm1(Math.abs(e))*t}}function pt(t){var e=1,n=t(dt(e),ht(e));return n.constant=function(n){return arguments.length?t(dt(e=+n),ht(e)):e},tt(n)}function mt(){var t=pt(G());return t.copy=function(){return W(t,mt()).constant(t.constant())},E.apply(t,arguments)}function vt(t){return function(e){return e<0?-Math.pow(-e,t):Math.pow(e,t)}}function gt(t){return t<0?-Math.sqrt(-t):Math.sqrt(t)}function _t(t){return t<0?-t*t:t*t}function yt(t){var e=t(F,F),n=1;function r(){return 1===n?t(F,F):.5===n?t(gt,_t):t(vt(n),vt(1/n))}return e.exponent=function(t){return arguments.length?(n=+t,r()):n},tt(e)}function bt(){var t=yt(G());return t.copy=function(){return W(t,bt()).exponent(t.exponent())},E.apply(t,arguments),t}function wt(){return bt.apply(null,arguments).exponent(.5)}function xt(){var t,e=[],n=[],r=[];function a(){var t=0,a=Math.max(1,n.length);for(r=new Array(a-1);++t<a;)r[t-1]=Object(i.r)(e,t/a);return o}function o(e){return isNaN(e=+e)?t:n[Object(i.b)(r,e)]}return o.invertExtent=function(t){var i=n.indexOf(t);return i<0?[NaN,NaN]:[i>0?r[i-1]:e[0],i<r.length?r[i]:e[e.length-1]]},o.domain=function(t){if(!arguments.length)return e.slice();e=[];for(var n,r=0,o=t.length;r<o;++r)null==(n=t[r])||isNaN(n=+n)||e.push(n);return e.sort(i.a),a()},o.range=function(t){return arguments.length?(n=O.call(t),a()):n.slice()},o.unknown=function(e){return arguments.length?(t=e,o):t},o.quantiles=function(){return r.slice()},o.copy=function(){return xt().domain(e).range(n).unknown(t)},E.apply(o,arguments)}function kt(){var t,e=0,n=1,r=1,a=[.5],o=[0,1];function u(e){return e<=e?o[Object(i.b)(a,e,0,r)]:t}function s(){var t=-1;for(a=new Array(r);++t<r;)a[t]=((t+1)*n-(t-r)*e)/(r+1);return u}return u.domain=function(t){return arguments.length?(e=+t[0],n=+t[1],s()):[e,n]},u.range=function(t){return arguments.length?(r=(o=O.call(t)).length-1,s()):o.slice()},u.invertExtent=function(t){var i=o.indexOf(t);return i<0?[NaN,NaN]:i<1?[e,a[0]]:i>=r?[a[r-1],n]:[a[i-1],a[i]]},u.unknown=function(e){return arguments.length?(t=e,u):u},u.thresholds=function(){return a.slice()},u.copy=function(){return kt().domain([e,n]).range(o).unknown(t)},E.apply(tt(u),arguments)}function Mt(){var t,e=[.5],n=[0,1],r=1;function a(a){return a<=a?n[Object(i.b)(e,a,0,r)]:t}return a.domain=function(t){return arguments.length?(e=O.call(t),r=Math.min(e.length,n.length-1),a):e.slice()},a.range=function(t){return arguments.length?(n=O.call(t),r=Math.min(e.length,n.length-1),a):n.slice()},a.invertExtent=function(t){var r=n.indexOf(t);return[e[r-1],e[r]]},a.unknown=function(e){return arguments.length?(t=e,a):t},a.copy=function(){return Mt().domain(e).range(n).unknown(t)},E.apply(a,arguments)}var Et=n(64),St=n(180),Tt=n(12),jt=n(127),Ot=n(181),At=n(182),Lt=n(112),Ct=n(113),Dt=n(41);function Nt(t){return new Date(t)}function Pt(t){return t instanceof Date?+t:+new Date(+t)}function Rt(t,e,n,r,a,o,u,s,c){var l=V(F,F),f=l.invert,d=l.domain,h=c(".%L"),p=c(":%S"),m=c("%I:%M"),v=c("%I %p"),g=c("%a %d"),_=c("%b %d"),y=c("%B"),b=c("%Y"),w=[[u,1,1e3],[u,5,5e3],[u,15,15e3],[u,30,3e4],[o,1,6e4],[o,5,3e5],[o,15,9e5],[o,30,18e5],[a,1,36e5],[a,3,108e5],[a,6,216e5],[a,12,432e5],[r,1,864e5],[r,2,1728e5],[n,1,6048e5],[e,1,2592e6],[e,3,7776e6],[t,1,31536e6]];function x(i){return(u(i)<i?h:o(i)<i?p:a(i)<i?m:r(i)<i?v:e(i)<i?n(i)<i?g:_:t(i)<i?y:b)(i)}function k(e,n,r,a){if(null==e&&(e=10),"number"==typeof e){var o=Math.abs(r-n)/e,u=Object(i.e)((function(t){return t[2]})).right(w,o);u===w.length?(a=Object(i.A)(n/31536e6,r/31536e6,e),e=t):u?(a=(u=w[o/w[u-1][2]<w[u][2]/o?u-1:u])[1],e=u[0]):(a=Math.max(Object(i.A)(n,r,e),1),e=s)}return null==a?e:e.every(a)}return l.invert=function(t){return new Date(f(t))},l.domain=function(t){return arguments.length?d(j.call(t,Pt)):d().map(Nt)},l.ticks=function(t,e){var n,r=d(),i=r[0],a=r[r.length-1],o=a<i;return o&&(n=i,i=a,a=n),n=(n=k(t,i,a,e))?n.range(i,a+1):[],o?n.reverse():n},l.tickFormat=function(t,e){return null==e?x:c(e)},l.nice=function(t,e){var n=d();return(t=k(t,n[0],n[n.length-1],e))?d(rt(n,t)):l},l.copy=function(){return W(l,Rt(t,e,n,r,a,o,u,s,c))},l}var Yt=function(){return E.apply(Rt(Et.a,St.a,Tt.g,jt.b,Ot.a,At.a,Lt.a,Ct.a,Dt.b).domain([new Date(2e3,0,1),new Date(2e3,0,2)]),arguments)},It=n(65),zt=n(183),Ft=n(13),Ht=n(128),Bt=n(184),qt=n(185),Ut=function(){return E.apply(Rt(It.a,zt.a,Ft.g,Ht.a,Bt.a,qt.a,Lt.a,Ct.a,Dt.d).domain([Date.UTC(2e3,0,1),Date.UTC(2e3,0,2)]),arguments)};function Wt(){var t,e,n,r,i,a=0,o=1,u=F,s=!1;function c(e){return isNaN(e=+e)?i:u(0===n?.5:(e=(r(e)-t)*n,s?Math.max(0,Math.min(1,e)):e))}return c.domain=function(i){return arguments.length?(t=r(a=+i[0]),e=r(o=+i[1]),n=t===e?0:1/(e-t),c):[a,o]},c.clamp=function(t){return arguments.length?(s=!!t,c):s},c.interpolator=function(t){return arguments.length?(u=t,c):u},c.unknown=function(t){return arguments.length?(i=t,c):i},function(i){return r=i,t=i(a),e=i(o),n=t===e?0:1/(e-t),c}}function Gt(t,e){return e.domain(t.domain()).interpolator(t.interpolator()).clamp(t.clamp()).unknown(t.unknown())}function Vt(){var t=tt(Wt()(F));return t.copy=function(){return Gt(t,Vt())},S.apply(t,arguments)}function $t(){var t=lt(Wt()).domain([1,10]);return t.copy=function(){return Gt(t,$t()).base(t.base())},S.apply(t,arguments)}function Xt(){var t=pt(Wt());return t.copy=function(){return Gt(t,Xt()).constant(t.constant())},S.apply(t,arguments)}function Jt(){var t=yt(Wt());return t.copy=function(){return Gt(t,Jt()).exponent(t.exponent())},S.apply(t,arguments)}function Zt(){return Jt.apply(null,arguments).exponent(.5)}function Kt(){var t=[],e=F;function n(n){if(!isNaN(n=+n))return e((Object(i.b)(t,n)-1)/(t.length-1))}return n.domain=function(e){if(!arguments.length)return t.slice();t=[];for(var r,a=0,o=e.length;a<o;++a)null==(r=e[a])||isNaN(r=+r)||t.push(r);return t.sort(i.a),n},n.interpolator=function(t){return arguments.length?(e=t,n):e},n.copy=function(){return Kt(e).domain(t)},S.apply(n,arguments)}function Qt(){var t,e,n,r,i,a,o,u=0,s=.5,c=1,l=F,f=!1;function d(t){return isNaN(t=+t)?o:(t=.5+((t=+a(t))-e)*(t<e?r:i),l(f?Math.max(0,Math.min(1,t)):t))}return d.domain=function(o){return arguments.length?(t=a(u=+o[0]),e=a(s=+o[1]),n=a(c=+o[2]),r=t===e?0:.5/(e-t),i=e===n?0:.5/(n-e),d):[u,s,c]},d.clamp=function(t){return arguments.length?(f=!!t,d):f},d.interpolator=function(t){return arguments.length?(l=t,d):l},d.unknown=function(t){return arguments.length?(o=t,d):o},function(o){return a=o,t=o(u),e=o(s),n=o(c),r=t===e?0:.5/(e-t),i=e===n?0:.5/(n-e),d}}function te(){var t=tt(Qt()(F));return t.copy=function(){return Gt(t,te())},S.apply(t,arguments)}function ee(){var t=lt(Qt()).domain([.1,1,10]);return t.copy=function(){return Gt(t,ee()).base(t.base())},S.apply(t,arguments)}function ne(){var t=pt(Qt());return t.copy=function(){return Gt(t,ne()).constant(t.constant())},S.apply(t,arguments)}function re(){var t=yt(Qt());return t.copy=function(){return Gt(t,re()).exponent(t.exponent())},S.apply(t,arguments)}function ie(){return re.apply(null,arguments).exponent(.5)}var ae=n(124),oe=n(16),ue=n(123),se=n(46),ce=n(40),le=n(77),fe=n(32),de=n(150),he=n(155)},function(t,e,n){"use strict";n.d(e,"a",(function(){return v})),n.d(e,"b",(function(){return s.a})),n.d(e,"c",(function(){return s.b}));var r=n(252),i=n(42),a=n(255),o=n(106),u=n(254),s=n(99),c=n(33),l=function(t){return function(){return t}};function f(t,e,n,r,i,a,o,u,s,c){this.target=t,this.type=e,this.subject=n,this.identifier=r,this.active=i,this.x=a,this.y=o,this.dx=u,this.dy=s,this._=c}function d(){return!i.c.ctrlKey&&!i.c.button}function h(){return this.parentNode}function p(t){return null==t?{x:i.c.x,y:i.c.y}:t}function m(){return navigator.maxTouchPoints||"ontouchstart"in this}f.prototype.on=function(){var t=this._.on.apply(this._,arguments);return t===this._?this:t};var v=function(){var t,e,n,v,g=d,_=h,y=p,b=m,w={},x=Object(r.a)("start","drag","end"),k=0,M=0;function E(t){t.on("mousedown.drag",S).filter(b).on("touchstart.drag",O).on("touchmove.drag",A).on("touchend.drag touchcancel.drag",L).style("touch-action","none").style("-webkit-tap-highlight-color","rgba(0,0,0,0)")}function S(){if(!v&&g.apply(this,arguments)){var r=C("mouse",_.apply(this,arguments),a.a,this,arguments);r&&(Object(o.a)(i.c.view).on("mousemove.drag",T,!0).on("mouseup.drag",j,!0),Object(s.a)(i.c.view),Object(c.b)(),n=!1,t=i.c.clientX,e=i.c.clientY,r("start"))}}function T(){if(Object(c.a)(),!n){var r=i.c.clientX-t,a=i.c.clientY-e;n=r*r+a*a>M}w.mouse("drag")}function j(){Object(o.a)(i.c.view).on("mousemove.drag mouseup.drag",null),Object(s.b)(i.c.view,n),Object(c.a)(),w.mouse("end")}function O(){if(g.apply(this,arguments)){var t,e,n=i.c.changedTouches,r=_.apply(this,arguments),a=n.length;for(t=0;t<a;++t)(e=C(n[t].identifier,r,u.a,this,arguments))&&(Object(c.b)(),e("start"))}}function A(){var t,e,n=i.c.changedTouches,r=n.length;for(t=0;t<r;++t)(e=w[n[t].identifier])&&(Object(c.a)(),e("drag"))}function L(){var t,e,n=i.c.changedTouches,r=n.length;for(v&&clearTimeout(v),v=setTimeout((function(){v=null}),500),t=0;t<r;++t)(e=w[n[t].identifier])&&(Object(c.b)(),e("end"))}function C(t,e,n,r,a){var o,u,s,c=n(e,t),l=x.copy();if(Object(i.a)(new f(E,"beforestart",o,t,k,c[0],c[1],0,0,l),(function(){return null!=(i.c.subject=o=y.apply(r,a))&&(u=o.x-c[0]||0,s=o.y-c[1]||0,!0)})))return function d(h){var p,m=c;switch(h){case"start":w[t]=d,p=k++;break;case"end":delete w[t],--k;case"drag":c=n(e,t),p=k}Object(i.a)(new f(E,h,o,t,p,c[0]+u,c[1]+s,c[0]-m[0],c[1]-m[1],l),l.apply,l,[h,r,a])}}return E.filter=function(t){return arguments.length?(g="function"==typeof t?t:l(!!t),E):g},E.container=function(t){return arguments.length?(_="function"==typeof t?t:l(t),E):_},E.subject=function(t){return arguments.length?(y="function"==typeof t?t:l(t),E):y},E.touchable=function(t){return arguments.length?(b="function"==typeof t?t:l(!!t),E):b},E.on=function(){var t=x.on.apply(x,arguments);return t===x?E:t},E.clickDistance=function(t){return arguments.length?(M=(t=+t)*t,E):Math.sqrt(M)},E}},function(t,e,n){"use strict";n.r(e),function(t){n.d(e,"default",(function(){return g})),n.d(e,"VERSION",(function(){return _})),n.d(e,"iteratee",(function(){return w})),n.d(e,"restArguments",(function(){return k})),n.d(e,"each",(function(){return L})),n.d(e,"forEach",(function(){return L})),n.d(e,"map",(function(){return C})),n.d(e,"collect",(function(){return C})),n.d(e,"reduce",(function(){return N})),n.d(e,"foldl",(function(){return N})),n.d(e,"inject",(function(){return N})),n.d(e,"reduceRight",(function(){return P})),n.d(e,"foldr",(function(){return P})),n.d(e,"find",(function(){return R})),n.d(e,"detect",(function(){return R})),n.d(e,"filter",(function(){return Y})),n.d(e,"select",(function(){return Y})),n.d(e,"reject",(function(){return I})),n.d(e,"every",(function(){return z})),n.d(e,"all",(function(){return z})),n.d(e,"some",(function(){return F})),n.d(e,"any",(function(){return F})),n.d(e,"contains",(function(){return H})),n.d(e,"includes",(function(){return H})),n.d(e,"include",(function(){return H})),n.d(e,"invoke",(function(){return B})),n.d(e,"pluck",(function(){return q})),n.d(e,"where",(function(){return U})),n.d(e,"findWhere",(function(){return W})),n.d(e,"max",(function(){return G})),n.d(e,"min",(function(){return V})),n.d(e,"shuffle",(function(){return $})),n.d(e,"sample",(function(){return X})),n.d(e,"sortBy",(function(){return J})),n.d(e,"groupBy",(function(){return K})),n.d(e,"indexBy",(function(){return Q})),n.d(e,"countBy",(function(){return tt})),n.d(e,"toArray",(function(){return nt})),n.d(e,"size",(function(){return rt})),n.d(e,"partition",(function(){return it})),n.d(e,"first",(function(){return at})),n.d(e,"head",(function(){return at})),n.d(e,"take",(function(){return at})),n.d(e,"initial",(function(){return ot})),n.d(e,"last",(function(){return ut})),n.d(e,"rest",(function(){return st})),n.d(e,"tail",(function(){return st})),n.d(e,"drop",(function(){return st})),n.d(e,"compact",(function(){return ct})),n.d(e,"flatten",(function(){return ft})),n.d(e,"without",(function(){return dt})),n.d(e,"uniq",(function(){return ht})),n.d(e,"unique",(function(){return ht})),n.d(e,"union",(function(){return pt})),n.d(e,"intersection",(function(){return mt})),n.d(e,"difference",(function(){return vt})),n.d(e,"unzip",(function(){return gt})),n.d(e,"zip",(function(){return _t})),n.d(e,"object",(function(){return yt})),n.d(e,"findIndex",(function(){return wt})),n.d(e,"findLastIndex",(function(){return xt})),n.d(e,"sortedIndex",(function(){return kt})),n.d(e,"indexOf",(function(){return Et})),n.d(e,"lastIndexOf",(function(){return St})),n.d(e,"range",(function(){return Tt})),n.d(e,"chunk",(function(){return jt})),n.d(e,"bind",(function(){return At})),n.d(e,"partial",(function(){return Lt})),n.d(e,"bindAll",(function(){return Ct})),n.d(e,"memoize",(function(){return Dt})),n.d(e,"delay",(function(){return Nt})),n.d(e,"defer",(function(){return Pt})),n.d(e,"throttle",(function(){return Rt})),n.d(e,"debounce",(function(){return Yt})),n.d(e,"wrap",(function(){return It})),n.d(e,"negate",(function(){return zt})),n.d(e,"compose",(function(){return Ft})),n.d(e,"after",(function(){return Ht})),n.d(e,"before",(function(){return Bt})),n.d(e,"once",(function(){return qt})),n.d(e,"keys",(function(){return Vt})),n.d(e,"allKeys",(function(){return $t})),n.d(e,"values",(function(){return Xt})),n.d(e,"mapObject",(function(){return Jt})),n.d(e,"pairs",(function(){return Zt})),n.d(e,"invert",(function(){return Kt})),n.d(e,"functions",(function(){return Qt})),n.d(e,"methods",(function(){return Qt})),n.d(e,"extend",(function(){return ee})),n.d(e,"extendOwn",(function(){return ne})),n.d(e,"assign",(function(){return ne})),n.d(e,"findKey",(function(){return re})),n.d(e,"pick",(function(){return ae})),n.d(e,"omit",(function(){return oe})),n.d(e,"defaults",(function(){return ue})),n.d(e,"create",(function(){return se})),n.d(e,"clone",(function(){return ce})),n.d(e,"tap",(function(){return le})),n.d(e,"isMatch",(function(){return fe})),n.d(e,"isEqual",(function(){return he})),n.d(e,"isEmpty",(function(){return pe})),n.d(e,"isElement",(function(){return me})),n.d(e,"isArray",(function(){return ge})),n.d(e,"isObject",(function(){return _e})),n.d(e,"isArguments",(function(){return ye})),n.d(e,"isFunction",(function(){return be})),n.d(e,"isString",(function(){return we})),n.d(e,"isNumber",(function(){return xe})),n.d(e,"isDate",(function(){return ke})),n.d(e,"isRegExp",(function(){return Me})),n.d(e,"isError",(function(){return Ee})),n.d(e,"isSymbol",(function(){return Se})),n.d(e,"isMap",(function(){return Te})),n.d(e,"isWeakMap",(function(){return je})),n.d(e,"isSet",(function(){return Oe})),n.d(e,"isWeakSet",(function(){return Ae})),n.d(e,"isFinite",(function(){return Ce})),n.d(e,"isNaN",(function(){return De})),n.d(e,"isBoolean",(function(){return Ne})),n.d(e,"isNull",(function(){return Pe})),n.d(e,"isUndefined",(function(){return Re})),n.d(e,"has",(function(){return Ye})),n.d(e,"identity",(function(){return Ie})),n.d(e,"constant",(function(){return ze})),n.d(e,"noop",(function(){return Fe})),n.d(e,"property",(function(){return He})),n.d(e,"propertyOf",(function(){return Be})),n.d(e,"matcher",(function(){return qe})),n.d(e,"matches",(function(){return qe})),n.d(e,"times",(function(){return Ue})),n.d(e,"random",(function(){return We})),n.d(e,"now",(function(){return Ge})),n.d(e,"escape",(function(){return Je})),n.d(e,"unescape",(function(){return Ze})),n.d(e,"result",(function(){return Ke})),n.d(e,"uniqueId",(function(){return tn})),n.d(e,"templateSettings",(function(){return en})),n.d(e,"template",(function(){return un})),n.d(e,"chain",(function(){return sn})),n.d(e,"mixin",(function(){return ln}));var r="object"==typeof self&&self.self===self&&self||"object"==typeof t&&t.global===t&&t||Function("return this")()||{},i=Array.prototype,a=Object.prototype,o="undefined"!=typeof Symbol?Symbol.prototype:null,u=i.push,s=i.slice,c=a.toString,l=a.hasOwnProperty,f=Array.isArray,d=Object.keys,h=Object.create,p=r.isNaN,m=r.isFinite,v=function(){};function g(t){return t instanceof g?t:this instanceof g?void(this._wrapped=t):new g(t)}var _=g.VERSION="1.10.2";function y(t,e,n){if(void 0===e)return t;switch(null==n?3:n){case 1:return function(n){return t.call(e,n)};case 3:return function(n,r,i){return t.call(e,n,r,i)};case 4:return function(n,r,i,a){return t.call(e,n,r,i,a)}}return function(){return t.apply(e,arguments)}}function b(t,e,n){return null==t?Ie:be(t)?y(t,e,n):_e(t)&&!ge(t)?qe(t):He(t)}function w(t,e){return b(t,e,1/0)}function x(t,e,n){return g.iteratee!==w?g.iteratee(t,e):b(t,e,n)}function k(t,e){return e=null==e?t.length-1:+e,function(){for(var n=Math.max(arguments.length-e,0),r=Array(n),i=0;i<n;i++)r[i]=arguments[i+e];switch(e){case 0:return t.call(this,r);case 1:return t.call(this,arguments[0],r);case 2:return t.call(this,arguments[0],arguments[1],r)}var a=Array(e+1);for(i=0;i<e;i++)a[i]=arguments[i];return a[e]=r,t.apply(this,a)}}function M(t){if(!_e(t))return{};if(h)return h(t);v.prototype=t;var e=new v;return v.prototype=null,e}function E(t){return function(e){return null==e?void 0:e[t]}}function S(t,e){return null!=t&&l.call(t,e)}function T(t,e){for(var n=e.length,r=0;r<n;r++){if(null==t)return;t=t[e[r]]}return n?t:void 0}g.iteratee=w;var j=Math.pow(2,53)-1,O=E("length");function A(t){var e=O(t);return"number"==typeof e&&e>=0&&e<=j}function L(t,e,n){var r,i;if(e=y(e,n),A(t))for(r=0,i=t.length;r<i;r++)e(t[r],r,t);else{var a=Vt(t);for(r=0,i=a.length;r<i;r++)e(t[a[r]],a[r],t)}return t}function C(t,e,n){e=x(e,n);for(var r=!A(t)&&Vt(t),i=(r||t).length,a=Array(i),o=0;o<i;o++){var u=r?r[o]:o;a[o]=e(t[u],u,t)}return a}function D(t){var e=function(e,n,r,i){var a=!A(e)&&Vt(e),o=(a||e).length,u=t>0?0:o-1;for(i||(r=e[a?a[u]:u],u+=t);u>=0&&u<o;u+=t){var s=a?a[u]:u;r=n(r,e[s],s,e)}return r};return function(t,n,r,i){var a=arguments.length>=3;return e(t,y(n,i,4),r,a)}}var N=D(1),P=D(-1);function R(t,e,n){var r=(A(t)?wt:re)(t,e,n);if(void 0!==r&&-1!==r)return t[r]}function Y(t,e,n){var r=[];return e=x(e,n),L(t,(function(t,n,i){e(t,n,i)&&r.push(t)})),r}function I(t,e,n){return Y(t,zt(x(e)),n)}function z(t,e,n){e=x(e,n);for(var r=!A(t)&&Vt(t),i=(r||t).length,a=0;a<i;a++){var o=r?r[a]:a;if(!e(t[o],o,t))return!1}return!0}function F(t,e,n){e=x(e,n);for(var r=!A(t)&&Vt(t),i=(r||t).length,a=0;a<i;a++){var o=r?r[a]:a;if(e(t[o],o,t))return!0}return!1}function H(t,e,n,r){return A(t)||(t=Xt(t)),("number"!=typeof n||r)&&(n=0),Et(t,e,n)>=0}var B=k((function(t,e,n){var r,i;return be(e)?i=e:ge(e)&&(r=e.slice(0,-1),e=e[e.length-1]),C(t,(function(t){var a=i;if(!a){if(r&&r.length&&(t=T(t,r)),null==t)return;a=t[e]}return null==a?a:a.apply(t,n)}))}));function q(t,e){return C(t,He(e))}function U(t,e){return Y(t,qe(e))}function W(t,e){return R(t,qe(e))}function G(t,e,n){var r,i,a=-1/0,o=-1/0;if(null==e||"number"==typeof e&&"object"!=typeof t[0]&&null!=t)for(var u=0,s=(t=A(t)?t:Xt(t)).length;u<s;u++)null!=(r=t[u])&&r>a&&(a=r);else e=x(e,n),L(t,(function(t,n,r){((i=e(t,n,r))>o||i===-1/0&&a===-1/0)&&(a=t,o=i)}));return a}function V(t,e,n){var r,i,a=1/0,o=1/0;if(null==e||"number"==typeof e&&"object"!=typeof t[0]&&null!=t)for(var u=0,s=(t=A(t)?t:Xt(t)).length;u<s;u++)null!=(r=t[u])&&r<a&&(a=r);else e=x(e,n),L(t,(function(t,n,r){((i=e(t,n,r))<o||i===1/0&&a===1/0)&&(a=t,o=i)}));return a}function $(t){return X(t,1/0)}function X(t,e,n){if(null==e||n)return A(t)||(t=Xt(t)),t[We(t.length-1)];var r=A(t)?ce(t):Xt(t),i=O(r);e=Math.max(Math.min(e,i),0);for(var a=i-1,o=0;o<e;o++){var u=We(o,a),s=r[o];r[o]=r[u],r[u]=s}return r.slice(0,e)}function J(t,e,n){var r=0;return e=x(e,n),q(C(t,(function(t,n,i){return{value:t,index:r++,criteria:e(t,n,i)}})).sort((function(t,e){var n=t.criteria,r=e.criteria;if(n!==r){if(n>r||void 0===n)return 1;if(n<r||void 0===r)return-1}return t.index-e.index})),"value")}function Z(t,e){return function(n,r,i){var a=e?[[],[]]:{};return r=x(r,i),L(n,(function(e,i){var o=r(e,i,n);t(a,e,o)})),a}}var K=Z((function(t,e,n){S(t,n)?t[n].push(e):t[n]=[e]})),Q=Z((function(t,e,n){t[n]=e})),tt=Z((function(t,e,n){S(t,n)?t[n]++:t[n]=1})),et=/[^\ud800-\udfff]|[\ud800-\udbff][\udc00-\udfff]|[\ud800-\udfff]/g;function nt(t){return t?ge(t)?s.call(t):we(t)?t.match(et):A(t)?C(t,Ie):Xt(t):[]}function rt(t){return null==t?0:A(t)?t.length:Vt(t).length}var it=Z((function(t,e,n){t[n?0:1].push(e)}),!0);function at(t,e,n){return null==t||t.length<1?null==e?void 0:[]:null==e||n?t[0]:ot(t,t.length-e)}function ot(t,e,n){return s.call(t,0,Math.max(0,t.length-(null==e||n?1:e)))}function ut(t,e,n){return null==t||t.length<1?null==e?void 0:[]:null==e||n?t[t.length-1]:st(t,Math.max(0,t.length-e))}function st(t,e,n){return s.call(t,null==e||n?1:e)}function ct(t){return Y(t,Boolean)}function lt(t,e,n,r){for(var i=(r=r||[]).length,a=0,o=O(t);a<o;a++){var u=t[a];if(A(u)&&(ge(u)||ye(u)))if(e)for(var s=0,c=u.length;s<c;)r[i++]=u[s++];else lt(u,e,n,r),i=r.length;else n||(r[i++]=u)}return r}function ft(t,e){return lt(t,e,!1)}var dt=k((function(t,e){return vt(t,e)}));function ht(t,e,n,r){Ne(e)||(r=n,n=e,e=!1),null!=n&&(n=x(n,r));for(var i=[],a=[],o=0,u=O(t);o<u;o++){var s=t[o],c=n?n(s,o,t):s;e&&!n?(o&&a===c||i.push(s),a=c):n?H(a,c)||(a.push(c),i.push(s)):H(i,s)||i.push(s)}return i}var pt=k((function(t){return ht(lt(t,!0,!0))}));function mt(t){for(var e=[],n=arguments.length,r=0,i=O(t);r<i;r++){var a=t[r];if(!H(e,a)){var o;for(o=1;o<n&&H(arguments[o],a);o++);o===n&&e.push(a)}}return e}var vt=k((function(t,e){return e=lt(e,!0,!0),Y(t,(function(t){return!H(e,t)}))}));function gt(t){for(var e=t&&G(t,O).length||0,n=Array(e),r=0;r<e;r++)n[r]=q(t,r);return n}var _t=k(gt);function yt(t,e){for(var n={},r=0,i=O(t);r<i;r++)e?n[t[r]]=e[r]:n[t[r][0]]=t[r][1];return n}function bt(t){return function(e,n,r){n=x(n,r);for(var i=O(e),a=t>0?0:i-1;a>=0&&a<i;a+=t)if(n(e[a],a,e))return a;return-1}}var wt=bt(1),xt=bt(-1);function kt(t,e,n,r){for(var i=(n=x(n,r,1))(e),a=0,o=O(t);a<o;){var u=Math.floor((a+o)/2);n(t[u])<i?a=u+1:o=u}return a}function Mt(t,e,n){return function(r,i,a){var o=0,u=O(r);if("number"==typeof a)t>0?o=a>=0?a:Math.max(a+u,o):u=a>=0?Math.min(a+1,u):a+u+1;else if(n&&a&&u)return r[a=n(r,i)]===i?a:-1;if(i!=i)return(a=e(s.call(r,o,u),De))>=0?a+o:-1;for(a=t>0?o:u-1;a>=0&&a<u;a+=t)if(r[a]===i)return a;return-1}}var Et=Mt(1,wt,kt),St=Mt(-1,xt);function Tt(t,e,n){null==e&&(e=t||0,t=0),n||(n=e<t?-1:1);for(var r=Math.max(Math.ceil((e-t)/n),0),i=Array(r),a=0;a<r;a++,t+=n)i[a]=t;return i}function jt(t,e){if(null==e||e<1)return[];for(var n=[],r=0,i=t.length;r<i;)n.push(s.call(t,r,r+=e));return n}function Ot(t,e,n,r,i){if(!(r instanceof e))return t.apply(n,i);var a=M(t.prototype),o=t.apply(a,i);return _e(o)?o:a}var At=k((function(t,e,n){if(!be(t))throw new TypeError("Bind must be called on a function");var r=k((function(i){return Ot(t,r,e,this,n.concat(i))}));return r})),Lt=k((function(t,e){var n=Lt.placeholder,r=function(){for(var i=0,a=e.length,o=Array(a),u=0;u<a;u++)o[u]=e[u]===n?arguments[i++]:e[u];for(;i<arguments.length;)o.push(arguments[i++]);return Ot(t,r,this,this,o)};return r}));Lt.placeholder=g;var Ct=k((function(t,e){var n=(e=lt(e,!1,!1)).length;if(n<1)throw new Error("bindAll must be passed function names");for(;n--;){var r=e[n];t[r]=At(t[r],t)}}));function Dt(t,e){var n=function(r){var i=n.cache,a=""+(e?e.apply(this,arguments):r);return S(i,a)||(i[a]=t.apply(this,arguments)),i[a]};return n.cache={},n}var Nt=k((function(t,e,n){return setTimeout((function(){return t.apply(null,n)}),e)})),Pt=Lt(Nt,g,1);function Rt(t,e,n){var r,i,a,o,u=0;n||(n={});var s=function(){u=!1===n.leading?0:Ge(),r=null,o=t.apply(i,a),r||(i=a=null)},c=function(){var c=Ge();u||!1!==n.leading||(u=c);var l=e-(c-u);return i=this,a=arguments,l<=0||l>e?(r&&(clearTimeout(r),r=null),u=c,o=t.apply(i,a),r||(i=a=null)):r||!1===n.trailing||(r=setTimeout(s,l)),o};return c.cancel=function(){clearTimeout(r),u=0,r=i=a=null},c}function Yt(t,e,n){var r,i,a=function(e,n){r=null,n&&(i=t.apply(e,n))},o=k((function(o){if(r&&clearTimeout(r),n){var u=!r;r=setTimeout(a,e),u&&(i=t.apply(this,o))}else r=Nt(a,e,this,o);return i}));return o.cancel=function(){clearTimeout(r),r=null},o}function It(t,e){return Lt(e,t)}function zt(t){return function(){return!t.apply(this,arguments)}}function Ft(){var t=arguments,e=t.length-1;return function(){for(var n=e,r=t[e].apply(this,arguments);n--;)r=t[n].call(this,r);return r}}function Ht(t,e){return function(){if(--t<1)return e.apply(this,arguments)}}function Bt(t,e){var n;return function(){return--t>0&&(n=e.apply(this,arguments)),t<=1&&(e=null),n}}var qt=Lt(Bt,2),Ut=!{toString:null}.propertyIsEnumerable("toString"),Wt=["valueOf","isPrototypeOf","toString","propertyIsEnumerable","hasOwnProperty","toLocaleString"];function Gt(t,e){var n=Wt.length,r=t.constructor,i=be(r)&&r.prototype||a,o="constructor";for(S(t,o)&&!H(e,o)&&e.push(o);n--;)(o=Wt[n])in t&&t[o]!==i[o]&&!H(e,o)&&e.push(o)}function Vt(t){if(!_e(t))return[];if(d)return d(t);var e=[];for(var n in t)S(t,n)&&e.push(n);return Ut&&Gt(t,e),e}function $t(t){if(!_e(t))return[];var e=[];for(var n in t)e.push(n);return Ut&&Gt(t,e),e}function Xt(t){for(var e=Vt(t),n=e.length,r=Array(n),i=0;i<n;i++)r[i]=t[e[i]];return r}function Jt(t,e,n){e=x(e,n);for(var r=Vt(t),i=r.length,a={},o=0;o<i;o++){var u=r[o];a[u]=e(t[u],u,t)}return a}function Zt(t){for(var e=Vt(t),n=e.length,r=Array(n),i=0;i<n;i++)r[i]=[e[i],t[e[i]]];return r}function Kt(t){for(var e={},n=Vt(t),r=0,i=n.length;r<i;r++)e[t[n[r]]]=n[r];return e}function Qt(t){var e=[];for(var n in t)be(t[n])&&e.push(n);return e.sort()}function te(t,e){return function(n){var r=arguments.length;if(e&&(n=Object(n)),r<2||null==n)return n;for(var i=1;i<r;i++)for(var a=arguments[i],o=t(a),u=o.length,s=0;s<u;s++){var c=o[s];e&&void 0!==n[c]||(n[c]=a[c])}return n}}var ee=te($t),ne=te(Vt);function re(t,e,n){e=x(e,n);for(var r,i=Vt(t),a=0,o=i.length;a<o;a++)if(e(t[r=i[a]],r,t))return r}function ie(t,e,n){return e in n}var ae=k((function(t,e){var n={},r=e[0];if(null==t)return n;be(r)?(e.length>1&&(r=y(r,e[1])),e=$t(t)):(r=ie,e=lt(e,!1,!1),t=Object(t));for(var i=0,a=e.length;i<a;i++){var o=e[i],u=t[o];r(u,o,t)&&(n[o]=u)}return n})),oe=k((function(t,e){var n,r=e[0];return be(r)?(r=zt(r),e.length>1&&(n=e[1])):(e=C(lt(e,!1,!1),String),r=function(t,n){return!H(e,n)}),ae(t,r,n)})),ue=te($t,!0);function se(t,e){var n=M(t);return e&&ne(n,e),n}function ce(t){return _e(t)?ge(t)?t.slice():ee({},t):t}function le(t,e){return e(t),t}function fe(t,e){var n=Vt(e),r=n.length;if(null==t)return!r;for(var i=Object(t),a=0;a<r;a++){var o=n[a];if(e[o]!==i[o]||!(o in i))return!1}return!0}function de(t,e,n,r){if(t===e)return 0!==t||1/t==1/e;if(null==t||null==e)return!1;if(t!=t)return e!=e;var i=typeof t;return("function"===i||"object"===i||"object"==typeof e)&&function(t,e,n,r){t instanceof g&&(t=t._wrapped);e instanceof g&&(e=e._wrapped);var i=c.call(t);if(i!==c.call(e))return!1;switch(i){case"[object RegExp]":case"[object String]":return""+t==""+e;case"[object Number]":return+t!=+t?+e!=+e:0==+t?1/+t==1/e:+t==+e;case"[object Date]":case"[object Boolean]":return+t==+e;case"[object Symbol]":return o.valueOf.call(t)===o.valueOf.call(e)}var a="[object Array]"===i;if(!a){if("object"!=typeof t||"object"!=typeof e)return!1;var u=t.constructor,s=e.constructor;if(u!==s&&!(be(u)&&u instanceof u&&be(s)&&s instanceof s)&&"constructor"in t&&"constructor"in e)return!1}r=r||[];var l=(n=n||[]).length;for(;l--;)if(n[l]===t)return r[l]===e;if(n.push(t),r.push(e),a){if((l=t.length)!==e.length)return!1;for(;l--;)if(!de(t[l],e[l],n,r))return!1}else{var f,d=Vt(t);if(l=d.length,Vt(e).length!==l)return!1;for(;l--;)if(f=d[l],!S(e,f)||!de(t[f],e[f],n,r))return!1}return n.pop(),r.pop(),!0}(t,e,n,r)}function he(t,e){return de(t,e)}function pe(t){return null==t||(A(t)&&(ge(t)||we(t)||ye(t))?0===t.length:0===Vt(t).length)}function me(t){return!(!t||1!==t.nodeType)}function ve(t){return function(e){return c.call(e)==="[object "+t+"]"}}var ge=f||ve("Array");function _e(t){var e=typeof t;return"function"===e||"object"===e&&!!t}var ye=ve("Arguments"),be=ve("Function"),we=ve("String"),xe=ve("Number"),ke=ve("Date"),Me=ve("RegExp"),Ee=ve("Error"),Se=ve("Symbol"),Te=ve("Map"),je=ve("WeakMap"),Oe=ve("Set"),Ae=ve("WeakSet");!function(){ye(arguments)||(ye=function(t){return S(t,"callee")})}();var Le=r.document&&r.document.childNodes;function Ce(t){return!Se(t)&&m(t)&&!p(parseFloat(t))}function De(t){return xe(t)&&p(t)}function Ne(t){return!0===t||!1===t||"[object Boolean]"===c.call(t)}function Pe(t){return null===t}function Re(t){return void 0===t}function Ye(t,e){if(!ge(e))return S(t,e);for(var n=e.length,r=0;r<n;r++){var i=e[r];if(null==t||!l.call(t,i))return!1;t=t[i]}return!!n}function Ie(t){return t}function ze(t){return function(){return t}}function Fe(){}function He(t){return ge(t)?function(e){return T(e,t)}:E(t)}function Be(t){return null==t?function(){}:function(e){return ge(e)?T(t,e):t[e]}}function qe(t){return t=ne({},t),function(e){return fe(e,t)}}function Ue(t,e,n){var r=Array(Math.max(0,t));e=y(e,n,1);for(var i=0;i<t;i++)r[i]=e(i);return r}function We(t,e){return null==e&&(e=t,t=0),t+Math.floor(Math.random()*(e-t+1))}"object"!=typeof Int8Array&&"function"!=typeof Le&&(be=function(t){return"function"==typeof t||!1});var Ge=Date.now||function(){return(new Date).getTime()},Ve={"&":"&amp;","<":"&lt;",">":"&gt;",'"':"&quot;","'":"&#x27;","`":"&#x60;"},$e=Kt(Ve);function Xe(t){var e=function(e){return t[e]},n="(?:"+Vt(t).join("|")+")",r=RegExp(n),i=RegExp(n,"g");return function(t){return t=null==t?"":""+t,r.test(t)?t.replace(i,e):t}}var Je=Xe(Ve),Ze=Xe($e);function Ke(t,e,n){ge(e)||(e=[e]);var r=e.length;if(!r)return be(n)?n.call(t):n;for(var i=0;i<r;i++){var a=null==t?void 0:t[e[i]];void 0===a&&(a=n,i=r),t=be(a)?a.call(t):a}return t}var Qe=0;function tn(t){var e=++Qe+"";return t?t+e:e}var en=g.templateSettings={evaluate:/<%([\s\S]+?)%>/g,interpolate:/<%=([\s\S]+?)%>/g,escape:/<%-([\s\S]+?)%>/g},nn=/(.)^/,rn={"'":"'","\\":"\\","\r":"r","\n":"n","\u2028":"u2028","\u2029":"u2029"},an=/\\|'|\r|\n|\u2028|\u2029/g,on=function(t){return"\\"+rn[t]};function un(t,e,n){!e&&n&&(e=n),e=ue({},e,g.templateSettings);var r,i=RegExp([(e.escape||nn).source,(e.interpolate||nn).source,(e.evaluate||nn).source].join("|")+"|$","g"),a=0,o="__p+='";t.replace(i,(function(e,n,r,i,u){return o+=t.slice(a,u).replace(an,on),a=u+e.length,n?o+="'+\n((__t=("+n+"))==null?'':_.escape(__t))+\n'":r?o+="'+\n((__t=("+r+"))==null?'':__t)+\n'":i&&(o+="';\n"+i+"\n__p+='"),e})),o+="';\n",e.variable||(o="with(obj||{}){\n"+o+"}\n"),o="var __t,__p='',__j=Array.prototype.join,print=function(){__p+=__j.call(arguments,'');};\n"+o+"return __p;\n";try{r=new Function(e.variable||"obj","_",o)}catch(t){throw t.source=o,t}var u=function(t){return r.call(this,t,g)},s=e.variable||"obj";return u.source="function("+s+"){\n"+o+"}",u}function sn(t){var e=g(t);return e._chain=!0,e}function cn(t,e){return t._chain?g(e).chain():e}function ln(t){return L(Qt(t),(function(e){var n=g[e]=t[e];g.prototype[e]=function(){var t=[this._wrapped];return u.apply(t,arguments),cn(this,n.apply(g,t))}})),g}L(["pop","push","reverse","shift","sort","splice","unshift"],(function(t){var e=i[t];g.prototype[t]=function(){var n=this._wrapped;return e.apply(n,arguments),"shift"!==t&&"splice"!==t||0!==n.length||delete n[0],cn(this,n)}})),L(["concat","join","slice"],(function(t){var e=i[t];g.prototype[t]=function(){return cn(this,e.apply(this._wrapped,arguments))}})),g.prototype.value=function(){return this._wrapped},g.prototype.valueOf=g.prototype.toJSON=g.prototype.value,g.prototype.toString=function(){return String(this._wrapped)}}.call(this,n(6))},function(t,e,n){"use strict";function r(t,e,n,r,i){var a=t*t,o=a*t;return((1-3*t+3*a-o)*e+(4-6*a+3*o)*n+(1+3*t+3*a-3*o)*r+o*i)/6}n.d(e,"a",(function(){return r})),e.b=function(t){var e=t.length-1;return function(n){var i=n<=0?n=0:n>=1?(n=1,e-1):Math.floor(n*e),a=t[i],o=t[i+1],u=i>0?t[i-1]:2*a-o,s=i<e-1?t[i+2]:2*o-a;return r((n-i/e)*e,u,a,o,s)}}},function(t,e,n){"use strict";var r=n(62);e.a=function(t){var e=t+="",n=e.indexOf(":");return n>=0&&"xmlns"!==(e=t.slice(0,n))&&(t=t.slice(n+1)),r.a.hasOwnProperty(e)?{space:r.a[e],local:t}:t}},function(t,e,n){"use strict";function r(){}e.a=function(t){return null==t?r:function(){return this.querySelector(t)}}},function(t,e,n){"use strict";e.a=function(t){return t.ownerDocument&&t.ownerDocument.defaultView||t.document&&t||t.defaultView}},function(t,e,n){"use strict";var r=n(14),i=n(78),a=n(165),o=n(164),u=n(25),s=n(166),c=n(160),l=n(107),f=n(63);e.a=function(t,e){var n,d=typeof e;return null==e||"boolean"===d?Object(l.a)(e):("number"===d?u.a:"string"===d?(n=Object(r.e)(e))?(e=n,i.a):c.a:e instanceof r.e?i.a:e instanceof Date?o.a:Object(f.b)(e)?f.a:Array.isArray(e)?a.b:"function"!=typeof e.valueOf&&"function"!=typeof e.toString||isNaN(e)?s.a:u.a)(t,e)}},function(t,e,n){"use strict";n.d(e,"b",(function(){return a}));var r=n(106),i=n(33);function a(t,e){var n=t.document.documentElement,a=Object(r.a)(t).on("dragstart.drag",null);e&&(a.on("click.drag",i.a,!0),setTimeout((function(){a.on("click.drag",null)}),0)),"onselectstart"in n?a.on("selectstart.drag",null):(n.style.MozUserSelect=n.__noselect,delete n.__noselect)}e.a=function(t){var e=t.document.documentElement,n=Object(r.a)(t).on("dragstart.drag",i.a,!0);"onselectstart"in e?n.on("selectstart.drag",i.a,!0):(e.__noselect=e.style.MozUserSelect,e.style.MozUserSelect="none")}},function(t,e,n){"use strict";var r={},i={};function a(t){return new Function("d","return {"+t.map((function(t,e){return JSON.stringify(t)+": d["+e+'] || ""'})).join(",")+"}")}function o(t){var e=Object.create(null),n=[];return t.forEach((function(t){for(var r in t)r in e||n.push(e[r]=r)})),n}function u(t,e){var n=t+"",r=n.length;return r<e?new Array(e-r+1).join(0)+n:n}function s(t){var e,n=t.getUTCHours(),r=t.getUTCMinutes(),i=t.getUTCSeconds(),a=t.getUTCMilliseconds();return isNaN(t)?"Invalid Date":((e=t.getUTCFullYear())<0?"-"+u(-e,6):e>9999?"+"+u(e,6):u(e,4))+"-"+u(t.getUTCMonth()+1,2)+"-"+u(t.getUTCDate(),2)+(a?"T"+u(n,2)+":"+u(r,2)+":"+u(i,2)+"."+u(a,3)+"Z":i?"T"+u(n,2)+":"+u(r,2)+":"+u(i,2)+"Z":r||n?"T"+u(n,2)+":"+u(r,2)+"Z":"")}e.a=function(t){var e=new RegExp('["'+t+"\n\r]"),n=t.charCodeAt(0);function u(t,e){var a,o=[],u=t.length,s=0,c=0,l=u<=0,f=!1;function d(){if(l)return i;if(f)return f=!1,r;var e,a,o=s;if(34===t.charCodeAt(o)){for(;s++<u&&34!==t.charCodeAt(s)||34===t.charCodeAt(++s););return(e=s)>=u?l=!0:10===(a=t.charCodeAt(s++))?f=!0:13===a&&(f=!0,10===t.charCodeAt(s)&&++s),t.slice(o+1,e-1).replace(/""/g,'"')}for(;s<u;){if(10===(a=t.charCodeAt(e=s++)))f=!0;else if(13===a)f=!0,10===t.charCodeAt(s)&&++s;else if(a!==n)continue;return t.slice(o,e)}return l=!0,t.slice(o,u)}for(10===t.charCodeAt(u-1)&&--u,13===t.charCodeAt(u-1)&&--u;(a=d())!==i;){for(var h=[];a!==r&&a!==i;)h.push(a),a=d();e&&null==(h=e(h,c++))||o.push(h)}return o}function c(e,n){return e.map((function(e){return n.map((function(t){return f(e[t])})).join(t)}))}function l(e){return e.map(f).join(t)}function f(t){return null==t?"":t instanceof Date?s(t):e.test(t+="")?'"'+t.replace(/"/g,'""')+'"':t}return{parse:function(t,e){var n,r,i=u(t,(function(t,i){if(n)return n(t,i-1);r=t,n=e?function(t,e){var n=a(t);return function(r,i){return e(n(r),i,t)}}(t,e):a(t)}));return i.columns=r||[],i},parseRows:u,format:function(e,n){return null==n&&(n=o(e)),[n.map(f).join(t)].concat(c(e,n)).join("\n")},formatBody:function(t,e){return null==e&&(e=o(t)),c(t,e).join("\n")},formatRows:function(t){return t.map(l).join("\n")},formatRow:l,formatValue:f}}},function(t,e,n){"use strict";n.d(e,"b",(function(){return i})),n.d(e,"a",(function(){return a}));var r=/^(?:(.)?([<>=^]))?([+\-( ])?([$#])?(0)?(\d+)?(,)?(\.\d+)?(~)?([a-z%])?$/i;function i(t){if(!(e=r.exec(t)))throw new Error("invalid format: "+t);var e;return new a({fill:e[1],align:e[2],sign:e[3],symbol:e[4],zero:e[5],width:e[6],comma:e[7],precision:e[8]&&e[8].slice(1),trim:e[9],type:e[10]})}function a(t){this.fill=void 0===t.fill?" ":t.fill+"",this.align=void 0===t.align?">":t.align+"",this.sign=void 0===t.sign?"-":t.sign+"",this.symbol=void 0===t.symbol?"":t.symbol+"",this.zero=!!t.zero,this.width=void 0===t.width?void 0:+t.width,this.comma=!!t.comma,this.precision=void 0===t.precision?void 0:+t.precision,this.trim=!!t.trim,this.type=void 0===t.type?"":t.type+""}i.prototype=a.prototype,a.prototype.toString=function(){return this.fill+this.align+this.sign+this.symbol+(this.zero?"0":"")+(void 0===this.width?"":Math.max(1,0|this.width))+(this.comma?",":"")+(void 0===this.precision?"":"."+Math.max(0,0|this.precision))+(this.trim?"~":"")+this.type}},function(t,e,n){"use strict";var r=n(228),i=n(5),a=n(61),o=n(50);e.a=function(){var t=o.a,e=o.b,n=Object(i.a)(!0),u=null,s=a.a,c=null;function l(i){var a,o,l,f=i.length,d=!1;for(null==u&&(c=s(l=Object(r.a)())),a=0;a<=f;++a)!(a<f&&n(o=i[a],a,i))===d&&((d=!d)?c.lineStart():c.lineEnd()),d&&c.point(+t(o,a,i),+e(o,a,i));if(l)return c=null,l+""||null}return l.x=function(e){return arguments.length?(t="function"==typeof e?e:Object(i.a)(+e),l):t},l.y=function(t){return arguments.length?(e="function"==typeof t?t:Object(i.a)(+t),l):e},l.defined=function(t){return arguments.length?(n="function"==typeof t?t:Object(i.a)(!!t),l):n},l.curve=function(t){return arguments.length?(s=t,null!=u&&(c=s(u)),l):s},l.context=function(t){return arguments.length?(null==t?u=c=null:c=s(u=t),l):u},l}},function(t,e,n){"use strict";n.r(e),function(t){n.d(e,"default",(function(){return g})),n.d(e,"VERSION",(function(){return _})),n.d(e,"iteratee",(function(){return w})),n.d(e,"restArguments",(function(){return k})),n.d(e,"each",(function(){return L})),n.d(e,"forEach",(function(){return L})),n.d(e,"map",(function(){return C})),n.d(e,"collect",(function(){return C})),n.d(e,"reduce",(function(){return N})),n.d(e,"foldl",(function(){return N})),n.d(e,"inject",(function(){return N})),n.d(e,"reduceRight",(function(){return P})),n.d(e,"foldr",(function(){return P})),n.d(e,"find",(function(){return R})),n.d(e,"detect",(function(){return R})),n.d(e,"filter",(function(){return Y})),n.d(e,"select",(function(){return Y})),n.d(e,"reject",(function(){return I})),n.d(e,"every",(function(){return z})),n.d(e,"all",(function(){return z})),n.d(e,"some",(function(){return F})),n.d(e,"any",(function(){return F})),n.d(e,"contains",(function(){return H})),n.d(e,"includes",(function(){return H})),n.d(e,"include",(function(){return H})),n.d(e,"invoke",(function(){return B})),n.d(e,"pluck",(function(){return q})),n.d(e,"where",(function(){return U})),n.d(e,"findWhere",(function(){return W})),n.d(e,"max",(function(){return G})),n.d(e,"min",(function(){return V})),n.d(e,"shuffle",(function(){return $})),n.d(e,"sample",(function(){return X})),n.d(e,"sortBy",(function(){return J})),n.d(e,"groupBy",(function(){return K})),n.d(e,"indexBy",(function(){return Q})),n.d(e,"countBy",(function(){return tt})),n.d(e,"toArray",(function(){return nt})),n.d(e,"size",(function(){return rt})),n.d(e,"partition",(function(){return it})),n.d(e,"first",(function(){return at})),n.d(e,"head",(function(){return at})),n.d(e,"take",(function(){return at})),n.d(e,"initial",(function(){return ot})),n.d(e,"last",(function(){return ut})),n.d(e,"rest",(function(){return st})),n.d(e,"tail",(function(){return st})),n.d(e,"drop",(function(){return st})),n.d(e,"compact",(function(){return ct})),n.d(e,"flatten",(function(){return ft})),n.d(e,"without",(function(){return dt})),n.d(e,"uniq",(function(){return ht})),n.d(e,"unique",(function(){return ht})),n.d(e,"union",(function(){return pt})),n.d(e,"intersection",(function(){return mt})),n.d(e,"difference",(function(){return vt})),n.d(e,"unzip",(function(){return gt})),n.d(e,"zip",(function(){return _t})),n.d(e,"object",(function(){return yt})),n.d(e,"findIndex",(function(){return wt})),n.d(e,"findLastIndex",(function(){return xt})),n.d(e,"sortedIndex",(function(){return kt})),n.d(e,"indexOf",(function(){return Et})),n.d(e,"lastIndexOf",(function(){return St})),n.d(e,"range",(function(){return Tt})),n.d(e,"chunk",(function(){return jt})),n.d(e,"bind",(function(){return At})),n.d(e,"partial",(function(){return Lt})),n.d(e,"bindAll",(function(){return Ct})),n.d(e,"memoize",(function(){return Dt})),n.d(e,"delay",(function(){return Nt})),n.d(e,"defer",(function(){return Pt})),n.d(e,"throttle",(function(){return Rt})),n.d(e,"debounce",(function(){return Yt})),n.d(e,"wrap",(function(){return It})),n.d(e,"negate",(function(){return zt})),n.d(e,"compose",(function(){return Ft})),n.d(e,"after",(function(){return Ht})),n.d(e,"before",(function(){return Bt})),n.d(e,"once",(function(){return qt})),n.d(e,"keys",(function(){return Vt})),n.d(e,"allKeys",(function(){return $t})),n.d(e,"values",(function(){return Xt})),n.d(e,"mapObject",(function(){return Jt})),n.d(e,"pairs",(function(){return Zt})),n.d(e,"invert",(function(){return Kt})),n.d(e,"functions",(function(){return Qt})),n.d(e,"methods",(function(){return Qt})),n.d(e,"extend",(function(){return ee})),n.d(e,"extendOwn",(function(){return ne})),n.d(e,"assign",(function(){return ne})),n.d(e,"findKey",(function(){return re})),n.d(e,"pick",(function(){return ae})),n.d(e,"omit",(function(){return oe})),n.d(e,"defaults",(function(){return ue})),n.d(e,"create",(function(){return se})),n.d(e,"clone",(function(){return ce})),n.d(e,"tap",(function(){return le})),n.d(e,"isMatch",(function(){return fe})),n.d(e,"isEqual",(function(){return he})),n.d(e,"isEmpty",(function(){return pe})),n.d(e,"isElement",(function(){return me})),n.d(e,"isArray",(function(){return ge})),n.d(e,"isObject",(function(){return _e})),n.d(e,"isArguments",(function(){return ye})),n.d(e,"isFunction",(function(){return be})),n.d(e,"isString",(function(){return we})),n.d(e,"isNumber",(function(){return xe})),n.d(e,"isDate",(function(){return ke})),n.d(e,"isRegExp",(function(){return Me})),n.d(e,"isError",(function(){return Ee})),n.d(e,"isSymbol",(function(){return Se})),n.d(e,"isMap",(function(){return Te})),n.d(e,"isWeakMap",(function(){return je})),n.d(e,"isSet",(function(){return Oe})),n.d(e,"isWeakSet",(function(){return Ae})),n.d(e,"isFinite",(function(){return Ce})),n.d(e,"isNaN",(function(){return De})),n.d(e,"isBoolean",(function(){return Ne})),n.d(e,"isNull",(function(){return Pe})),n.d(e,"isUndefined",(function(){return Re})),n.d(e,"has",(function(){return Ye})),n.d(e,"identity",(function(){return Ie})),n.d(e,"constant",(function(){return ze})),n.d(e,"noop",(function(){return Fe})),n.d(e,"property",(function(){return He})),n.d(e,"propertyOf",(function(){return Be})),n.d(e,"matcher",(function(){return qe})),n.d(e,"matches",(function(){return qe})),n.d(e,"times",(function(){return Ue})),n.d(e,"random",(function(){return We})),n.d(e,"now",(function(){return Ge})),n.d(e,"escape",(function(){return Je})),n.d(e,"unescape",(function(){return Ze})),n.d(e,"result",(function(){return Ke})),n.d(e,"uniqueId",(function(){return tn})),n.d(e,"templateSettings",(function(){return en})),n.d(e,"template",(function(){return un})),n.d(e,"chain",(function(){return sn})),n.d(e,"mixin",(function(){return ln}));var r="object"==typeof self&&self.self===self&&self||"object"==typeof t&&t.global===t&&t||Function("return this")()||{},i=Array.prototype,a=Object.prototype,o="undefined"!=typeof Symbol?Symbol.prototype:null,u=i.push,s=i.slice,c=a.toString,l=a.hasOwnProperty,f=Array.isArray,d=Object.keys,h=Object.create,p=r.isNaN,m=r.isFinite,v=function(){};function g(t){return t instanceof g?t:this instanceof g?void(this._wrapped=t):new g(t)}var _=g.VERSION="1.10.2";function y(t,e,n){if(void 0===e)return t;switch(null==n?3:n){case 1:return function(n){return t.call(e,n)};case 3:return function(n,r,i){return t.call(e,n,r,i)};case 4:return function(n,r,i,a){return t.call(e,n,r,i,a)}}return function(){return t.apply(e,arguments)}}function b(t,e,n){return null==t?Ie:be(t)?y(t,e,n):_e(t)&&!ge(t)?qe(t):He(t)}function w(t,e){return b(t,e,1/0)}function x(t,e,n){return g.iteratee!==w?g.iteratee(t,e):b(t,e,n)}function k(t,e){return e=null==e?t.length-1:+e,function(){for(var n=Math.max(arguments.length-e,0),r=Array(n),i=0;i<n;i++)r[i]=arguments[i+e];switch(e){case 0:return t.call(this,r);case 1:return t.call(this,arguments[0],r);case 2:return t.call(this,arguments[0],arguments[1],r)}var a=Array(e+1);for(i=0;i<e;i++)a[i]=arguments[i];return a[e]=r,t.apply(this,a)}}function M(t){if(!_e(t))return{};if(h)return h(t);v.prototype=t;var e=new v;return v.prototype=null,e}function E(t){return function(e){return null==e?void 0:e[t]}}function S(t,e){return null!=t&&l.call(t,e)}function T(t,e){for(var n=e.length,r=0;r<n;r++){if(null==t)return;t=t[e[r]]}return n?t:void 0}g.iteratee=w;var j=Math.pow(2,53)-1,O=E("length");function A(t){var e=O(t);return"number"==typeof e&&e>=0&&e<=j}function L(t,e,n){var r,i;if(e=y(e,n),A(t))for(r=0,i=t.length;r<i;r++)e(t[r],r,t);else{var a=Vt(t);for(r=0,i=a.length;r<i;r++)e(t[a[r]],a[r],t)}return t}function C(t,e,n){e=x(e,n);for(var r=!A(t)&&Vt(t),i=(r||t).length,a=Array(i),o=0;o<i;o++){var u=r?r[o]:o;a[o]=e(t[u],u,t)}return a}function D(t){var e=function(e,n,r,i){var a=!A(e)&&Vt(e),o=(a||e).length,u=t>0?0:o-1;for(i||(r=e[a?a[u]:u],u+=t);u>=0&&u<o;u+=t){var s=a?a[u]:u;r=n(r,e[s],s,e)}return r};return function(t,n,r,i){var a=arguments.length>=3;return e(t,y(n,i,4),r,a)}}var N=D(1),P=D(-1);function R(t,e,n){var r=(A(t)?wt:re)(t,e,n);if(void 0!==r&&-1!==r)return t[r]}function Y(t,e,n){var r=[];return e=x(e,n),L(t,(function(t,n,i){e(t,n,i)&&r.push(t)})),r}function I(t,e,n){return Y(t,zt(x(e)),n)}function z(t,e,n){e=x(e,n);for(var r=!A(t)&&Vt(t),i=(r||t).length,a=0;a<i;a++){var o=r?r[a]:a;if(!e(t[o],o,t))return!1}return!0}function F(t,e,n){e=x(e,n);for(var r=!A(t)&&Vt(t),i=(r||t).length,a=0;a<i;a++){var o=r?r[a]:a;if(e(t[o],o,t))return!0}return!1}function H(t,e,n,r){return A(t)||(t=Xt(t)),("number"!=typeof n||r)&&(n=0),Et(t,e,n)>=0}var B=k((function(t,e,n){var r,i;return be(e)?i=e:ge(e)&&(r=e.slice(0,-1),e=e[e.length-1]),C(t,(function(t){var a=i;if(!a){if(r&&r.length&&(t=T(t,r)),null==t)return;a=t[e]}return null==a?a:a.apply(t,n)}))}));function q(t,e){return C(t,He(e))}function U(t,e){return Y(t,qe(e))}function W(t,e){return R(t,qe(e))}function G(t,e,n){var r,i,a=-1/0,o=-1/0;if(null==e||"number"==typeof e&&"object"!=typeof t[0]&&null!=t)for(var u=0,s=(t=A(t)?t:Xt(t)).length;u<s;u++)null!=(r=t[u])&&r>a&&(a=r);else e=x(e,n),L(t,(function(t,n,r){((i=e(t,n,r))>o||i===-1/0&&a===-1/0)&&(a=t,o=i)}));return a}function V(t,e,n){var r,i,a=1/0,o=1/0;if(null==e||"number"==typeof e&&"object"!=typeof t[0]&&null!=t)for(var u=0,s=(t=A(t)?t:Xt(t)).length;u<s;u++)null!=(r=t[u])&&r<a&&(a=r);else e=x(e,n),L(t,(function(t,n,r){((i=e(t,n,r))<o||i===1/0&&a===1/0)&&(a=t,o=i)}));return a}function $(t){return X(t,1/0)}function X(t,e,n){if(null==e||n)return A(t)||(t=Xt(t)),t[We(t.length-1)];var r=A(t)?ce(t):Xt(t),i=O(r);e=Math.max(Math.min(e,i),0);for(var a=i-1,o=0;o<e;o++){var u=We(o,a),s=r[o];r[o]=r[u],r[u]=s}return r.slice(0,e)}function J(t,e,n){var r=0;return e=x(e,n),q(C(t,(function(t,n,i){return{value:t,index:r++,criteria:e(t,n,i)}})).sort((function(t,e){var n=t.criteria,r=e.criteria;if(n!==r){if(n>r||void 0===n)return 1;if(n<r||void 0===r)return-1}return t.index-e.index})),"value")}function Z(t,e){return function(n,r,i){var a=e?[[],[]]:{};return r=x(r,i),L(n,(function(e,i){var o=r(e,i,n);t(a,e,o)})),a}}var K=Z((function(t,e,n){S(t,n)?t[n].push(e):t[n]=[e]})),Q=Z((function(t,e,n){t[n]=e})),tt=Z((function(t,e,n){S(t,n)?t[n]++:t[n]=1})),et=/[^\ud800-\udfff]|[\ud800-\udbff][\udc00-\udfff]|[\ud800-\udfff]/g;function nt(t){return t?ge(t)?s.call(t):we(t)?t.match(et):A(t)?C(t,Ie):Xt(t):[]}function rt(t){return null==t?0:A(t)?t.length:Vt(t).length}var it=Z((function(t,e,n){t[n?0:1].push(e)}),!0);function at(t,e,n){return null==t||t.length<1?null==e?void 0:[]:null==e||n?t[0]:ot(t,t.length-e)}function ot(t,e,n){return s.call(t,0,Math.max(0,t.length-(null==e||n?1:e)))}function ut(t,e,n){return null==t||t.length<1?null==e?void 0:[]:null==e||n?t[t.length-1]:st(t,Math.max(0,t.length-e))}function st(t,e,n){return s.call(t,null==e||n?1:e)}function ct(t){return Y(t,Boolean)}function lt(t,e,n,r){for(var i=(r=r||[]).length,a=0,o=O(t);a<o;a++){var u=t[a];if(A(u)&&(ge(u)||ye(u)))if(e)for(var s=0,c=u.length;s<c;)r[i++]=u[s++];else lt(u,e,n,r),i=r.length;else n||(r[i++]=u)}return r}function ft(t,e){return lt(t,e,!1)}var dt=k((function(t,e){return vt(t,e)}));function ht(t,e,n,r){Ne(e)||(r=n,n=e,e=!1),null!=n&&(n=x(n,r));for(var i=[],a=[],o=0,u=O(t);o<u;o++){var s=t[o],c=n?n(s,o,t):s;e&&!n?(o&&a===c||i.push(s),a=c):n?H(a,c)||(a.push(c),i.push(s)):H(i,s)||i.push(s)}return i}var pt=k((function(t){return ht(lt(t,!0,!0))}));function mt(t){for(var e=[],n=arguments.length,r=0,i=O(t);r<i;r++){var a=t[r];if(!H(e,a)){var o;for(o=1;o<n&&H(arguments[o],a);o++);o===n&&e.push(a)}}return e}var vt=k((function(t,e){return e=lt(e,!0,!0),Y(t,(function(t){return!H(e,t)}))}));function gt(t){for(var e=t&&G(t,O).length||0,n=Array(e),r=0;r<e;r++)n[r]=q(t,r);return n}var _t=k(gt);function yt(t,e){for(var n={},r=0,i=O(t);r<i;r++)e?n[t[r]]=e[r]:n[t[r][0]]=t[r][1];return n}function bt(t){return function(e,n,r){n=x(n,r);for(var i=O(e),a=t>0?0:i-1;a>=0&&a<i;a+=t)if(n(e[a],a,e))return a;return-1}}var wt=bt(1),xt=bt(-1);function kt(t,e,n,r){for(var i=(n=x(n,r,1))(e),a=0,o=O(t);a<o;){var u=Math.floor((a+o)/2);n(t[u])<i?a=u+1:o=u}return a}function Mt(t,e,n){return function(r,i,a){var o=0,u=O(r);if("number"==typeof a)t>0?o=a>=0?a:Math.max(a+u,o):u=a>=0?Math.min(a+1,u):a+u+1;else if(n&&a&&u)return r[a=n(r,i)]===i?a:-1;if(i!=i)return(a=e(s.call(r,o,u),De))>=0?a+o:-1;for(a=t>0?o:u-1;a>=0&&a<u;a+=t)if(r[a]===i)return a;return-1}}var Et=Mt(1,wt,kt),St=Mt(-1,xt);function Tt(t,e,n){null==e&&(e=t||0,t=0),n||(n=e<t?-1:1);for(var r=Math.max(Math.ceil((e-t)/n),0),i=Array(r),a=0;a<r;a++,t+=n)i[a]=t;return i}function jt(t,e){if(null==e||e<1)return[];for(var n=[],r=0,i=t.length;r<i;)n.push(s.call(t,r,r+=e));return n}function Ot(t,e,n,r,i){if(!(r instanceof e))return t.apply(n,i);var a=M(t.prototype),o=t.apply(a,i);return _e(o)?o:a}var At=k((function(t,e,n){if(!be(t))throw new TypeError("Bind must be called on a function");var r=k((function(i){return Ot(t,r,e,this,n.concat(i))}));return r})),Lt=k((function(t,e){var n=Lt.placeholder,r=function(){for(var i=0,a=e.length,o=Array(a),u=0;u<a;u++)o[u]=e[u]===n?arguments[i++]:e[u];for(;i<arguments.length;)o.push(arguments[i++]);return Ot(t,r,this,this,o)};return r}));Lt.placeholder=g;var Ct=k((function(t,e){var n=(e=lt(e,!1,!1)).length;if(n<1)throw new Error("bindAll must be passed function names");for(;n--;){var r=e[n];t[r]=At(t[r],t)}}));function Dt(t,e){var n=function(r){var i=n.cache,a=""+(e?e.apply(this,arguments):r);return S(i,a)||(i[a]=t.apply(this,arguments)),i[a]};return n.cache={},n}var Nt=k((function(t,e,n){return setTimeout((function(){return t.apply(null,n)}),e)})),Pt=Lt(Nt,g,1);function Rt(t,e,n){var r,i,a,o,u=0;n||(n={});var s=function(){u=!1===n.leading?0:Ge(),r=null,o=t.apply(i,a),r||(i=a=null)},c=function(){var c=Ge();u||!1!==n.leading||(u=c);var l=e-(c-u);return i=this,a=arguments,l<=0||l>e?(r&&(clearTimeout(r),r=null),u=c,o=t.apply(i,a),r||(i=a=null)):r||!1===n.trailing||(r=setTimeout(s,l)),o};return c.cancel=function(){clearTimeout(r),u=0,r=i=a=null},c}function Yt(t,e,n){var r,i,a=function(e,n){r=null,n&&(i=t.apply(e,n))},o=k((function(o){if(r&&clearTimeout(r),n){var u=!r;r=setTimeout(a,e),u&&(i=t.apply(this,o))}else r=Nt(a,e,this,o);return i}));return o.cancel=function(){clearTimeout(r),r=null},o}function It(t,e){return Lt(e,t)}function zt(t){return function(){return!t.apply(this,arguments)}}function Ft(){var t=arguments,e=t.length-1;return function(){for(var n=e,r=t[e].apply(this,arguments);n--;)r=t[n].call(this,r);return r}}function Ht(t,e){return function(){if(--t<1)return e.apply(this,arguments)}}function Bt(t,e){var n;return function(){return--t>0&&(n=e.apply(this,arguments)),t<=1&&(e=null),n}}var qt=Lt(Bt,2),Ut=!{toString:null}.propertyIsEnumerable("toString"),Wt=["valueOf","isPrototypeOf","toString","propertyIsEnumerable","hasOwnProperty","toLocaleString"];function Gt(t,e){var n=Wt.length,r=t.constructor,i=be(r)&&r.prototype||a,o="constructor";for(S(t,o)&&!H(e,o)&&e.push(o);n--;)(o=Wt[n])in t&&t[o]!==i[o]&&!H(e,o)&&e.push(o)}function Vt(t){if(!_e(t))return[];if(d)return d(t);var e=[];for(var n in t)S(t,n)&&e.push(n);return Ut&&Gt(t,e),e}function $t(t){if(!_e(t))return[];var e=[];for(var n in t)e.push(n);return Ut&&Gt(t,e),e}function Xt(t){for(var e=Vt(t),n=e.length,r=Array(n),i=0;i<n;i++)r[i]=t[e[i]];return r}function Jt(t,e,n){e=x(e,n);for(var r=Vt(t),i=r.length,a={},o=0;o<i;o++){var u=r[o];a[u]=e(t[u],u,t)}return a}function Zt(t){for(var e=Vt(t),n=e.length,r=Array(n),i=0;i<n;i++)r[i]=[e[i],t[e[i]]];return r}function Kt(t){for(var e={},n=Vt(t),r=0,i=n.length;r<i;r++)e[t[n[r]]]=n[r];return e}function Qt(t){var e=[];for(var n in t)be(t[n])&&e.push(n);return e.sort()}function te(t,e){return function(n){var r=arguments.length;if(e&&(n=Object(n)),r<2||null==n)return n;for(var i=1;i<r;i++)for(var a=arguments[i],o=t(a),u=o.length,s=0;s<u;s++){var c=o[s];e&&void 0!==n[c]||(n[c]=a[c])}return n}}var ee=te($t),ne=te(Vt);function re(t,e,n){e=x(e,n);for(var r,i=Vt(t),a=0,o=i.length;a<o;a++)if(e(t[r=i[a]],r,t))return r}function ie(t,e,n){return e in n}var ae=k((function(t,e){var n={},r=e[0];if(null==t)return n;be(r)?(e.length>1&&(r=y(r,e[1])),e=$t(t)):(r=ie,e=lt(e,!1,!1),t=Object(t));for(var i=0,a=e.length;i<a;i++){var o=e[i],u=t[o];r(u,o,t)&&(n[o]=u)}return n})),oe=k((function(t,e){var n,r=e[0];return be(r)?(r=zt(r),e.length>1&&(n=e[1])):(e=C(lt(e,!1,!1),String),r=function(t,n){return!H(e,n)}),ae(t,r,n)})),ue=te($t,!0);function se(t,e){var n=M(t);return e&&ne(n,e),n}function ce(t){return _e(t)?ge(t)?t.slice():ee({},t):t}function le(t,e){return e(t),t}function fe(t,e){var n=Vt(e),r=n.length;if(null==t)return!r;for(var i=Object(t),a=0;a<r;a++){var o=n[a];if(e[o]!==i[o]||!(o in i))return!1}return!0}function de(t,e,n,r){if(t===e)return 0!==t||1/t==1/e;if(null==t||null==e)return!1;if(t!=t)return e!=e;var i=typeof t;return("function"===i||"object"===i||"object"==typeof e)&&function(t,e,n,r){t instanceof g&&(t=t._wrapped);e instanceof g&&(e=e._wrapped);var i=c.call(t);if(i!==c.call(e))return!1;switch(i){case"[object RegExp]":case"[object String]":return""+t==""+e;case"[object Number]":return+t!=+t?+e!=+e:0==+t?1/+t==1/e:+t==+e;case"[object Date]":case"[object Boolean]":return+t==+e;case"[object Symbol]":return o.valueOf.call(t)===o.valueOf.call(e)}var a="[object Array]"===i;if(!a){if("object"!=typeof t||"object"!=typeof e)return!1;var u=t.constructor,s=e.constructor;if(u!==s&&!(be(u)&&u instanceof u&&be(s)&&s instanceof s)&&"constructor"in t&&"constructor"in e)return!1}r=r||[];var l=(n=n||[]).length;for(;l--;)if(n[l]===t)return r[l]===e;if(n.push(t),r.push(e),a){if((l=t.length)!==e.length)return!1;for(;l--;)if(!de(t[l],e[l],n,r))return!1}else{var f,d=Vt(t);if(l=d.length,Vt(e).length!==l)return!1;for(;l--;)if(f=d[l],!S(e,f)||!de(t[f],e[f],n,r))return!1}return n.pop(),r.pop(),!0}(t,e,n,r)}function he(t,e){return de(t,e)}function pe(t){return null==t||(A(t)&&(ge(t)||we(t)||ye(t))?0===t.length:0===Vt(t).length)}function me(t){return!(!t||1!==t.nodeType)}function ve(t){return function(e){return c.call(e)==="[object "+t+"]"}}var ge=f||ve("Array");function _e(t){var e=typeof t;return"function"===e||"object"===e&&!!t}var ye=ve("Arguments"),be=ve("Function"),we=ve("String"),xe=ve("Number"),ke=ve("Date"),Me=ve("RegExp"),Ee=ve("Error"),Se=ve("Symbol"),Te=ve("Map"),je=ve("WeakMap"),Oe=ve("Set"),Ae=ve("WeakSet");!function(){ye(arguments)||(ye=function(t){return S(t,"callee")})}();var Le=r.document&&r.document.childNodes;function Ce(t){return!Se(t)&&m(t)&&!p(parseFloat(t))}function De(t){return xe(t)&&p(t)}function Ne(t){return!0===t||!1===t||"[object Boolean]"===c.call(t)}function Pe(t){return null===t}function Re(t){return void 0===t}function Ye(t,e){if(!ge(e))return S(t,e);for(var n=e.length,r=0;r<n;r++){var i=e[r];if(null==t||!l.call(t,i))return!1;t=t[i]}return!!n}function Ie(t){return t}function ze(t){return function(){return t}}function Fe(){}function He(t){return ge(t)?function(e){return T(e,t)}:E(t)}function Be(t){return null==t?function(){}:function(e){return ge(e)?T(t,e):t[e]}}function qe(t){return t=ne({},t),function(e){return fe(e,t)}}function Ue(t,e,n){var r=Array(Math.max(0,t));e=y(e,n,1);for(var i=0;i<t;i++)r[i]=e(i);return r}function We(t,e){return null==e&&(e=t,t=0),t+Math.floor(Math.random()*(e-t+1))}"object"!=typeof Int8Array&&"function"!=typeof Le&&(be=function(t){return"function"==typeof t||!1});var Ge=Date.now||function(){return(new Date).getTime()},Ve={"&":"&amp;","<":"&lt;",">":"&gt;",'"':"&quot;","'":"&#x27;","`":"&#x60;"},$e=Kt(Ve);function Xe(t){var e=function(e){return t[e]},n="(?:"+Vt(t).join("|")+")",r=RegExp(n),i=RegExp(n,"g");return function(t){return t=null==t?"":""+t,r.test(t)?t.replace(i,e):t}}var Je=Xe(Ve),Ze=Xe($e);function Ke(t,e,n){ge(e)||(e=[e]);var r=e.length;if(!r)return be(n)?n.call(t):n;for(var i=0;i<r;i++){var a=null==t?void 0:t[e[i]];void 0===a&&(a=n,i=r),t=be(a)?a.call(t):a}return t}var Qe=0;function tn(t){var e=++Qe+"";return t?t+e:e}var en=g.templateSettings={evaluate:/<%([\s\S]+?)%>/g,interpolate:/<%=([\s\S]+?)%>/g,escape:/<%-([\s\S]+?)%>/g},nn=/(.)^/,rn={"'":"'","\\":"\\","\r":"r","\n":"n","\u2028":"u2028","\u2029":"u2029"},an=/\\|'|\r|\n|\u2028|\u2029/g,on=function(t){return"\\"+rn[t]};function un(t,e,n){!e&&n&&(e=n),e=ue({},e,g.templateSettings);var r,i=RegExp([(e.escape||nn).source,(e.interpolate||nn).source,(e.evaluate||nn).source].join("|")+"|$","g"),a=0,o="__p+='";t.replace(i,(function(e,n,r,i,u){return o+=t.slice(a,u).replace(an,on),a=u+e.length,n?o+="'+\n((__t=("+n+"))==null?'':_.escape(__t))+\n'":r?o+="'+\n((__t=("+r+"))==null?'':__t)+\n'":i&&(o+="';\n"+i+"\n__p+='"),e})),o+="';\n",e.variable||(o="with(obj||{}){\n"+o+"}\n"),o="var __t,__p='',__j=Array.prototype.join,print=function(){__p+=__j.call(arguments,'');};\n"+o+"return __p;\n";try{r=new Function(e.variable||"obj","_",o)}catch(t){throw t.source=o,t}var u=function(t){return r.call(this,t,g)},s=e.variable||"obj";return u.source="function("+s+"){\n"+o+"}",u}function sn(t){var e=g(t);return e._chain=!0,e}function cn(t,e){return t._chain?g(e).chain():e}function ln(t){return L(Qt(t),(function(e){var n=g[e]=t[e];g.prototype[e]=function(){var t=[this._wrapped];return u.apply(t,arguments),cn(this,n.apply(g,t))}})),g}L(["pop","push","reverse","shift","sort","splice","unshift"],(function(t){var e=i[t];g.prototype[t]=function(){var n=this._wrapped;return e.apply(n,arguments),"shift"!==t&&"splice"!==t||0!==n.length||delete n[0],cn(this,n)}})),L(["concat","join","slice"],(function(t){var e=i[t];g.prototype[t]=function(){return cn(this,e.apply(this._wrapped,arguments))}})),g.prototype.value=function(){return this._wrapped},g.prototype.valueOf=g.prototype.toJSON=g.prototype.value,g.prototype.toString=function(){return String(this._wrapped)}}.call(this,n(6))},function(t,e,n){"use strict";n.r(e),function(t){n.d(e,"default",(function(){return g})),n.d(e,"VERSION",(function(){return _})),n.d(e,"iteratee",(function(){return w})),n.d(e,"restArguments",(function(){return k})),n.d(e,"each",(function(){return L})),n.d(e,"forEach",(function(){return L})),n.d(e,"map",(function(){return C})),n.d(e,"collect",(function(){return C})),n.d(e,"reduce",(function(){return N})),n.d(e,"foldl",(function(){return N})),n.d(e,"inject",(function(){return N})),n.d(e,"reduceRight",(function(){return P})),n.d(e,"foldr",(function(){return P})),n.d(e,"find",(function(){return R})),n.d(e,"detect",(function(){return R})),n.d(e,"filter",(function(){return Y})),n.d(e,"select",(function(){return Y})),n.d(e,"reject",(function(){return I})),n.d(e,"every",(function(){return z})),n.d(e,"all",(function(){return z})),n.d(e,"some",(function(){return F})),n.d(e,"any",(function(){return F})),n.d(e,"contains",(function(){return H})),n.d(e,"includes",(function(){return H})),n.d(e,"include",(function(){return H})),n.d(e,"invoke",(function(){return B})),n.d(e,"pluck",(function(){return q})),n.d(e,"where",(function(){return U})),n.d(e,"findWhere",(function(){return W})),n.d(e,"max",(function(){return G})),n.d(e,"min",(function(){return V})),n.d(e,"shuffle",(function(){return $})),n.d(e,"sample",(function(){return X})),n.d(e,"sortBy",(function(){return J})),n.d(e,"groupBy",(function(){return K})),n.d(e,"indexBy",(function(){return Q})),n.d(e,"countBy",(function(){return tt})),n.d(e,"toArray",(function(){return nt})),n.d(e,"size",(function(){return rt})),n.d(e,"partition",(function(){return it})),n.d(e,"first",(function(){return at})),n.d(e,"head",(function(){return at})),n.d(e,"take",(function(){return at})),n.d(e,"initial",(function(){return ot})),n.d(e,"last",(function(){return ut})),n.d(e,"rest",(function(){return st})),n.d(e,"tail",(function(){return st})),n.d(e,"drop",(function(){return st})),n.d(e,"compact",(function(){return ct})),n.d(e,"flatten",(function(){return ft})),n.d(e,"without",(function(){return dt})),n.d(e,"uniq",(function(){return ht})),n.d(e,"unique",(function(){return ht})),n.d(e,"union",(function(){return pt})),n.d(e,"intersection",(function(){return mt})),n.d(e,"difference",(function(){return vt})),n.d(e,"unzip",(function(){return gt})),n.d(e,"zip",(function(){return _t})),n.d(e,"object",(function(){return yt})),n.d(e,"findIndex",(function(){return wt})),n.d(e,"findLastIndex",(function(){return xt})),n.d(e,"sortedIndex",(function(){return kt})),n.d(e,"indexOf",(function(){return Et})),n.d(e,"lastIndexOf",(function(){return St})),n.d(e,"range",(function(){return Tt})),n.d(e,"chunk",(function(){return jt})),n.d(e,"bind",(function(){return At})),n.d(e,"partial",(function(){return Lt})),n.d(e,"bindAll",(function(){return Ct})),n.d(e,"memoize",(function(){return Dt})),n.d(e,"delay",(function(){return Nt})),n.d(e,"defer",(function(){return Pt})),n.d(e,"throttle",(function(){return Rt})),n.d(e,"debounce",(function(){return Yt})),n.d(e,"wrap",(function(){return It})),n.d(e,"negate",(function(){return zt})),n.d(e,"compose",(function(){return Ft})),n.d(e,"after",(function(){return Ht})),n.d(e,"before",(function(){return Bt})),n.d(e,"once",(function(){return qt})),n.d(e,"keys",(function(){return Vt})),n.d(e,"allKeys",(function(){return $t})),n.d(e,"values",(function(){return Xt})),n.d(e,"mapObject",(function(){return Jt})),n.d(e,"pairs",(function(){return Zt})),n.d(e,"invert",(function(){return Kt})),n.d(e,"functions",(function(){return Qt})),n.d(e,"methods",(function(){return Qt})),n.d(e,"extend",(function(){return ee})),n.d(e,"extendOwn",(function(){return ne})),n.d(e,"assign",(function(){return ne})),n.d(e,"findKey",(function(){return re})),n.d(e,"pick",(function(){return ae})),n.d(e,"omit",(function(){return oe})),n.d(e,"defaults",(function(){return ue})),n.d(e,"create",(function(){return se})),n.d(e,"clone",(function(){return ce})),n.d(e,"tap",(function(){return le})),n.d(e,"isMatch",(function(){return fe})),n.d(e,"isEqual",(function(){return he})),n.d(e,"isEmpty",(function(){return pe})),n.d(e,"isElement",(function(){return me})),n.d(e,"isArray",(function(){return ge})),n.d(e,"isObject",(function(){return _e})),n.d(e,"isArguments",(function(){return ye})),n.d(e,"isFunction",(function(){return be})),n.d(e,"isString",(function(){return we})),n.d(e,"isNumber",(function(){return xe})),n.d(e,"isDate",(function(){return ke})),n.d(e,"isRegExp",(function(){return Me})),n.d(e,"isError",(function(){return Ee})),n.d(e,"isSymbol",(function(){return Se})),n.d(e,"isMap",(function(){return Te})),n.d(e,"isWeakMap",(function(){return je})),n.d(e,"isSet",(function(){return Oe})),n.d(e,"isWeakSet",(function(){return Ae})),n.d(e,"isFinite",(function(){return Ce})),n.d(e,"isNaN",(function(){return De})),n.d(e,"isBoolean",(function(){return Ne})),n.d(e,"isNull",(function(){return Pe})),n.d(e,"isUndefined",(function(){return Re})),n.d(e,"has",(function(){return Ye})),n.d(e,"identity",(function(){return Ie})),n.d(e,"constant",(function(){return ze})),n.d(e,"noop",(function(){return Fe})),n.d(e,"property",(function(){return He})),n.d(e,"propertyOf",(function(){return Be})),n.d(e,"matcher",(function(){return qe})),n.d(e,"matches",(function(){return qe})),n.d(e,"times",(function(){return Ue})),n.d(e,"random",(function(){return We})),n.d(e,"now",(function(){return Ge})),n.d(e,"escape",(function(){return Je})),n.d(e,"unescape",(function(){return Ze})),n.d(e,"result",(function(){return Ke})),n.d(e,"uniqueId",(function(){return tn})),n.d(e,"templateSettings",(function(){return en})),n.d(e,"template",(function(){return un})),n.d(e,"chain",(function(){return sn})),n.d(e,"mixin",(function(){return ln}));var r="object"==typeof self&&self.self===self&&self||"object"==typeof t&&t.global===t&&t||Function("return this")()||{},i=Array.prototype,a=Object.prototype,o="undefined"!=typeof Symbol?Symbol.prototype:null,u=i.push,s=i.slice,c=a.toString,l=a.hasOwnProperty,f=Array.isArray,d=Object.keys,h=Object.create,p=r.isNaN,m=r.isFinite,v=function(){};function g(t){return t instanceof g?t:this instanceof g?void(this._wrapped=t):new g(t)}var _=g.VERSION="1.10.2";function y(t,e,n){if(void 0===e)return t;switch(null==n?3:n){case 1:return function(n){return t.call(e,n)};case 3:return function(n,r,i){return t.call(e,n,r,i)};case 4:return function(n,r,i,a){return t.call(e,n,r,i,a)}}return function(){return t.apply(e,arguments)}}function b(t,e,n){return null==t?Ie:be(t)?y(t,e,n):_e(t)&&!ge(t)?qe(t):He(t)}function w(t,e){return b(t,e,1/0)}function x(t,e,n){return g.iteratee!==w?g.iteratee(t,e):b(t,e,n)}function k(t,e){return e=null==e?t.length-1:+e,function(){for(var n=Math.max(arguments.length-e,0),r=Array(n),i=0;i<n;i++)r[i]=arguments[i+e];switch(e){case 0:return t.call(this,r);case 1:return t.call(this,arguments[0],r);case 2:return t.call(this,arguments[0],arguments[1],r)}var a=Array(e+1);for(i=0;i<e;i++)a[i]=arguments[i];return a[e]=r,t.apply(this,a)}}function M(t){if(!_e(t))return{};if(h)return h(t);v.prototype=t;var e=new v;return v.prototype=null,e}function E(t){return function(e){return null==e?void 0:e[t]}}function S(t,e){return null!=t&&l.call(t,e)}function T(t,e){for(var n=e.length,r=0;r<n;r++){if(null==t)return;t=t[e[r]]}return n?t:void 0}g.iteratee=w;var j=Math.pow(2,53)-1,O=E("length");function A(t){var e=O(t);return"number"==typeof e&&e>=0&&e<=j}function L(t,e,n){var r,i;if(e=y(e,n),A(t))for(r=0,i=t.length;r<i;r++)e(t[r],r,t);else{var a=Vt(t);for(r=0,i=a.length;r<i;r++)e(t[a[r]],a[r],t)}return t}function C(t,e,n){e=x(e,n);for(var r=!A(t)&&Vt(t),i=(r||t).length,a=Array(i),o=0;o<i;o++){var u=r?r[o]:o;a[o]=e(t[u],u,t)}return a}function D(t){var e=function(e,n,r,i){var a=!A(e)&&Vt(e),o=(a||e).length,u=t>0?0:o-1;for(i||(r=e[a?a[u]:u],u+=t);u>=0&&u<o;u+=t){var s=a?a[u]:u;r=n(r,e[s],s,e)}return r};return function(t,n,r,i){var a=arguments.length>=3;return e(t,y(n,i,4),r,a)}}var N=D(1),P=D(-1);function R(t,e,n){var r=(A(t)?wt:re)(t,e,n);if(void 0!==r&&-1!==r)return t[r]}function Y(t,e,n){var r=[];return e=x(e,n),L(t,(function(t,n,i){e(t,n,i)&&r.push(t)})),r}function I(t,e,n){return Y(t,zt(x(e)),n)}function z(t,e,n){e=x(e,n);for(var r=!A(t)&&Vt(t),i=(r||t).length,a=0;a<i;a++){var o=r?r[a]:a;if(!e(t[o],o,t))return!1}return!0}function F(t,e,n){e=x(e,n);for(var r=!A(t)&&Vt(t),i=(r||t).length,a=0;a<i;a++){var o=r?r[a]:a;if(e(t[o],o,t))return!0}return!1}function H(t,e,n,r){return A(t)||(t=Xt(t)),("number"!=typeof n||r)&&(n=0),Et(t,e,n)>=0}var B=k((function(t,e,n){var r,i;return be(e)?i=e:ge(e)&&(r=e.slice(0,-1),e=e[e.length-1]),C(t,(function(t){var a=i;if(!a){if(r&&r.length&&(t=T(t,r)),null==t)return;a=t[e]}return null==a?a:a.apply(t,n)}))}));function q(t,e){return C(t,He(e))}function U(t,e){return Y(t,qe(e))}function W(t,e){return R(t,qe(e))}function G(t,e,n){var r,i,a=-1/0,o=-1/0;if(null==e||"number"==typeof e&&"object"!=typeof t[0]&&null!=t)for(var u=0,s=(t=A(t)?t:Xt(t)).length;u<s;u++)null!=(r=t[u])&&r>a&&(a=r);else e=x(e,n),L(t,(function(t,n,r){((i=e(t,n,r))>o||i===-1/0&&a===-1/0)&&(a=t,o=i)}));return a}function V(t,e,n){var r,i,a=1/0,o=1/0;if(null==e||"number"==typeof e&&"object"!=typeof t[0]&&null!=t)for(var u=0,s=(t=A(t)?t:Xt(t)).length;u<s;u++)null!=(r=t[u])&&r<a&&(a=r);else e=x(e,n),L(t,(function(t,n,r){((i=e(t,n,r))<o||i===1/0&&a===1/0)&&(a=t,o=i)}));return a}function $(t){return X(t,1/0)}function X(t,e,n){if(null==e||n)return A(t)||(t=Xt(t)),t[We(t.length-1)];var r=A(t)?ce(t):Xt(t),i=O(r);e=Math.max(Math.min(e,i),0);for(var a=i-1,o=0;o<e;o++){var u=We(o,a),s=r[o];r[o]=r[u],r[u]=s}return r.slice(0,e)}function J(t,e,n){var r=0;return e=x(e,n),q(C(t,(function(t,n,i){return{value:t,index:r++,criteria:e(t,n,i)}})).sort((function(t,e){var n=t.criteria,r=e.criteria;if(n!==r){if(n>r||void 0===n)return 1;if(n<r||void 0===r)return-1}return t.index-e.index})),"value")}function Z(t,e){return function(n,r,i){var a=e?[[],[]]:{};return r=x(r,i),L(n,(function(e,i){var o=r(e,i,n);t(a,e,o)})),a}}var K=Z((function(t,e,n){S(t,n)?t[n].push(e):t[n]=[e]})),Q=Z((function(t,e,n){t[n]=e})),tt=Z((function(t,e,n){S(t,n)?t[n]++:t[n]=1})),et=/[^\ud800-\udfff]|[\ud800-\udbff][\udc00-\udfff]|[\ud800-\udfff]/g;function nt(t){return t?ge(t)?s.call(t):we(t)?t.match(et):A(t)?C(t,Ie):Xt(t):[]}function rt(t){return null==t?0:A(t)?t.length:Vt(t).length}var it=Z((function(t,e,n){t[n?0:1].push(e)}),!0);function at(t,e,n){return null==t||t.length<1?null==e?void 0:[]:null==e||n?t[0]:ot(t,t.length-e)}function ot(t,e,n){return s.call(t,0,Math.max(0,t.length-(null==e||n?1:e)))}function ut(t,e,n){return null==t||t.length<1?null==e?void 0:[]:null==e||n?t[t.length-1]:st(t,Math.max(0,t.length-e))}function st(t,e,n){return s.call(t,null==e||n?1:e)}function ct(t){return Y(t,Boolean)}function lt(t,e,n,r){for(var i=(r=r||[]).length,a=0,o=O(t);a<o;a++){var u=t[a];if(A(u)&&(ge(u)||ye(u)))if(e)for(var s=0,c=u.length;s<c;)r[i++]=u[s++];else lt(u,e,n,r),i=r.length;else n||(r[i++]=u)}return r}function ft(t,e){return lt(t,e,!1)}var dt=k((function(t,e){return vt(t,e)}));function ht(t,e,n,r){Ne(e)||(r=n,n=e,e=!1),null!=n&&(n=x(n,r));for(var i=[],a=[],o=0,u=O(t);o<u;o++){var s=t[o],c=n?n(s,o,t):s;e&&!n?(o&&a===c||i.push(s),a=c):n?H(a,c)||(a.push(c),i.push(s)):H(i,s)||i.push(s)}return i}var pt=k((function(t){return ht(lt(t,!0,!0))}));function mt(t){for(var e=[],n=arguments.length,r=0,i=O(t);r<i;r++){var a=t[r];if(!H(e,a)){var o;for(o=1;o<n&&H(arguments[o],a);o++);o===n&&e.push(a)}}return e}var vt=k((function(t,e){return e=lt(e,!0,!0),Y(t,(function(t){return!H(e,t)}))}));function gt(t){for(var e=t&&G(t,O).length||0,n=Array(e),r=0;r<e;r++)n[r]=q(t,r);return n}var _t=k(gt);function yt(t,e){for(var n={},r=0,i=O(t);r<i;r++)e?n[t[r]]=e[r]:n[t[r][0]]=t[r][1];return n}function bt(t){return function(e,n,r){n=x(n,r);for(var i=O(e),a=t>0?0:i-1;a>=0&&a<i;a+=t)if(n(e[a],a,e))return a;return-1}}var wt=bt(1),xt=bt(-1);function kt(t,e,n,r){for(var i=(n=x(n,r,1))(e),a=0,o=O(t);a<o;){var u=Math.floor((a+o)/2);n(t[u])<i?a=u+1:o=u}return a}function Mt(t,e,n){return function(r,i,a){var o=0,u=O(r);if("number"==typeof a)t>0?o=a>=0?a:Math.max(a+u,o):u=a>=0?Math.min(a+1,u):a+u+1;else if(n&&a&&u)return r[a=n(r,i)]===i?a:-1;if(i!=i)return(a=e(s.call(r,o,u),De))>=0?a+o:-1;for(a=t>0?o:u-1;a>=0&&a<u;a+=t)if(r[a]===i)return a;return-1}}var Et=Mt(1,wt,kt),St=Mt(-1,xt);function Tt(t,e,n){null==e&&(e=t||0,t=0),n||(n=e<t?-1:1);for(var r=Math.max(Math.ceil((e-t)/n),0),i=Array(r),a=0;a<r;a++,t+=n)i[a]=t;return i}function jt(t,e){if(null==e||e<1)return[];for(var n=[],r=0,i=t.length;r<i;)n.push(s.call(t,r,r+=e));return n}function Ot(t,e,n,r,i){if(!(r instanceof e))return t.apply(n,i);var a=M(t.prototype),o=t.apply(a,i);return _e(o)?o:a}var At=k((function(t,e,n){if(!be(t))throw new TypeError("Bind must be called on a function");var r=k((function(i){return Ot(t,r,e,this,n.concat(i))}));return r})),Lt=k((function(t,e){var n=Lt.placeholder,r=function(){for(var i=0,a=e.length,o=Array(a),u=0;u<a;u++)o[u]=e[u]===n?arguments[i++]:e[u];for(;i<arguments.length;)o.push(arguments[i++]);return Ot(t,r,this,this,o)};return r}));Lt.placeholder=g;var Ct=k((function(t,e){var n=(e=lt(e,!1,!1)).length;if(n<1)throw new Error("bindAll must be passed function names");for(;n--;){var r=e[n];t[r]=At(t[r],t)}}));function Dt(t,e){var n=function(r){var i=n.cache,a=""+(e?e.apply(this,arguments):r);return S(i,a)||(i[a]=t.apply(this,arguments)),i[a]};return n.cache={},n}var Nt=k((function(t,e,n){return setTimeout((function(){return t.apply(null,n)}),e)})),Pt=Lt(Nt,g,1);function Rt(t,e,n){var r,i,a,o,u=0;n||(n={});var s=function(){u=!1===n.leading?0:Ge(),r=null,o=t.apply(i,a),r||(i=a=null)},c=function(){var c=Ge();u||!1!==n.leading||(u=c);var l=e-(c-u);return i=this,a=arguments,l<=0||l>e?(r&&(clearTimeout(r),r=null),u=c,o=t.apply(i,a),r||(i=a=null)):r||!1===n.trailing||(r=setTimeout(s,l)),o};return c.cancel=function(){clearTimeout(r),u=0,r=i=a=null},c}function Yt(t,e,n){var r,i,a=function(e,n){r=null,n&&(i=t.apply(e,n))},o=k((function(o){if(r&&clearTimeout(r),n){var u=!r;r=setTimeout(a,e),u&&(i=t.apply(this,o))}else r=Nt(a,e,this,o);return i}));return o.cancel=function(){clearTimeout(r),r=null},o}function It(t,e){return Lt(e,t)}function zt(t){return function(){return!t.apply(this,arguments)}}function Ft(){var t=arguments,e=t.length-1;return function(){for(var n=e,r=t[e].apply(this,arguments);n--;)r=t[n].call(this,r);return r}}function Ht(t,e){return function(){if(--t<1)return e.apply(this,arguments)}}function Bt(t,e){var n;return function(){return--t>0&&(n=e.apply(this,arguments)),t<=1&&(e=null),n}}var qt=Lt(Bt,2),Ut=!{toString:null}.propertyIsEnumerable("toString"),Wt=["valueOf","isPrototypeOf","toString","propertyIsEnumerable","hasOwnProperty","toLocaleString"];function Gt(t,e){var n=Wt.length,r=t.constructor,i=be(r)&&r.prototype||a,o="constructor";for(S(t,o)&&!H(e,o)&&e.push(o);n--;)(o=Wt[n])in t&&t[o]!==i[o]&&!H(e,o)&&e.push(o)}function Vt(t){if(!_e(t))return[];if(d)return d(t);var e=[];for(var n in t)S(t,n)&&e.push(n);return Ut&&Gt(t,e),e}function $t(t){if(!_e(t))return[];var e=[];for(var n in t)e.push(n);return Ut&&Gt(t,e),e}function Xt(t){for(var e=Vt(t),n=e.length,r=Array(n),i=0;i<n;i++)r[i]=t[e[i]];return r}function Jt(t,e,n){e=x(e,n);for(var r=Vt(t),i=r.length,a={},o=0;o<i;o++){var u=r[o];a[u]=e(t[u],u,t)}return a}function Zt(t){for(var e=Vt(t),n=e.length,r=Array(n),i=0;i<n;i++)r[i]=[e[i],t[e[i]]];return r}function Kt(t){for(var e={},n=Vt(t),r=0,i=n.length;r<i;r++)e[t[n[r]]]=n[r];return e}function Qt(t){var e=[];for(var n in t)be(t[n])&&e.push(n);return e.sort()}function te(t,e){return function(n){var r=arguments.length;if(e&&(n=Object(n)),r<2||null==n)return n;for(var i=1;i<r;i++)for(var a=arguments[i],o=t(a),u=o.length,s=0;s<u;s++){var c=o[s];e&&void 0!==n[c]||(n[c]=a[c])}return n}}var ee=te($t),ne=te(Vt);function re(t,e,n){e=x(e,n);for(var r,i=Vt(t),a=0,o=i.length;a<o;a++)if(e(t[r=i[a]],r,t))return r}function ie(t,e,n){return e in n}var ae=k((function(t,e){var n={},r=e[0];if(null==t)return n;be(r)?(e.length>1&&(r=y(r,e[1])),e=$t(t)):(r=ie,e=lt(e,!1,!1),t=Object(t));for(var i=0,a=e.length;i<a;i++){var o=e[i],u=t[o];r(u,o,t)&&(n[o]=u)}return n})),oe=k((function(t,e){var n,r=e[0];return be(r)?(r=zt(r),e.length>1&&(n=e[1])):(e=C(lt(e,!1,!1),String),r=function(t,n){return!H(e,n)}),ae(t,r,n)})),ue=te($t,!0);function se(t,e){var n=M(t);return e&&ne(n,e),n}function ce(t){return _e(t)?ge(t)?t.slice():ee({},t):t}function le(t,e){return e(t),t}function fe(t,e){var n=Vt(e),r=n.length;if(null==t)return!r;for(var i=Object(t),a=0;a<r;a++){var o=n[a];if(e[o]!==i[o]||!(o in i))return!1}return!0}function de(t,e,n,r){if(t===e)return 0!==t||1/t==1/e;if(null==t||null==e)return!1;if(t!=t)return e!=e;var i=typeof t;return("function"===i||"object"===i||"object"==typeof e)&&function(t,e,n,r){t instanceof g&&(t=t._wrapped);e instanceof g&&(e=e._wrapped);var i=c.call(t);if(i!==c.call(e))return!1;switch(i){case"[object RegExp]":case"[object String]":return""+t==""+e;case"[object Number]":return+t!=+t?+e!=+e:0==+t?1/+t==1/e:+t==+e;case"[object Date]":case"[object Boolean]":return+t==+e;case"[object Symbol]":return o.valueOf.call(t)===o.valueOf.call(e)}var a="[object Array]"===i;if(!a){if("object"!=typeof t||"object"!=typeof e)return!1;var u=t.constructor,s=e.constructor;if(u!==s&&!(be(u)&&u instanceof u&&be(s)&&s instanceof s)&&"constructor"in t&&"constructor"in e)return!1}r=r||[];var l=(n=n||[]).length;for(;l--;)if(n[l]===t)return r[l]===e;if(n.push(t),r.push(e),a){if((l=t.length)!==e.length)return!1;for(;l--;)if(!de(t[l],e[l],n,r))return!1}else{var f,d=Vt(t);if(l=d.length,Vt(e).length!==l)return!1;for(;l--;)if(f=d[l],!S(e,f)||!de(t[f],e[f],n,r))return!1}return n.pop(),r.pop(),!0}(t,e,n,r)}function he(t,e){return de(t,e)}function pe(t){return null==t||(A(t)&&(ge(t)||we(t)||ye(t))?0===t.length:0===Vt(t).length)}function me(t){return!(!t||1!==t.nodeType)}function ve(t){return function(e){return c.call(e)==="[object "+t+"]"}}var ge=f||ve("Array");function _e(t){var e=typeof t;return"function"===e||"object"===e&&!!t}var ye=ve("Arguments"),be=ve("Function"),we=ve("String"),xe=ve("Number"),ke=ve("Date"),Me=ve("RegExp"),Ee=ve("Error"),Se=ve("Symbol"),Te=ve("Map"),je=ve("WeakMap"),Oe=ve("Set"),Ae=ve("WeakSet");!function(){ye(arguments)||(ye=function(t){return S(t,"callee")})}();var Le=r.document&&r.document.childNodes;function Ce(t){return!Se(t)&&m(t)&&!p(parseFloat(t))}function De(t){return xe(t)&&p(t)}function Ne(t){return!0===t||!1===t||"[object Boolean]"===c.call(t)}function Pe(t){return null===t}function Re(t){return void 0===t}function Ye(t,e){if(!ge(e))return S(t,e);for(var n=e.length,r=0;r<n;r++){var i=e[r];if(null==t||!l.call(t,i))return!1;t=t[i]}return!!n}function Ie(t){return t}function ze(t){return function(){return t}}function Fe(){}function He(t){return ge(t)?function(e){return T(e,t)}:E(t)}function Be(t){return null==t?function(){}:function(e){return ge(e)?T(t,e):t[e]}}function qe(t){return t=ne({},t),function(e){return fe(e,t)}}function Ue(t,e,n){var r=Array(Math.max(0,t));e=y(e,n,1);for(var i=0;i<t;i++)r[i]=e(i);return r}function We(t,e){return null==e&&(e=t,t=0),t+Math.floor(Math.random()*(e-t+1))}"object"!=typeof Int8Array&&"function"!=typeof Le&&(be=function(t){return"function"==typeof t||!1});var Ge=Date.now||function(){return(new Date).getTime()},Ve={"&":"&amp;","<":"&lt;",">":"&gt;",'"':"&quot;","'":"&#x27;","`":"&#x60;"},$e=Kt(Ve);function Xe(t){var e=function(e){return t[e]},n="(?:"+Vt(t).join("|")+")",r=RegExp(n),i=RegExp(n,"g");return function(t){return t=null==t?"":""+t,r.test(t)?t.replace(i,e):t}}var Je=Xe(Ve),Ze=Xe($e);function Ke(t,e,n){ge(e)||(e=[e]);var r=e.length;if(!r)return be(n)?n.call(t):n;for(var i=0;i<r;i++){var a=null==t?void 0:t[e[i]];void 0===a&&(a=n,i=r),t=be(a)?a.call(t):a}return t}var Qe=0;function tn(t){var e=++Qe+"";return t?t+e:e}var en=g.templateSettings={evaluate:/<%([\s\S]+?)%>/g,interpolate:/<%=([\s\S]+?)%>/g,escape:/<%-([\s\S]+?)%>/g},nn=/(.)^/,rn={"'":"'","\\":"\\","\r":"r","\n":"n","\u2028":"u2028","\u2029":"u2029"},an=/\\|'|\r|\n|\u2028|\u2029/g,on=function(t){return"\\"+rn[t]};function un(t,e,n){!e&&n&&(e=n),e=ue({},e,g.templateSettings);var r,i=RegExp([(e.escape||nn).source,(e.interpolate||nn).source,(e.evaluate||nn).source].join("|")+"|$","g"),a=0,o="__p+='";t.replace(i,(function(e,n,r,i,u){return o+=t.slice(a,u).replace(an,on),a=u+e.length,n?o+="'+\n((__t=("+n+"))==null?'':_.escape(__t))+\n'":r?o+="'+\n((__t=("+r+"))==null?'':__t)+\n'":i&&(o+="';\n"+i+"\n__p+='"),e})),o+="';\n",e.variable||(o="with(obj||{}){\n"+o+"}\n"),o="var __t,__p='',__j=Array.prototype.join,print=function(){__p+=__j.call(arguments,'');};\n"+o+"return __p;\n";try{r=new Function(e.variable||"obj","_",o)}catch(t){throw t.source=o,t}var u=function(t){return r.call(this,t,g)},s=e.variable||"obj";return u.source="function("+s+"){\n"+o+"}",u}function sn(t){var e=g(t);return e._chain=!0,e}function cn(t,e){return t._chain?g(e).chain():e}function ln(t){return L(Qt(t),(function(e){var n=g[e]=t[e];g.prototype[e]=function(){var t=[this._wrapped];return u.apply(t,arguments),cn(this,n.apply(g,t))}})),g}L(["pop","push","reverse","shift","sort","splice","unshift"],(function(t){var e=i[t];g.prototype[t]=function(){var n=this._wrapped;return e.apply(n,arguments),"shift"!==t&&"splice"!==t||0!==n.length||delete n[0],cn(this,n)}})),L(["concat","join","slice"],(function(t){var e=i[t];g.prototype[t]=function(){return cn(this,e.apply(this._wrapped,arguments))}})),g.prototype.value=function(){return this._wrapped},g.prototype.valueOf=g.prototype.toJSON=g.prototype.value,g.prototype.toString=function(){return String(this._wrapped)}}.call(this,n(6))},function(t,e,n){"use strict";var r=n(42);e.a=function(){for(var t,e=r.c;t=e.sourceEvent;)e=t;return e}},function(t,e,n){"use strict";var r=n(44);e.a=function(t){return"string"==typeof t?new r.a([[document.querySelector(t)]],[document.documentElement]):new r.a([[t]],r.c)}},function(t,e,n){"use strict";e.a=function(t){return function(){return t}}},function(t,e,n){"use strict";n.d(e,"a",(function(){return r})),n.d(e,"b",(function(){return i}));var r=Math.PI/180,i=180/Math.PI},function(t,e){t.exports=Array.isArray||function(t){return"[object Array]"==Object.prototype.toString.call(t)}},function(t,e,n){"use strict";function r(t){return t*t*t}function i(t){return--t*t*t+1}function a(t){return((t*=2)<=1?t*t*t:(t-=2)*t*t+2)/2}n.d(e,"a",(function(){return r})),n.d(e,"c",(function(){return i})),n.d(e,"b",(function(){return a}))},function(t,e,n){"use strict";n.d(e,"b",(function(){return c})),n.d(e,"a",(function(){return l})),n.d(e,"d",(function(){return g})),n.d(e,"c",(function(){return _}));var r=n(26),i=n(14),a=n(108),o=6/29,u=3*o*o;function s(t){if(t instanceof f)return new f(t.l,t.a,t.b,t.opacity);if(t instanceof y)return b(t);t instanceof i.b||(t=Object(i.h)(t));var e,n,r=m(t.r),a=m(t.g),o=m(t.b),u=d((.2225045*r+.7168786*a+.0606169*o)/1);return r===a&&a===o?e=n=u:(e=d((.4360747*r+.3850649*a+.1430804*o)/.96422),n=d((.0139322*r+.0971045*a+.7141733*o)/.82521)),new f(116*u-16,500*(e-u),200*(u-n),t.opacity)}function c(t,e){return new f(t,0,0,null==e?1:e)}function l(t,e,n,r){return 1===arguments.length?s(t):new f(t,e,n,null==r?1:r)}function f(t,e,n,r){this.l=+t,this.a=+e,this.b=+n,this.opacity=+r}function d(t){return t>.008856451679035631?Math.pow(t,1/3):t/u+4/29}function h(t){return t>o?t*t*t:u*(t-4/29)}function p(t){return 255*(t<=.0031308?12.92*t:1.055*Math.pow(t,1/2.4)-.055)}function m(t){return(t/=255)<=.04045?t/12.92:Math.pow((t+.055)/1.055,2.4)}function v(t){if(t instanceof y)return new y(t.h,t.c,t.l,t.opacity);if(t instanceof f||(t=s(t)),0===t.a&&0===t.b)return new y(NaN,0<t.l&&t.l<100?0:NaN,t.l,t.opacity);var e=Math.atan2(t.b,t.a)*a.b;return new y(e<0?e+360:e,Math.sqrt(t.a*t.a+t.b*t.b),t.l,t.opacity)}function g(t,e,n,r){return 1===arguments.length?v(t):new y(n,e,t,null==r?1:r)}function _(t,e,n,r){return 1===arguments.length?v(t):new y(t,e,n,null==r?1:r)}function y(t,e,n,r){this.h=+t,this.c=+e,this.l=+n,this.opacity=+r}function b(t){if(isNaN(t.h))return new f(t.l,0,0,t.opacity);var e=t.h*a.a;return new f(t.l,Math.cos(e)*t.c,Math.sin(e)*t.c,t.opacity)}Object(r.a)(f,l,Object(r.b)(i.a,{brighter:function(t){return new f(this.l+18*(null==t?1:t),this.a,this.b,this.opacity)},darker:function(t){return new f(this.l-18*(null==t?1:t),this.a,this.b,this.opacity)},rgb:function(){var t=(this.l+16)/116,e=isNaN(this.a)?t:t+this.a/500,n=isNaN(this.b)?t:t-this.b/200;return e=.96422*h(e),t=1*h(t),n=.82521*h(n),new i.b(p(3.1338561*e-1.6168667*t-.4906146*n),p(-.9787684*e+1.9161415*t+.033454*n),p(.0719453*e-.2289914*t+1.4052427*n),this.opacity)}})),Object(r.a)(y,_,Object(r.b)(i.a,{brighter:function(t){return new y(this.h,this.c,this.l+18*(null==t?1:t),this.opacity)},darker:function(t){return new y(this.h,this.c,this.l-18*(null==t?1:t),this.opacity)},rgb:function(){return b(this).rgb()}}))},function(t,e,n){"use strict";n.d(e,"b",(function(){return o}));var r=n(7),i=n(9),a=Object(r.a)((function(t){t.setTime(t-t.getMilliseconds())}),(function(t,e){t.setTime(+t+e*i.d)}),(function(t,e){return(e-t)/i.d}),(function(t){return t.getUTCSeconds()}));e.a=a;var o=a.range},function(t,e,n){"use strict";n.d(e,"b",(function(){return a}));var r=n(7),i=Object(r.a)((function(){}),(function(t,e){t.setTime(+t+e)}),(function(t,e){return e-t}));i.every=function(t){return t=Math.floor(t),isFinite(t)&&t>0?t>1?Object(r.a)((function(e){e.setTime(Math.floor(e/t)*t)}),(function(e,n){e.setTime(+e+n*t)}),(function(e,n){return(n-e)/t})):i:null},e.a=i;var a=i.range},function(t,e){t.exports=function(t,e){return{enumerable:!(1&t),configurable:!(2&t),writable:!(4&t),value:e}}},function(t,e,n){var r=n(267),i=n(198);t.exports=Object.keys||function(t){return r(t,i)}},function(t,e){var n={}.toString;t.exports=function(t){return n.call(t).slice(8,-1)}},function(t,e){t.exports=!0},function(t,e,n){t.exports=n(275)},function(t,e,n){var r=n(120);function i(t,e,n,i,a,o,u){try{var s=t[o](u),c=s.value}catch(t){return void n(t)}s.done?e(c):r.resolve(c).then(i,a)}t.exports=function(t){return function(){var e=this,n=arguments;return new r((function(r,a){var o=t.apply(e,n);function u(t){i(o,r,a,u,s,"next",t)}function s(t){i(o,r,a,u,s,"throw",t)}u(void 0)}))}}},function(t,e,n){t.exports=n(556)},function(t,e,n){"use strict";var r=n(557)(!0);n(277)(String,"String",(function(t){this._t=String(t),this._i=0}),(function(){var t,e=this._t,n=this._i;return n>=e.length?{value:void 0,done:!0}:(t=r(e,n),this._i+=t.length,{value:t,done:!1})}))},function(t,e,n){(function(t){function n(t){return Object.prototype.toString.call(t)}e.isArray=function(t){return Array.isArray?Array.isArray(t):"[object Array]"===n(t)},e.isBoolean=function(t){return"boolean"==typeof t},e.isNull=function(t){return null===t},e.isNullOrUndefined=function(t){return null==t},e.isNumber=function(t){return"number"==typeof t},e.isString=function(t){return"string"==typeof t},e.isSymbol=function(t){return"symbol"==typeof t},e.isUndefined=function(t){return void 0===t},e.isRegExp=function(t){return"[object RegExp]"===n(t)},e.isObject=function(t){return"object"==typeof t&&null!==t},e.isDate=function(t){return"[object Date]"===n(t)},e.isError=function(t){return"[object Error]"===n(t)||t instanceof Error},e.isFunction=function(t){return"function"==typeof t},e.isPrimitive=function(t){return null===t||"boolean"==typeof t||"number"==typeof t||"string"==typeof t||"symbol"==typeof t||void 0===t},e.isBuffer=t.isBuffer}).call(this,n(28).Buffer)},function(t,e,n){"use strict";n.d(e,"a",(function(){return M})),n.d(e,"b",(function(){return j})),n.d(e,"v",(function(){return S.a})),n.d(e,"A",(function(){return L})),n.d(e,"c",(function(){return Y})),n.d(e,"C",(function(){return Y})),n.d(e,"w",(function(){return R})),n.d(e,"D",(function(){return R})),n.d(e,"B",(function(){return I})),n.d(e,"x",(function(){return G})),n.d(e,"z",(function(){return V})),n.d(e,"y",(function(){return $})),n.d(e,"Q",(function(){return dt})),n.d(e,"Y",(function(){return ft})),n.d(e,"R",(function(){return X})),n.d(e,"S",(function(){return J})),n.d(e,"T",(function(){return Q})),n.d(e,"U",(function(){return it})),n.d(e,"V",(function(){return rt})),n.d(e,"W",(function(){return ot})),n.d(e,"X",(function(){return lt})),n.d(e,"e",(function(){return _t})),n.d(e,"f",(function(){return bt})),n.d(e,"d",(function(){return vt})),n.d(e,"g",(function(){return xt})),n.d(e,"i",(function(){return Tt})),n.d(e,"j",(function(){return Ot})),n.d(e,"h",(function(){return Et})),n.d(e,"l",(function(){return Nt})),n.d(e,"m",(function(){return Rt})),n.d(e,"k",(function(){return Ct})),n.d(e,"o",(function(){return It})),n.d(e,"n",(function(){return E.a})),n.d(e,"p",(function(){return Gt})),n.d(e,"q",(function(){return Vt})),n.d(e,"r",(function(){return Jt})),n.d(e,"s",(function(){return Kt})),n.d(e,"t",(function(){return te})),n.d(e,"u",(function(){return Qt})),n.d(e,"E",(function(){return ie})),n.d(e,"G",(function(){return ae})),n.d(e,"F",(function(){return oe})),n.d(e,"H",(function(){return ee})),n.d(e,"I",(function(){return ue})),n.d(e,"J",(function(){return se})),n.d(e,"K",(function(){return ce})),n.d(e,"L",(function(){return fe})),n.d(e,"M",(function(){return he})),n.d(e,"N",(function(){return pe})),n.d(e,"O",(function(){return ne})),n.d(e,"P",(function(){return me}));var r=n(228),i=n(5),a=Math.abs,o=Math.atan2,u=Math.cos,s=Math.max,c=Math.min,l=Math.sin,f=Math.sqrt,d=Math.PI,h=d/2,p=2*d;function m(t){return t>1?0:t<-1?d:Math.acos(t)}function v(t){return t>=1?h:t<=-1?-h:Math.asin(t)}function g(t){return t.innerRadius}function _(t){return t.outerRadius}function y(t){return t.startAngle}function b(t){return t.endAngle}function w(t){return t&&t.padAngle}function x(t,e,n,r,i,a,o,u){var s=n-t,c=r-e,l=o-i,f=u-a,d=f*s-l*c;if(!(d*d<1e-12))return[t+(d=(l*(e-a)-f*(t-i))/d)*s,e+d*c]}function k(t,e,n,r,i,a,o){var u=t-n,c=e-r,l=(o?a:-a)/f(u*u+c*c),d=l*c,h=-l*u,p=t+d,m=e+h,v=n+d,g=r+h,_=(p+v)/2,y=(m+g)/2,b=v-p,w=g-m,x=b*b+w*w,k=i-a,M=p*g-v*m,E=(w<0?-1:1)*f(s(0,k*k*x-M*M)),S=(M*w-b*E)/x,T=(-M*b-w*E)/x,j=(M*w+b*E)/x,O=(-M*b+w*E)/x,A=S-_,L=T-y,C=j-_,D=O-y;return A*A+L*L>C*C+D*D&&(S=j,T=O),{cx:S,cy:T,x01:-d,y01:-h,x11:S*(i/k-1),y11:T*(i/k-1)}}var M=function(){var t=g,e=_,n=Object(i.a)(0),s=null,M=y,E=b,S=w,T=null;function j(){var i,g,_=+t.apply(this,arguments),y=+e.apply(this,arguments),b=M.apply(this,arguments)-h,w=E.apply(this,arguments)-h,j=a(w-b),O=w>b;if(T||(T=i=Object(r.a)()),y<_&&(g=y,y=_,_=g),y>1e-12)if(j>p-1e-12)T.moveTo(y*u(b),y*l(b)),T.arc(0,0,y,b,w,!O),_>1e-12&&(T.moveTo(_*u(w),_*l(w)),T.arc(0,0,_,w,b,O));else{var A,L,C=b,D=w,N=b,P=w,R=j,Y=j,I=S.apply(this,arguments)/2,z=I>1e-12&&(s?+s.apply(this,arguments):f(_*_+y*y)),F=c(a(y-_)/2,+n.apply(this,arguments)),H=F,B=F;if(z>1e-12){var q=v(z/_*l(I)),U=v(z/y*l(I));(R-=2*q)>1e-12?(N+=q*=O?1:-1,P-=q):(R=0,N=P=(b+w)/2),(Y-=2*U)>1e-12?(C+=U*=O?1:-1,D-=U):(Y=0,C=D=(b+w)/2)}var W=y*u(C),G=y*l(C),V=_*u(P),$=_*l(P);if(F>1e-12){var X,J=y*u(D),Z=y*l(D),K=_*u(N),Q=_*l(N);if(j<d&&(X=x(W,G,K,Q,J,Z,V,$))){var tt=W-X[0],et=G-X[1],nt=J-X[0],rt=Z-X[1],it=1/l(m((tt*nt+et*rt)/(f(tt*tt+et*et)*f(nt*nt+rt*rt)))/2),at=f(X[0]*X[0]+X[1]*X[1]);H=c(F,(_-at)/(it-1)),B=c(F,(y-at)/(it+1))}}Y>1e-12?B>1e-12?(A=k(K,Q,W,G,y,B,O),L=k(J,Z,V,$,y,B,O),T.moveTo(A.cx+A.x01,A.cy+A.y01),B<F?T.arc(A.cx,A.cy,B,o(A.y01,A.x01),o(L.y01,L.x01),!O):(T.arc(A.cx,A.cy,B,o(A.y01,A.x01),o(A.y11,A.x11),!O),T.arc(0,0,y,o(A.cy+A.y11,A.cx+A.x11),o(L.cy+L.y11,L.cx+L.x11),!O),T.arc(L.cx,L.cy,B,o(L.y11,L.x11),o(L.y01,L.x01),!O))):(T.moveTo(W,G),T.arc(0,0,y,C,D,!O)):T.moveTo(W,G),_>1e-12&&R>1e-12?H>1e-12?(A=k(V,$,J,Z,_,-H,O),L=k(W,G,K,Q,_,-H,O),T.lineTo(A.cx+A.x01,A.cy+A.y01),H<F?T.arc(A.cx,A.cy,H,o(A.y01,A.x01),o(L.y01,L.x01),!O):(T.arc(A.cx,A.cy,H,o(A.y01,A.x01),o(A.y11,A.x11),!O),T.arc(0,0,_,o(A.cy+A.y11,A.cx+A.x11),o(L.cy+L.y11,L.cx+L.x11),O),T.arc(L.cx,L.cy,H,o(L.y11,L.x11),o(L.y01,L.x01),!O))):T.arc(0,0,_,P,N,O):T.lineTo(V,$)}else T.moveTo(0,0);if(T.closePath(),i)return T=null,i+""||null}return j.centroid=function(){var n=(+t.apply(this,arguments)+ +e.apply(this,arguments))/2,r=(+M.apply(this,arguments)+ +E.apply(this,arguments))/2-d/2;return[u(r)*n,l(r)*n]},j.innerRadius=function(e){return arguments.length?(t="function"==typeof e?e:Object(i.a)(+e),j):t},j.outerRadius=function(t){return arguments.length?(e="function"==typeof t?t:Object(i.a)(+t),j):e},j.cornerRadius=function(t){return arguments.length?(n="function"==typeof t?t:Object(i.a)(+t),j):n},j.padRadius=function(t){return arguments.length?(s=null==t?null:"function"==typeof t?t:Object(i.a)(+t),j):s},j.startAngle=function(t){return arguments.length?(M="function"==typeof t?t:Object(i.a)(+t),j):M},j.endAngle=function(t){return arguments.length?(E="function"==typeof t?t:Object(i.a)(+t),j):E},j.padAngle=function(t){return arguments.length?(S="function"==typeof t?t:Object(i.a)(+t),j):S},j.context=function(t){return arguments.length?(T=null==t?null:t,j):T},j},E=n(61),S=n(102),T=n(50),j=function(){var t=T.a,e=null,n=Object(i.a)(0),a=T.b,o=Object(i.a)(!0),u=null,s=E.a,c=null;function l(i){var l,f,d,h,p,m=i.length,v=!1,g=new Array(m),_=new Array(m);for(null==u&&(c=s(p=Object(r.a)())),l=0;l<=m;++l){if(!(l<m&&o(h=i[l],l,i))===v)if(v=!v)f=l,c.areaStart(),c.lineStart();else{for(c.lineEnd(),c.lineStart(),d=l-1;d>=f;--d)c.point(g[d],_[d]);c.lineEnd(),c.areaEnd()}v&&(g[l]=+t(h,l,i),_[l]=+n(h,l,i),c.point(e?+e(h,l,i):g[l],a?+a(h,l,i):_[l]))}if(p)return c=null,p+""||null}function f(){return Object(S.a)().defined(o).curve(s).context(u)}return l.x=function(n){return arguments.length?(t="function"==typeof n?n:Object(i.a)(+n),e=null,l):t},l.x0=function(e){return arguments.length?(t="function"==typeof e?e:Object(i.a)(+e),l):t},l.x1=function(t){return arguments.length?(e=null==t?null:"function"==typeof t?t:Object(i.a)(+t),l):e},l.y=function(t){return arguments.length?(n="function"==typeof t?t:Object(i.a)(+t),a=null,l):n},l.y0=function(t){return arguments.length?(n="function"==typeof t?t:Object(i.a)(+t),l):n},l.y1=function(t){return arguments.length?(a=null==t?null:"function"==typeof t?t:Object(i.a)(+t),l):a},l.lineX0=l.lineY0=function(){return f().x(t).y(n)},l.lineY1=function(){return f().x(t).y(a)},l.lineX1=function(){return f().x(e).y(n)},l.defined=function(t){return arguments.length?(o="function"==typeof t?t:Object(i.a)(!!t),l):o},l.curve=function(t){return arguments.length?(s=t,null!=u&&(c=s(u)),l):s},l.context=function(t){return arguments.length?(null==t?u=c=null:c=s(u=t),l):u},l},O=function(t,e){return e<t?-1:e>t?1:e>=t?0:NaN},A=function(t){return t},L=function(){var t=A,e=O,n=null,r=Object(i.a)(0),a=Object(i.a)(p),o=Object(i.a)(0);function u(i){var u,s,c,l,f,d=i.length,h=0,m=new Array(d),v=new Array(d),g=+r.apply(this,arguments),_=Math.min(p,Math.max(-p,a.apply(this,arguments)-g)),y=Math.min(Math.abs(_)/d,o.apply(this,arguments)),b=y*(_<0?-1:1);for(u=0;u<d;++u)(f=v[m[u]=u]=+t(i[u],u,i))>0&&(h+=f);for(null!=e?m.sort((function(t,n){return e(v[t],v[n])})):null!=n&&m.sort((function(t,e){return n(i[t],i[e])})),u=0,c=h?(_-d*b)/h:0;u<d;++u,g=l)s=m[u],l=g+((f=v[s])>0?f*c:0)+b,v[s]={data:i[s],index:u,value:f,startAngle:g,endAngle:l,padAngle:y};return v}return u.value=function(e){return arguments.length?(t="function"==typeof e?e:Object(i.a)(+e),u):t},u.sortValues=function(t){return arguments.length?(e=t,n=null,u):e},u.sort=function(t){return arguments.length?(n=t,e=null,u):n},u.startAngle=function(t){return arguments.length?(r="function"==typeof t?t:Object(i.a)(+t),u):r},u.endAngle=function(t){return arguments.length?(a="function"==typeof t?t:Object(i.a)(+t),u):a},u.padAngle=function(t){return arguments.length?(o="function"==typeof t?t:Object(i.a)(+t),u):o},u},C=N(E.a);function D(t){this._curve=t}function N(t){function e(e){return new D(t(e))}return e._curve=t,e}function P(t){var e=t.curve;return t.angle=t.x,delete t.x,t.radius=t.y,delete t.y,t.curve=function(t){return arguments.length?e(N(t)):e()._curve},t}D.prototype={areaStart:function(){this._curve.areaStart()},areaEnd:function(){this._curve.areaEnd()},lineStart:function(){this._curve.lineStart()},lineEnd:function(){this._curve.lineEnd()},point:function(t,e){this._curve.point(e*Math.sin(t),e*-Math.cos(t))}};var R=function(){return P(Object(S.a)().curve(C))},Y=function(){var t=j().curve(C),e=t.curve,n=t.lineX0,r=t.lineX1,i=t.lineY0,a=t.lineY1;return t.angle=t.x,delete t.x,t.startAngle=t.x0,delete t.x0,t.endAngle=t.x1,delete t.x1,t.radius=t.y,delete t.y,t.innerRadius=t.y0,delete t.y0,t.outerRadius=t.y1,delete t.y1,t.lineStartAngle=function(){return P(n())},delete t.lineX0,t.lineEndAngle=function(){return P(r())},delete t.lineX1,t.lineInnerRadius=function(){return P(i())},delete t.lineY0,t.lineOuterRadius=function(){return P(a())},delete t.lineY1,t.curve=function(t){return arguments.length?e(N(t)):e()._curve},t},I=function(t,e){return[(e=+e)*Math.cos(t-=Math.PI/2),e*Math.sin(t)]},z=Array.prototype.slice;function F(t){return t.source}function H(t){return t.target}function B(t){var e=F,n=H,a=T.a,o=T.b,u=null;function s(){var i,s=z.call(arguments),c=e.apply(this,s),l=n.apply(this,s);if(u||(u=i=Object(r.a)()),t(u,+a.apply(this,(s[0]=c,s)),+o.apply(this,s),+a.apply(this,(s[0]=l,s)),+o.apply(this,s)),i)return u=null,i+""||null}return s.source=function(t){return arguments.length?(e=t,s):e},s.target=function(t){return arguments.length?(n=t,s):n},s.x=function(t){return arguments.length?(a="function"==typeof t?t:Object(i.a)(+t),s):a},s.y=function(t){return arguments.length?(o="function"==typeof t?t:Object(i.a)(+t),s):o},s.context=function(t){return arguments.length?(u=null==t?null:t,s):u},s}function q(t,e,n,r,i){t.moveTo(e,n),t.bezierCurveTo(e=(e+r)/2,n,e,i,r,i)}function U(t,e,n,r,i){t.moveTo(e,n),t.bezierCurveTo(e,n=(n+i)/2,r,n,r,i)}function W(t,e,n,r,i){var a=I(e,n),o=I(e,n=(n+i)/2),u=I(r,n),s=I(r,i);t.moveTo(a[0],a[1]),t.bezierCurveTo(o[0],o[1],u[0],u[1],s[0],s[1])}function G(){return B(q)}function V(){return B(U)}function $(){var t=B(W);return t.angle=t.x,delete t.x,t.radius=t.y,delete t.y,t}var X={draw:function(t,e){var n=Math.sqrt(e/d);t.moveTo(n,0),t.arc(0,0,n,0,p)}},J={draw:function(t,e){var n=Math.sqrt(e/5)/2;t.moveTo(-3*n,-n),t.lineTo(-n,-n),t.lineTo(-n,-3*n),t.lineTo(n,-3*n),t.lineTo(n,-n),t.lineTo(3*n,-n),t.lineTo(3*n,n),t.lineTo(n,n),t.lineTo(n,3*n),t.lineTo(-n,3*n),t.lineTo(-n,n),t.lineTo(-3*n,n),t.closePath()}},Z=Math.sqrt(1/3),K=2*Z,Q={draw:function(t,e){var n=Math.sqrt(e/K),r=n*Z;t.moveTo(0,-n),t.lineTo(r,0),t.lineTo(0,n),t.lineTo(-r,0),t.closePath()}},tt=Math.sin(d/10)/Math.sin(7*d/10),et=Math.sin(p/10)*tt,nt=-Math.cos(p/10)*tt,rt={draw:function(t,e){var n=Math.sqrt(.8908130915292852*e),r=et*n,i=nt*n;t.moveTo(0,-n),t.lineTo(r,i);for(var a=1;a<5;++a){var o=p*a/5,u=Math.cos(o),s=Math.sin(o);t.lineTo(s*n,-u*n),t.lineTo(u*r-s*i,s*r+u*i)}t.closePath()}},it={draw:function(t,e){var n=Math.sqrt(e),r=-n/2;t.rect(r,r,n,n)}},at=Math.sqrt(3),ot={draw:function(t,e){var n=-Math.sqrt(e/(3*at));t.moveTo(0,2*n),t.lineTo(-at*n,-n),t.lineTo(at*n,-n),t.closePath()}},ut=Math.sqrt(3)/2,st=1/Math.sqrt(12),ct=3*(st/2+1),lt={draw:function(t,e){var n=Math.sqrt(e/ct),r=n/2,i=n*st,a=r,o=n*st+n,u=-a,s=o;t.moveTo(r,i),t.lineTo(a,o),t.lineTo(u,s),t.lineTo(-.5*r-ut*i,ut*r+-.5*i),t.lineTo(-.5*a-ut*o,ut*a+-.5*o),t.lineTo(-.5*u-ut*s,ut*u+-.5*s),t.lineTo(-.5*r+ut*i,-.5*i-ut*r),t.lineTo(-.5*a+ut*o,-.5*o-ut*a),t.lineTo(-.5*u+ut*s,-.5*s-ut*u),t.closePath()}},ft=[X,J,Q,it,rt,ot,lt],dt=function(){var t=Object(i.a)(X),e=Object(i.a)(64),n=null;function a(){var i;if(n||(n=i=Object(r.a)()),t.apply(this,arguments).draw(n,+e.apply(this,arguments)),i)return n=null,i+""||null}return a.type=function(e){return arguments.length?(t="function"==typeof e?e:Object(i.a)(e),a):t},a.size=function(t){return arguments.length?(e="function"==typeof t?t:Object(i.a)(+t),a):e},a.context=function(t){return arguments.length?(n=null==t?null:t,a):n},a},ht=function(){};function pt(t,e,n){t._context.bezierCurveTo((2*t._x0+t._x1)/3,(2*t._y0+t._y1)/3,(t._x0+2*t._x1)/3,(t._y0+2*t._y1)/3,(t._x0+4*t._x1+e)/6,(t._y0+4*t._y1+n)/6)}function mt(t){this._context=t}mt.prototype={areaStart:function(){this._line=0},areaEnd:function(){this._line=NaN},lineStart:function(){this._x0=this._x1=this._y0=this._y1=NaN,this._point=0},lineEnd:function(){switch(this._point){case 3:pt(this,this._x1,this._y1);case 2:this._context.lineTo(this._x1,this._y1)}(this._line||0!==this._line&&1===this._point)&&this._context.closePath(),this._line=1-this._line},point:function(t,e){switch(t=+t,e=+e,this._point){case 0:this._point=1,this._line?this._context.lineTo(t,e):this._context.moveTo(t,e);break;case 1:this._point=2;break;case 2:this._point=3,this._context.lineTo((5*this._x0+this._x1)/6,(5*this._y0+this._y1)/6);default:pt(this,t,e)}this._x0=this._x1,this._x1=t,this._y0=this._y1,this._y1=e}};var vt=function(t){return new mt(t)};function gt(t){this._context=t}gt.prototype={areaStart:ht,areaEnd:ht,lineStart:function(){this._x0=this._x1=this._x2=this._x3=this._x4=this._y0=this._y1=this._y2=this._y3=this._y4=NaN,this._point=0},lineEnd:function(){switch(this._point){case 1:this._context.moveTo(this._x2,this._y2),this._context.closePath();break;case 2:this._context.moveTo((this._x2+2*this._x3)/3,(this._y2+2*this._y3)/3),this._context.lineTo((this._x3+2*this._x2)/3,(this._y3+2*this._y2)/3),this._context.closePath();break;case 3:this.point(this._x2,this._y2),this.point(this._x3,this._y3),this.point(this._x4,this._y4)}},point:function(t,e){switch(t=+t,e=+e,this._point){case 0:this._point=1,this._x2=t,this._y2=e;break;case 1:this._point=2,this._x3=t,this._y3=e;break;case 2:this._point=3,this._x4=t,this._y4=e,this._context.moveTo((this._x0+4*this._x1+t)/6,(this._y0+4*this._y1+e)/6);break;default:pt(this,t,e)}this._x0=this._x1,this._x1=t,this._y0=this._y1,this._y1=e}};var _t=function(t){return new gt(t)};function yt(t){this._context=t}yt.prototype={areaStart:function(){this._line=0},areaEnd:function(){this._line=NaN},lineStart:function(){this._x0=this._x1=this._y0=this._y1=NaN,this._point=0},lineEnd:function(){(this._line||0!==this._line&&3===this._point)&&this._context.closePath(),this._line=1-this._line},point:function(t,e){switch(t=+t,e=+e,this._point){case 0:this._point=1;break;case 1:this._point=2;break;case 2:this._point=3;var n=(this._x0+4*this._x1+t)/6,r=(this._y0+4*this._y1+e)/6;this._line?this._context.lineTo(n,r):this._context.moveTo(n,r);break;case 3:this._point=4;default:pt(this,t,e)}this._x0=this._x1,this._x1=t,this._y0=this._y1,this._y1=e}};var bt=function(t){return new yt(t)};function wt(t,e){this._basis=new mt(t),this._beta=e}wt.prototype={lineStart:function(){this._x=[],this._y=[],this._basis.lineStart()},lineEnd:function(){var t=this._x,e=this._y,n=t.length-1;if(n>0)for(var r,i=t[0],a=e[0],o=t[n]-i,u=e[n]-a,s=-1;++s<=n;)r=s/n,this._basis.point(this._beta*t[s]+(1-this._beta)*(i+r*o),this._beta*e[s]+(1-this._beta)*(a+r*u));this._x=this._y=null,this._basis.lineEnd()},point:function(t,e){this._x.push(+t),this._y.push(+e)}};var xt=function t(e){function n(t){return 1===e?new mt(t):new wt(t,e)}return n.beta=function(e){return t(+e)},n}(.85);function kt(t,e,n){t._context.bezierCurveTo(t._x1+t._k*(t._x2-t._x0),t._y1+t._k*(t._y2-t._y0),t._x2+t._k*(t._x1-e),t._y2+t._k*(t._y1-n),t._x2,t._y2)}function Mt(t,e){this._context=t,this._k=(1-e)/6}Mt.prototype={areaStart:function(){this._line=0},areaEnd:function(){this._line=NaN},lineStart:function(){this._x0=this._x1=this._x2=this._y0=this._y1=this._y2=NaN,this._point=0},lineEnd:function(){switch(this._point){case 2:this._context.lineTo(this._x2,this._y2);break;case 3:kt(this,this._x1,this._y1)}(this._line||0!==this._line&&1===this._point)&&this._context.closePath(),this._line=1-this._line},point:function(t,e){switch(t=+t,e=+e,this._point){case 0:this._point=1,this._line?this._context.lineTo(t,e):this._context.moveTo(t,e);break;case 1:this._point=2,this._x1=t,this._y1=e;break;case 2:this._point=3;default:kt(this,t,e)}this._x0=this._x1,this._x1=this._x2,this._x2=t,this._y0=this._y1,this._y1=this._y2,this._y2=e}};var Et=function t(e){function n(t){return new Mt(t,e)}return n.tension=function(e){return t(+e)},n}(0);function St(t,e){this._context=t,this._k=(1-e)/6}St.prototype={areaStart:ht,areaEnd:ht,lineStart:function(){this._x0=this._x1=this._x2=this._x3=this._x4=this._x5=this._y0=this._y1=this._y2=this._y3=this._y4=this._y5=NaN,this._point=0},lineEnd:function(){switch(this._point){case 1:this._context.moveTo(this._x3,this._y3),this._context.closePath();break;case 2:this._context.lineTo(this._x3,this._y3),this._context.closePath();break;case 3:this.point(this._x3,this._y3),this.point(this._x4,this._y4),this.point(this._x5,this._y5)}},point:function(t,e){switch(t=+t,e=+e,this._point){case 0:this._point=1,this._x3=t,this._y3=e;break;case 1:this._point=2,this._context.moveTo(this._x4=t,this._y4=e);break;case 2:this._point=3,this._x5=t,this._y5=e;break;default:kt(this,t,e)}this._x0=this._x1,this._x1=this._x2,this._x2=t,this._y0=this._y1,this._y1=this._y2,this._y2=e}};var Tt=function t(e){function n(t){return new St(t,e)}return n.tension=function(e){return t(+e)},n}(0);function jt(t,e){this._context=t,this._k=(1-e)/6}jt.prototype={areaStart:function(){this._line=0},areaEnd:function(){this._line=NaN},lineStart:function(){this._x0=this._x1=this._x2=this._y0=this._y1=this._y2=NaN,this._point=0},lineEnd:function(){(this._line||0!==this._line&&3===this._point)&&this._context.closePath(),this._line=1-this._line},point:function(t,e){switch(t=+t,e=+e,this._point){case 0:this._point=1;break;case 1:this._point=2;break;case 2:this._point=3,this._line?this._context.lineTo(this._x2,this._y2):this._context.moveTo(this._x2,this._y2);break;case 3:this._point=4;default:kt(this,t,e)}this._x0=this._x1,this._x1=this._x2,this._x2=t,this._y0=this._y1,this._y1=this._y2,this._y2=e}};var Ot=function t(e){function n(t){return new jt(t,e)}return n.tension=function(e){return t(+e)},n}(0);function At(t,e,n){var r=t._x1,i=t._y1,a=t._x2,o=t._y2;if(t._l01_a>1e-12){var u=2*t._l01_2a+3*t._l01_a*t._l12_a+t._l12_2a,s=3*t._l01_a*(t._l01_a+t._l12_a);r=(r*u-t._x0*t._l12_2a+t._x2*t._l01_2a)/s,i=(i*u-t._y0*t._l12_2a+t._y2*t._l01_2a)/s}if(t._l23_a>1e-12){var c=2*t._l23_2a+3*t._l23_a*t._l12_a+t._l12_2a,l=3*t._l23_a*(t._l23_a+t._l12_a);a=(a*c+t._x1*t._l23_2a-e*t._l12_2a)/l,o=(o*c+t._y1*t._l23_2a-n*t._l12_2a)/l}t._context.bezierCurveTo(r,i,a,o,t._x2,t._y2)}function Lt(t,e){this._context=t,this._alpha=e}Lt.prototype={areaStart:function(){this._line=0},areaEnd:function(){this._line=NaN},lineStart:function(){this._x0=this._x1=this._x2=this._y0=this._y1=this._y2=NaN,this._l01_a=this._l12_a=this._l23_a=this._l01_2a=this._l12_2a=this._l23_2a=this._point=0},lineEnd:function(){switch(this._point){case 2:this._context.lineTo(this._x2,this._y2);break;case 3:this.point(this._x2,this._y2)}(this._line||0!==this._line&&1===this._point)&&this._context.closePath(),this._line=1-this._line},point:function(t,e){if(t=+t,e=+e,this._point){var n=this._x2-t,r=this._y2-e;this._l23_a=Math.sqrt(this._l23_2a=Math.pow(n*n+r*r,this._alpha))}switch(this._point){case 0:this._point=1,this._line?this._context.lineTo(t,e):this._context.moveTo(t,e);break;case 1:this._point=2;break;case 2:this._point=3;default:At(this,t,e)}this._l01_a=this._l12_a,this._l12_a=this._l23_a,this._l01_2a=this._l12_2a,this._l12_2a=this._l23_2a,this._x0=this._x1,this._x1=this._x2,this._x2=t,this._y0=this._y1,this._y1=this._y2,this._y2=e}};var Ct=function t(e){function n(t){return e?new Lt(t,e):new Mt(t,0)}return n.alpha=function(e){return t(+e)},n}(.5);function Dt(t,e){this._context=t,this._alpha=e}Dt.prototype={areaStart:ht,areaEnd:ht,lineStart:function(){this._x0=this._x1=this._x2=this._x3=this._x4=this._x5=this._y0=this._y1=this._y2=this._y3=this._y4=this._y5=NaN,this._l01_a=this._l12_a=this._l23_a=this._l01_2a=this._l12_2a=this._l23_2a=this._point=0},lineEnd:function(){switch(this._point){case 1:this._context.moveTo(this._x3,this._y3),this._context.closePath();break;case 2:this._context.lineTo(this._x3,this._y3),this._context.closePath();break;case 3:this.point(this._x3,this._y3),this.point(this._x4,this._y4),this.point(this._x5,this._y5)}},point:function(t,e){if(t=+t,e=+e,this._point){var n=this._x2-t,r=this._y2-e;this._l23_a=Math.sqrt(this._l23_2a=Math.pow(n*n+r*r,this._alpha))}switch(this._point){case 0:this._point=1,this._x3=t,this._y3=e;break;case 1:this._point=2,this._context.moveTo(this._x4=t,this._y4=e);break;case 2:this._point=3,this._x5=t,this._y5=e;break;default:At(this,t,e)}this._l01_a=this._l12_a,this._l12_a=this._l23_a,this._l01_2a=this._l12_2a,this._l12_2a=this._l23_2a,this._x0=this._x1,this._x1=this._x2,this._x2=t,this._y0=this._y1,this._y1=this._y2,this._y2=e}};var Nt=function t(e){function n(t){return e?new Dt(t,e):new St(t,0)}return n.alpha=function(e){return t(+e)},n}(.5);function Pt(t,e){this._context=t,this._alpha=e}Pt.prototype={areaStart:function(){this._line=0},areaEnd:function(){this._line=NaN},lineStart:function(){this._x0=this._x1=this._x2=this._y0=this._y1=this._y2=NaN,this._l01_a=this._l12_a=this._l23_a=this._l01_2a=this._l12_2a=this._l23_2a=this._point=0},lineEnd:function(){(this._line||0!==this._line&&3===this._point)&&this._context.closePath(),this._line=1-this._line},point:function(t,e){if(t=+t,e=+e,this._point){var n=this._x2-t,r=this._y2-e;this._l23_a=Math.sqrt(this._l23_2a=Math.pow(n*n+r*r,this._alpha))}switch(this._point){case 0:this._point=1;break;case 1:this._point=2;break;case 2:this._point=3,this._line?this._context.lineTo(this._x2,this._y2):this._context.moveTo(this._x2,this._y2);break;case 3:this._point=4;default:At(this,t,e)}this._l01_a=this._l12_a,this._l12_a=this._l23_a,this._l01_2a=this._l12_2a,this._l12_2a=this._l23_2a,this._x0=this._x1,this._x1=this._x2,this._x2=t,this._y0=this._y1,this._y1=this._y2,this._y2=e}};var Rt=function t(e){function n(t){return e?new Pt(t,e):new jt(t,0)}return n.alpha=function(e){return t(+e)},n}(.5);function Yt(t){this._context=t}Yt.prototype={areaStart:ht,areaEnd:ht,lineStart:function(){this._point=0},lineEnd:function(){this._point&&this._context.closePath()},point:function(t,e){t=+t,e=+e,this._point?this._context.lineTo(t,e):(this._point=1,this._context.moveTo(t,e))}};var It=function(t){return new Yt(t)};function zt(t){return t<0?-1:1}function Ft(t,e,n){var r=t._x1-t._x0,i=e-t._x1,a=(t._y1-t._y0)/(r||i<0&&-0),o=(n-t._y1)/(i||r<0&&-0),u=(a*i+o*r)/(r+i);return(zt(a)+zt(o))*Math.min(Math.abs(a),Math.abs(o),.5*Math.abs(u))||0}function Ht(t,e){var n=t._x1-t._x0;return n?(3*(t._y1-t._y0)/n-e)/2:e}function Bt(t,e,n){var r=t._x0,i=t._y0,a=t._x1,o=t._y1,u=(a-r)/3;t._context.bezierCurveTo(r+u,i+u*e,a-u,o-u*n,a,o)}function qt(t){this._context=t}function Ut(t){this._context=new Wt(t)}function Wt(t){this._context=t}function Gt(t){return new qt(t)}function Vt(t){return new Ut(t)}function $t(t){this._context=t}function Xt(t){var e,n,r=t.length-1,i=new Array(r),a=new Array(r),o=new Array(r);for(i[0]=0,a[0]=2,o[0]=t[0]+2*t[1],e=1;e<r-1;++e)i[e]=1,a[e]=4,o[e]=4*t[e]+2*t[e+1];for(i[r-1]=2,a[r-1]=7,o[r-1]=8*t[r-1]+t[r],e=1;e<r;++e)n=i[e]/a[e-1],a[e]-=n,o[e]-=n*o[e-1];for(i[r-1]=o[r-1]/a[r-1],e=r-2;e>=0;--e)i[e]=(o[e]-i[e+1])/a[e];for(a[r-1]=(t[r]+i[r-1])/2,e=0;e<r-1;++e)a[e]=2*t[e+1]-i[e+1];return[i,a]}qt.prototype={areaStart:function(){this._line=0},areaEnd:function(){this._line=NaN},lineStart:function(){this._x0=this._x1=this._y0=this._y1=this._t0=NaN,this._point=0},lineEnd:function(){switch(this._point){case 2:this._context.lineTo(this._x1,this._y1);break;case 3:Bt(this,this._t0,Ht(this,this._t0))}(this._line||0!==this._line&&1===this._point)&&this._context.closePath(),this._line=1-this._line},point:function(t,e){var n=NaN;if(e=+e,(t=+t)!==this._x1||e!==this._y1){switch(this._point){case 0:this._point=1,this._line?this._context.lineTo(t,e):this._context.moveTo(t,e);break;case 1:this._point=2;break;case 2:this._point=3,Bt(this,Ht(this,n=Ft(this,t,e)),n);break;default:Bt(this,this._t0,n=Ft(this,t,e))}this._x0=this._x1,this._x1=t,this._y0=this._y1,this._y1=e,this._t0=n}}},(Ut.prototype=Object.create(qt.prototype)).point=function(t,e){qt.prototype.point.call(this,e,t)},Wt.prototype={moveTo:function(t,e){this._context.moveTo(e,t)},closePath:function(){this._context.closePath()},lineTo:function(t,e){this._context.lineTo(e,t)},bezierCurveTo:function(t,e,n,r,i,a){this._context.bezierCurveTo(e,t,r,n,a,i)}},$t.prototype={areaStart:function(){this._line=0},areaEnd:function(){this._line=NaN},lineStart:function(){this._x=[],this._y=[]},lineEnd:function(){var t=this._x,e=this._y,n=t.length;if(n)if(this._line?this._context.lineTo(t[0],e[0]):this._context.moveTo(t[0],e[0]),2===n)this._context.lineTo(t[1],e[1]);else for(var r=Xt(t),i=Xt(e),a=0,o=1;o<n;++a,++o)this._context.bezierCurveTo(r[0][a],i[0][a],r[1][a],i[1][a],t[o],e[o]);(this._line||0!==this._line&&1===n)&&this._context.closePath(),this._line=1-this._line,this._x=this._y=null},point:function(t,e){this._x.push(+t),this._y.push(+e)}};var Jt=function(t){return new $t(t)};function Zt(t,e){this._context=t,this._t=e}Zt.prototype={areaStart:function(){this._line=0},areaEnd:function(){this._line=NaN},lineStart:function(){this._x=this._y=NaN,this._point=0},lineEnd:function(){0<this._t&&this._t<1&&2===this._point&&this._context.lineTo(this._x,this._y),(this._line||0!==this._line&&1===this._point)&&this._context.closePath(),this._line>=0&&(this._t=1-this._t,this._line=1-this._line)},point:function(t,e){switch(t=+t,e=+e,this._point){case 0:this._point=1,this._line?this._context.lineTo(t,e):this._context.moveTo(t,e);break;case 1:this._point=2;default:if(this._t<=0)this._context.lineTo(this._x,e),this._context.lineTo(t,e);else{var n=this._x*(1-this._t)+t*this._t;this._context.lineTo(n,this._y),this._context.lineTo(n,e)}}this._x=t,this._y=e}};var Kt=function(t){return new Zt(t,.5)};function Qt(t){return new Zt(t,0)}function te(t){return new Zt(t,1)}var ee=function(t,e){if((i=t.length)>1)for(var n,r,i,a=1,o=t[e[0]],u=o.length;a<i;++a)for(r=o,o=t[e[a]],n=0;n<u;++n)o[n][1]+=o[n][0]=isNaN(r[n][1])?r[n][0]:r[n][1]},ne=function(t){for(var e=t.length,n=new Array(e);--e>=0;)n[e]=e;return n};function re(t,e){return t[e]}var ie=function(){var t=Object(i.a)([]),e=ne,n=ee,r=re;function a(i){var a,o,u=t.apply(this,arguments),s=i.length,c=u.length,l=new Array(c);for(a=0;a<c;++a){for(var f,d=u[a],h=l[a]=new Array(s),p=0;p<s;++p)h[p]=f=[0,+r(i[p],d,p,i)],f.data=i[p];h.key=d}for(a=0,o=e(l);a<c;++a)l[o[a]].index=a;return n(l,o),l}return a.keys=function(e){return arguments.length?(t="function"==typeof e?e:Object(i.a)(z.call(e)),a):t},a.value=function(t){return arguments.length?(r="function"==typeof t?t:Object(i.a)(+t),a):r},a.order=function(t){return arguments.length?(e=null==t?ne:"function"==typeof t?t:Object(i.a)(z.call(t)),a):e},a.offset=function(t){return arguments.length?(n=null==t?ee:t,a):n},a},ae=function(t,e){if((r=t.length)>0){for(var n,r,i,a=0,o=t[0].length;a<o;++a){for(i=n=0;n<r;++n)i+=t[n][a][1]||0;if(i)for(n=0;n<r;++n)t[n][a][1]/=i}ee(t,e)}},oe=function(t,e){if((u=t.length)>0)for(var n,r,i,a,o,u,s=0,c=t[e[0]].length;s<c;++s)for(a=o=0,n=0;n<u;++n)(i=(r=t[e[n]][s])[1]-r[0])>0?(r[0]=a,r[1]=a+=i):i<0?(r[1]=o,r[0]=o+=i):(r[0]=0,r[1]=i)},ue=function(t,e){if((n=t.length)>0){for(var n,r=0,i=t[e[0]],a=i.length;r<a;++r){for(var o=0,u=0;o<n;++o)u+=t[o][r][1]||0;i[r][1]+=i[r][0]=-u/2}ee(t,e)}},se=function(t,e){if((i=t.length)>0&&(r=(n=t[e[0]]).length)>0){for(var n,r,i,a=0,o=1;o<r;++o){for(var u=0,s=0,c=0;u<i;++u){for(var l=t[e[u]],f=l[o][1]||0,d=(f-(l[o-1][1]||0))/2,h=0;h<u;++h){var p=t[e[h]];d+=(p[o][1]||0)-(p[o-1][1]||0)}s+=f,c+=d*f}n[o-1][1]+=n[o-1][0]=a,s&&(a-=c/s)}n[o-1][1]+=n[o-1][0]=a,ee(t,e)}},ce=function(t){var e=t.map(le);return ne(t).sort((function(t,n){return e[t]-e[n]}))};function le(t){for(var e,n=-1,r=0,i=t.length,a=-1/0;++n<i;)(e=+t[n][1])>a&&(a=e,r=n);return r}var fe=function(t){var e=t.map(de);return ne(t).sort((function(t,n){return e[t]-e[n]}))};function de(t){for(var e,n=0,r=-1,i=t.length;++r<i;)(e=+t[r][1])&&(n+=e);return n}var he=function(t){return fe(t).reverse()},pe=function(t){var e,n,r=t.length,i=t.map(de),a=ce(t),o=0,u=0,s=[],c=[];for(e=0;e<r;++e)n=a[e],o<u?(o+=i[n],s.push(n)):(u+=i[n],c.push(n));return c.reverse().concat(s)},me=function(t){return ne(t).reverse()}},function(t,e,n){"use strict";n.d(e,"R",(function(){return r.a})),n.d(e,"M",(function(){return a})),n.d(e,"S",(function(){return o})),n.d(e,"Z",(function(){return u})),n.d(e,"ab",(function(){return s})),n.d(e,"bb",(function(){return c})),n.d(e,"ob",(function(){return l})),n.d(e,"pb",(function(){return f})),n.d(e,"qb",(function(){return d})),n.d(e,"sb",(function(){return h})),n.d(e,"b",(function(){return g})),n.d(e,"O",(function(){return v})),n.d(e,"o",(function(){return y})),n.d(e,"Y",(function(){return _})),n.d(e,"p",(function(){return w})),n.d(e,"cb",(function(){return b})),n.d(e,"t",(function(){return k})),n.d(e,"fb",(function(){return x})),n.d(e,"x",(function(){return E})),n.d(e,"ib",(function(){return M})),n.d(e,"y",(function(){return T})),n.d(e,"jb",(function(){return S})),n.d(e,"A",(function(){return O})),n.d(e,"lb",(function(){return j})),n.d(e,"B",(function(){return L})),n.d(e,"mb",(function(){return A})),n.d(e,"E",(function(){return D})),n.d(e,"rb",(function(){return C})),n.d(e,"c",(function(){return P})),n.d(e,"P",(function(){return N})),n.d(e,"d",(function(){return Y})),n.d(e,"Q",(function(){return R})),n.d(e,"h",(function(){return z})),n.d(e,"T",(function(){return I})),n.d(e,"m",(function(){return H})),n.d(e,"W",(function(){return F})),n.d(e,"s",(function(){return q})),n.d(e,"eb",(function(){return B})),n.d(e,"r",(function(){return W})),n.d(e,"db",(function(){return U})),n.d(e,"u",(function(){return V})),n.d(e,"gb",(function(){return G})),n.d(e,"z",(function(){return X})),n.d(e,"kb",(function(){return $})),n.d(e,"J",(function(){return Z})),n.d(e,"ub",(function(){return J})),n.d(e,"I",(function(){return Q})),n.d(e,"tb",(function(){return K})),n.d(e,"K",(function(){return et})),n.d(e,"vb",(function(){return tt})),n.d(e,"L",(function(){return rt})),n.d(e,"wb",(function(){return nt})),n.d(e,"a",(function(){return at})),n.d(e,"N",(function(){return it})),n.d(e,"i",(function(){return ut})),n.d(e,"U",(function(){return ot})),n.d(e,"j",(function(){return ct})),n.d(e,"V",(function(){return st})),n.d(e,"v",(function(){return ft})),n.d(e,"hb",(function(){return lt})),n.d(e,"C",(function(){return ht})),n.d(e,"nb",(function(){return dt})),n.d(e,"n",(function(){return mt})),n.d(e,"X",(function(){return pt})),n.d(e,"e",(function(){return vt})),n.d(e,"g",(function(){return yt})),n.d(e,"w",(function(){return kt})),n.d(e,"H",(function(){return bt})),n.d(e,"f",(function(){return wt})),n.d(e,"D",(function(){return jt})),n.d(e,"F",(function(){return Ot})),n.d(e,"G",(function(){return Lt})),n.d(e,"l",(function(){return Ct})),n.d(e,"k",(function(){return Dt})),n.d(e,"q",(function(){return Nt}));var r=n(261),i=n(3),a=Object(i.a)("7fc97fbeaed4fdc086ffff99386cb0f0027fbf5b17666666"),o=Object(i.a)("1b9e77d95f027570b3e7298a66a61ee6ab02a6761d666666"),u=Object(i.a)("a6cee31f78b4b2df8a33a02cfb9a99e31a1cfdbf6fff7f00cab2d66a3d9affff99b15928"),s=Object(i.a)("fbb4aeb3cde3ccebc5decbe4fed9a6ffffcce5d8bdfddaecf2f2f2"),c=Object(i.a)("b3e2cdfdcdaccbd5e8f4cae4e6f5c9fff2aef1e2cccccccc"),l=Object(i.a)("e41a1c377eb84daf4a984ea3ff7f00ffff33a65628f781bf999999"),f=Object(i.a)("66c2a5fc8d628da0cbe78ac3a6d854ffd92fe5c494b3b3b3"),d=Object(i.a)("8dd3c7ffffb3bebadafb807280b1d3fdb462b3de69fccde5d9d9d9bc80bdccebc5ffed6f"),h=Object(i.a)("4e79a7f28e2ce1575976b7b259a14fedc949af7aa1ff9da79c755fbab0ab"),p=n(78),m=function(t){return Object(p.b)(t[t.length-1])},v=new Array(3).concat("d8b365f5f5f55ab4ac","a6611adfc27d80cdc1018571","a6611adfc27df5f5f580cdc1018571","8c510ad8b365f6e8c3c7eae55ab4ac01665e","8c510ad8b365f6e8c3f5f5f5c7eae55ab4ac01665e","8c510abf812ddfc27df6e8c3c7eae580cdc135978f01665e","8c510abf812ddfc27df6e8c3f5f5f5c7eae580cdc135978f01665e","5430058c510abf812ddfc27df6e8c3c7eae580cdc135978f01665e003c30","5430058c510abf812ddfc27df6e8c3f5f5f5c7eae580cdc135978f01665e003c30").map(i.a),g=m(v),_=new Array(3).concat("af8dc3f7f7f77fbf7b","7b3294c2a5cfa6dba0008837","7b3294c2a5cff7f7f7a6dba0008837","762a83af8dc3e7d4e8d9f0d37fbf7b1b7837","762a83af8dc3e7d4e8f7f7f7d9f0d37fbf7b1b7837","762a839970abc2a5cfe7d4e8d9f0d3a6dba05aae611b7837","762a839970abc2a5cfe7d4e8f7f7f7d9f0d3a6dba05aae611b7837","40004b762a839970abc2a5cfe7d4e8d9f0d3a6dba05aae611b783700441b","40004b762a839970abc2a5cfe7d4e8f7f7f7d9f0d3a6dba05aae611b783700441b").map(i.a),y=m(_),b=new Array(3).concat("e9a3c9f7f7f7a1d76a","d01c8bf1b6dab8e1864dac26","d01c8bf1b6daf7f7f7b8e1864dac26","c51b7de9a3c9fde0efe6f5d0a1d76a4d9221","c51b7de9a3c9fde0eff7f7f7e6f5d0a1d76a4d9221","c51b7dde77aef1b6dafde0efe6f5d0b8e1867fbc414d9221","c51b7dde77aef1b6dafde0eff7f7f7e6f5d0b8e1867fbc414d9221","8e0152c51b7dde77aef1b6dafde0efe6f5d0b8e1867fbc414d9221276419","8e0152c51b7dde77aef1b6dafde0eff7f7f7e6f5d0b8e1867fbc414d9221276419").map(i.a),w=m(b),x=new Array(3).concat("998ec3f7f7f7f1a340","5e3c99b2abd2fdb863e66101","5e3c99b2abd2f7f7f7fdb863e66101","542788998ec3d8daebfee0b6f1a340b35806","542788998ec3d8daebf7f7f7fee0b6f1a340b35806","5427888073acb2abd2d8daebfee0b6fdb863e08214b35806","5427888073acb2abd2d8daebf7f7f7fee0b6fdb863e08214b35806","2d004b5427888073acb2abd2d8daebfee0b6fdb863e08214b358067f3b08","2d004b5427888073acb2abd2d8daebf7f7f7fee0b6fdb863e08214b358067f3b08").map(i.a),k=m(x),M=new Array(3).concat("ef8a62f7f7f767a9cf","ca0020f4a58292c5de0571b0","ca0020f4a582f7f7f792c5de0571b0","b2182bef8a62fddbc7d1e5f067a9cf2166ac","b2182bef8a62fddbc7f7f7f7d1e5f067a9cf2166ac","b2182bd6604df4a582fddbc7d1e5f092c5de4393c32166ac","b2182bd6604df4a582fddbc7f7f7f7d1e5f092c5de4393c32166ac","67001fb2182bd6604df4a582fddbc7d1e5f092c5de4393c32166ac053061","67001fb2182bd6604df4a582fddbc7f7f7f7d1e5f092c5de4393c32166ac053061").map(i.a),E=m(M),S=new Array(3).concat("ef8a62ffffff999999","ca0020f4a582bababa404040","ca0020f4a582ffffffbababa404040","b2182bef8a62fddbc7e0e0e09999994d4d4d","b2182bef8a62fddbc7ffffffe0e0e09999994d4d4d","b2182bd6604df4a582fddbc7e0e0e0bababa8787874d4d4d","b2182bd6604df4a582fddbc7ffffffe0e0e0bababa8787874d4d4d","67001fb2182bd6604df4a582fddbc7e0e0e0bababa8787874d4d4d1a1a1a","67001fb2182bd6604df4a582fddbc7ffffffe0e0e0bababa8787874d4d4d1a1a1a").map(i.a),T=m(S),j=new Array(3).concat("fc8d59ffffbf91bfdb","d7191cfdae61abd9e92c7bb6","d7191cfdae61ffffbfabd9e92c7bb6","d73027fc8d59fee090e0f3f891bfdb4575b4","d73027fc8d59fee090ffffbfe0f3f891bfdb4575b4","d73027f46d43fdae61fee090e0f3f8abd9e974add14575b4","d73027f46d43fdae61fee090ffffbfe0f3f8abd9e974add14575b4","a50026d73027f46d43fdae61fee090e0f3f8abd9e974add14575b4313695","a50026d73027f46d43fdae61fee090ffffbfe0f3f8abd9e974add14575b4313695").map(i.a),O=m(j),A=new Array(3).concat("fc8d59ffffbf91cf60","d7191cfdae61a6d96a1a9641","d7191cfdae61ffffbfa6d96a1a9641","d73027fc8d59fee08bd9ef8b91cf601a9850","d73027fc8d59fee08bffffbfd9ef8b91cf601a9850","d73027f46d43fdae61fee08bd9ef8ba6d96a66bd631a9850","d73027f46d43fdae61fee08bffffbfd9ef8ba6d96a66bd631a9850","a50026d73027f46d43fdae61fee08bd9ef8ba6d96a66bd631a9850006837","a50026d73027f46d43fdae61fee08bffffbfd9ef8ba6d96a66bd631a9850006837").map(i.a),L=m(A),C=new Array(3).concat("fc8d59ffffbf99d594","d7191cfdae61abdda42b83ba","d7191cfdae61ffffbfabdda42b83ba","d53e4ffc8d59fee08be6f59899d5943288bd","d53e4ffc8d59fee08bffffbfe6f59899d5943288bd","d53e4ff46d43fdae61fee08be6f598abdda466c2a53288bd","d53e4ff46d43fdae61fee08bffffbfe6f598abdda466c2a53288bd","9e0142d53e4ff46d43fdae61fee08be6f598abdda466c2a53288bd5e4fa2","9e0142d53e4ff46d43fdae61fee08bffffbfe6f598abdda466c2a53288bd5e4fa2").map(i.a),D=m(C),N=new Array(3).concat("e5f5f999d8c92ca25f","edf8fbb2e2e266c2a4238b45","edf8fbb2e2e266c2a42ca25f006d2c","edf8fbccece699d8c966c2a42ca25f006d2c","edf8fbccece699d8c966c2a441ae76238b45005824","f7fcfde5f5f9ccece699d8c966c2a441ae76238b45005824","f7fcfde5f5f9ccece699d8c966c2a441ae76238b45006d2c00441b").map(i.a),P=m(N),R=new Array(3).concat("e0ecf49ebcda8856a7","edf8fbb3cde38c96c688419d","edf8fbb3cde38c96c68856a7810f7c","edf8fbbfd3e69ebcda8c96c68856a7810f7c","edf8fbbfd3e69ebcda8c96c68c6bb188419d6e016b","f7fcfde0ecf4bfd3e69ebcda8c96c68c6bb188419d6e016b","f7fcfde0ecf4bfd3e69ebcda8c96c68c6bb188419d810f7c4d004b").map(i.a),Y=m(R),I=new Array(3).concat("e0f3dba8ddb543a2ca","f0f9e8bae4bc7bccc42b8cbe","f0f9e8bae4bc7bccc443a2ca0868ac","f0f9e8ccebc5a8ddb57bccc443a2ca0868ac","f0f9e8ccebc5a8ddb57bccc44eb3d32b8cbe08589e","f7fcf0e0f3dbccebc5a8ddb57bccc44eb3d32b8cbe08589e","f7fcf0e0f3dbccebc5a8ddb57bccc44eb3d32b8cbe0868ac084081").map(i.a),z=m(I),F=new Array(3).concat("fee8c8fdbb84e34a33","fef0d9fdcc8afc8d59d7301f","fef0d9fdcc8afc8d59e34a33b30000","fef0d9fdd49efdbb84fc8d59e34a33b30000","fef0d9fdd49efdbb84fc8d59ef6548d7301f990000","fff7ecfee8c8fdd49efdbb84fc8d59ef6548d7301f990000","fff7ecfee8c8fdd49efdbb84fc8d59ef6548d7301fb300007f0000").map(i.a),H=m(F),B=new Array(3).concat("ece2f0a6bddb1c9099","f6eff7bdc9e167a9cf02818a","f6eff7bdc9e167a9cf1c9099016c59","f6eff7d0d1e6a6bddb67a9cf1c9099016c59","f6eff7d0d1e6a6bddb67a9cf3690c002818a016450","fff7fbece2f0d0d1e6a6bddb67a9cf3690c002818a016450","fff7fbece2f0d0d1e6a6bddb67a9cf3690c002818a016c59014636").map(i.a),q=m(B),U=new Array(3).concat("ece7f2a6bddb2b8cbe","f1eef6bdc9e174a9cf0570b0","f1eef6bdc9e174a9cf2b8cbe045a8d","f1eef6d0d1e6a6bddb74a9cf2b8cbe045a8d","f1eef6d0d1e6a6bddb74a9cf3690c00570b0034e7b","fff7fbece7f2d0d1e6a6bddb74a9cf3690c00570b0034e7b","fff7fbece7f2d0d1e6a6bddb74a9cf3690c00570b0045a8d023858").map(i.a),W=m(U),G=new Array(3).concat("e7e1efc994c7dd1c77","f1eef6d7b5d8df65b0ce1256","f1eef6d7b5d8df65b0dd1c77980043","f1eef6d4b9dac994c7df65b0dd1c77980043","f1eef6d4b9dac994c7df65b0e7298ace125691003f","f7f4f9e7e1efd4b9dac994c7df65b0e7298ace125691003f","f7f4f9e7e1efd4b9dac994c7df65b0e7298ace125698004367001f").map(i.a),V=m(G),$=new Array(3).concat("fde0ddfa9fb5c51b8a","feebe2fbb4b9f768a1ae017e","feebe2fbb4b9f768a1c51b8a7a0177","feebe2fcc5c0fa9fb5f768a1c51b8a7a0177","feebe2fcc5c0fa9fb5f768a1dd3497ae017e7a0177","fff7f3fde0ddfcc5c0fa9fb5f768a1dd3497ae017e7a0177","fff7f3fde0ddfcc5c0fa9fb5f768a1dd3497ae017e7a017749006a").map(i.a),X=m($),J=new Array(3).concat("edf8b17fcdbb2c7fb8","ffffcca1dab441b6c4225ea8","ffffcca1dab441b6c42c7fb8253494","ffffccc7e9b47fcdbb41b6c42c7fb8253494","ffffccc7e9b47fcdbb41b6c41d91c0225ea80c2c84","ffffd9edf8b1c7e9b47fcdbb41b6c41d91c0225ea80c2c84","ffffd9edf8b1c7e9b47fcdbb41b6c41d91c0225ea8253494081d58").map(i.a),Z=m(J),K=new Array(3).concat("f7fcb9addd8e31a354","ffffccc2e69978c679238443","ffffccc2e69978c67931a354006837","ffffccd9f0a3addd8e78c67931a354006837","ffffccd9f0a3addd8e78c67941ab5d238443005a32","ffffe5f7fcb9d9f0a3addd8e78c67941ab5d238443005a32","ffffe5f7fcb9d9f0a3addd8e78c67941ab5d238443006837004529").map(i.a),Q=m(K),tt=new Array(3).concat("fff7bcfec44fd95f0e","ffffd4fed98efe9929cc4c02","ffffd4fed98efe9929d95f0e993404","ffffd4fee391fec44ffe9929d95f0e993404","ffffd4fee391fec44ffe9929ec7014cc4c028c2d04","ffffe5fff7bcfee391fec44ffe9929ec7014cc4c028c2d04","ffffe5fff7bcfee391fec44ffe9929ec7014cc4c02993404662506").map(i.a),et=m(tt),nt=new Array(3).concat("ffeda0feb24cf03b20","ffffb2fecc5cfd8d3ce31a1c","ffffb2fecc5cfd8d3cf03b20bd0026","ffffb2fed976feb24cfd8d3cf03b20bd0026","ffffb2fed976feb24cfd8d3cfc4e2ae31a1cb10026","ffffccffeda0fed976feb24cfd8d3cfc4e2ae31a1cb10026","ffffccffeda0fed976feb24cfd8d3cfc4e2ae31a1cbd0026800026").map(i.a),rt=m(nt),it=new Array(3).concat("deebf79ecae13182bd","eff3ffbdd7e76baed62171b5","eff3ffbdd7e76baed63182bd08519c","eff3ffc6dbef9ecae16baed63182bd08519c","eff3ffc6dbef9ecae16baed64292c62171b5084594","f7fbffdeebf7c6dbef9ecae16baed64292c62171b5084594","f7fbffdeebf7c6dbef9ecae16baed64292c62171b508519c08306b").map(i.a),at=m(it),ot=new Array(3).concat("e5f5e0a1d99b31a354","edf8e9bae4b374c476238b45","edf8e9bae4b374c47631a354006d2c","edf8e9c7e9c0a1d99b74c47631a354006d2c","edf8e9c7e9c0a1d99b74c47641ab5d238b45005a32","f7fcf5e5f5e0c7e9c0a1d99b74c47641ab5d238b45005a32","f7fcf5e5f5e0c7e9c0a1d99b74c47641ab5d238b45006d2c00441b").map(i.a),ut=m(ot),st=new Array(3).concat("f0f0f0bdbdbd636363","f7f7f7cccccc969696525252","f7f7f7cccccc969696636363252525","f7f7f7d9d9d9bdbdbd969696636363252525","f7f7f7d9d9d9bdbdbd969696737373525252252525","fffffff0f0f0d9d9d9bdbdbd969696737373525252252525","fffffff0f0f0d9d9d9bdbdbd969696737373525252252525000000").map(i.a),ct=m(st),lt=new Array(3).concat("efedf5bcbddc756bb1","f2f0f7cbc9e29e9ac86a51a3","f2f0f7cbc9e29e9ac8756bb154278f","f2f0f7dadaebbcbddc9e9ac8756bb154278f","f2f0f7dadaebbcbddc9e9ac8807dba6a51a34a1486","fcfbfdefedf5dadaebbcbddc9e9ac8807dba6a51a34a1486","fcfbfdefedf5dadaebbcbddc9e9ac8807dba6a51a354278f3f007d").map(i.a),ft=m(lt),dt=new Array(3).concat("fee0d2fc9272de2d26","fee5d9fcae91fb6a4acb181d","fee5d9fcae91fb6a4ade2d26a50f15","fee5d9fcbba1fc9272fb6a4ade2d26a50f15","fee5d9fcbba1fc9272fb6a4aef3b2ccb181d99000d","fff5f0fee0d2fcbba1fc9272fb6a4aef3b2ccb181d99000d","fff5f0fee0d2fcbba1fc9272fb6a4aef3b2ccb181da50f1567000d").map(i.a),ht=m(dt),pt=new Array(3).concat("fee6cefdae6be6550d","feeddefdbe85fd8d3cd94701","feeddefdbe85fd8d3ce6550da63603","feeddefdd0a2fdae6bfd8d3ce6550da63603","feeddefdd0a2fdae6bfd8d3cf16913d948018c2d04","fff5ebfee6cefdd0a2fdae6bfd8d3cf16913d948018c2d04","fff5ebfee6cefdd0a2fdae6bfd8d3cf16913d94801a636037f2704").map(i.a),mt=m(pt),vt=function(t){return t=Math.max(0,Math.min(1,t)),"rgb("+Math.max(0,Math.min(255,Math.round(-4.54-t*(35.34-t*(2381.73-t*(6402.7-t*(7024.72-2710.57*t)))))))+", "+Math.max(0,Math.min(255,Math.round(32.49+t*(170.73+t*(52.82-t*(131.46-t*(176.58-67.37*t)))))))+", "+Math.max(0,Math.min(255,Math.round(81.24+t*(442.36-t*(2482.43-t*(6167.24-t*(6614.94-2475.67*t)))))))+")"},gt=n(171),_t=n(179),yt=Object(_t.a)(Object(gt.a)(300,.5,0),Object(gt.a)(-240,.5,1)),bt=Object(_t.a)(Object(gt.a)(-100,.75,.35),Object(gt.a)(80,1.5,.8)),wt=Object(_t.a)(Object(gt.a)(260,.75,.35),Object(gt.a)(80,1.5,.8)),xt=Object(gt.a)(),kt=function(t){(t<0||t>1)&&(t-=Math.floor(t));var e=Math.abs(t-.5);return xt.h=360*t-100,xt.s=1.5-1.5*e,xt.l=.8-.9*e,xt+""},Mt=n(14),Et=Object(Mt.g)(),St=Math.PI/3,Tt=2*Math.PI/3,jt=function(t){var e;return t=(.5-t)*Math.PI,Et.r=255*(e=Math.sin(t))*e,Et.g=255*(e=Math.sin(t+St))*e,Et.b=255*(e=Math.sin(t+Tt))*e,Et+""},Ot=function(t){return t=Math.max(0,Math.min(1,t)),"rgb("+Math.max(0,Math.min(255,Math.round(34.61+t*(1172.33-t*(10793.56-t*(33300.12-t*(38394.49-14825.05*t)))))))+", "+Math.max(0,Math.min(255,Math.round(23.31+t*(557.33+t*(1225.33-t*(3574.96-t*(1073.77+707.56*t)))))))+", "+Math.max(0,Math.min(255,Math.round(27.2+t*(3211.1-t*(15327.97-t*(27814-t*(22569.18-6838.66*t)))))))+")"};function At(t){var e=t.length;return function(n){return t[Math.max(0,Math.min(e-1,Math.floor(n*e)))]}}var Lt=At(Object(i.a)("44015444025645045745055946075a46085c460a5d460b5e470d60470e6147106347116447136548146748166848176948186a481a6c481b6d481c6e481d6f481f70482071482173482374482475482576482677482878482979472a7a472c7a472d7b472e7c472f7d46307e46327e46337f463480453581453781453882443983443a83443b84433d84433e85423f854240864241864142874144874045884046883f47883f48893e49893e4a893e4c8a3d4d8a3d4e8a3c4f8a3c508b3b518b3b528b3a538b3a548c39558c39568c38588c38598c375a8c375b8d365c8d365d8d355e8d355f8d34608d34618d33628d33638d32648e32658e31668e31678e31688e30698e306a8e2f6b8e2f6c8e2e6d8e2e6e8e2e6f8e2d708e2d718e2c718e2c728e2c738e2b748e2b758e2a768e2a778e2a788e29798e297a8e297b8e287c8e287d8e277e8e277f8e27808e26818e26828e26828e25838e25848e25858e24868e24878e23888e23898e238a8d228b8d228c8d228d8d218e8d218f8d21908d21918c20928c20928c20938c1f948c1f958b1f968b1f978b1f988b1f998a1f9a8a1e9b8a1e9c891e9d891f9e891f9f881fa0881fa1881fa1871fa28720a38620a48621a58521a68522a78522a88423a98324aa8325ab8225ac8226ad8127ad8128ae8029af7f2ab07f2cb17e2db27d2eb37c2fb47c31b57b32b67a34b67935b77937b87838b9773aba763bbb753dbc743fbc7340bd7242be7144bf7046c06f48c16e4ac16d4cc26c4ec36b50c46a52c56954c56856c66758c7655ac8645cc8635ec96260ca6063cb5f65cb5e67cc5c69cd5b6ccd5a6ece5870cf5773d05675d05477d1537ad1517cd2507fd34e81d34d84d44b86d54989d5488bd6468ed64590d74393d74195d84098d83e9bd93c9dd93ba0da39a2da37a5db36a8db34aadc32addc30b0dd2fb2dd2db5de2bb8de29bade28bddf26c0df25c2df23c5e021c8e020cae11fcde11dd0e11cd2e21bd5e21ad8e219dae319dde318dfe318e2e418e5e419e7e419eae51aece51befe51cf1e51df4e61ef6e620f8e621fbe723fde725")),Ct=At(Object(i.a)("00000401000501010601010802010902020b02020d03030f03031204041405041606051806051a07061c08071e0907200a08220b09240c09260d0a290e0b2b100b2d110c2f120d31130d34140e36150e38160f3b180f3d19103f1a10421c10441d11471e114920114b21114e22115024125325125527125829115a2a115c2c115f2d11612f116331116533106734106936106b38106c390f6e3b0f703d0f713f0f72400f74420f75440f764510774710784910784a10794c117a4e117b4f127b51127c52137c54137d56147d57157e59157e5a167e5c167f5d177f5f187f601880621980641a80651a80671b80681c816a1c816b1d816d1d816e1e81701f81721f817320817521817621817822817922827b23827c23827e24828025828125818326818426818627818827818928818b29818c29818e2a81902a81912b81932b80942c80962c80982d80992d809b2e7f9c2e7f9e2f7fa02f7fa1307ea3307ea5317ea6317da8327daa337dab337cad347cae347bb0357bb2357bb3367ab5367ab73779b83779ba3878bc3978bd3977bf3a77c03a76c23b75c43c75c53c74c73d73c83e73ca3e72cc3f71cd4071cf4070d0416fd2426fd3436ed5446dd6456cd8456cd9466bdb476adc4869de4968df4a68e04c67e24d66e34e65e44f64e55064e75263e85362e95462ea5661eb5760ec5860ed5a5fee5b5eef5d5ef05f5ef1605df2625df2645cf3655cf4675cf4695cf56b5cf66c5cf66e5cf7705cf7725cf8745cf8765cf9785df9795df97b5dfa7d5efa7f5efa815ffb835ffb8560fb8761fc8961fc8a62fc8c63fc8e64fc9065fd9266fd9467fd9668fd9869fd9a6afd9b6bfe9d6cfe9f6dfea16efea36ffea571fea772fea973feaa74feac76feae77feb078feb27afeb47bfeb67cfeb77efeb97ffebb81febd82febf84fec185fec287fec488fec68afec88cfeca8dfecc8ffecd90fecf92fed194fed395fed597fed799fed89afdda9cfddc9efddea0fde0a1fde2a3fde3a5fde5a7fde7a9fde9aafdebacfcecaefceeb0fcf0b2fcf2b4fcf4b6fcf6b8fcf7b9fcf9bbfcfbbdfcfdbf")),Dt=At(Object(i.a)("00000401000501010601010802010a02020c02020e03021004031204031405041706041907051b08051d09061f0a07220b07240c08260d08290e092b10092d110a30120a32140b34150b37160b39180c3c190c3e1b0c411c0c431e0c451f0c48210c4a230c4c240c4f260c51280b53290b552b0b572d0b592f0a5b310a5c320a5e340a5f3609613809623909633b09643d09653e0966400a67420a68440a68450a69470b6a490b6a4a0c6b4c0c6b4d0d6c4f0d6c510e6c520e6d540f6d550f6d57106e59106e5a116e5c126e5d126e5f136e61136e62146e64156e65156e67166e69166e6a176e6c186e6d186e6f196e71196e721a6e741a6e751b6e771c6d781c6d7a1d6d7c1d6d7d1e6d7f1e6c801f6c82206c84206b85216b87216b88226a8a226a8c23698d23698f24699025689225689326679526679727669827669a28659b29649d29649f2a63a02a63a22b62a32c61a52c60a62d60a82e5fa92e5eab2f5ead305dae305cb0315bb1325ab3325ab43359b63458b73557b93556ba3655bc3754bd3853bf3952c03a51c13a50c33b4fc43c4ec63d4dc73e4cc83f4bca404acb4149cc4248ce4347cf4446d04545d24644d34743d44842d54a41d74b3fd84c3ed94d3dda4e3cdb503bdd513ade5238df5337e05536e15635e25734e35933e45a31e55c30e65d2fe75e2ee8602de9612bea632aeb6429eb6628ec6726ed6925ee6a24ef6c23ef6e21f06f20f1711ff1731df2741cf3761bf37819f47918f57b17f57d15f67e14f68013f78212f78410f8850ff8870ef8890cf98b0bf98c0af98e09fa9008fa9207fa9407fb9606fb9706fb9906fb9b06fb9d07fc9f07fca108fca309fca50afca60cfca80dfcaa0ffcac11fcae12fcb014fcb216fcb418fbb61afbb81dfbba1ffbbc21fbbe23fac026fac228fac42afac62df9c72ff9c932f9cb35f8cd37f8cf3af7d13df7d340f6d543f6d746f5d949f5db4cf4dd4ff4df53f4e156f3e35af3e55df2e661f2e865f2ea69f1ec6df1ed71f1ef75f1f179f2f27df2f482f3f586f3f68af4f88ef5f992f6fa96f8fb9af9fc9dfafda1fcffa4")),Nt=At(Object(i.a)("0d088710078813078916078a19068c1b068d1d068e20068f2206902406912605912805922a05932c05942e05952f059631059733059735049837049938049a3a049a3c049b3e049c3f049c41049d43039e44039e46039f48039f4903a04b03a14c02a14e02a25002a25102a35302a35502a45601a45801a45901a55b01a55c01a65e01a66001a66100a76300a76400a76600a76700a86900a86a00a86c00a86e00a86f00a87100a87201a87401a87501a87701a87801a87a02a87b02a87d03a87e03a88004a88104a78305a78405a78606a68707a68808a68a09a58b0aa58d0ba58e0ca48f0da4910ea3920fa39410a29511a19613a19814a099159f9a169f9c179e9d189d9e199da01a9ca11b9ba21d9aa31e9aa51f99a62098a72197a82296aa2395ab2494ac2694ad2793ae2892b02991b12a90b22b8fb32c8eb42e8db52f8cb6308bb7318ab83289ba3388bb3488bc3587bd3786be3885bf3984c03a83c13b82c23c81c33d80c43e7fc5407ec6417dc7427cc8437bc9447aca457acb4679cc4778cc4977cd4a76ce4b75cf4c74d04d73d14e72d24f71d35171d45270d5536fd5546ed6556dd7566cd8576bd9586ada5a6ada5b69db5c68dc5d67dd5e66de5f65de6164df6263e06363e16462e26561e26660e3685fe4695ee56a5de56b5de66c5ce76e5be76f5ae87059e97158e97257ea7457eb7556eb7655ec7754ed7953ed7a52ee7b51ef7c51ef7e50f07f4ff0804ef1814df1834cf2844bf3854bf3874af48849f48948f58b47f58c46f68d45f68f44f79044f79143f79342f89441f89540f9973ff9983ef99a3efa9b3dfa9c3cfa9e3bfb9f3afba139fba238fca338fca537fca636fca835fca934fdab33fdac33fdae32fdaf31fdb130fdb22ffdb42ffdb52efeb72dfeb82cfeba2cfebb2bfebd2afebe2afec029fdc229fdc328fdc527fdc627fdc827fdca26fdcb26fccd25fcce25fcd025fcd225fbd324fbd524fbd724fad824fada24f9dc24f9dd25f8df25f8e125f7e225f7e425f6e626f6e826f5e926f5eb27f4ed27f3ee27f3f027f2f227f1f426f1f525f0f724f0f921"))},function(t,e,n){"use strict";function r(t){return+t}function i(t){return t*t}function a(t){return t*(2-t)}function o(t){return((t*=2)<=1?t*t:--t*(2-t)+1)/2}n.d(e,"y",(function(){return r})),n.d(e,"D",(function(){return o})),n.d(e,"E",(function(){return i})),n.d(e,"G",(function(){return a})),n.d(e,"F",(function(){return o})),n.d(e,"m",(function(){return u.b})),n.d(e,"n",(function(){return u.a})),n.d(e,"p",(function(){return u.c})),n.d(e,"o",(function(){return u.b})),n.d(e,"z",(function(){return l})),n.d(e,"A",(function(){return s})),n.d(e,"C",(function(){return c})),n.d(e,"B",(function(){return l})),n.d(e,"H",(function(){return m})),n.d(e,"I",(function(){return h})),n.d(e,"K",(function(){return p})),n.d(e,"J",(function(){return m})),n.d(e,"u",(function(){return _})),n.d(e,"v",(function(){return v})),n.d(e,"x",(function(){return g})),n.d(e,"w",(function(){return _})),n.d(e,"i",(function(){return w})),n.d(e,"j",(function(){return y})),n.d(e,"l",(function(){return b})),n.d(e,"k",(function(){return w})),n.d(e,"e",(function(){return M})),n.d(e,"f",(function(){return k})),n.d(e,"h",(function(){return M})),n.d(e,"g",(function(){return E})),n.d(e,"a",(function(){return j})),n.d(e,"b",(function(){return S})),n.d(e,"d",(function(){return T})),n.d(e,"c",(function(){return j})),n.d(e,"q",(function(){return L})),n.d(e,"r",(function(){return A})),n.d(e,"t",(function(){return L})),n.d(e,"s",(function(){return C}));var u=n(110),s=function t(e){function n(t){return Math.pow(t,e)}return e=+e,n.exponent=t,n}(3),c=function t(e){function n(t){return 1-Math.pow(1-t,e)}return e=+e,n.exponent=t,n}(3),l=function t(e){function n(t){return((t*=2)<=1?Math.pow(t,e):2-Math.pow(2-t,e))/2}return e=+e,n.exponent=t,n}(3),f=Math.PI,d=f/2;function h(t){return 1-Math.cos(t*d)}function p(t){return Math.sin(t*d)}function m(t){return(1-Math.cos(f*t))/2}function v(t){return Math.pow(2,10*t-10)}function g(t){return 1-Math.pow(2,-10*t)}function _(t){return((t*=2)<=1?Math.pow(2,10*t-10):2-Math.pow(2,10-10*t))/2}function y(t){return 1-Math.sqrt(1-t*t)}function b(t){return Math.sqrt(1- --t*t)}function w(t){return((t*=2)<=1?1-Math.sqrt(1-t*t):Math.sqrt(1-(t-=2)*t)+1)/2}var x=7.5625;function k(t){return 1-M(1-t)}function M(t){return(t=+t)<4/11?x*t*t:t<8/11?x*(t-=6/11)*t+3/4:t<10/11?x*(t-=9/11)*t+15/16:x*(t-=21/22)*t+63/64}function E(t){return((t*=2)<=1?1-M(1-t):M(t-1)+1)/2}var S=function t(e){function n(t){return t*t*((e+1)*t-e)}return e=+e,n.overshoot=t,n}(1.70158),T=function t(e){function n(t){return--t*t*((e+1)*t+e)+1}return e=+e,n.overshoot=t,n}(1.70158),j=function t(e){function n(t){return((t*=2)<1?t*t*((e+1)*t-e):(t-=2)*t*((e+1)*t+e)+2)/2}return e=+e,n.overshoot=t,n}(1.70158),O=2*Math.PI,A=function t(e,n){var r=Math.asin(1/(e=Math.max(1,e)))*(n/=O);function i(t){return e*Math.pow(2,10*--t)*Math.sin((r-t)/n)}return i.amplitude=function(e){return t(e,n*O)},i.period=function(n){return t(e,n)},i}(1,.3),L=function t(e,n){var r=Math.asin(1/(e=Math.max(1,e)))*(n/=O);function i(t){return 1-e*Math.pow(2,-10*(t=+t))*Math.sin((t+r)/n)}return i.amplitude=function(e){return t(e,n*O)},i.period=function(n){return t(e,n)},i}(1,.3),C=function t(e,n){var r=Math.asin(1/(e=Math.max(1,e)))*(n/=O);function i(t){return((t=2*t-1)<0?e*Math.pow(2,10*t)*Math.sin((r-t)/n):2-e*Math.pow(2,-10*t)*Math.sin((r+t)/n))/2}return i.amplitude=function(e){return t(e,n*O)},i.period=function(n){return t(e,n)},i}(1,.3)},function(t,e,n){"use strict";n.d(e,"i",(function(){return r.a})),n.d(e,"g",(function(){return i.f})),n.d(e,"h",(function(){return i.g})),n.d(e,"b",(function(){return i.a})),n.d(e,"c",(function(){return i.b})),n.d(e,"e",(function(){return i.d})),n.d(e,"d",(function(){return i.c})),n.d(e,"f",(function(){return i.e})),n.d(e,"o",(function(){return a.f})),n.d(e,"p",(function(){return a.g})),n.d(e,"j",(function(){return a.a})),n.d(e,"k",(function(){return a.b})),n.d(e,"m",(function(){return a.d})),n.d(e,"l",(function(){return a.c})),n.d(e,"n",(function(){return a.e})),n.d(e,"a",(function(){return o}));var r=n(100),i=n(51),a=n(52);function o(t){for(var e in t){var n,r,i=t[e].trim();if(i)if("true"===i)i=!0;else if("false"===i)i=!1;else if("NaN"===i)i=NaN;else if(isNaN(n=+i)){if(!(r=i.match(/^([-+]\d{2})?\d{4}(-\d{2}(-\d{2})?)?(T\d{2}:\d{2}(:\d{2}(\.\d{3})?)?(Z|[-+]\d{2}:\d{2})?)?$/)))continue;u&&r[4]&&!r[7]&&(i=i.replace(/-/g,"/").replace(/T/," ")),i=new Date(i)}else i=n;else i=null;t[e]=i}return t}var u=new Date("2019-01-01T00:00").getHours()||new Date("2019-07-01T00:00").getHours()},function(t,e,n){"use strict";n.d(e,"a",(function(){return o}));var r=n(7),i=n(9),a=Object(r.a)((function(t){t.setHours(0,0,0,0)}),(function(t,e){t.setDate(t.getDate()+e)}),(function(t,e){return(e-t-(e.getTimezoneOffset()-t.getTimezoneOffset())*i.c)/i.a}),(function(t){return t.getDate()-1}));e.b=a;var o=a.range},function(t,e,n){"use strict";n.d(e,"b",(function(){return o}));var r=n(7),i=n(9),a=Object(r.a)((function(t){t.setUTCHours(0,0,0,0)}),(function(t,e){t.setUTCDate(t.getUTCDate()+e)}),(function(t,e){return(e-t)/i.a}),(function(t){return t.getUTCDate()-1}));e.a=a;var o=a.range},function(t,e,n){function r(){var t;try{t=e.storage.debug}catch(t){}return t}(e=t.exports=n(475)).log=function(){return"object"==typeof console&&console.log&&Function.prototype.apply.call(console.log,console,arguments)},e.formatArgs=function(){var t=arguments,n=this.useColors;if(t[0]=(n?"%c":"")+this.namespace+(n?" %c":" ")+t[0]+(n?"%c ":" ")+"+"+e.humanize(this.diff),!n)return t;var r="color: "+this.color;t=[t[0],r,"color: inherit"].concat(Array.prototype.slice.call(t,1));var i=0,a=0;return t[0].replace(/%[a-z%]/g,(function(t){"%%"!==t&&(i++,"%c"===t&&(a=i))})),t.splice(a,0,r),t},e.save=function(t){try{null==t?e.storage.removeItem("debug"):e.storage.debug=t}catch(t){}},e.load=r,e.useColors=function(){return"WebkitAppearance"in document.documentElement.style||window.console&&(console.firebug||console.exception&&console.table)||navigator.userAgent.toLowerCase().match(/firefox\/(\d+)/)&&parseInt(RegExp.$1,10)>=31},e.storage="undefined"!=typeof chrome&&void 0!==chrome.storage?chrome.storage.local:function(){try{return window.localStorage}catch(t){}}(),e.colors=["lightseagreen","forestgreen","goldenrod","dodgerblue","darkorchid","crimson"],e.formatters.j=function(t){return JSON.stringify(t)},e.enable(r())},function(t,e){function n(t){if(t)return function(t){for(var e in n.prototype)t[e]=n.prototype[e];return t}(t)}t.exports=n,n.prototype.on=n.prototype.addEventListener=function(t,e){return this._callbacks=this._callbacks||{},(this._callbacks[t]=this._callbacks[t]||[]).push(e),this},n.prototype.once=function(t,e){var n=this;function r(){n.off(t,r),e.apply(this,arguments)}return this._callbacks=this._callbacks||{},r.fn=e,this.on(t,r),this},n.prototype.off=n.prototype.removeListener=n.prototype.removeAllListeners=n.prototype.removeEventListener=function(t,e){if(this._callbacks=this._callbacks||{},0==arguments.length)return this._callbacks={},this;var n,r=this._callbacks[t];if(!r)return this;if(1==arguments.length)return delete this._callbacks[t],this;for(var i=0;i<r.length;i++)if((n=r[i])===e||n.fn===e){r.splice(i,1);break}return this},n.prototype.emit=function(t){this._callbacks=this._callbacks||{};var e=[].slice.call(arguments,1),n=this._callbacks[t];if(n)for(var r=0,i=(n=n.slice(0)).length;r<i;++r)n[r].apply(this,e);return this},n.prototype.listeners=function(t){return this._callbacks=this._callbacks||{},this._callbacks[t]||[]},n.prototype.hasListeners=function(t){return!!this.listeners(t).length}},function(t,e){t.exports=function(t,e){var n=function(){};n.prototype=e.prototype,t.prototype=new n,t.prototype.constructor=t}},function(t,e,n){function r(){var t;try{t=e.storage.debug}catch(t){}return t}(e=t.exports=n(494)).log=function(){return"object"==typeof console&&console.log&&Function.prototype.apply.call(console.log,console,arguments)},e.formatArgs=function(){var t=arguments,n=this.useColors;if(t[0]=(n?"%c":"")+this.namespace+(n?" %c":" ")+t[0]+(n?"%c ":" ")+"+"+e.humanize(this.diff),!n)return t;var r="color: "+this.color;t=[t[0],r,"color: inherit"].concat(Array.prototype.slice.call(t,1));var i=0,a=0;return t[0].replace(/%[a-z%]/g,(function(t){"%%"!==t&&(i++,"%c"===t&&(a=i))})),t.splice(a,0,r),t},e.save=function(t){try{null==t?e.storage.removeItem("debug"):e.storage.debug=t}catch(t){}},e.load=r,e.useColors=function(){return"WebkitAppearance"in document.documentElement.style||window.console&&(console.firebug||console.exception&&console.table)||navigator.userAgent.toLowerCase().match(/firefox\/(\d+)/)&&parseInt(RegExp.$1,10)>=31},e.storage="undefined"!=typeof chrome&&void 0!==chrome.storage?chrome.storage.local:function(){try{return window.localStorage}catch(t){}}(),e.colors=["lightseagreen","forestgreen","goldenrod","dodgerblue","darkorchid","crimson"],e.formatters.j=function(t){return JSON.stringify(t)},e.enable(r())},function(t,e,n){"use strict";
/*
object-assign
(c) Sindre Sorhus
@license MIT
```
