## supplementary data file 3 for "Repeated truncation of a modular antimicrobial peptide gene for neural context": widgets.js

```
Function&&Function.prototype&&Function.prototype.bind&&(/(MSIE ([6789]|10|11))|Trident/.test(navigator.userAgent)||(window.__twttr&&window.__twttr.widgets&&window.__twttr.widgets.loaded&&window.twttr.widgets.load&&window.twttr.widgets.load(),window.__twttr&&window.__twttr.widgets&&window.__twttr.widgets.init||function(t){function e(e){for(var n,i,o=e[0],s=e[1],a=0,c=[];a<o.length;a++)i=o[a],r[i]&&c.push(r[i][0]),r[i]=0;for(n in s)Object.prototype.hasOwnProperty.call(s,n)&&(t[n]=s[n]);for(u&&u(e);c.length;)c.shift()()}var n={},r={1:0};function i(e){if(n[e])return n[e].exports;var r=n[e]={i:e,l:!1,exports:{}};return t[e].call(r.exports,r,r.exports,i),r.l=!0,r.exports}i.e=function(t){var e=[],n=r[t];if(0!==n)if(n)e.push(n[2]);else{var o=new Promise(function(e,i){n=r[t]=[e,i]});e.push(n[2]=o);var s,a=document.getElementsByTagName("head")[0],u=document.createElement("script");u.charset="utf-8",u.timeout=120,i.nc&&u.setAttribute("nonce",i.nc),u.src=function(t){return i.p+"js/"+({0:"moment~timeline",2:"dm_button",3:"button",4:"moment",5:"periscope_on_air",6:"horizon_timeline",7:"timeline",8:"horizon_tweet"}[t]||t)+"."+{0:"bcb1cafa923482f4826e32741fe16a98",2:"35421d81b9479e1a0a333bbe879f3f6f",3:"5573c974dc31bbdab5ea7923a0bd5cf3",4:"72a230b36d8b9d78def80f272bcf895f",5:"5dc814ad2a165a01f994cc7dd36dcd3b",6:"f52b1b8c6d48352e19119d2f606b03be",7:"28ecda9667eeb8e1b18898b99fee6c31",8:"2bd42981e3af03ce9186a5655508da28"}[t]+".js"}(t),s=function(e){u.onerror=u.onload=null,clearTimeout(c);var n=r[t];if(0!==n){if(n){var i=e&&("load"===e.type?"missing":e.type),o=e&&e.target&&e.target.src,s=new Error("Loading chunk "+t+" failed.\n("+i+": "+o+")");s.type=i,s.request=o,n[1](s)}r[t]=void 0}};var c=setTimeout(function(){s({type:"timeout",target:u})},12e4);u.onerror=u.onload=s,a.appendChild(u)}return Promise.all(e)},i.m=t,i.c=n,i.d=function(t,e,n){i.o(t,e)||Object.defineProperty(t,e,{enumerable:!0,get:n})},i.r=function(t){"undefined"!=typeof Symbol&&Symbol.toStringTag&&Object.defineProperty(t,Symbol.toStringTag,{value:"Module"}),Object.defineProperty(t,"__esModule",{value:!0})},i.t=function(t,e){if(1&e&&(t=i(t)),8&e)return t;if(4&e&&"object"==typeof t&&t&&t.__esModule)return t;var n=Object.create(null);if(i.r(n),Object.defineProperty(n,"default",{enumerable:!0,value:t}),2&e&&"string"!=typeof t)for(var r in t)i.d(n,r,function(e){return t[e]}.bind(null,r));return n},i.n=function(t){var e=t&&t.__esModule?function(){return t.default}:function(){return t};return i.d(e,"a",e),e},i.o=function(t,e){return Object.prototype.hasOwnProperty.call(t,e)},i.p="https://platform.twitter.com/",i.oe=function(t){throw console.error(t),t};var o=window.__twttrll=window.__twttrll||[],s=o.push.bind(o);o.push=e,o=o.slice();for(var a=0;a<o.length;a++)e(o[a]);var u=s;i(i.s=91)}([function(t,e,n){var r=n(1);function i(t,e){var n;for(n in t)t.hasOwnProperty&&!t.hasOwnProperty(n)||e(n,t[n]);return t}function o(t){return{}.toString.call(t).match(/\s([a-zA-Z]+)/)[1].toLowerCase()}function s(t){return t===Object(t)}function a(t){var e;if(!s(t))return!1;if(Object.keys)return!Object.keys(t).length;for(e in t)if(t.hasOwnProperty(e))return!1;return!0}function u(t){return t?Array.prototype.slice.call(t):[]}t.exports={aug:function(t){return u(arguments).slice(1).forEach(function(e){i(e,function(e,n){t[e]=n})}),t},async:function(t,e){r.setTimeout(function(){t.call(e||null)},0)},compact:function t(e){return i(e,function(n,r){s(r)&&(t(r),a(r)&&delete e[n]),void 0!==r&&null!==r&&""!==r||delete e[n]}),e},contains:function(t,e){return!(!t||!t.indexOf)&&t.indexOf(e)>-1},forIn:i,isObject:s,isEmptyObject:a,toType:o,isType:function(t,e){return t==o(e)},toRealArray:u}},function(t,e){t.exports=window},function(t,e,n){var r=n(6);t.exports=function(){var t=this;this.promise=new r(function(e,n){t.resolve=e,t.reject=n})}},function(t,e,n){var r=n(11),i=/(?:^|(?:https?:)?\/\/(?:www\.)?twitter\.com(?::\d+)?(?:\/intent\/(?:follow|user)\/?\?screen_name=|(?:\/#!)?\/))@?([\w]+)(?:\?|&|$)/i,o=/(?:^|(?:https?:)?\/\/(?:www\.)?twitter\.com(?::\d+)?\/(?:#!\/)?[\w_]+\/status(?:es)?\/)(\d+)/i,s=/^http(s?):\/\/(\w+\.)*twitter\.com([:/]|$)/i,a=/^http(s?):\/\/(ton|pbs)\.twimg\.com/,u=/^#?([^.,<>!\s/#\-()'"]+)$/,c=/twitter\.com(?::\d{2,4})?\/intent\/(\w+)/,d=/^https?:\/\/(?:www\.)?twitter\.com\/\w+\/timelines\/(\d+)/i,l=/^https?:\/\/(?:www\.)?twitter\.com\/i\/moments\/(\d+)/i,f=/^https?:\/\/(?:www\.)?twitter\.com\/(\w+)\/(?:likes|favorites)/i,h=/^https?:\/\/(?:www\.)?twitter\.com\/(\w+)\/lists\/([\w-%]+)/i,p=/^https?:\/\/(?:www\.)?twitter\.com\/i\/live\/(\d+)/i,m=/^https?:\/\/syndication\.twitter\.com\/settings/i,v=/^https?:\/\/(localhost|platform)\.twitter\.com(?::\d+)?\/widgets\/widget_iframe\.(.+)/i,g=/^https?:\/\/(?:www\.)?twitter\.com\/search\?q=(\w+)/i;function w(t){return"string"==typeof t&&i.test(t)&&RegExp.$1.length<=20}function y(t){if(w(t))return RegExp.$1}function b(t,e){var n=r.decodeURL(t);if(e=e||!1,n.screen_name=y(t),n.screen_name)return r.url("https://twitter.com/intent/"+(e?"follow":"user"),n)}function _(t){return"string"==typeof t&&u.test(t)}function E(t){return"string"==typeof t&&o.test(t)}t.exports={isHashTag:_,hashTag:function(t,e){if(e=void 0===e||e,_(t))return(e?"#":"")+RegExp.$1},isScreenName:w,screenName:y,isStatus:E,status:function(t){return E(t)&&RegExp.$1},intentForProfileURL:b,intentForFollowURL:function(t){return b(t,!0)},isTwitterURL:function(t){return s.test(t)},isTwimgURL:function(t){return a.test(t)},isIntentURL:function(t){return c.test(t)},isSettingsURL:function(t){return m.test(t)},isWidgetIframeURL:function(t){return v.test(t)},isSearchUrl:function(t){return g.test(t)},regexen:{profile:i},momentId:function(t){return l.test(t)&&RegExp.$1},collectionId:function(t){return d.test(t)&&RegExp.$1},intentType:function(t){return c.test(t)&&RegExp.$1},likesScreenName:function(t){return f.test(t)&&RegExp.$1},listScreenNameAndSlug:function(t){var e,n,r;if(h.test(t)){e=RegExp.$1,n=RegExp.$2;try{r=decodeURIComponent(n)}catch(t){}return{ownerScreenName:e,slug:r||n}}return!1},eventId:function(t){return p.test(t)&&RegExp.$1}}},function(t,e){t.exports=document},function(t,e,n){var r=n(0),i=[!0,1,"1","on","ON","true","TRUE","yes","YES"],o=[!1,0,"0","off","OFF","false","FALSE","no","NO"];function s(t){return void 0!==t&&null!==t&&""!==t}function a(t){return c(t)&&t%1==0}function u(t){return c(t)&&!a(t)}function c(t){return s(t)&&!isNaN(t)}function d(t){return r.contains(o,t)}function l(t){return r.contains(i,t)}t.exports={hasValue:s,isInt:a,isFloat:u,isNumber:c,isString:function(t){return"string"===r.toType(t)},isArray:function(t){return s(t)&&"array"==r.toType(t)},isTruthValue:l,isFalseValue:d,asInt:function(t){if(a(t))return parseInt(t,10)},asFloat:function(t){if(u(t))return t},asNumber:function(t){if(c(t))return t},asBoolean:function(t){return!(!s(t)||!l(t)&&(d(t)||!t))}}},function(t,e,n){var r=n(1),i=n(21),o=n(49);i.hasPromiseSupport()||(r.Promise=o),t.exports=r.Promise},function(t,e,n){var r=n(0);t.exports=function(t,e){var n=Array.prototype.slice.call(arguments,2);return function(){var i=r.toRealArray(arguments);return t.apply(e,n.concat(i))}}},function(t,e,n){var r=n(51);t.exports=new r("__twttr")},function(t,e){t.exports=location},function(t,e,n){var r=n(0),i=/\b([\w-_]+)\b/g;function o(t){return new RegExp("\\b"+t+"\\b","g")}function s(t,e){t.classList?t.classList.add(e):o(e).test(t.className)||(t.className+=" "+e)}function a(t,e){t.classList?t.classList.remove(e):t.className=t.className.replace(o(e)," ")}function u(t,e){return t.classList?t.classList.contains(e):r.contains(c(t),e)}function c(t){return r.toRealArray(t.classList?t.classList:t.className.match(i))}t.exports={add:s,remove:a,replace:function(t,e,n){if(t.classList&&u(t,e))return a(t,e),void s(t,n);t.className=t.className.replace(o(e),n)},toggle:function(t,e,n){return void 0===n&&t.classList&&t.classList.toggle?t.classList.toggle(e,n):(n?s(t,e):a(t,e),n)},present:u,list:c}},function(t,e,n){var r=n(5),i=n(0);function o(t){return encodeURIComponent(t).replace(/\+/g,"%2B").replace(/'/g,"%27")}function s(t){return decodeURIComponent(t)}function a(t){var e=[];return i.forIn(t,function(t,n){var s=o(t);i.isType("array",n)||(n=[n]),n.forEach(function(t){r.hasValue(t)&&e.push(s+"="+o(t))})}),e.sort().join("&")}function u(t){var e={};return t?(t.split("&").forEach(function(t){var n=t.split("="),r=s(n[0]),o=s(n[1]);if(2==n.length){if(!i.isType("array",e[r]))return r in e?(e[r]=[e[r]],void e[r].push(o)):void(e[r]=o);e[r].push(o)}}),e):{}}t.exports={url:function(t,e){return a(e).length>0?i.contains(t,"?")?t+"&"+a(e):t+"?"+a(e):t},decodeURL:function(t){var e=t&&t.split("?");return 2==e.length?u(e[1]):{}},decode:u,encode:a,encodePart:o,decodePart:s}},function(t,e,n){var r=n(9),i=n(1),o=n(0),s={},a=o.contains(r.href,"tw_debug=true");function u(){}function c(){}function d(){return i.performance&&+i.performance.now()||+new Date}function l(t,e){if(i.console&&i.console[t])switch(e.length){case 1:i.console[t](e[0]);break;case 2:i.console[t](e[0],e[1]);break;case 3:i.console[t](e[0],e[1],e[2]);break;case 4:i.console[t](e[0],e[1],e[2],e[3]);break;case 5:i.console[t](e[0],e[1],e[2],e[3],e[4]);break;default:0!==e.length&&i.console.warn&&i.console.warn("too many params passed to logger."+t)}}t.exports={devError:u,devInfo:c,devObject:function(t,e){},publicError:function(){l("error",o.toRealArray(arguments))},publicLog:function(){l("info",o.toRealArray(arguments))},publicWarn:function(){l("warn",o.toRealArray(arguments))},time:function(t){a&&(s[t]=d())},timeEnd:function(t){a&&s[t]&&(d(),s[t])}}},function(t,e,n){var r=n(19),i=n(5),o=n(11),s=n(0),a=n(115);t.exports=function(t){var e=t.href&&t.href.split("?")[1],n=e?o.decode(e):{},u={lang:a(t),width:t.getAttribute("data-width")||t.getAttribute("width"),height:t.getAttribute("data-height")||t.getAttribute("height"),related:t.getAttribute("data-related"),partner:t.getAttribute("data-partner")};return i.asBoolean(t.getAttribute("data-dnt"))&&r.setOn(),s.forIn(u,function(t,e){var r=n[t];n[t]=i.hasValue(r)?r:e}),s.compact(n)}},function(t,e,n){var r=n(77),i=n(22);t.exports=function(){var t="data-twitter-extracted-"+i.generate();return function(e,n){return r(e,n).filter(function(e){return!e.hasAttribute(t)}).map(function(e){return e.setAttribute(t,"true"),e})}}},function(t,e){function n(t,e,n,r,i,o,s){this.factory=t,this.Sandbox=e,this.srcEl=o,this.targetEl=i,this.parameters=r,this.className=n,this.options=s}n.prototype.destroy=function(){this.srcEl=this.targetEl=null},t.exports=n},function(t,e){t.exports={DM_BUTTON:"twitter-dm-button",FOLLOW_BUTTON:"twitter-follow-button",HASHTAG_BUTTON:"twitter-hashtag-button",MENTION_BUTTON:"twitter-mention-button",MOMENT:"twitter-moment",PERISCOPE:"periscope-on-air",SHARE_BUTTON:"twitter-share-button",TIMELINE:"twitter-timeline",TWEET:"twitter-tweet"}},function(t,e,n){var r=n(6),i=n(19),o=n(53),s=n(36),a=n(5),u=n(0);t.exports=function(t,e,n){var c;return t=t||[],e=e||{},c="ƒ("+t.join(", ")+", target, [options]);",function(){var d,l,f,h,p=Array.prototype.slice.apply(arguments,[0,t.length]),m=Array.prototype.slice.apply(arguments,[t.length]);return m.forEach(function(t){t&&(t.nodeType!==Node.ELEMENT_NODE?u.isType("function",t)?d=t:u.isType("object",t)&&(l=t):f=t)}),p.length!==t.length||0===m.length?(d&&u.async(function(){d(!1)}),r.reject(new Error("Not enough parameters. Expected: "+c))):f?(l=u.aug({},l||{},e),t.forEach(function(t){l[t]=p.shift()}),a.asBoolean(l.dnt)&&i.setOn(),h=s.getExperiments().then(function(t){return o.addWidget(n(l,f,void 0,t))}),d&&h.then(d,function(){d(!1)}),h):(d&&u.async(function(){d(!1)}),r.reject(new Error("No target element specified. Expected: "+c)))}}},function(t,e,n){var r=n(98),i=n(2),o=n(0);function s(t,e){return function(){try{e.resolve(t.call(this))}catch(t){e.reject(t)}}}t.exports={sync:function(t,e){t.call(e)},read:function(t,e){var n=new i;return r.read(s(t,n),e),n.promise},write:function(t,e){var n=new i;return r.write(s(t,n),e),n.promise},defer:function(t,e,n){var a=new i;return o.isType("function",t)&&(n=e,e=t,t=1),r.defer(t,s(e,a),n),a.promise}}},function(t,e,n){var r=n(4),i=n(9),o=n(38),s=n(102),a=n(5),u=n(33),c=!1,d=/https?:\/\/([^/]+).*/i;t.exports={setOn:function(){c=!0},enabled:function(t,e){return!!(c||a.asBoolean(u.val("dnt"))||s.isUrlSensitive(e||i.host)||o.isFramed()&&s.isUrlSensitive(o.rootDocumentLocation())||(t=d.test(t||r.referrer)&&RegExp.$1)&&s.isUrlSensitive(t))}}},function(t,e,n){var r=n(8),i=n(59),o="https://platform.twitter.com",s=["https://syndication.twitter.com","https://cdn.syndication.twimg.com","https://localhost.twitter.com:8444"],a=["https://syndication.twitter.com","https://localhost.twitter.com:8445"],u=["https://platform.twitter.com","https://localhost.twitter.com",/^https:\/\/ton\.smf1\.twitter\.com\/syndication-internal\/embed-iframe\/[0-9A-Za-z_-]+\/app/],c=function(t,e){return t.some(function(t){return t instanceof RegExp?t.test(e):t===e})},d=function(){var t=r.get("backendHost");return t&&c(s,t)?t:"https://cdn.syndication.twimg.com"},l=function(){var t=r.get("settingsSvcHost");return t&&c(a,t)?t:"https://syndication.twitter.com"};function f(t,e){var n=[t];return e.forEach(function(t){n.push(function(t){var e=(t||"").toString(),n="/"===e.slice(0,1)?1:0,r=function(t){return"/"===t.slice(-1)}(e)?-1:void 0;return e.slice(n,r)}(t))}),n.join("/")}t.exports={cookieConsent:function(t){var e=t||[];return e.unshift("cookie/consent"),f(l(),e)},embedIframe:function(t,e){var n=t||[],s=o,a=r.get("embedIframeURL");return a&&c(u,a)?f(a,n)+".html":(n.unshift(i.getBaseURLPath(e)),f(s,n)+".html")},eventVideo:function(t){var e=t||[];return e.unshift("video/event"),f(d(),e)},grid:function(t){var e=t||[];return e.unshift("grid/collection"),f(d(),e)},moment:function(t){var e=t||[];return e.unshift("moments"),f(d(),e)},settings:function(t){var e=t||[];return e.unshift("settings"),f(l(),e)},timeline:function(t){var e=t||[];return e.unshift("timeline"),f(d(),e)},tweetBatch:function(t){var e=t||[];return e.unshift("tweets.json"),f(d(),e)},video:function(t){var e=t||[];return e.unshift("widgets/video"),f(d(),e)}}},function(t,e,n){var r=n(4),i=n(92),o=n(1),s=n(0),a=i.userAgent;function u(t){return/(Trident|MSIE|Edge[/ ]?\d)/.test(t=t||a)}t.exports={retina:function(t){return(t=t||o).devicePixelRatio?t.devicePixelRatio>=1.5:!!t.matchMedia&&t.matchMedia("only screen and (min-resolution: 144dpi)").matches},anyIE:u,ie9:function(t){return/MSIE 9/.test(t=t||a)},ie10:function(t){return/MSIE 10/.test(t=t||a)},ios:function(t){return/(iPad|iPhone|iPod)/.test(t=t||a)},android:function(t){return/^Mozilla\/5\.0 \(Linux; (U; )?Android/.test(t=t||a)},canPostMessage:function(t,e){return t=t||o,e=e||a,t.postMessage&&!(u(e)&&t.opener)},touch:function(t,e,n){return t=t||o,e=e||i,n=n||a,"ontouchstart"in t||/Opera Mini/.test(n)||e.msMaxTouchPoints>0},cssTransitions:function(){var t=r.body.style;return void 0!==t.transition||void 0!==t.webkitTransition||void 0!==t.mozTransition||void 0!==t.oTransition||void 0!==t.msTransition},hasPromiseSupport:function(){return!!(o.Promise&&o.Promise.resolve&&o.Promise.reject&&o.Promise.all&&o.Promise.race&&(new o.Promise(function(e){t=e}),s.isType("function",t)));var t},hasIntersectionObserverSupport:function(){return!!o.IntersectionObserver},hasPerformanceInformation:function(){return o.performance&&o.performance.getEntriesByType}}},function(t,e){var n="i",r=0,i=0;t.exports={generate:function(){return n+String(+new Date)+Math.floor(1e5*Math.random())+r++},deterministic:function(){return n+String(i++)}}},function(t,e,n){var r=n(50),i=n(52),o=n(0);t.exports=o.aug(r.get("events")||{},i.Emitter)},function(t,e,n){var r=n(6),i=n(2);function o(t,e){return t.then(e,e)}function s(t){return t instanceof r}t.exports={always:o,allResolved:function(t){var e;return void 0===t?r.reject(new Error("undefined is not an object")):Array.isArray(t)?(e=t.length)?new r(function(n,r){var i=0,o=[];function a(){(i+=1)===e&&(0===o.length?r():n(o))}function u(t){o.push(t),a()}t.forEach(function(t){s(t)?t.then(u,a):u(t)})}):r.resolve([]):r.reject(new Error("Type error"))},some:function(t){var e;return e=(t=t||[]).length,t=t.filter(s),e?e!==t.length?r.reject("non-Promise passed to .some"):new r(function(e,n){var r=0;function i(){(r+=1)===t.length&&n()}t.forEach(function(t){t.then(e,i)})}):r.reject("no promises passed to .some")},isPromise:s,allSettled:function(t){function e(){}return r.all((t||[]).map(function(t){return o(t,e)}))},timeout:function(t,e){var n=new i;return setTimeout(function(){n.reject(new Error("Promise timed out"))},e),t.then(function(t){n.resolve(t)},function(t){n.reject(t)}),n.promise}}},function(t,e,n){var r=n(1).JSON;t.exports={stringify:r.stringify||r.encode,parse:r.parse||r.decode}},function(t,e,n){var r=n(27),i=n(108);t.exports=r.build([i])},function(t,e,n){var r=n(39),i=n(105),o=n(7);(r=Object.create(r)).build=o(r.build,null,i),t.exports=r},function(t,e,n){var r=n(39),i=n(40),o=n(7);(r=Object.create(r)).build=o(r.build,null,i),t.exports=r},function(t,e,n){var r=n(79),i=n(80),o=n(81),s=n(9),a=n(71),u=n(82),c=n(19),d=n(5),l=n(22),f=n(0);function h(t){if(!t||!t.headers)throw new Error("unexpected response schema");return{html:t.body,config:t.config,pollInterval:1e3*parseInt(t.headers.xPolling,10)||null,maxCursorPosition:t.headers.maxPosition,minCursorPosition:t.headers.minPosition}}function p(t){if(t&&t.headers)throw new Error(t.headers.status);throw t instanceof Error?t:new Error(t)}t.exports=function(t){t.params({instanceId:{required:!0,fallback:l.deterministic},lang:{required:!0,transform:a.matchLanguage,fallback:"en"},tweetLimit:{transform:d.asInt}}),t.defineProperty("endpoint",{get:function(){throw new Error("endpoint not specified")}}),t.defineProperty("pollEndpoint",{get:function(){return this.endpoint}}),t.define("cbId",function(t){var e=t?"_new":"_old";return"tl_"+this.params.instanceId+"_"+this.id+e}),t.define("queryParams",function(){return{lang:this.params.lang,tz:u.getTimezoneOffset(),t:r(),domain:s.host,tweet_limit:this.params.tweetLimit,dnt:c.enabled()}}),t.define("fetch",function(){return i.fetch(this.endpoint,this.queryParams(),o,this.cbId()).then(h,p)}),t.define("poll",function(t,e){var n,r;return n={since_id:(t=t||{}).sinceId,max_id:t.maxId,min_position:t.minPosition,max_position:t.maxPosition},r=f.aug(this.queryParams(),n),i.fetch(this.pollEndpoint,r,o,this.cbId(e)).then(h,p)})}},function(t,e,n){var r=n(52).makeEmitter();t.exports={emitter:r,START:"start",ALL_WIDGETS_RENDER_START:"all_widgets_render_start",ALL_WIDGETS_RENDER_END:"all_widgets_render_end",ALL_WIDGETS_AND_IMAGES_LOADED:"all_widgets_and_images_loaded"}},function(t,e,n){var r=n(4),i=n(0);t.exports=function(t,e,n){var o;if(n=n||r,t=t||{},e=e||{},t.name){try{o=n.createElement('<iframe name="'+t.name+'"></iframe>')}catch(e){(o=n.createElement("iframe")).name=t.name}delete t.name}else o=n.createElement("iframe");return t.id&&(o.id=t.id,delete t.id),o.allowtransparency="true",o.scrolling="no",o.setAttribute("frameBorder",0),o.setAttribute("allowTransparency",!0),i.forIn(t,function(t,e){o.setAttribute(t,e)}),i.forIn(e,function(t,e){o.style[t]=e}),o}},function(t,e,n){var r=n(27),i=n(122);t.exports=r.build([i])},function(t,e,n){var r,i=n(4);function o(t){var e,n,o,s=0;for(r={},e=(t=t||i).getElementsByTagName("meta");e[s];s++){if(n=e[s],/^twitter:/.test(n.getAttribute("name")))o=n.getAttribute("name").replace(/^twitter:/,"");else{if(!/^twitter:/.test(n.getAttribute("property")))continue;o=n.getAttribute("property").replace(/^twitter:/,"")}r[o]=n.getAttribute("content")||n.getAttribute("value")}}o(),t.exports={init:o,val:function(t){return r[t]}}},function(t,e,n){var r=n(0),i=n(45);t.exports={closest:function t(e,n,o){var s;if(n)return o=o||n&&n.ownerDocument,s=r.isType("function",e)?e:function(t){return function(e){return!!e.tagName&&i(e,t)}}(e),n===o?s(n)?n:void 0:s(n)?n:t(s,n.parentNode,o)}}},function(t,e,n){var r=n(10),i={},o=-1,s={};function a(t){var e=t.getAttribute("data-twitter-event-id");return e||(t.setAttribute("data-twitter-event-id",++o),o)}function u(t,e,n){var r=0,i=t&&t.length||0;for(r=0;r<i;r++)if(t[r].call(e,n,e),n.ceaseImmediately)return!1}function c(t,e,n){for(var i=n||t.target||t.srcElement,o=r.list(i).map(function(t){return"."+t}).concat(i.tagName),s=0,a=o.length;s<a;s++)if(!1===u(e[o[s]],i,t))return;t.cease||i!==this&&c.call(this,t,e,i.parentElement||i.parentNode)}function d(t,e,n,r){function i(r){c.call(t,r,n[e])}!function(t,e,n,r){t.id&&(s[t.id]=s[t.id]||[],s[t.id].push({el:t,listener:e,type:n,rootId:r}))}(t,i,e,r),t.addEventListener(e,i,!1)}function l(t){t&&t.preventDefault?t.preventDefault():t.returnValue=!1}function f(t){t&&(t.cease=!0)&&t.stopPropagation?t.stopPropagation():t.cancelBubble=!0}t.exports={stop:function(t){return f(t),l(t),!1},stopPropagation:f,stopImmediatePropagation:function(t){t&&(t.ceaseImmediately=!0,f(t),t.stopImmediatePropagation())},preventDefault:l,delegate:function(t,e,n,r){var o=a(t);i[o]=i[o]||{},i[o][e]||(i[o][e]={},d(t,e,i[o],o)),i[o][e][n]=i[o][e][n]||[],i[o][e][n].push(r)},simulate:function(t,e,n){var r=a(e),o=i[r]&&i[r];c.call(e,{target:n},o[t])},removeDelegatesForWidget:function(t){var e=s[t];e&&(e.forEach(function(t){t.el.removeEventListener(t.type,t.listener,!1),delete i[t.rootId]}),delete s[t])}}},function(t,e,n){var r=n(111),i=n(25),o=n(1);function s(t){var e="";if(!t)return"";try{return e=i.stringify(t),o.btoa(e)}catch(t){return""}}function a(t){return r.settingsLoaded().then(function(e){return e[t]})}function u(){return a("features")}t.exports={encodeFeatures:s,shouldObtainCookieConsent:function(){return a("shouldObtainCookieConsent")},getExperiments:u,getExperiment:function(t){return u().then(function(e){if(!e[t])throw new Error("Experiment not found");return e[t]})},getHorizonSettings:function(){return r.settingsLoaded().then(function(t){return{features:t.features,sessionId:t.sessionId}})},getActiveExperimentDataString:function(){return u().then(function(t){return s(t)})},getExperimentKeys:function(){return u().then(function(t){return Object.keys(t)})},load:function(){r.load()}}},function(t,e,n){var r=n(46),i=n(4),o=n(19),s=n(34),a=n(73),u=n(25),c=n(8),d=n(3),l=n(0),f=1,h=r.version,p=c.get("clientEventEndpoint")||"https://syndication.twitter.com/i/jot";function m(t){return l.aug({client:"tfw"},t||{})}function v(t,e,n){return e=e||{},l.aug({},e,{_category_:t,triggered_on:e.triggered_on||+new Date,dnt:o.enabled(n)})}t.exports={extractTermsFromDOM:function t(e,n){var r;return n=n||{},e&&e.nodeType===Node.ELEMENT_NODE?((r=e.getAttribute("data-scribe"))&&r.split(" ").forEach(function(t){var e=t.trim().split(":"),r=e[0],i=e[1];r&&i&&!n[r]&&(n[r]=i)}),t(e.parentNode,n)):n},clickEventElement:function(t){var e=s.closest("[data-expanded-url]",t),n=e&&e.getAttribute("data-expanded-url");return n&&d.isTwitterURL(n)?"twitter_url":"url"},flattenClientEventPayload:function(t,e){return l.aug({},e,{event_namespace:t})},formatGenericEventData:v,formatClientEventData:function(t,e,n){var r=t&&t.widget_origin||i.referrer;return(t=v("tfw_client_event",t,r)).client_version=h,t.format_version=void 0!==n?n:1,e||(t.widget_origin=r),t},formatClientEventNamespace:m,formatHorizonTweetData:function(t){var e={item_ids:[],item_details:{}};return e.item_ids.push(t),e.item_details[t]={item_type:a.TWEET},e},formatTweetAssociation:function(t,e){var n={};return(e=e||{}).association_namespace=m(t),n[f]=e,n},noticeSeen:function(t){return"notice"===t.element&&"seen"===t.action},splitLogEntry:function(t){var e,n,r,i,o;return t.item_ids&&t.item_ids.length>1?(e=Math.floor(t.item_ids.length/2),n=t.item_ids.slice(0,e),r={},i=t.item_ids.slice(e),o={},n.forEach(function(e){r[e]=t.item_details[e]}),i.forEach(function(e){o[e]=t.item_details[e]}),[l.aug({},t,{item_ids:n,item_details:r}),l.aug({},t,{item_ids:i,item_details:o})]):[t]},stringify:function(t){var e,n=Array.prototype.toJSON;return delete Array.prototype.toJSON,e=u.stringify(t),n&&(Array.prototype.toJSON=n),e},CLIENT_EVENT_ENDPOINT:p,RUFOUS_REDIRECT:"https://platform.twitter.com/jot.html"}},function(t,e,n){var r=n(9),i=n(75),o=n(0),s=i.getCanonicalURL()||r.href,a=s;t.exports={isFramed:function(){return s!==a},rootDocumentLocation:function(t){return t&&o.isType("string",t)&&(s=t),s},currentDocumentLocation:function(){return a}}},function(t,e,n){var r=n(103),i=n(104),o=n(0);t.exports={couple:function(){return o.toRealArray(arguments)},build:function(t,e,n){var o=new t;return(e=i(r(e||[]))).forEach(function(t){t.call(null,o)}),o.build(n)}}},function(t,e,n){var r=n(106),i=n(0),o=n(41);function s(){this.Component=this.factory(),this._adviceArgs=[],this._lastArgs=[]}i.aug(s.prototype,{factory:o,build:function(t){var e=this;return this.Component,i.aug(this.Component.prototype.boundParams,t),this._adviceArgs.concat(this._lastArgs).forEach(function(t){(function(t,e,n){var r=this[e];if(!r)throw new Error(e+" does not exist");this[e]=t(r,n)}).apply(e.Component.prototype,t)}),delete this._lastArgs,delete this._adviceArgs,this.Component},params:function(t){var e=this.Component.prototype.paramConfigs;t=t||{},this.Component.prototype.paramConfigs=i.aug({},t,e)},define:function(t,e){if(t in this.Component.prototype)throw new Error(t+" has previously been defined");this.override(t,e)},defineStatic:function(t,e){this.Component[t]=e},override:function(t,e){this.Component.prototype[t]=e},defineProperty:function(t,e){if(t in this.Component.prototype)throw new Error(t+" has previously been defined");this.overrideProperty(t,e)},overrideProperty:function(t,e){var n=i.aug({configurable:!0},e);Object.defineProperty(this.Component.prototype,t,n)},before:function(t,e){this._adviceArgs.push([r.before,t,e])},after:function(t,e){this._adviceArgs.push([r.after,t,e])},around:function(t,e){this._adviceArgs.push([r.around,t,e])},last:function(t,e){this._lastArgs.push([r.after,t,e])}}),t.exports=s},function(t,e,n){var r=n(0);function i(){return!0}function o(t){return t}t.exports=function(){function t(t){var e=this;t=t||{},this.params=Object.keys(this.paramConfigs).reduce(function(n,s){var a=[],u=e.boundParams,c=e.paramConfigs[s],d=c.validate||i,l=c.transform||o;if(s in u&&a.push(u[s]),s in t&&a.push(t[s]),a="fallback"in c?a.concat(c.fallback):a,n[s]=function(t,e,n){var i=null;return t.some(function(t){if(t=r.isType("function",t)?t():t,e(t))return i=n(t),!0}),i}(a,d,l),c.required&&null==n[s])throw new Error(s+" is a required parameter");return n},{}),this.initialize()}return r.aug(t.prototype,{paramConfigs:{},boundParams:{},initialize:function(){}}),t}},function(t,e,n){var r=n(101),i=n(76),o=new(n(110))(function(t){(!function(t){return 1===t.length&&i.canFlushOneItem(t[0])}(t)?function(t){r.init(),t.forEach(function(t){var e=t.input.namespace,n=t.input.data,i=t.input.offsite,o=t.input.version;r.clientEvent(e,n,i,o)}),r.flush().then(function(){t.forEach(function(t){t.taskDoneDeferred.resolve()})},function(){t.forEach(function(t){t.taskDoneDeferred.reject()})})}:function(t){t.forEach(function(t){var e=t.input.namespace,n=t.input.data,r=t.input.offsite,o=t.input.version;i.clientEvent(e,n,r,o),t.taskDoneDeferred.resolve()})})(t)});t.exports={scribe:function(t,e,n,r){return o.add({namespace:t,data:e,offsite:n,version:r})},pause:function(){o.pause()},resume:function(){o.resume()}}},function(t,e,n){var r,i=n(10),o=n(4),s=n(1),a=n(33),u=n(54),c=n(5),d=n(22),l="csptest";t.exports={inlineStyle:function(){var t=l+d.generate(),e=o.createElement("div"),n=o.createElement("style"),f="."+t+" { visibility: hidden; }";return!!o.body&&(c.asBoolean(a.val("widgets:csp"))&&(r=!1),void 0!==r?r:(e.style.display="none",i.add(e,t),n.type="text/css",n.appendChild(o.createTextNode(f)),o.body.appendChild(n),o.body.appendChild(e),r="hidden"===s.getComputedStyle(e).visibility,u(e),u(n),r))}}},function(t,e,n){var r=n(1);t.exports=function(t,e,n){var i,o=0;return n=n||null,function s(){var a=n||this,u=arguments,c=+new Date;if(r.clearTimeout(i),c-o>e)return o=c,void t.apply(a,u);i=r.setTimeout(function(){s.apply(a,u)},e)}}},function(t,e,n){var r=n(1).HTMLElement,i=r.prototype.matches||r.prototype.matchesSelector||r.prototype.webkitMatchesSelector||r.prototype.mozMatchesSelector||r.prototype.msMatchesSelector||r.prototype.oMatchesSelector;t.exports=function(t,e){if(i)return i.call(t,e)}},function(t){t.exports={version:"82e1070:1619632193066"}},function(t,e){t.exports=function(t){var e=t.getBoundingClientRect();return{width:e.width,height:e.height}}},function(t,e,n){var r=n(12).publicWarn;t.exports=function(){r("Warning: This Timeline type belongs to a group that will not be supported in the future (Likes, Collections, & Moments). It is not recommended for use. \n\t","* Twitter will continue to support Profile and List Timelines \n\t","* You can learn more about this change in our announcement: \n\t","https://twittercommunity.com/t/removing-support-for-embedded-like-collection-and-moment-timelines/150313 \n\t","* In order to create a new Embedded Timeline, visit: https://publish.twitter.com")}},function(t,e,n){
/*!
 * @overview es6-promise - a tiny implementation of Promises/A+.
 * @copyright Copyright (c) 2014 Yehuda Katz, Tom Dale, Stefan Penner and contributors (Conversion to ES6 API by Jake Archibald)
 * @license   Licensed under MIT license
 *            See https://raw.githubusercontent.com/stefanpenner/es6-promise/master/LICENSE
 * @version   v4.2.5+7f2b526d
 */var r;r=function(){"use strict";function t(t){return"function"==typeof t}var e=Array.isArray?Array.isArray:function(t){return"[object Array]"===Object.prototype.toString.call(t)},n=0,r=void 0,i=void 0,o=function(t,e){f[n]=t,f[n+1]=e,2===(n+=2)&&(i?i(h):w())},s="undefined"!=typeof window?window:void 0,a=s||{},u=a.MutationObserver||a.WebKitMutationObserver,c="undefined"==typeof self&&"undefined"!=typeof process&&"[object process]"==={}.toString.call(process),d="undefined"!=typeof Uint8ClampedArray&&"undefined"!=typeof importScripts&&"undefined"!=typeof MessageChannel;function l(){var t=setTimeout;return function(){return t(h,1)}}var f=new Array(1e3);function h(){for(var t=0;t<n;t+=2)(0,f[t])(f[t+1]),f[t]=void 0,f[t+1]=void 0;n=0}var p,m,v,g,w=void 0;function y(t,e){var n=this,r=new this.constructor(E);void 0===r[_]&&k(r);var i=n._state;if(i){var s=arguments[i-1];o(function(){return D(i,r,s,n._result)})}else j(n,r,t,e);return r}function b(t){if(t&&"object"==typeof t&&t.constructor===this)return t;var e=new this(E);return C(e,t),e}c?w=function(){return process.nextTick(h)}:u?(m=0,v=new u(h),g=document.createTextNode(""),v.observe(g,{characterData:!0}),w=function(){g.data=m=++m%2}):d?((p=new MessageChannel).port1.onmessage=h,w=function(){return p.port2.postMessage(0)}):w=void 0===s?function(){try{var t=Function("return this")().require("vertx");return void 0!==(r=t.runOnLoop||t.runOnContext)?function(){r(h)}:l()}catch(t){return l()}}():l();var _=Math.random().toString(36).substring(2);function E(){}var x=void 0,T=1,A=2,S={error:null};function R(t){try{return t.then}catch(t){return S.error=t,S}}function N(e,n,r){n.constructor===e.constructor&&r===y&&n.constructor.resolve===b?function(t,e){e._state===T?L(t,e._result):e._state===A?I(t,e._result):j(e,void 0,function(e){return C(t,e)},function(e){return I(t,e)})}(e,n):r===S?(I(e,S.error),S.error=null):void 0===r?L(e,n):t(r)?function(t,e,n){o(function(t){var r=!1,i=function(t,e,n,r){try{t.call(e,n,r)}catch(t){return t}}(n,e,function(n){r||(r=!0,e!==n?C(t,n):L(t,n))},function(e){r||(r=!0,I(t,e))},t._label);!r&&i&&(r=!0,I(t,i))},t)}(e,n,r):L(e,n)}function C(t,e){var n,r;t===e?I(t,new TypeError("You cannot resolve a promise with itself")):(r=typeof(n=e),null===n||"object"!==r&&"function"!==r?L(t,e):N(t,e,R(e)))}function P(t){t._onerror&&t._onerror(t._result),O(t)}function L(t,e){t._state===x&&(t._result=e,t._state=T,0!==t._subscribers.length&&o(O,t))}function I(t,e){t._state===x&&(t._state=A,t._result=e,o(P,t))}function j(t,e,n,r){var i=t._subscribers,s=i.length;t._onerror=null,i[s]=e,i[s+T]=n,i[s+A]=r,0===s&&t._state&&o(O,t)}function O(t){var e=t._subscribers,n=t._state;if(0!==e.length){for(var r=void 0,i=void 0,o=t._result,s=0;s<e.length;s+=3)r=e[s],i=e[s+n],r?D(n,r,i,o):i(o);t._subscribers.length=0}}function D(e,n,r,i){var o=t(r),s=void 0,a=void 0,u=void 0,c=void 0;if(o){if((s=function(t,e){try{return t(e)}catch(t){return S.error=t,S}}(r,i))===S?(c=!0,a=s.error,s.error=null):u=!0,n===s)return void I(n,new TypeError("A promises callback cannot return that same promise."))}else s=i,u=!0;n._state!==x||(o&&u?C(n,s):c?I(n,a):e===T?L(n,s):e===A&&I(n,s))}var z=0;function k(t){t[_]=z++,t._state=void 0,t._result=void 0,t._subscribers=[]}var M=function(){function t(t,n){this._instanceConstructor=t,this.promise=new t(E),this.promise[_]||k(this.promise),e(n)?(this.length=n.length,this._remaining=n.length,this._result=new Array(this.length),0===this.length?L(this.promise,this._result):(this.length=this.length||0,this._enumerate(n),0===this._remaining&&L(this.promise,this._result))):I(this.promise,new Error("Array Methods must be provided an Array"))}return t.prototype._enumerate=function(t){for(var e=0;this._state===x&&e<t.length;e++)this._eachEntry(t[e],e)},t.prototype._eachEntry=function(t,e){var n=this._instanceConstructor,r=n.resolve;if(r===b){var i=R(t);if(i===y&&t._state!==x)this._settledAt(t._state,e,t._result);else if("function"!=typeof i)this._remaining--,this._result[e]=t;else if(n===U){var o=new n(E);N(o,t,i),this._willSettleAt(o,e)}else this._willSettleAt(new n(function(e){return e(t)}),e)}else this._willSettleAt(r(t),e)},t.prototype._settledAt=function(t,e,n){var r=this.promise;r._state===x&&(this._remaining--,t===A?I(r,n):this._result[e]=n),0===this._remaining&&L(r,this._result)},t.prototype._willSettleAt=function(t,e){var n=this;j(t,void 0,function(t){return n._settledAt(T,e,t)},function(t){return n._settledAt(A,e,t)})},t}(),U=function(){function e(t){this[_]=z++,this._result=this._state=void 0,this._subscribers=[],E!==t&&("function"!=typeof t&&function(){throw new TypeError("You must pass a resolver function as the first argument to the promise constructor")}(),this instanceof e?function(t,e){try{e(function(e){C(t,e)},function(e){I(t,e)})}catch(e){I(t,e)}}(this,t):function(){throw new TypeError("Failed to construct 'Promise': Please use the 'new' operator, this object constructor cannot be called as a function.")}())}return e.prototype.catch=function(t){return this.then(null,t)},e.prototype.finally=function(e){var n=this.constructor;return t(e)?this.then(function(t){return n.resolve(e()).then(function(){return t})},function(t){return n.resolve(e()).then(function(){throw t})}):this.then(e,e)},e}();return U.prototype.then=y,U.all=function(t){return new M(this,t).promise},U.race=function(t){var n=this;return e(t)?new n(function(e,r){for(var i=t.length,o=0;o<i;o++)n.resolve(t[o]).then(e,r)}):new n(function(t,e){return e(new TypeError("You must pass an array to race."))})},U.resolve=b,U.reject=function(t){var e=new this(E);return I(e,t),e},U._setScheduler=function(t){i=t},U._setAsap=function(t){o=t},U._asap=o,U.polyfill=function(){var t=void 0;if("undefined"!=typeof global)t=global;else if("undefined"!=typeof self)t=self;else try{t=Function("return this")()}catch(t){throw new Error("polyfill failed because global object is unavailable in this environment")}var e=t.Promise;if(e){var n=null;try{n=Object.prototype.toString.call(e.resolve())}catch(t){}if("[object Promise]"===n&&!e.cast)return}t.Promise=U},U.Promise=U,U},t.exports=r()},function(t,e,n){var r=n(51);t.exports=new r("twttr")},function(t,e,n){var r=n(1),i=n(0);function o(t){return i.isType("string",t)?t.split("."):i.isType("array",t)?t:[]}function s(t,e){(e=e||r)[t]=e[t]||{},Object.defineProperty(this,"base",{value:e[t]}),Object.defineProperty(this,"name",{value:t})}i.aug(s.prototype,{get:function(t){return o(t).reduce(function(t,e){if(i.isObject(t))return t[e]},this.base)},set:function(t,e,n){var r=o(t),s=function(t,e){var n=o(e).slice(0,-1);return n.reduce(function(t,e,r){if(t[e]=t[e]||{},!i.isObject(t[e]))throw new Error(n.slice(0,r+1).join(".")+" is already defined with a value.");return t[e]},t)}(this.base,t),a=r.slice(-1);return n&&a in s?s[a]:s[a]=e},init:function(t,e){return this.set(t,e,!0)},unset:function(t){var e=o(t),n=this.get(e.slice(0,-1));n&&delete n[e.slice(-1)]},aug:function(t){var e=this.get(t),n=i.toRealArray(arguments).slice(1);if(e=void 0!==e?e:{},n.unshift(e),!n.every(i.isObject))throw new Error("Cannot augment non-object.");return this.set(t,i.aug.apply(null,n))},call:function(t){var e=this.get(t),n=i.toRealArray(arguments).slice(1);if(!i.isType("function",e))throw new Error("Function "+t+"does not exist.");return e.apply(null,n)},fullPath:function(t){var e=o(t);return e.unshift(this.name),e.join(".")}}),t.exports=s},function(t,e,n){var r=n(0),i=n(7),o={bind:function(t,e){return this._handlers=this._handlers||{},this._handlers[t]=this._handlers[t]||[],this._handlers[t].push(e)},unbind:function(t,e){var n;this._handlers&&this._handlers[t]&&(e?(n=this._handlers[t].indexOf(e))>=0&&this._handlers[t].splice(n,1):this._handlers[t]=[])},trigger:function(t,e){var n=this._handlers&&this._handlers[t];(e=e||{}).type=t,n&&n.forEach(function(t){r.async(i(t,this,e))})}};t.exports={Emitter:o,makeEmitter:function(){return r.aug(function(){},o)}}},function(t,e,n){var r=n(97),i=n(99),o=n(6),s=n(24),a=n(7),u=n(0),c=new i(function(t){var e=function(t){return t.reduce(function(t,e){return t[e._className]=t[e._className]||[],t[e._className].push(e),t},{})}(t.map(r.fromRawTask));u.forIn(e,function(t,e){s.allSettled(e.map(function(t){return t.initialize()})).then(function(){e.forEach(function(t){o.all([t.hydrate(),t.insertIntoDom()]).then(a(t.render,t)).then(a(t.success,t),a(t.fail,t))})})})});t.exports={addWidget:function(t){return c.add(t)}}},function(t,e,n){var r=n(18);t.exports=function(t){return r.write(function(){t&&t.parentNode&&t.parentNode.removeChild(t)})}},function(t,e,n){n(12),t.exports={log:function(t,e){}}},function(t,e,n){var r=n(1);function i(t){return(t=t||r).getSelection&&t.getSelection()}t.exports={getSelection:i,getSelectedText:function(t){var e=i(t);return e?e.toString():""}}},function(t,e,n){var r=n(4),i=n(1),o=n(2),s=2e4;t.exports=function(t){var e=new o,n=r.createElement("img");return n.onload=n.onerror=function(){i.setTimeout(e.resolve,50)},n.src=t,i.setTimeout(e.reject,s),e.promise}},function(t,e,n){var r=n(109);t.exports=function(t){t.define("createElement",r),t.define("createFragment",r),t.define("htmlToElement",r),t.define("hasSelectedText",r),t.define("addRootClass",r),t.define("removeRootClass",r),t.define("hasRootClass",r),t.define("prependStyleSheet",r),t.define("appendStyleSheet",r),t.define("prependCss",r),t.define("appendCss",r),t.define("makeVisible",r),t.define("injectWidgetEl",r),t.define("matchHeightToContent",r),t.define("matchWidthToContent",r)}},function(t,e){var n="tfw_horizon_tweet_embed_9555";t.exports={getBaseURLPath:function(t){switch(t&&t.tfw_team_holdback_11929&&t.tfw_team_holdback_11929.bucket){case"control":return"embed-holdback";case"holdback_prod":return"embed-holdback-prod";default:return"embed"}},isHorizonTweetEnabled:function(t){return!(t&&t[n]&&"control"===t[n].bucket)},isHorizonTimelineEnabled:function(){return!1}}},function(t,e){t.exports=function(t){var e,n=!1;return function(){return n?e:(n=!0,e=t.apply(this,arguments))}}},function(t,e,n){var r=n(15),i=n(116),o=n(117),s=n(16);t.exports=function(t,e,n){return new r(i,o,s.DM_BUTTON,t,e,n)}},function(t,e,n){var r=n(27),i=n(118);t.exports=r.build([i])},function(t,e,n){var r=n(15),i=n(121),o=n(32),s=n(16);t.exports=function(t,e,n){return new r(i,o,s.FOLLOW_BUTTON,t,e,n)}},function(t,e,n){var r=n(15),i=n(129),o=n(26),s=n(16);t.exports=function(t,e,n){return new r(i,o,s.MOMENT,t,e,n)}},function(t,e,n){var r=n(15),i=n(131),o=n(26),s=n(16);t.exports=function(t,e,n){return new r(i,o,s.PERISCOPE,t,e,n)}},function(t,e,n){var r=n(78),i=n(133),o=n(137),s=n(139),a=n(141),u=n(143),c={collection:i,event:o,likes:s,list:a,profile:u,url:l},d=[u,s,i,a,o];function l(t){return r(d,function(e){try{return new e(t)}catch(t){}})}t.exports=function(t){return t?function(t){var e,n;return e=(t.sourceType+"").toLowerCase(),(n=c[e])?new n(t):null}(t)||l(t):null}},function(t,e,n){var r=n(4),i=n(59),o=n(15),s=n(32),a=n(145),u=n(26),c=n(146),d=n(16);t.exports=function(t,e,n,l){var f;return i.isHorizonTimelineEnabled(l,t)?(f=r.createElement("div"),new o(a,s,d.TIMELINE,t,e,n,{sandboxWrapperEl:f})):new o(c,u,d.TIMELINE,t,e,n)}},function(t,e,n){var r=n(4),i=n(15),o=n(32),s=n(148),a=n(16);t.exports=function(t,e,n){var u=r.createElement("div");return new i(s,o,a.TWEET,t,e,n,{sandboxWrapperEl:u})}},function(t,e,n){var r=n(15),i=n(150),o=n(32),s=n(16);t.exports=function(t,e,n){var a=t&&t.type||"share",u="hashtag"==a?s.HASHTAG_BUTTON:"mention"==a?s.MENTION_BUTTON:s.SHARE_BUTTON;return new r(i,o,u,t,e,n)}},function(t,e,n){var r=n(42),i=n(38),o=n(0);t.exports=function(t){var e={widget_origin:i.rootDocumentLocation(),widget_frame:i.isFramed()?i.currentDocumentLocation():null,duration_ms:t.duration,item_ids:t.widgetIds||[]},n=o.aug(t.namespace,{page:"page",component:"performance"});r.scribe(n,e)}},function(t,e,n){var r=n(0),i=n(134),o=["ar","fa","he","ur"];t.exports={isRtlLang:function(t){return t=String(t).toLowerCase(),r.contains(o,t)},matchLanguage:function(t){return t=(t=(t||"").toLowerCase()).replace("_","-"),i(t)?t:(t=t.replace(/-.*/,""),i(t)?t:"en")}}},function(t){t.exports={tweetButtonHtmlPath:"/widgets/tweet_button.06c6ee58c3810956b7509218508c7b56.{{lang}}.html",followButtonHtmlPath:"/widgets/follow_button.06c6ee58c3810956b7509218508c7b56.{{lang}}.html",hubHtmlPath:"/widgets/hub.html",widgetIframeHtmlPath:"/widgets/widget_iframe.06c6ee58c3810956b7509218508c7b56.html",resourceBaseUrl:"https://platform.twitter.com"}},function(t){t.exports={TWEET:0,RETWEET:10,CUSTOM_TIMELINE:17,LIVE_VIDEO_EVENT:28,QUOTE_TWEET:23}},function(t,e,n){var r=n(3),i=n(95),o=n(23),s=n(11),a={favorite:["favorite","like"],follow:["follow"],like:["favorite","like"],retweet:["retweet"],tweet:["tweet"]};function u(t){this.srcEl=[],this.element=t}u.open=function(t,e,n){var u=(r.intentType(t)||"").toLowerCase();r.isTwitterURL(t)&&(function(t,e){i.open(t,e)}(t,n),e&&o.trigger("click",{target:e,region:"intent",type:"click",data:{}}),e&&a[u]&&a[u].forEach(function(n){o.trigger(n,{target:e,region:"intent",type:n,data:function(t,e){var n=s.decodeURL(e);switch(t){case"favorite":case"like":return{tweet_id:n.tweet_id};case"follow":return{screen_name:n.screen_name,user_id:n.user_id};case"retweet":return{source_tweet_id:n.tweet_id};default:return{}}}(u,t)})}))},t.exports=u},function(t,e,n){var r=n(4),i=n(9),o=n(3);function s(t,e){var n,r;return e=e||i,/^https?:\/\//.test(t)?t:/^\/\//.test(t)?e.protocol+t:(n=e.host+(e.port.length?":"+e.port:""),0!==t.indexOf("/")&&((r=e.pathname.split("/")).pop(),r.push(t),t="/"+r.join("/")),[e.protocol,"//",n,t].join(""))}t.exports={absolutize:s,getCanonicalURL:function(){for(var t,e=r.getElementsByTagName("link"),n=0;e[n];n++)if("canonical"==(t=e[n]).rel)return s(t.href)},getScreenNameFromPage:function(){for(var t,e,n,i=[r.getElementsByTagName("a"),r.getElementsByTagName("link")],s=0,a=0,u=/\bme\b/;t=i[s];s++)for(a=0;e=t[a];a++)if(u.test(e.rel)&&(n=o.screenName(e.href)))return n}}},function(t,e,n){var r=n(19),i=n(55),o=n(11),s=n(37),a=n(0),u=n(8).get("scribeCallback"),c=2083,d=[],l=o.url(s.CLIENT_EVENT_ENDPOINT,{dnt:0,l:""}),f=encodeURIComponent(l).length;function h(t,e,n,r,i){var o=!a.isObject(t),c=!!e&&!a.isObject(e);o||c||(u&&u(arguments),p(s.formatClientEventNamespace(t),s.formatClientEventData(e,n,r),s.CLIENT_EVENT_ENDPOINT,i))}function p(t,e,n,r){var u;n&&a.isObject(t)&&a.isObject(e)&&(i.log(t,e),u=s.flattenClientEventPayload(t,e),r=a.aug({},r,{l:s.stringify(u)}),u.dnt&&(r.dnt=1),w(o.url(n,r)))}function m(t,e,n,r){var i=!a.isObject(t),o=!!e&&!a.isObject(e);if(!i&&!o)return v(s.flattenClientEventPayload(s.formatClientEventNamespace(t),s.formatClientEventData(e,n,r)))}function v(t){return d.push(t),d}function g(t){return encodeURIComponent(t).length+3}function w(t){return(new Image).src=t}t.exports={canFlushOneItem:function(t){var e=g(s.stringify(t));return f+e<c},_enqueueRawObject:v,scribe:p,clientEvent:h,clientEvent2:function(t,e,n){return h(t,e,n,2)},enqueueClientEvent:m,flushClientEvents:function(){var t;return d.length>1&&m({page:"widgets_js",component:"scribe_pixel",action:"batch_log"},{}),t=d,d=[],t.reduce(function(e,n,r){var i=e.length,o=i&&e[i-1];return r+1==t.length&&n.event_namespace&&"batch_log"==n.event_namespace.action&&(n.message=["entries:"+r,"requests:"+i].join("/")),function t(e){return Array.isArray(e)||(e=[e]),e.reduce(function(e,n){var r,i=s.stringify(n),o=g(i);return f+o<c?e=e.concat(i):(r=s.splitLogEntry(n)).length>1&&(e=e.concat(t(r))),e},[])}(n).forEach(function(t){var n=g(t);(!o||o.urlLength+n>c)&&(o={urlLength:f,items:[]},e.push(o)),o.urlLength+=n,o.items.push(t)}),e},[]).map(function(t){var e={l:t.items};return r.enabled()&&(e.dnt=1),w(o.url(s.CLIENT_EVENT_ENDPOINT,e))})},interaction:function(t,e,n,r){var i=s.extractTermsFromDOM(t.target||t.srcElement);i.action=r||"click",h(i,e,n)}}},function(t,e,n){var r=n(0),i=n(45);t.exports=function(t,e){return i(t,e)?[t]:r.toRealArray(t.querySelectorAll(e))}},function(t,e){t.exports=function(t,e,n){for(var r,i=0;i<t.length;i++)if(r=e.call(n,t[i],i,t))return r}},function(t,e){t.exports=function(){return Math.floor(+new Date/9e5)}},function(t,e,n){var r=n(4),i=n(8),o=n(2),s=n(0),a=n(11),u="cb",c=0;t.exports={fetch:function(t,e,n,d){var l,f,h;return d=function(t){if(t)return t.replace(/[^\w$]/g,"_")}(d||u+c++),l=i.fullPath(["callbacks",d]),f=r.createElement("script"),h=new o,e=s.aug({},e,{callback:l,suppress_response_codes:!0}),i.set(["callbacks",d],function(t){var e;t=(e=n(t||!1)).resp,e.success?h.resolve(t):h.reject(t),f.onload=f.onreadystatechange=null,f.parentNode&&f.parentNode.removeChild(f),i.unset(["callbacks",d])}),f.onerror=function(){h.reject(new Error("failed to fetch "+f.src))},f.src=a.url(t,e),f.async="async",r.body.appendChild(f),h.promise}}},function(t,e,n){var r=n(12);t.exports=function(t){var e,n;return e=t.headers&&t.headers.status,!(n=t&&!t.error&&200===e)&&t.headers&&t.headers.message&&r.publicError(t.headers.message),{success:n,resp:t}}},function(t,e){t.exports={getTimezoneOffset:function(){var t=(new Date).toString().match(/(GMT[+-]?\d+)/);return t&&t[0]||"GMT"}}},,,,,,,,,function(t,e,n){var r,i=n(2),o=n(4),s=n(93),a=n(50),u=n(8),c=n(94),d=n(23),l=n(96),f=n(151),h=n(159),p=n(160),m=n(30),v=n(36);n(161),m.emitter.trigger(m.START),u.set("widgets.init",!0),a.set("init",!0),p(),r=new i,s.exposeReadyPromise(r.promise,a.base,"_e"),a.set("widgets",f),a.set("widgets.load",l.load),a.set("events",d),h(function(){v.load(),r.resolve(a.base),c.attachTo(o),l.loadPage()})},function(t,e){t.exports=navigator},function(t,e,n){var r=n(7);t.exports={exposeReadyPromise:function(t,e,n){e.ready=r(t.then,t),n&&Array.isArray(e[n])&&(e[n].forEach(r(t.then,t)),delete e[n])}}},function(t,e,n){var r=n(9),i=n(35),o=n(34),s=n(74),a=n(3);function u(t){var e,n,u;t.altKey||t.metaKey||t.shiftKey||(e=o.closest(function(t){return"A"===t.tagName||"AREA"===t.tagName},t.target))&&a.isIntentURL(e.href)&&(n=(n=(n=[u=e.href,"original_referer="+r.href].join(-1==u.indexOf("?")?"?":"&")).replace(/^http[:]/,"https:")).replace(/^\/\//,"https://"),s.open(n,e),i.preventDefault(t))}t.exports={attachTo:function(t){t.addEventListener("click",u,!1)}}},function(t,e,n){var r=n(1),i=n(35),o=n(34),s=n(21),a=n(3),u=n(22),c="intent_";function d(t){this.win=t}d.prototype.open=function(t,e){var n=e&&"click"==e.type&&o.closest("a",e.target),r=e&&(e.altKey||e.metaKey||e.shiftKey),d=n&&(s.ios()||s.android());if(a.isTwitterURL(t))return r||d?this:(this.name=c+u.generate(),this.popup=this.win.open(t,this.name),e&&i.preventDefault(e),this)},d.open=function(t,e){return new d(r).open(t,e)},t.exports=d},function(t,e,n){var r=n(4),i=n(6),o=n(24),s=n(53),a=n(33),u=n(8),c=n(42),d=n(23),l=n(5),f=n(0),h=n(36),p=n(113),m=n(30);function v(){var t=a.val("widgets:autoload")||!0;return!l.isFalseValue(t)&&(l.isTruthValue(t)?r.body:r.querySelectorAll(t))}function g(t,e){var n,i;return t=(t=t||r.body).length?f.toRealArray(t):[t],c.pause(),i=function(t,e){return t.reduce(function(t,n){return t.concat(p.reduce(function(t,r){return t.concat(r(n,e))},[]))},[])}(t,e),m.emitter.trigger(m.ALL_WIDGETS_RENDER_START,{widgets:i}),n=o.allResolved(i.map(function(t){return s.addWidget(t)})).then(function(t){d.trigger("loaded",{widgets:t}),t&&t.length&&m.emitter.trigger(m.ALL_WIDGETS_RENDER_END,{widgets:t})}),o.always(n,function(){c.resume()}),n}function w(t){return h.getExperiments().then(function(e){return g(t,e)})}t.exports={load:w,loadPage:function(){var t=v();return!1===t?i.resolve():(u.set("widgets.loaded",!0),w(t))},_getPageLoadTarget:v}},function(t,e,n){var r=n(10),i=n(18),o=n(23),s=n(54),a=n(6),u=n(24);function c(t,e){this._widget=null,this._sandbox=null,this._hydrated=!1,this._insertedIntoDom=!1,this._Sandbox=t.Sandbox,this._factory=t.factory,this._widgetParams=t.parameters,this._resolve=e,this._className=t.className,this._renderedClassName=t.className+"-rendered",this._errorClassName=t.className+"-error",this._srcEl=t.srcEl,this._targetGlobal=function(t){return(t.srcEl||t.targetEl).ownerDocument.defaultView}(t),this._sandboxWrapperEl=t.options?t.options.sandboxWrapperEl:null,this._insertionStrategy=function(e){var n,r=t.srcEl,i=t.targetEl,o=t.className,s=t.className+"-rendered",a=t.options?t.options.sandboxWrapperEl:null;a?(a.appendChild(e),a.classList.add(o,s),n=a):n=e,r?i.insertBefore(n,r):i.appendChild(n)}}c.fromRawTask=function(t){return new c(t.input,t.taskDoneDeferred.resolve)},c.prototype.initialize=function(){var t=this,e=new this._Sandbox(this._targetGlobal);return this._factory(this._widgetParams,e).then(function(n){return t._widget=n,t._sandbox=e,n._sandboxWrapperEl=t._sandboxWrapperEl,n})},c.prototype.insertIntoDom=function(){var t=this,e=this._sandboxWrapperEl?"":[this._className,this._renderedClassName].join(" ");return this._widget?this._sandbox.insert(this._widget.id,{class:e},null,this._insertionStrategy).then(function(){t._insertedIntoDom=!0}):a.reject(new Error("cannot insert widget into DOM before it is initialized"))},c.prototype.hydrate=function(){var t=this;return this._widget?this._widget.hydrate().then(function(){t._hydrated=!0}):a.reject(new Error("cannot hydrate widget before it is initialized"))},c.prototype.render=function(){var t=this;function e(e){var n=t._sandboxWrapperEl?t._sandboxWrapperEl:t._sandbox.sandboxEl;return s(n).then(function(){return a.reject(e)})}return this._hydrated?this._insertedIntoDom?t._widget.render(t._sandbox).then(function(){return t._sandbox.onResize(function(){return t._widget.resize().then(function(){var e=t._sandboxWrapperEl?t._sandboxWrapperEl:t._sandbox.sandboxEl;o.trigger("resize",{target:e})})}),t._widget.show()}).then(function(){return s(t._srcEl).then(function(){return t._sandbox.sandboxEl})},e):e(new Error("cannot render widget before DOM insertion")):e(new Error("cannot render widget before hydration"))},c.prototype.fail=function(){var t=this,e=t._errorClassName,n=t._sandboxWrapperEl?t._sandboxWrapperEl:t._srcEl;return this._srcEl?u.always(i.write(function(){r.add(n,e)}),function(){o.trigger("rendered",{target:n}),t._resolve(n)}):(t._resolve(),a.resolve())},c.prototype.success=function(){var t=this._sandboxWrapperEl?this._sandboxWrapperEl:this._sandbox.sandboxEl;o.trigger("rendered",{target:t}),this._resolve(t)},t.exports=c},function(t,e,n){var r;!function(){"use strict";var i=window.requestAnimationFrame||window.webkitRequestAnimationFrame||window.mozRequestAnimationFrame||window.msRequestAnimationFrame||function(t){return window.setTimeout(t,1e3/60)};function o(){this.frames=[],this.lastId=0,this.raf=i,this.batch={hash:{},read:[],write:[],mode:null}}o.prototype.read=function(t,e){var n=this.add("read",t,e),r=n.id;return this.batch.read.push(n.id),"reading"===this.batch.mode||this.batch.scheduled?r:(this.scheduleBatch(),r)},o.prototype.write=function(t,e){var n=this.add("write",t,e),r=this.batch.mode,i=n.id;return this.batch.write.push(n.id),"writing"===r||"reading"===r||this.batch.scheduled?i:(this.scheduleBatch(),i)},o.prototype.defer=function(t,e,n){"function"==typeof t&&(n=e,e=t,t=1);var r=this,i=t-1;return this.schedule(i,function(){r.run({fn:e,ctx:n})})},o.prototype.clear=function(t){if("function"==typeof t)return this.clearFrame(t);t=Number(t);var e=this.batch.hash[t];if(e){var n=this.batch[e.type],r=n.indexOf(t);delete this.batch.hash[t],~r&&n.splice(r,1)}},o.prototype.clearFrame=function(t){var e=this.frames.indexOf(t);~e&&this.frames.splice(e,1)},o.prototype.scheduleBatch=function(){var t=this;this.schedule(0,function(){t.batch.scheduled=!1,t.runBatch()}),this.batch.scheduled=!0},o.prototype.uniqueId=function(){return++this.lastId},o.prototype.flush=function(t){for(var e;e=t.shift();)this.run(this.batch.hash[e])},o.prototype.runBatch=function(){try{this.batch.mode="reading",this.flush(this.batch.read),this.batch.mode="writing",this.flush(this.batch.write),this.batch.mode=null}catch(t){throw this.runBatch(),t}},o.prototype.add=function(t,e,n){var r=this.uniqueId();return this.batch.hash[r]={id:r,fn:e,ctx:n,type:t}},o.prototype.run=function(t){var e=t.ctx||this,n=t.fn;if(delete this.batch.hash[t.id],!this.onError)return n.call(e);try{n.call(e)}catch(t){this.onError(t)}},o.prototype.loop=function(){var t,e=this,n=this.raf,r=!1;function i(){var t=e.frames.shift();e.frames.length?n(i):e.looping=!1,t&&t()}this.looping||(t=setTimeout(function(){r=!0,i()},500),n(function(){r||(clearTimeout(t),i())}),this.looping=!0)},o.prototype.schedule=function(t,e){return this.frames[t]?this.schedule(t+1,e):(this.loop(),this.frames[t]=e)};var s=new o;void 0!==t&&t.exports?t.exports=s:void 0===(r=function(){return s}.call(e,n,e,t))||(t.exports=r)}()},function(t,e,n){var r=n(2),i=n(100),o=n(7);function s(t){this._inputsQueue=[],this._task=t,this._hasFlushBeenScheduled=!1}s.prototype.add=function(t){var e=new r;return this._inputsQueue.push({input:t,taskDoneDeferred:e}),this._hasFlushBeenScheduled||(this._hasFlushBeenScheduled=!0,i(o(this._flush,this))),e.promise},s.prototype._flush=function(){try{this._task.call(null,this._inputsQueue)}catch(t){this._inputsQueue.forEach(function(e){e.taskDoneDeferred.reject(t)})}this._inputsQueue=[],this._hasFlushBeenScheduled=!1},t.exports=s},function(t,e,n){var r=n(49).Promise;t.exports=r._asap},function(t,e,n){var r,i,o,s=n(4),a=n(1),u=n(31),c=n(19),d=n(2),l=n(6),f=n(55),h=n(37),p=n(0),m=n(26),v=n(8).get("scribeCallback"),g=Math.floor(1e3*Math.random())+"_",w="rufous-frame-"+g+"-",y="rufous-form-"+g+"-",b=0,_=!1,E=new d;function x(){var t=o.createElement("form"),e=o.createElement("input"),n=o.createElement("input");return b++,t.action=h.CLIENT_EVENT_ENDPOINT,t.method="POST",t.target=w+b,t.id=y+b,e.type="hidden",e.name="dnt",e.value=c.enabled(),n.type="hidden",n.name="tfw_redirect",n.value=h.RUFOUS_REDIRECT,t.appendChild(e),t.appendChild(n),t}function T(){var t=w+b;return u({id:t,name:t,width:0,height:0,border:0},{display:"none"},o.doc)}t.exports={clientEvent:function(t,e,n,i){(function(t,e){var n=!p.isObject(t),r=!!e&&!p.isObject(e),i=n||r;return i})(t,e)||(v&&v(arguments),E.promise.then(function(){!function(t,e){var n,i,s;p.isObject(t)&&p.isObject(e)&&(f.log(t,e),s=h.flattenClientEventPayload(t,e),(n=r.firstChild).value=+(+n.value||s.dnt||0),(i=o.createElement("input")).type="hidden",i.name="l",i.value=h.stringify(s),r.appendChild(i))}(h.formatClientEventNamespace(t),h.formatClientEventData(e,n,i))}))},flush:function(){return E.promise.then(function(){var t;return r.children.length<=2?l.reject():(t=l.all([o.doc.body.appendChild(r),o.doc.body.appendChild(i)]).then(function(t){var e=t[0],n=t[1];return n.addEventListener("load",function(){!function(t,e){return function(){var n=t.parentNode;n&&(n.removeChild(t),n.removeChild(e))}}(e,n)()}),e.submit(),t}),r=x(),i=T(),t)})},init:function(){return _?E.promise:((o=new m(a)).insert("rufous-sandbox",null,{display:"none"},function(t){s.body.appendChild(t)}).then(function(){o.setTitle("Twitter analytics iframe"),r=x(),i=T(),E.resolve([r,i])}),_=!0,E.promise)}}},function(t,e,n){var r=n(9),i=/^[^#?]*\.(gov|mil)(:\d+)?([#?].*)?$/i,o={};function s(t){return t in o?o[t]:o[t]=i.test(t)}t.exports={isUrlSensitive:s,isHostPageSensitive:function(){return s(r.host)}}},function(t,e,n){var r=n(0);t.exports=function t(e){var n=[];return e.forEach(function(e){var i=r.isType("array",e)?t(e):[e];n=n.concat(i)}),n}},function(t,e){t.exports=function(t){return t.filter(function(e,n){return t.indexOf(e)===n})}},function(t,e,n){var r=n(40),i=n(0),o=n(107);function s(){r.apply(this,arguments)}s.prototype=Object.create(r.prototype),i.aug(s.prototype,{factory:o}),t.exports=s},function(t,e,n){var r=n(24),i=n(0),o=n(7);t.exports={before:function(t,e){return function(){var n,i=this,o=arguments;return n=e.apply(this,arguments),r.isPromise(n)?n.then(function(){return t.apply(i,o)}):t.apply(this,arguments)}},after:function(t,e){return function(){var n,i=this,o=arguments;function s(t,e){return r.isPromise(e)?e.then(function(){return t}):t}return n=t.apply(this,arguments),r.isPromise(n)?n.then(function(t){return s(t,e.apply(i,o))}):s(n,e.apply(this,arguments))}},around:function(t,e){return function(){var n=i.toRealArray(arguments);return n.unshift(o(t,this)),e.apply(this,n)}}}},function(t,e,n){var r=n(10),i=n(18),o=n(41),s=n(6),a=n(0);t.exports=function(){var t=o();function e(e){t.apply(this,arguments),Object.defineProperty(this,"targetGlobal",{value:e})}return e.prototype=Object.create(t.prototype),a.aug(e.prototype,{id:null,initialized:!1,width:0,height:0,sandboxEl:null,insert:function(){return s.reject()},onResize:function(){},addClass:function(t){var e=this.sandboxEl;return t=Array.isArray(t)?t:[t],i.write(function(){t.forEach(function(t){r.add(e,t)})})},removeClass:function(t){var e=this.sandboxEl;return t=Array.isArray(t)?t:[t],i.write(function(){t.forEach(function(t){r.remove(e,t)})})},styleSelf:function(t){var e=this;return i.write(function(){a.forIn(t,function(t,n){e.sandboxEl.style[t]=n})})}}),e}},function(t,e,n){var r=n(4),i=n(10),o=n(18),s=n(56),a=n(27),u=n(57),c=n(43),d=n(44),l=n(31),f=n(12),h=n(47),p=n(2),m=n(6),v=n(0),g=n(8),w=n(22),y=n(7),b={allowfullscreen:"true"},_={position:"absolute",visibility:"hidden",display:"block",width:"0px",height:"0px",padding:"0",border:"none"},E={position:"static",visibility:"visible"},x="SandboxRoot",T=".SandboxRoot { display: none; }",A=50;function S(t,e,n,r){return e=v.aug({id:t},b,e),n=v.aug({},_,n),l(e,n,r)}function R(t,e,n,i,s){var a=new p,u=w.generate(),c=S(t,e,n,s);return g.set(["sandbox",u],function(){var t=c.contentWindow.document;o.write(function(){t.write("<!DOCTYPE html><html><head></head><body></body></html>")}).then(function(){t.close(),a.resolve(c)})}),c.src=["javascript:",'document.write("");',"try { window.parent.document; }",'catch (e) { document.domain="'+r.domain+'"; }',"window.parent."+g.fullPath(["sandbox",u])+"();"].join(""),c.addEventListener("error",a.reject,!1),o.write(function(){i.parentNode.replaceChild(c,i)}),a.promise}t.exports=a.couple(n(58),function(t){t.overrideProperty("id",{get:function(){return this.sandboxEl&&this.sandboxEl.id}}),t.overrideProperty("initialized",{get:function(){return!!this.win}}),t.overrideProperty("width",{get:function(){return this._width}}),t.overrideProperty("height",{get:function(){return this._height}}),t.overrideProperty("sandboxEl",{get:function(){return this.iframeEl}}),t.defineProperty("iframeEl",{get:function(){return this._iframe}}),t.defineProperty("rootEl",{get:function(){return this.doc&&this.doc.documentElement}}),t.defineProperty("widgetEl",{get:function(){return this.doc&&this.doc.body.firstElementChild}}),t.defineProperty("win",{get:function(){return this.iframeEl&&this.iframeEl.contentWindow}}),t.defineProperty("doc",{get:function(){return this.win&&this.win.document}}),t.define("_updateCachedDimensions",function(){var t=this;return o.read(function(){var e,n=h(t.sandboxEl);"visible"==t.sandboxEl.style.visibility?t._width=n.width:(e=h(t.sandboxEl.parentElement).width,t._width=Math.min(n.width,e)),t._height=n.height})}),t.define("_setTargetToBlank",function(){var t=this.createElement("base");t.target="_blank",this.doc.head.appendChild(t)}),t.define("_didResize",function(){var t=this,e=this._resizeHandlers.slice(0);return this._updateCachedDimensions().then(function(){e.forEach(function(e){e(t)})})}),t.define("setTitle",function(t){this.iframeEl.title=t}),t.override("createElement",function(t){return this.doc.createElement(t)}),t.override("createFragment",function(){return this.doc.createDocumentFragment()}),t.override("htmlToElement",function(t){var e;return(e=this.createElement("div")).innerHTML=t,e.firstElementChild}),t.override("hasSelectedText",function(){return!!s.getSelectedText(this.win)}),t.override("addRootClass",function(t){var e=this.rootEl;return t=Array.isArray(t)?t:[t],this.initialized?o.write(function(){t.forEach(function(t){i.add(e,t)})}):m.reject(new Error("sandbox not initialized"))}),t.override("removeRootClass",function(t){var e=this.rootEl;return t=Array.isArray(t)?t:[t],this.initialized?o.write(function(){t.forEach(function(t){i.remove(e,t)})}):m.reject(new Error("sandbox not initialized"))}),t.override("hasRootClass",function(t){return i.present(this.rootEl,t)}),t.define("addStyleSheet",function(t,e){var n,r=new p;return this.initialized?((n=this.createElement("link")).type="text/css",n.rel="stylesheet",n.href=t,n.addEventListener("load",r.resolve,!1),n.addEventListener("error",r.reject,!1),o.write(y(e,null,n)).then(function(){return u(t).then(r.resolve,r.reject),r.promise})):m.reject(new Error("sandbox not initialized"))}),t.override("prependStyleSheet",function(t){var e=this.doc;return this.addStyleSheet(t,function(t){var n=e.head.firstElementChild;return n?e.head.insertBefore(t,n):e.head.appendChild(t)})}),t.override("appendStyleSheet",function(t){var e=this.doc;return this.addStyleSheet(t,function(t){return e.head.appendChild(t)})}),t.define("addCss",function(t,e){var n;return c.inlineStyle()?((n=this.createElement("style")).type="text/css",n.appendChild(this.doc.createTextNode(t)),o.write(y(e,null,n))):(f.devError("CSP enabled; cannot embed inline styles"),m.resolve())}),t.override("prependCss",function(t){var e=this.doc;return this.addCss(t,function(t){var n=e.head.firstElementChild;return n?e.head.insertBefore(t,n):e.head.appendChild(t)})}),t.override("appendCss",function(t){var e=this.doc;return this.addCss(t,function(t){return e.head.appendChild(t)})}),t.override("makeVisible",function(){var t=this;return this.styleSelf(E).then(function(){t._updateCachedDimensions()})}),t.override("injectWidgetEl",function(t){var e=this;return this.initialized?this.widgetEl?m.reject(new Error("widget already injected")):o.write(function(){e.doc.body.appendChild(t)}):m.reject(new Error("sandbox not initialized"))}),t.override("matchHeightToContent",function(){var t,e=this;return o.read(function(){t=e.widgetEl?h(e.widgetEl).height:0}),o.write(function(){e.sandboxEl.style.height=t+"px"}).then(function(){return e._updateCachedDimensions()})}),t.override("matchWidthToContent",function(){var t,e=this;return o.read(function(){t=e.widgetEl?h(e.widgetEl).width:0}),o.write(function(){e.sandboxEl.style.width=t+"px"}).then(function(){return e._updateCachedDimensions()})}),t.after("initialize",function(){this._iframe=null,this._width=this._height=0,this._resizeHandlers=[]}),t.override("insert",function(t,e,n,r){var i=this,s=new p,a=this.targetGlobal.document,u=S(t,e,n,a);return o.write(y(r,null,u)),u.addEventListener("load",function(){(function(t){try{t.contentWindow.document}catch(t){return m.reject(t)}return m.resolve(t)})(u).then(null,y(R,null,t,e,n,u,a)).then(s.resolve,s.reject)},!1),u.addEventListener("error",s.reject,!1),s.promise.then(function(t){var e=d(i._didResize,A,i);return i._iframe=t,i.win.addEventListener("resize",e,!1),m.all([i._setTargetToBlank(),i.addRootClass(x),i.prependCss(T)])})}),t.override("onResize",function(t){this._resizeHandlers.push(t)}),t.after("styleSelf",function(){return this._updateCachedDimensions()})})},function(t,e){t.exports=function(){throw new Error("unimplemented method")}},function(t,e,n){var r=n(2),i=n(7),o=100,s=3e3;function a(t,e){this._inputsQueue=[],this._task=t,this._isPaused=!1,this._flushDelay=e&&e.flushDelay||o,this._pauseLength=e&&e.pauseLength||s,this._flushTimeout=void 0}a.prototype.add=function(t){var e=new r;return this._inputsQueue.push({input:t,taskDoneDeferred:e}),this._scheduleFlush(),e.promise},a.prototype._scheduleFlush=function(){this._isPaused||(clearTimeout(this._flushTimeout),this._flushTimeout=setTimeout(i(this._flush,this),this._flushDelay))},a.prototype._flush=function(){try{this._task.call(null,this._inputsQueue)}catch(t){this._inputsQueue.forEach(function(e){e.taskDoneDeferred.reject(t)})}this._inputsQueue=[],this._flushTimeout=void 0},a.prototype.pause=function(t){clearTimeout(this._flushTimeout),this._isPaused=!0,!t&&this._pauseLength&&setTimeout(i(this.resume,this),this._pauseLength)},a.prototype.resume=function(){this._isPaused=!1,this._scheduleFlush()},t.exports=a},function(t,e,n){var r=n(72),i=n(31),o=n(2),s=n(4),a=n(20),u=n(21),c=n(25),d=n(9),l=n(12),f=n(112),h=n(60),p=n(8),m=n(11),v=n(3),g=n(0),w=n(1),y=h(function(){return new o}),b={shouldObtainCookieConsent:!1,features:{}};t.exports={load:function(){var t,e,n,o;if(u.ie9()||u.ie10()||"http:"!==d.protocol&&"https:"!==d.protocol)return l.devError("Using default settings due to unsupported browser or protocol."),void y().resolve();t={origin:d.origin},a.settings().indexOf("localhost")>-1&&(t.localSettings=!0),e=m.url(r.resourceBaseUrl+r.widgetIframeHtmlPath,t),n=function(t){var n,r,i,o;if(r=v.isTwitterURL(t.origin),i=e.substr(0,t.origin.length)===t.origin,o=v.isTwimgURL(t.origin),i&&r||o)try{(n="string"==typeof t.data?c.parse(t.data):t.data).namespace===f.settings&&(b=g.aug(b,{features:n.settings.features,sessionId:n.sessionId}),y().resolve())}catch(t){l.devError(t)}},w.addEventListener("message",n),o=i({src:e,title:"Twitter settings iframe"},{display:"none"}),s.body.appendChild(o)},settingsLoaded:function(){var t,e;return t=p.get("experimentOverride"),y().promise.then(function(){return t&&t.name&&t.assignment&&((e={})[t.name]={bucket:t.assignment},b.features=g.aug(b.features,e)),b})}}},function(t,e){t.exports={settings:"twttr.settings"}},function(t,e,n){t.exports=[n(114),n(120),n(128),n(130),n(132),n(147),n(149)]},function(t,e,n){var r=n(11),i=n(5),o=n(0),s=n(13),a=n(14)(),u=n(61),c="a.twitter-dm-button";t.exports=function(t){return a(t,c).map(function(t){return u(function(t){var e=t.getAttribute("data-show-screen-name"),n=s(t),a=t.getAttribute("href"),u=t.getAttribute("data-screen-name"),c=e?i.asBoolean(e):null,d=t.getAttribute("data-size"),l=r.decodeURL(a),f=l.recipient_id,h=t.getAttribute("data-text")||l.text,p=t.getAttribute("data-welcome-message-id")||l.welcomeMessageId;return o.aug(n,{screenName:u,showScreenName:c,size:d,text:h,userId:f,welcomeMessageId:p})}(t),t.parentNode,t)})}},function(t,e,n){var r=n(0);t.exports=function t(e){var n;if(e)return n=e.lang||e.getAttribute("data-lang"),r.isType("string",n)?n:t(e.parentElement)}},function(t,e,n){var r=n(2);t.exports=function(t,e){var i=new r;return n.e(2).then(function(r){var o;try{o=n(83),i.resolve(new o(t,e))}catch(t){i.reject(t)}}.bind(null,n)).catch(function(t){i.reject(t)}),i.promise}},function(t,e,n){var r=n(62),i=n(26);t.exports=r.isSupported()?r:i},function(t,e,n){var r=n(119),i=n(1),o=n(10),s=n(35),a=n(18),u=n(56),c=n(27),d=n(57),l=n(43),f=n(47),h=n(7),p=n(44),m=n(6),v=n(0),g=50,w={position:"absolute",visibility:"hidden",display:"block",transform:"rotate(0deg)"},y={position:"static",visibility:"visible"},b="twitter-widget",_="open",E="SandboxRoot",x=".SandboxRoot { display: none; max-height: 10000px; }";t.exports=c.couple(n(58),function(t){t.defineStatic("isSupported",function(){return!!i.HTMLElement.prototype.attachShadow&&l.inlineStyle()}),t.overrideProperty("id",{get:function(){return this.sandboxEl&&this.sandboxEl.id}}),t.overrideProperty("initialized",{get:function(){return!!this._shadowHost}}),t.overrideProperty("width",{get:function(){return this._width}}),t.overrideProperty("height",{get:function(){return this._height}}),t.overrideProperty("sandboxEl",{get:function(){return this._shadowHost}}),t.define("_updateCachedDimensions",function(){var t=this;return a.read(function(){var e,n=f(t.sandboxEl);"visible"==t.sandboxEl.style.visibility?t._width=n.width:(e=f(t.sandboxEl.parentElement).width,t._width=Math.min(n.width,e)),t._height=n.height})}),t.define("_didResize",function(){var t=this,e=this._resizeHandlers.slice(0);return this._updateCachedDimensions().then(function(){e.forEach(function(e){e(t)})})}),t.override("createElement",function(t){return this.targetGlobal.document.createElement(t)}),t.override("createFragment",function(){return this.targetGlobal.document.createDocumentFragment()}),t.override("htmlToElement",function(t){var e;return(e=this.createElement("div")).innerHTML=t,e.firstElementChild}),t.override("hasSelectedText",function(){return!!u.getSelectedText(this.targetGlobal)}),t.override("addRootClass",function(t){var e=this._shadowRootBody;return t=Array.isArray(t)?t:[t],this.initialized?a.write(function(){t.forEach(function(t){o.add(e,t)})}):m.reject(new Error("sandbox not initialized"))}),t.override("removeRootClass",function(t){var e=this._shadowRootBody;return t=Array.isArray(t)?t:[t],this.initialized?a.write(function(){t.forEach(function(t){o.remove(e,t)})}):m.reject(new Error("sandbox not initialized"))}),t.override("hasRootClass",function(t){return o.present(this._shadowRootBody,t)}),t.override("addStyleSheet",function(t,e){return this.addCss('@import url("'+t+'");',e).then(function(){return d(t)})}),t.override("prependStyleSheet",function(t){var e=this._shadowRoot;return this.addStyleSheet(t,function(t){var n=e.firstElementChild;return n?e.insertBefore(t,n):e.appendChild(t)})}),t.override("appendStyleSheet",function(t){var e=this._shadowRoot;return this.addStyleSheet(t,function(t){return e.appendChild(t)})}),t.override("addCss",function(t,e){var n;return this.initialized?l.inlineStyle()?((n=this.createElement("style")).type="text/css",n.appendChild(this.targetGlobal.document.createTextNode(t)),a.write(h(e,null,n))):m.resolve():m.reject(new Error("sandbox not initialized"))}),t.override("prependCss",function(t){var e=this._shadowRoot;return this.addCss(t,function(t){var n=e.firstElementChild;return n?e.insertBefore(t,n):e.appendChild(t)})}),t.override("appendCss",function(t){var e=this._shadowRoot;return this.addCss(t,function(t){return e.appendChild(t)})}),t.override("makeVisible",function(){return this.styleSelf(y)}),t.override("injectWidgetEl",function(t){var e=this;return this.initialized?this._shadowRootBody.firstElementChild?m.reject(new Error("widget already injected")):a.write(function(){e._shadowRootBody.appendChild(t)}).then(function(){return e._updateCachedDimensions()}).then(function(){var t=p(e._didResize,g,e);new r(e._shadowRootBody,t)}):m.reject(new Error("sandbox not initialized"))}),t.override("matchHeightToContent",function(){return m.resolve()}),t.override("matchWidthToContent",function(){return m.resolve()}),t.override("insert",function(t,e,n,r){var i=this.targetGlobal.document,o=this._shadowHost=i.createElement(b),u=this._shadowRoot=o.attachShadow({mode:_}),c=this._shadowRootBody=i.createElement("div");return v.forIn(e||{},function(t,e){o.setAttribute(t,e)}),o.id=t,u.appendChild(c),s.delegate(c,"click","A",function(t,e){e.hasAttribute("target")||e.setAttribute("target","_blank")}),m.all([this.styleSelf(w),this.addRootClass(E),this.prependCss(x),a.write(r.bind(null,o))])}),t.override("onResize",function(t){this._resizeHandlers.push(t)}),t.after("initialize",function(){this._shadowHost=this._shadowRoot=this._shadowRootBody=null,this._width=this._height=0,this._resizeHandlers=[]}),t.after("styleSelf",function(){return this._updateCachedDimensions()})})},function(t,e){var n;(n=function(t,e){function r(t,e){if(t.resizedAttached){if(t.resizedAttached)return void t.resizedAttached.add(e)}else t.resizedAttached=new function(){var t,e;this.q=[],this.add=function(t){this.q.push(t)},this.call=function(){for(t=0,e=this.q.length;t<e;t++)this.q[t].call()}},t.resizedAttached.add(e);t.resizeSensor=document.createElement("div"),t.resizeSensor.className="resize-sensor";var n="position: absolute; left: 0; top: 0; right: 0; bottom: 0; overflow: hidden; z-index: -1; visibility: hidden;",r="position: absolute; left: 0; top: 0; transition: 0s;";t.resizeSensor.style.cssText=n,t.resizeSensor.innerHTML='<div class="resize-sensor-expand" style="'+n+'"><div style="'+r+'"></div></div><div class="resize-sensor-shrink" style="'+n+'"><div style="'+r+' width: 200%; height: 200%"></div></div>',t.appendChild(t.resizeSensor),{fixed:1,absolute:1}[function(t,e){return t.currentStyle?t.currentStyle[e]:window.getComputedStyle?window.getComputedStyle(t,null).getPropertyValue(e):t.style[e]}(t,"position")]||(t.style.position="relative");var i,o,s=t.resizeSensor.childNodes[0],a=s.childNodes[0],u=t.resizeSensor.childNodes[1],c=(u.childNodes[0],function(){a.style.width=s.offsetWidth+10+"px",a.style.height=s.offsetHeight+10+"px",s.scrollLeft=s.scrollWidth,s.scrollTop=s.scrollHeight,u.scrollLeft=u.scrollWidth,u.scrollTop=u.scrollHeight,i=t.offsetWidth,o=t.offsetHeight});c();var d=function(t,e,n){t.attachEvent?t.attachEvent("on"+e,n):t.addEventListener(e,n)},l=function(){t.offsetWidth==i&&t.offsetHeight==o||t.resizedAttached&&t.resizedAttached.call(),c()};d(s,"scroll",l),d(u,"scroll",l)}var i=Object.prototype.toString.call(t),o="[object Array]"===i||"[object NodeList]"===i||"[object HTMLCollection]"===i||"undefined"!=typeof jQuery&&t instanceof jQuery||"undefined"!=typeof Elements&&t instanceof Elements;if(o)for(var s=0,a=t.length;s<a;s++)r(t[s],e);else r(t,e);this.detach=function(){if(o)for(var e=0,r=t.length;e<r;e++)n.detach(t[e]);else n.detach(t)}}).detach=function(t){t.resizeSensor&&(t.removeChild(t.resizeSensor),delete t.resizeSensor,delete t.resizedAttached)},void 0!==t&&void 0!==t.exports?t.exports=n:window.ResizeSensor=n},function(t,e,n){var r=n(3),i=n(0),o=n(13),s=n(14)(),a=n(63),u=n(5),c="a.twitter-follow-button";t.exports=function(t){return s(t,c).map(function(t){return a(function(t){var e=o(t),n={screenName:r.screenName(t.href),showScreenName:"false"!==t.getAttribute("data-show-screen-name"),showCount:"false"!==t.getAttribute("data-show-count"),size:t.getAttribute("data-size"),count:t.getAttribute("data-count"),preview:t.getAttribute("data-preview")};return i.forIn(n,function(t,n){var r=e[t];e[t]=u.hasValue(r)?r:n}),e.screenName=e.screenName||e.screen_name,e}(t),t.parentNode,t)})}},function(t,e,n){var r=n(2);t.exports=function(t,e){var i=new r;return n.e(3).then(function(r){var o;try{o=n(84),i.resolve(new o(t,e))}catch(t){i.reject(t)}}.bind(null,n)).catch(function(t){i.reject(t)}),i.promise}},function(t,e,n){var r=n(18),i=n(123),o=n(60),s=n(31),a=n(2),u=n(6),c=n(7),d=n(0),l={allowfullscreen:"true"},f={position:"absolute",visibility:"hidden",width:"0px",height:"0px"},h={position:"static",visibility:"visible"},p={};i(function(t,e,n){var r=p[t];if(r)return e=e||1,n=n||1,r.styleSelf({width:e+"px",height:n+"px"}).then(function(){r.didResize()})},function(t){var e=p[t];if(e)return e._results.resolve()},function(t){var e=p[t];if(e)return e._rendered.resolve()},function(t,e){var n=p[t];n&&e&&n.setIframeVersion(e)},function(t){var e=p[t];if(e)return e._results.reject()}),t.exports=function(t){t.overrideProperty("id",{get:function(){return this.sandboxEl&&this.sandboxEl.id}}),t.overrideProperty("initialized",{get:function(){return!!this.iframeEl}}),t.overrideProperty("width",{get:function(){return this._width}}),t.overrideProperty("height",{get:function(){return this._height}}),t.overrideProperty("sandboxEl",{get:function(){return this.iframeEl}}),t.defineProperty("iframeEl",{get:function(){return this._iframe}}),t.defineProperty("iframeVersion",{get:function(){return this._iframeVersion}}),t.define("updateCachedDimensions",function(){var t=this;return this.initialized?r.read(function(){t._width=t.sandboxEl.offsetWidth,t._height=t.sandboxEl.offsetHeight}):u.resolve()}),t.define("setTitle",function(t){this.iframeEl.title=t}),t.define("setWaitToSwapUntilRendered",function(t){this._waitToSwapUntilRendered=t}),t.define("setIframeVersion",function(t){this._iframeVersion=t}),t.define("getResultsPromise",function(){return this._results.promise}),t.define("getRenderedPromise",function(){return this._rendered.promise}),t.define("makeVisible",function(){return this.styleSelf(h)}),t.define("didResize",function(){var t=this,e=t._resizeHandlers.length>0;return this.updateCachedDimensions().then(function(){e&&t._resizeHandlers.forEach(function(e){e(t)})})}),t.define("loadDocument",function(t){var e=new a;return this.initialized?this.iframeEl.src?u.reject(new Error("widget already loaded")):(this.iframeEl.addEventListener("load",e.resolve,!1),this.iframeEl.addEventListener("error",e.reject,!1),this.iframeEl.src=t,e.promise):u.reject(new Error("sandbox not initialized"))}),t.after("initialize",function(){var t=new a,e=new a;this._iframe=null,this._iframeVersion=null,this._width=this._height=0,this._resizeHandlers=[],this._rendered=t,this._results=e,this._waitToSwapUntilRendered=!1}),t.override("insert",function(t,e,n,i){var a=this;return e=d.aug({id:t},l,e),n=d.aug({},f,n),this._iframe=s(e,n),p[t]=this,a._waitToSwapUntilRendered||this.onResize(o(function(){a.makeVisible()})),r.write(c(i,null,this._iframe))}),t.override("onResize",function(t){this._resizeHandlers.push(t)}),t.after("styleSelf",function(){return this.updateCachedDimensions()})}},function(t,e,n){var r=n(1),i=n(124),o=n(126),s=n(23),a=n(5),u=n(127);t.exports=function(t,e,n,c,d){function l(t){var e=u(this);s.trigger(t.type,{target:e,region:t.region,type:t.type,data:t.data||{}})}function f(e){var n=u(this),r=n&&n.id,i=a.asInt(e.width),o=a.asInt(e.height);r&&void 0!==i&&void 0!==o&&t(r,i,o)}(new i).attachReceiver(new o.Receiver(r,"twttr.button")).bind("twttr.private.trigger",l).bind("twttr.private.resizeButton",f),(new i).attachReceiver(new o.Receiver(r,"twttr.embed")).bind("twttr.private.initialized",function(t){var e=u(this),n=e&&e.id,r=t.iframe_version;n&&r&&c&&c(n,r)}).bind("twttr.private.trigger",l).bind("twttr.private.results",function(){var t=u(this),n=t&&t.id;n&&e&&e(n)}).bind("twttr.private.rendered",function(){var t=u(this),e=t&&t.id;e&&n&&n(e)}).bind("twttr.private.no_results",function(){var t=u(this),e=t&&t.id;e&&d&&d(e)}).bind("twttr.private.resize",f)}},function(t,e,n){var r=n(25),i=n(125),o=n(0),s=n(6),a=n(24),u="2.0";function c(t){this.registry=t||{}}function d(t){var e,n;return e=o.isType("string",t),n=o.isType("number",t),e||n||null===t}function l(t,e){return{jsonrpc:u,id:d(t)?t:null,error:e}}c.prototype._invoke=function(t,e){var n,r,i;n=this.registry[t.method],r=t.params||[],r=o.isType("array",r)?r:[r];try{i=n.apply(e.source||null,r)}catch(t){i=s.reject(t.message)}return a.isPromise(i)?i:s.resolve(i)},c.prototype._processRequest=function(t,e){var n,r;return function(t){var e,n,r;return!!o.isObject(t)&&(e=t.jsonrpc===u,n=o.isType("string",t.method),r=!("id"in t)||d(t.id),e&&n&&r)}(t)?(n="params"in t&&(r=t.params,!o.isObject(r)||o.isType("function",r))?s.resolve(l(t.id,i.INVALID_PARAMS)):this.registry[t.method]?this._invoke(t,{source:e}).then(function(e){return n=t.id,{jsonrpc:u,id:n,result:e};var n},function(){return l(t.id,i.INTERNAL_ERROR)}):s.resolve(l(t.id,i.METHOD_NOT_FOUND)),null!=t.id?n:s.resolve()):s.resolve(l(t.id,i.INVALID_REQUEST))},c.prototype.attachReceiver=function(t){return t.attachTo(this),this},c.prototype.bind=function(t,e){return this.registry[t]=e,this},c.prototype.receive=function(t,e){var n,a,u,c=this;try{u=t,t=o.isType("string",u)?r.parse(u):u}catch(t){return s.resolve(l(null,i.PARSE_ERROR))}return e=e||null,a=((n=o.isType("array",t))?t:[t]).map(function(t){return c._processRequest(t,e)}),n?function(t){return s.all(t).then(function(t){return(t=t.filter(function(t){return void 0!==t})).length?t:void 0})}(a):a[0]},t.exports=c},function(t){t.exports={PARSE_ERROR:{code:-32700,message:"Parse error"},INVALID_REQUEST:{code:-32600,message:"Invalid Request"},INVALID_PARAMS:{code:-32602,message:"Invalid params"},METHOD_NOT_FOUND:{code:-32601,message:"Method not found"},INTERNAL_ERROR:{code:-32603,message:"Internal error"}}},function(t,e,n){var r=n(9),i=n(1),o=n(25),s=n(2),a=n(21),u=n(0),c=n(3),d=n(7),l=a.ie9();function f(t,e,n){var r;t&&t.postMessage&&(l?r=(n||"")+o.stringify(e):n?(r={})[n]=e:r=e,t.postMessage(r,"*"))}function h(t){return u.isType("string",t)?t:"JSONRPC"}function p(t,e){return e?u.isType("string",t)&&0===t.indexOf(e)?t.substring(e.length):t&&t[e]?t[e]:void 0:t}function m(t,e){var n=t.document;this.filter=h(e),this.server=null,this.isTwitterFrame=c.isTwitterURL(n.location.href),t.addEventListener("message",d(this._onMessage,this),!1)}function v(t,e){this.pending={},this.target=t,this.isTwitterHost=c.isTwitterURL(r.href),this.filter=h(e),i.addEventListener("message",d(this._onMessage,this),!1)}u.aug(m.prototype,{_onMessage:function(t){var e,n=this;this.server&&(this.isTwitterFrame&&!c.isTwitterURL(t.origin)||(e=p(t.data,this.filter))&&this.server.receive(e,t.source).then(function(e){e&&f(t.source,e,n.filter)}))},attachTo:function(t){this.server=t},detach:function(){this.server=null}}),u.aug(v.prototype,{_processResponse:function(t){var e=this.pending[t.id];e&&(e.resolve(t),delete this.pending[t.id])},_onMessage:function(t){var e;if((!this.isTwitterHost||c.isTwitterURL(t.origin))&&(e=p(t.data,this.filter))){if(u.isType("string",e))try{e=o.parse(e)}catch(t){return}(e=u.isType("array",e)?e:[e]).forEach(d(this._processResponse,this))}},send:function(t){var e=new s;return t.id?this.pending[t.id]=e:e.resolve(),f(this.target,t,this.filter),e.promise}}),t.exports={Receiver:m,Dispatcher:v,_stringifyPayload:function(t){return arguments.length>0&&(l=!!t),l}}},function(t,e,n){var r=n(4);t.exports=function(t){for(var e,n=r.getElementsByTagName("iframe"),i=0;n[i];i++)if((e=n[i]).contentWindow===t)return e}},function(t,e,n){var r=n(5),i=n(0),o=n(3),s=n(13),a=n(14)(),u=n(64),c="a.twitter-moment";t.exports=function(t){return a(t,c).map(function(t){return u(function(t){var e=s(t),n={momentId:o.momentId(t.href),chrome:t.getAttribute("data-chrome"),limit:t.getAttribute("data-limit")};return i.forIn(n,function(t,n){var i=e[t];e[t]=r.hasValue(i)?i:n}),e}(t),t.parentNode,t)})}},function(t,e,n){var r=n(2);t.exports=function(t,e){var i=new r;return Promise.all([n.e(0),n.e(4)]).then(function(r){var o;try{o=n(85),i.resolve(new o(t,e))}catch(t){i.reject(t)}}.bind(null,n)).catch(function(t){i.reject(t)}),i.promise}},function(t,e,n){var r=n(0),i=n(13),o=n(14)(),s=n(65),a="a.periscope-on-air",u=/^https?:\/\/(?:www\.)?(?:periscope|pscp)\.tv\/@?([a-zA-Z0-9_]+)\/?$/i;t.exports=function(t){return o(t,a).map(function(t){return s(function(t){var e=i(t),n=t.getAttribute("href"),o=t.getAttribute("data-size"),s=u.exec(n)[1];return r.aug(e,{username:s,size:o})}(t),t.parentNode,t)})}},function(t,e,n){var r=n(2);t.exports=function(t,e){var i=new r;return n.e(5).then(function(r){var o;try{o=n(86),i.resolve(new o(t,e))}catch(t){i.reject(t)}}.bind(null,n)).catch(function(t){i.reject(t)}),i.promise}},function(t,e,n){var r=n(5),i=n(0),o=n(66),s=n(13),a=n(14)(),u=n(67),c=n(3),d=n(12),l="a.twitter-timeline,div.twitter-timeline,a.twitter-grid",f="Embedded Search timelines have been deprecated. See https://twittercommunity.com/t/deprecating-widget-settings/102295.",h="You may have been affected by an update to settings in embedded timelines. See https://twittercommunity.com/t/deprecating-widget-settings/102295.",p="Embedded grids have been deprecated and will now render as timelines. Please update your embed code to use the twitter-timeline class. More info: https://twittercommunity.com/t/update-on-the-embedded-grid-display-type/119564.";t.exports=function(t){return a(t,l).map(function(t){return u(function(t){var e=s(t),n=t.getAttribute("data-show-replies"),a={isPreconfigured:!!t.getAttribute("data-widget-id"),chrome:t.getAttribute("data-chrome"),tweetLimit:t.getAttribute("data-tweet-limit")||t.getAttribute("data-limit"),ariaLive:t.getAttribute("data-aria-polite"),theme:t.getAttribute("data-theme"),borderColor:t.getAttribute("data-border-color"),showReplies:n?r.asBoolean(n):null,profileScreenName:t.getAttribute("data-screen-name"),profileUserId:t.getAttribute("data-user-id"),favoritesScreenName:t.getAttribute("data-favorites-screen-name"),favoritesUserId:t.getAttribute("data-favorites-user-id"),likesScreenName:t.getAttribute("data-likes-screen-name"),likesUserId:t.getAttribute("data-likes-user-id"),listOwnerScreenName:t.getAttribute("data-list-owner-screen-name"),listOwnerUserId:t.getAttribute("data-list-owner-id"),listId:t.getAttribute("data-list-id"),listSlug:t.getAttribute("data-list-slug"),customTimelineId:t.getAttribute("data-custom-timeline-id"),staticContent:t.getAttribute("data-static-content"),url:t.href};return a.isPreconfigured&&(c.isSearchUrl(a.url)?d.publicError(f,t):d.publicLog(h,t)),"twitter-grid"===t.className&&d.publicLog(p,t),(a=i.aug(a,e)).dataSource=o(a),a.id=a.dataSource&&a.dataSource.id,a}(t),t.parentNode,t)})}},function(t,e,n){var r=n(28);t.exports=r.build([n(29),n(136)])},function(t,e,n){var r=n(0),i=n(135);t.exports=function(t){return"en"===t||r.contains(i,t)}},function(t,e){t.exports=["hi","zh-cn","fr","zh-tw","msa","fil","fi","sv","pl","ja","ko","de","it","pt","es","ru","id","tr","da","no","nl","hu","fa","ar","ur","he","th","cs","uk","vi","ro","bn","el","en-gb","gu","kn","mr","ta","bg","ca","hr","sr","sk"]},function(t,e,n){var r=n(3),i=n(0),o=n(20),s=n(48),a="collection:";function u(t,e){return r.collectionId(t)||e}t.exports=function(t){t.params({id:{},url:{}}),t.overrideProperty("id",{get:function(){var t=u(this.params.url,this.params.id);return a+t}}),t.overrideProperty("endpoint",{get:function(){return o.timeline(["collection"])}}),t.around("queryParams",function(t){return i.aug(t(),{collection_id:u(this.params.url,this.params.id)})}),t.before("initialize",function(){if(!u(this.params.url,this.params.id))throw new Error("one of url or id is required");s()})}},function(t,e,n){var r=n(28);t.exports=r.build([n(29),n(138)])},function(t,e,n){var r=n(3),i=n(0),o=n(20),s="event:";function a(t,e){return r.eventId(t)||e}t.exports=function(t){t.params({id:{},url:{}}),t.overrideProperty("id",{get:function(){var t=a(this.params.url,this.params.id);return s+t}}),t.overrideProperty("endpoint",{get:function(){return o.timeline(["event"])}}),t.around("queryParams",function(t){return i.aug(t(),{event_id:a(this.params.url,this.params.id)})}),t.before("initialize",function(){if(!a(this.params.url,this.params.id))throw new Error("one of url or id is required")})}},function(t,e,n){var r=n(28);t.exports=r.build([n(29),n(140)])},function(t,e,n){var r=n(3),i=n(0),o=n(20),s=n(48),a="likes:";function u(t){return r.likesScreenName(t.url)||t.screenName}t.exports=function(t){t.params({screenName:{},userId:{},url:{}}),t.overrideProperty("id",{get:function(){var t=u(this.params)||this.params.userId;return a+t}}),t.overrideProperty("endpoint",{get:function(){return o.timeline(["likes"])}}),t.define("_getLikesQueryParam",function(){var t=u(this.params);return t?{screen_name:t}:{user_id:this.params.userId}}),t.around("queryParams",function(t){return i.aug(t(),this._getLikesQueryParam())}),t.before("initialize",function(){if(!u(this.params)&&!this.params.userId)throw new Error("screen name or user id is required");s()})}},function(t,e,n){var r=n(28);t.exports=r.build([n(29),n(142)])},function(t,e,n){var r=n(3),i=n(0),o=n(20),s="list:";function a(t){var e=r.listScreenNameAndSlug(t.url)||t;return i.compact({screen_name:e.ownerScreenName,user_id:e.ownerUserId,list_slug:e.slug})}t.exports=function(t){t.params({id:{},ownerScreenName:{},ownerUserId:{},slug:{},url:{}}),t.overrideProperty("id",{get:function(){var t,e,n;return this.params.id?s+this.params.id:(e=(t=a(this.params))&&t.list_slug.replace(/-/g,"_"),n=t&&(t.screen_name||t.user_id),s+(n+":")+e)}}),t.overrideProperty("endpoint",{get:function(){return o.timeline(["list"])}}),t.define("_getListQueryParam",function(){return this.params.id?{list_id:this.params.id}:a(this.params)}),t.around("queryParams",function(t){return i.aug(t(),this._getListQueryParam())}),t.before("initialize",function(){var t=a(this.params);if(i.isEmptyObject(t)&&!this.params.id)throw new Error("qualified slug or list id required")})}},function(t,e,n){var r=n(28);t.exports=r.build([n(29),n(144)])},function(t,e,n){var r=n(3),i=n(5),o=n(0),s=n(20),a="profile:";function u(t,e){return r.screenName(t)||e}t.exports=function(t){t.params({showReplies:{fallback:!1,transform:i.asBoolean},screenName:{},userId:{},url:{}}),t.overrideProperty("id",{get:function(){var t=u(this.params.url,this.params.screenName);return a+(t||this.params.userId)}}),t.overrideProperty("endpoint",{get:function(){return s.timeline(["profile"])}}),t.define("_getProfileQueryParam",function(){var t=u(this.params.url,this.params.screenName),e=t?{screen_name:t}:{user_id:this.params.userId};return o.aug(e,{with_replies:this.params.showReplies?"true":"false"})}),t.around("queryParams",function(t){return o.aug(t(),this._getProfileQueryParam())}),t.before("initialize",function(){if(!u(this.params.url,this.params.screenName)&&!this.params.userId)throw new Error("screen name or user id is required")})}},function(t,e,n){var r=n(2);t.exports=function(t,e){var i=new r;return n.e(6).then(function(r){var o;try{o=n(87),i.resolve(new o(t,e))}catch(t){i.reject(t)}}.bind(null,n)).catch(function(t){i.reject(t)}),i.promise}},function(t,e,n){var r=n(2);t.exports=function(t,e){var i=new r;return Promise.all([n.e(0),n.e(7)]).then(function(r){var o;try{o=n(88),i.resolve(new o(t,e))}catch(t){i.reject(t)}}.bind(null,n)).catch(function(t){i.reject(t)}),i.promise}},function(t,e,n){var r=n(10),i=n(3),o=n(0),s=n(13),a=n(14)(),u=n(68),c="blockquote.twitter-tweet, blockquote.twitter-video",d=/\btw-align-(left|right|center)\b/;t.exports=function(t,e){return a(t,c).map(function(t){return u(function(t){var e=s(t),n=t.getElementsByTagName("A"),a=n&&n[n.length-1],u=a&&i.status(a.href),c=t.getAttribute("data-conversation"),l="none"==c||"hidden"==c||r.present(t,"tw-hide-thread"),f=t.getAttribute("data-cards"),h="none"==f||"hidden"==f||r.present(t,"tw-hide-media"),p=t.getAttribute("data-align")||t.getAttribute("align"),m=t.getAttribute("data-theme");return!p&&d.test(t.className)&&(p=RegExp.$1),o.aug(e,{tweetId:u,hideThread:l,hideCard:h,align:p,theme:m,id:u})}(t),t.parentNode,t,e)})}},function(t,e,n){var r=n(2);t.exports=function(t,e){var i=new r;return n.e(8).then(function(r){var o;try{o=n(89),i.resolve(new o(t,e))}catch(t){i.reject(t)}}.bind(null,n)).catch(function(t){i.reject(t)}),i.promise}},function(t,e,n){var r=n(10),i=n(0),o=n(13),s=n(14)(),a=n(69),u=n(5),c="a.twitter-share-button, a.twitter-mention-button, a.twitter-hashtag-button",d="twitter-hashtag-button",l="twitter-mention-button";t.exports=function(t){return s(t,c).map(function(t){return a(function(t){var e=o(t),n={screenName:t.getAttribute("data-button-screen-name"),text:t.getAttribute("data-text"),type:t.getAttribute("data-type"),size:t.getAttribute("data-size"),url:t.getAttribute("data-url"),hashtags:t.getAttribute("data-hashtags"),via:t.getAttribute("data-via"),buttonHashtag:t.getAttribute("data-button-hashtag")};return i.forIn(n,function(t,n){var r=e[t];e[t]=u.hasValue(r)?r:n}),e.screenName=e.screenName||e.screen_name,e.buttonHashtag=e.buttonHashtag||e.button_hashtag||e.hashtag,r.present(t,d)&&(e.type="hashtag"),r.present(t,l)&&(e.type="mention"),e}(t),t.parentNode,t)})}},function(t,e,n){var r=n(2);t.exports=function(t,e){var i=new r;return n.e(3).then(function(r){var o;try{o=n(90),i.resolve(new o(t,e))}catch(t){i.reject(t)}}.bind(null,n)).catch(function(t){i.reject(t)}),i.promise}},function(t,e,n){var r=n(0);t.exports=r.aug({},n(152),n(153),n(154),n(155),n(156),n(157),n(158))},function(t,e,n){var r=n(61),i=n(17)(["userId"],{},r);t.exports={createDMButton:i}},function(t,e,n){var r=n(63),i=n(17)(["screenName"],{},r);t.exports={createFollowButton:i}},function(t,e,n){var r=n(64),i=n(17)(["momentId"],{},r);t.exports={createMoment:i}},function(t,e,n){var r=n(65),i=n(17)(["username"],{},r);t.exports={createPeriscopeOnAirButton:i}},function(t,e,n){var r=n(9),i=n(12),o=n(3),s=n(0),a=n(5),u=n(66),c=n(67),d=n(17)([],{},c),l=n(6),f="Embedded grids have been deprecated. Please use twttr.widgets.createTimeline instead. More info: https://twittercommunity.com/t/update-on-the-embedded-grid-display-type/119564.",h={createTimeline:p,createGridFromCollection:function(t){var e=s.toRealArray(arguments).slice(1),n={sourceType:"collection",id:t};return e.unshift(n),i.publicLog(f),p.apply(this,e)}};function p(t){var e,n=s.toRealArray(arguments).slice(1);return a.isString(t)||a.isNumber(t)?l.reject("Embedded timelines with widget settings have been deprecated. See https://twittercommunity.com/t/deprecating-widget-settings/102295."):s.isObject(t)?(t=t||{},n.forEach(function(t){s.isType("object",t)&&function(t){t.ariaLive=t.ariaPolite}(e=t)}),e||(e={},n.push(e)),t.lang=e.lang,t.tweetLimit=e.tweetLimit,t.showReplies=e.showReplies,e.dataSource=u(t),d.apply(this,n)):l.reject("data source must be an object.")}o.isTwitterURL(r.href)&&(h.createTimelinePreview=function(t,e,n){var r={previewParams:t,useLegacyDefaults:!0,isPreviewTimeline:!0};return r.dataSource=u(r),d(e,r,n)}),t.exports=h},function(t,e,n){var r,i=n(0),o=n(68),s=n(17),a=(r=s(["tweetId"],{},o),function(){return i.toRealArray(arguments).slice(1).forEach(function(t){i.isType("object",t)&&(t.hideCard="none"==t.cards||"hidden"==t.cards,t.hideThread="none"==t.conversation||"hidden"==t.conversation)}),r.apply(this,arguments)});t.exports={createTweet:a,createTweetEmbed:a,createVideo:a}},function(t,e,n){var r=n(0),i=n(69),o=n(17),s=o(["url"],{type:"share"},i),a=o(["buttonHashtag"],{type:"hashtag"},i),u=o(["screenName"],{type:"mention"},i);function c(t){return function(){return r.toRealArray(arguments).slice(1).forEach(function(t){r.isType("object",t)&&(t.screenName=t.screenName||t.screen_name,t.buttonHashtag=t.buttonHashtag||t.button_hashtag||t.hashtag)}),t.apply(this,arguments)}}t.exports={createShareButton:c(s),createHashtagButton:c(a),createMentionButton:c(u)}},function(t,e,n){var r,i,o,s=n(4),a=n(1),u=0,c=[],d=s.createElement("a");function l(){var t,e;for(u=1,t=0,e=c.length;t<e;t++)c[t]()}/^loade|c/.test(s.readyState)&&(u=1),s.addEventListener&&s.addEventListener("DOMContentLoaded",i=function(){s.removeEventListener("DOMContentLoaded",i,!1),l()},!1),d.doScroll&&s.attachEvent("onreadystatechange",r=function(){/^c/.test(s.readyState)&&(s.detachEvent("onreadystatechange",r),l())}),o=d.doScroll?function(t){a.self!=a.top?u?t():c.push(t):function(){try{d.doScroll("left")}catch(e){return setTimeout(function(){o(t)},50)}t()}()}:function(t){u?t():c.push(t)},t.exports=o},function(t,e,n){var r=n(46),i=n(8);t.exports=function(){i.set("buildVersion",r.version)}},function(t,e,n){n(162),n(165),n(166)},function(t,e,n){var r=n(163),i=n(30),o=n(70),s=new r,a=function(t){t.widgets&&1===t.widgets.length&&(s.start(),i.emitter.unbind(i.ALL_WIDGETS_RENDER_START,a))},u=function(t){var e;t.widgets&&1===t.widgets.length&&(e=t.widgets[0],s.end(),e.dataset&&e.dataset.tweetId&&o({duration:s.duration(),namespace:{element:"tweet",action:"render"},widgetIds:[e.dataset.tweetId]})),i.emitter.unbind(i.ALL_WIDGETS_RENDER_END,u)};i.emitter.bind(i.ALL_WIDGETS_RENDER_START,a),i.emitter.bind(i.ALL_WIDGETS_RENDER_END,u)},function(t,e,n){var r=n(164);function i(){}i.prototype.start=function(){this._startTime=r()},i.prototype.end=function(){this._duration=r()-this._startTime},i.prototype.duration=function(){return this._duration},t.exports=i},function(t,e,n){var r=n(1);t.exports=function(){return r.performance&&r.performance.now?r.performance.now():Date.now()}},function(t,e,n){var r,i=n(30),o=0;function s(){r&&r.length===o&&(i.emitter.trigger(i.ALL_WIDGETS_AND_IMAGES_LOADED,r),r=null)}i.emitter.bind(i.ALL_WIDGETS_RENDER_END,function(t){r=t.widgets,s()}),t.exports={reportImagesLoadForAWidget:function(){o++,s()}}},function(t,e,n){var r=n(30),i=n(70),o=n(167),s=n(3),a=n(1),u=n(0),c=n(21),d=n(62);function l(t){return t.performance.getEntriesByType("resource").filter(function(t){return s.isTwimgURL(t.name)||s.isTwitterURL(t.name)}).reduce(function(t,e){return t[e.name]=e.duration,t},{})}r.emitter.bind(r.ALL_WIDGETS_AND_IMAGES_LOADED,function(t){var e,n,r=[];c.hasPerformanceInformation()&&(e=l(a),d.isSupported()||(r=function(t){return t.reduce(function(t,e){return u.aug(t,l(e.contentDocument.defaultView))},{})}(t)),n=u.aug({},e,r),Object.keys(o).forEach(function(t){!function(t,e,n){var r=Object.keys(t).reduce(function(e,r){return n(r)?e+t[r]:e},0);i({duration:r,namespace:{element:e,action:"resource"}})}(n,t,o[t])}))})},function(t,e,n){var r=n(3),i={all:function(){return!0},image:function(t){return r.isTwimgURL(t)},settings:function(t){return r.isSettingsURL(t)},widget_iframe:function(t){return r.isWidgetIframeURL(t)}};t.exports=i}])));
```
