## supplementary data file 3 for "Repeated truncation of a modular antimicrobial peptide gene for neural context": Datamonkey Adaptive Evolution Server.html

Methods and Tools

aBSREL
SpiderMonkey/BGM
BUSTED
Contrast-FEL
FADE
FEL
FUBAR
GARD
HIV-TRACE
MULTI-HIT
MEME
RELAX
SLAC
All Methods

Job Queue
Usage statistics

API

API Info
Get API Key
Check Key Status

Citations
Help
COVID-19
Blog
 Classic

Methods and Tools

aBSREL
BUSTED
FADE Beta
FEL
FUBAR
GARD
HIV-TRACE
MEME
RELAX
SLAC
All Methods

Job Queue
Usage statistics
Citations
Help
 Classic

- summary
- model statistics
- tree
- phylo alignment

×Close**Error!**

##### Branch-site Unrestricted Statistical Test for Episodic Diversification results summary

INPUT DATA |6095a673238adf71a515dd06|23 sequences |58 sites

Export

- Original file
- Analysis log
- View MSA
- Save JSON
- View JSON

###### Alignment viewer

×

Close

BUSTED with synyonymous rate variation **found evidence** (LRT, p-value = 0.008 ≤ .05) of gene-wide episodic diversifying selection in the selected test branches of your phylogeny. Therefore, there is evidence that at least one site on at least one test branch has experienced diversifying selection.

---

See here for more information about this method.  
Please cite PMID 25701167 if you use this result in a publication, presentation, or other scientific work.

###### Model fits

| Model | *log* L | #. params | AIC_c_ | CV(SRV) | Branch set | ω_1_ | ω_2_ | ω_3_ |
| --- | --- | --- | --- | --- | --- | --- | --- | --- |
| Unconstrained model | -1548.0 | 67 | 3237.3 | 1.538 | Test | 0.00 (63.41%) | 0.15 (35.86%) | 482.99 (0.73%) |
| Constrained model | -1552.2 | 66 | 3243.3 | 1.448 | Test | 0.00 (65.83%) | 0.12 (29.06%) | 1.00 (5.10%) |

###### BUSTED Site Proportion Chart

×

###### ω distribution

### **Unconstrained model, Test branches**

SVG PNG

Neutrality (ω=1)ω0.000010.00010.0010.010.1110100100010000Proportion of sites0%10%20%30%40%50%60%70%80%90%100%

Close

This table reports a statistical summary of the models fit to the data. Here, **Unconstrained model** refers to the BUSTED alternative model for selection, and **Constrained model** refers to the BUSTED null model for selection.

###### Model Test Statistics Per Site

Export Chart to SVG Export Chart to PNG

1020304050Site index-2024682\*Log Evidence RatioConstrainedOptimized Null

Constrained Test Statistic

Optimized Null Test Statistic

Showing entries 1 through 20 out of 58.

Export Table to CSV

| Site index | Unconstrained likelihood | Constrained likelihood | Optimized Null Likelihood | Constrained Statistic | Optimized Null Statistic |
| --- | --- | --- | --- | --- | --- |
| 1 | -17.86 | -17.71 | -18.07 | -0.31 | 0.42 |
| 2 | -54.33 | -54.3 | -54.8 | -0.06 | 0.96 |
| 3 | -20.28 | -20.11 | -20.23 | -0.33 | -0.1 |
| 4 | -17.36 | -17.2 | -17.7 | -0.32 | 0.68 |
| 5 | -12.41 | -12.24 | -12.42 | -0.34 | 0.03 |
| 6 | -19.96 | -19.78 | -19.89 | -0.37 | -0.14 |
| 7 | -24.61 | -24.44 | -24.92 | -0.35 | 0.62 |
| 8 | -29.96 | -29.86 | -30.39 | -0.18 | 0.86 |
| 9 | -32.8 | -32.65 | -33.78 | -0.3 | 1.96 |
| 10 | -26.04 | -25.87 | -26.2 | -0.34 | 0.33 |
| 11 | -10.43 | -10.25 | -10.3 | -0.34 | -0.26 |
| 12 | -27.83 | -27.67 | -27.83 | -0.33 | -0.01 |
| 13 | -18.04 | -17.88 | -18.23 | -0.32 | 0.38 |
| 14 | -20.84 | -20.66 | -20.9 | -0.36 | 0.11 |
| 15 | -76.04 | -75.92 | -77.27 | -0.25 | 2.46 |
| 16 | -20.71 | -20.53 | -20.88 | -0.36 | 0.34 |
| 17 | -16.61 | -16.43 | -16.62 | -0.36 | 0.01 |
| 18 | -29.06 | -28.88 | -29.53 | -0.36 | 0.94 |
| 19 | -19.78 | -19.58 | -19.75 | -0.39 | -0.06 |
| 20 | -18.97 | -18.79 | -18.9 | -0.35 | -0.13 |

###### Fitted tree

Options

- Partitions
- 1
- Models
- Unconstrained model
- Constrained model

- Hide Legend
- GrayScale

Export 

- PNG
- SVG
- Newick File

TestBackgroundLength = 0.4752626882759278Length = 0.7294966191475027Length = 0.3179963713660124Length = 0.0001047171503513057Length = 0.06334564774616787Length = 0.2010201368567486Length = 0.1959087398902796Length = 0.004005772685970319Length = 0.2658733493109541Length = 0.009488572382786725Length = 1.238677726399081Length = 0.4659237516207727Length = 0.6472824875393706Length = 0.4512261798931614Length = 1.291402685492405Length = 0.4368046286855465Length = 0.0917642663911017Length = 1.3920130721216Length = 0Length = 2.308889295464793Length = 1.287487960651553Length = 0.9197870044947707Length = 0Length = 0.5103682277234906Length = 1.047902704925222Length = 0.9708963596584242Length = 1.431436878434739Length = 3.472532930755869Length = 0Length = 1.574942815015225Length = 0Length = 4.063287530114751Length = 0Length = 0.05691523083409536Length = 0.5596856265489762Length = 0.4495533813579038Length = 0.009527800387170684Length = 0.4426840171202434Length = 0Length = 0.383668276263302Length = 1.569347863108978Length = 2.104557846972568Length = 10.72805306929661DINS\_BARACDTRO\_BARACDWIL\_GK10645DPAU\_BARAC\_STR\_L06\_DPAU\_BARAC\_STR\_L12\_DPSE\_GA24235DSER\_XM\_020962164DBIP\_XM\_017235265DANA\_GF13690DRHO\_XM\_017120938DFIC\_XM\_017195800DEUG\_XM\_017221794DELE\_XM\_017276537DTAK\_XM\_017145699DSUZ\_XM\_017088027DBIA\_XM\_017101808DMEL\_BARAA\_CG33470\_AND\_CG18279\_DSIM\_GD10965\_W501\_DSEC\_GM21466\_XM\_002033668DMAU\_XM\_033298434\_LOC117137134DERE\_GG20379DYAK\_GE12540SLEB\_XM\_030519534

###### Phylogenetic alignment evidence ratio plot

Current site:Add siteRemove sitePNG

ConstrainedOptimized Null2.62.42.22.01.81.61.41.21.00.80.60.40.20.0log(1+evidence ratio)SLEB\_XM\_030519534DINS\_BARACDPSE\_GA24235DTRO\_BARACDSER\_XM\_020962164DWIL\_GK10645DPAU\_BARAC\_STR\_L06\_DPAU\_BARAC\_STR\_L12\_DBIP\_XM\_017235265DANA\_GF13690DRHO\_XM\_017120938DFIC\_XM\_017195800DEUG\_XM\_017221794DELE\_XM\_017276537DTAK\_XM\_017145699DSUZ\_XM\_017088027DBIA\_XM\_017101808DMEL\_BARAA\_CG33470\_ADERE\_GG20379DYAK\_GE12540DSIM\_GD10965\_W501\_DSEC\_GM21466\_XM\_0020DMAU\_XM\_033298434\_LOCAACAACAACAACAACAACAACAGCAGCAGCAACAACAGCAACAACAACAACAACAGCAGCAGCAGCAGQQQQQQQQQQQQQQQQQQQQQQQCodon 1CAACAACAACAACAACAACAACAGCAGCAGCAACAACAGCAACAACAACAACAACAGCAGCAGCAGCAGQQQQQQQQQQQQQQQQQQQQQQQCodon 1GAAGAAGAAGAAGAAGAAGAAGAAGAAGAGGAAGAAGAGGAAGAAGAAGAAGAGGCGGCGCCGGAGGAGEEEEEEEEEEEEEEEEEEAAPEECodon 27CAACAACAACAACAACAACAACAGCAGCAGCAACAACAGCAACAACAACAACAACAGCAGCAGCAGCAGQQQQQQQQQQQQQQQQQQQQQQQCodon 1CAACAACAACAACAACAACAACAGCAGCAGCAACAACAGCAACAACAACAACAACAGCAGCAGCAGCAGQQQQQQQQQQQQQQQQQQQQQQQCodon 1

×

###### Error

This is my error message

Close

Datamonkey is funded jointly by MIDAS and NIH award R01 GM093939
