## supplementary data file 3 for "Repeated truncation of a modular antimicrobial peptide gene for neural context": log.txt

Analysis Description
--------------------
BUSTED (branch-site unrestricted statistical test of episodic
diversification) uses a random effects branch-site model fitted jointly
to all or a subset of tree branches in order to test for alignment-wide
evidence of episodic diversifying selection. Assuming there is evidence
of positive selection (i.e. there is an omega > 1), BUSTED will also
perform a quick evidence-ratio style analysis to explore which
individual sites may have been subject to selection. v2.0 adds support
for synonymous rate variation, and relaxes the test statistic to 0.5
(chi^2_0 + chi^2_2). Version 2.1 adds a grid search for the initial
starting point. Version 2.2 changes the grid search to LHC, and adds an
initial search phase to use adaptive Nedler-Mead. Version 3.0 implements
the option for branch-site variation in synonymous substitution rates.
Version 3.1 adds HMM auto-correlation option for SRV, and binds SRV
distributions for multiple branch sets 

- __Requirements__: in-frame codon alignment and a phylogenetic tree (optionally annotated
with {})

- __Citation__: *Gene-wide identification of episodic selection*, Mol Biol Evol.
32(5):1365-71

- __Written by__: Sergei L Kosakovsky Pond

- __Contact Information__:

- __Analysis Version__: 3.1


>code –> Universal

>Loaded a multiple sequence alignment with **23** sequences, **58** codons, and **1** partitions from `/home/datamonkey/datamonkey-js-server/production/app/busted/output/6095a673238adf71a515dd06`

>branches –> All

>srv –> Yes
The number omega rate classes to include in the model (permissible range = [1,10], default value = 3, integer): 
>rates –> 3
The number omega rate classes to include in the model (permissible range = [1,10], default value = 3, integer): 
>syn-rates –> 3
The number of points in the initial distributional guess for likelihood fitting (permissible range = [1,10000], default value = 250, integer): 
>grid-size –> 250
The number of initial random guesses to 'seed' rate values optimization (permissible range = [1,25], default value = 1, integer): 
>starting-points –> 1


### Branches to test for selection in the BUSTED analysis
* Selected 43 branches to test in the BUSTED analysis: `DINS_BARAC, DTRO_BARAC, DWIL_GK10645, DPAU_BARAC_STR_L06_, DPAU_BARAC_STR_L12_, Node9, Node7, Node5, Node3, DPSE_GA24235, DSER_XM_020962164, DBIP_XM_017235265, DANA_GF13690, Node17, DRHO_XM_017120938, DFIC_XM_017195800, Node21, DEUG_XM_017221794, DELE_XM_017276537, DTAK_XM_017145699, DSUZ_XM_017088027, DBIA_XM_017101808, Node31, Node29, DMEL_BARAA_CG33470_AND_CG18279_, DSIM_GD10965_W501_, DSEC_GM21466_XM_002033668, DMAU_XM_033298434_LOC117137134, Node39, Node37, Node35, DERE_GG20379, DYAK_GE12540, Node42, Node34, Node28, Node26, Node24, Node20, Node16, Node14, Node12, SLEB_XM_030519534`


### Obtaining branch lengths and nucleotide substitution biases under the nucleotide GTR model

>kill-zero-lengths –> No
* Log(L) = -1799.78, AIC-c =  3702.90 (51 estimated parameters)

### Obtaining the global omega estimate based on relative GTR branch lengths and nucleotide substitution biases
* Log(L) = -1618.81, AIC-c =  3358.98 (58 estimated parameters)
* non-synonymous/synonymous rate ratio for *test* =   0.1353

### Improving branch lengths, nucleotide substitution biases, and global dN/dS ratios under a full codon model
* Log(L) = -1591.68, AIC-c =  3304.72 (58 estimated parameters)
* non-synonymous/synonymous rate ratio for *test* =   0.0942

### Performing the full (dN/dS > 1 allowed) branch-site model fit
* Log(L) = -1548.03, AIC-c =  3237.25 (67 estimated parameters)
* For *test* branches, the following rate distribution for branch-site combinations was inferred

|          Selection mode           |     dN/dS     |Proportion, %|               Notes               |
|-----------------------------------|---------------|-------------|-----------------------------------|
|        Negative selection         |     0.000     |   63.410    |                                   |
|        Negative selection         |     0.153     |   35.862    |                                   |
|      Diversifying selection       |    482.986    |    0.728    |                                   |

* The following rate distribution for site-to-site **synonymous** rate variation was inferred

|               Rate                | Proportion, % |               Notes               |
|-----------------------------------|---------------|-----------------------------------|
|               0.527               |    76.331     |                                   |
|               1.144               |    18.468     |                                   |
|               7.429               |     5.202     |                                   |


### Performing the constrained (dN/dS > 1 not allowed) model fit
