## supplementary data file 3 for "Repeated truncation of a modular antimicrobial peptide gene for neural context": Summary ATP8A locus.txt

FEL found evidence of
 pervasive positive/diversifying selection at 1 sites
 pervasive negative/purifying selection at 51 sites
with p-value threshold of .1

(site 29, p = 0.011, omega approaches infinity)

FUBAR found evidence of
 episodic positive/diversifying selection at 2 sites
 episodic negative/purifying selection at 46 sites
with posterior probability of .9

(site 29, p = 0.013)
(site 31, p = 0.026)

SLAC found evidence of pervasive
 positive/diversifying selection at 1 sites
 negative/purifying selection at 38 sites
with p-value threshold of .1

(site 29, p = 0.039)

aBSREL found evidence of episodic diversifying selection on 1 out of 43 branches in your phylogeny. A total of 43 branches were formally tested for diversifying selection. Significance was assessed using the Likelihood Ratio Test at a threshold of p ≤ 0.05, after correcting for multiple testing. Significance and number of rate categories inferred at each branch are provided in the detailed results table.

(node 12, p-adj = 0.0435, uncorrected p = 0.001, w1 = 1(94%), w2 = 14400(6%))
- Node 12 corresponds with the divergence of Willistoni clade Baramicins from rest of subgenus Drosophila

BUSTED found evidence (LRT, p-value = 0.008 < .05) of gene-wide episodic diversifying selection in the selected test branches of your phylogeny. Therefore, there is evidence that at least one site on at least one test branch has experienced diversifying selection.
