## supplementary data file 3 for "Repeated truncation of a modular antimicrobial peptide gene for neural context": Datamonkey Adaptive Evolution Server2.html

Datamonkey Adaptive Evolution Server

Methods and Tools

aBSREL
SpiderMonkey/BGM
BUSTED
Contrast-FEL
FADE
FEL
FUBAR
GARD
HIV-TRACE
MULTI-HIT
MEME
RELAX
SLAC
All Methods

Job Queue
Usage statistics

API

API Info
Get API Key
Check Key Status

Citations
Help
COVID-19
Blog
 Classic

Methods and Tools

aBSREL
BUSTED
FADE Beta
FEL
FUBAR
GARD
HIV-TRACE
MEME
RELAX
SLAC
All Methods

Job Queue
Usage statistics
Citations
Help
 Classic

- summary
- tree
- table
- model fits

×Close**Error!**

##### adaptive Branch Site REL results summary

INPUT DATA |609599f7238adf71a515d790|12 sequences |58 sites

Export

- Original file
- Analysis log
- Save JSON
- View JSON

###### Alignment viewer

×

Close

aBSREL **found no evidence** of episodic diversifying selection in your phylogeny.

A total of **20** branches were formally tested for diversifying selection. Significance was assessed using the Likelihood Ratio Test at a threshold of p ≤ 0.05, after correcting for multiple testing. Significance and number of rate categories inferred at each branch are provided in the detailed results table.

---

See here for more information about this method.  
Please cite PMID 25697341 if you use this result in a publication, presentation, or other scientific work.

###### Tree summary

| ω rate classes | # of branches | % of branches | % of tree length | # under selection |
| --- | --- | --- | --- | --- |
| 1 | 17 | 85% | 35% | 0 |
| 2 | 3 | 15% | 65% | 0 |

This table contains a summary of the inferred aBSREL model complexity. Each row provides information about the branches that were best described by the given number of ω rate categories.

###### Fitted tree

Options

- Models
- Full adaptive model
- Baseline MG94xREV

- Hide Legend
- GrayScale

Export 

- PNG
- SVG
- Newick File

00.010.10.512510ωLength = 0.01154866513569438Length = 0Length = 0.005600594903804572Length = 0.007548538887811502Length = 0.04030500831720442Length = 0.04101327896547582Length = 0.1280806310895582Length = 0.0126474870096584Length = 0.3570605722723381Length = 0.070569281009682Length = 0.06130222323532414Length = 0.02217104786657896Length = 0.02760124343573041Length = 0Length = 0.1197379330832979Length = 0.1245164311130385Length = 0.1155429530839606Length = 0.8792230584498528Length = 0.005971980012955655Length = 0.2473692157549807DBIP\_XM\_017237939DANA\_XM\_014907649DRHO\_XM\_017134245DSUZ\_XM\_017084964DBIA\_XM\_017102001DEUG\_XM\_017210758DYAK\_GE19241\_BARAB\_DMEL\_BARAB\_CG13749\_DSIM\_BARAB\_XM\_002080640\_3\_173\_346DMAU\_BARAB\_XM\_033300133\_1\_282\_455DSEC\_BARAB\_XM\_002032966\_2\_61\_234SLEB\_IM24

###### Detailed results

| Name | B | LRT | Test p-value | Uncorrected p-value | ω distribution over sites |
| --- | --- | --- | --- | --- | --- |
| DANA\_XM\_014907649 | 0.0000 | 0.0000 | 1.0000 | 1.0000 | ω_1_ = 0.0384 (100%) |
| DBIA\_XM\_017102001 | 0.0000 | 0.0000 | 1.0000 | 1.0000 | ω_1_ = 0.0750 (100%) |
| DBIP\_XM\_017237939 | 0.0000 | 0.0000 | 1.0000 | 1.0000 | ω_1_ = 0.300 (100%) |
| DEUG\_XM\_017210758 | 0.0000 | 1.3720 | 1.0000 | 0.1994 | ω_1_ = 0.0818 (93%) ω_2_ = 8.51 (6.6%) |
| DMAU\_BARAB\_XM\_033300133\_1\_282\_455 | 0.0000 | 0.0000 | 1.0000 | 1.0000 | ω_1_ = 1.00 (100%) |
| DMEL\_BARAB\_CG13749\_ | 0.0000 | 0.0000 | 1.0000 | 1.0000 | ω_1_ = 0.528 (100%) |
| DRHO\_XM\_017134245 | 0.0000 | 0.0000 | 1.0000 | 1.0000 | ω_1_ = 0.0195 (100%) |
| DSEC\_BARAB\_XM\_002032966\_2\_61\_234 | 0.0000 | 0.0000 | 1.0000 | 1.0000 | ω_1_ = 0.389 (100%) |
| DSIM\_BARAB\_XM\_002080640\_3\_173\_346 | 0.0000 | 0.0000 | 1.0000 | 1.0000 | ω_1_ = 0.00 (100%) |
| DSUZ\_XM\_017084964 | 0.0000 | 0.0000 | 1.0000 | 1.0000 | ω_1_ = 0.0889 (100%) |
| DYAK\_GE19241\_BARAB\_ | 0.0000 | 0.0000 | 1.0000 | 1.0000 | ω_1_ = 0.678 (100%) |
| Node12 | 0.0000 | 0.0000 | 1.0000 | 1.0000 | ω_1_ = 0.00 (100%) |
| Node14 | 0.0000 | 0.1351 | 1.0000 | 0.4230 | ω_1_ = 10000000000 (100%) |
| Node16 | 0.0000 | 0.0000 | 1.0000 | 1.0000 | ω_1_ = 0.0532 (100%) |
| Node18 | 0.0000 | 0.0000 | 1.0000 | 1.0000 | ω_1_ = 0.00 (100%) |
| Node3 | 0.0000 | 2.7302 | 1.0000 | 0.0963 | ω_1_ = 0.00 (90%) ω_2_ = 4.34 (9.8%) |
| Node6 | 0.0000 | 0.0000 | 1.0000 | 1.0000 | ω_1_ = 0.00 (100%) |
| Node8 | 0.0000 | 0.0000 | 1.0000 | 1.0000 | ω_1_ = 1.00 (100%) |
| Node9 | 0.0000 | 0.0000 | 1.0000 | 1.0000 | ω_1_ = 0.100 (100%) |
| SLEB\_IM24 | 0.0000 | 1.0764 | 1.0000 | 0.2350 | ω_1_ = 0.0484 (89%) ω_2_ = 4.24 (11%) |

###### aBSREL Site Proportion Chart

×

###### ω distribution

### **DANA\_XM\_014907649**

SVG PNG

Neutrality (ω=1)ω0.000010.00010.0010.010.1110100100010000Proportion of sites0%10%20%30%40%50%60%70%80%90%100%

Close

###### Model fits

| Model | AIC_C_ | log L | Parameters |
| --- | --- | --- | --- |
| Nucleotide GTR | 2249.49 | -1095.32 | 29 |
| Baseline MG94xREV | 2102.48 | -992.61 | 54 |
| Full adaptive model | 2086.22 | -977.34 | 60 |

This table reports a statistical summary of the models fit to the data. Here, **Baseline MG94xREV** refers to the MG94xREV baseline model that infers a single ω rate category per branch. **Full adaptive model** refers to the adaptive aBSREL model that infers an optimized number of ω rate categories per branch.

×

###### Error

This is my error message

Close

Datamonkey is funded jointly by MIDAS and NIH award R01 GM093939
