## supplementary data file 3 for "Repeated truncation of a modular antimicrobial peptide gene for neural context": Datamonkey Adaptive Evolution Server.html

Methods and Tools

aBSREL
SpiderMonkey/BGM
BUSTED
Contrast-FEL
FADE
FEL
FUBAR
GARD
HIV-TRACE
MULTI-HIT
MEME
RELAX
SLAC
All Methods

Job Queue
Usage statistics

API

API Info
Get API Key
Check Key Status

Citations
Help
COVID-19
Blog
 Classic

Methods and Tools

aBSREL
BUSTED
FADE Beta
FEL
FUBAR
GARD
HIV-TRACE
MEME
RELAX
SLAC
All Methods

Job Queue
Usage statistics
Citations
Help
 Classic

- summary
- model statistics
- tree
- phylo alignment

×Close**Error!**

##### Branch-site Unrestricted Statistical Test for Episodic Diversification results summary

INPUT DATA |60959a0c238adf71a515d7ac|12 sequences |58 sites

Export

- Original file
- Analysis log
- View MSA
- Save JSON
- View JSON

###### Alignment viewer

×

Close

BUSTED with synyonymous rate variation **found no evidence** (LRT, p-value = 0.500 ≥ .05) of gene-wide episodic diversifying selection in the selected test branches of your phylogeny. Therefore, there is no evidence that any sites have experienced diversifying selection along the test branch(es).

---

See here for more information about this method.  
Please cite PMID 25701167 if you use this result in a publication, presentation, or other scientific work.

###### Model fits

| Model | *log* L | #. params | AIC_c_ | CV(SRV) | Branch set | ω_1_ | ω_2_ | ω_3_ |
| --- | --- | --- | --- | --- | --- | --- | --- | --- |
| Unconstrained model | -997.5 | 45 | 2091.3 | 0.172 | Test | 0.03 (76.98%) | 0.05 (8.09%) | 1.00 (14.92%) |

###### BUSTED Site Proportion Chart

×

###### ω distribution

### **Unconstrained model, Test branches**

SVG PNG

Neutrality (ω=1)ω0.000010.00010.0010.010.1110100100010000Proportion of sites0%10%20%30%40%50%60%70%80%90%100%

Close

This table reports a statistical summary of the models fit to the data. Here, **Unconstrained model** refers to the BUSTED alternative model for selection, and **Constrained model** refers to the BUSTED null model for selection.

###### Model Test Statistics Per Site

Export Chart to SVG Export Chart to PNG

Constrained Test Statistic

Optimized Null Test Statistic

| Site index | Unconstrained likelihood | Constrained likelihood | Optimized Null Likelihood | Constrained Statistic | Optimized Null Statistic |
| --- | --- | --- | --- | --- | --- |

###### Fitted tree

Options

- Partitions
- 1
- Models
- Unconstrained model

- Hide Legend
- GrayScale

Export 

- PNG
- SVG
- Newick File

TestBackgroundLength = 0.01153316267878406Length = 0Length = 0Length = 0.005705761010031209Length = 0.005265764562897056Length = 0.04174330005672557Length = 0.03598131731981331Length = 0.1295128824847573Length = 0.02557137528977479Length = 0.2280372273028681Length = 0.06826745532000279Length = 0.06606650353960025Length = 0.007632869692932015Length = 0.03201621101617723Length = 0Length = 0.120750447191074Length = 0.09611625369474466Length = 0.1435002616034837Length = 0.6401036760668959Length = 0Length = 0.2265925730750245DBIP\_XM\_017237939DANA\_XM\_014907649DRHO\_XM\_017134245DSUZ\_XM\_017084964DBIA\_XM\_017102001DEUG\_XM\_017210758DYAK\_GE19241\_BARAB\_DMEL\_BARAB\_CG13749\_DSIM\_BARAB\_XM\_002080640\_3\_173\_346DMAU\_BARAB\_XM\_033300133\_1\_282\_455DSEC\_BARAB\_XM\_002032966\_2\_61\_234SLEB\_IM24

###### Phylogenetic alignment evidence ratio plot

Phylogenetic Alignment cannot be rendered for this job.

In order to view the phylogenetic alignment plot, this job must be completed and rendered on datamonkey.org. Hyphy-Vision will not render this plot.

If this job was completed on datamonkey.org, and this message is being displayed, then this job did not present the required data for a successful plot.

This is generally caused by failing to reject the null hypothesis under the unconstrained model, rendering future tests moot to conduct (e.g. constrained model).

×

###### Error

This is my error message

Close

Datamonkey is funded jointly by MIDAS and NIH award R01 GM093939
