## supplementary data file 3 for "Repeated truncation of a modular antimicrobial peptide gene for neural context": FELDatamonkey Adaptive Evolution Server.html

Datamonkey Adaptive Evolution Server


Methods and Tools

aBSREL
SpiderMonkey/BGM
BUSTED
Contrast-FEL
FADE
FEL
FUBAR
GARD
HIV-TRACE
MULTI-HIT
MEME
RELAX
SLAC
All Methods

Job Queue
Usage statistics

API

API Info
Get API Key
Check Key Status

Citations
Help
COVID-19
Blog
 Classic

Methods and Tools

aBSREL
BUSTED
FADE Beta
FEL
FUBAR
GARD
HIV-TRACE
MEME
RELAX
SLAC
All Methods

Job Queue
Usage statistics
Citations
Help
 Classic

- summary
- table
- plot
- tree
- fits

×Close**Error!**

### Fixed Effects Likelihood results summary

INPUT DATA |609599bf238adf71a515d73a|12 sequences |58 sites

Export

- Original file
- Analysis log
- Save JSON
- View JSON

#### Alignment viewer

×

Close

×Close**Error!**

FEL  **found evidence** of

pervasive positive/diversifying selection at 0 sites

pervasive negative/purifying selection at 34 sites

with p-value threshold of.

---

See here for more information about this method.  
Please cite PMID 15703242 if you use this result in a publication, presentation, or other scientific work.

#### FEL Table

Positively selected sites with evidence are highlighted in green.

Negatively selected sites with evidence are highlighted in black.

Showing entries 1 through 20 out of 58.

Export Table to CSV

| Site | Partition | alpha | beta | omega | alpha=beta | LRT | p-value | Total branch length |
| --- | --- | --- | --- | --- | --- | --- | --- | --- |
| 1 | 1 | 3.632 | 0.000 | 0.000 | 0.312 | 10.800 | 0.001 | 0.000 |
| 2 | 1 | 1.246 | 0.410 | 0.329 | 0.559 | 0.925 | 0.336 | 0.000 |
| 3 | 1 | 0.443 | 0.521 | 1.177 | 0.499 | 0.020 | 0.886 | 0.000 |
| 4 | 1 | 1.405 | 0.126 | 0.090 | 0.323 | 3.596 | 0.058 | 0.000 |
| 5 | 1 | 1.058 | 0.231 | 0.219 | 0.338 | 1.276 | 0.259 | 0.000 |
| 6 | 1 | 2.885 | 0.119 | 0.041 | 0.621 | 10.335 | 0.001 | 0.000 |
| 7 | 1 | 0.397 | 0.430 | 1.082 | 0.421 | 0.004 | 0.948 | 0.000 |
| 8 | 1 | 0.960 | 0.206 | 0.215 | 0.326 | 1.342 | 0.247 | 0.000 |
| 9 | 1 | 1.891 | 0.000 | 0.000 | 0.375 | 12.271 | 0.000 | 0.000 |
| 10 | 1 | 10.000 | 0.000 | 0.000 | 0.426 | 14.797 | 0.000 | 0.000 |
| 11 | 1 | 363.437 | 0.101 | 0.000 | 0.481 | 17.285 | 0.000 | 0.000 |
| 12 | 1 | 0.509 | 0.116 | 0.228 | 0.240 | 1.497 | 0.221 | 0.000 |
| 13 | 1 | 0.000 | 0.085 | Infinity | 0.072 | 0.295 | 0.587 | 0.000 |
| 14 | 1 | 0.909 | 0.000 | 0.000 | 0.256 | 7.258 | 0.007 | 0.000 |
| 15 | 1 | 1.307 | 0.230 | 0.176 | 0.483 | 3.424 | 0.064 | 0.000 |
| 16 | 1 | 0.953 | 0.000 | 0.000 | 0.245 | 7.366 | 0.007 | 0.000 |
| 17 | 1 | 0.695 | 0.000 | 0.000 | 0.167 | 5.362 | 0.021 | 0.000 |
| 18 | 1 | 4.669 | 0.194 | 0.042 | 0.566 | 9.417 | 0.002 | 0.000 |
| 19 | 1 | 1.997 | 0.123 | 0.062 | 0.344 | 4.700 | 0.030 | 0.000 |
| 20 | 1 | 0.573 | 0.114 | 0.199 | 0.244 | 1.764 | 0.184 | 0.000 |

### FEL Site Plot

Y-axis: 

- alpha
- beta
- omega
- alpha=beta
- LRT
- p-value
- Total branch length

alpha

Export to PNG Export to SVG

5101520253035404550550.00.51.01.52.02.53.03.54.04.55.05.56.0Sitealpha

#### Fitted tree

Options

- Partitions
- 1
- Models
- Global MG94xREV
- Nucleotide GTR

- Hide Legend
- GrayScale

Export 

- PNG
- SVG
- Newick File

TestBackgroundLength = 0.01141837775089771Length = 0Length = 0.005650374309668017Length = 0.00496341314512962Length = 0.04373990789177799Length = 0.03007321549212623Length = 0.1506555712403421Length = 0.0309952609328117Length = 0.2076996068377695Length = 0.06305017330720755Length = 0.06350079960994272Length = 0.005627606443674139Length = 0.03230291042122855Length = 0Length = 0.1034702021771859Length = 0.08442176986698845Length = 0.1459672809071342Length = 0.3719747857665682Length = 0.05978762109857879Length = 0.1456153545830473DBIP\_XM\_017237939DANA\_XM\_014907649DRHO\_XM\_017134245DSUZ\_XM\_017084964DBIA\_XM\_017102001DEUG\_XM\_017210758DYAK\_GE19241\_BARAB\_DMEL\_BARAB\_CG13749\_DSIM\_BARAB\_XM\_002080640\_3\_173\_346DMAU\_BARAB\_XM\_033300133\_1\_282\_455DSEC\_BARAB\_XM\_002032966\_2\_61\_234SLEB\_IM240.0500.100.150.200.250.300.35

#### Model fits

| Model | AIC_C_ | log L | Parameters | Rate distributions |
| --- | --- | --- | --- | --- |
| Nucleotide GTR | 2249.49 | -1095.32 | 29 |  |
| Global MG94xREV | 2081.33 | -1003.75 | 35 | non-synonymous/synonymous rate ratio for \*test\* \| \|; 100% \| @ \| 0.137 |

This table reports a statistical summary of the models fit to the data. Here, **MG94** refers to the MG94xREV baseline model that infers a single ω rate category per branch.

×

#### Error

This is my error message

Close

Datamonkey is funded jointly by MIDAS and NIH award R01 GM093939
