## supplementary data file 3 for "Repeated truncation of a modular antimicrobial peptide gene for neural context": FUBARDatamonkey Adaptive Evolution Server.html

Datamonkey Adaptive Evolution Server


Methods and Tools

aBSREL
SpiderMonkey/BGM
BUSTED
Contrast-FEL
FADE
FEL
FUBAR
GARD
HIV-TRACE
MULTI-HIT
MEME
RELAX
SLAC
All Methods

Job Queue
Usage statistics

API

API Info
Get API Key
Check Key Status

Citations
Help
COVID-19
Blog
 Classic

Methods and Tools

aBSREL
BUSTED
FADE Beta
FEL
FUBAR
GARD
HIV-TRACE
MEME
RELAX
SLAC
All Methods

Job Queue
Usage statistics
Citations
Help
 Classic

- summary
- plot
- table
- tree
- fits

×Close**Error!**

### Fast Unconstrained Bayesian AppRoximation results summary

INPUT DATA |609599e0238adf71a515d74e|12 sequences |58 sites

Export

- Original file
- Analysis log
- Save JSON
- View JSON

#### Alignment viewer

×

Close

FUBAR **found evidence** of

episodic positive/diversifying selection at 0 sites

episodic negative/purifying selection at 29 sites

with posterior probability of.

---

See here for more information about the FUBAR method.  
Please cite PMID 23420840 if you use this result in a publication, presentation, or other scientific work.

#### Posterior rate distribution

Site

Export Chart to SVG Export Chart to PNG

00.070.140.210.290.360.430.50.570.640.710.790.8611.232.817.1215.5229.365000.070.140.210.290.360.430.50.570.640.710.790.8611.232.817.1215.5229.3650Synonymous substitution rateNon-synonymous substitution rateω0110

This graph shows the posterior distribution over the discretized rate grid. The size of a dot is proportional to the posterior weight allocated to that gridpoint, and the color shows the intensity of selection. Site-specific distributions can be viewed by entering a site number in the input box above the figure. When this is empty, the alignment-wide distribution will be shown.

#### FUBAR Site Table

Positively selected sites with evidence are highlighted in green.

Negatively selected sites with evidence are highlighted in black.

Showing entries 1 through 20 out of 58.

Export Table to CSV

| Site | Partition | α | β | β-α | Prob[α>β] | Prob[α<β] | BayesFactor[α<β] |  |  |
| --- | --- | --- | --- | --- | --- | --- | --- | --- | --- |
| 1 | 1 | 14.857 | 0.255 | -14.602 | 0.996 | 0.003 | 0.004 | 0.000 | 0.000 |
| 2 | 1 | 3.174 | 0.884 | -2.290 | 0.594 | 0.339 | 0.751 | 0.000 | 0.000 |
| 3 | 1 | 1.023 | 1.048 | 0.025 | 0.362 | 0.550 | 1.792 | 0.000 | 0.000 |
| 4 | 1 | 3.883 | 0.466 | -3.417 | 0.879 | 0.090 | 0.145 | 0.000 | 0.000 |
| 5 | 1 | 2.154 | 0.663 | -1.491 | 0.628 | 0.308 | 0.652 | 0.000 | 0.000 |
| 6 | 1 | 13.100 | 0.493 | -12.607 | 0.999 | 0.000 | 0.001 | 0.000 | 0.000 |
| 7 | 1 | 0.818 | 0.907 | 0.089 | 0.357 | 0.561 | 1.876 | 0.000 | 0.000 |
| 8 | 1 | 2.039 | 0.653 | -1.387 | 0.631 | 0.306 | 0.646 | 0.000 | 0.000 |
| 9 | 1 | 8.000 | 0.232 | -7.768 | 0.997 | 0.002 | 0.003 | 0.000 | 0.000 |
| 10 | 1 | 16.160 | 0.213 | -15.947 | 0.999 | 0.000 | 0.001 | 0.000 | 0.000 |
| 11 | 1 | 26.887 | 0.443 | -26.444 | 0.999 | 0.000 | 0.001 | 0.000 | 0.000 |
| 12 | 1 | 0.976 | 0.476 | -0.501 | 0.760 | 0.185 | 0.333 | 0.000 | 0.000 |
| 13 | 1 | 0.594 | 0.384 | -0.209 | 0.519 | 0.412 | 1.026 | 0.000 | 0.000 |
| 14 | 1 | 2.365 | 0.243 | -2.122 | 0.967 | 0.023 | 0.034 | 0.000 | 0.000 |
| 15 | 1 | 5.171 | 0.619 | -4.551 | 0.922 | 0.054 | 0.084 | 0.000 | 0.000 |
| 16 | 1 | 2.191 | 0.210 | -1.981 | 0.970 | 0.021 | 0.031 | 0.000 | 0.000 |
| 17 | 1 | 1.466 | 0.247 | -1.219 | 0.924 | 0.055 | 0.085 | 0.000 | 0.000 |
| 18 | 1 | 14.589 | 0.594 | -13.996 | 0.991 | 0.005 | 0.008 | 0.000 | 0.000 |
| 19 | 1 | 8.209 | 0.505 | -7.704 | 0.929 | 0.053 | 0.082 | 0.000 | 0.000 |
| 20 | 1 | 1.174 | 0.469 | -0.705 | 0.788 | 0.163 | 0.285 | 0.000 | 0.000 |

#### Fitted tree

Options

- Partitions
- 1
- Models
- Nucleotide GTR

- Hide Legend
- GrayScale

Export 

- PNG
- SVG
- Newick File

Length = 0.0117436838787748Length = 0Length = 0.005815703920995877Length = 0.006842644445110527Length = 0.04146386631620481Length = 0.03272920128077526Length = 0.1306435053898901Length = 0.02058964353255576Length = 0.1794897481724929Length = 0.05628725964363076Length = 0.06160794883899332Length = 0.01161909852728024Length = 0.03199148293000968Length = 0.02229854931370254Length = 0.07757382083785726Length = 0.08306270775386806Length = 0.1178408038675837Length = 0.2581073504870456Length = 0.06414733385054974Length = 0.09838510644657933DBIP\_XM\_017237939DANA\_XM\_014907649DRHO\_XM\_017134245DSUZ\_XM\_017084964DBIA\_XM\_017102001DEUG\_XM\_017210758DYAK\_GE19241\_BARAB\_DMEL\_BARAB\_CG13749\_DSIM\_BARAB\_XM\_002080640\_3\_173\_346DMAU\_BARAB\_XM\_033300133\_1\_282\_455DSEC\_BARAB\_XM\_002032966\_2\_61\_234SLEB\_IM240.0200.0400.0600.0800.100.120.140.160.180.200.220.240.26

#### Model fits

| Model | AIC_C_ | log L | Parameters | Rate distributions |
| --- | --- | --- | --- | --- |
| Nucleotide GTR | 2249.49 | -1095.32 | 29 |  |

This table reports a statistical summary of the models fit to the data. Here, **MG94** refers to the MG94xREV baseline model that infers a single ω rate category per branch.

×

#### Error

This is my error message

Close

Datamonkey is funded jointly by MIDAS and NIH award R01 GM093939
