## supplementary data file 3 for "Repeated truncation of a modular antimicrobial peptide gene for neural context": SLACDatamonkey Adaptive Evolution Server.html

Datamonkey Adaptive Evolution Server


Methods and Tools

aBSREL
SpiderMonkey/BGM
BUSTED
Contrast-FEL
FADE
FEL
FUBAR
GARD
HIV-TRACE
MULTI-HIT
MEME
RELAX
SLAC
All Methods

Job Queue
Usage statistics

API

API Info
Get API Key
Check Key Status

Citations
Help
COVID-19
Blog
 Classic

Methods and Tools

aBSREL
BUSTED
FADE Beta
FEL
FUBAR
GARD
HIV-TRACE
MEME
RELAX
SLAC
All Methods

Job Queue
Usage statistics
Citations
Help
 Classic

- summary
- information
- table
- graph
- tree
- phylo alignment

×Close**Error!**

### Single-Likelihood Ancestor Counting results summary

INPUT DATA |609599ed238adf71a515d760|12 sequences |58 sites

Export

- Original file
- Analysis log
- Save JSON
- View JSON

#### Alignment viewer

×

Close

SLAC **found evidence** of pervasive

positive/diversifying selection at 0 sites

negative/purifying selection at 20 sites

with p-value threshold of.

---

See here for more information about the SLAC method.  
Please cite PMID 15703242 if you use this result in a publication, presentation, or other scientific work.

#### Partition information

| Partition | Sites | Branches | | Branch Length | | | Selected at p≤0.1 | |
| --- | --- | --- | --- | --- | --- | --- | --- | --- |
|  |  | Tested | Total | Tested | % of total | Total | Positive | Negative |
| 1 | 58 | 21 | 21 | 1.21 | 78.0% | 1.55 | 0 | 20 |

#### Model fits

| Model | AIC_C_ | log L | Parameters | Rate distributions |
| --- | --- | --- | --- | --- |
| Nucleotide GTR | 2249.49 | -1095.32 | 29 |  |
| Global MG94xREV | 2094.77 | -1009.36 | 36 | non-synonymous/synonymous rate ratio for \*test\* \| \|; 100% \| @ \| 0.162 |

#### Execution time

| Task | Time | % |
| --- | --- | --- |
| Total time | 00:00:17 |  |
| Model fitting | 00:00:01 | 5.88% |
| Primary SLAC analysis | 00:00:01 | 5.88% |
| Ancestor sampling analysis | 00:00:15 | 88.24% |

#### SLAC Site Table

- Variable sites only
- Color cells based on MLE-median

Display

- AVERAGED
- RESOLVED

Ambiguities

- Partition
- Site
- ES
- EN
- S
- N
- P[S]
- dS
- dN
- dN-dS
- P [dN/dS > 1]
- P [dN/dS < 1]
- Total branch length

Site

is in [

,]

- AND
- OR

AND

Add filter

× 

**Color legend:** MLE is  is much less is the same as is much greater  than the sampled median.

Default table shading is used to indicate the magnitude of difference between the estimate of a specific quantity using the MLE ancestral state reconstruction, and the median of the estimate using a sample from the distribution of ancestral state reconstructions.

You can mouse over the cells to see individual sampling intervals.

Showing entries 1 through 20 out of 58.

Export Table to CSV

| Partition | Site | ES | EN | S | N | P[S] | dS | dN | dN-dS | P [dN/dS > 1] | P [dN/dS < 1] | Total branch length |
| --- | --- | --- | --- | --- | --- | --- | --- | --- | --- | --- | --- | --- |
| 1 | 1 | 0.594 | 1.85 | 3.00 | 0.00 | 0.243 | 5.05 | 0.00 | -3.26 | 1.00 | 0.0143 | 1.55 |
| 1 | 2 | 0.624 | 1.98 | 2.00 | 3.00 | 0.240 | 3.20 | 1.52 | -1.09 | 0.907 | 0.346 | 1.55 |
| 1 | 3 | 0.888 | 2.10 | 1.50 | 4.50 | 0.297 | 1.69 | 2.14 | 0.290 | 0.589 | 0.726 | 1.55 |
| 1 | 4 | 0.579 | 2.00 | 2.00 | 1.00 | 0.225 | 3.45 | 0.500 | -1.91 | 0.989 | 0.129 | 1.55 |
| 1 | 5 | 0.590 | 2.40 | 1.00 | 2.00 | 0.197 | 1.69 | 0.832 | -0.557 | 0.899 | 0.482 | 1.55 |
| 1 | 6 | 0.989 | 1.97 | 5.00 | 1.00 | 0.335 | 5.06 | 0.508 | -2.94 | 0.999 | 0.0181 | 1.55 |
| 1 | 7 | 0.907 | 2.07 | 1.50 | 3.50 | 0.305 | 1.65 | 1.69 | 0.0252 | 0.674 | 0.660 | 1.55 |
| 1 | 8 | 0.600 | 2.40 | 1.50 | 2.50 | 0.200 | 2.50 | 1.04 | -0.940 | 0.896 | 0.386 | 1.55 |
| 1 | 9 | 1.00 | 1.99 | 4.00 | 0.00 | 0.334 | 4.00 | 0.00 | -2.58 | 1.00 | 0.0124 | 1.55 |
| 1 | 10 | 1.00 | 2.00 | 5.00 | 0.00 | 0.333 | 5.00 | 0.00 | -3.23 | 1.00 | 0.00412 | 1.55 |
| 1 | 11 | 0.577 | 2.42 | 4.00 | 1.00 | 0.192 | 6.93 | 0.413 | -4.21 | 1.00 | 0.00579 | 1.55 |
| 1 | 12 | 1.00 | 2.00 | 2.00 | 1.00 | 0.333 | 2.00 | 0.500 | -0.968 | 0.963 | 0.259 | 1.55 |
| 1 | 13 | 0.584 | 2.40 | 0.00 | 1.00 | 0.196 | 0.00 | 0.416 | 0.269 | 0.804 | 1.00 | 1.55 |
| 1 | 14 | 1.00 | 1.87 | 3.00 | 0.00 | 0.348 | 3.00 | 0.00 | -1.94 | 1.00 | 0.0422 | 1.55 |
| 1 | 15 | 0.993 | 2.01 | 4.00 | 2.00 | 0.331 | 4.03 | 0.997 | -1.96 | 0.983 | 0.0980 | 1.55 |
| 1 | 16 | 1.00 | 2.00 | 3.00 | 0.00 | 0.333 | 3.00 | 0.00 | -1.94 | 1.00 | 0.0370 | 1.55 |
| 1 | 17 | 1.00 | 1.99 | 2.00 | 0.00 | 0.335 | 2.00 | 0.00 | -1.29 | 1.00 | 0.112 | 1.55 |
| 1 | 18 | 0.574 | 2.43 | 4.00 | 2.00 | 0.191 | 6.97 | 0.824 | -3.97 | 0.999 | 0.0144 | 1.55 |
| 1 | 19 | 0.621 | 1.85 | 2.00 | 1.00 | 0.251 | 3.22 | 0.539 | -1.73 | 0.984 | 0.157 | 1.55 |
| 1 | 20 | 0.979 | 2.02 | 2.00 | 1.00 | 0.326 | 2.04 | 0.495 | -0.999 | 0.965 | 0.250 | 1.55 |

#### SLAC Site Graph

X-axis:

- Site
- ES
- EN
- S
- N
- P[S]
- dS
- dN
- dN-dS
- P [dN/dS > 1]
- P [dN/dS < 1]
- Total branch length

Site

Y-axis:

- ES
- EN
- S
- N
- P[S]
- dS
- dN
- dN-dS
- P [dN/dS > 1]
- P [dN/dS < 1]
- Total branch length

dN-dS

Ambiguities 

- AVERAGED
- RESOLVED

RESOLVED

Export to PNG Export to SVG

510152025303540455055-4.0-3.5-3.0-2.5-2.0-1.5-1.0-0.50.00.5SitedN-dS

#### Fitted tree

Options

- Partitions
- 1
- Models
- Global MG94xREV
- Nucleotide GTR

- Hide Legend
- GrayScale

Export 

- PNG
- SVG
- Newick File

TestBackgroundLength = 0.01386203916957873Length = 0Length = 0Length = 0.006864755249178939Length = 0.008076937892806961Length = 0.04894322300655905Length = 0.03863297708167415Length = 0.1542093100989249Length = 0.02430365531652305Length = 0.2118665612416413Length = 0.06644049737565492Length = 0.07272094589560989Length = 0.01371498079844063Length = 0.03776218723583095Length = 0.02632081782872176Length = 0.09156678211781427Length = 0.09804576828710719Length = 0.13909722501467Length = 0.3046654047691284Length = 0.07571839157451288Length = 0.1161320598667153DBIP\_XM\_017237939DANA\_XM\_014907649DRHO\_XM\_017134245DSUZ\_XM\_017084964DBIA\_XM\_017102001DEUG\_XM\_017210758DYAK\_GE19241\_BARAB\_DMEL\_BARAB\_CG13749\_DSIM\_BARAB\_XM\_002080640\_3\_173\_346DMAU\_BARAB\_XM\_033300133\_1\_282\_455DSEC\_BARAB\_XM\_002032966\_2\_61\_234SLEB\_IM240.0500.100.150.200.250.30

#### SLAC Phylogenetic Alignment

Current site:Show amino acidcodonnone on treePNG

TotalNonsynonymousSynonymous3.02.82.62.42.22.01.81.61.41.21.00.80.60.40.20.0Substitution countsNode3Node6SLEB\_IM24DBIP\_XM\_017237939DANA\_XM\_014907649DRHO\_XM\_017134245Node8Node9Node12DSUZ\_XM\_017084964DBIA\_XM\_017102001DEUG\_XM\_017210758Node14DYAK\_GE19241\_BARAB\_Node16DMEL\_BARAB\_CG13749\_Node18DSIM\_BARAB\_XM\_002080Node20DMAU\_BARAB\_XM\_033300DSEC\_BARAB\_XM\_002032CAGCAGCAGCAACAGCAACAGCAGCAGCAGCAGCAGCAGCAGCAGCAGCAGCAGCAGCAGCAAQQQQQQQQQQQQQQQQQQQQQCodon 1

×

#### Error

This is my error message

Close

Datamonkey is funded jointly by MIDAS and NIH award R01 GM093939
