## supplementary data file 3 for "Repeated truncation of a modular antimicrobial peptide gene for neural context": README.rtf

HyPhy analysis outputs (Datamonkey webserver outputs) are stored for each gene locus in a separate folder. 

A summary of key results is also provided in each folder, copied from the Datamonkey webserver output (and these results can be found within excel files of these folders).
