## supplementary data file 3 for "Repeated truncation of a modular antimicrobial peptide gene for neural context": Datamonkey Adaptive Evolution Server.html

Methods and Tools

aBSREL
SpiderMonkey/BGM
BUSTED
Contrast-FEL
FADE
FEL
FUBAR
GARD
HIV-TRACE
MULTI-HIT
MEME
RELAX
SLAC
All Methods

Job Queue
Usage statistics

API

API Info
Get API Key
Check Key Status

Citations
Help
COVID-19
Blog
 Classic

Methods and Tools

aBSREL
BUSTED
FADE Beta
FEL
FUBAR
GARD
HIV-TRACE
MEME
RELAX
SLAC
All Methods

Job Queue
Usage statistics
Citations
Help
 Classic

- summary
- tree
- table
- model fits

×Close**Error!**

##### adaptive Branch Site REL results summary

INPUT DATA |6095a6bb238adf71a515dd81|24 sequences |58 sites

Export

- Original file
- Analysis log
- Save JSON
- View JSON

###### Alignment viewer

×

Close

aBSREL **found evidence** of episodic diversifying selection on **1** out of **44** branches in your phylogeny.

A total of **44** branches were formally tested for diversifying selection. Significance was assessed using the Likelihood Ratio Test at a threshold of p ≤ 0.05, after correcting for multiple testing. Significance and number of rate categories inferred at each branch are provided in the detailed results table.

---

See here for more information about this method.  
Please cite PMID 25697341 if you use this result in a publication, presentation, or other scientific work.

###### Tree summary

| ω rate classes | # of branches | % of branches | % of tree length | # under selection |
| --- | --- | --- | --- | --- |
| 1 | 38 | 86% | 0.24% | 0 |
| 2 | 6 | 14% | 100% | 1 |

This table contains a summary of the inferred aBSREL model complexity. Each row provides information about the branches that were best described by the given number of ω rate categories.

###### Fitted tree

Options

- Models
- Full adaptive model
- Baseline MG94xREV

- Hide Legend
- GrayScale

Export 

- PNG
- SVG
- Newick File

00.010.10.512510ωLength = 0.005490720627682237Length = 0Length = 0Length = 0.005467984237327723Length = 0.01090958376533652Length = 0.0240761621464922Length = 0.02760316393966488Length = 0.09229115499336737Length = 0.07871355773648374Length = 0.02404413679551904Length = 0.04462780932208216Length = 0.118306181882418Length = 0.04624042368125375Length = 0.05866879246710729Length = 0.04685718830904137Length = 0.06676149397853735Length = 0.009181529407681193Length = 0.03492034534912974Length = 0.00463816401504084Length = 0.09628870443410553Length = 4.142592437824455Length = 0.04840163669227976Length = 0.02424243971526773Length = 0.00569801466415147Length = 0Length = 534.8999320325061Length = 0.03501851782570966Length = 0.02295668837904526Length = 0.01428316951234084Length = 0.01169095336310318Length = 0.04119199325766765Length = 0.09517154808757743Length = 0.1450298734673382Length = 0Length = 36.59361919336702Length = 0.02648733544535357Length = 0.02087275140755344Length = 0.01181734680304538Length = 0.06891161209741042Length = 0.04220328989368644Length = 0Length = 0.6677835432172854Length = 0.1188735511168711Length = 0.2808483260593987DTRO\_BARAADINS\_BARAADWIL\_GK10648DPAU\_BARAA\_STR\_L06\_2DPAU\_BARAA\_STR\_L12\_2DSUB\_XM\_034794836\_LOC117890147\_CDSDGUA\_XM\_034286156\_LOC117592412\_CDSDOBS\_XM\_022367329\_LOC111074527\_CDSDPSE\_GA15751DMIR\_XM\_017294356\_LOC108160378\_CDSDPER\_XM\_002015800\_LOC6590356\_CDSDFIC\_XM\_017198865\_LOC108096915\_CDSDRHO\_XM\_017121875\_LOC108043260\_CDSDELE\_XM\_017265456\_LOC108141863\_CDSDEUG\_XM\_017223845\_LOC108113294\_CDSDTAK\_XM\_017141080\_LOC108057013\_CDSDSUZ\_XM\_017075092\_LOC108010239\_CDSDBIA\_XM\_017090984\_LOC108022142\_CDSDYAK\_GE13684\_CDSDMEL\_CG30285\_BARAC\_REVERSED\_DSEC\_GM15695\_CDSDSIM\_GD25174\_CDSDMAU\_XM\_033296995\_LOC117136220\_CDSSLEB\_XM\_030519534

###### Detailed results

| Name | B | LRT | Test p-value | Uncorrected p-value | ω distribution over sites |
| --- | --- | --- | --- | --- | --- |
| DPAU\_BARAA\_STR\_L06\_2 | 0.0000 | 13.8412 | 0.0148 | 0.0003 | ω_1_ = 0.00 (98%) ω_2_ = 100000 (1.9%) |
| Node38 | 0.0000 | 5.6999 | 0.8922 | 0.0207 | ω_1_ = 0.00 (98%) ω_2_ = 3850 (1.8%) |
| DBIA\_XM\_017090984\_LOC108022142\_CDS | 0.0000 | 0.0000 | 1.0000 | 1.0000 | ω_1_ = 0.124 (100%) |
| DELE\_XM\_017265456\_LOC108141863\_CDS | 0.0000 | 0.0000 | 1.0000 | 1.0000 | ω_1_ = 0.177 (100%) |
| DEUG\_XM\_017223845\_LOC108113294\_CDS | 0.0000 | 0.0000 | 1.0000 | 1.0000 | ω_1_ = 0.126 (100%) |
| DFIC\_XM\_017198865\_LOC108096915\_CDS | 0.0000 | 0.0000 | 1.0000 | 1.0000 | ω_1_ = 0.0414 (100%) |
| DGUA\_XM\_034286156\_LOC117592412\_CDS | 0.0000 | 0.0000 | 1.0000 | 1.0000 | ω_1_ = 0.0971 (100%) |
| DINS\_BARAA | 0.0000 | 0.0000 | 1.0000 | 1.0000 | ω_1_ = 0.0266 (100%) |
| DMAU\_XM\_033296995\_LOC117136220\_CDS | 0.0000 | 0.0000 | 1.0000 | 1.0000 | ω_1_ = 0.00 (100%) |
| DMEL\_CG30285\_BARAC\_REVERSED\_ | 0.0000 | 0.0000 | 1.0000 | 1.0000 | ω_1_ = 0.277 (100%) |
| DMIR\_XM\_017294356\_LOC108160378\_CDS | 0.0000 | 0.0000 | 1.0000 | 1.0000 | ω_1_ = 0.00 (100%) |
| DOBS\_XM\_022367329\_LOC111074527\_CDS | 0.0000 | 0.0000 | 1.0000 | 1.0000 | ω_1_ = 0.00 (100%) |
| DPAU\_BARAA\_STR\_L12\_2 | 0.0000 | 0.0000 | 1.0000 | 1.0000 | ω_1_ = 1.00 (100%) |
| DPER\_XM\_002015800\_LOC6590356\_CDS | 0.0000 | 2.1630 | 1.0000 | 0.1300 | ω_1_ = 10000000000 (100%) |
| DPSE\_GA15751 | 0.0000 | 0.0000 | 1.0000 | 1.0000 | ω_1_ = 1.00 (100%) |
| DRHO\_XM\_017121875\_LOC108043260\_CDS | 0.0000 | 0.0000 | 1.0000 | 1.0000 | ω_1_ = 0.0614 (100%) |
| DSEC\_GM15695\_CDS | 0.0000 | 0.0000 | 1.0000 | 1.0000 | ω_1_ = 0.00 (100%) |
| DSIM\_GD25174\_CDS | 0.0000 | 0.0000 | 1.0000 | 1.0000 | ω_1_ = 1.00 (100%) |
| DSUB\_XM\_034794836\_LOC117890147\_CDS | 0.0000 | 0.0000 | 1.0000 | 1.0000 | ω_1_ = 0.00 (100%) |
| DSUZ\_XM\_017075092\_LOC108010239\_CDS | 0.0000 | 0.0000 | 1.0000 | 1.0000 | ω_1_ = 0.0926 (100%) |
| DTAK\_XM\_017141080\_LOC108057013\_CDS | 0.0000 | 2.8604 | 1.0000 | 0.0899 | ω_1_ = 0.163 (93%) ω_2_ = 11.4 (7.2%) |
| DTRO\_BARAA | 0.0000 | 0.0000 | 1.0000 | 1.0000 | ω_1_ = 1.00 (100%) |
| DWIL\_GK10648 | 0.0000 | 0.0000 | 1.0000 | 1.0000 | ω_1_ = 0.111 (100%) |
| DYAK\_GE13684\_CDS | 0.0000 | 0.0000 | 1.0000 | 1.0000 | ω_1_ = 0.204 (100%) |
| Node12 | 0.0000 | 0.0000 | 1.0000 | 1.0000 | ω_1_ = 0.268 (100%) |
| Node13 | 0.0000 | 0.0000 | 1.0000 | 1.0000 | ω_1_ = 0.0436 (100%) |
| Node14 | 0.0000 | 0.0000 | 1.0000 | 1.0000 | ω_1_ = 0.00 (100%) |
| Node17 | 0.0000 | 0.3616 | 1.0000 | 0.3599 | ω_1_ = 10000000000 (100%) |
| Node19 | 0.0000 | 5.0412 | 1.0000 | 0.0291 | ω_1_ = 0.0870 (96%) ω_2_ = 100000 (3.8%) |
| Node24 | 0.0000 | 0.0000 | 1.0000 | 1.0000 | ω_1_ = 0.0178 (100%) |
| Node25 | 0.0000 | 0.0000 | 1.0000 | 1.0000 | ω_1_ = 0.00 (100%) |
| Node27 | 0.0000 | 0.0000 | 1.0000 | 1.0000 | ω_1_ = 0.00 (100%) |
| Node28 | 0.0000 | 0.0000 | 1.0000 | 1.0000 | ω_1_ = 0.00 (100%) |
| Node3 | 0.0000 | 0.3719 | 1.0000 | 0.3575 | ω_1_ = 0.00 (81%) ω_2_ = 1.68 (19%) |
| Node31 | 0.0000 | 0.0000 | 1.0000 | 1.0000 | ω_1_ = 0.00 (100%) |
| Node33 | 0.0000 | 0.0000 | 1.0000 | 1.0000 | ω_1_ = 0.182 (100%) |
| Node35 | 0.0000 | 0.0000 | 1.0000 | 1.0000 | ω_1_ = 0.00 (100%) |
| Node40 | 0.0000 | 0.0000 | 1.0000 | 1.0000 | ω_1_ = 0.00 (100%) |
| Node42 | 0.0000 | 0.0000 | 1.0000 | 1.0000 | ω_1_ = 0.00 (100%) |
| Node44 | 0.0000 | 0.0000 | 1.0000 | 1.0000 | ω_1_ = 1.00 (100%) |
| Node5 | 0.0000 | 0.0000 | 1.0000 | 1.0000 | ω_1_ = 0.00 (100%) |
| Node7 | 0.0000 | 0.5023 | 1.0000 | 0.3291 | ω_1_ = 10000000000 (100%) |
| Node9 | 0.0000 | 0.0000 | 1.0000 | 1.0000 | ω_1_ = 0.583 (100%) |
| SLEB\_XM\_030519534 | 0.0000 | 0.4112 | 1.0000 | 0.3485 | ω_1_ = 0.00 (99%) ω_2_ = 45.2 (1.4%) |

###### aBSREL Site Proportion Chart

×

###### ω distribution

### **DBIA\_XM\_017090984\_LOC108022142\_CDS**

SVG PNG

Neutrality (ω=1)ω0.000010.00010.0010.010.1110100100010000Proportion of sites0%10%20%30%40%50%60%70%80%90%100%

Close

###### Model fits

| Model | AIC_C_ | log L | Parameters |
| --- | --- | --- | --- |
| Nucleotide GTR | 3267.69 | -1580.15 | 53 |
| Baseline MG94xREV | 3011.30 | -1395.50 | 102 |
| Full adaptive model | 2939.95 | -1345.71 | 114 |

This table reports a statistical summary of the models fit to the data. Here, **Baseline MG94xREV** refers to the MG94xREV baseline model that infers a single ω rate category per branch. **Full adaptive model** refers to the adaptive aBSREL model that infers an optimized number of ω rate categories per branch.

×

###### Error

This is my error message

Close

Datamonkey is funded jointly by MIDAS and NIH award R01 GM093939
