## supplementary data file 3 for "Repeated truncation of a modular antimicrobial peptide gene for neural context": Summary stum locus.txt

FEL found evidence of
 pervasive positive/diversifying selection at 0 sites
 pervasive negative/purifying selection at 47 sites
with p-value threshold of 0.1


FUBAR found evidence of
 episodic positive/diversifying selection at 0 sites
 episodic negative/purifying selection at 39 sites
with posterior probability of 0.9


SLAC found evidence of pervasive
 positive/diversifying selection at 0 sites
 negative/purifying selection at 36 sites
with p-value threshold of 0.1


aBSREL found evidence of episodic diversifying selection on 1 out of 44 branches in your phylogeny. A total of 44 branches were formally tested for diversifying selection. Significance was assessed using the Likelihood Ratio Test at a threshold of p ≤ 0.05, after correcting for multiple testing. Significance and number of rate categories inferred at each branch are provided in the detailed results table.

DPAU_BARAA_STR_L06_2
0.0000
13.8412
0.0148
0.0003
ω1 = 0.00 (98%)
ω2 = 100000 (1.9%)

BUSTED found evidence (LRT, p-value = 0.006 < .05) of gene-wide episodic diversifying selection in the selected test branches of your phylogeny. Therefore, there is evidence that at least one site on at least one test branch has experienced diversifying selection.
