## Supplementary figures and images for "Repeated truncation of a modular antimicrobial peptide gene for neural context"

### busted-chart.png

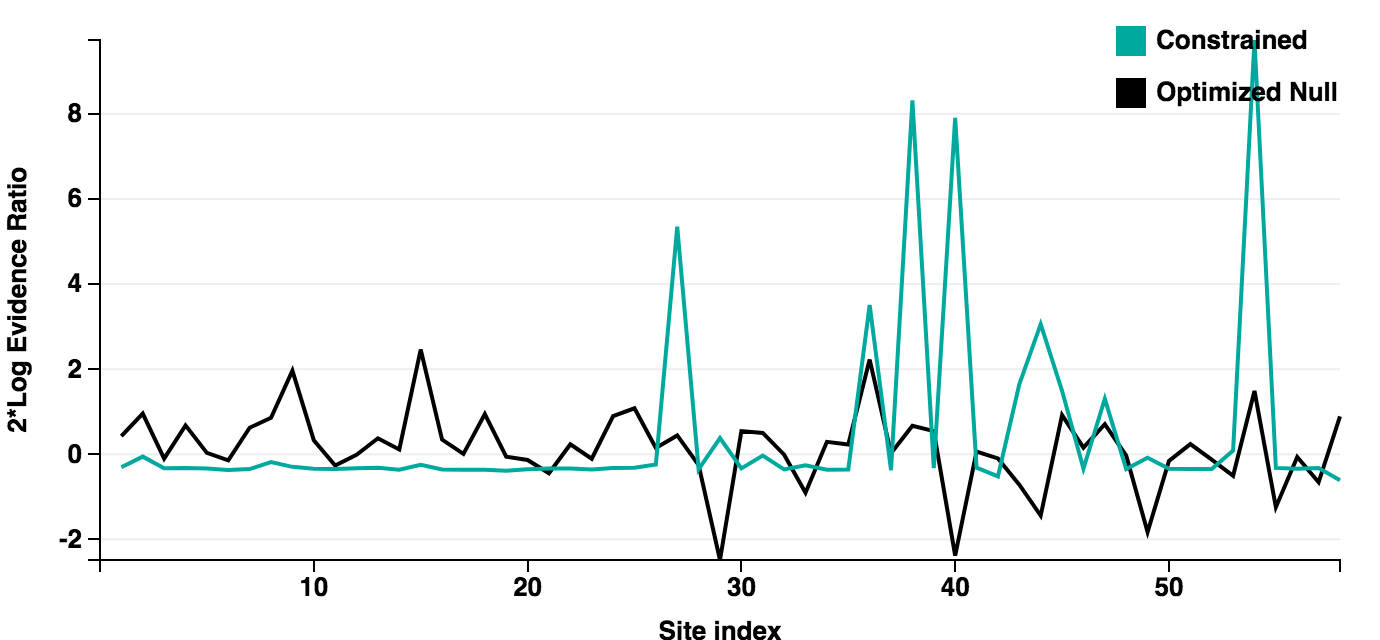
