## supplementary figures and tables for "Repeated truncation of a modular antimicrobial peptide gene for neural context"

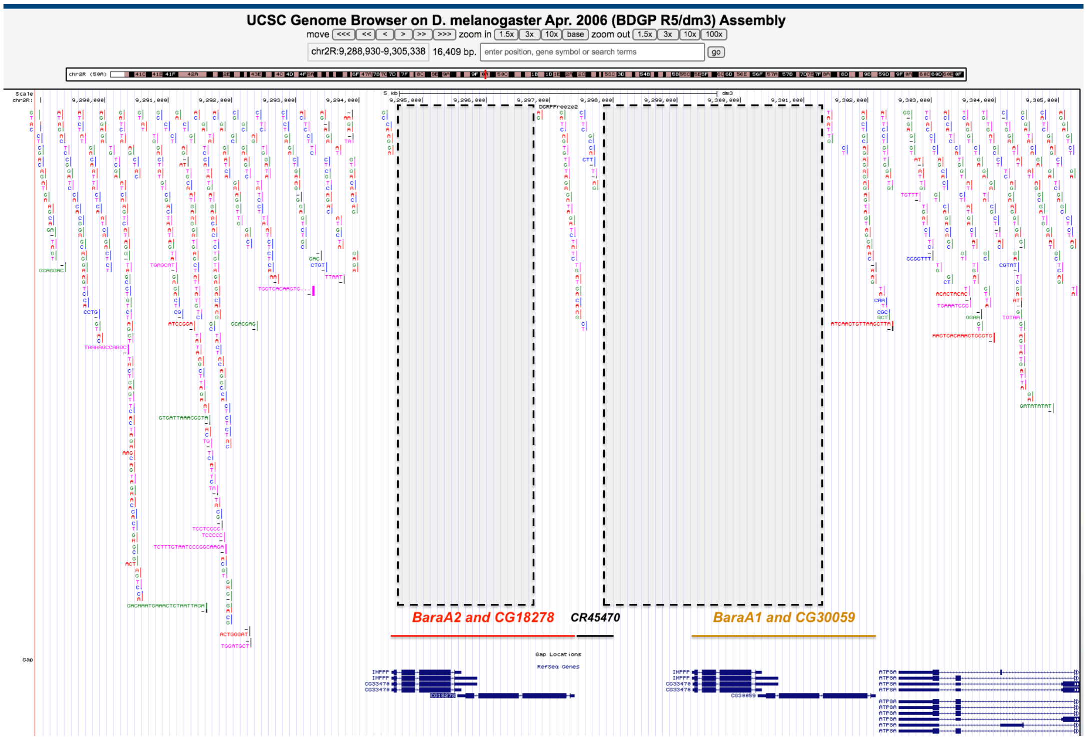


**Figure S1: The *BaraA* locus is poorly resolved in DGRP genome assemblies.** The *BaraA1* and *BaraA2* gene regions are totally devoid of mapped variants (dashed boxes). We speculate this is due to an artefact during genomic assembly, where reads mapping equally to the two identical *BaraA* genes were discarded as non-specific. This would explain why *BaraA* is typically discarded in RNAseq datasets using such measures in their pipeline, but not in microarray data from De Gregorio et al. (De Gregorio et al. 2002) where it is called "IM10."

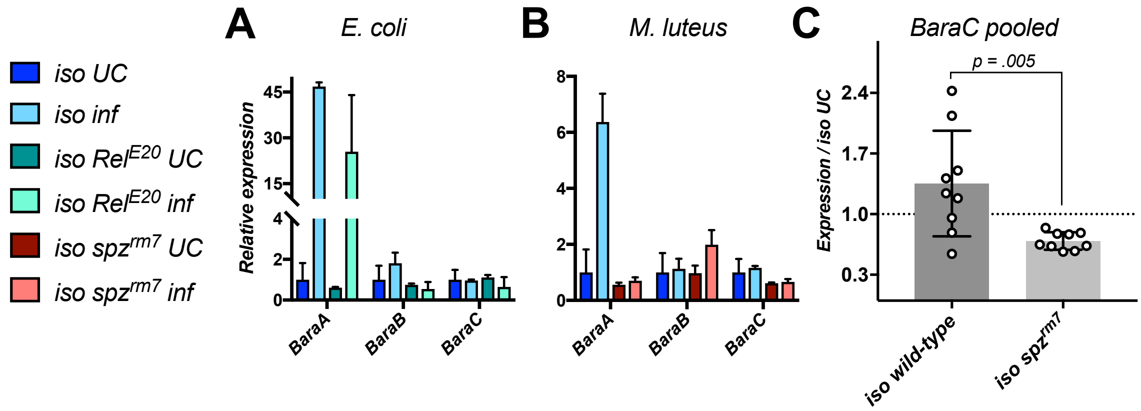


**Figure S2: Additional assays of Baramicin expression upon infection. A)** Neither *BaraB* nor *BaraC* are regulated by the Imd pathway, which is specifically stimulated by *E. coli* infection. **B)** Neither *BaraB* nor *BaraC* are induced after infection by *M. luteus*. **C)** *BaraC* levels were consistently depressed in $spz^{rm7}$ flies in the unchallenged condition (UC) or upon infection with *C. albicans* (**Fig.** 3B) or *M. luteus* (**Fig.** S4B). Data here are pooled for *iso* wild type or *iso* $spz^{rm7}$ flies without regard for infection treatment (student's t, p = .005).

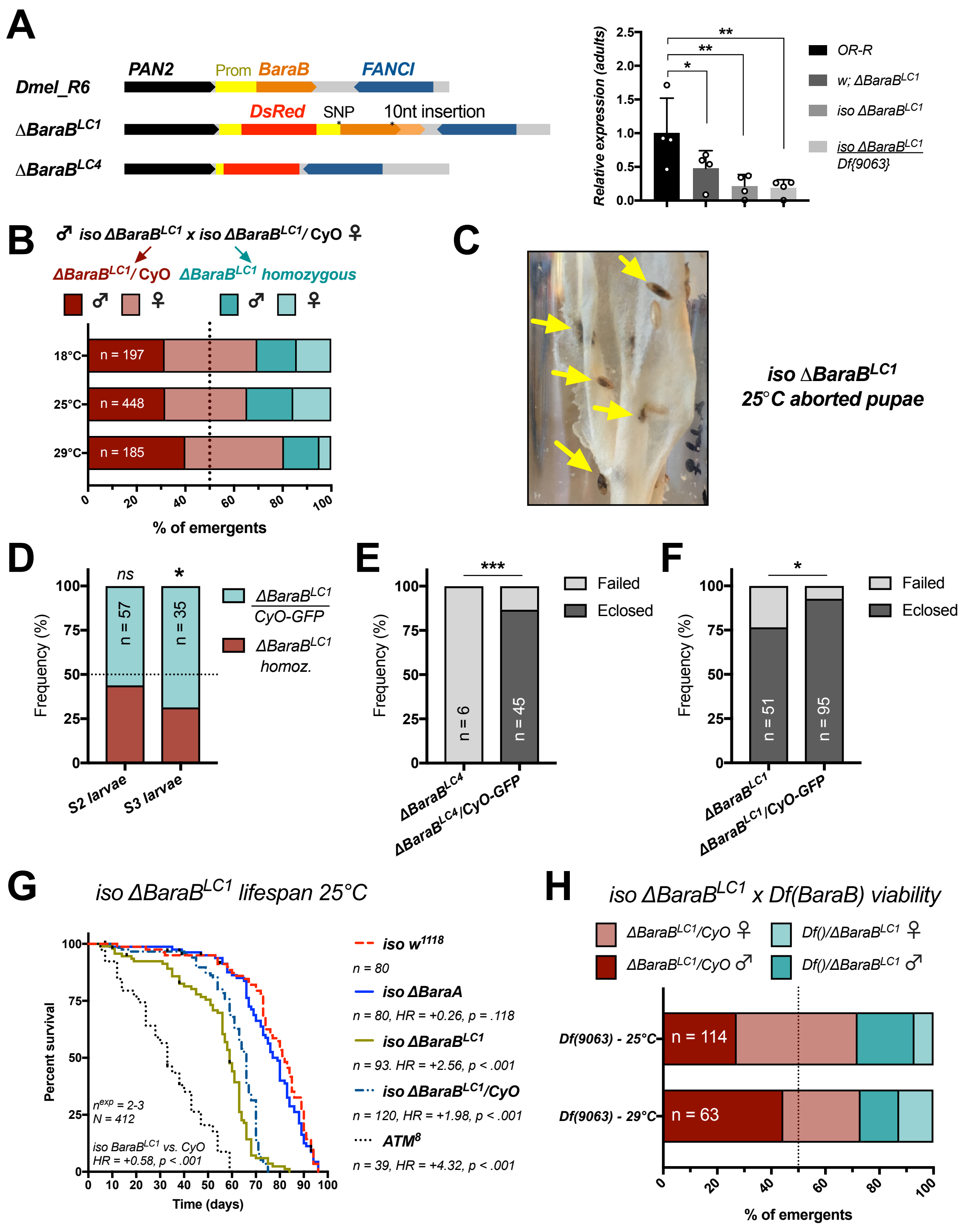


**Figure S3: *BaraB* mutation is highly deleterious, even in *ΔBaraB*$^{LC1}$ hypomorphs. A)** Diagram of *BaraB* mutant loci and qPCR showing the *ΔBaraB*$^{LC1}$ is a hypomorph mutation. Under our normal qPCR assay conditions, *BaraB* expression is not detected in *ΔBaraB*$^{LC1}$ homozygotes. However using highly concentrated cDNA beyond our assay's valid range (100ng/10µL reaction), we could detect *BaraB* transcript in *ΔBaraB*$^{LC1}$ flies. Quantification shown here is intended only to show that *BaraB* transcript can be recovered from *ΔBaraB*$^{LC1}$ homozygotes, and to give a sense of relative whole-fly expression levels. **B)** Emergent frequencies of *ΔBaraB*$^{LC1}$ flies at different temperatures. **C)** Aborted pupae (yellow arrows) are a common occurrence in *ΔBaraB* vials, and sometimes contain fully-developed adults that simply never eclosed. In **D-F:** ns = not significant, * = p < .05, *** p = < .001. **D)** The ratio of *ΔBaraB*$^{LC1}$/CyO-GFP to *ΔBaraB*$^{LC1}$ homozygous larvae drops between the S2 and S3 larval stages ($\chi^2$, p = .515 and p = .012 respectively). **E)** Frequency of successfully eclosing adults using *BaraB*$^{LC4}$/CyO-GFP flies. **F)** Frequency of successfully eclosing adults using *BaraB*$^{LC1}$/CyO-GFP flies. **G)** *BaraB* mutation negatively affects lifespan. *iso ΔBaraB*$^{LC1}$ homozygotes suffer reduced lifespan even relative to their *iso ΔBaraB*$^{LC1}$*/CyO* siblings. By comparison, *iso ΔBaraA* flies that used the same vector for mutant generation live as wild-type. ATM$^{8}$ flies suffer precocious neurodegeneration and are included as short-lived controls (Petersen et al. 2013). **H)** *ΔBaraB*$^{LC1}$ crossed to the genomic deficiency line (*Df(9063)*) supports a partial-lethal effect of *BaraB* mutation.

| Parents and temperature | Offspring | Sex | # eclosed | $\chi^2$ | p-value |
|---|---|---|---|---|---|
| ***TRiP*** | | | | | |
| y1 v1; P{TRiP.HMJ23624}attP40/CyO | Act-Gal4 / CyO | m | 26 | 25.687 | **p < .001** |
| Act-Gal4/CyO-GFP ; + | | f | 30 | | |
| 25°C | TRiP / CyO | m | 26 | | |
| | | f | 29 | | |
| | Act-Gal4 > BaraB-IR{TRiP} | m | 2 | | |
| | | f | 18 | | |
| y1 v1; P{TRiP.HMJ23624}attP40/CyO male | elav>Dcr2 ; +/CyO | m | 67 | 10.037 | **p < .02** |
| elav>Dcr2 ;; female | | f | 62 | | |
| 25°C | elav>Dcr2 ; +/BaraB-IR | m | 38 | | |
| | | f | 47 | | |
| y1 v1; P{TRiP.HMJ23624}attP40/CyO male | elav-Gal4; +/CyO | m | 29 | 13.593 | **p < .01** |
| elav-Gal4> ;; female | | f | 46 | | |
| 25°C | elav-Gal4 ; +/BaraB-IR | m | 21 | | |
| | | f | 22 | | |
| ***KK*** | | | | | |
| P{KK112854}VIE-260B | Act-Gal4 ; +/CyO | m | 27 | 45.187 | **p < .001** |
| Act-Gal4/CyO-GFP ; + | | f | 55 | | |
| 25°C | Act-Gal4 > BaraB-IR{KK} | m | 14 | | |
| | | f | 11 | | |
| P{KK112854}VIE-260B male | elav>Dcr2 ; +/BaraB-IR | m | 0 | 11.000 | **p < .001** |
| elav>Dcr2 ;; female | | f | 11 | | |
| 25°C | | | | | |
| P{KK112854}VIE-260B male | normal wing | m | 0 | 87.766 | **p < .001** |
| elav-Gal4> ;; female | | f | 0 | | |
| 29°C | nubbin-like wing | m | 28 | | |
| | | f | 57 | | |
| P{KK112854}VIE-260B male | normal wing | m | 37 | 32.133 | **p < .001** |
| elav-Gal4> ;; female | | f | 47 | | |
| 25°C | nubbin-like wing | m | 31 | | |
| | | f | 5 | | |

**Table S1: *BaraB* RNAi summary statistics.** Crosses used either the TRiP or KK *BaraB-IR* lines, driven by either *Actin5C-Gal4* or *elav-Gal4,* sometimes including *UAS-Dcr2*. Rearing at 29°C and inclusion of *UAS-Dcr2* increases the strength of RNA silencing. In the event there was no lethality, it was expected that emerging *elav>TRiP-IR* flies would follow simple mendelian inheritance. However both *elav>TRiP-IR* and *elav>Dcr2, TRiP-IR* resulted in partial lethality and occasional nubbin-like wings ($\chi^2$ p < .02). Crosses using *KK-IR* used homozygous flies, and so we did not assess lethality using mendelian inheritance. However using this construct, no adults emerged when *elav>Dcr2, KK-IR* flies were reared at 29°C. Rare emergents (N = 11 after three experiments) occurred at 25°C, all of which bore nubbin-like wings. Using *elav-Gal4* at 29°C without *Dcr2*, we observed greater numbers of emerging adults, but 100% of flies had nubbin-like wings. Finally, *elav>KK-IR* flies at 25°C suffered both partial lethality and nubbin-like wings, but normal-winged flies began emerging ($\chi^2$ p < .001).

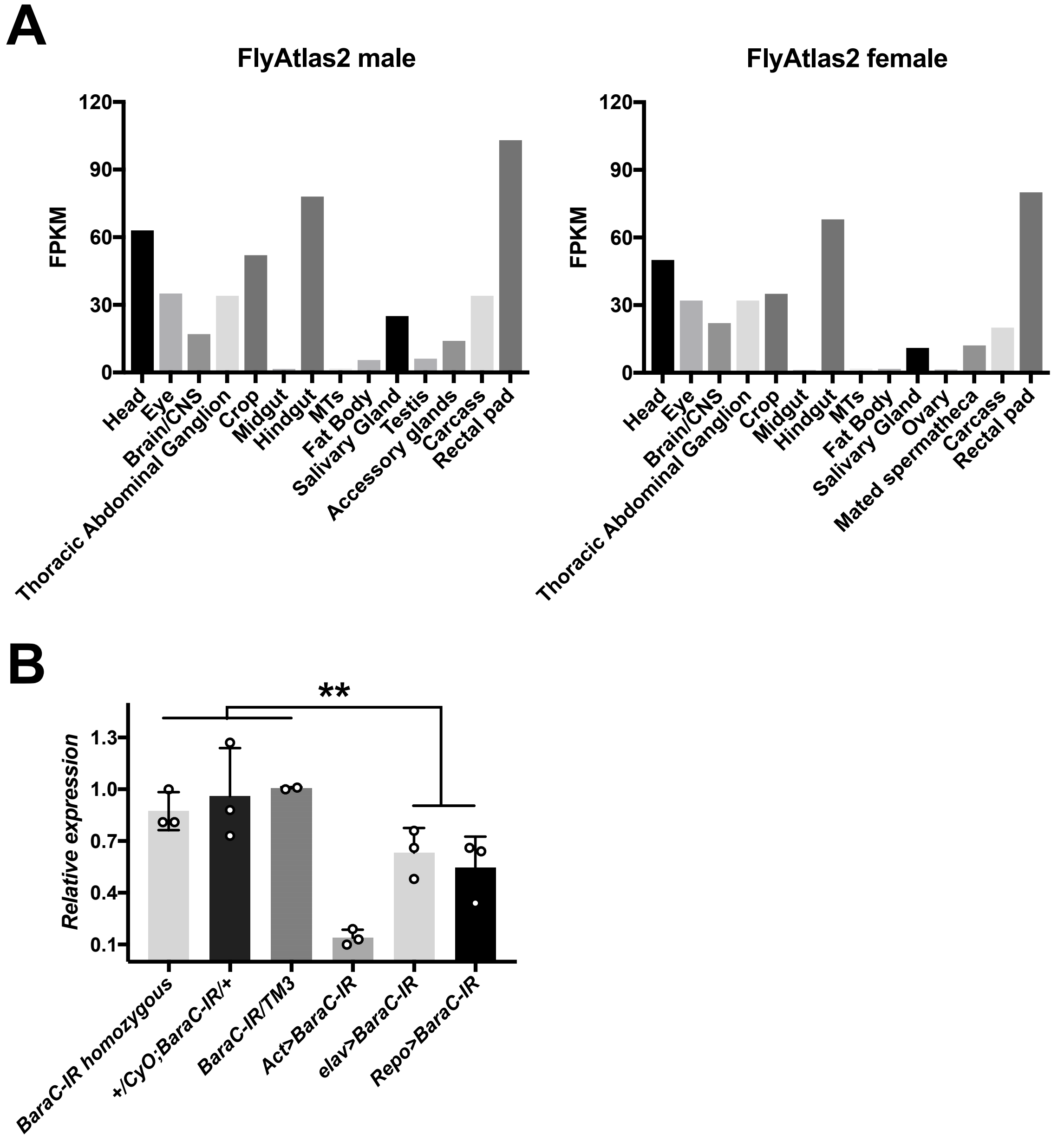


**Figure S4: *BaraC* is expressed in the nervous system, but also the hindgut and rectal pads. A)** FlyAtlas2 expression data for *BaraC*. **B)** RT-qPCR of *BaraC* in whole flies using different Gal4 drivers to express *BaraC* RNAi. *BaraC* is knocked down by both the *elav-Gal4* and *Repo-Gal4* nervous system drivers. Cumulatively, nervous system drivers significantly depress *BaraC* expression compared to *BaraC-IR* controls (student's t, $p < .01$). Ubiquitous knockdown using *Act>BaraC-IR* provides a comparative knockdown to better understand the strength of nervous system-specific knockdowns at the whole fly level.

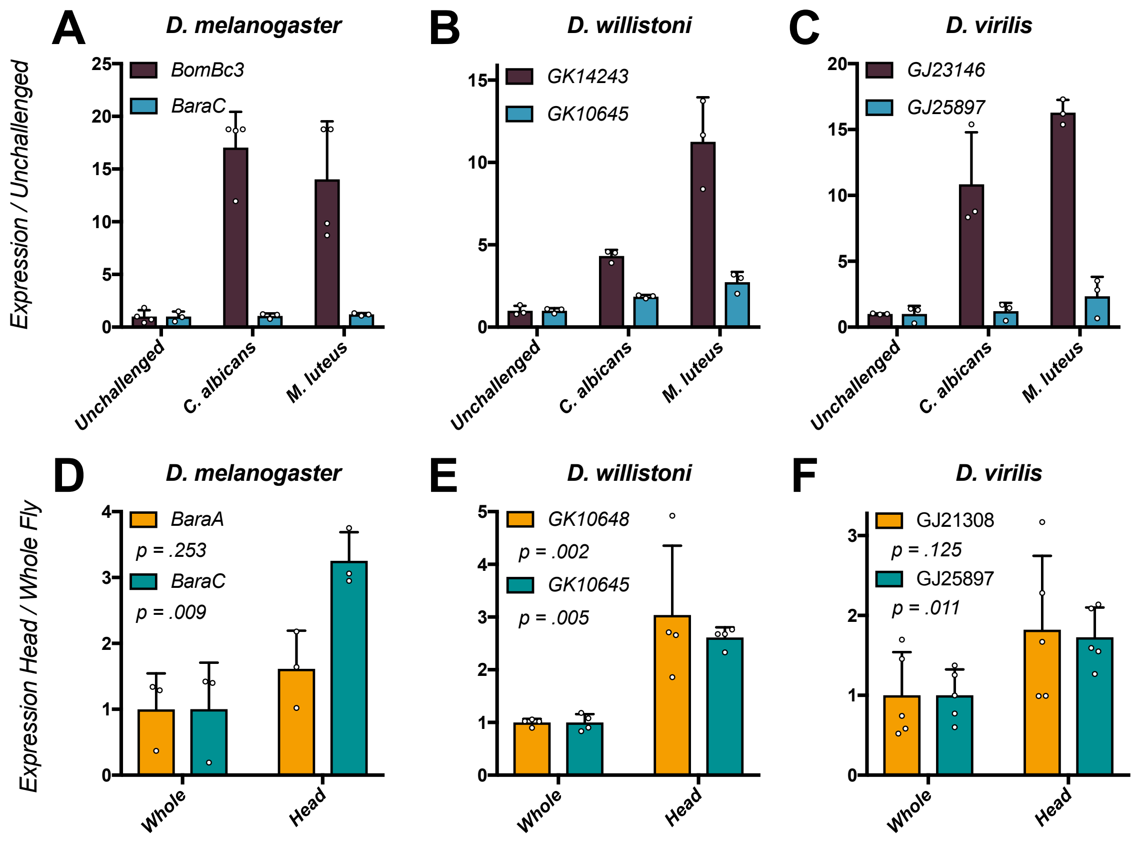


**Figure S5: RT-qPCR of *Baramicin* genes in diverse species. A-C)** Independent IM24-specific genes in *D. melanogaster* (A), *D. willistoni* (B), and *D. virilis* (C) are not induced by infection. *BomBc3* is included as an immune-induced control. **D-F)** The independent IM24-specific genes (blue) of *D. melanogaster* (D), *D. willistoni* (E), and *D. virilis* (F) are each enriched in the head relative to whole flies. *BaraA-like* genes (orange) were expressed more stochastically in the head, but also generally showed an enrichment pattern relative to whole flies (not always significant). Each data point represents an independent pooled sample from 20 male flies. Data were analyzed using one-way ANOVA with Holm's-Sidak multiple test correction.

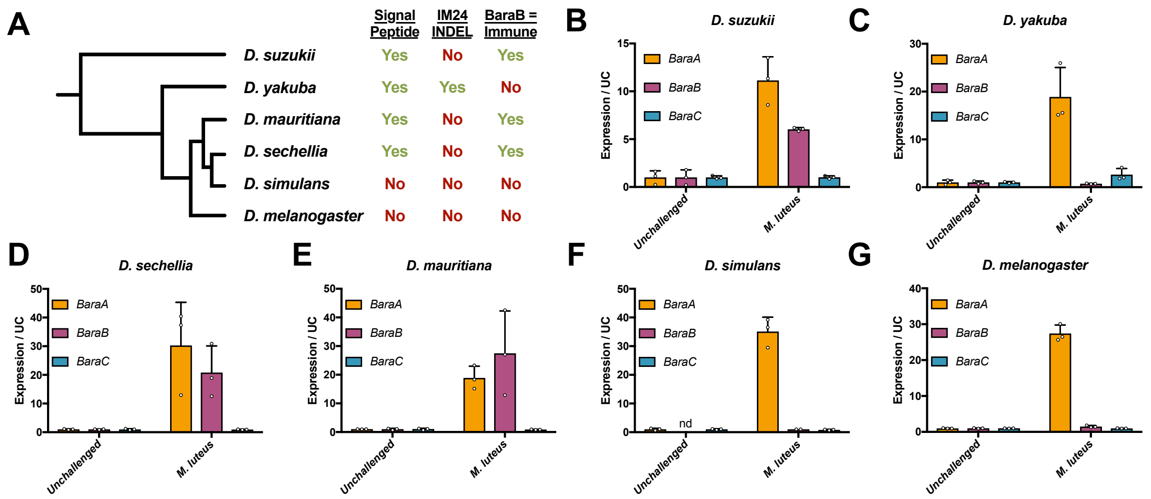


**Figure S6: The *D. melanogaster BaraB* gene acquired its non-immune role only recently. A)** Cladogram of the Melanogaster species group. The presence of a functional signal peptide (Fig. 5A), and the disruption of the *D. yakuba* IM24 peptide by an in-frame insertion is noted. A summary of whether *BaraB* is an immune-induced orthologue (B-G) is annotated. **B-G)** *Baramicin* expression data from Melanogaster group flies either unchallenged or infected with *M. luteus*. *BaraB* is immune-induced in *D. suzukii, D. sechellia,* and *D. mauritiana*, but not in *D. simulans* and *D. melanogaster,* which both lack signal peptide structures. *Drosophila yakuba BaraB* is not immune-induced (C), has an insertion event in its IM24 peptide (Fig. 5A), and its sister species *D. erecta* has pseudogenized its *BaraB* orthologue (Fig. 5B), suggesting pseudogenization may explain the lack of immune induction in *D. yakuba BaraB*.
